# Ligand-induced changes in dynamics mediate long-range allostery in the *lac* repressor

**DOI:** 10.1101/2021.11.30.470682

**Authors:** Anum Glasgow, Helen T. Hobbs, Zion R. Perry, Susan Marqusee, Tanja Kortemme

**Affiliations:** Department of Bioengineering and Therapeutic Sciences, University of California, San Francisco, CA 94158; Department of Biochemistry and Molecular Biophysics, Columbia University, New York, NY 10032; Department of Chemistry, University of California, Berkeley, Berkeley CA 94720; Department of Molecular Biophysics and Biochemistry, Yale University, New Haven, CT 06511; Department of Molecular & Cell Biology, University of California, Berkeley, Berkeley CA 94720

## Abstract

Allostery, broadly defined as a protein’s functional response to distal perturbations, is fundamental to biological regulation. In classical models, allosteric ligand binding produces a defined set of structural changes in the protein, resulting in a different low-energy conformation. Proteins that undergo ligand-induced allostery with few observable structural changes therefore frustrate interpretations by classical models. Here we used hydrogen-deuterium exchange with mass spectrometry (HDX/MS) to map the allosteric effects in a paradigm ligand-responsive allosteric transcription factor, the *lac* repressor (LacI). X-ray crystal structures of the core domain of LacI bound to different small molecule ligands, or the DNA operator, show less than 1.5 Å difference in the protein all-atom root-mean-square-deviation (RMSD) between any two structures. Despite this high degree of similarity among static structures, our HDX/MS experiments reveal widespread and unexpected differences in the flexibility of secondary structures in the LacI core domain in each functional state. We propose a model in which ligand binding allosterically switches the functional response of the repressor by selectively changing the dynamics of particular secondary structure elements relative to each other, shifting the conformational ensemble of the protein between mutually incompatible DNA-bound and inducer-bound states. Our model also provides a mechanistic context for the altered functions of thousands of documented LacI mutants. Furthermore, our approach provides a platform for characterizing and engineering allosteric responses in proteins.

## Introduction

Biological regulation ubiquitously depends on protein allostery, but few allosteric mechanisms are understood in structural detail. In the absence of high-resolution structural information, the Monod-Wyman-Changeux (MWC) (Monod et al., 1965) and Koshland-Nemethy-Filmer (KNF) (Koshland et al., 1966) models for allostery have been used to explain the functional behavior of diverse allosteric systems. In a few cases, detailed structural information has enabled descriptions of allosteric mechanisms for proteins, both in terms of specific, sequential conformational changes resulting from perturbation (Daily and Gray, 2009; Lockless and Ranganathan, 1999; Yu and Koshland, 2001) and as shifts in the average conformational ensemble (Cooper and Dryden, 1984; McLeish et al., 2015; Pan et al., 2000). However, deducing an atomic-level mechanism for allosteric effects on protein function is challenging in several common scenarios: when the beginning- and end-state average structures are structurally indistinguishable; when the structural change occurs over a large distance with few structural differences in the allosteric site; or when the protein is disordered in one or both states.

A paradigm example for these challenging scenarios is the *E. coli lac* repressor (LacI), a model allosteric transcription factor that regulates expression of the genes of the *lac* operon. Wild-type (WT) LacI forms a dimer of dimers, and each dimer can bind to one of three DNA operator sequences in the *E. coli* genome. The MWC and KNF models for protein allostery were established in part from studies of LacI after observing bacterial responses to sugar ligands (Jacob et al., 1965). The ligands bind in a pocket in the core domain of LacI, 40 Å away from the DNA-binding surface of the DNA-binding domain (Fig. 1A), and allosterically modulate the binding affinity of LacI for its DNA operator. Interestingly, X-ray crystal structures of the core domain are essentially identical in different ligand-bound states as well as the apo state (0.5-1 Å heavy atom RMSD) (Fig. 1B, C). The high structural similarity of the core domains, combined with a lack of X-ray crystal structures for a non-operator-bound DNA-binding domain, complicates mechanistic understanding of ligand-regulated allostery in LacI despite more than five decades of study using structural (Bell and Lewis, 2000; Daber et al., 2007; Friedman et al., 1995; Lewis et al., 1996), genetic (Coulondre and Miller, 1977a, 1977b; Farabaugh et al., 1978; Nelson et al., 1997; Suckow et al., 1996; Swerdlow and Schaaper, 2014; Swint-Kruse et al., 2003), computational (Daily and Gray, 2009; Flynn et al., 2003; Parente and Swint-Kruse, 2013; Villa et al., 2005), and biophysical (Adler et al., 1972; Kalodimos et al., 2002; Lamerichs et al., 1990; Morgan et al., 2005; O’Gorman et al., 1980; Royer et al., 1990; Wick and Matthews, 1991; Wilson et al., 2007, 2005) methods.

**Figure 1.**
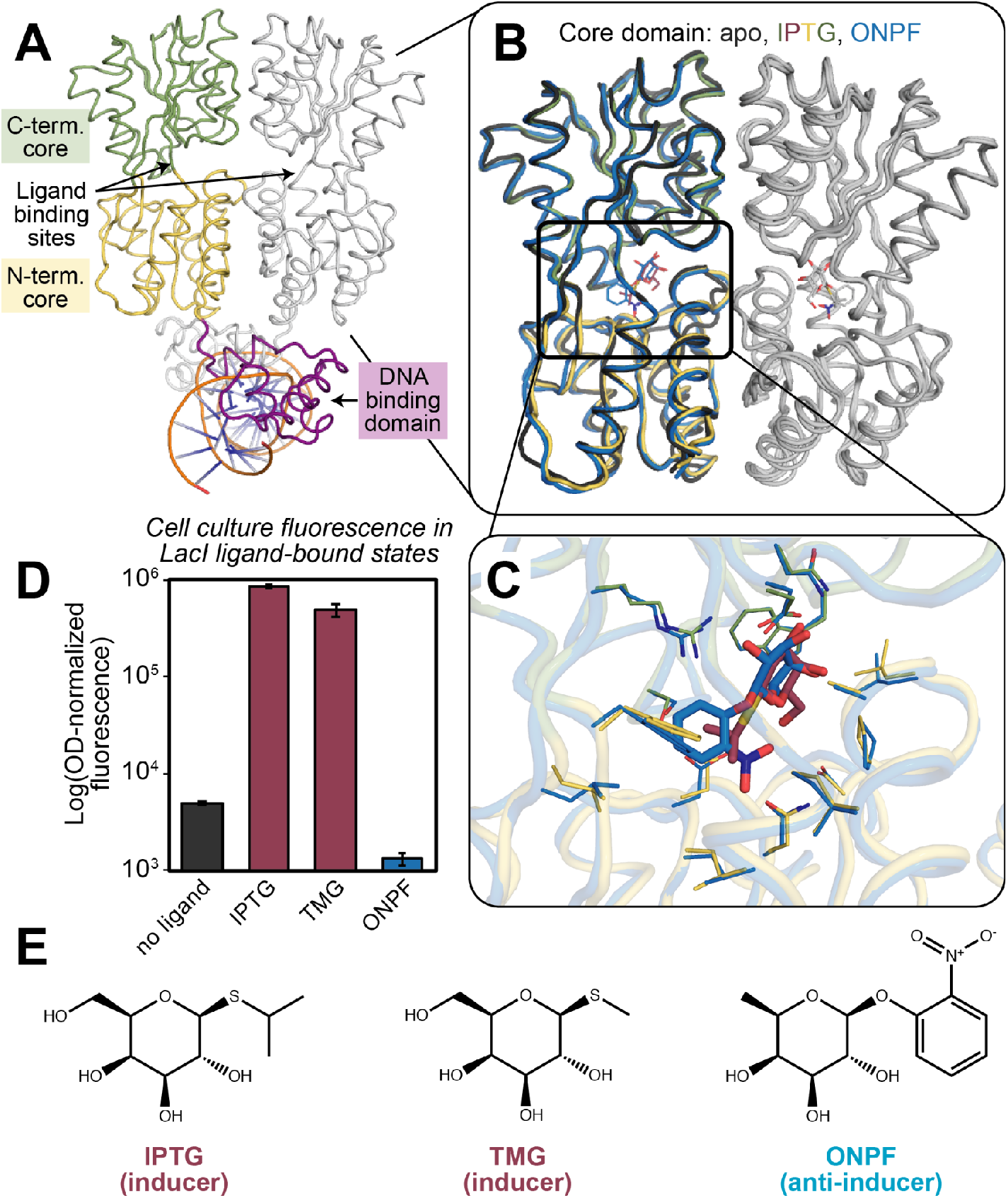
Ligands bind without dramatic structural changes in the core domain to control the structure of the DNA-binding domain allosterically. **(A)** X-ray crystal structure of dimeric LacI bound to the DNA operator. One subunit is shown in color (green: C-terminal subdomain; yellow: N-terminal subdomain; purple: DNA-binding domain) and the other subunit is shown in gray. PDB ID: 1EFA. **(B)** Overlaid structures of the core domains show similar backbone structures for apo-LacI (dark gray), IPTG-LacI (green, yellow, maroon), and ONPF-LacI (blue). PDB IDs: 1LBI, 2P9H, 2PAF. **(C)** Overlaid structures of the ligand binding site in IPTG-LacI and ONPF-LacI reveal similar sidechain-atom positions, colored as in (B). Thin sticks show amino acid sidechains. Thick sticks show ligands. **(D)** A cell culture fluorescence assay shows ligand-modulated gene expression by LacI. TMG- and IPTG-binding to LacI in cells results in GFP expression and fluorescence, but ONPF-binding results in increased repression. **(E)** Chemical structures of IPTG, TMG, and ONPF.

Adding to this challenge, chemically similar molecules binding in the core domain can have opposite effects on LacI-regulated gene expression (Fig. 1D, E). When bound to the inducers allolactose or isopropyl-β-D-1-thiogalactopyranoside (IPTG), LacI binds approximately 1000-fold less efficiently to the DNA operator than apo-LacI, allowing for the transcription of downstream genes. However, when bound to the anti-inducer *ortho*-nitrophenyl-β-D-fucoside (ONPF), LacI is stabilized in the operator-bound state by 2- to 5-fold over apo-LacI. Other ligands such as *ortho*-nitrophenyl-β-galactoside (ONPG) bind LacI but produce no observable functional effects. Inducer potency has been hypothesized to depend on the molecule’s binding affinity for LacI, the presence of an O6 hydroxyl group on the sugar ring, an optimal size for the C1 substituent, and the flexibility of the substituent linkage to the sugar ring (Barkley et al., 1975; Chakerian et al., 1987).

Several models have been proposed to explain the inducer-specific effects on gene expression by LacI. Crystallography and molecular dynamics simulations support a model in which inducer binding leads to new intramolecular interactions in the N-terminal subdomain of the core and partial unfolding of the DNA-binding domain, with the C-terminal subdomain of the core remaining unchanged (Daber et al., 2007; Richards et al., 2017; Rondon and Wilson, 2019).

Crystallographic studies of LacI bound to the inducer IPTG or to DNA revealed similar structures of the core domain, and a small shift (all-atom RMSD 1.5 Å) of the N-terminal subdomains of the core closer together in the IPTG-bound structure as compared to DNA-bound LacI (Fig. S1) (Daber et al., 2007). However, because of the structural similarity of the core domain in the apo, DNA-bound, inducer-bound, and anti-inducer-bound states of the protein, a detailed molecular mechanism for allosteric structural changes in the core domain upon ligand binding remains elusive.

Here, we map allosteric changes in the core domain of LacI in different functional states via hydrogen-deuterium exchange with mass spectrometry (HDX/MS). We define “functional states’’ by binding partner, where IPTG-LacI denotes the IPTG-bound protein; ONPF-LacI the ONPF-bound protein; ONPF-DNA-LacI the protein bound to ONPF and the *lac* operator at the same time; and apo-LacI the unbound protein. We propose a model for mutually exclusive low-energy conformational ensembles of LacI in inducer- and DNA-bound states. Using a combined computational and experimental approach, we show that protein, ligand, and solvent atoms together play a central role in modulating the rigidity of secondary-structure elements in the ligand-binding core domain of LacI to control its functional state, without dramatic conformational changes. Our model for allosteric regulation in LacI can inform future efforts to re-engineer the protein to control gene expression in response to new ligands, and establishes a method for characterizing how allostery in other proteins is conserved, evolved, or affected in disease.

## Results

We set out to determine the mechanism of allosteric changes in dimeric LacI by comparing structural and functional differences between DNA-bound states and inducer-bound states using HDX/MS. The DNA-bound and inducer-bound states are the key functionally relevant states of LacI. To study inducer-bound states, we complexed LacI with IPTG, a strong inducer molecule (LacI-IPTG K_D_ ∼ 5 × 10^−6^ M), and thiomethyl β-D-galactoside (TMG), a weaker inducer molecule (LacI-TMG K_D_ ∼ 7 × 10^−4^ M) (Barkley et al., 1975). To study DNA-bound states, we used the LacI-*lac* operator complex, with and without the anti-inducer molecule ONPF (LacI-operator K_D_ < 10^−12^ M, LacI-ONPF K_D_ ∼ 7 × 10^−3^ M) (Barkley et al., 1975). This strategy generated a set of LacI states with a range of binding affinities for the operator DNA (in order from unmeasurably weak to strongest): IPTG-LacI; TMG-LacI; DNA-LacI; and the ternary ONPF-DNA-LacI. We confirmed the effects on gene expression of IPTG, TMG and ONPF ligands binding to LacI using a plate-based cell culture assay, in which LacI regulates the expression of a green fluorescent protein (GFP) that is encoded downstream of the LacI DNA operator sequence on a plasmid (Fig. 1D, E).

For each state, we monitored amide hydrogen-deuterium exchange at the peptide level over a four hour time period using mass spectrometry (see Methods). From the full set of mass spectra representing more than 300 total peptides, we curated a set of 57 peptides from the core domain of LacI for which we could collect replicate data in six different states as well as a fully deuterated control HDX/MS experiment (in order to correct for back-exchange; see Methods). The curated dataset had 97% peptide coverage for the core domain, excluding the extreme C-terminal 22 amino acids (Fig. S2) for which we could not collect a complete dataset (data for each time point and each state with back-exchange).

### Comparison of inducer-bound and DNA-bound states

We first compared exchange patterns for core domain peptides in the IPTG-LacI and DNA-LacI states (Figs. 2, S3). We quantified the level of exchange in both states by calculating the deuterium uptake as a function of time for each peptide (Fig. S4), and fit these data to a nonlinear regression model to estimate the number of slow-, medium-, fast-, and non-exchanging protons in each peptide/state, and the resulting rates associated with each group (slow, medium, and fast) (Methods). In the deuterium-uptake plots in Figure 2A, increased rates of exchange for a peptide indicate increased structural flexibility associated with decreased participation in secondary structure hydrogen bonds by the peptide’s backbone amide groups. Figure 2B shows this behavior for the curated set of 57 peptides in the core domain, and Figure 2C illustrates these differences in H-D exchange between IPTG-LacI and DNA-LacI mapped onto the structure of the core domain.

**Figure 2.**
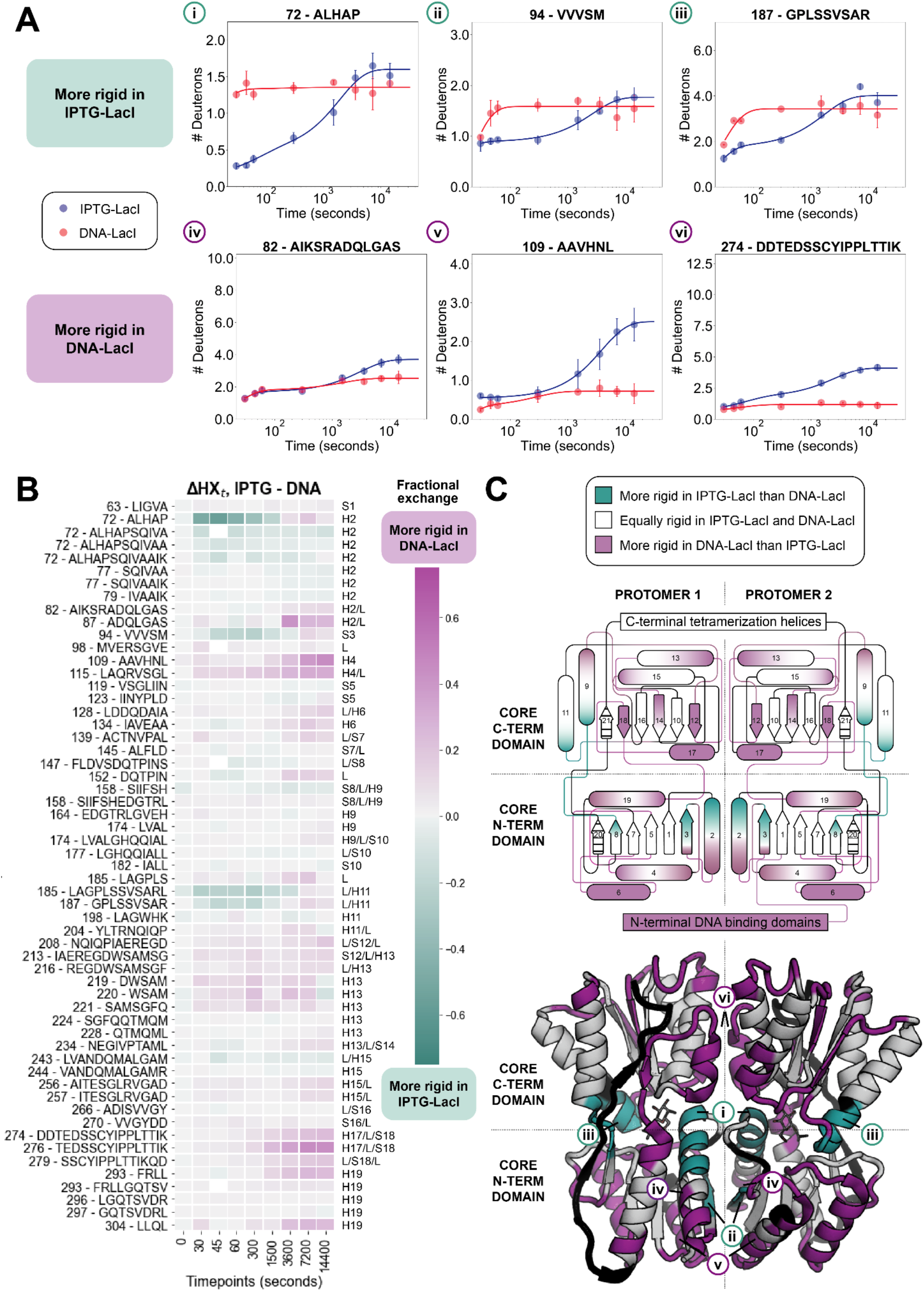
HDX/MS highlights key differences in the conformational ensembles of IPTG-LacI and DNA-LacI. **(A)** Deuterium uptake plots corresponding to the peptides i-vi marked in the bottom panel of (C). Peptides that are more rigid in IPTG-LacI are shown in the top row, and peptides that are more rigid in DNA-LacI are shown in the bottom row. The data points represent averaged, back exchange-corrected HDX/MS data for *n* = 2-5 individual measurements. The error bars are standard deviations. Lines are fit to averaged experimental data using a nonlinear regression model as described in the Methods. **(B)** Comparison of the fractional difference in H-D exchange at all timepoints in IPTG-LacI minus DNA-LacI for a curated set of 57 peptides. Purple indicates less exchange in DNA-LacI, while teal indicates less exchange in IPTG-LacI. Labels on the right side of the heatmap correspond to the labeled secondary structure elements in the top panel of (C). **(C)** Schematic (top) and structural model (bottom) of differential H-D exchange in IPTG-LacI and DNA-LacI, colored as in (B). Striped regions in the schematic and dark gray regions in the structural model indicate no data. Residues were labeled “more rigid” in either state if a larger than 20% difference in exchange was observed at three or more timepoints in HDX/MS peptides that included that residue. PDB ID: 2P9H.

When comparing IPTG-LacI to DNA-LacI, we observed both regions of decreased deuterium uptake (less exchange in IPTG-LacI relative to DNA-LacI, teal, Fig. 2A,C, i-iii, Fig. S5A) and regions of increased deuterium uptake (more exchange in IPTG-LacI, purple, Fig. 2A, C, iv-vii). Regions with *decreased* deuterium uptake are located near the ligand-binding pocket (Fig. 2A, C, i, N-terminal end of helix 2) and the monomer-monomer interface in the N-terminal subdomain (Fig. 2A, C, ii, beta strand 3). We hypothesize that the decreased deuterium uptake for beta strand 3 reports on the formation of a hydrogen bond between the backbone atoms of position 96 across the beta sheets of both N-terminal subdomains in the LacI dimer (Fig. S6). We also observed decreased deuterium uptake in regions distal to the monomer-monomer interface (Fig. 2C, iii, N-terminal portion of helix 11). Other regions in this category include the loop connecting beta strand 8 (N-terminal subdomain) and helix 9 (C-terminal subdomain) and residues 159-162 (Fig. 2C).

Conversely, regions with *increased* deuterium uptake (purple, Fig. 2) in IPTG-LacI compared to DNA-LacI included alternating beta strands in the C-terminal subdomain beta sheet (strands 12, 14 and 18) and all of the N-terminal subdomain helices (C-terminal end of helix 2, and helices 4, 6 and 19), which provide an interacting surface for the DNA-binding domain. Peptides from N-terminal subdomain helices 2 and 4 are shown in Fig. 2C, panels iv and v, respectively. Other regions in this category included residues in the C-terminal subdomain at the monomer-monomer interface (helix 13, residues 221-235, and helix 17, residues 276-282, Fig. 2C, vi, Fig. S5B). Of the nine loops that flank each ligand binding site, five loops exchanged more in IPTG-LacI than DNA-LacI.

Taken together, these results suggest a model where DNA-bound and inducer-bound states of the *lac* repressor adopt mutually incompatible low-energy conformational ensembles (Fig. 3A). DNA-LacI is characterized by an overall rigid structure of both the N- and C-terminal core domains, where the N-terminal subdomain makes extensive interactions with a well-folded operator-bound DNA-binding domain in the DNA-LacI crystal structure. Upon binding of the inducer IPTG, the ligand-binding pocket periphery as well as several secondary structure elements at the monomer-monomer interface are further rigidified, resulting in increased interactions between the LacI core subdomains. Importantly, this rigidification upon ligand binding leads to a simultaneous *increase* in flexibility in all of the N-terminal subdomain helices at the interface with the DNA-binding domain, even in the absence of the operator DNA (Fig. S7, see ΔHX_t_ IPTG-DNA plot). In addition, the C-terminal core subdomain helices 13 and 17, which flank the central helix 15 at the monomer-monomer interface, become more flexible in the inducer-bound states, in concert with increased flexibility in every other beta strand in the C-terminal subdomain beta sheet.

**Figure 3.**
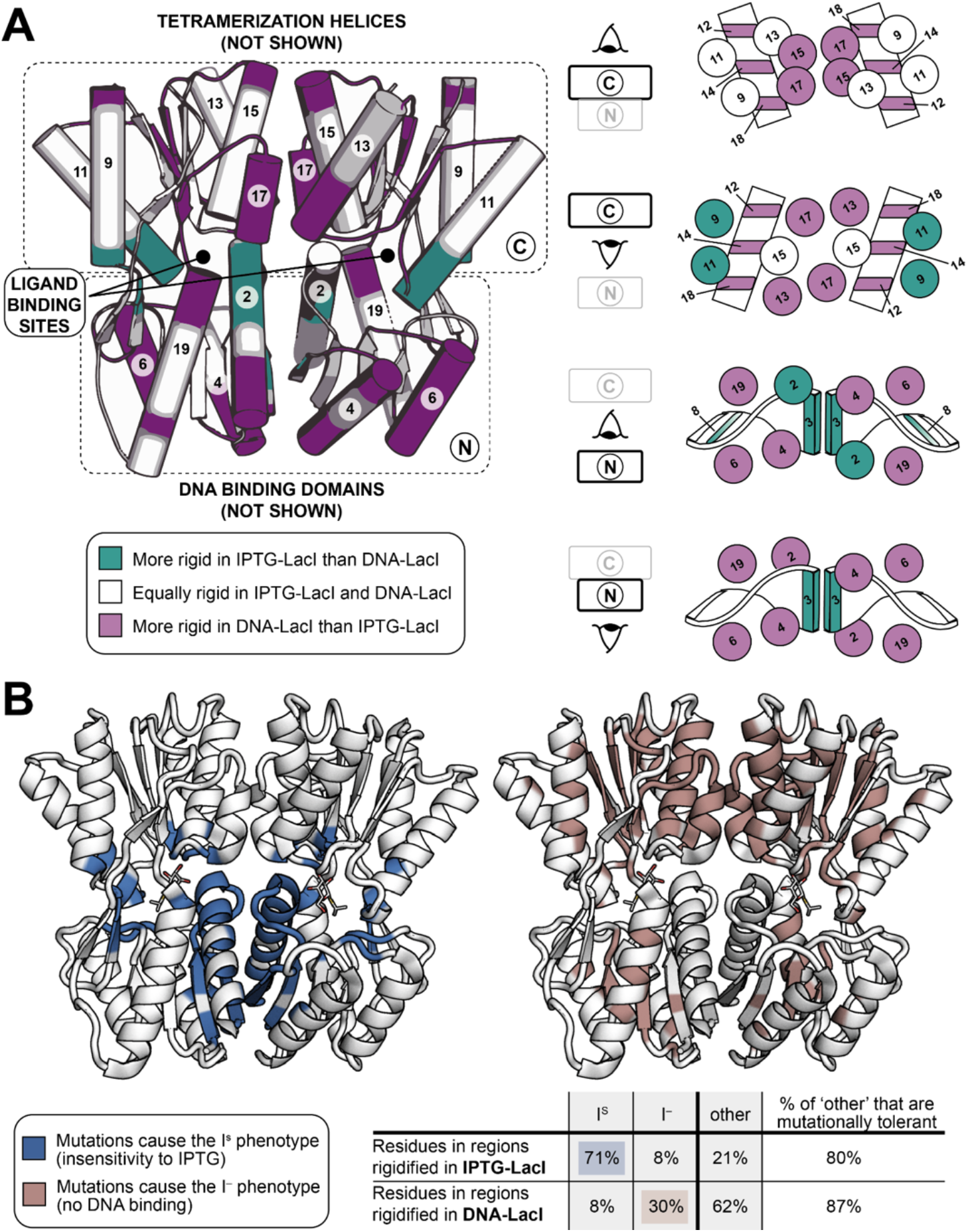
Structural differences correspond to functional differences in the two mutually incompatible low-energy states of LacI. **(A)** Differential H-D exchange between IPTG-LacI and DNA-LacI, visualized by core subdomains. Diagrams on the right are positioned and cross-sectioned according to the schematic of LacI on the left. Secondary structure elements are labeled according to the schematic in the top panel of Fig. 2C, with helices represented as cylinders or circles and sheets represented as arrows, rectangles, or twisted rectangles. Purple indicates less exchange in DNA-LacI, while teal indicates less exchange in IPTG-LacI. **(B)** LacI structural models colored according to the two main classes of observed mutational phenotypes: I^S^ (constitutive repression) and I^-^ (constitutive expression) (Suckow et al., 1996). The table summarizes the mutational phenotype data as percentages of residues in differentially rigidified regions observed in HDX/MS experiments. The final column shows the percentage of mutationally tolerant residues in the regions that are rigidified in IPTG-LacI or DNA-LacI but do not cause an I^S^ or I^-^ phenotype. PDB ID: 2P9H.

To further characterize differences in the rigidity of secondary structure elements between inducer- and DNA-bound states, we also used an alternate inducer. With few exceptions localized to the ligand binding pocket, we found that H-D exchange in TMG-LacI was similar to IPTG-LacI (Fig. S7). Likewise, H-D exchange in ONPF-DNA-LacI was similar to DNA-LacI (Fig. S7).

### Comparison to mutational data

We next compared our model for inducer-triggered conformational changes in the LacI core domain (Fig. 3A) to the extensive mutational phenotype data for LacI, where “phenotype” is defined by the effect of point mutations on gene expression. In a landmark study, 4000 LacI point mutants were tested for effects on gene expression (Suckow et al., 1996). The authors binned each WT amino-acid residue position according to its tolerance to mutation, and mapped these classifications onto the first crystal structure of LacI (Friedman et al., 1995) (Table S1, Fig. S8). The main mutational phenotypes observed in the study were: similar inducer response to WT LacI; unresponsive to inducer binding (I^S^); or constitutively inducing (I^-^).

We find a striking correlation between the phenotypic effects of these point mutants and our H-D exchange data. We found that the regions rigidified in IPTG-LacI (teal in Figure 2C, see Methods) correlated with positions at which mutations commonly cause an I^s^ phenotype, whereas regions rigidified in DNA-LacI (purple in Figure 2C) correlated with positions at which mutations cause an I^-^ phenotype. Mutations at 71% of the core domain residues that we observed to be rigidified in IPTG-LacI cause the I^S^ phenotype (Fig. 3B), but mutations to only 8% of these rigidified residues cause the I^-^ phenotype. Conversely, mutations at 30% of the core domain residues in regions that we observed to be rigidified in DNA-LacI cause the I^-^ phenotype, while mutations to only 8% of these residues cause the I^S^ phenotype. There were also mutations in regions with altered rigidity that cause neither phenotype, but the majority of these positions are tolerant to substitutions (Fig. 3B, last two columns in the table). Overall, we find considerable agreement between Suckow’s phenotype assignment groups and differences in exchange between DNA-LacI and IPTG-LacI. The HDX/MS and mutational phenotype data for LacI together support a model for allosteric response in which the differential rigidification of secondary structures relative to one another in each state modulates protein function. Mutations in regions that are uniquely rigidified in IPTG-LacI may destabilize the conformational ensemble of that state, leading to constitutive repression. Conversely, mutations in regions that are more rigid in DNA-LacI may destabilize the DNA-LacI conformational ensemble, leading to constitutive transcription. Notably, the regions of the protein that are rigidified in either state are not confined to the ligand binding pocket or the DNA-binding surface.

We also compared the HDX/MS data to functional data from decades of LacI biochemistry studies, focusing on the effects of mutations in regions of the core domain that respond to IPTG- and DNA-binding in different ways: the monomer-monomer interface and the binding-pocket periphery.

At the monomer-monomer interface in helix 2, mutations at A81 and K84 (Fig. S9A) changed the IPTG-LacI and DNA-LacI binding affinities, as well as the protein stability (Chakerian et al., 1985; Chang et al., 1993; Markiewicz et al., 1994; Nichols and Matthews, 1997; Swint-Kruse et al., 2005). One hypothesis informed by crystal structures is that an inter-subunit, inter-subdomain salt bridge between H74 in helix 2 and D278 in helix 17 of the other subunit (Fig. S9B) stabilizes the conformation of IPTG-LacI (Lewis et al., 1996) although mutations to these residues do not necessarily disrupt IPTG binding (Barry and Matthews, 1999). Our H-D exchange data alongside these mutational data suggest that several quaternary contacts at the monomer-monomer interface in helices 2, 13, and 17, and beta strand 3, facilitate the IPTG-specific shifts in the conformational ensemble (Figs. 2B-C, S5).

At the N-terminal subdomain monomer-monomer interface, mutations to residues 94-98 in beta strand 3 (Fig. S9C) have also previously been shown to disrupt LacI allostery (Zhan et al., 2010, 2009). This observation is consistent with our hypothesis that hydrogen bonds across the beta sheets in both subunits rigidify the inducer-bound states, which arises from the observation of decreased H-D exchange in beta strand 3 in IPTG-LacI as compared to DNA-LacI (Fig. 2A, ii, and see also peptide 94 - VVVSM in Fig. S7, ΔHX_t_ DNA-TMG and ΔHX_t_ IPTG-DNA plots). Crystal structures additionally show a decrease in the distance between the beta strands, which corresponds to the formation of direct hydrogen bonds between V96 backbone atoms in the IPTG state, rather than water-mediated hydrogen bonds in the DNA state in which the sidechain of K84 interacts with the carbonyl oxygens of V94 and V96 (Fig. S6).

Finally, we considered residues in the binding pocket-peripheral loops that interact directly with IPTG, TMG, and ONPF. Mutagenesis to residues L148 and S151 (Fig. S9D) resulted in shifts in the binding affinities of LacI for both IPTG and the DNA operator (Swint-Kruse et al., 2003). In line with the shift in binding affinity for IPTG noted in these mutational studies, we observed attenuated H-D exchange in the pocket-peripheral loops for IPTG as compared to the weaker-affinity ligand TMG (Fig. S7, ΔHX_t_ IPTG-TMG plot, peptides 98 - MVERSGVE, 158 - SIIFSH, 147 - FLDVSDQTPINS and 185 - LAGPLSSVSARL) (Barkley et al., 1975).

### Effect of anti-inducer binding

We next sought to identify differences in the conformational ensembles between IPTG-LacI and ONPF-LacI. The two ligands IPTG and ONPF have opposite functional effects, where IPTG destabilizes and ONPF stabilizes the DNA-bound state (Barkley et al., 1975; Daber et al., 2007). To facilitate the comparison and because IPTG-LacI does not show measurable binding to DNA, we compared H-D exchange for IPTG-LacI and ONPF-LacI without the DNA operator to the apo state (Fig. 4). We found that, compared to the apo state, rigidification (less exchange) of particular secondary structure elements near the ligand binding pocket and in the interface between the N- and C-terminal core subdomains (the N-terminal part of helix 2, strand 3, strand 8 and the loop C-terminal to strand 8, helix 11, helix 15 and the loop N-terminal to helix 15) was unique to IPTG-LacI and not observed in ONPF-LacI (Figs. 4A, B, iv-vi). In contrast, ONPF binding rigidified regions distal to the ligand binding site in the N-terminal subdomain (Fig. 4A, B, i-iii, the C-terminal end of helix 2 and the loop connecting it to strand 3, the loop between strand 3 and helix 4, the loop between helix 4 and strand 5, the loop between strand 5 and helix 6, the loop between helix 6 and strand 7, the loop connecting strands 7 and 8, and the C-terminal ends of helices 6 and 19). Many of these regions interface directly with the DNA-binding domain (although DNA is absent in these HDX/MS experiments). Taken together, these results suggest that IPTG and ONPF selectively stabilize different secondary structure elements in the core domain to shift the conformational ensemble of LacI to different, mutually exclusive low-energy states. Unlike IPTG binding, ONPF binding does not cause dramatic changes to the structure of LacI that are incompatible with the observed conformational ensemble of DNA-LacI (Fig. S7, compare ΔHX_t_ ONPF-APO, ΔHX_t_ IPTG-APO, and ΔHX_t_ APO-DNA plots).

**Figure 4.**
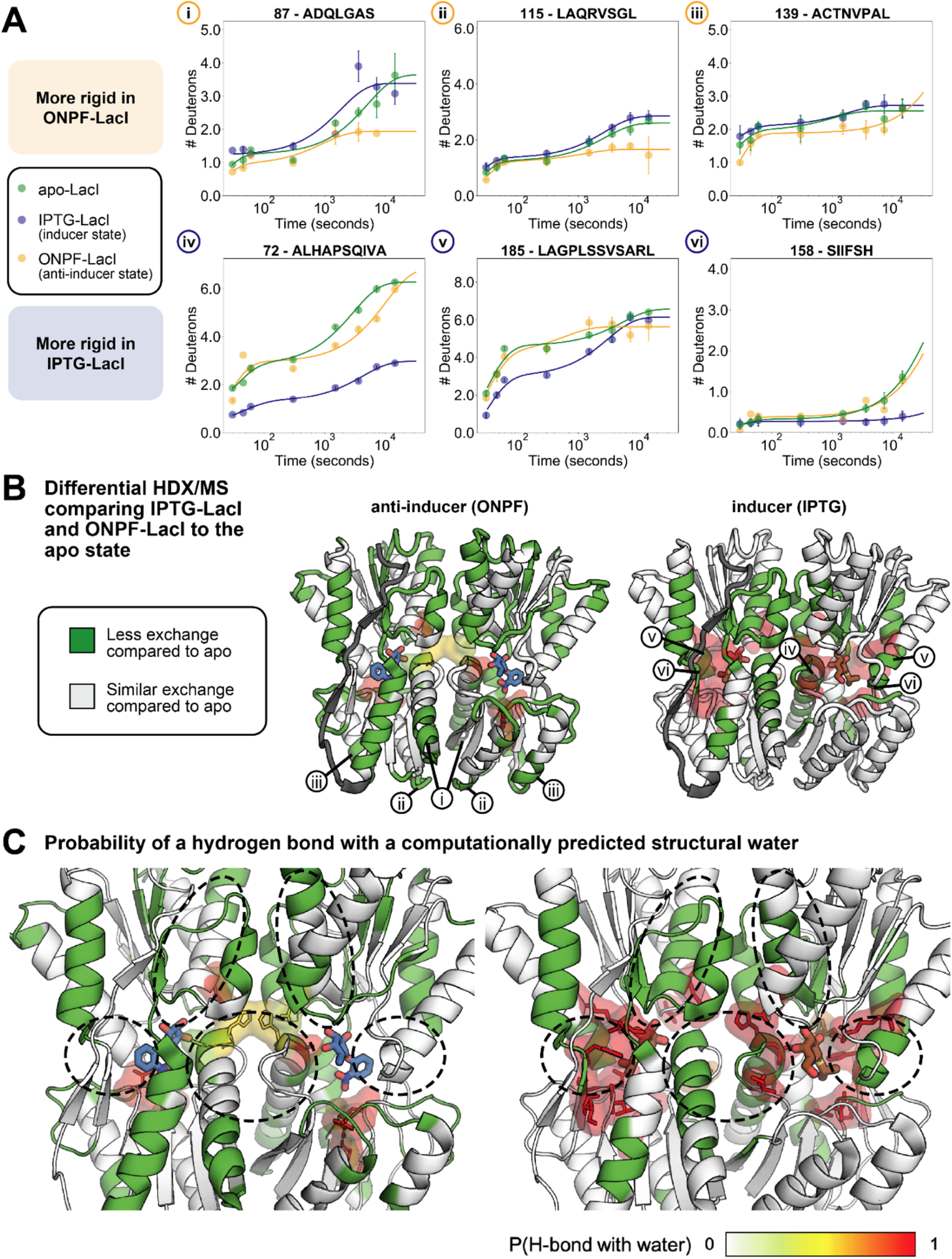
Computationally predicted solvent-mediated hydrogen bonding networks in the ligand-binding pocket of LacI correspond with differences in H-D exchange between IPTG-LacI and ONPF-LacI. **(A)** Deuterium uptake plots for the peptides i-vi marked in (B). Peptides that are more rigid in ONPF-LacI are shown in the top row (yellow line), and peptides that are more rigid in IPTG-LacI are shown in the bottom row (blue line), with apo-LacI (green line) shown for reference. The data points show averaged, back exchange-corrected HDX/MS data for *n* = 2-5 individual measurements and error bars show standard deviations. Lines are fit to averaged experimental data using a nonlinear regression model as described in the Methods. **(B)** Crystal structures of the core domain of ONPF-LacI (left) and IPTG-LacI (right), colored according to differences in H-D exchange compared to apo-LacI. Green regions are rigidified in each ligand-bound state as compared to apo-LacI. Dark gray regions indicate no H-D exchange data. **(C)** Overlay of models of the ligand-binding pockets with ONPF (left) and IPTG (right) bound, with rigidified residues colored green in the cartoon representation as in (B), and surface representations showing the predicted probabilities of residue interactions with structural water molecules by our semi-explicit solvation protocol. Sidechains that are predicted to interact with structural waters are shown as sticks and surfaces and colored according to probabilities calculated from 100 trials. Secondary structure elements that are rigidified in IPTG-LacI but not ONPF-LacI are highlighted in the dashed circles.

The distinctive rigidification of the N-terminal subdomain in ONPF-LacI in the absence of DNA suggests that ONPF binding could preorganize this subdomain and the DNA-binding domain for DNA-binding. To test this idea, we compared HDX/MS data for peptides in the DNA-binding domain in different functional states of LacI. The structure of the DNA-binding domain has only been solved by X-ray crystallography in the DNA-LacI state (Fig. S10A). In these crystal structures, the N-terminal subdomain beta strands and helices form electrostatic interactions with a well-folded DNA-binding domain. Interestingly, while HDX/MS confirmed that the DNA-binding domain is well-structured in DNA-LacI and ONPF-DNA-LacI, we observed a range of hydrogen exchange behavior in the DNA-binding domain for the non-DNA-bound ligand-bound states. The DNA-binding domain underwent the most H-D exchange across timepoints in IPTG-LacI, particularly in the hinge helix (residues 50-57) that connects the DNA-binding domain to the core domain (Fig. S10B), and was therefore the least structured among the states. However, decreased H-D exchange in the DNA-binding domain for ONPF-LacI when compared to IPTG-LacI even in the absence of DNA suggests that ONPF binding preorganizes the DNA-binding domain, thereby improving the binding affinity of LacI for the DNA operator when compared to apo-LacI (Fig. S10C).

### Water-mediated interactions in the ligand binding site

Several secondary structure elements near the ligand binding pocket that show decreased exchange only in IPTG-LacI and/or TMG-LacI, such as helix 2, strand 8 and helix 11 (Figs. S3, S7), do not make direct contacts with ligands in X-ray crystal structures (Fig. S11). We hypothesized that decreased exchange in these regions may result from the formation of additional solvent-mediated hydrogen bonds in the ligand binding pocket that form upon binding inducers, but not anti-inducers. Crystal structures of IPTG-LacI show such solvent-mediated interactions (Daber et al., 2007). In addition, mutations at binding pocket residues D149 and N125, which form water-mediated hydrogen bonds to IPTG in the crystal structure, decrease the protein’s affinity for IPTG (Xu et al., 2011).

To explore the potential role of water-mediated hydrogen bonding networks in the ligand binding pocket, we applied a computational “semi-explicit” solvation protocol in the macromolecular modeling program Rosetta (Pavlovicz et al., 2020) to predict the positions of structural water molecules in IPTG-LacI, ONPF-LacI, and apo-LacI. We defined “structural waters” as water molecules that make three or four hydrogen bonds with atoms in the protein or ligand. As inputs, we first relaxed the experimentally solved crystal structures of IPTG-LacI and ONPF-LacI (Supplemental computational methods). The computational protocol then places water molecules in hydrogen bonding geometries relative to protein polar groups. Highly coordinated water molecules in the protein structure are kept, with or without “solvated” sidechains allowed to change rotameric conformations in Monte Carlo simulations. Only one high-resolution structure of LacI that includes electron densities for water molecules in the ligand binding sites is available (protein databank (PDB) ID 2P9H, the core domain of IPTG-LacI). Using our computational strategy without sidechain repacking, we first confirmed that the positions of our computationally predicted structural water molecules in IPTG-LacI are within 0.5 Å of several water molecules observed in this structure (Fig. S12). We then placed structural water molecules in the ligand binding pockets of ONPF-LacI and apo-LacI from crystal structures using the same protocol (Figs. 4C, S13). Based on 100 models for each LacI state, we calculated the probability that an atom in the protein or ligand makes a hydrogen bond with a computationally placed structural water in the ligand binding pocket (Fig. S13).

We observed a distinctive correspondence between regions with ligand-induced rigidification of secondary structures as measured by HDX/MS (Figs. 4A, B) and the calculated probability for water-mediated hydrogen bonding among residues in the ligand binding pocket (Fig. 4C, circled regions correspond to regions iv, v, and vi highlighted in Fig. 4A, bottom row, and 4B, right panel, as well as helix 15). Most strikingly, we predict an extensive water-mediated hydrogen bonding network for IPTG-LacI (Fig. 4C, right). In contrast, this network is not observed in the ONPF-LacI simulations (Fig. 4C, left). This result supports a model in which structural water molecules play a role in differentially rigidifying LacI in different functional states, where water-mediated hydrogen bonding networks contribute to increased interactions in the ligand binding pocket between the N- and C-terminal core subdomains in IPTG-LacI but not ONPF-LacI.

## Discussion

We used a computational and experimental approach to map the allosteric response of LacI to different binding partners in six different functional states, at secondary structure resolution. Because the X-ray crystal structures of the core domains of DNA-LacI and IPTG-LacI are almost identical, our approach addressed the long-standing question of how allosteric effects in LacI are transmitted across more than 40 Å in the absence of sizeable conformational changes between functional states. Our model for inducer-triggered relief of DNA operator binding centers on selective dynamics of particular secondary structure elements relative to each other, shifting the conformational ensemble of the protein between mutually incompatible DNA-bound and inducer-bound states. Our results provide a detailed molecular mechanism consistent with data from structural and biochemical studies, expanding on the original MWC and KWF phenomenological models. The differences in conformational ensembles between the inducer-bound states and DNA-bound states that we observed by HDX/MS and predicted using computational methods also provide a mechanistic context for the altered functions of thousands of documented LacI mutants.

Figure 5A illustrates the key differences in the mutually incompatible conformational ensembles of the core domains in the IPTG-LacI and DNA-LacI states. The main features of IPTG-LacI, compared to DNA-LacI, are: rigidification of specific secondary structure elements near the ligand binding pocket (teal); increased specific inter-subdomain and inter-subunit interactions in the periphery of the ligand binding site (orange lines), some of which are supported by hydrogen bonding of protein and ligand atoms with structural water molecules (red circles and lines); and increased flexibility of other secondary structures throughout the C-terminal subdomain and in the N-terminal subdomain adjacent to the DNA-binding domain (purple). In contrast, DNA-LacI is characterized by increased rigidification in both core subdomains away from the ligand-binding site and at the interface with the DNA binding domain. We propose that IPTG binding rigidifies the N-terminal end of helix 2 and the inter-subunit contacts at the expense of the rigidity of neighboring helices. These IPTG-triggered shifts in the conformational ensemble in the core domain lead to increased flexibility in the hinge helices that in turn increase flexibility in the DNA-binding domain and ultimately weaken the binding affinity of LacI for the DNA operator.

**Figure 5.**
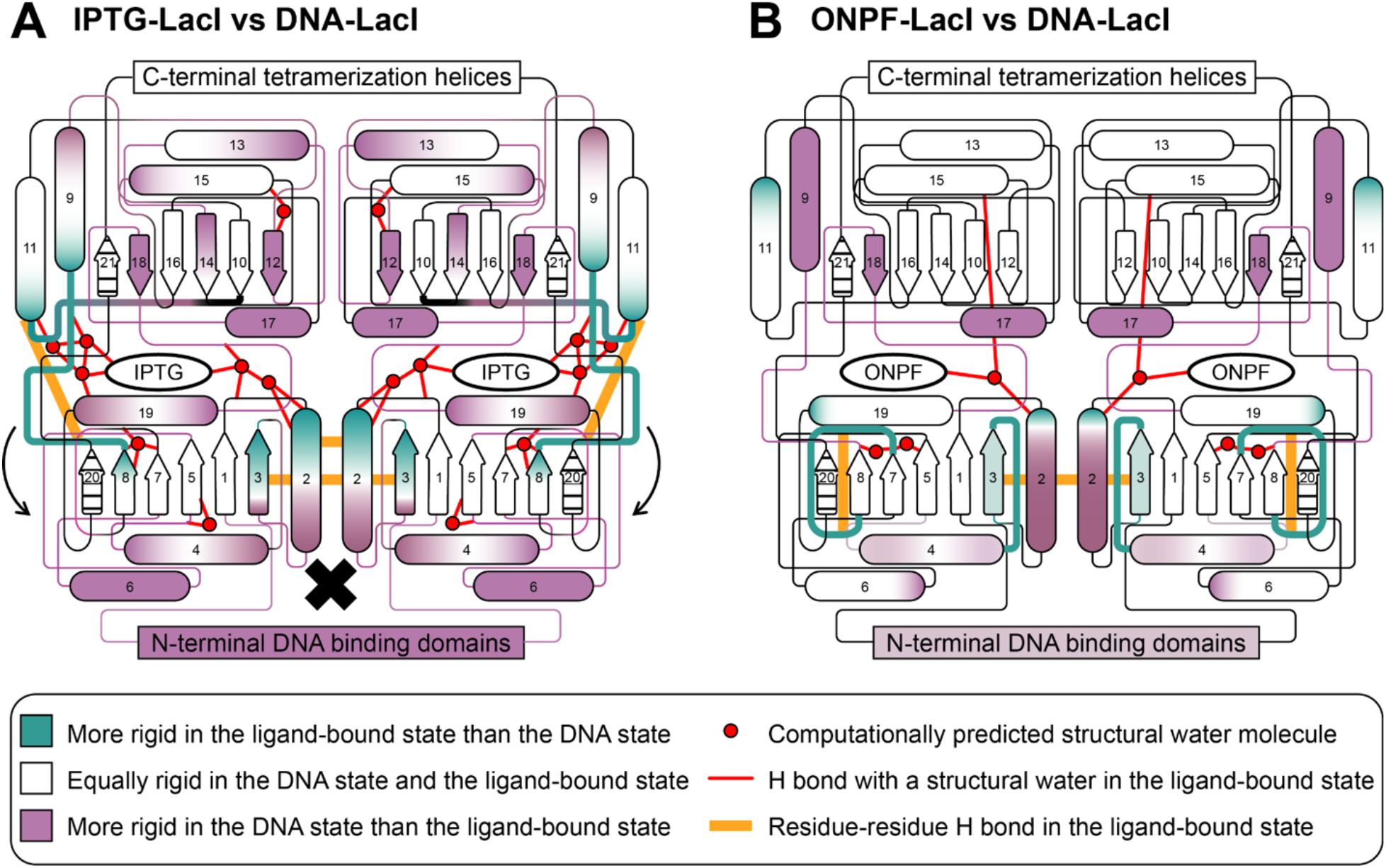
Consolidated models for ligand-specific changes in the conformational ensemble in IPTG-LacI and ONPF-LacI, as compared to DNA-LacI. Schematics summarizing the rigidity of LacI core domain secondary structure elements in IPTG-LacI and ONPF-LacI as compared to DNA-LacI, with rigidified regions in each ligand-bound state shown in teal, de-rigidified regions shown in purple, and regions for which there is no data shown with stripes. Red circles show the positions of predicted structural water molecules, and red lines show the interactions of the predicted structural water molecules with atoms in ligands or amino acid residues. Orange lines represent hydrogen bonds uniquely formed among protein atoms in each ligand-bound state as compared to DNA-LacI. Rigidified loops in each schematic are thick for emphasis. **(A)** In IPTG-LacI, hydrogen bond formation at the ligand-binding pocket periphery promotes interactions between the N- and C-terminal subdomains of the core, which causes small structural rearrangements in the N-terminal subdomains (arrows) to increase flexibility in the hinge helix (black X). **(B)** In ONPF-LacI, the C-terminal core subdomain is not extensively perturbed as compared to DNA-LacI. However, N-terminal core subdomain loops become more structured, which may stabilize the interface of the core domain with the DNA-binding domain to bind the operator.

Figure 5B shows the key differences in the conformational ensembles of the core domains of ONPF-LacI and DNA-LacI. The main feature of ONPF-LacI is increased rigidity (teal) in loops in the N-terminal subdomain of the core distal from the ligand binding site. We propose that because ONPF binding does not increase the flexibility of key secondary structure elements in the core domain to the same extent as inducer binding, ONPF binding is not incompatible with the DNA-bound conformational ensemble. The main functional consequence of ONPF binding may be that the binding interface of the core domain with the DNA-binding domain is preorganized in ONPF-LacI as compared to apo-LacI, leading to an increase in the binding affinity of LacI for the DNA operator.

One interesting aspect of our model is that it includes allosteric differences in exchange not only in the N-terminal subdomain but also in the C-terminal subdomain of the core (Figs. 2B-C, 3A). This observation is in contrast to previous models because the C-terminal subdomain in this family of transcription factors has long been considered conformationally fixed (Lewis et al., 1996; Mowbray and Björkman, 1999; Schumacher et al., 2004, 1995; Swint-Kruse and Matthews, 2009), with “virtually no difference in the hydrogen bond patterns or the number of hydrogen bonds in the C-terminal subdomains” (Daber et al., 2007). Targeted molecular dynamics simulations have built on this assumption, focusing solely on dynamics in the N-terminal subdomain while anchoring the C-terminal subdomain in place (Flynn et al., 2003). However, differences in HDX on the C-terminal side of the binding pocket, at the C-terminal monomer-monomer interface, in the beta sheet spanning the C-terminal subdomain, and in C-terminal subdomain solvent-exposed loops far from the ligand binding pocket highlight conformational changes that uniquely result from inducer binding. These observations provide a structural explanation for why mutations to these residues, for example D274, disrupt the response of LacI to binding IPTG (Chang and Matthews, 1995; Markiewicz et al., 1994; Suckow et al., 1996).

Engineering allosteric regulation in proteins remains challenging because allosteric mechanisms are typically incompletely understood. Our study provides mechanistic understanding of differences in conformational ensembles that can inform efforts to reengineer natural allosteric proteins to respond to user-defined perturbations. Transcription factors regulate gene expression through two types of binding interactions: binding to a specific DNA sequence, and binding to a set of small molecule or protein partners that control DNA binding. Domain-swapping (Jiang et al., 2021; Tungtur et al., 2010), chimeric fusions (Juárez et al., 2018), directed evolution (Richards et al., 2017; Taylor et al., 2016), saturation mutagenesis (Suckow et al., 1996; Wu et al., 2017), and deep mutational scanning (Tack et al., 2021) have resulted in LacI variants that control gene transcription in unexpected and interesting ways that depart from WT behavior, or in response to other sugars. However, none of these approaches are predictive with respect to tuning ligand response or allostery. Our strategy for computationally predicting interactions between protein, ligand, and water molecules and observing changes in conformational flexibility can help pinpoint structural changes that govern allosteric behavior in transcription factors and other proteins towards predictively and precisely designing the genotype-phenotype landscape.

## Materials and methods

### Cell culture fluorescence assay

The *E. coli* DH10B Δ*lacI*:220 strain was chemically transformed with a plasmid encoding superfolder GFP (sfGFP) under the control of the pLlacO promoter and a plasmid encoding the WT LacI (pEZ22G-GFP-kan and pSc101-lacI-specR, gifts from the Raman lab at U. Wisconsin) (Taylor et al., 2016) and plated on Lysogeny broth (LB) agar with 50 µl/ml kanamycin and 50 µl/ml spectinomycin. A culture was grown overnight in LB medium with antibiotics at the same concentrations from a single colony. New 5 ml cultures were grown at 37 °C and shaking at 225 rpm for 16 hours in blocks by subculturing the overnight culture using a 1:100 dilution in fresh LB medium. IPTG, TMG, and ONPF (Sigma-Aldrich, St. Louis, MO) were added to separate cultures to final concentrations of 100 µM, 100 µM, and 3 mM, respectively, at the two hour mark. Fully grown cultures were spun down at 3,000 rpm for ten minutes and the supernatant was decanted. Pellets were washed two times in 5 ml 1X phosphate-buffered saline (PBS), resuspended in 5 ml 1X PBS, and diluted 1:20 in 200 µl PBS in a clear-bottom black 96 well microtiter plate (Costar). Fluorescence from sfGFP expression was read from 7 mm above the plate from all sample wells using an excitation wavelength of 485 nm and an emission wavelength of 528 nm. All fluorescence readings were normalized by culture growth, measured by absorbance at 600 nm.

### Protein expression and purification

A BL21(AI) Δ*lacI*-ZYA strain was generated by the method of Datsenko and Wanner (Datsenko and Wanner, 2000) and BL21(AI) Δ*lacI*-ZYA cells were transformed with the pET9a-6×His-LacI(1-331) vector. This construct encodes for dimeric LacI, without the tetramerization domain. All HDX/MS experiments were conducted with dimeric LacI. Cells were grown at 37 °C while shaking at 225 rpm for 12-16 hours overnight in LB medium with 50 µg/ml kanamycin (LB-kan). The overnight culture was subcultured at a 1:50 dilution in LB-kan with 0.2% glucose and grown at 37 °C while shaking at 225 rpm for two hours, until the OD600 value of the culture was between 0.4 and 0.5. The temperature was then decreased to 16 °C, protein expression was induced by the addition of 0.2% arabinose, and cells were grown for an additional 16 hours. The culture was centrifuged at 6,000 × *g* for 20 minutes at 4 °C and the pellet was frozen at −80 °C for at least two hours. The pellet was then thawed on ice for 15 minutes and resuspended in 40 ml lysis buffer (50 mM Tris, 150 mM NaCl, pH 8.0, with 5 mM MgCl_2_, 1 mM MnCl_2_, 100 µM CaCl_2_, a dissolved Pierce™ protease inhibitor cocktail tablet (Thermo Fisher Scientific, Waltham, MA), and 40 µg DNaseI (Thermo Fisher Scientific)) by gentle vortexing for 20 minutes. The resuspended cells were lysed by two passes of microfluidization. The lysed cells solution was centrifuged at 27,000 × *g* for 30 minutes at 4 °C. The soluble cell fraction was decanted, supplemented with 20 mM imidazole, stored on ice for one hour, centrifuged at 3,270 × *g* for 10 minutes at 4 °C, and decanted again. We added 2 ml Ni^2+^-NTA slurry (HisPur Ni-NTA resin, Thermo Fisher Scientific) in binding buffer (50 mM Tris, 150 mM NaCl, 20 mM imidazole, pH 8.0) to the soluble cell fraction and nutated the protein-resin solution at 4 °C for one hour. The protein-resin solution was then centrifuged at 500 × *g* for five minutes at 4 °C and the supernatant was removed. The pelleted resin was resuspended in 2 ml binding buffer and applied to a 20 ml chromatography column. The resin was washed in the column with 50 ml binding buffer, then the protein was eluted in 14 ml elution buffer (50 mM Tris, 150 mM NaCl, 250 mM imidazole, pH 8.0). The eluted protein solution was spin-concentrated to a volume of 1.5 ml using Amicon™ Ultra Centrifugal Filter Units (MilliporeSigma, Burlington, MA) with a 10,000 Da molecular weight cutoff (MWCO). A volume of 3 µl from a 2 mg/ml stock solution of tobacco etch virus (TEV) protease with an N-terminal 6× histidine tag and 0.5 mM tris(2-carboxyethyl)phosphine (TCEP) was added to the concentrated protein solution. The reaction was incubated at 30 °C for two hours. Then, we diluted the reaction to 3 ml in LacI in dialysis buffer (50 mM MOPS, 200 mM NaCl, 0.5 mM TCEP, pH 7.0) and dialyzed the reaction in dialysis buffer at 4 °C while stirring, using a dialysis cassette with a 3,000 Da MWCO (Slide-A-Lyzer, Thermo Fisher Scientific). The dialysis buffer was changed in 1 L increments of approximately 2 hours, 16 hours, and 2 hours. The dialyzed protein solution was diluted to 17 ml in dialysis buffer with 20 mM imidazole and supplemented with 2 ml Ni^2+^-NTA slurry that had been pre-equilibrated in dialysis buffer, nutated at 4 °C for one hour, and then applied to a 20 ml chromatography column. The flowthrough containing purified LacI was collected. An additional 5 ml dialysis buffer was applied to the column and the flowthrough was also collected. The flowthrough fractions were combined and spin-concentrated to a final concentration of 10 µM. A small volume of the final LacI sample was run on a denaturing SDS-PAGE gel to confirm the protein size and purity (>95%). The oligomerization state was confirmed by size-exclusion chromatography and protein secondary structure assessed by circular dichroism.

### Sample preparation for HDX/MS

Exchange buffer (50 mM MOPS, 200 mM NaCl, 0.5 mM TCEP, pH 7.0) was prepared in deionized water. To make a D_2_O exchange buffer, the H_2_O exchange buffer was lyophilized and resuspended in the same volume of D_2_O three times. LacI samples were prepared at 10 µM in 100 µl H_2_O exchange buffer, with a small molecule ligand, DNA operator, or both added to separate protein samples at the following concentrations: IPTG, 10 µM; ONPF, 3 mM; TMG, 10 µM; DNA operator, 25 µM. These ligand concentrations were chosen to be saturating and to maintain solubility. Samples were equilibrated at 4 °C with binding partners for at least 30 minutes before the HDX/MS experiment. Fully deuterated samples were prepared at 10 µM in D_2_O buffer with 8 M urea, diluted in D_2_O exchange buffer 1:10, incubated at room temperature for 16 hours, and lyophilized two times for at least 16 hours, adding back an equal volume of D_2_O each time. Then the fully deuterated samples were incubated at room temperature for an additional 24 hours.

### HDX/MS experimental protocol

All HDX labeling was performed in solution using a LEAP technologies HDX PAL robot (Carrboro, NC), with a temperature-controlled sample vials and buffer holder held at 4 °C and refrigerated compartments held at 1 °C. Samples were injected on a Thermo UltiMate 3000 LC that was connected in-line with an Q Exactive mass spectrometer equipped with an electrospray ionization source (Thermo Fisher Scientific). The samples were diluted 1:10 in the D_2_O exchange buffer (50 mM MOPS, 200 mM NaCl, 0.5 mM TCEP, pH 7.0) to allow for hydrogen atoms in the protein backbone amide groups to exchange for deuterons in the buffer. Aliquots of the diluted samples were collected at nine timepoints (0 sec, 30 sec, 45 sec, 1 min, 5 min, 25 min, 1 hr, 2 hr, 4 hr), quenched 1:1 in an acidic quench buffer (6 M urea, 200 mM arginine, 100 mM TCEP, pH 2.0), and cooled to 1 °C to slow the HDX reaction. This range of timepoints was chosen to allow for the measurement of fast and slow exchange events in all functional states of LacI. Each sample was then enzymatically proteolyzed on a Waters Enzymate™ BEH Pepsin Column (30 mm length × 2.1 mm diameter, 5 μm particle size, Waters Corporation, Milford, MA), or a combination of pepsin and fungal protease type XIII (manually conjugated to POROS™ 20 AL Aldehyde Activated Resin, Thermo Fisher Scientific, and packed in a 2 mm ID x 2 cm IDEX C-130B column, Fisher Scientific, Hampton, NH), loaded on a C18 analytical column (Hypersil Gold, 10 mm length × 2.1 mm diameter, 3 µm particle size, Thermo Fisher Scientific) with a guard column attached to the inlet (Thermo Fisher Scientific), and injected in a mass spectrometer using a 100 µL sample loop (Thermo Q Exactive) in solvent A (defined below). Desalting in sample A occurred over 2 minutes at 300 µl/min. Five independently generated exchange reactions (technical replicates) for all states were performed from four separately expressed and purified LacI samples (biological replicates) to ensure adequate technical and biological replicates, except for the TMG state, for which two biological replicates were performed. Back exchange was calculated on a peptide basis by conducting an additional HDX/MS experiment on an apo LacI “maximally labeled” or “fully deuterated” control sample (Masson et al., 2019; Zhang and Smith, 1993). The average deuteration for all fully deuterated peptides over the whole protein was 70% with a standard deviation of 15.7%, and 11.4% of fully deuterated peptides had 50% deuteration or less (44 out of 386 total peptides). In the curated set of 57 core domain peptides included in this study, the average deuteration for fully deuterated peptides was 70.5% with a standard deviation of 14.6% (Table S2).

Acetonitrile, formic acid (1 mL ampules, Fisher Optima grade, 99.9%), and MS-grade water (Thermo Fisher Scientific) were used to prepare mobile phase solvents. Solvent A was 99.9% water/0.1% formic acid and solvent B was 90% acetonitrile/10% water with 0.1% formic acid added (v/v). The elution program consisted of isocratic flow at 10% B for 4 min, a linear gradient to 45% B over 9 min, isocratic flow at 100% B for one min, isocratic flow at 10% B for one min, a linear gradient to 45% B over one min, isocratic flow at 100% B for one min, and then isocratic flow at 10% B for 3 min, at a flow rate of 20 mL/min. Full-scan mass spectra were acquired in the positive ion mode over the range m/z = 400 to 2000 using the Orbitrap mass analyzer, with a mass resolution setting of 70,000, AGC target of 1.00e6, and maximum IT set to 50 s. We also ran three tandem mass spectrometry (MS/MS) experiments for each sample with the same full MS settings as described above, and dd-MS2 settings as follows: resolution 17,500, AGC target 1e5, maximum IT 100 ms, loop count 6, isolation window 2.0 m/z, NCE 28, charge state 1 and ≥7 excluded, dynamic exclusion of 20 seconds. Data acquisition was controlled using Xcalibur software (version 4.1, Thermo Fisher Scientific).

### HDX/MS data analysis

For each sample, peptides were identified using combined MS/MS raw data by searching against the LacI amino acid sequence (UniProtKB, www.uniprot.org, accessed 6/18/2019) and the pepsin amino acid sequence using Proteome Discoverer software (version 1.3, SEQUEST algorithm, Thermo Fisher Scientific) or Byonic software (Protein Metrics, Cupertino, CA). The protein search configuration used in the search were: cleavage residues, FWYLI;FWYLI; digest cutter, N-terminal and C-terminal cutter; and spectrum-level FDR, autocut. In a typical sample MS/MS, 476 unique forward peptides were found; 1027 spectra were matched to forward peptides; coverage was 98.18% of the 330-amino acid dimeric LacI sequence; and estimated spectrum-level false-discovery rate (FDR) on true proteins was 0.0%. The following filters were applied when identifying peptides: minimum precursor mass, 350 Da; maximum precursor mass, 5000 Da; minimum peak count, 1; maximum collision energy, 1000; S/N threshold, 1.5; minimum peptide length, 4; maximum peptide length, 144; maximum number of peptides reported, 10; precursor mass tolerance, 10 ppm; and fragment mass tolerance, 0.02 Da. The exported peptide lists (sequence, charge state, and retention time) from separate samples were imported into HD Examiner 2.0 (Sierra Analytics, Modesto, CA) and combined. All peptide isotope distributions at each exchange timepoint were fit in HD Examiner and manually checked for correct assignments. For each peptide in each functional state, deuteration levels at each timepoint were calculated by subtracting mass centroids of undeuterated peptides (*m*_*0*_) from mass centroids of deuterated peptides (*m*_*t*_), dividing by the difference between the mass centroid of the fully deuterated peptide (*m*_*∞*_) and the undeuterated peptide, and multiplying by the theoretical maximum number of exchangeable protons (*max_protons*):

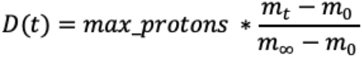

For each peptide, the charge state with the highest quality data as determined by HD Examiner was used. We note that the fully deuterated sample was in 100% D exchange buffer, while the experimental samples at each timepoint were in 90% D exchange buffer after 1:10 dilution, which would result in an increase in the NE and C parameters (described below).

Uptake plots were constructed by graphing the calculated deuteration levels against time for each peptide, with the maximum number of exchangeable protons as the limit on the *y* axis and the error bars showing the standard deviation of *n* = 2 to 5 replicates. The uptake data were fit to a nonlinear regression model using the SciPy optimize.curve_fit function. All calculations for fractional exchange as shown in Fig. 2B and Fig. S7 were performed using averaged centroid data at each timepoint, not the fitting functions; the nonlinear regression function fits are shown as lines in uptake plots to guide the eye. The nonlinear regression function was:

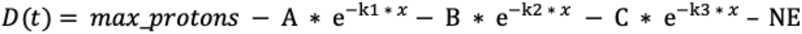

where

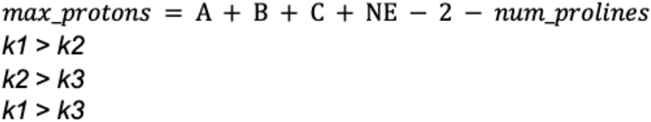

*D(t)*: number of deuterons exchanged for protons as a function of time

*max_protons*: theoretical maximum number of exchangeable protons for the peptide

*A, B, C*: number of fast-, medium-, and slow-exchanging protons, respectively

*k1, k2, k3*: rates of exchange for fast-, medium-, and slow-exchanging protons, respectively

*NE*: number of non-exchanging protons

*num_prolines*: number of prolines in the peptide

For slow-exchanging peptides in which the last timepoint had an experimentally observed value of less than 0.5 D, data were fit to a modified regression function with A = 0 and k1 = 0.

The fractional difference in H-D exchange at all timepoints and the associated uncertainty were calculated as described previously using averaged centroid data (Weis, 2021). Appendix 1 contains the calculated parameters for H-D exchange fitting to experimental data points for all peptides, corresponding to each line for the uptake plots in Fig. S3. Appendix 2 contains the calculated uncertainties for the differential exchange plots shown in Fig. S7. Appendix 3 contains the raw MS centroids for all replicates at each timepoint for each peptide and functional state.

### Computational prediction of structural water molecules

To predict the positions of structural water molecules, we applied a semi-explicit solvation method implemented in the macromolecular modeling program Rosetta (Pavlovicz et al., 2020) on each pre-relaxed LacI crystal structure (PDB 2PAF, 2P9H, 1LBI) using the beta_nov16 energy function. Predicted structural water molecules within a 6 Å shell of any atom of the ligand were retained. All hydrogen bonds within a heavy-atom distance of 3.2 Angstroms were calculated between water oxygen atoms and protein or ligand residues. The probability of hydrogen bonding was calculated as the number of times that a residue could form a hydrogen bond with a predicted structural water molecule out of 100 total output structures. See the Supplemental computational methods for the detailed protocol, scripts, and command lines.

## Acknowledgements

We thank members of the Kortemme and Marqusee Labs at UC San Francisco and UC Berkeley for discussions, the Raman Lab at the University of Wisconsin, Madison for the *E. coli* Δ*lacI*:220 strain and the pZE22G-GFP-kan and pSC101-LacI expression vectors, Shawn Costello for suggestions on the manuscript, and Monita Muralidharan for advice on HDX/MS data analysis. Funding: This work was supported by grants from the National Institutes of Health (K99GM135529 to AG and R01GM050945 to SM), and fellowships from the UCSF IRACDA program (K12GM081266) and the UC Office of the President through the UCSF Chancellor’s Fellowship to AG. ZP was supported by the Rosetta Commons Research Experience for Undergraduates Summer Internship program (NSF Award 1950697). SM and TK are Chan Zuckerberg Biohub Investigators.

## Author contributions

AG conceived the idea for the project, developed the core questions, and performed the majority of the experimental studies and analysis. HTH and AG carried out the HDX/MS experiments. ZRP performed the Rosetta simulations under the guidance of AG. SM and TK provided discussion, mentorship, and resources. AG wrote the manuscript with contributions from the other authors.

