## Supplemental information for "Ligand-induced changes in dynamics mediate long-range allostery in the *lac* repressor"

#### Supplemental computational methods: command lines, input files and scripts

We used Rosetta 3.9 (version 2019.19) to predict structural water molecule placement, as described in the Methods.

##### (1) Preparation of LacI structures for solvation

Relax command line:
~/Path/to/rosetta/source/bin/relax.linuxgccrelease -database ~/Path/to/rosetta//database -s INPUT.pdb -nstruct 2 -constrain_relax_to_start_coords -coord_constrain_sidechains -beta -ex1 -ex2 -use_input_sc -flip_HNQ -no_optH false -optimization::default_max_cycles 200

##### (2) Semi-explicit solvation

Command line to solvate the relaxed LacI structure 2P9H using the beta_nov16 energy function and output 100 solvated structures:

~/Desktop/Rosetta/rosetta/main/source/bin/rosetta_scripts.macosclangrelease @flags_NOWATERS.txt -parser:protocol solvate_NOpack.xml -database ~/Desktop/Rosetta/rosetta/main/database -in:file:s 2p9h_R_align_relax.pdb -out:suffix _solvate

Example XML file implementing semi-explicit solvation (Pavlovicz et al., 2020) without packing:

```
<ROSETTASCRIPTS>
  <SCOREFXNS>
          <ScoreFunction name="beta" weights="beta_nov16"/>
     </SCOREFXNS>
     <TASKOPERATIONS>
          <PreventRepacking name="prevent_repacking" resnum="62A-330A,62B-330B"/>
     </TASKOPERATIONS>
     <MOVERS>
          <PackRotamersMover
               name="pack"
               scorefxn="beta"
               task_operations="prevent_repacking"/>
          <WaterBoxMover
               name="solvate"
               mode="replace"
               gen_fixed="1"
               scorefxn="beta"
               task_operations="prevent_repacking"/>
     </MOVERS>
     <PROTOCOLS>
          <Add mover_name="solvate"/>
```

```
            <Add mover_name="pack"/>
      </PROTOCOLS>
</ROSETTASCRIPTS>
```

Example flags file to specify input file, packing parameters, energy function, and number of output structures:

```
-in
  -file
    -s 2p9h_R_r.pdb
-packing
  -ex1
  -ex1aro
  -extrachi_cutoff 0
  -ex2
-nstruct 100
-beta_nov16
-parser
  -view
-mute core.util.prof
-mute core.io.database
```

**(3) Downstream analysis and WaterMap generation performed in Python and PyMOL.**

All Python scripts are available on Github (https://github.com/anumazam/watermaps_analysis_scripts):

delete_waters.py
bin_waters.py
hbnets.py
run_hbnets.py
data2bfactor.py
color_b.py

**Figure S1. X-ray crystal structures of LacI.** ONPF-DNA-LacI (PDB 1EFA, gray, resolution 2.6 Å) (Bell and Lewis, 2000) and IPTG-LacI (PDB ID 2P9H, red, resolution 2.0 Å) (Daber et al., 2007) states are aligned to show a small shift towards a more “pinched” conformation of the N-terminal subdomain of the core.

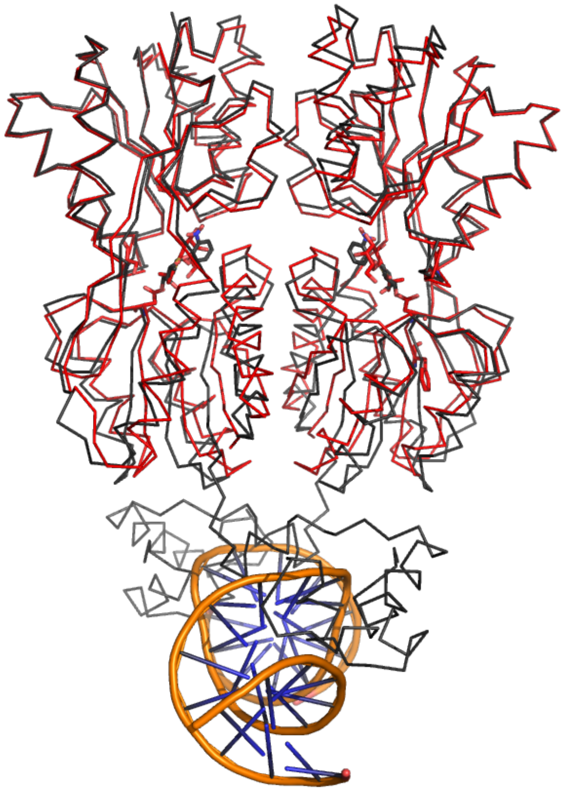

**Figure S2. Summary of HDX/MS peptide coverage for the curated set of peptides for the core domain of LacI. (A)** Coverage map showing individual peptides (gray bars) by amino acid position (small squares) in the core domain (residues 63-330). Blue boxes mark every tenth residue in each row. Brown boxes mark residues that were not present in any of the peptides. The extreme C-terminal sequence of the core domain pertaining to beta strands 20-21 (as labeled in Fig. 2) and the loop connecting them were not included in the dataset because of incomplete or noisy data for at least one of the functional states in this region of the protein. Excluding this sequence, HDX/MS coverage of the core domain was 97% with several overlapping peptides for some regions. **(B)** Structure of the core domain (PDB ID 2P9H) with brown residues corresponding to brown amino acid positions in (A). **(C)** Peptide statistics. The majority of peptides (65%) were 4 to 9 residues in length. Peptides were evenly distributed between the N- and C-terminal subdomains of the core.

**A** **HDX/MS peptide coverage in the curated dataset for the LacI core domain**

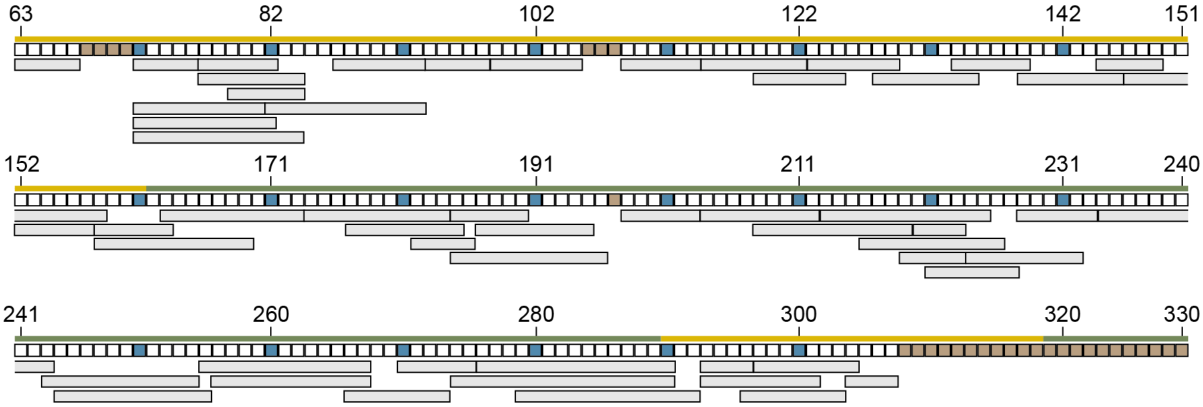


**B**

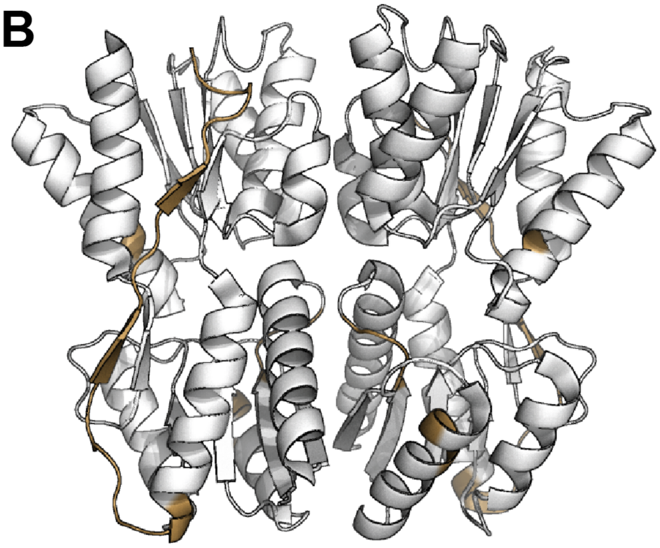

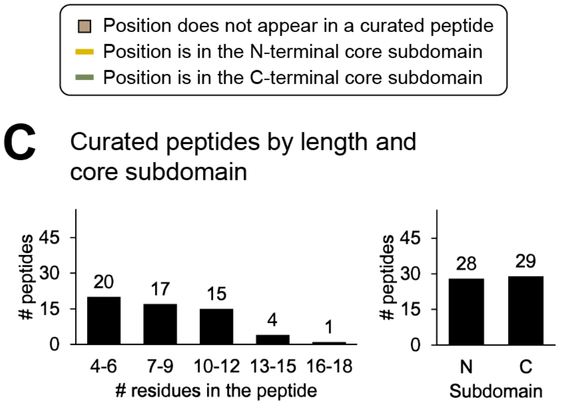


| | |
|---|---|
| Total number of curated peptides | 57 |
| Length of peptide (# residues), average | 8.5 |
| Length of peptide, standard deviation | 3.1 |

**Figure S3. Uptake plots for 57 LacI peptides in six functional states.** Plots are organized from N- to C-terminus of LacI. Gray labels in the corner of the plots indicate the LacI secondary structure element(s) represented in the peptide, as in Fig. 2.

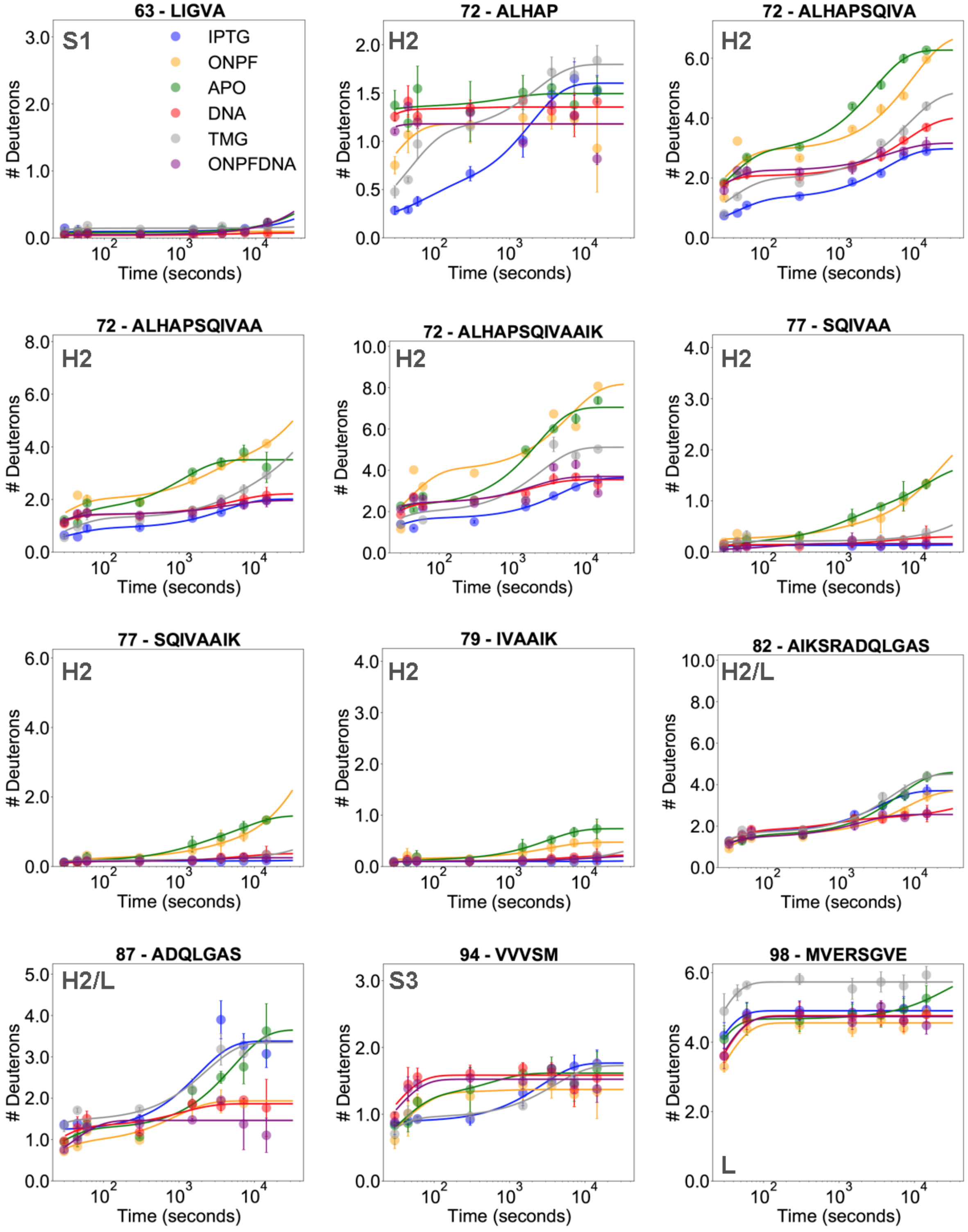

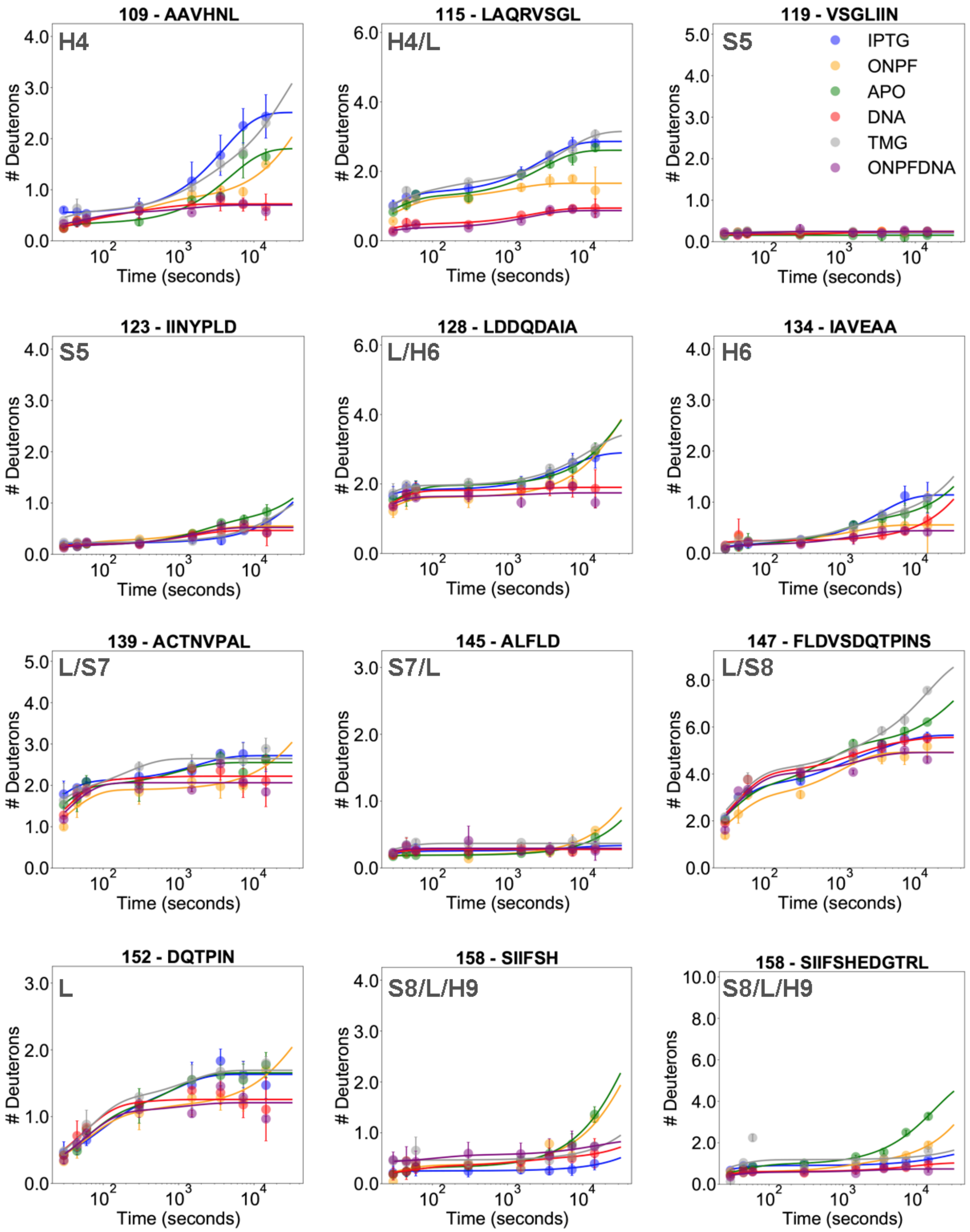
109 - AAVHNL
H4
115 - LAQRVSGL
H4/L
119 - VSGLIIN
S5
IPTG
ONPF
APO
DNA
TMG
ONPFDNA
123 - IINYPLD
S5
128 - LDDQDAIA
L/H6
134 - IAVEAA
H6
139 - ACTNVPAL
L/S7
145 - ALFLD
S7/L
147 - FLDVSDQTPINS
L/S8
152 - DQTPIN
L
158 - SIIFSH
S8/L/H9
158 - SIIFSHEDGTRL
S8/L/H9
### Deuterons
Time (seconds)

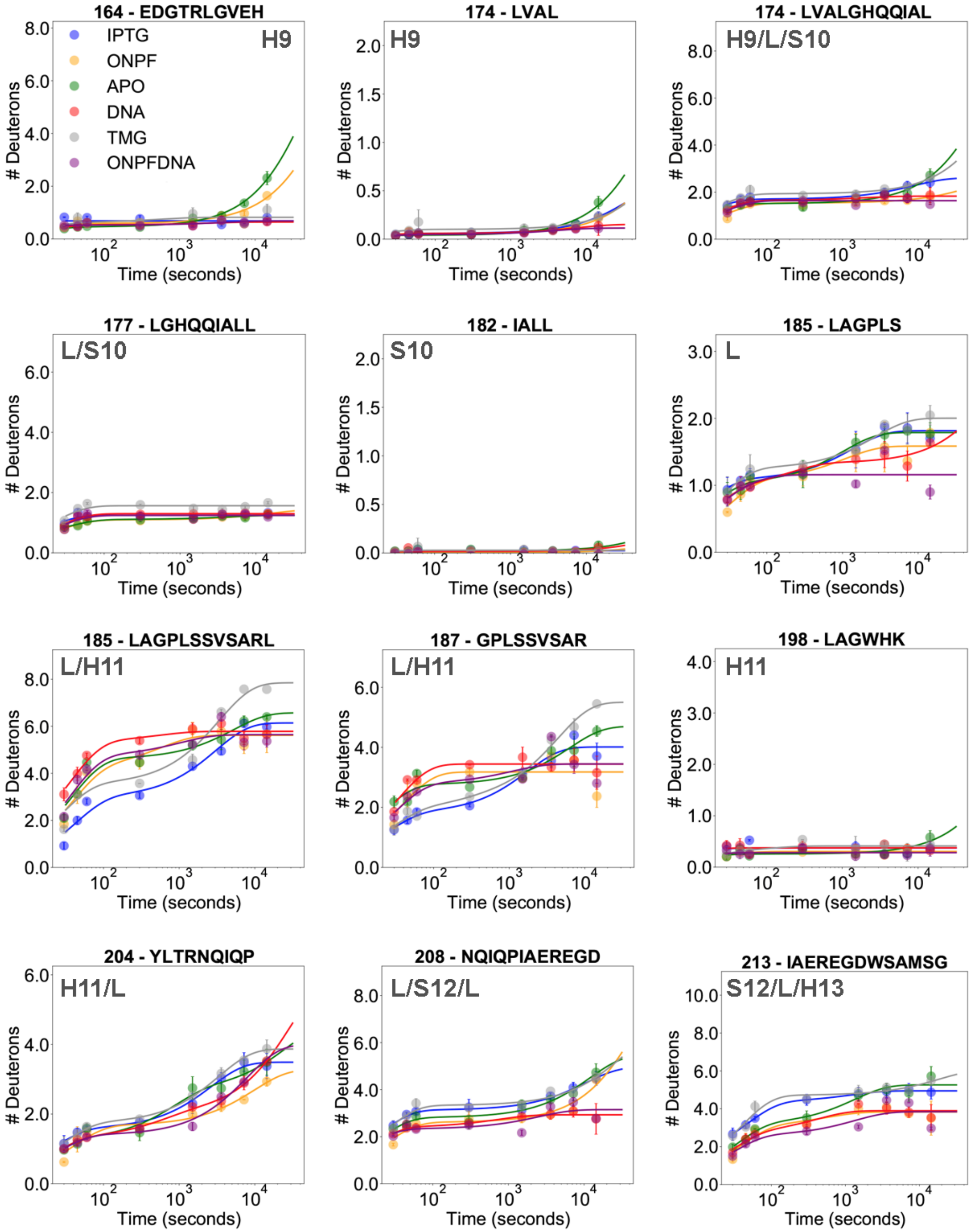

164 - EDGTRLGVEH
H9
IPTG
ONPF
APO
DNA
TMG
ONPFDNA
174 - LVAL
H9
174 - LVALGHQQIAL
H9/L/S10
177 - LGHQQIALL
L/S10
182 - IALL
S10
185 - LAGPLS
L
185 - LAGPLSSVSARL
L/H11
187 - GPLSSVSAR
L/H11
198 - LAGWHK
H11
204 - YLTRNQIQP
H11/L
208 - NQIQPIAEREGD
L/S12/L
213 - IAEREGDWSAMSG
S12/L/H13
### Deuterons
Time (seconds)

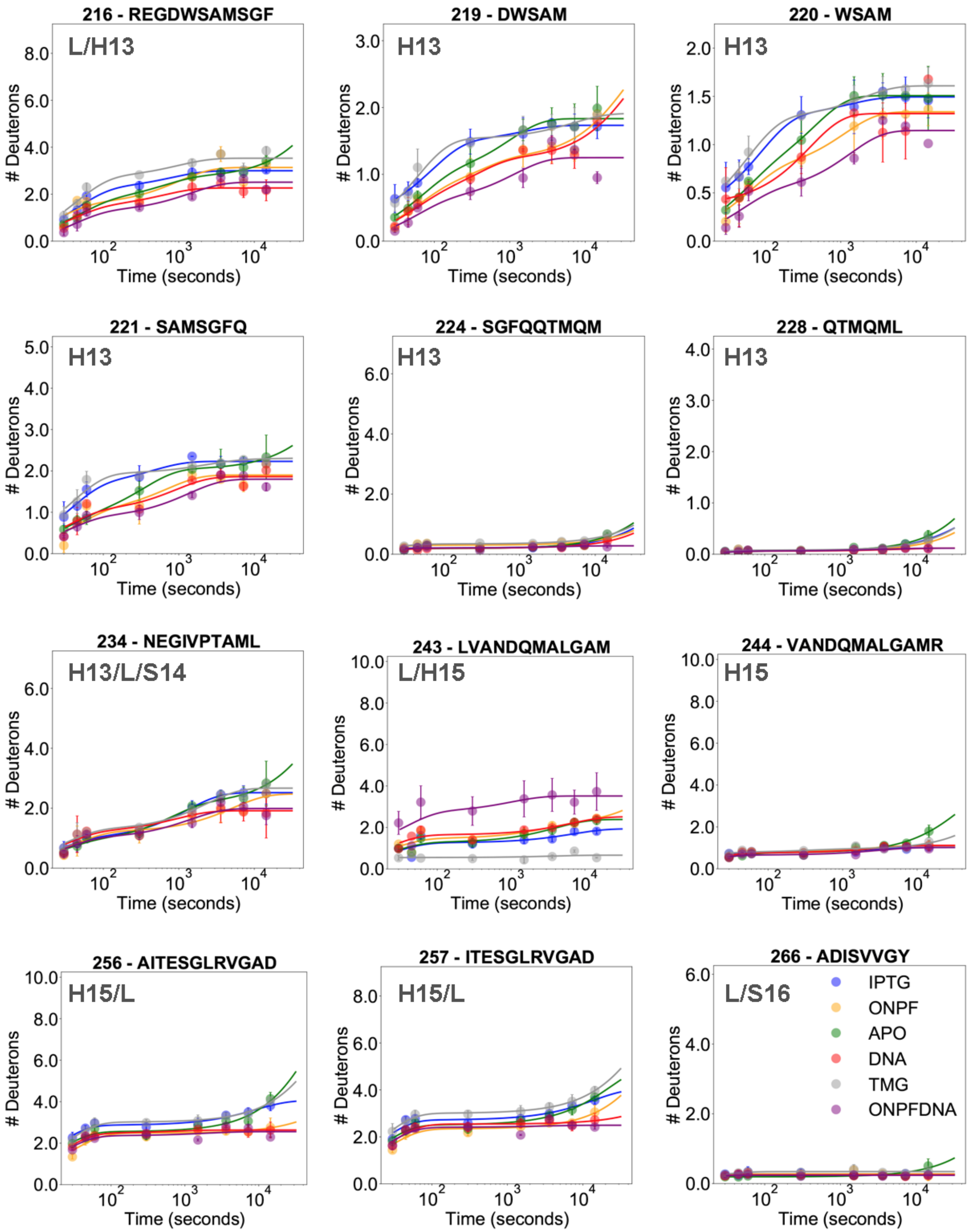

216 - REGDWSAMSGF
L/H13
219 - DWSAM
H13
220 - WSAM
H13
221 - SAMSGFQ
H13
224 - SGFQQTMQM
H13
228 - QTMQML
H13
234 - NEGIVPTAML
H13/L/S14
243 - LVANDQMALGAM
L/H15
244 - VANDQMALGAMR
H15
256 - AITESGLRVGAD
H15/L
257 - ITESGLRVGAD
H15/L
266 - ADISVVGY
L/S16
### Deuterons
Time (seconds)
IPTG
ONPF
APO
DNA
TMG
ONPFDNA

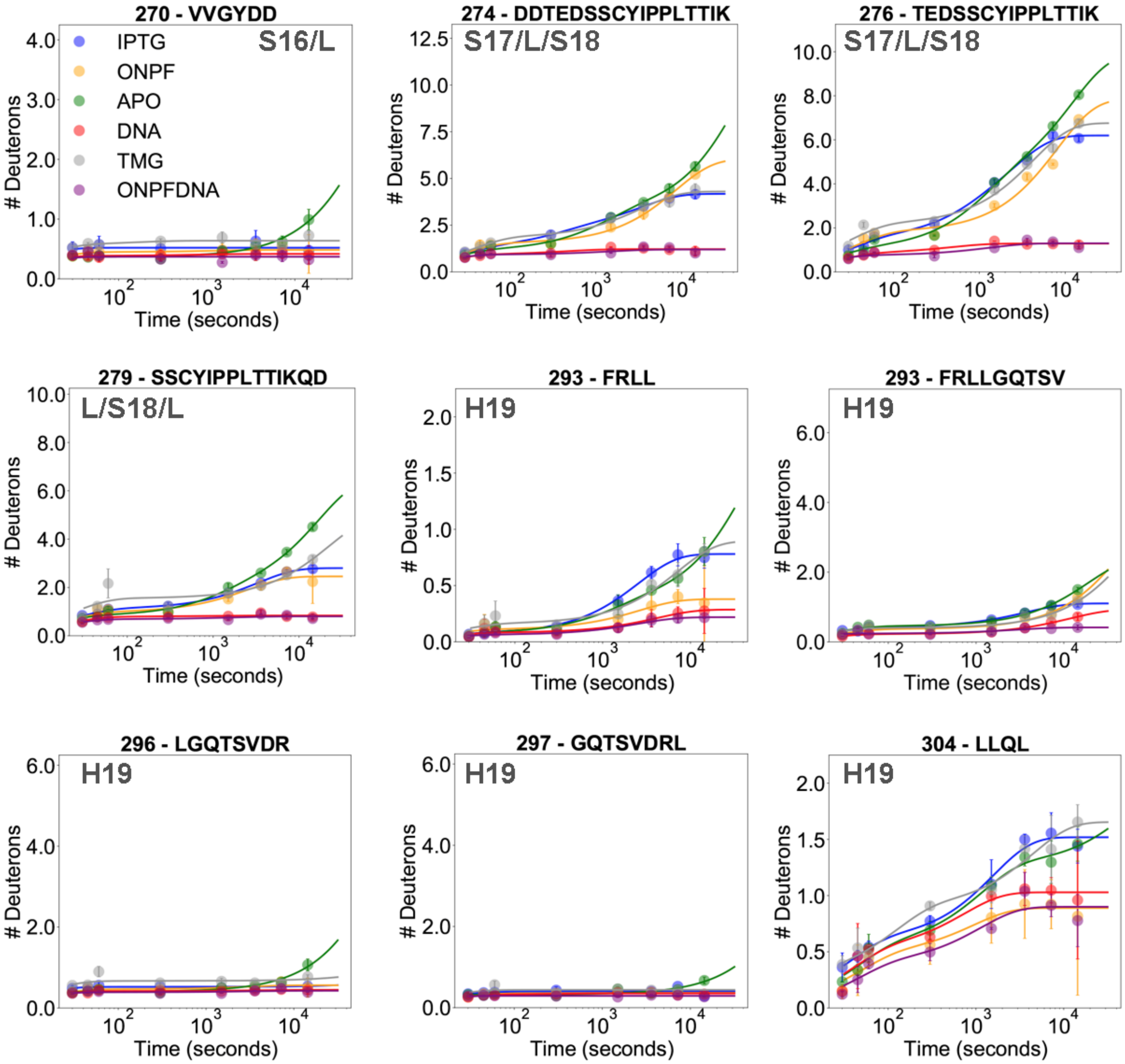
270 - VVGYDD
S16/L
IPTG
ONPF
APO
DNA
TMG
ONPFDNA
274 - DDTEDSSCYIPPLTTIK
S17/L/S18
276 - TEDSSCYIPPLTTIK
S17/L/S18
279 - SSCYIPPLTTIKQD
L/S18/L
293 - FRLL
H19
293 - FRLLGQTSV
H19
296 - LGQTSVDR
H19
297 - GQTSVDRL
H19
304 - LLQL
H19
### Deuterons
Time (seconds)

**Figure S4. Example mass spectra for all states for peptide 115 - LAQRVSGL.**

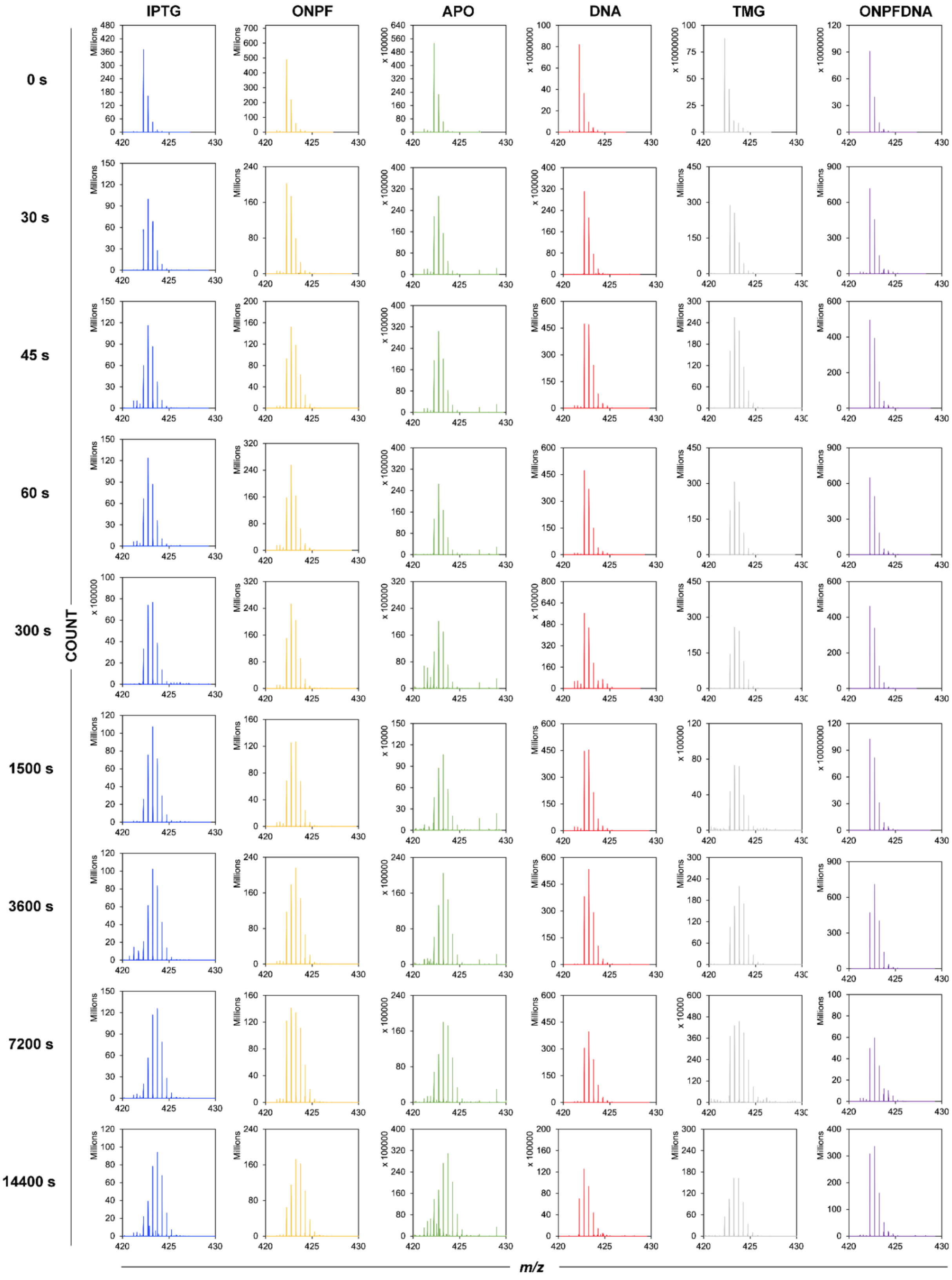

**Figure S5. H-D exchange at interface helices.** Peptides in the structural models for LacI on the left are colored green, magenta and navy blue to match the heading (peptide starting position and sequence) of the respective deuterium uptake plots on the right. **(A)** H-D exchange in helix 2 at the N-terminal subdomain monomer-monomer interface of LacI (PDB ID 2P9H). The N-terminal region of the helix (green) shows decreased exchange in IPTG-LacI (blue line), whereas the C-terminal region of the helix (navy blue) shows decreased exchange in DNA-LacI (red line). The central region of the helix (magenta) is rigid in IPTG-LacI and DNA-LacI, but shows increased exchange in ONPF-LacI (orange line). **(B)** The outward-facing alpha helices 13 (green) and 17 (navy blue) at the C-terminal subdomain monomer-monomer interface of LacI show decreased exchange in DNA-LacI as compared to IPTG-LacI.

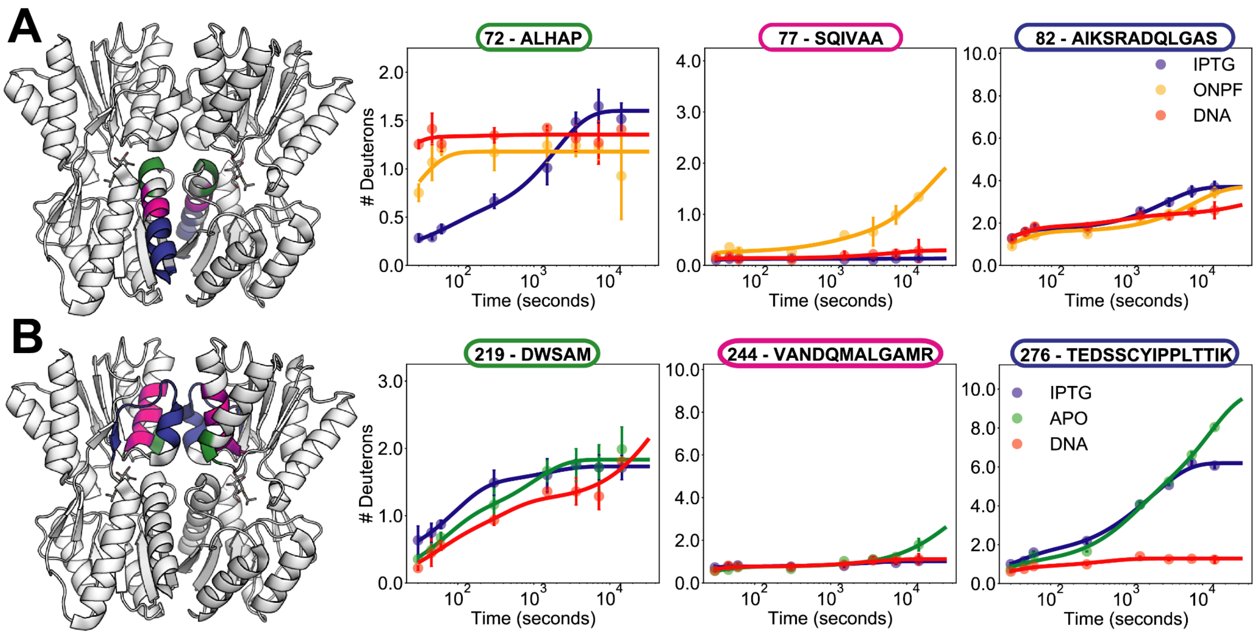

**Figure S6. Reduced inter-beta-sheet distance in the N-terminal subdomain monomer-monomer interface in IPTG-LacI compared to DNA-LacI. (A)** A water molecule bridges the distance between beta strand residues in each subunit in the structure of DNA-LacI (PDB ID 1EFA). **(B)** In IPTG-LacI, valines in the interface beta strands on each subunit form direct backbone-backbone hydrogen bonds (PDB ID 2P9H).

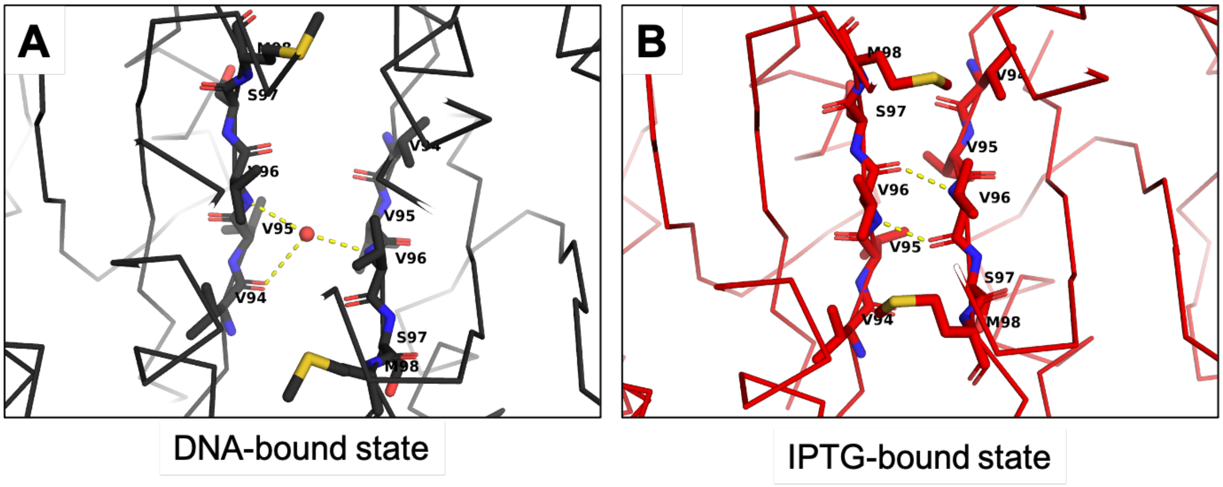

**Figure S7. Fractional differences in H-D exchange between LacI states at single HX timepoints based on maximum theoretical exchange and corrected for back-exchange.** The fractional difference was calculated as $\Delta HX_t = \Delta D_t / (N_H m_\delta)$, where $\Delta D_t = m_{A,t} - m_{B,t}$ (the difference between averaged centroids at time t for states A and B); $N_H$ is the number of exchangeable amide hydrogens in the peptide (total length - 2 - number of prolines); and $m_\delta \approx 1$ Da, the difference between the mass of deuterium and protium. The heading of each heatmap shows the states compared in each heatmap and the direction of the comparison. For example, in the plot with the heading "APO-DNA," the blue boxes denote less exchange in apo-LacI, and the red boxes denote less exchange in DNA-LacI.

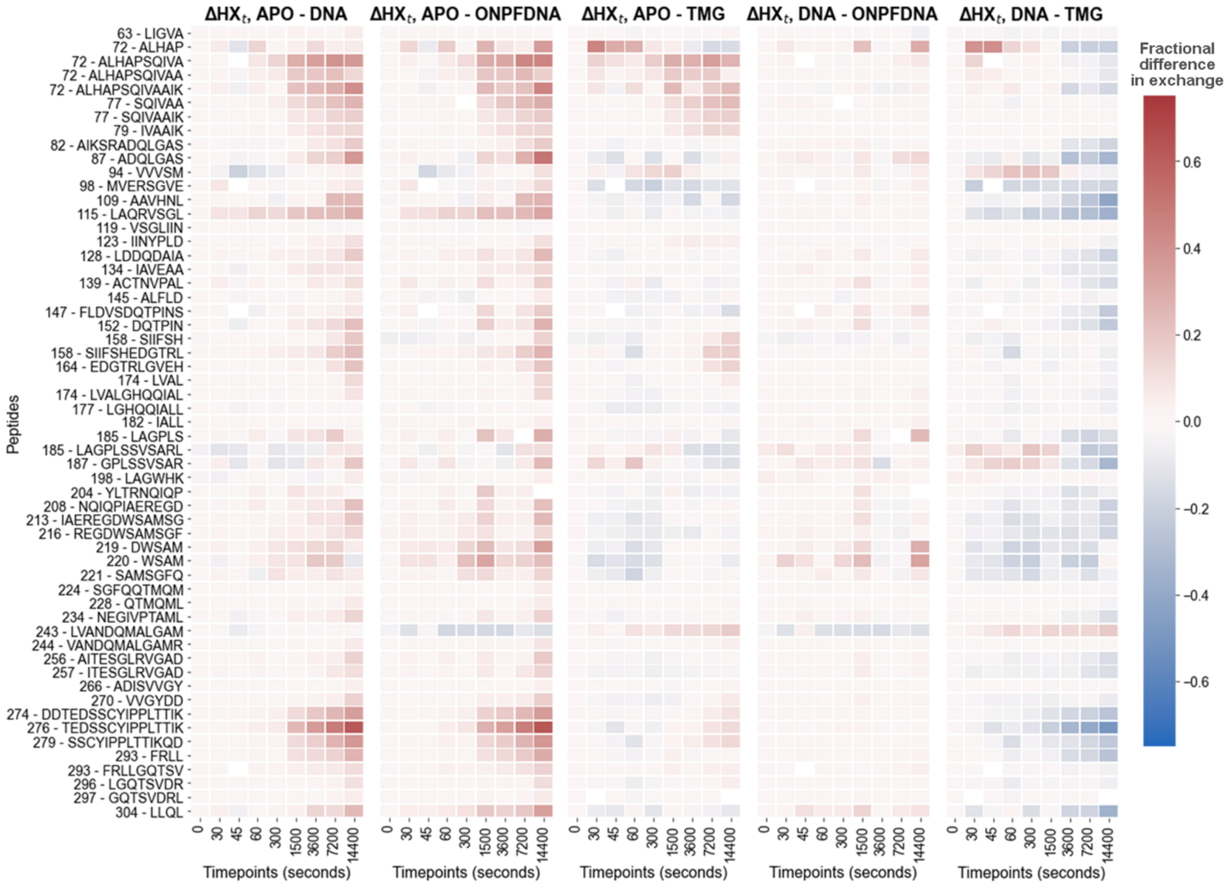

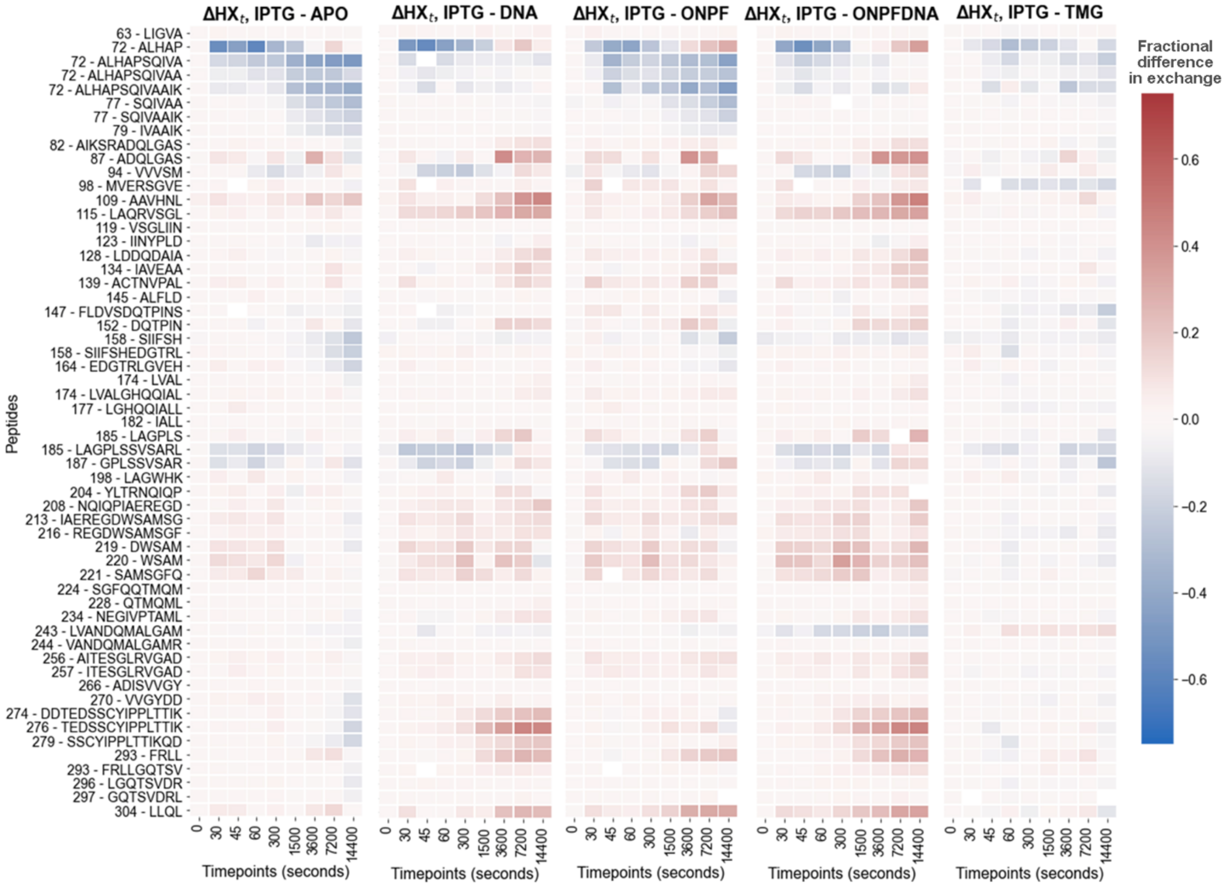

ΔHXt, IPTG - APO
ΔHXt, IPTG - DNA
ΔHXt, IPTG - ONPF
ΔHXt, IPTG - ONPFDNA
ΔHXt, IPTG - TMG
Fractional difference in exchange
Peptides
Timepoints (seconds)

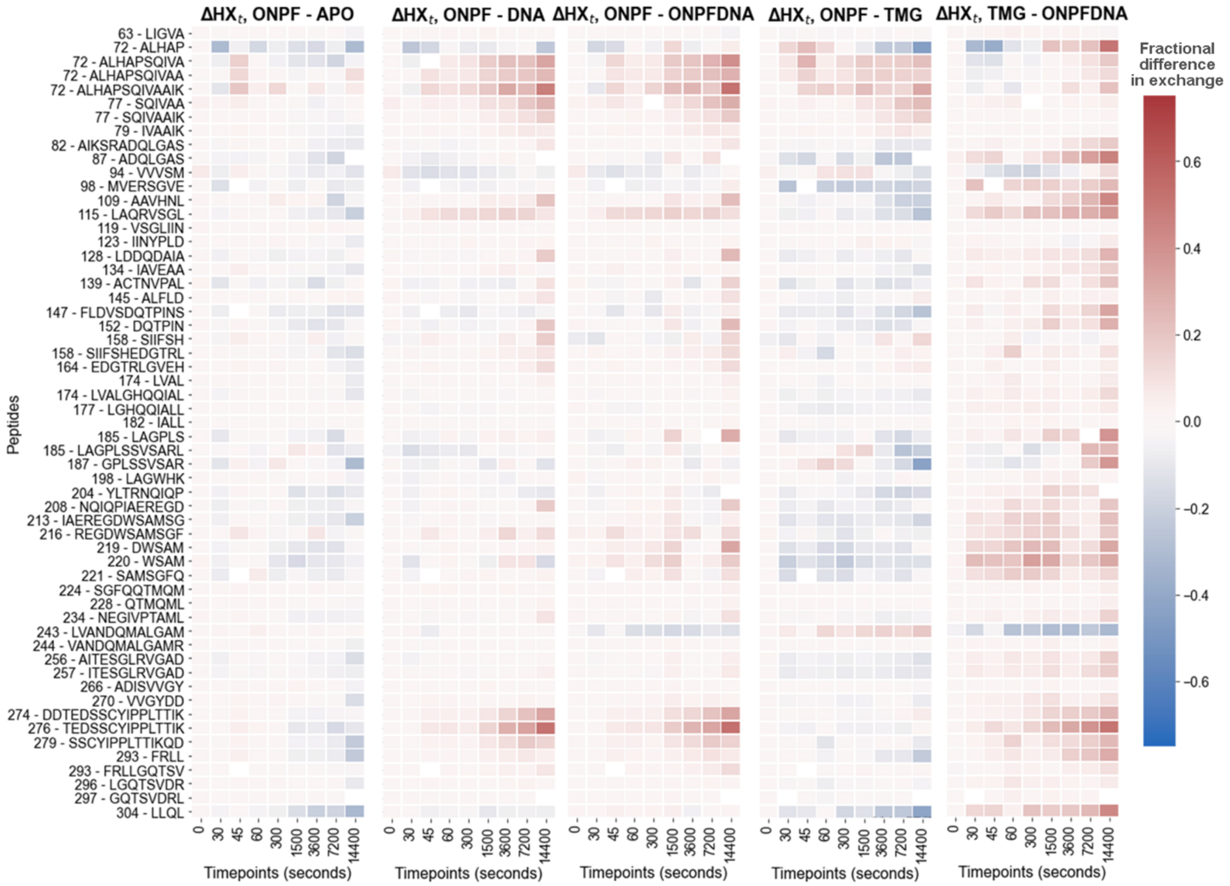

ΔHXt, ONPF - APO
ΔHXt, ONPF - DNA
ΔHXt, ONPF - ONPFDNA
ΔHXt, ONPF - TMG
ΔHXt, TMG - ONPFDNA
Peptides
63 - LIGVA
72 - ALHAP
72 - ALHAPSQIVA
72 - ALHAPSQIVAA
72 - ALHAPSQIVAAIK
77 - SQIVAA
77 - SQIVAAIK
79 - IVAAIK
82 - AIKSRADQLGAS
87 - ADQLGAS
94 - VVVSM
98 - MVERSGVE
109 - AAVHNL
115 - LAQRVSGL
119 - VSGLIIN
123 - IINYPLD
128 - LDDQDAIA
134 - IAVEAA
139 - ACTNVPAL
145 - ALFLD
147 - FLDVSDQTPINS
152 - DQTPIN
158 - SIIFSH
158 - SIIFSHEDGTRL
164 - EDGTRLGVEH
174 - LVAL
174 - LVALGHQQIAL
177 - LGHQQIALL
182 - IALL
185 - LAGPLS
185 - LAGPLSSVSARL
187 - GPLSSVSAR
198 - LAGWHK
204 - YLTRNQIQP
208 - NQIQPIAEREGD
213 - IAEREGDWSAMSG
216 - REGDWSAMSGF
219 - DWSAM
220 - WSAM
221 - SAMSGFQ
224 - SGFQQTMQM
228 - QTMQML
234 - NEGIVPTAML
243 - LVANDQMALGAM
244 - VANDQMALGAMR
256 - AITESGLRVGAD
257 - ITESGLRVGAD
266 - ADISVVGY
270 - VVGYDD
274 - DDTEDSSCYIPPLTTIK
276 - TEDSSCYIPPLTTIK
279 - SSCYIPPLTTIKQD
293 - FRLL
293 - FRLLGQTSV
296 - LGQTSVDR
297 - GQTSVDRL
304 - LLQL
0 30 45 60 300 1500 3600 7200 14400
Timepoints (seconds)
Fractional difference in exchange
0.6
0.4
0.2
0.0
−0.2
−0.4
−0.6

**Figure S8. LacI core domain colored according to effects of point mutations on gene expression.** Each residue position was classified in one of fifteen categories according to its position in the protein structure as well as its most common mutational phenotype. The structure is colored according to these classifications as shown in Table S1. Data from Suckow *et al.* (Suckow et al., 1996) (PDB ID: 2P9H.)

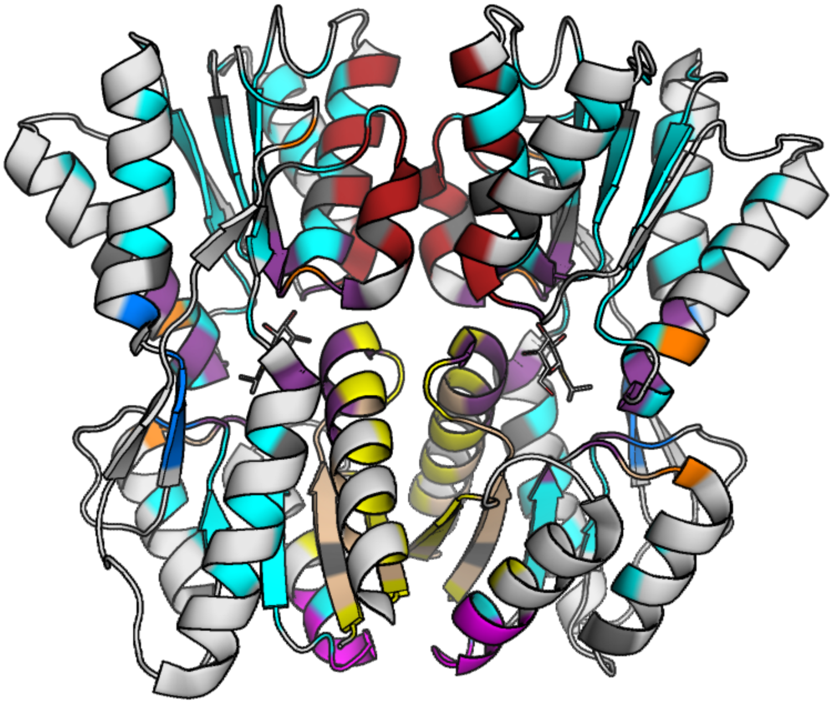

**Figure S9. Comparison of HDX/MS data to functional data from individual LacI mutants.** H-D exchange patterns can be compared to functional data for residues at the monomer-monomer interface **(A-C)**, which play a role in switching the conformational ensemble of LacI from the IPTG-bound state to the DNA-bound state, and in a pocket peripheral loop **(D)**, which modulates the inducer affinity. Structures are colored according to the color scheme in Figs. 2-3.

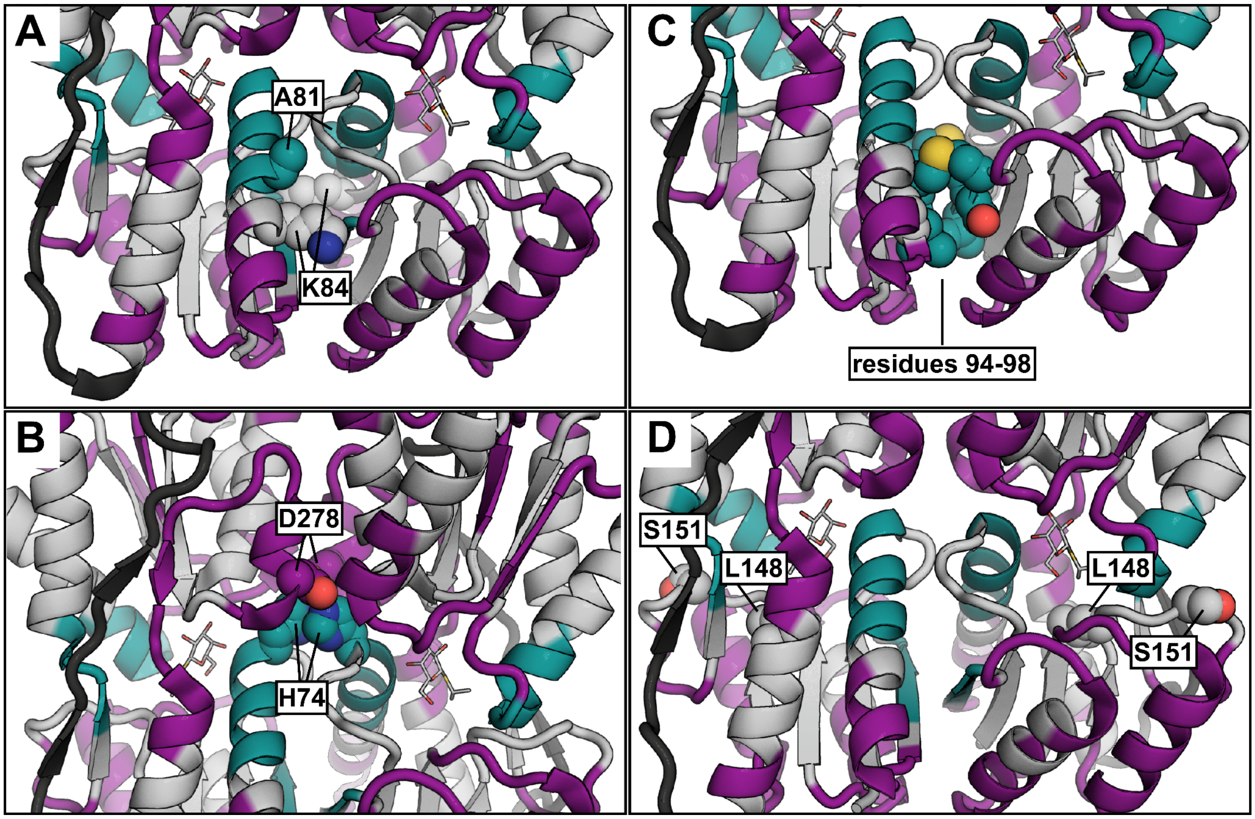

**Figure S10. Compiled HDX/MS data for the DNA-binding domain of LacI (residues 1-61). (A)** The DNA-binding domain has a helix-turn-helix structure when bound to DNA. One subunit is shown in color and the other is shown in gray. In DNA-LacI, the hinge regions (darkest blue) have a helical structure and interact with each other (PDB ID: 1EFA). **(B-C)** Woods plots show that DNA-LacI (red) and ONPF-LacI (orange) exchange less than IPTG-LacI (blue) in the DNA-binding domain over the time course.

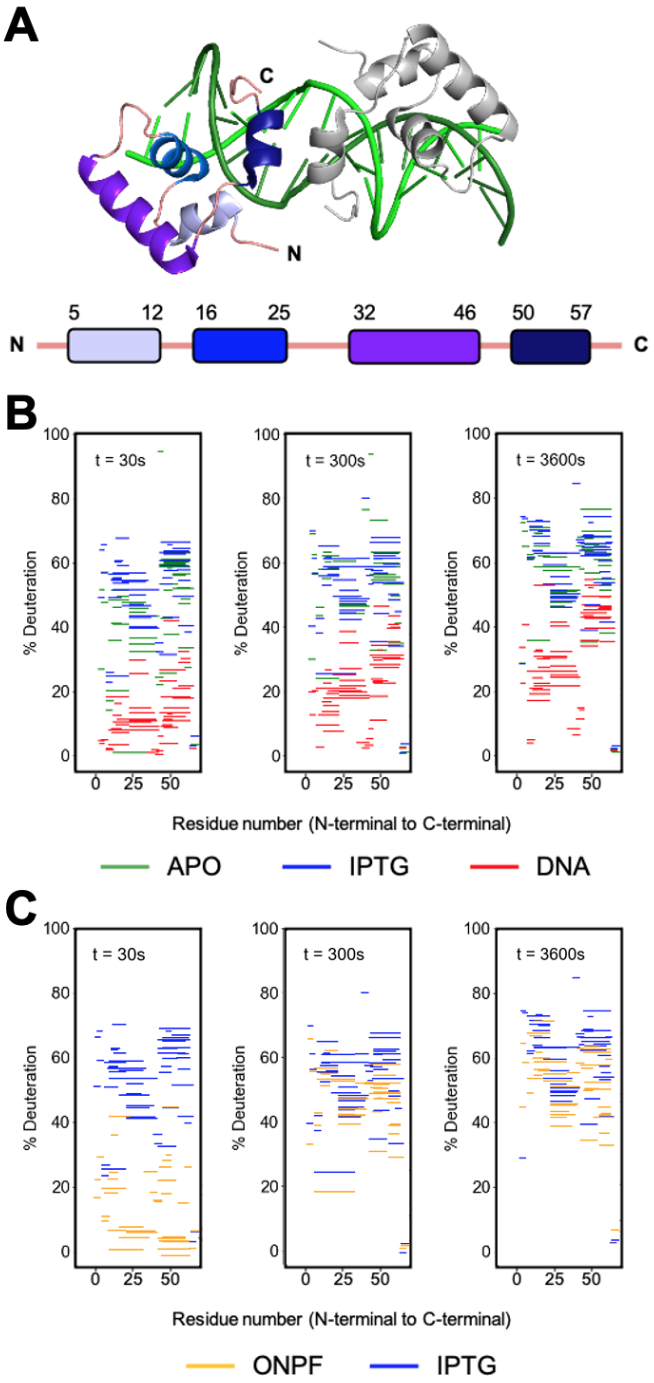

**Figure S11. Direct interactions of ligands with LacI residues in crystal structures.** Residues within 6 Å of any atom in the ligand are shown as sticks, with atoms that are within 3.5 Å of the ligand colored black, for the **(A)** ONPF-bound and **(B)** IPTG-bound states of LacI. Structures are colored as in the color scheme in Fig. 4. PDB 2PAF for (A) and 2P9H for (B).

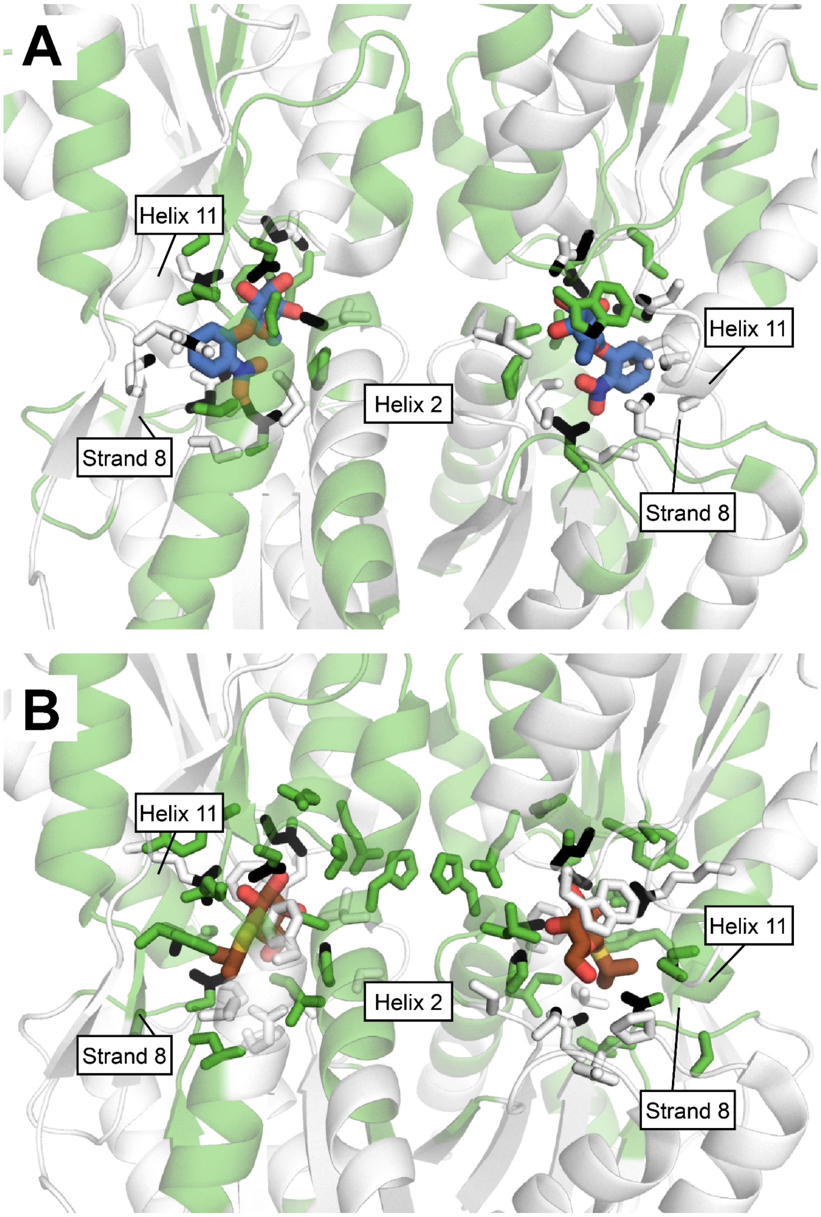

**Figures S12. Computationally predicted structural waters. (A)** Representative models showing the placement of structural water molecules, which make 3-4 hydrogen bonds with ligand or protein atoms, in the ligand binding pocket of LacI. A semi-explicit water molecule placement protocol implemented in Rosetta (Pavlovicz et al., 2020) was used to solvate apo-LacI, DNA-LacI, and IPTG-LacI (Methods). LacI models were based on X-ray crystal structures (PDB IDs: 1LBI, 2PAF, 2P9H) (Daber et al., 2007; Lewis et al., 1996). **(B)** Several predicted structural water molecules are within 0.5 Å of assigned water molecules (structural and transient) in the X-ray crystal structure for a high-resolution (2.0 Å) structure of IPTG-LacI.

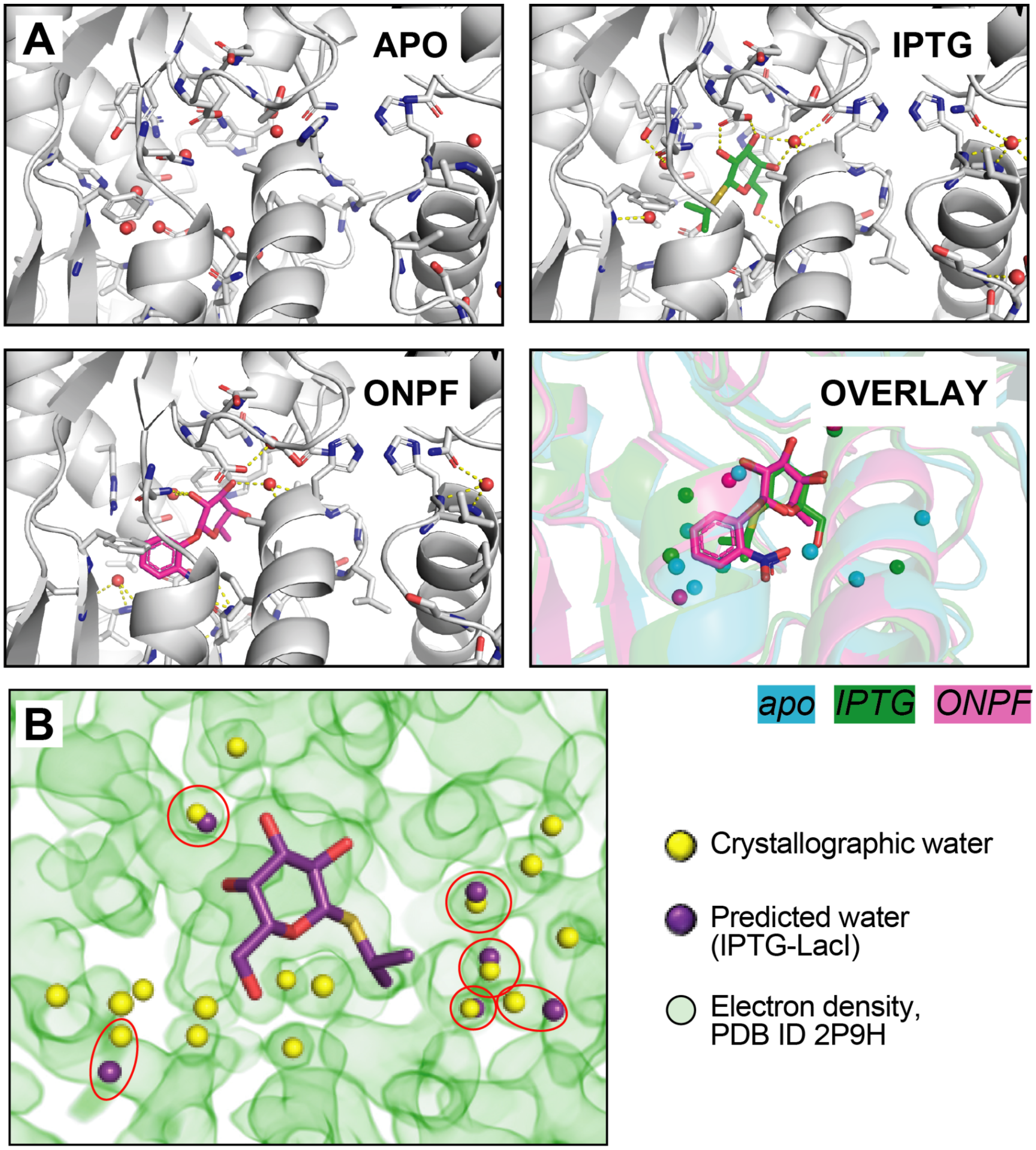

**Figure S13. Frequency of residue interactions with predicted structural water molecules in different ligand-bound states of LacI. (A)** Percentage of simulations in which a predicted structural water molecule is observed within 6 Å of any ligand or protein atom in the binding pocket. Data is shown for both LacI subunits, with and without allowing sidechain repacking (change in rotamer conformation). **(B)** Structural models of apo-LacI, IPTG-LacI, and ONPF-LacI from structural water placement simulations without sidechain repacking (top) and with sidechain repacking (bottom). Sidechains in the ligand binding pocket are colored according to the probability that they make a hydrogen bond with a predicted structural water molecule.

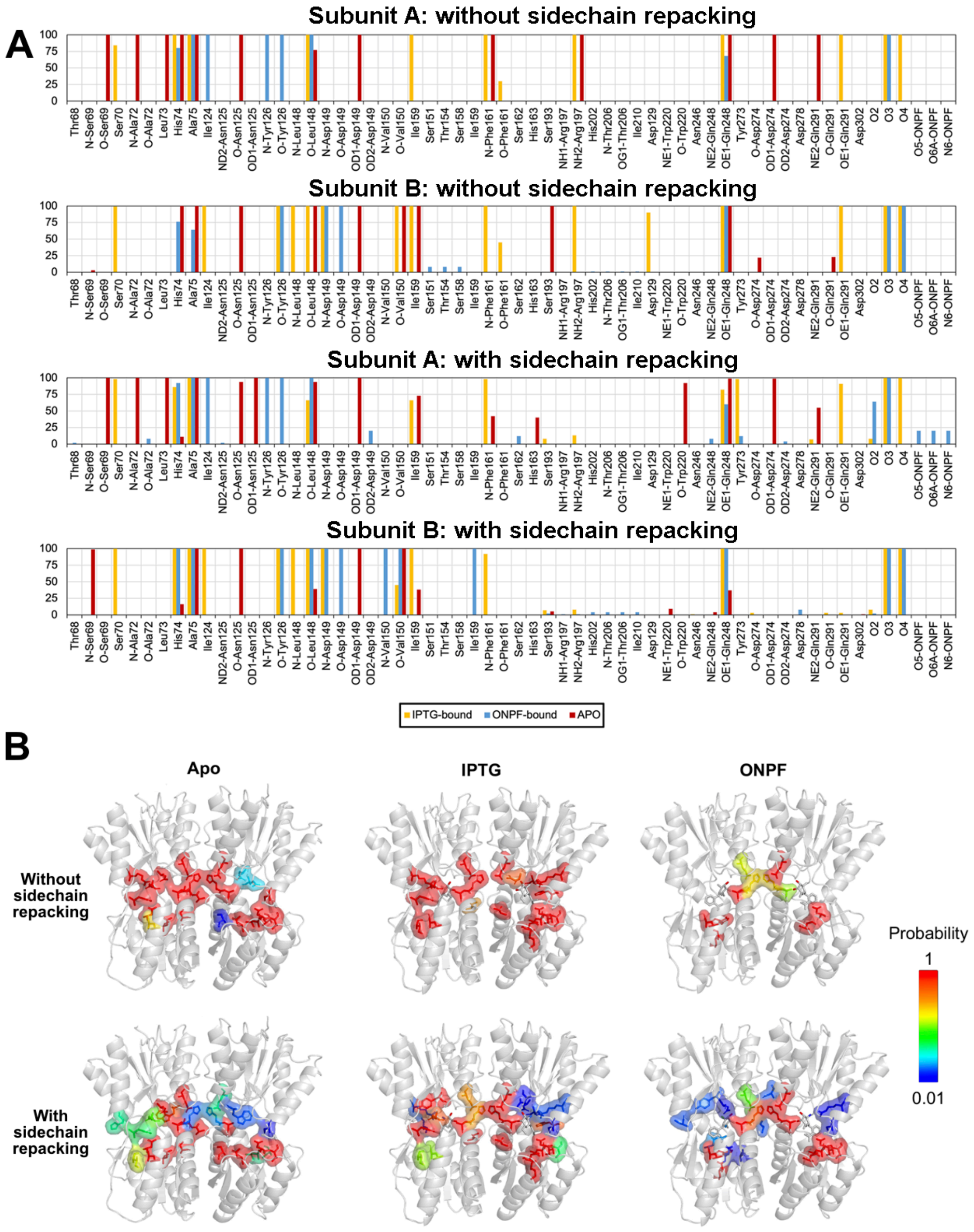

**Table S1. Mutational phenotypes in LacI.** Classification scheme for effects on gene expression of point mutations in the core domain of LacI from Suckow *et al*. (Suckow et al., 1996).

| Group | Location, type of amino acid | Remarks |
|---|---|---|
| **1** | Solvent exposed | Mostly tolerant to substitutions |
| **2** | Spacer regions | Tolerant to substitutions; Ala stretch tolerated |
| **3** | Buried | Tolerant to substitutions |
| **4** | Solvent exposed | Intolerant to substitutions: involved in salt bridges |
| **5** | Buried and small | Only small amino acids are tolerated ($I^-$) |
| **6** | Solvent exposed and small | Preferably small amino acids are tolerated |
| **7** | Buried | Intolerant to substitutions ($I^-$) |
| **8** | C-terminal subdomain and monomer-monomer interface | Intolerant to substitutions; substitutions result in $I^-$ phenotype |
| **9** | DNA contacts | Intolerant to substitutions $I^-$ |
| **10** | Stabilization of the helix-turn-helix motif in the DNA binding domain | Conserved amino acids $I^-$ |
| **11** | IPTG contacts | Substitutions result in $I^s$ phenotype |
| **12** | N-terminal subdomain and monomer-monomer interface | Substitutions result in $I^s$ phenotype |
| **13** | N-terminal subdomain of the core, near the hinge connecting both core domains | Substitutions result in $I^s$ phenotype |
| **14** | N-terminal subdomain of the core, near the center of the domain | Substitutions result in weak $I^s$ phenotype |
| **15** | Interface between DNA binding domain A and core B, or the interface between DNA binding domains A and B | Intolerant to substitutions |

**Table S2. Fully deuterated LacI peptides.** Percent deuteration observed in maximally labeled HDX/MS controls for the 57 peptides in the curated peptide dataset.

| First residue | Last residue | Peptide | %D |
|---|---|---|---|
| 63 | 67 | LIGVA | 62.375 |
| 72 | 76 | ALHAP | 53.456 |
| 72 | 81 | ALHAPSQIVA | 69.843 |
| 72 | 82 | ALHAPSQIVAA | 66.865 |
| 72 | 84 | ALHAPSQIVAAIK | 71.838 |
| 77 | 82 | SQIVAA | 82.949 |
| 77 | 84 | SQIVAAIK | 88.284 |
| 79 | 84 | IVAAIK | 91.593 |
| 82 | 93 | AIKSRADQLGAS | 62.077 |
| 87 | 93 | ADQLGAS | 57.699 |
| 94 | 98 | VVVSM | 82.723 |
| 98 | 105 | MVERSGVE | 52.327 |
| 109 | 114 | AAVHNL | 54.412 |
| 115 | 122 | LAQRVSGL | 74.479 |
| 119 | 125 | VSGLIIN | 84.658 |
| 123 | 129 | IINYPLD | 75.582 |
| 128 | 135 | LDDQDAIA | 54.55 |
| 134 | 139 | IAVEAA | 104.619 |
| 139 | 146 | ACTNVPAL | 71.379 |
| 145 | 149 | ALFLD | 64.994 |
| 147 | 158 | FLDVSDQTPINS | 79.973 |
| 152 | 157 | DQTPIN | 79.801 |
| 158 | 163 | SIIFSH | 62.602 |
| 158 | 169 | SIIFSHEDGTRL | 32.071 |
| 164 | 173 | EDGTRLGVEH | 54.708 |
| 174 | 177 | LVAL | 82.243 |
| 174 | 184 | LVALGHQQIAL | 69.995 |
| 177 | 185 | LGHQQIALL | 60.627 |
| 182 | 185 | IALL | 92.705 |

| | | | |
|---|---|---|---|
| 185 | 190 | LAGPLS | 71.843 |
| 185 | 196 | LAGPLSSVSARL | 46.157 |
| 187 | 195 | GPLSSVSAR | 77.371 |
| 198 | 203 | LAGWHK | 50.69 |
| 204 | 212 | YLTRNQIQP | 85.554 |
| 208 | 219 | NQIQPIAEREGD | 71.614 |
| 213 | 225 | IAEREGDWSAMSG | 59.832 |
| 216 | 226 | REGDWSAMSGF | 60.662 |
| 219 | 223 | DWSAM | 71.97 |
| 220 | 223 | WSAM | 81.576 |
| 221 | 227 | SAMSGFQ | 78.404 |
| 224 | 232 | SGFQQTMQM | 67.03 |
| 228 | 233 | QTMQML | 82.819 |
| 234 | 243 | NEGIVPTAML | 83.32 |
| 243 | 254 | LVANDQMALGAM | 43.976 |
| 244 | 255 | VANDQMALGAMR | 73.363 |
| 256 | 267 | AITESGLRVGAD | 68.841 |
| 257 | 267 | ITESGLRVGAD | 66.687 |
| 266 | 273 | ADISVVGY | 86.737 |
| 270 | 275 | VVGYDD | 48.3 |
| 274 | 290 | DDTEDSSCYIPPLTTIK | 97.302 |
| 276 | 290 | TEDSSCYIPPLTTIK | 43.726 |
| 279 | 292 | SSCYIPPLTTIKQD | 79.388 |
| 293 | 296 | FRLL | 82.69 |
| 293 | 301 | FRLLGQTSV | 79.012 |
| 296 | 303 | LGQTSVDR | 68.554 |
| 297 | 304 | GQTSVDRL | 70.732 |
| 304 | 307 | LLQL | 80.627 |
| | | Average | 70.52989 |
| | | Standard deviation | 14.61508 |
