## Appendix 1 for "Ligand-induced changes in dynamics mediate long-range allostery in the *lac* repressor"

**Appendix 1. Fitted HDX parameters for 57 LacI peptides in 6 functional states.** Nonlinear regression parameters and errors were calculated for deuterium incorporation in each peptide over 9 timepoints (0 s, 30 s, 45 s, 60 s, 300 s, 1500 s, 3600 s, 7200 s, 14400 s) with back-exchange correction, as described in the Methods, with *n* = 2-5 for each peptide.

**LIGVA**

| | IPTG | ONPF | APO | DNA | TMG | ONPFDNA |
|---|---|---|---|---|---|---|
| **A** | 0 | 0 | 0 | 0 | 0 | 0 |
| **B** | 0.115463185 | 0.10024544 | 0.084606856 | 0.057214699 | 0.126436209 | 0.060692427 |
| **C** | 2.903053752 | 0.021290284 | 2.919684509 | 0.036125307 | 2.853918913 | 2.951084938 |
| **k1** | 0 | 0 | 0 | 0 | 0 | 0 |
| **k2** | 0.1 | 0.044754511 | 0.1 | 0.1 | 0.060004711 | 0.1 |
| **k3** | 2.06E-06 | 0.000630601 | 3.16E-06 | 0.000178678 | 2.15E-07 | 3.90E-06 |
| **NE** | 1.07E-09 | 2.901638814 | 1.92E-15 | 2.923818648 | 1.85E-09 | 2.54E-13 |

**ALHAP**

| | IPTG | ONPF | APO | DNA | TMG | ONPFDNA |
|---|---|---|---|---|---|---|
| **A** | 0.123193928 | 0.006262936 | 1.238802193 | 1.131581426 | 0.006546529 | 1.181419608 |
| **B** | 0.351875901 | 1.172436108 | 0.11524941 | 0.183472391 | 1.094517786 | 7.25E-07 |
| **C** | 1.126203602 | 3.36E-10 | 0.139649633 | 0.025229198 | 0.700671853 | 9.97E-34 |
| **k1** | 0.524464625 | 0.04363626 | 0.533346564 | 0.106267903 | 0.000600259 | 0.121281617 |
| **k2** | 0.014963257 | 0.043605679 | 0.054518936 | 0.1 | 0.022049753 | 0.1 |
| **k3** | 0.000519497 | 0.009982929 | 0.001141383 | 0.003490958 | 0.000444242 | 0.009999927 |
| **NE** | 0.398726564 | 0.821997051 | 0.506305495 | 0.645219376 | 0.204051819 | 0.819977149 |

**ALHAPSQIVA**

| | IPTG | ONPF | APO | DNA | TMG | ONPFDNA |
|---|---|---|---|---|---|---|
| **A** | 0.608610214 | 1.456936105 | 1.135677283 | 1.040043545 | 1.002925004 | 0.009847622 |
| **B** | 0.689746335 | 1.577539849 | 1.647584895 | 0.997625866 | 1.010614997 | 2.290805372 |
| **C** | 1.67564032 | 3.847833538 | 3.443422379 | 1.97016621 | 2.927034271 | 0.9269073 |
| **k1** | 0.025531109 | 0.039488113 | 0.033201389 | 0.070622063 | 0.031176988 | 0.052819182 |
| **k2** | 0.025534119 | 0.039488938 | 0.033209768 | 0.070622662 | 0.031180283 | 0.05275461 |
| **k3** | 0.000235924 | 0.000105089 | 0.000340287 | 0.000124227 | 0.00012261 | 0.00020674 |
| **NE** | 4.026432816 | 0.243396721 | 0.731273023 | 2.974413159 | 2.127368912 | 3.838758762 |

**ALHAPSQIVAA**

| | IPTG | ONPF | APO | DNA | TMG | ONPFDNA |
|---|---|---|---|---|---|---|
| **A** | 0.115927425 | 1.235830587 | 1.03E-14 | 0.731672612 | 5.806072679 | 2.14E-20 |
| **B** | 0.785434444 | 2.018917173 | 1.544045727 | 0.68026695 | 1.354085478 | 1.435818163 |
| **C** | 1.117347634 | 4.781128946 | 1.958549635 | 0.796386112 | 0.907194978 | 0.528713045 |
| **k1** | 0.581306417 | 0.000461237 | 1.96E-08 | 0.058609689 | 1.00E-05 | 0.068994538 |
| **k2** | 0.031246548 | 0.045103819 | 0.042840175 | 0.058604254 | 0.033399343 | 0.063122291 |
| **k3** | 0.000250091 | 1.44E-05 | 0.000886733 | 0.000194576 | 0.00020307 | 0.00022259 |
| **NE** | 5.985075983 | 2.36E-06 | 4.490601757 | 5.787843993 | 5.84E-05 | 6.038582298 |

**ALHAPSQIVAAIK**

| | IPTG | ONPF | APO | DNA | TMG | ONPFDNA |
|---|---|---|---|---|---|---|
| **A** | 5.70E-08 | 1.211780028 | 1.489704313 | 0.037474194 | 0.97793931 | 1.172953579 |
| **B** | 1.667429187 | 2.87128947 | 0.771586853 | 2.461995998 | 1.029258773 | 1.275651649 |
| **C** | 1.97475263 | 4.20117811 | 4.801947359 | 1.12421584 | 3.173918309 | 1.297695788 |
| **k1** | 3.07E-13 | 0.028196061 | 0.695009352 | 0.062672971 | 0.0638817 | 0.090041492 |
| **k2** | 0.056266605 | 0.028184805 | 0.075704363 | 0.062676722 | 0.06386423 | 0.087466182 |
| **k3** | 0.000217541 | 0.000169244 | 0.000462763 | 0.000545103 | 0.000412208 | 0.000530332 |
| **NE** | 6.371634737 | 1.820169237 | 2.954439825 | 6.456075672 | 4.886353701 | 6.301700233 |

**SQIVAA**

| | IPTG | ONPF | APO | DNA | TMG | ONPFDNA |
|---|---|---|---|---|---|---|
| **A** | 0 | 1.903763247 | 0.18313929 | 0 | 0 | 0 |
| **B** | 0.115957623 | 0.099598123 | 0.502763023 | 0.175895504 | 0.166778701 | 0.083152763 |
| **C** | 2.654925357 | 0.172133353 | 1.009579505 | 0.155604193 | 3.788948354 | 0.052574738 |
| **k1** | 0 | 4.55E-05 | 0.028969449 | 0 | 0 | 0 |
| **k2** | 0.061657834 | 0.05819577 | 0.000929625 | 0.1 | 0.062244996 | 0.007090137 |
| **k3** | 9.54E-08 | 0.001306893 | 7.04E-05 | 0.000163912 | 2.72E-06 | 0.007090759 |
| **NE** | 1.2156987 | 1.665641177 | 2.304169448 | 3.704015553 | 7.46E-05 | 3.836956277 |

**SQIVAAIK**

| | IPTG | ONPF | APO | DNA | TMG | ONPFDNA |
|---|---|---|---|---|---|---|
| **A** | 0 | 5.537306063 | 0.351152842 | 0 | 0 | 0 |
| **B** | 0.143640165 | 0.225000924 | 0.163409705 | 0.16920706 | 0.195456788 | 0.140838974 |
| **C** | 5.066022292 | 0.230048155 | 0.952347961 | 0.221048546 | 5.815410109 | 0.109109728 |
| **k1** | 0 | 1.17E-05 | 0.000931962 | 0 | 0 | 0 |
| **k2** | 0.051301531 | 0.032640504 | 0.040861788 | 0.039116795 | 0.058349578 | 0.070265966 |
| **k3** | 1.13E-07 | 0.000626699 | 0.000143032 | 0.000144413 | 1.59E-06 | 0.000364743 |
| **NE** | 0.778685584 | 1.59E-07 | 4.540086417 | 5.630376788 | 1.94E-05 | 5.750595319 |

**IVAAIK**

| | IPTG | ONPF | APO | DNA | TMG | ONPFDNA |
|---|---|---|---|---|---|---|
| **A** | 0 | 0 | 0.052942939 | 0 | 0 | 0 |
| **B** | 0.085508162 | 0.1328334 | 0.069818316 | 0.115968651 | 0.102891276 | 0.085070831 |
| **C** | 2.060572909 | 0.31938008 | 0.616697563 | 0.108795413 | 3.88914118 | 0.105973012 |
| **k1** | 0 | 0 | 0.041627817 | 0 | 0 | 0 |
| **k2** | 0.1 | 0.058227506 | 0.041623648 | 0.068043244 | 0.063996414 | 0.056323651 |
| **k3** | 1.31E-07 | 0.000304008 | 0.000309073 | 0.000138481 | 1.50E-06 | 7.33E-05 |
| **NE** | 1.841098281 | 3.52605758 | 3.262407428 | 3.780085069 | 2.23E-11 | 3.792308186 |

**AIKSRADQLGAS**

| | IPTG | ONPF | APO | DNA | TMG | ONPFDNA |
|---|---|---|---|---|---|---|
| **A** | 0.796523583 | 0.764854755 | 0.410636859 | 0.573195865 | 0.878778056 | 0.092981211 |
| **B** | 0.909725471 | 0.81332074 | 1.148168485 | 1.803028645 | 0.801106255 | 1.392101106 |
| **C** | 2.015315075 | 2.119531443 | 3.040286173 | 1.070211474 | 2.837112475 | 1.107085914 |
| **k1** | 0.052151284 | 0.034877505 | 0.923020065 | 0.00117518 | 0.056479014 | 0.053037015 |
| **k2** | 0.052152345 | 0.034878026 | 0.038469576 | 0.045574613 | 0.056464872 | 0.05303494 |
| **k3** | 0.00031234 | 0.000131416 | 0.000165173 | 1.96E-05 | 0.00020577 | 0.000571319 |
| **NE** | 6.292882309 | 6.290874352 | 5.400908483 | 6.58451683 | 5.490919886 | 7.440012269 |

**ADQLGAS**

| | IPTG | ONPF | APO | DNA | TMG | ONPFDNA |
|---|---|---|---|---|---|---|
| **A** | 1.277310383 | 0.476842341 | 0.400900755 | 0.744298362 | 0.696158716 | 0.094652199 |
| **B** | 6.72E-21 | 0.508540949 | 0.850047764 | 0.614103824 | 0.686466283 | 1.492752577 |
| **C** | 2.165714948 | 1.01827824 | 2.402999313 | 0.565743345 | 1.896636338 | 2.16E-05 |
| **k1** | 0.613894278 | 0.056524963 | 0.048922639 | 0.058147887 | 0.094863476 | 0.029203938 |
| **k2** | 0.099999996 | 0.05651833 | 0.048921939 | 0.058153603 | 0.094861878 | 0.029212727 |
| **k3** | 0.000552794 | 0.001108472 | 0.0001914 | 0.000914279 | 0.000438481 | 0.009984332 |
| **NE** | 1.61901846 | 3.065369674 | 1.347819149 | 3.133102958 | 1.649722921 | 3.535664345 |

**VVVSM**

| | IPTG | ONPF | APO | DNA | TMG | ONPFDNA |
|---|---|---|---|---|---|---|
| **A** | 0.889374411 | 0.053846039 | 0.055554334 | 7.51E-17 | 0.439497915 | 0.750022816 |
| **B** | 3.24E-10 | 1.147233562 | 1.173432691 | 1.597353486 | 0.444208983 | 0.790756436 |
| **C** | 0.879287144 | 0.000178266 | 0.386837832 | 4.98E-14 | 0.782149771 | 5.61E-07 |
| **k1** | 0.109287069 | 0.001762475 | 0.705808477 | 0.420444554 | 0.055715628 | 0.038865706 |
| **k2** | 3.02E-05 | 0.02542359 | 0.031876867 | 0.041649717 | 0.055716125 | 0.038858113 |
| **k3** | 0.000370352 | 0.009999014 | 0.001823584 | 0.01 | 0.000250272 | 0.01 |
| **NE** | 1.233946022 | 1.628900383 | 1.384175143 | 1.41511241 | 1.270642131 | 1.476138362 |

**MVERSGVE**

| | IPTG | ONPF | APO | DNA | TMG | ONPFDNA |
|---|---|---|---|---|---|---|
| **A** | 9.54E-18 | 3.94E-15 | 2.02E-24 | 2.05E-14 | 3.866699762 | 3.94E-17 |
| **B** | 4.905780467 | 4.503340327 | 4.647058998 | 4.771765542 | 1.805249269 | 4.764014171 |
| **C** | 7.78E-09 | 1.47E-12 | 1.336214495 | 4.48E-12 | 1.90E-07 | 7.71E-21 |
| **k1** | 0.354801242 | 0.076670814 | 0.005239193 | 2.96E-07 | 0.064181297 | 0.519134169 |
| **k2** | 0.064752254 | 0.044340463 | 0.070397855 | 0.048695019 | 0.064175535 | 0.051104873 |
| **k3** | 0.01 | 0.01 | 3.74E-05 | 0.01 | 0.009999985 | 0.01 |
| **NE** | 1.09744742 | 1.446622774 | 6.23E-17 | 1.239604972 | 0.268546386 | 1.264383838 |

**AAVHNL**

| | IPTG | ONPF | APO | DNA | TMG | ONPFDNA |
|---|---|---|---|---|---|---|
| **A** | 0.373970342 | 0.30056858 | 0.627652463 | 0.294829105 | 0.510661242 | 0.04566784 |
| **B** | 3.76E-17 | 0.408254562 | 0.333422263 | 0.191088711 | 0.616100213 | 0.365927749 |
| **C** | 1.972108503 | 3.214145864 | 0.847341923 | 0.196558799 | 2.832108186 | 0.170024572 |
| **k1** | 0.838801221 | 0.037395362 | 0.00020367 | 0.03323381 | 0.051234275 | 0.528135611 |
| **k2** | 0.097025636 | 0.002465809 | 0.059949222 | 0.007780216 | 0.00063238 | 0.019592558 |
| **k3** | 0.00025262 | 1.54E-05 | 0.000203578 | 0.001660453 | 3.52E-05 | 0.000760475 |
| **NE** | 1.487373452 | 1.64E-05 | 2.194568839 | 3.27775267 | 1.30E-19 | 3.300575611 |

**LAQRVSGL**

| | IPTG | ONPF | APO | DNA | TMG | ONPFDNA |
|---|---|---|---|---|---|---|
| **A** | 0.598162865 | 0.539963789 | 0.618603179 | 0.203425174 | 0.444046563 | 0.17551411 |
| **B** | 0.746344426 | 0.642884777 | 0.629120888 | 0.470578527 | 1.159581308 | 0.173256373 |
| **C** | 1.494121085 | 0.457168101 | 1.358474769 | 0.265758191 | 1.53810156 | 0.51212521 |
| **k1** | 0.046263771 | 0.03431201 | 0.040964084 | 0.000461825 | 0.009576998 | 0.056828975 |
| **k2** | 0.046259652 | 0.034309149 | 0.040967333 | 0.049553999 | 0.053722125 | 0.05682989 |
| **k3** | 0.000350193 | 0.000873771 | 0.000344046 | 0.000461875 | 0.00017795 | 0.000533649 |
| **NE** | 3.139432539 | 4.343436037 | 3.393865419 | 5.058401656 | 2.846823552 | 5.129212223 |

**VSGLIIN**

| | IPTG | ONPF | APO | DNA | TMG | ONPFDNA |
|---|---|---|---|---|---|---|
| **A** | 0 | 0 | 0 | 0 | 0 | 0 |
| **B** | 0.189154947 | 0.135414781 | 0.156185885 | 0.112710688 | 0.223787881 | 0.079469029 |
| **C** | 2.47E-08 | 3.64E-11 | 1.49E-12 | 0.073868366 | 4.67E-12 | 0.052615274 |
| **k1** | 0 | 0 | 0 | 0 | 0 | 0 |
| **k2** | 0.061834741 | 0.056566463 | 0.1 | 0.1 | 0.045156806 | 0.1 |
| **k3** | 5.02E-26 | 1.35E-25 | 0.009999994 | 0.000507169 | 0.01 | 0.01 |
| **NE** | 4.783193112 | 4.771411086 | 4.843578023 | 4.749795297 | 4.756545192 | 4.752238941 |

**IINYPLD**

| | IPTG | ONPF | APO | DNA | TMG | ONPFDNA |
|---|---|---|---|---|---|---|
| **A** | 0.193663035 | 0.122805744 | 0.423201595 | 0 | 0.19536132 | 0 |
| **B** | 0.028125507 | 0.13149759 | 0.165586405 | 0.164718793 | 0.005033882 | 0.185218311 |
| **C** | 2.340592524 | 0.300047402 | 3.411208632 | 0.286916018 | 1.144755849 | 0.358274546 |
| **k1** | 0.114391229 | 0.030009903 | 0.000513014 | 0 | 0.558051646 | 0 |
| **k2** | 0.012664453 | 0.030008838 | 0.080150217 | 0.081421847 | 0.017517334 | 0.058101251 |
| **k3** | 1.31E-05 | 0.000497073 | 5.00E-06 | 0.000706389 | 3.26E-05 | 0.000630873 |
| **NE** | 1.440635444 | 3.44970887 | 6.69E-13 | 3.534418856 | 2.624016089 | 3.475327386 |

**LDDQDAIA**

| | IPTG | ONPF | APO | DNA | TMG | ONPFDNA |
|---|---|---|---|---|---|---|
| **A** | 0.020020434 | 1.97E-24 | 0.689361077 | 0.093503677 | 0.87913265 | 0.00561492 |
| **B** | 1.765892688 | 1.525721405 | 1.25627043 | 1.829493441 | 1.043446901 | 1.631870354 |
| **C** | 1.075590959 | 4.391920576 | 4.054368493 | 9.19E-08 | 1.544313663 | 0.111809676 |
| **k1** | 0.08890734 | 5.73E-05 | 0.703146201 | 0.000455895 | 0.076936021 | 0.248689726 |
| **k2** | 0.088939956 | 0.057135557 | 0.036070089 | 0.054952683 | 0.076926233 | 0.067771498 |
| **k3** | 0.000156782 | 2.26E-05 | 1.97E-05 | 0.009975747 | 8.58E-05 | 0.000549956 |
| **NE** | 3.099244354 | 7.00E-08 | 2.49E-13 | 4.097455398 | 2.505831699 | 4.255907383 |

**IAVEAA**

| | IPTG | ONPF | APO | DNA | TMG | ONPFDNA |
|---|---|---|---|---|---|---|
| **A** | 0.0465787 | 0.104715233 | 0.463397018 | 0.126898329 | 0.426937401 | 0 |
| **B** | 0.124269749 | 0.105425446 | 0.150015984 | 0.132783267 | 0.207174188 | 0.165867326 |
| **C** | 0.964030348 | 0.332441228 | 3.37812064 | 3.67004259 | 3.35460235 | 0.280304219 |
| **k1** | 0.580332526 | 0.057364284 | 0.000937436 | 0.069623679 | 0.000619248 | 0 |
| **k2** | 0.029422832 | 0.057358542 | 0.041924392 | 0.06963508 | 0.050714409 | 0.042852772 |
| **k3** | 0.000298106 | 0.000639335 | 7.21E-06 | 7.83E-06 | 9.23E-06 | 0.000595788 |
| **NE** | 2.858573734 | 3.445672008 | 0.012487874 | 0.085139981 | 2.56E-05 | 3.559497934 |

**ACTNVPAL**

| | IPTG | ONPF | APO | DNA | TMG | ONPFDNA |
|---|---|---|---|---|---|---|
| **A** | 0.364699779 | 2.04E-39 | 0.978231958 | 3.91E-06 | 0.069231417 | 8.97E-15 |
| **B** | 1.710973271 | 1.783178422 | 0.995264638 | 2.13069922 | 1.654506492 | 2.074150143 |
| **C** | 0.626247839 | 3.113098653 | 0.581525118 | 0.095817843 | 0.902524104 | 2.69E-09 |
| **k1** | 0.501317405 | 0.382436701 | 0.04916163 | 0.200339911 | 0.07035891 | 0.202837257 |
| **k2** | 0.055732053 | 0.032511023 | 0.049164264 | 0.035419348 | 0.070414716 | 0.034319749 |
| **k3** | 0.000573025 | 1.45E-05 | 0.000980126 | 0.002364933 | 0.004900921 | 6.82E-15 |
| **NE** | 2.281983477 | 8.43E-10 | 2.448489263 | 2.779518407 | 2.353178755 | 2.936764625 |

**ALFLD**

| | IPTG | ONPF | APO | DNA | TMG | ONPFDNA |
|---|---|---|---|---|---|---|
| **A** | 0 | 0.15087794 | 0 | 0 | 0 | 0 |
| **B** | 0.23389919 | 4.67E-31 | 0.196807542 | 0.300152764 | 0.339067605 | 0.285164983 |
| **C** | 0.083939947 | 2.810116311 | 2.811401434 | 4.94E-15 | 1.43E-16 | 1.07E-11 |
| **k1** | 0 | 0.751636433 | 0 | 0 | 0 | 0 |
| **k2** | 0.058837133 | 0.009303356 | 0.1 | 0.060058588 | 0.043920054 | 0.064318355 |
| **k3** | 0.000138692 | 9.17E-06 | 6.43E-06 | 0.01 | 0.00992111 | 0.01 |
| **NE** | 2.66413954 | 2.50E-06 | 2.08E-16 | 2.722627655 | 2.632379968 | 2.706146186 |

**FLDVSDQTPINS**

| | IPTG | ONPF | APO | DNA | TMG | ONPFDNA |
|---|---|---|---|---|---|---|
| **A** | 1.017564257 | 0.089592264 | 3.725418739 | 0.626409312 | 0.792880346 | 3.89E-05 |
| **B** | 3.445625195 | 2.969562498 | 3.265938104 | 4.107879909 | 4.162204668 | 4.062761346 |
| **C** | 1.236769179 | 1.986953976 | 1.92966237 | 0.938512451 | 4.134993293 | 0.993621634 |
| **k1** | 0.001384256 | 0.031538745 | 2.28E-05 | 0.000185538 | 0.001012129 | 0.053713378 |
| **k2** | 0.036866439 | 0.031526374 | 0.034438 | 0.028720234 | 0.02991451 | 0.028179203 |
| **k3** | 0.000220854 | 0.000932483 | 0.001601261 | 0.000811593 | 6.87E-05 | 0.000550167 |
| **NE** | 3.349989822 | 4.079665333 | 0.0960493 | 3.441957445 | 5.63E-06 | 4.088093832 |

**DQTPIN**

| | IPTG | ONPF | APO | DNA | TMG | ONPFDNA |
|---|---|---|---|---|---|---|
| **A** | 0.187363319 | 0.92655919 | 0.430712196 | 0.107571881 | 0.568657969 | 0.513088677 |
| **B** | 0.842426595 | 0.175475644 | 0.566393125 | 1.171942155 | 0.655757543 | 0.564219611 |
| **C** | 0.612223672 | 1.858670632 | 0.670790508 | 0.0001038 | 0.476657422 | 0.145236757 |
| **k1** | 0.391737753 | 0.016936288 | 0.018411769 | 0.003388128 | 0.016997567 | 0.017641928 |
| **k2** | 0.011699243 | 0.002944995 | 0.018410945 | 0.019394403 | 0.016997229 | 0.017641197 |
| **k3** | 0.001152022 | 2.06E-05 | 0.001165372 | 0.01 | 0.000782871 | 0.000838086 |
| **NE** | 1.36720502 | 2.42E-05 | 1.345496304 | 1.743419736 | 1.306461096 | 1.791716565 |

**SIIFSH**

| | IPTG | ONPF | APO | DNA | TMG | ONPFDNA |
|---|---|---|---|---|---|---|
| **A** | 0 | 0.230293942 | 2.46E-33 | 0.13316651 | 4.95E-31 | 0.126194363 |
| **B** | 0.347829674 | 0.195958286 | 0.321957275 | 0.225035825 | 0.267592601 | 0.160917931 |
| **C** | 3.748467516 | 3.641543661 | 3.682492777 | 2.594855891 | 3.536519202 | 0.304749037 |
| **k1** | 0 | 0.038821274 | 0.844333445 | 0.000708488 | 0.999978135 | 0.911151028 |
| **k2** | 0.078960117 | 0.038797191 | 0.034323239 | 0.020460123 | 0.1 | 0.009064348 |
| **k3** | 2.23E-06 | 1.78E-05 | 2.20E-05 | 3.11E-06 | 4.64E-06 | 6.40E-05 |
| **NE** | 8.46E-13 | 0.000222259 | 1.91E-13 | 0.932168122 | 2.07E-06 | 3.136016429 |

**SIIFSHEDGTRL**

| | IPTG | ONPF | APO | DNA | TMG | ONPFDNA |
|---|---|---|---|---|---|---|
| **A** | 7.87E-21 | 0.597997908 | 0.473058106 | 0.320252847 | 0.612440301 | 0.291815217 |
| **B** | 0.644375436 | 0.383039418 | 0.463240647 | 0.308504066 | 0.629329747 | 0.284011368 |
| **C** | 0.935099002 | 9.03731504 | 4.228375675 | 0.426640156 | 2.927329924 | 0.164005941 |
| **k1** | 0.003798515 | 0.999973223 | 0.029320224 | 0.05810532 | 0.041928044 | 0.046237008 |
| **k2** | 0.043295386 | 0.000876652 | 0.029326626 | 0.058112895 | 0.041920509 | 0.046241334 |
| **k3** | 2.69E-05 | 7.41E-06 | 5.66E-05 | 0.00011134 | 5.36E-06 | 0.000381893 |
| **NE** | 8.165991582 | 8.42E-14 | 4.824183864 | 8.965576376 | 5.891689088 | 9.262060228 |

**EDGTRLGVEH**

| | IPTG | ONPF | APO | DNA | TMG | ONPFDNA |
|---|---|---|---|---|---|---|
| **A** | 0.648486304 | 6.29E-37 | 0.245191416 | 0.118025008 | 0.282850366 | 0.406508145 |
| **B** | 5.38E-16 | 0.569295351 | 0.225543262 | 0.454355339 | 0.26195077 | 0.129099048 |
| **C** | 1.83E-15 | 7.409614422 | 7.543303528 | 0.104830718 | 0.195781134 | 0.158678107 |
| **k1** | 0.857651676 | 0.999995578 | 0.070126411 | 0.060911268 | 0.060538502 | 0.73197984 |
| **k2** | 0.1 | 0.064054484 | 0.070096235 | 0.060903984 | 0.060564154 | 0.1 |
| **k3** | 0.01 | 9.89E-06 | 1.90E-05 | 0.000378485 | 0.001764975 | 0.000273685 |
| **NE** | 7.314992793 | 1.16E-09 | 1.66E-06 | 7.356001128 | 7.178356778 | 7.317247617 |

**LVAL**

| | IPTG | ONPF | APO | DNA | TMG | ONPFDNA |
|---|---|---|---|---|---|---|
| **A** | 0 | 0 | 0 | 0 | 0 | 0 |
| **B** | 0.059621227 | 0.055672589 | 0.037509119 | 0.058649798 | 0.102930432 | 0.050676079 |
| **C** | 0.580934104 | 1.94291822 | 1.960561225 | 0.094293327 | 1.899922927 | 0.066991936 |
| **k1** | 0 | 0 | 0 | 0 | 0 | 0 |
| **k2** | 0.050860871 | 0.073395644 | 0.1 | 0.064494529 | 0.054794493 | 0.060336866 |
| **k3** | 2.43E-05 | 5.55E-06 | 1.20E-05 | 0.000132985 | 4.62E-06 | 0.000247882 |
| **NE** | 1.359652403 | 7.12E-08 | 7.65E-16 | 1.848147295 | 2.71E-06 | 1.885867196 |

**LVALGHQQIAL**

| | IPTG | ONPF | APO | DNA | TMG | ONPFDNA |
|---|---|---|---|---|---|---|
| **A** | 1.45E-13 | 0.758287515 | 4.06E-39 | 0.646405145 | 4.14E-37 | 0.811850664 |
| **B** | 1.684423368 | 0.683407196 | 1.511241013 | 0.981890447 | 1.904395205 | 0.829936302 |
| **C** | 0.908838213 | 1.170578031 | 7.490699234 | 0.178735409 | 7.074580815 | 7.10E-07 |
| **k1** | 0.174794019 | 0.039174421 | 0.828670689 | 0.050265355 | 0.153879979 | 0.045657633 |
| **k2** | 0.069153592 | 0.039167336 | 0.070233447 | 0.050266884 | 0.054065883 | 0.045644847 |
| **k3** | 0.00011544 | 1.73E-05 | 1.16E-05 | 0.000497698 | 6.83E-06 | 0.009999913 |
| **NE** | 6.395382322 | 6.293111623 | 2.41E-14 | 7.171818104 | 2.76E-09 | 7.365406305 |

**LGHQQIALL**

| | IPTG | ONPF | APO | DNA | TMG | ONPFDNA |
|---|---|---|---|---|---|---|
| **A** | 0.555016987 | 1.92E-13 | 0.239806071 | 0.001716008 | 0.000498243 | 1.37E-21 |
| **B** | 0.649382542 | 0.932510005 | 0.773292328 | 1.262086989 | 1.515846641 | 1.242013591 |
| **C** | 1.10E-08 | 0.345555112 | 0.138966262 | 1.08E-14 | 1.11E-11 | 8.01E-19 |
| **k1** | 0.061276226 | 0.233259794 | 0.638144953 | 1.50E-11 | 0.047856806 | 0.999994513 |
| **k2** | 0.061275571 | 0.040137927 | 0.035491504 | 0.045154644 | 0.047810461 | 0.046422921 |
| **k3** | 0.00999939 | 6.30E-05 | 0.000221177 | 0.01 | 0.009999865 | 2.32E-05 |
| **NE** | 5.753649447 | 5.560332755 | 5.754185337 | 5.698565175 | 5.436398477 | 5.7565278 |

**IALL**

| | IPTG | ONPF | APO | DNA | TMG | ONPFDNA |
|---|---|---|---|---|---|---|
| **A** | 0 | 0 | 0 | 0 | 0 | 0 |
| **B** | 0.044403303 | 0.03475756 | 0.025537734 | 0.037737075 | 0.057343316 | 0.042997412 |
| **C** | 3.68E-07 | 1.996181297 | 1.979511973 | 1.983777643 | 1.69E-06 | 0.009789165 |
| **k1** | 0 | 0 | 0 | 0 | 0 | 0 |
| **k2** | 0.058207591 | 0.1 | 0.1 | 0.1 | 0.062770274 | 0.1 |
| **k3** | 1.74E-17 | 5.92E-07 | 1.37E-06 | 9.98E-07 | 3.99E-19 | 2.79E-17 |
| **NE** | 1.976181682 | 7.18E-09 | 1.65E-08 | 1.65E-10 | 1.971981519 | 1.998815444 |

**LAGPLS**

| | IPTG | ONPF | APO | DNA | TMG | ONPFDNA |
|---|---|---|---|---|---|---|
| **A** | 0.030410272 | 2.45E-14 | 0.00069302 | 0.952854304 | 5.20E-19 | 0.319363189 |
| **B** | 1.045284228 | 1.119738663 | 1.027895334 | 0.393654032 | 1.279377435 | 0.87358147 |
| **C** | 0.73890566 | 0.511206276 | 0.762190077 | 1.669809223 | 0.75783735 | 6.15E-07 |
| **k1** | 0.069819103 | 0.140057622 | 0.000980188 | 0.05571887 | 0.051528942 | 0.018148255 |
| **k2** | 0.069777956 | 0.034217355 | 0.060700663 | 0.005011054 | 0.044256391 | 0.053344885 |
| **k3** | 0.000735776 | 0.000916143 | 0.000978015 | 1.06E-05 | 0.000370646 | 0.01 |
| **NE** | 1.184566691 | 1.414230523 | 1.211568475 | 1.64E-10 | 0.997448336 | 1.841506943 |

**LAGPLSSVSARL**

| | IPTG | ONPF | APO | DNA | TMG | ONPFDNA |
|---|---|---|---|---|---|---|
| **A** | 1.54050119 | 0.853503947 | 0.541270942 | 0.019027757 | 0.890859498 | 3.37E-06 |
| **B** | 1.670762808 | 3.443687937 | 4.182111694 | 4.922694511 | 2.791709296 | 4.970964098 |
| **C** | 3.191916954 | 1.460527929 | 1.969824993 | 0.441242614 | 4.34616904 | 0.937156962 |
| **k1** | 0.024857872 | 0.02779967 | 0.027926206 | 0.001837496 | 0.038149495 | 0.171877808 |
| **k2** | 0.024859913 | 0.027791112 | 0.027925773 | 0.028971166 | 0.038139096 | 0.029453678 |
| **k3** | 0.000328499 | 0.001769204 | 0.000192352 | 0.001832589 | 0.000294522 | 0.000968049 |
| **NE** | 2.861074957 | 3.383659887 | 2.429024358 | 3.217628024 | 1.152934638 | 3.358790693 |

**GPLSSVSAR**

| | IPTG | ONPF | APO | DNA | TMG | ONPFDNA |
|---|---|---|---|---|---|---|
| **A** | 0.859443238 | 1.440520906 | 0.256128644 | 1.759420358 | 1.850543745 | 1.402688979 |
| **B** | 0.916876224 | 1.698264101 | 2.49157497 | 1.692970158 | 0.334364827 | 1.370503254 |
| **C** | 2.216572758 | 3.23E-05 | 1.949815168 | 1.24E-05 | 3.283085193 | 0.673918648 |
| **k1** | 0.04144469 | 0.027389273 | 0.051798035 | 0.031866851 | 0.038870426 | 0.03486246 |
| **k2** | 0.041444283 | 0.02737765 | 0.051799576 | 0.031860452 | 0.004578258 | 0.034872652 |
| **k3** | 0.00057292 | 0.01 | 0.000159213 | 0.01 | 0.000228332 | 0.000929605 |
| **NE** | 2.988429908 | 3.822166655 | 2.302200136 | 3.557201013 | 1.497944506 | 3.554651635 |

**LAGWHK**

| | IPTG | ONPF | APO | DNA | TMG | ONPFDNA |
|---|---|---|---|---|---|---|
| **A** | 0 | 0 | 2.23E-39 | 0 | 0 | 0 |
| **B** | 0.332800966 | 0.264157666 | 0.246481681 | 0.221216944 | 0.282667992 | 0.35791492 |
| **C** | 5.45E-14 | 1.18E-12 | 3.745898793 | 0.000739596 | 0.138784878 | 6.53E-09 |
| **k1** | 0 | 0 | 0.999932028 | 0 | 0 | 0 |
| **k2** | 0.1 | 0.041406634 | 0.091281845 | 0.1 | 0.06006399 | 0.1 |
| **k3** | 0.01 | 2.24E-05 | 4.86E-06 | 0.003903854 | 0.01 | 0.01 |
| **NE** | 3.62705792 | 3.695817726 | 9.61E-07 | 3.623947163 | 3.58759062 | 3.717669977 |

**YLTRNQIQP**

| | IPTG | ONPF | APO | DNA | TMG | ONPFDNA |
|---|---|---|---|---|---|---|
| **A** | 0.772915696 | 0.823031392 | 1.413802731 | 1.320235883 | 0.832798477 | 0.481425364 |
| **B** | 0.78865275 | 0.865668883 | 1.263338 | 0.692660105 | 0.896194205 | 0.933613179 |
| **C** | 1.923320569 | 1.611585743 | 1.875704669 | 4.002754741 | 2.143303399 | 2.605306363 |
| **k1** | 0.048197007 | 0.026081991 | 0.001214568 | 0.048602319 | 0.037344915 | 0.042890946 |
| **k2** | 0.048196523 | 0.026081929 | 0.05178588 | 0.001778038 | 0.03734505 | 0.042892918 |
| **k3** | 0.000379582 | 0.000102417 | 4.07E-05 | 3.35E-05 | 0.000299082 | 0.000128098 |
| **NE** | 2.510158832 | 2.723215774 | 1.454130862 | 4.75E-10 | 2.131013725 | 2.003221669 |

**NQIQPIAEREGD**

| | IPTG | ONPF | APO | DNA | TMG | ONPFDNA |
|---|---|---|---|---|---|---|
| **A** | 0.669673008 | 1.032057982 | 0.047063393 | 1.181268698 | 3.85E-22 | 0.000365378 |
| **B** | 2.425017651 | 1.541846783 | 2.749185435 | 1.214702902 | 3.317408862 | 2.306204608 |
| **C** | 1.835814737 | 6.392284058 | 2.676683655 | 0.537114542 | 3.12762041 | 0.823992712 |
| **k1** | 0.054819362 | 0.04423923 | 0.590934514 | 0.072178845 | 0.998936591 | 0.048981502 |
| **k2** | 0.054816078 | 0.044235375 | 0.056398782 | 0.072165825 | 0.046119752 | 0.081628067 |
| **k3** | 9.52E-05 | 1.98E-05 | 8.37E-05 | 0.001360318 | 3.37E-05 | 0.000400764 |
| **NE** | 4.039241446 | 0.000284915 | 3.518275592 | 6.070342562 | 2.546907332 | 5.852013244 |

**IAEREGDWSAMSG**

| | IPTG | ONPF | APO | DNA | TMG | ONPFDNA |
|---|---|---|---|---|---|---|
| A | 1.09278301 | 1.21324451 | 1.525813483 | 1.389419915 | 2.283963192 | 1.280033604 |
| B | 3.289359918 | 1.821159768 | 1.658426594 | 1.310298408 | 2.435755513 | 1.333949032 |
| C | 0.603330535 | 0.77327294 | 2.08918821 | 1.222545508 | 1.343981752 | 1.272523727 |
| k1 | 0.853007346 | 0.027327018 | 0.031787583 | 0.036627746 | 0.02688829 | 0.035344709 |
| k2 | 0.022173356 | 0.027320879 | 0.03178888 | 0.036628342 | 0.026889124 | 0.03535311 |
| k3 | 0.000853081 | 0.001773977 | 0.000750966 | 0.002429563 | 5.42E-05 | 0.00075032 |
| NE | 6.05311072 | 7.168555489 | 5.738771143 | 7.100340932 | 4.942591622 | 7.155596381 |

**REGDWSAMSGF**

| | IPTG | ONPF | APO | DNA | TMG | ONPFDNA |
|---|---|---|---|---|---|---|
| A | 6.28E-31 | 0.732037422 | 1.05908464 | 0.631748442 | 3.44E-33 | 0.789180229 |
| B | 2.283411719 | 0.806877634 | 1.674944863 | 0.864890449 | 2.880992748 | 0.771320489 |
| C | 0.810340114 | 1.589343925 | 6.2915073 | 0.858255045 | 0.866981987 | 1.277754857 |
| k1 | 0.314372789 | 0.038187747 | 0.001704235 | 0.027450482 | 0.816457675 | 0.026007743 |
| k2 | 0.022862292 | 0.038187018 | 0.021599762 | 0.027446838 | 0.023334847 | 0.026004962 |
| k3 | 0.001277868 | 0.001001285 | 7.64E-06 | 0.001666991 | 0.001046827 | 0.00069767 |
| NE | 6.002661283 | 5.863697457 | 0.002224032 | 6.735176851 | 5.468083675 | 6.489419656 |

**DWSAM**

| | IPTG | ONPF | APO | DNA | TMG | ONPFDNA |
|---|---|---|---|---|---|---|
| A | 0.306059729 | 0.522145805 | 0.497212042 | 0.644552371 | 0.506678855 | 0.297833783 |
| B | 1.188960277 | 0.753403511 | 0.467130178 | 0.624932431 | 1.068448002 | 0.349213547 |
| C | 0.265847906 | 1.776753031 | 0.879089002 | 1.789904127 | 0.386543081 | 0.668592321 |
| k1 | 0.861845153 | 0.002628106 | 0.015718459 | 0.021599515 | 0.017249188 | 0.018046987 |
| k2 | 0.011230655 | 0.018799913 | 0.015716134 | 0.002883016 | 0.017243682 | 0.018048103 |
| k3 | 0.000694447 | 2.76E-05 | 0.001011156 | 2.27E-05 | 0.000143797 | 0.000868555 |
| NE | 1.266828029 | 2.07E-06 | 1.167028032 | 3.41E-06 | 1.088867192 | 1.750638766 |

**WSAM**

| | IPTG | ONPF | APO | DNA | TMG | ONPFDNA |
|---|---|---|---|---|---|---|
| A | 0.229657155 | 0.362994763 | 0.647811775 | 0.328764125 | 0.096097743 | 0.25035886 |
| B | 1.051219904 | 0.341060491 | 0.349477362 | 0.408365515 | 1.140120974 | 0.283869584 |
| C | 0.212629311 | 0.618517686 | 0.520309636 | 0.545114422 | 0.342006154 | 0.654359563 |
| k1 | 0.922106483 | 0.017537918 | 0.021435056 | 0.768782289 | 0.837623399 | 0.021718917 |
| k2 | 0.011810276 | 0.017535662 | 0.002245761 | 0.002574208 | 0.017308744 | 0.021712774 |
| k3 | 0.000786665 | 0.00089846 | 0.002245773 | 0.002574298 | 0.000528667 | 0.000753177 |
| NE | 0.505837593 | 0.660404445 | 0.492763925 | 0.678365804 | 0.390884561 | 0.853832355 |

**SAMSGFQ**

| | IPTG | ONPF | APO | DNA | TMG | ONPFDNA |
|---|---|---|---|---|---|---|
| **A** | 0.817895584 | 0.488928341 | 0.772376281 | 0.489605476 | 0.049530513 | 9.33E-08 |
| **B** | 0.892663227 | 0.516437532 | 1.26275698 | 0.545512549 | 1.878784545 | 0.882408839 |
| **C** | 0.599651819 | 0.922003875 | 2.971475734 | 0.861108989 | 0.330285175 | 0.943807137 |
| **k1** | 0.028718308 | 0.023907666 | 0.035168795 | 0.032169218 | 8.61E-05 | 0.057005631 |
| **k2** | 0.028718959 | 0.023903645 | 0.002971568 | 0.032163493 | 0.025942288 | 0.030835896 |
| **k3** | 0.002226156 | 0.001451563 | 6.82E-06 | 0.001097514 | 0.000451116 | 0.000740391 |
| **NE** | 2.769831695 | 3.100458129 | 1.14E-17 | 3.139506399 | 2.692927345 | 3.200659784 |

**SGFQQTMQM**

| | IPTG | ONPF | APO | DNA | TMG | ONPFDNA |
|---|---|---|---|---|---|---|
| **A** | 0.201458592 | 0.149073884 | 0.197696442 | 0 | 0.154255961 | 0 |
| **B** | 1.69E-22 | 0.086564563 | 0.009492574 | 0.156635046 | 0.146005365 | 0.184369685 |
| **C** | 3.903102556 | 5.988255194 | 3.682442956 | 6.801094187 | 6.657740446 | 0.073029108 |
| **k1** | 0.126441893 | 0.069615403 | 0.067822636 | 0 | 0.052017936 | 0 |
| **k2** | 0.092596978 | 0.069603303 | 8.23E-06 | 0.044591317 | 0.051997947 | 0.066931362 |
| **k3** | 5.79E-06 | 2.65E-06 | 8.20E-06 | 2.37E-06 | 3.07E-06 | 0.000296931 |
| **NE** | 2.893916274 | 0.713449751 | 3.11908714 | 1.14E-09 | 0.002448401 | 6.721291935 |

**QTMQML**

| | IPTG | ONPF | APO | DNA | TMG | ONPFDNA |
|---|---|---|---|---|---|---|
| **A** | 0 | 0 | 0 | 0 | 0 | 0 |
| **B** | 0.067733126 | 0.068668748 | 0.065706128 | 0.096485739 | 0.095350899 | 0.089465891 |
| **C** | 1.103119401 | 3.933559802 | 3.938716518 | 0.058510662 | 3.922683359 | 0.055010039 |
| **k1** | 0 | 0 | 0 | 0 | 0 | 0 |
| **k2** | 0.074468762 | 0.051269459 | 0.037935455 | 0.062834292 | 0.06442549 | 0.041532834 |
| **k3** | 1.65E-05 | 2.94E-06 | 5.45E-06 | 0.000173948 | 3.65E-06 | 0.000262549 |
| **NE** | 2.836894144 | 1.01E-08 | 2.71E-16 | 3.88200463 | 4.72E-10 | 3.882709071 |

**NEGIVPTAML**

| | IPTG | ONPF | APO | DNA | TMG | ONPFDNA |
|---|---|---|---|---|---|---|
| **A** | 0.499353769 | 0.558750093 | 0.8760085 | 0.503253667 | 0.591599884 | 0.44438584 |
| **B** | 0.442723526 | 0.544540385 | 1.263623642 | 0.651568355 | 0.651315857 | 0.554419566 |
| **C** | 1.549133499 | 1.267787051 | 4.879299909 | 0.734591292 | 1.406102797 | 1.008811422 |
| **k1** | 0.037484018 | 0.023562684 | 0.036860544 | 0.033360153 | 0.033155472 | 0.034245481 |
| **k2** | 0.037484006 | 0.023560477 | 0.001292489 | 0.033361642 | 0.033158397 | 0.034243508 |
| **k3** | 0.000761687 | 0.000192215 | 1.02E-05 | 0.001076975 | 0.000426153 | 0.00070928 |
| **NE** | 4.482152523 | 4.524667478 | 0.000775835 | 5.083958962 | 4.330783267 | 5.009898286 |

LVANDQMALGAM

| | IPTG | ONPF | APO | DNA | TMG | ONPFDNA |
|---|---|---|---|---|---|---|
| A | 9.84E-07 | 0.664527344 | 7.39E-15 | 0.770978811 | 0.61895426 | 0.221318456 |
| B | 1.271630028 | 1.34218574 | 1.25644212 | 0.829609451 | 0.008959238 | 2.05691936 |
| C | 0.673619656 | 7.872920739 | 1.11009661 | 0.880002688 | 0.113892347 | 0.793765169 |
| k1 | 0.000131001 | 0.000353361 | 0.246300028 | 0.046421671 | 0.995395282 | 0.945482152 |
| k2 | 0.039658422 | 0.041544042 | 0.041377504 | 0.04642527 | 0.099999999 | 0.028469744 |
| k3 | 0.000146389 | 2.87E-06 | 0.000249286 | 0.000155719 | 0.000263815 | 0.000999426 |
| NE | 8.066368391 | 0.003726608 | 7.615214706 | 7.484030505 | 9.337099454 | 6.483744517 |

VANDQMALGAMR

| | IPTG | ONPF | APO | DNA | TMG | ONPFDNA |
|---|---|---|---|---|---|---|
| A | 0.777093139 | 0.412767204 | 0.680011682 | 1.95E-06 | 0.771783762 | 2.59E-09 |
| B | 1.14E-10 | 0.323850227 | 0.094776717 | 0.772702924 | 0.152320104 | 0.663216695 |
| C | 0.249747264 | 0.310418385 | 3.247746174 | 0.357993604 | 7.880379133 | 0.363972754 |
| k1 | 0.101751627 | 0.04647227 | 0.040219879 | 7.57E-20 | 0.091744149 | 0.284557692 |
| k2 | 0.058665503 | 0.046471162 | 0.1 | 0.062602804 | 0.00298608 | 0.072484508 |
| k3 | 0.000382351 | 0.000419209 | 2.59E-05 | 0.000331344 | 2.69E-06 | 0.000325297 |
| NE | 8.980481576 | 8.893479145 | 5.990157253 | 8.882640228 | 1.185968765 | 8.982081063 |

AITESGLRVGAD

| | IPTG | ONPF | APO | DNA | TMG | ONPFDNA |
|---|---|---|---|---|---|---|
| A | 8.67E-13 | 0.068686367 | 1.401215382 | 8.65E-05 | 2.769032201 | 1.163308342 |
| B | 2.863265144 | 2.29091339 | 1.152813709 | 2.449359492 | 0.203557686 | 1.196269467 |
| C | 1.195812313 | 2.98844917 | 7.450265431 | 0.154920531 | 6.99133431 | 0.199728929 |
| k1 | 0.094512125 | 0.034553433 | 0.053231167 | 0.056599668 | 0.044938924 | 0.049701018 |
| k2 | 0.058299347 | 0.034531923 | 0.053231582 | 0.048733824 | 0.044946327 | 0.049634128 |
| k3 | 0.000109479 | 6.18E-06 | 1.55E-05 | 0.001405322 | 1.03E-05 | 0.000501596 |
| NE | 5.942626373 | 4.528080441 | 0.001255561 | 7.368005745 | 0.000424431 | 7.445686452 |

ITESGLRVGAD

| | IPTG | ONPF | APO | DNA | TMG | ONPFDNA |
|---|---|---|---|---|---|---|
| A | 5.74E-05 | 2.17E-38 | 0.02377937 | 1.45E-34 | 7.45E-36 | 0.087911409 |
| B | 2.777345383 | 2.306987164 | 2.502435343 | 2.592285727 | 2.968377184 | 2.506166677 |
| C | 1.333638442 | 6.66342357 | 2.613240002 | 6.454632899 | 5.349668855 | 0.005965124 |
| k1 | 5.80E-19 | 0.999976535 | 0.049993062 | 0.8563855 | 0.90711752 | 0.000330476 |
| k2 | 0.053076179 | 0.041279201 | 0.049985322 | 0.041331451 | 0.047531842 | 0.048498131 |
| k3 | 7.22E-05 | 7.54E-06 | 4.22E-05 | 1.59E-06 | 1.39E-05 | 0.009999464 |
| NE | 4.951148638 | 1.17E-07 | 3.885743066 | 0.002963174 | 0.649661005 | 6.505721618 |

**ADISVVGY**

| | IPTG | ONPF | APO | DNA | TMG | ONPFDNA |
|---|---|---|---|---|---|---|
| **A** | 0 | 0 | 0.188003895 | 0 | 0 | 0 |
| **B** | 0.250685123 | 0.250124534 | 1.44E-23 | 0.236359397 | 0.325807978 | 0.241159565 |
| **C** | 1.27E-17 | 1.35E-15 | 3.74887402 | 1.24E-10 | 0.001984495 | 8.40E-10 |
| **k1** | 0 | 0 | 0.998187931 | 0 | 0 | 0 |
| **k2** | 0.1 | 0.060997781 | 0.070085065 | 0.099999998 | 0.048994472 | 0.1 |
| **k3** | 0.01 | 0.01 | 4.86E-06 | 0.01 | 0.00252465 | 1.55E-15 |
| **NE** | 5.728404055 | 5.725429641 | 2.058267716 | 5.752750557 | 5.656644204 | 5.761757141 |

**VVGYDD**

| | IPTG | ONPF | APO | DNA | TMG | ONPFDNA |
|---|---|---|---|---|---|---|
| **A** | 0 | 0 | 0.361217868 | 0 | 8.80E-13 | 0 |
| **B** | 0.509111788 | 0.444089118 | 8.83E-24 | 0.396418528 | 0.566667113 | 0.378578257 |
| **C** | 4.12E-17 | 0.036964802 | 3.633308416 | 0.033412104 | 0.072768251 | 2.07E-17 |
| **k1** | 0 | 0 | 0.76764394 | 0 | 0.910958338 | 0 |
| **k2** | 0.1 | 0.075193887 | 0.099999313 | 0.1 | 0.082468379 | 0.1 |
| **k3** | 0.01 | 0.000778152 | 1.25E-05 | 0.001257325 | 0.006595416 | 0.01 |
| **NE** | 3.479103865 | 3.51536697 | 2.63E-07 | 3.582918685 | 3.361801284 | 3.630268415 |

**DDTEDSSCYIPPLTTIK**

| | IPTG | ONPF | APO | DNA | TMG | ONPFDNA |
|---|---|---|---|---|---|---|
| **A** | 1.254318015 | 0.711136491 | 1.137629332 | 0.000777005 | 0.994361263 | 0.003499609 |
| **B** | 0.971889241 | 0.858260932 | 2.018220641 | 0.9627777 | 0.870949204 | 0.976371651 |
| **C** | 1.976806505 | 4.498190745 | 9.849176788 | 0.301747965 | 2.470465535 | 0.276954691 |
| **k1** | 0.041613426 | 0.03661087 | 0.050381563 | 0.216598986 | 0.029115321 | 0.60581454 |
| **k2** | 0.003108388 | 0.03661014 | 0.000826123 | 0.062922765 | 0.029112518 | 0.070278791 |
| **k3** | 0.000299264 | 0.000117216 | 2.02E-05 | 0.002178815 | 0.000293302 | 0.000672477 |
| **NE** | 8.822910939 | 6.973983343 | 2.25E-18 | 11.78344966 | 8.697763017 | 11.80225004 |

**TEDSSCYIPPLTTIK**

| | IPTG | ONPF | APO | DNA | TMG | ONPFDNA |
|---|---|---|---|---|---|---|
| **A** | 1.556158942 | 0.905079637 | 1.030343206 | 0.441543018 | 1.196164657 | 0.379321104 |
| **B** | 0.809604118 | 0.974444615 | 2.896849146 | 0.489022162 | 0.981607126 | 0.389762086 |
| **C** | 3.910578968 | 6.024632724 | 5.979150867 | 0.462655758 | 4.620472821 | 0.554258046 |
| **k1** | 0.034007018 | 0.029547049 | 0.045930275 | 0.048652283 | 0.034876185 | 0.067820847 |
| **k2** | 0.001817886 | 0.029541352 | 0.00095218 | 0.048647935 | 0.034876224 | 0.067864963 |
| **k3** | 0.000393407 | 0.000121969 | 8.19E-05 | 0.002200478 | 0.000237001 | 0.000775912 |
| **NE** | 4.802841215 | 3.164265721 | 1.104426569 | 9.7124752 | 4.240824452 | 9.701689373 |

**SSCYIPPLTTIKQD**

| | IPTG | ONPF | APO | DNA | TMG | ONPFDNA |
|---|---|---|---|---|---|---|
| **A** | 0.130810187 | 0.642413033 | 0.823054684 | 0.00021676 | 0.761662413 | 0.375536537 |
| **B** | 0.967598641 | 0.299906808 | 0.987173756 | 0.76784736 | 0.829350946 | 0.296699986 |
| **C** | 1.691973435 | 1.489285742 | 4.722252288 | 0.044575752 | 3.394668487 | 0.115675819 |
| **k1** | 0.74434456 | 0.051072324 | 0.066054389 | 0.09767127 | 0.043331748 | 0.056387979 |
| **k2** | 0.038955714 | 0.0510693 | 0.000824975 | 0.050729061 | 0.043326971 | 0.056504494 |
| **k3** | 0.000307053 | 0.000393708 | 5.90E-05 | 0.001113702 | 4.54E-05 | 0.000574887 |
| **NE** | 7.199014689 | 7.542752085 | 3.469629204 | 9.163341998 | 5.058566125 | 9.197139363 |

**FRLL**

| | IPTG | ONPF | APO | DNA | TMG | ONPFDNA |
|---|---|---|---|---|---|---|
| **A** | 0.110105893 | 0 | 0.076546111 | 0 | 0.108237439 | 0 |
| **B** | 0.066996739 | 0.100867429 | 0.256463103 | 0.08420405 | 0.06257733 | 0.063455588 |
| **C** | 0.59712447 | 0.272010478 | 1.668627967 | 0.205558266 | 0.736102452 | 0.157206027 |
| **k1** | 0.00038976 | 0 | 0.056945022 | 0 | 0.046334155 | 0 |
| **k2** | 0.065844227 | 0.059710012 | 0.000820678 | 0.064540749 | 0.046327873 | 0.064289434 |
| **k3** | 0.000389705 | 0.000377778 | 2.26E-05 | 0.000242122 | 0.000139956 | 0.000391565 |
| **NE** | 1.21977678 | 1.621227098 | 9.86E-06 | 1.714069599 | 1.105613665 | 1.781732619 |

**FRLLGQTSV**

| | IPTG | ONPF | APO | DNA | TMG | ONPFDNA |
|---|---|---|---|---|---|---|
| **A** | 0.139736718 | 0.192301081 | 0.217320196 | 0.18431024 | 0.238234793 | 0 |
| **B** | 0.304726555 | 0.212774957 | 0.239855393 | 0.122859513 | 0.230572895 | 0.121385781 |
| **C** | 0.689332146 | 3.760681575 | 1.978761923 | 0.711898237 | 3.38538576 | 0.176937078 |
| **k1** | 0.896304155 | 0.0392138 | 0.040312634 | 0.057785273 | 0.054147405 | 0 |
| **k2** | 0.034869593 | 0.039202939 | 0.040312145 | 0.057788315 | 0.054141733 | 0.094787671 |
| **k3** | 0.000268502 | 1.91E-05 | 5.23E-05 | 8.50E-05 | 1.78E-05 | 0.000347577 |
| **NE** | 5.8966946 | 2.874319436 | 4.572492082 | 6.066823556 | 3.215708387 | 6.585360841 |

**LGQTSVDR**

| | IPTG | ONPF | APO | DNA | TMG | ONPFDNA |
|---|---|---|---|---|---|---|
| **A** | 5.00E-39 | 3.03E-08 | 0.401804817 | 0 | 0.072945415 | 0 |
| **B** | 0.499979276 | 0.437077541 | 4.68E-15 | 0.426872423 | 0.589450811 | 0.394185904 |
| **C** | 4.652834157 | 0.097970795 | 5.598232396 | 0.031760722 | 2.280335178 | 0.038096873 |
| **k1** | 0.855896632 | 0.977514284 | 0.129523616 | 0 | 0.068171797 | 0 |
| **k2** | 0.090333269 | 0.062588725 | 9.52E-06 | 0.075104944 | 0.068162898 | 0.1 |
| **k3** | 2.96E-07 | 0.000343452 | 8.15E-06 | 0.0002399 | 1.34E-06 | 0.000312438 |
| **NE** | 0.82514454 | 5.438054166 | 1.73E-06 | 5.552239898 | 3.051786285 | 5.573989807 |

**GQTSVDRL**

| | IPTG | ONPF | APO | DNA | TMG | ONPFDNA |
|---|---|---|---|---|---|---|
| **A** | 0 | 0 | 0.292953513 | 0 | 0 | 0 |
| **B** | 0.33451389 | 0.320702877 | 0.022253958 | 0.32717771 | 0.452396532 | 0.285105524 |
| **C** | 0.027192927 | 0.010336733 | 5.67734544 | 0.030290974 | 5.82E-09 | 6.81E-08 |
| **k1** | 0 | 0 | 0.845399583 | 0 | 0 | 0 |
| **k2** | 0.066486252 | 0.048971692 | 0.045401786 | 0.058643081 | 0.063971803 | 0.099999999 |
| **k3** | 0.01 | 0.001793366 | 4.05E-06 | 0.003851939 | 4.15E-13 | 1.02E-09 |
| **NE** | 5.596515114 | 5.634517757 | 2.92E-06 | 5.64880139 | 5.560479158 | 5.70858039 |

**LLQL**

| | IPTG | ONPF | APO | DNA | TMG | ONPFDNA |
|---|---|---|---|---|---|---|
| **A** | 0.424998015 | 0.223092342 | 0.557166438 | 0.255350703 | 0.261636585 | 0.191567915 |
| **B** | 0.179738592 | 0.246744715 | 0.73088921 | 0.282527942 | 0.623866593 | 0.237726609 |
| **C** | 0.920235119 | 0.417486013 | 0.731379744 | 0.520407526 | 0.765584232 | 0.509704477 |
| **k1** | 0.02958332 | 0.021901569 | 0.023639393 | 0.027449606 | 0.109529557 | 0.023587116 |
| **k2** | 0.029586161 | 0.021901531 | 0.000893312 | 0.027447244 | 0.008533518 | 0.023584066 |
| **k3** | 0.000632798 | 0.001075579 | 1.85E-05 | 0.001369187 | 0.000223998 | 0.000887792 |
| **NE** | 0.48099988 | 1.110742548 | 3.14E-13 | 0.971072018 | 0.345807287 | 1.09987796 |
