## Appendix 2 for "Ligand-induced changes in dynamics mediate long-range allostery in the *lac* repressor"

### Appendix 2. Calculated uncertainties for the fractional differences at single HX timepoints based on maximum theoretical exchange and corrected for back-exchange.

### APO vs DNA

| Peptide | 0 s | 30 s | 45 s | 60 s | 300 s | 1500 s | 3600 s | 7200 s | 14400 s |
|---|---|---|---|---|---|---|---|---|---|
| LIGVA | 3.32E-06 | 3.62E-05 | 1.75E-05 | 3.79E-05 | 5.77E-05 | 3.22E-06 | 3.48E-06 | 8.91E-06 | 4.94E-05 |
| ALHAP | 1.51E-06 | 2.41E-04 | 3.08E-04 | 5.53E-04 | 9.37E-04 | 2.91E-04 | 4.98E-04 | 8.55E-04 | nan |
| ALHAPSQIVA | 2.50E-06 | 3.31E-05 | nan | 1.00E-04 | 4.85E-05 | 3.20E-05 | 1.73E-04 | 1.06E-04 | 5.26E-05 |
| ALHAPSQIVAA | 7.63E-06 | 7.02E-05 | 2.24E-04 | 1.27E-04 | 1.39E-04 | 6.40E-05 | 2.49E-04 | 4.38E-04 | 1.65E-03 |
| ALHAPSQIVAAIK | 5.04E-06 | nan | 2.63E-04 | 1.61E-04 | 3.36E-05 | 6.40E-05 | 6.36E-04 | 7.15E-04 | 2.24E-03 |
| SQIVAA | 2.42E-06 | 1.76E-05 | 1.38E-05 | 8.87E-05 | 7.88E-05 | 3.28E-04 | 1.14E-06 | 8.25E-04 | 5.31E-04 |
| SQIVAAIK | 4.22E-06 | 1.91E-05 | nan | 1.78E-04 | 2.32E-05 | 4.54E-04 | 9.74E-05 | 3.09E-04 | 5.38E-04 |
| IVAAIK | 5.93E-06 | 4.25E-06 | 5.01E-05 | 1.41E-04 | 5.73E-06 | 2.18E-04 | 5.05E-05 | 8.61E-05 | 5.14E-04 |
| AIKSRADQLGAS | 2.30E-06 | 1.36E-04 | 3.38E-04 | 1.67E-04 | 2.29E-04 | 1.50E-04 | 3.06E-04 | 3.83E-04 | 1.19E-03 |
| ADQLGAS | 1.28E-05 | 3.90E-05 | 1.17E-03 | 3.14E-04 | 2.14E-05 | 1.78E-04 | 3.00E-04 | nan | 8.32E-03 |
| VVVSM | 8.41E-07 | 3.62E-04 | 7.40E-04 | 2.09E-04 | 1.17E-03 | 2.93E-04 | 4.68E-04 | 1.26E-03 | 1.24E-03 |
| MVERSGVE | 6.49E-07 | 9.14E-04 | nan | 1.35E-03 | 1.16E-03 | 7.61E-04 | 1.19E-03 | 1.11E-03 | 9.17E-04 |
| AAVHNL | 2.54E-05 | 1.95E-05 | 2.20E-04 | 6.18E-07 | 2.52E-04 | 3.60E-06 | 4.50E-04 | 2.08E-03 | 7.94E-04 |
| LAQRVSGL | 1.82E-07 | 4.53E-05 | 2.78E-04 | 2.23E-06 | 1.15E-04 | 2.95E-06 | 1.82E-04 | 3.58E-04 | 7.66E-04 |
| VSGLIIN | 5.91E-05 | 1.79E-05 | 6.37E-05 | 9.76E-05 | 2.61E-05 | 9.35E-06 | 1.38E-04 | 1.57E-04 | 1.28E-05 |
| IINYPLD | 5.30E-06 | 6.46E-06 | 1.24E-05 | 2.40E-06 | 2.10E-05 | 1.49E-05 | 4.05E-05 | 1.18E-04 | 7.46E-04 |
| LDDQDAIA | 5.91E-08 | 5.49E-04 | 1.07E-03 | 8.62E-04 | nan | 2.27E-04 | 1.05E-03 | 1.50E-03 | 3.16E-03 |
| IAVEAA | 5.91E-07 | nan | 8.96E-04 | 2.01E-04 | 5.28E-05 | 1.45E-04 | 1.72E-04 | 4.42E-04 | 3.04E-04 |
| ACTNVPAL | 2.98E-05 | 6.74E-04 | 1.17E-03 | 2.16E-04 | 1.55E-03 | 1.87E-04 | 6.14E-04 | 2.47E-03 | 4.09E-03 |
| ALFLD | 3.64E-08 | 3.75E-05 | 1.61E-04 | 1.70E-05 | 4.56E-05 | 3.14E-05 | 5.91E-05 | 1.79E-05 | 1.21E-04 |
| FLDVSDQTPINS | 8.28E-06 | 9.66E-05 | nan | 2.85E-04 | 1.35E-04 | 3.71E-04 | 3.07E-04 | 5.20E-04 | 1.94E-04 |
| DQTPIN | 5.40E-06 | 7.40E-06 | 4.66E-04 | 4.79E-05 | 6.41E-04 | 1.69E-05 | 4.96E-04 | 8.56E-04 | 2.33E-03 |
| SIIFSH | 3.77E-05 | 1.54E-04 | 3.28E-04 | 1.85E-04 | nan | 2.20E-04 | 3.48E-04 | 3.79E-04 | 9.78E-04 |
| SIIFSHEDGTRL | 1.96E-06 | 2.06E-05 | 1.09E-05 | 1.82E-05 | 3.60E-05 | 3.93E-05 | 5.78E-05 | 7.56E-05 | 1.66E-04 |
| EDGTRLGVEH | 5.50E-06 | 2.75E-05 | 1.25E-05 | 3.61E-05 | 6.02E-06 | 1.17E-05 | 1.32E-04 | 1.05E-04 | 7.07E-04 |
| LVAL | 1.18E-07 | 1.84E-06 | 1.54E-05 | 4.16E-06 | 1.62E-06 | 1.43E-05 | 9.49E-06 | 3.09E-06 | 1.29E-04 |
| LVALGHQQIAL | 4.45E-06 | 2.64E-04 | 3.39E-04 | 9.73E-05 | 2.16E-04 | 4.99E-05 | 3.51E-04 | 7.92E-04 | 1.63E-03 |

| | | | | | | | | | |
|---|---|---|---|---|---|---|---|---|---|
| **LGHQQIALL** | 4.01E-06 | 1.48E-05 | nan | 1.30E-05 | 5.02E-05 | 3.11E-05 | 4.39E-05 | 7.31E-05 | 4.38E-05 |
| **IALL** | 6.39E-06 | 8.75E-06 | nan | 6.59E-06 | 1.17E-05 | 1.42E-05 | 2.24E-06 | 6.26E-06 | 1.33E-05 |
| **LAGPLS** | 5.32E-06 | 2.20E-04 | 3.37E-04 | 6.90E-05 | 3.84E-04 | nan | 4.80E-04 | nan | nan |
| **LAGPLSSVSARL** | 3.17E-04 | 4.29E-04 | 4.43E-04 | 8.11E-05 | 2.04E-04 | 4.12E-04 | 9.66E-04 | 4.86E-04 | 7.77E-04 |
| **GPLSSVSAR** | 4.34E-06 | 3.61E-04 | 6.58E-05 | 1.16E-04 | nan | 1.03E-03 | 2.65E-03 | 4.03E-03 | 3.06E-03 |
| **LAGWHK** | 8.92E-05 | 6.26E-05 | 7.76E-05 | 7.54E-06 | 5.05E-05 | 1.71E-05 | 1.57E-05 | 5.00E-05 | 6.09E-05 |
| **YLTRNQIQP** | 1.93E-06 | 6.68E-05 | 2.82E-04 | 6.55E-05 | 5.77E-04 | 9.71E-04 | 4.94E-04 | 1.32E-03 | nan |
| **NQIQPIAEREGD** | 3.73E-06 | 3.91E-04 | 5.22E-04 | 1.77E-04 | 7.29E-04 | 3.21E-04 | 6.84E-04 | 1.35E-03 | 5.10E-03 |
| **IAEREGDWSAMSG** | 2.85E-06 | 2.78E-04 | 1.13E-03 | 1.08E-05 | 1.36E-03 | 8.53E-04 | 1.77E-03 | 2.47E-03 | 6.54E-03 |
| **REGDWSAMSGF** | 5.96E-06 | 2.64E-05 | 2.72E-04 | 8.96E-05 | 1.12E-04 | 4.89E-04 | 3.02E-04 | 4.49E-04 | 1.10E-03 |
| **DWSAM** | 5.84E-06 | 2.82E-05 | 4.68E-04 | 9.85E-05 | 5.97E-04 | 2.37E-04 | 9.10E-04 | 1.41E-03 | nan |
| **WSAM** | 9.28E-05 | 1.04E-03 | 1.12E-03 | 1.59E-05 | 9.26E-04 | nan | 1.31E-03 | nan | nan |
| **SAMSGFQ** | 2.21E-05 | 1.06E-04 | 1.10E-03 | nan | 4.83E-04 | 4.04E-05 | 1.41E-03 | 6.97E-04 | nan |
| **SGFQQTMQM** | 4.22E-06 | 8.75E-06 | 1.02E-05 | 4.46E-05 | 1.26E-05 | 8.95E-06 | 7.64E-06 | 1.01E-05 | 8.76E-05 |
| **QTMQML** | 1.23E-05 | 1.20E-06 | nan | 1.48E-05 | 2.84E-06 | 2.71E-06 | 3.17E-06 | 1.75E-05 | 9.64E-05 |
| **NEGIVPTAML** | 1.15E-05 | 3.41E-05 | 3.58E-03 | 3.15E-04 | 4.98E-04 | 2.11E-06 | 2.79E-03 | 2.06E-03 | 1.13E-02 |
| **LVANDQMALGAM** | 2.32E-05 | 1.88E-04 | nan | 4.01E-04 | 2.93E-04 | 3.64E-04 | 2.73E-04 | 2.69E-04 | 2.79E-04 |
| **VANDQMALGAMR** | 4.09E-07 | 6.72E-05 | 2.08E-04 | 7.03E-06 | 8.02E-05 | 1.65E-05 | 2.81E-04 | 2.23E-04 | 1.42E-03 |
| **AITESGLRVGAD** | 7.54E-06 | 3.66E-04 | 3.03E-04 | 1.80E-04 | 4.68E-04 | 8.34E-05 | 3.98E-04 | 1.46E-03 | 2.23E-03 |
| **ITESGLRVGAD** | 3.83E-06 | 2.38E-04 | 2.69E-04 | 2.59E-04 | 3.34E-04 | 2.53E-04 | 3.50E-04 | 4.28E-04 | 4.41E-04 |
| **ADISVVGY** | 5.55E-06 | 8.57E-05 | 1.10E-04 | 1.59E-04 | 7.55E-05 | 3.96E-05 | 5.71E-05 | 1.72E-05 | 4.31E-04 |
| **VVGYDD** | 4.41E-06 | 8.46E-05 | 4.92E-05 | 6.23E-05 | 9.11E-05 | 4.41E-05 | 1.49E-04 | 1.58E-04 | 4.76E-04 |
| **DDTEDSSCYIPPLTTIK** | 4.55E-07 | 8.57E-05 | 1.62E-04 | 2.79E-05 | 1.87E-04 | 1.77E-04 | 2.20E-04 | 6.32E-04 | 1.17E-03 |
| **TEDSSCYIPPLTTIK** | 1.39E-06 | 2.08E-05 | 2.51E-05 | 1.38E-04 | 3.75E-05 | 2.82E-04 | 7.70E-05 | 2.95E-04 | 4.38E-04 |
| **SSCYIPPLTTIKQD** | 1.68E-06 | 4.75E-05 | 1.22E-04 | 3.60E-05 | 4.78E-05 | 2.26E-04 | 1.53E-04 | 1.77E-04 | 3.34E-04 |
| **FRLL** | 2.64E-07 | 1.82E-06 | 6.37E-05 | 2.22E-05 | 3.43E-06 | 7.12E-05 | 5.61E-05 | 6.13E-05 | 5.06E-04 |
| **FRLLGQTSV** | 2.13E-05 | 1.47E-05 | nan | 3.21E-05 | 2.39E-05 | 3.43E-05 | 5.08E-05 | 2.86E-05 | 1.17E-04 |
| **LGQTSVDR** | 1.32E-07 | 3.27E-05 | 2.89E-05 | 5.07E-05 | 3.72E-05 | 1.49E-05 | 4.17E-05 | 6.96E-05 | 2.73E-04 |
| **GQTSVDRL** | 1.14E-07 | 1.60E-05 | 2.30E-05 | 1.43E-05 | 3.01E-05 | 3.46E-05 | 3.31E-05 | 7.30E-05 | 1.05E-04 |
| **LLQL** | 4.02E-06 | 2.05E-07 | 7.87E-04 | 1.54E-04 | 6.61E-05 | 5.53E-05 | 2.52E-04 | 4.46E-04 | 2.71E-03 |

### APO vs ONPFDNA

| Peptide | 0 s | 30 s | 45 s | 60 s | 300 s | 1500 s | 3600 s | 7200 s | 14400 s |
|---|---|---|---|---|---|---|---|---|---|
| LIGVA | 4.48E-06 | 4.58E-05 | 4.82E-06 | 3.52E-05 | 4.71E-05 | 6.85E-06 | 3.77E-06 | 9.18E-06 | 6.73E-05 |
| ALHAP | 3.32E-07 | 2.25E-04 | 6.85E-05 | 5.34E-04 | 9.45E-04 | 2.88E-04 | 3.31E-04 | 4.60E-04 | 1.84E-04 |
| ALHAPSQIVA | 4.74E-06 | 1.32E-04 | nan | 1.48E-04 | 5.32E-05 | 2.29E-04 | 1.99E-04 | 1.49E-04 | 3.40E-04 |
| ALHAPSQIVAA | 4.93E-06 | 1.26E-04 | nan | 1.37E-04 | 1.21E-04 | 1.64E-04 | 2.80E-04 | 3.91E-04 | 1.47E-03 |
| ALHAPSQIVAAIK | 4.59E-06 | nan | 4.71E-05 | 1.71E-04 | 4.85E-05 | 4.35E-05 | 2.46E-04 | 8.57E-04 | 2.29E-04 |
| SQIVAA | 2.44E-05 | 1.06E-05 | 2.50E-06 | 7.42E-05 | nan | 4.48E-04 | 8.52E-06 | 7.66E-04 | 8.94E-05 |
| SQIVAAIK | 7.27E-07 | 9.62E-06 | 1.84E-05 | 1.79E-04 | 1.55E-06 | 4.58E-04 | 1.76E-04 | 3.72E-04 | 5.07E-05 |
| IVAAIK | 3.78E-06 | 9.32E-07 | 5.26E-06 | 1.42E-04 | 1.86E-06 | 2.19E-04 | 4.07E-05 | 1.19E-04 | 3.41E-04 |
| AIKSRADQLGAS | 2.22E-06 | 1.65E-04 | 1.26E-04 | 2.35E-05 | 1.48E-04 | 2.91E-04 | 2.15E-04 | 2.61E-04 | 5.64E-04 |
| ADQLGAS | 3.06E-05 | 4.36E-05 | 5.14E-04 | 5.73E-05 | 4.21E-04 | 6.29E-05 | 2.30E-04 | 5.05E-03 | 5.55E-03 |
| VVVSM | 7.45E-07 | 3.87E-04 | 4.36E-04 | 1.47E-04 | 1.22E-03 | 3.60E-04 | 3.22E-04 | 8.31E-04 | 9.58E-04 |
| MVERSGVE | 1.07E-06 | 1.46E-03 | nan | 1.21E-03 | 6.51E-04 | 6.32E-04 | 7.00E-04 | 6.01E-04 | 1.00E-03 |
| AAVHNL | 5.17E-05 | 8.95E-07 | 7.14E-05 | 2.24E-05 | 2.17E-04 | 5.20E-06 | 1.16E-04 | nan | 2.09E-04 |
| LAQRVSGL | 3.27E-07 | 3.49E-05 | 1.56E-04 | 3.86E-07 | 1.11E-04 | 1.35E-05 | 1.41E-04 | 3.06E-04 | 1.24E-04 |
| VSGLIIN | 6.77E-05 | 2.33E-05 | 6.01E-05 | 9.08E-05 | 1.31E-04 | 1.01E-05 | 1.09E-04 | 1.24E-04 | 1.15E-05 |
| IINYPLD | 3.11E-06 | 1.05E-06 | 6.12E-06 | 1.43E-05 | 4.66E-06 | 3.45E-05 | 3.49E-05 | 7.22E-05 | 1.86E-04 |
| LDDQDAIA | 6.73E-07 | 5.02E-04 | 7.38E-04 | 4.93E-04 | nan | 2.43E-04 | 4.69E-04 | 7.88E-04 | 6.80E-04 |
| IAVEAA | 5.32E-07 | 4.62E-06 | 6.11E-05 | 2.12E-04 | 2.12E-05 | 2.04E-04 | 1.56E-04 | 4.32E-04 | 2.20E-04 |
| ACTNVPAL | 1.29E-05 | 7.20E-04 | 8.37E-04 | 2.32E-04 | 1.52E-03 | 1.93E-04 | 1.33E-04 | 1.37E-03 | 6.77E-04 |
| ALFLD | 1.49E-06 | 2.47E-05 | nan | 6.23E-05 | 4.34E-04 | 2.23E-05 | 1.05E-04 | 1.09E-04 | 2.85E-04 |
| FLDVSDQTPINS | 8.64E-06 | 2.07E-04 | nan | 1.16E-04 | 1.22E-04 | 2.33E-04 | 3.41E-04 | 1.91E-04 | 1.44E-04 |
| DQTPIN | 7.02E-06 | 2.64E-05 | 6.36E-05 | 5.20E-05 | 7.41E-04 | 2.75E-05 | 2.15E-04 | 5.37E-04 | 3.25E-04 |
| SIIFSH | 2.74E-04 | 2.52E-04 | 8.23E-04 | 4.48E-04 | nan | 4.23E-04 | 1.12E-03 | 1.20E-03 | nan |
| SIIFSHEDGTRL | 1.86E-06 | 1.94E-05 | nan | 1.03E-05 | 5.02E-05 | 3.46E-05 | 6.18E-05 | 9.02E-05 | 9.48E-05 |
| EDGTRLGVEH | 1.44E-06 | 3.61E-05 | 1.03E-05 | 7.03E-06 | 3.51E-06 | 2.46E-05 | 1.69E-04 | 1.00E-04 | 5.75E-04 |
| LVAL | 1.55E-07 | 3.09E-07 | 6.23E-07 | 4.20E-06 | 5.27E-07 | 1.76E-05 | 4.55E-06 | 3.86E-07 | 4.15E-05 |
| LVALGHQQIAL | 9.32E-07 | 2.60E-04 | 2.84E-04 | 8.75E-05 | 2.27E-04 | 3.76E-05 | 2.11E-04 | 7.08E-04 | 6.74E-04 |
| LGHQQIALL | 4.79E-06 | 3.59E-05 | nan | 5.55E-05 | 4.62E-05 | 1.46E-05 | 3.33E-05 | 5.48E-05 | 1.55E-05 |
| IALL | 4.79E-06 | 7.56E-06 | 9.06E-06 | 6.93E-06 | 8.30E-06 | 6.30E-06 | 8.82E-07 | 6.59E-06 | 3.54E-05 |

| | | | | | | | | | |
|---|---|---|---|---|---|---|---|---|---|
| **LAGPLS** | 2.19E-06 | 2.66E-04 | 1.68E-04 | 7.39E-05 | 3.77E-04 | 1.55E-04 | 1.46E-04 | nan | 3.64E-04 |
| **LAGPLSSVSARL** | 1.54E-05 | 1.96E-04 | 2.39E-04 | 5.21E-05 | 2.99E-04 | 2.66E-04 | 8.31E-04 | 6.46E-04 | 3.25E-04 |
| **GPLSSVSAR** | 3.39E-06 | 4.04E-04 | 2.81E-04 | 1.13E-04 | nan | 1.53E-05 | 2.50E-03 | 3.50E-03 | 2.91E-04 |
| **LAGWHK** | 3.70E-06 | 4.58E-05 | 1.86E-06 | 1.18E-05 | 2.75E-05 | 5.96E-06 | nan | 3.35E-05 | 1.48E-04 |
| **YLTRNQIQP** | 1.49E-06 | 1.12E-04 | 6.34E-05 | 1.14E-04 | 5.71E-04 | 1.09E-03 | 2.35E-04 | 9.75E-04 | nan |
| **NQIQPIAEREGD** | 7.27E-07 | 3.80E-04 | 4.44E-04 | 1.69E-04 | 7.48E-04 | 4.15E-04 | 3.38E-04 | 1.11E-03 | nan |
| **IAEREGDWSAMSG** | 2.97E-06 | 3.28E-04 | 7.38E-04 | 1.18E-05 | 1.53E-03 | 9.23E-04 | 9.61E-04 | 1.76E-03 | 2.38E-03 |
| **REGDWSAMSGF** | 2.64E-05 | 9.14E-05 | 3.94E-04 | 1.26E-04 | 9.83E-05 | 1.07E-04 | 2.10E-04 | 1.69E-04 | 3.03E-04 |
| **DWSAM** | 1.35E-06 | 2.16E-05 | 1.30E-04 | 5.30E-05 | 6.79E-04 | 4.25E-04 | 6.03E-04 | 1.04E-03 | 1.07E-03 |
| **WSAM** | 2.66E-05 | 1.16E-04 | 4.32E-04 | 9.45E-05 | 8.51E-04 | 3.87E-04 | 4.70E-04 | nan | nan |
| **SAMSGFQ** | 7.55E-05 | 7.57E-05 | 2.20E-04 | 3.22E-04 | 7.51E-04 | 6.31E-05 | 1.22E-03 | 6.16E-04 | 2.66E-03 |
| **SGFQQTMQM** | 6.77E-06 | 1.48E-06 | 2.53E-05 | nan | 2.97E-06 | 1.14E-05 | 1.68E-06 | 1.31E-05 | 3.90E-05 |
| **QTMQML** | 8.53E-06 | 1.11E-06 | 3.65E-06 | 1.29E-05 | 2.99E-06 | 1.14E-05 | 3.06E-06 | 1.78E-05 | 1.07E-04 |
| **NEGIVPTAML** | 3.46E-05 | 1.37E-04 | 8.22E-04 | 3.10E-04 | 4.39E-04 | 2.28E-04 | 1.07E-03 | 1.21E-03 | 5.80E-03 |
| **LVANDQMALGAM** | 1.42E-04 | 2.00E-03 | nan | 3.99E-03 | 3.09E-03 | 4.85E-03 | 4.05E-03 | 2.23E-03 | 5.18E-03 |
| **VANDQMALGAMR** | 4.82E-07 | 5.30E-05 | 5.01E-05 | 1.44E-05 | 4.32E-05 | 1.68E-06 | 1.88E-04 | 2.43E-04 | 8.48E-04 |
| **AITESGLRVGAD** | 1.33E-06 | 3.43E-04 | 7.56E-05 | 1.40E-04 | 4.75E-04 | 5.50E-05 | 1.41E-04 | 1.01E-03 | 8.72E-04 |
| **ITESGLRVGAD** | 7.39E-06 | 2.87E-04 | 1.95E-04 | 2.07E-04 | 1.91E-04 | 9.31E-05 | 1.99E-04 | 2.39E-04 | 1.13E-04 |
| **ADISVVGY** | 9.45E-07 | 4.21E-05 | 6.75E-05 | 4.70E-06 | 1.33E-05 | 2.12E-05 | 2.78E-05 | 3.86E-05 | 3.90E-04 |
| **VVGYDD** | 2.63E-06 | 5.18E-05 | 6.68E-05 | 2.16E-05 | 2.75E-05 | 2.14E-05 | 6.73E-05 | 1.66E-04 | 2.78E-04 |
| **DDTEDSSCYIPPLTTIK** | 5.14E-07 | 7.83E-05 | 2.43E-04 | 3.77E-05 | 2.44E-04 | 1.75E-04 | 1.92E-04 | 5.21E-04 | 5.94E-04 |
| **TEDSSCYIPPLTTIK** | 1.39E-06 | 1.10E-05 | 1.47E-05 | 1.37E-04 | 6.50E-05 | 2.83E-04 | 7.19E-05 | 2.56E-04 | 1.73E-04 |
| **SSCYIPPLTTIKQD** | 2.64E-07 | 6.50E-05 | 8.49E-05 | 3.76E-05 | 3.83E-05 | 2.29E-04 | 1.35E-05 | 1.39E-04 | 1.83E-04 |
| **FRLL** | 6.77E-07 | 1.03E-06 | 7.27E-06 | 2.05E-05 | 1.32E-07 | 8.36E-05 | 3.69E-05 | 4.74E-05 | 1.43E-04 |
| **FRLLGQTSV** | 7.57E-05 | 2.22E-05 | 1.50E-05 | 3.22E-05 | 1.59E-05 | 7.79E-05 | 1.46E-05 | 2.21E-05 | 4.60E-05 |
| **LGQTSVDR** | 1.82E-08 | 2.10E-05 | 2.50E-05 | 2.36E-05 | 2.12E-05 | 5.30E-06 | 2.03E-05 | 7.22E-05 | 1.97E-04 |
| **GQTSVDRL** | 2.77E-07 | 1.43E-05 | 1.46E-05 | 1.22E-05 | 1.26E-05 | 2.91E-05 | 3.85E-06 | 6.65E-05 | 6.68E-05 |
| **LLQL** | 1.93E-06 | 4.70E-06 | 1.42E-04 | 1.58E-04 | 1.12E-04 | 5.58E-05 | 3.10E-04 | 4.23E-04 | 7.08E-04 |

### APO vs TMG

| Peptide | 0 s | 30 s | 45 s | 60 s | 300 s | 1500 s | 3600 s | 7200 s | 14400 s |
|---|---|---|---|---|---|---|---|---|---|
| LIGVA | 3.77E-06 | 2.31E-05 | 5.27E-06 | 2.70E-05 | 3.35E-05 | 2.22E-06 | 1.71E-05 | 9.33E-06 | 4.63E-05 |
| ALHAP | 4.09E-07 | 2.56E-04 | 7.29E-05 | 7.12E-04 | 8.83E-04 | 4.26E-04 | 5.55E-04 | 6.77E-04 | 3.76E-04 |
| ALHAPSQIVA | 2.99E-06 | 4.66E-05 | nan | 7.08E-04 | 1.35E-05 | 2.26E-04 | 1.07E-04 | 4.27E-04 | 1.76E-04 |
| ALHAPSQIVAA | 5.76E-06 | 6.69E-05 | 6.97E-05 | 6.62E-04 | 8.38E-05 | 2.00E-04 | 9.69E-05 | 6.17E-04 | 1.39E-03 |
| ALHAPSQIVAAIK | 7.93E-06 | nan | 2.10E-04 | nan | 1.57E-04 | 3.75E-05 | 1.19E-03 | 6.63E-04 | 2.04E-04 |
| SQIVAA | 1.35E-05 | 1.21E-05 | 3.31E-05 | 1.50E-04 | nan | 2.80E-04 | 9.05E-05 | 7.82E-04 | 8.57E-05 |
| SQIVAAIK | 1.82E-06 | 9.94E-06 | 1.68E-05 | 2.65E-04 | 1.18E-06 | 4.82E-04 | 2.16E-05 | 2.52E-04 | 5.76E-05 |
| IVAAIK | 4.44E-06 | 1.64E-06 | 4.37E-06 | 1.84E-04 | 2.53E-06 | 2.33E-04 | 3.49E-05 | 8.84E-05 | 3.41E-04 |
| AIKSRADQLGAS | 2.92E-06 | 2.92E-04 | 1.11E-04 | 3.13E-05 | 1.28E-04 | 2.32E-04 | 4.03E-04 | 1.15E-03 | 5.73E-04 |
| ADQLGAS | 1.38E-05 | 1.97E-04 | 5.40E-04 | 3.00E-04 | 2.34E-05 | 6.66E-04 | 1.52E-03 | 2.67E-03 | 4.10E-03 |
| VVVSM | 6.15E-05 | 5.87E-04 | 2.72E-04 | 6.92E-05 | 1.02E-03 | 2.50E-04 | 2.96E-04 | 1.12E-03 | 6.89E-04 |
| MVERSGVE | 1.11E-05 | 2.99E-03 | nan | 1.05E-03 | 6.57E-04 | 1.23E-03 | 1.32E-03 | 1.35E-03 | 1.19E-03 |
| AAVHNL | 2.40E-05 | 1.18E-06 | 3.76E-04 | 3.54E-05 | 8.28E-05 | 4.46E-05 | 2.20E-04 | 2.10E-03 | 2.85E-04 |
| LAQRVSGL | 3.86E-07 | 1.98E-04 | 2.25E-04 | 4.39E-05 | 1.31E-04 | 5.18E-05 | 1.76E-04 | 6.32E-04 | 2.81E-04 |
| VSGLIIN | 5.11E-05 | 1.63E-05 | 8.09E-05 | 9.32E-05 | 2.70E-05 | 2.16E-05 | 1.11E-04 | 1.00E-04 | 1.60E-05 |
| IINYPLD | 1.87E-05 | 1.18E-05 | 2.02E-06 | nan | 1.82E-06 | 3.53E-05 | 2.36E-05 | 7.94E-05 | 1.76E-04 |
| LDDQDAIA | 6.24E-06 | 1.18E-03 | 6.42E-04 | 4.88E-04 | nan | 3.43E-04 | 4.67E-04 | 1.01E-03 | 5.12E-04 |
| IAVEAA | 3.73E-07 | 9.89E-06 | 1.09E-04 | 2.00E-04 | 2.48E-05 | 4.48E-05 | 1.45E-04 | 5.35E-04 | 2.06E-04 |
| ACTNVPAL | 3.65E-05 | 2.28E-03 | 8.38E-04 | 3.18E-04 | 1.66E-03 | 8.93E-04 | 5.11E-04 | 3.07E-03 | 1.22E-03 |
| ALFLD | 1.01E-05 | 3.53E-06 | 1.18E-04 | 1.81E-05 | 9.32E-06 | 7.27E-07 | 2.29E-05 | 6.85E-05 | 1.95E-04 |
| FLDVSDQTPINS | 1.35E-05 | 2.74E-04 | nan | 7.54E-04 | 3.57E-05 | 5.43E-04 | 3.36E-04 | 6.38E-04 | 1.01E-04 |
| DQTPIN | 2.65E-06 | 4.77E-05 | 1.75E-04 | 4.63E-04 | 6.15E-04 | 4.51E-04 | 3.94E-04 | 1.13E-03 | 3.95E-04 |
| SIIFSH | 3.42E-04 | 4.16E-05 | 9.17E-05 | 6.83E-04 | nan | 1.18E-04 | 1.17E-04 | 3.67E-04 | 2.45E-04 |
| SIIFSHEDGTRL | 2.01E-06 | 2.03E-04 | 5.52E-06 | 2.45E-04 | 5.34E-05 | 6.71E-05 | 2.90E-05 | 8.01E-05 | 1.11E-04 |
| EDGTRLGVEH | 1.75E-05 | 8.18E-05 | 3.22E-04 | nan | 1.02E-04 | 4.70E-04 | 1.87E-05 | 1.28E-04 | 9.96E-04 |
| LVAL | 1.86E-07 | 1.32E-07 | 6.64E-06 | 1.48E-04 | 4.59E-07 | 3.17E-05 | 6.42E-06 | 2.41E-07 | 3.79E-05 |
| LVALGHQQIAL | 4.98E-06 | 4.98E-04 | 2.35E-04 | 4.14E-04 | 1.84E-04 | 2.52E-04 | 1.56E-04 | 5.71E-04 | 7.47E-04 |
| LGHQQIALL | 1.08E-05 | 3.33E-05 | nan | 1.03E-05 | 2.68E-05 | 6.07E-05 | 3.46E-05 | 6.37E-05 | 2.82E-05 |
| IALL | 4.95E-06 | 1.35E-05 | 9.43E-06 | 6.80E-05 | 7.86E-06 | 2.20E-05 | 1.54E-06 | 9.54E-06 | 1.25E-05 |

| | | | | | | | | | |
|---|---|---|---|---|---|---|---|---|---|
| **LAGPLS** | 2.26E-05 | 6.59E-04 | 1.72E-04 | 4.88E-04 | 3.96E-04 | 3.13E-04 | 1.41E-04 | nan | 4.59E-04 |
| **LAGPLSSVSARL** | 3.93E-05 | 1.56E-04 | 9.43E-05 | 2.71E-04 | 1.32E-04 | 6.82E-04 | 7.83E-04 | nan | nan |
| **GPLSSVSAR** | 6.88E-06 | 7.47E-04 | 1.42E-04 | nan | nan | 4.13E-04 | 2.79E-03 | 3.40E-03 | nan |
| **LAGWHK** | 1.70E-05 | 1.87E-04 | 2.43E-04 | nan | 1.30E-07 | 5.08E-04 | 1.61E-05 | 3.01E-04 | 4.25E-04 |
| **YLTRNQIQP** | 2.79E-06 | 4.81E-04 | nan | 1.57E-04 | 5.26E-04 | nan | 4.85E-04 | nan | 1.87E-03 |
| **NQIQPIAEREGD** | 1.20E-06 | 1.28E-03 | 4.43E-04 | 5.23E-04 | nan | 7.84E-04 | nan | 1.29E-03 | 1.40E-03 |
| **IAEREGDWSAMSG** | 2.97E-06 | 8.18E-04 | 5.79E-04 | 7.04E-04 | 1.43E-03 | 1.78E-03 | 1.34E-03 | 2.70E-03 | 2.87E-03 |
| **REGDWSAMSGF** | 6.29E-06 | 3.79E-04 | 2.98E-04 | 3.11E-04 | nan | 2.75E-04 | nan | 8.80E-04 | 4.85E-04 |
| **DWSAM** | 1.41E-05 | 1.60E-04 | 2.05E-04 | 7.44E-04 | 6.03E-04 | 6.33E-04 | 8.29E-04 | 1.82E-03 | 1.28E-03 |
| **WSAM** | 1.48E-05 | 7.12E-05 | 4.77E-04 | 2.59E-04 | 6.81E-04 | 5.27E-04 | 7.05E-04 | nan | 1.26E-03 |
| **SAMSGFQ** | 2.91E-05 | 4.48E-04 | 8.73E-04 | 5.86E-04 | 4.99E-04 | 1.74E-04 | 1.29E-03 | nan | 2.68E-03 |
| **SGFQQTMQM** | 6.47E-06 | 2.64E-06 | 3.04E-05 | nan | 3.84E-06 | 1.36E-05 | 3.21E-06 | 3.51E-06 | 3.06E-05 |
| **QTMQML** | 2.71E-06 | 1.20E-05 | 1.59E-05 | nan | 3.07E-06 | 7.39E-06 | 4.22E-06 | 2.30E-05 | 8.29E-05 |
| **NEGIVPTAML** | 2.20E-05 | 2.78E-04 | 1.50E-03 | 3.18E-04 | 2.45E-04 | 2.36E-04 | 6.30E-04 | 2.22E-03 | 5.37E-03 |
| **LVANDQMALGAM** | 2.29E-05 | 2.45E-04 | 6.22E-06 | nan | 2.68E-04 | 4.68E-04 | 2.51E-04 | 2.16E-04 | 1.99E-04 |
| **VANDQMALGAMR** | 1.14E-06 | 1.20E-04 | 8.91E-07 | nan | 3.38E-05 | 1.97E-04 | 1.54E-05 | 1.22E-04 | 8.60E-04 |
| **AITESGLRVGAD** | 9.72E-06 | 1.09E-03 | 4.65E-06 | 6.40E-04 | 6.34E-04 | 7.19E-04 | 3.08E-04 | 1.49E-03 | 1.20E-03 |
| **ITESGLRVGAD** | 9.36E-06 | 1.17E-03 | 1.93E-04 | 4.12E-04 | 3.40E-04 | 6.96E-04 | 3.76E-04 | 9.44E-04 | 3.68E-04 |
| **ADISVVGY** | 1.31E-06 | 2.39E-05 | 6.10E-06 | nan | 1.31E-06 | 1.77E-04 | 5.06E-06 | 6.59E-06 | 4.24E-04 |
| **VVGYDD** | 2.77E-06 | 2.86E-05 | 2.53E-05 | nan | 2.95E-05 | 9.84E-05 | 3.62E-05 | 9.07E-05 | 3.25E-04 |
| **DDTEDSSCYIPPLTTIK** | 4.55E-07 | 2.17E-04 | 1.75E-04 | nan | 1.74E-04 | 4.35E-04 | 1.74E-04 | 1.13E-03 | 1.34E-03 |
| **TEDSSCYIPPLTTIK** | 3.78E-06 | 5.63E-05 | 1.25E-04 | nan | 7.80E-06 | 3.76E-04 | 2.53E-05 | 5.09E-04 | 4.61E-04 |
| **SSCYIPPLTTIKQD** | 1.55E-07 | 9.28E-05 | 1.96E-04 | 3.33E-03 | 3.28E-05 | 4.58E-04 | 2.02E-05 | 1.82E-04 | 3.73E-04 |
| **FRLL** | 4.45E-07 | 2.01E-06 | 5.38E-05 | 1.78E-04 | 3.19E-06 | 8.22E-05 | 1.33E-05 | 1.41E-04 | 1.87E-04 |
| **FRLLGQTSV** | 2.54E-05 | 6.55E-05 | 9.84E-05 | 1.32E-04 | 4.28E-05 | 8.54E-05 | nan | 3.90E-05 | 3.39E-05 |
| **LGQTSVDR** | 2.41E-07 | 2.48E-05 | 2.75E-05 | 1.71E-04 | 2.22E-05 | 5.73E-05 | 1.76E-05 | 5.46E-05 | 2.17E-04 |
| **GQTSVDRL** | 2.77E-07 | nan | nan | 8.64E-05 | nan | 8.47E-05 | 1.45E-06 | 6.06E-05 | nan |
| **LLQL** | 2.46E-06 | 5.60E-05 | 3.05E-04 | 2.13E-04 | 6.81E-05 | 1.79E-04 | 1.76E-04 | 1.17E-03 | 4.32E-04 |

### DNA vs ONPFDNA

| Peptide | 0 s | 30 s | 45 s | 60 s | 300 s | 1500 s | 3600 s | 7200 s | 14400 s |
|---|---|---|---|---|---|---|---|---|---|
| LIGVA | 1.17E-06 | 3.62E-05 | 1.79E-05 | 2.19E-05 | 3.91E-05 | 5.68E-06 | 6.18E-07 | 2.05E-06 | 3.47E-05 |
| ALHAP | 1.76E-06 | 2.57E-05 | 2.41E-04 | 9.10E-05 | 1.22E-04 | 1.09E-05 | 1.72E-04 | 5.24E-04 | nan |
| ALHAPSQIVA | 2.57E-06 | 1.18E-04 | nan | 1.93E-04 | 7.47E-05 | 2.25E-04 | 2.19E-04 | 1.02E-04 | 3.76E-04 |
| ALHAPSQIVAA | 2.82E-06 | 1.36E-04 | nan | 2.05E-04 | 1.15E-04 | 1.78E-04 | 3.42E-04 | 2.86E-04 | 6.67E-04 |
| ALHAPSQIVAAIK | 2.50E-06 | 4.84E-05 | 2.24E-04 | 3.09E-05 | 7.60E-05 | 7.59E-05 | 7.43E-04 | 5.70E-04 | 2.06E-03 |
| SQIVAA | 2.64E-05 | 2.15E-05 | 1.42E-05 | 2.76E-05 | nan | 2.73E-04 | 9.43E-06 | 6.34E-05 | 4.55E-04 |
| SQIVAAIK | 3.64E-06 | 2.00E-05 | nan | 4.20E-06 | 2.29E-05 | 4.20E-06 | 2.70E-04 | 1.79E-04 | 5.45E-04 |
| IVAAIK | 4.48E-06 | 4.86E-06 | 5.54E-05 | 1.11E-06 | 4.31E-06 | 1.31E-06 | 4.14E-05 | 4.71E-05 | 1.83E-04 |
| AIKSRADQLGAS | 2.16E-07 | 1.45E-04 | 2.46E-04 | 1.68E-04 | 1.95E-04 | 2.84E-04 | 2.33E-04 | 1.70E-04 | 1.31E-03 |
| ADQLGAS | 4.33E-05 | 6.42E-05 | 6.58E-04 | 3.48E-04 | 4.24E-04 | 1.41E-04 | 1.40E-04 | nan | 5.90E-03 |
| VVVSM | 1.00E-06 | 3.10E-05 | 8.68E-04 | 2.69E-04 | 3.46E-04 | 1.81E-04 | 2.18E-04 | 7.27E-04 | 1.02E-03 |
| MVERSGVE | 4.34E-07 | 1.00E-03 | nan | 5.43E-04 | 9.65E-04 | 6.13E-04 | 8.08E-04 | 7.90E-04 | 6.80E-04 |
| AAVHNL | 3.63E-05 | 1.86E-05 | 1.49E-04 | 2.27E-05 | 3.70E-04 | 1.68E-06 | 5.00E-04 | nan | 6.07E-04 |
| LAQRVSGL | 1.82E-07 | 1.91E-05 | 1.63E-04 | 2.17E-06 | 1.31E-05 | 1.31E-05 | 1.40E-04 | 6.36E-05 | 6.43E-04 |
| VSGLIIN | 3.22E-05 | 1.06E-05 | 2.91E-05 | 7.73E-06 | 1.06E-04 | 1.86E-06 | 2.86E-05 | 9.49E-05 | 7.44E-06 |
| IINYPLD | 8.33E-06 | 5.73E-06 | 1.74E-05 | 1.67E-05 | 2.20E-05 | 4.92E-05 | 2.83E-05 | 4.85E-05 | 5.79E-04 |
| LDDQDAIA | 6.95E-07 | 7.34E-05 | 5.52E-04 | 4.82E-04 | 5.19E-04 | 3.44E-04 | 7.69E-04 | 8.95E-04 | 2.94E-03 |
| IAVEAA | 3.86E-07 | nan | 9.49E-04 | 1.27E-05 | 3.56E-05 | 2.69E-04 | 4.31E-05 | 7.16E-05 | 1.38E-04 |
| ACTNVPAL | 1.81E-05 | 4.64E-05 | 3.34E-04 | 1.63E-05 | 2.69E-05 | 2.05E-05 | 4.91E-04 | 1.12E-03 | 3.53E-03 |
| ALFLD | 1.49E-06 | 5.60E-05 | nan | 7.70E-05 | 4.79E-04 | 5.36E-05 | 1.64E-04 | 9.21E-05 | 1.93E-04 |
| FLDVSDQTPINS | 3.82E-07 | 1.61E-04 | nan | 1.78E-04 | 1.92E-04 | 2.71E-04 | 4.90E-04 | 4.20E-04 | 3.35E-04 |
| DQTPIN | 1.15E-05 | 2.56E-05 | 5.11E-04 | 8.97E-06 | 1.57E-04 | 3.91E-05 | 2.83E-04 | 3.73E-04 | 2.03E-03 |
| SIIFSH | 2.57E-04 | 3.39E-04 | 9.67E-04 | 5.39E-04 | 4.40E-04 | 6.43E-04 | 1.26E-03 | 8.52E-04 | nan |
| SIIFSHEDGTRL | 2.44E-07 | 1.97E-05 | nan | 2.81E-05 | 4.05E-05 | 3.24E-05 | 6.18E-05 | 9.11E-05 | 9.72E-05 |
| EDGTRLGVEH | 5.84E-06 | 2.87E-05 | 8.33E-06 | 4.29E-05 | 5.14E-06 | 2.10E-05 | 2.67E-04 | 4.15E-05 | 1.36E-04 |
| LVAL | 4.55E-08 | 2.11E-06 | 1.58E-05 | 1.86E-07 | 1.24E-06 | 3.32E-06 | 1.11E-05 | 3.44E-06 | 9.55E-05 |
| LVALGHQQIAL | 3.60E-06 | 5.97E-05 | 1.63E-04 | 3.35E-05 | 8.68E-05 | 1.23E-05 | 2.96E-04 | 6.65E-04 | 9.54E-04 |
| LGHQQIALL | 1.05E-06 | 3.95E-05 | 1.01E-04 | 6.80E-05 | 5.15E-05 | 2.27E-05 | 3.34E-05 | 5.71E-05 | 3.04E-05 |
| IALL | 4.55E-06 | 1.76E-06 | nan | 1.28E-05 | 4.75E-06 | 8.09E-06 | 1.59E-06 | 4.09E-07 | 2.50E-05 |

| | | | | | | | | | |
|---|---|---|---|---|---|---|---|---|---|
| **LAGPLS** | 6.77E-06 | 8.08E-05 | 1.73E-04 | 1.20E-05 | 7.14E-06 | nan | 3.54E-04 | nan | nan |
| **LAGPLSSVSARL** | 3.04E-04 | 4.64E-04 | 4.95E-04 | 5.93E-05 | 2.39E-04 | 4.01E-04 | 4.26E-04 | 7.80E-04 | 1.04E-03 |
| **GPLSSVSAR** | 1.10E-06 | 5.70E-05 | 2.33E-04 | 2.20E-05 | 1.84E-06 | 1.04E-03 | 1.58E-04 | 2.12E-03 | 2.77E-03 |
| **LAGWHK** | 9.27E-05 | 6.98E-05 | 7.81E-05 | 7.10E-06 | 7.80E-05 | 2.17E-05 | nan | 2.20E-05 | 8.73E-05 |
| **YLTRNQIQP** | 4.73E-07 | 4.57E-05 | 2.84E-04 | 4.83E-05 | 1.82E-04 | 1.23E-04 | 2.61E-04 | 3.51E-04 | nan |
| **NQIQPIAEREGD** | 3.15E-06 | 1.64E-05 | 8.12E-05 | 7.68E-06 | 8.74E-05 | 1.21E-04 | 4.68E-04 | 8.87E-04 | nan |
| **IAEREGDWSAMSG** | 1.42E-06 | 1.62E-04 | 9.05E-04 | 1.15E-05 | 3.72E-04 | 7.13E-05 | 9.97E-04 | 7.06E-04 | 4.23E-03 |
| **REGDWSAMSGF** | 2.74E-05 | 7.26E-05 | 6.20E-04 | 1.75E-04 | 1.01E-04 | 4.83E-04 | 2.85E-04 | 3.13E-04 | 1.04E-03 |
| **DWSAM** | 5.40E-06 | 1.71E-05 | 4.27E-04 | 4.55E-05 | 1.81E-04 | 1.98E-04 | 4.37E-04 | 3.67E-04 | nan |
| **WSAM** | 8.98E-05 | 1.02E-03 | 9.32E-04 | 1.02E-04 | 4.57E-04 | nan | 8.60E-04 | 7.71E-04 | nan |
| **SAMSGFQ** | 7.10E-05 | 4.46E-05 | 1.09E-03 | nan | 2.69E-04 | 2.98E-05 | 2.07E-04 | 8.18E-05 | nan |
| **SGFQQTMQM** | 7.35E-06 | 7.28E-06 | 1.54E-05 | nan | 1.40E-05 | 1.77E-05 | 8.59E-06 | 2.06E-05 | 6.54E-05 |
| **QTMQML** | 1.90E-05 | 1.00E-06 | nan | 6.77E-06 | 5.91E-07 | 1.05E-05 | 1.42E-06 | 3.86E-07 | 3.97E-05 |
| **NEGIVPTAML** | 3.36E-05 | 1.27E-04 | 4.08E-03 | 5.55E-06 | 4.48E-04 | 2.26E-04 | 2.76E-03 | 8.80E-04 | 7.22E-03 |
| **LVANDQMALGAM** | 1.19E-04 | 1.94E-03 | nan | 3.96E-03 | 3.00E-03 | 4.74E-03 | 3.93E-03 | 2.13E-03 | 5.07E-03 |
| **VANDQMALGAMR** | 1.55E-07 | 5.15E-05 | 2.58E-04 | 2.12E-05 | 8.95E-05 | 1.68E-05 | 4.39E-04 | 2.65E-04 | 6.58E-04 |
| **AITESGLRVGAD** | 6.24E-06 | 3.67E-05 | 3.78E-04 | 4.15E-05 | 9.11E-05 | 6.80E-05 | 2.97E-04 | 9.40E-04 | 1.36E-03 |
| **ITESGLRVGAD** | 4.94E-06 | 2.55E-04 | 1.54E-04 | 2.24E-04 | 2.45E-04 | 1.66E-04 | 2.94E-04 | 2.92E-04 | 4.04E-04 |
| **ADISVVGY** | 5.91E-06 | 1.17E-04 | 1.73E-04 | 1.64E-04 | 8.68E-05 | 5.68E-05 | 8.47E-05 | 4.59E-05 | 4.59E-05 |
| **VVGYDD** | 2.23E-06 | 1.05E-04 | 7.15E-05 | 7.34E-05 | 9.01E-05 | 2.71E-05 | 1.45E-04 | 1.50E-04 | 1.99E-04 |
| **DDTEDSSCYIPPLTTIK** | 3.86E-07 | 2.42E-05 | 1.70E-04 | 1.97E-05 | 1.42E-04 | 6.57E-06 | 1.21E-04 | 1.87E-04 | 7.53E-04 |
| **TEDSSCYIPPLTTIK** | 4.45E-07 | 2.51E-05 | 3.41E-05 | 8.75E-06 | 8.86E-05 | 1.06E-06 | 1.15E-04 | 5.74E-05 | 3.13E-04 |
| **SSCYIPPLTTIKQD** | 1.86E-06 | 5.13E-05 | 1.12E-04 | 1.68E-05 | 3.90E-05 | 7.35E-06 | 1.42E-04 | 9.49E-05 | 1.54E-04 |
| **FRLL** | 4.95E-07 | 2.84E-06 | 6.17E-05 | 1.89E-06 | 3.33E-06 | 1.51E-05 | 8.49E-05 | 4.14E-05 | 3.64E-04 |
| **FRLLGQTSV** | 5.55E-05 | 1.19E-05 | nan | 5.75E-06 | 9.58E-06 | 4.35E-05 | 4.25E-05 | 6.59E-06 | 1.28E-04 |
| **LGQTSVDR** | 1.14E-07 | 2.10E-05 | 1.06E-05 | 3.70E-05 | 2.12E-05 | 9.66E-06 | 2.71E-05 | 4.72E-05 | 7.69E-05 |
| **GQTSVDRL** | 1.64E-07 | 9.39E-06 | 1.01E-05 | 9.71E-06 | 2.08E-05 | 8.39E-06 | 3.64E-05 | 3.93E-05 | 4.06E-05 |
| **LLQL** | 5.04E-06 | 4.82E-06 | 8.83E-04 | 3.40E-05 | 6.20E-05 | 1.06E-06 | 4.52E-04 | 2.07E-04 | 2.99E-03 |

### DNA vs TMG

| Peptide | 0 s | 30 s | 45 s | 60 s | 300 s | 1500 s | 3600 s | 7200 s | 14400 s |
|---|---|---|---|---|---|---|---|---|---|
| LIGVA | 4.59E-07 | 1.34E-05 | 1.84E-05 | 1.38E-05 | 2.55E-05 | 1.04E-06 | 1.39E-05 | 2.20E-06 | 1.37E-05 |
| ALHAP | 1.84E-06 | 5.70E-05 | 2.46E-04 | 2.69E-04 | 5.99E-05 | 1.49E-04 | 3.95E-04 | 7.41E-04 | nan |
| ALHAPSQIVA | 8.15E-07 | 3.23E-05 | nan | 7.53E-04 | 3.50E-05 | 2.22E-04 | 1.27E-04 | 3.81E-04 | 2.12E-04 |
| ALHAPSQIVAA | 3.65E-06 | 7.69E-05 | 1.80E-04 | 7.30E-04 | 7.78E-05 | 2.14E-04 | 1.58E-04 | 5.13E-04 | 5.89E-04 |
| ALHAPSQIVAAIK | 5.84E-06 | 1.43E-04 | 3.88E-04 | nan | 1.84E-04 | 6.99E-05 | 1.68E-03 | 3.76E-04 | 2.04E-03 |
| SQIVAA | 1.55E-05 | 2.31E-05 | 4.48E-05 | 1.04E-04 | nan | 1.06E-04 | 9.14E-05 | 7.92E-05 | 4.51E-04 |
| SQIVAAIK | 4.73E-06 | 2.03E-05 | nan | 9.11E-05 | 2.26E-05 | 2.78E-05 | 1.15E-04 | 5.92E-05 | 5.52E-04 |
| IVAAIK | 5.13E-06 | 5.56E-06 | 5.45E-05 | 4.30E-05 | 4.98E-06 | 1.51E-05 | 3.56E-05 | 1.61E-05 | 1.83E-04 |
| AIKSRADQLGAS | 9.16E-07 | 2.72E-04 | 2.31E-04 | 1.76E-04 | 1.75E-04 | 2.26E-04 | 4.22E-04 | 1.06E-03 | 1.32E-03 |
| ADQLGAS | 2.65E-05 | 2.17E-04 | 6.83E-04 | 5.91E-04 | 2.56E-05 | 7.44E-04 | 1.43E-03 | nan | 4.46E-03 |
| VVVSM | 6.17E-05 | 2.31E-04 | 7.04E-04 | 1.91E-04 | 1.44E-04 | 7.08E-05 | 1.91E-04 | 1.02E-03 | 7.49E-04 |
| MVERSGVE | 1.05E-05 | 2.53E-03 | nan | 3.82E-04 | 9.70E-04 | 1.21E-03 | 1.43E-03 | 1.54E-03 | 8.67E-04 |
| AAVHNL | 8.51E-06 | 1.89E-05 | 4.54E-04 | 3.57E-05 | 2.36E-04 | 4.11E-05 | 6.05E-04 | 3.30E-04 | 6.82E-04 |
| LAQRVSGL | 2.41E-07 | 1.82E-04 | 2.31E-04 | 4.57E-05 | 3.26E-05 | 5.14E-05 | 1.74E-04 | 3.89E-04 | 8.00E-04 |
| VSGLIIN | 1.56E-05 | 3.64E-06 | 5.00E-05 | 1.01E-05 | 2.02E-06 | 1.33E-05 | 3.02E-05 | 7.16E-05 | 1.20E-05 |
| IINYPLD | 2.39E-05 | 1.65E-05 | 1.33E-05 | nan | 1.92E-05 | 5.00E-05 | 1.70E-05 | 5.57E-05 | 5.70E-04 |
| LDDQDAIA | 6.26E-06 | 7.48E-04 | 4.55E-04 | 4.77E-04 | 3.78E-04 | 4.44E-04 | 7.68E-04 | 1.12E-03 | 2.77E-03 |
| IAVEAA | 2.27E-07 | nan | 9.97E-04 | 9.32E-07 | 3.92E-05 | 1.10E-04 | 3.29E-05 | 1.75E-04 | 1.23E-04 |
| ACTNVPAL | 4.17E-05 | 1.60E-03 | 3.35E-04 | 1.03E-04 | 1.65E-04 | 7.20E-04 | 8.68E-04 | 2.83E-03 | 4.07E-03 |
| ALFLD | 1.01E-05 | 3.49E-05 | 2.68E-04 | 3.27E-05 | 5.47E-05 | 3.20E-05 | 8.20E-05 | 5.15E-05 | 1.04E-04 |
| FLDVSDQTPINS | 5.27E-06 | 2.28E-04 | nan | 8.16E-04 | 1.06E-04 | 5.82E-04 | 4.86E-04 | 8.67E-04 | 2.92E-04 |
| DQTPIN | 7.15E-06 | 4.70E-05 | 6.23E-04 | 4.20E-04 | 3.10E-05 | 4.62E-04 | 4.62E-04 | 9.69E-04 | 2.10E-03 |
| SIIFSH | 3.24E-04 | 1.29E-04 | 2.36E-04 | 7.73E-04 | 4.55E-05 | 3.38E-04 | 2.63E-04 | 1.81E-05 | 8.02E-04 |
| SIIFSHEDGTRL | 3.92E-07 | 2.03E-04 | 5.83E-06 | 2.63E-04 | 4.37E-05 | 6.49E-05 | 2.89E-05 | 8.10E-05 | 1.13E-04 |
| EDGTRLGVEH | 2.19E-05 | 7.44E-05 | 3.20E-04 | nan | 1.03E-04 | 4.67E-04 | 1.17E-04 | 6.93E-05 | 5.57E-04 |
| LVAL | 7.73E-08 | 1.93E-06 | 2.19E-05 | 1.44E-04 | 1.17E-06 | 1.75E-05 | 1.30E-05 | 3.30E-06 | 9.19E-05 |
| LVALGHQQIAL | 7.65E-06 | 2.98E-04 | 1.14E-04 | 3.60E-04 | 4.39E-05 | 2.26E-04 | 2.42E-04 | 5.28E-04 | 1.03E-03 |
| LGHQQIALL | 7.05E-06 | 3.68E-05 | 2.51E-05 | 2.28E-05 | 3.22E-05 | 6.87E-05 | 3.48E-05 | 6.61E-05 | 4.31E-05 |
| IALL | 4.71E-06 | 7.70E-06 | nan | 7.39E-05 | 4.31E-06 | 2.38E-05 | 2.24E-06 | 3.35E-06 | 2.13E-06 |

| | | | | | | | | | |
|---|---|---|---|---|---|---|---|---|---|
| **LAGPLS** | 2.72E-05 | 4.74E-04 | 1.77E-04 | 4.26E-04 | 2.61E-05 | nan | 3.49E-04 | 9.37E-04 | nan |
| **LAGPLSSVSARL** | 3.28E-04 | 4.24E-04 | 3.50E-04 | 2.78E-04 | 7.20E-05 | 8.17E-04 | 3.78E-04 | nan | nan |
| **GPLSSVSAR** | 4.59E-06 | 4.00E-04 | 9.41E-05 | nan | 1.06E-05 | 1.44E-03 | 4.49E-04 | 2.03E-03 | nan |
| **LAGWHK** | 1.06E-04 | 2.11E-04 | 3.19E-04 | nan | 5.06E-05 | 5.23E-04 | 2.81E-05 | 2.90E-04 | 3.64E-04 |
| **YLTRNQIQP** | 1.77E-06 | 4.15E-04 | nan | 9.17E-05 | 1.37E-04 | nan | 5.11E-04 | nan | nan |
| **NQIQPIAEREGD** | 3.62E-06 | 9.14E-04 | 7.94E-05 | 3.61E-04 | nan | 4.90E-04 | nan | 1.07E-03 | 4.05E-03 |
| **IAEREGDWSAMSG** | 1.42E-06 | 6.52E-04 | 7.45E-04 | 7.04E-04 | 2.73E-04 | 9.29E-04 | 1.38E-03 | 1.64E-03 | 4.72E-03 |
| **REGDWSAMSGF** | 7.32E-06 | 3.60E-04 | 5.24E-04 | 3.60E-04 | nan | 6.50E-04 | nan | 1.02E-03 | 1.22E-03 |
| **DWSAM** | 1.82E-05 | 1.56E-04 | 5.02E-04 | 7.37E-04 | 1.06E-04 | 4.06E-04 | 6.63E-04 | 1.15E-03 | nan |
| **WSAM** | 7.80E-05 | 9.70E-04 | 9.77E-04 | 2.67E-04 | 2.87E-04 | nan | 1.09E-03 | nan | nan |
| **SAMSGFQ** | 2.46E-05 | 4.17E-04 | 1.74E-03 | nan | 1.65E-05 | 1.41E-04 | 2.78E-04 | nan | nan |
| **SGFQQTMQM** | 7.06E-06 | 8.44E-06 | 2.05E-05 | nan | 1.49E-05 | 1.99E-05 | 1.01E-05 | 1.10E-05 | 5.70E-05 |
| **QTMQML** | 1.32E-05 | 1.19E-05 | nan | nan | 6.77E-07 | 6.46E-06 | 2.59E-06 | 5.59E-06 | 1.61E-05 |
| **NEGIVPTAML** | 2.11E-05 | 2.67E-04 | 4.76E-03 | 1.33E-05 | 2.53E-04 | 2.35E-04 | 2.32E-03 | 1.89E-03 | 6.78E-03 |
| **LVANDQMALGAM** | 3.41E-07 | 1.81E-04 | nan | nan | 1.72E-04 | 3.52E-04 | 1.32E-04 | 1.14E-04 | 8.72E-05 |
| **VANDQMALGAMR** | 8.09E-07 | 1.18E-04 | 2.09E-04 | nan | 8.02E-05 | 2.12E-04 | 2.67E-04 | 1.43E-04 | 6.70E-04 |
| **AITESGLRVGAD** | 1.46E-05 | 7.86E-04 | 3.07E-04 | 5.42E-04 | 2.50E-04 | 7.33E-04 | 4.65E-04 | 1.42E-03 | 1.68E-03 |
| **ITESGLRVGAD** | 6.92E-06 | 1.14E-03 | 1.52E-04 | 4.29E-04 | 3.95E-04 | 7.68E-04 | 4.71E-04 | 9.98E-04 | 6.59E-04 |
| **ADISVVGY** | 6.28E-06 | 9.90E-05 | 1.12E-04 | nan | 7.48E-05 | 2.12E-04 | 6.20E-05 | 1.39E-05 | 8.05E-05 |
| **VVGYDD** | 2.37E-06 | 8.15E-05 | 3.00E-05 | nan | 9.21E-05 | 1.04E-04 | 1.14E-04 | 7.51E-05 | 2.46E-04 |
| **DDTEDSSCYIPPLTTIK** | 3.27E-07 | 1.63E-04 | 1.02E-04 | nan | 7.26E-05 | 2.66E-04 | 1.03E-04 | 7.92E-04 | 1.50E-03 |
| **TEDSSCYIPPLTTIK** | 2.84E-06 | 7.05E-05 | 1.45E-04 | nan | 3.15E-05 | 9.43E-05 | 6.85E-05 | 3.10E-04 | 6.01E-04 |
| **SSCYIPPLTTIKQD** | 1.75E-06 | 7.91E-05 | 2.23E-04 | 3.31E-03 | 3.34E-05 | 2.37E-04 | 1.49E-04 | 1.38E-04 | 3.43E-04 |
| **FRLL** | 2.64E-07 | 3.82E-06 | 1.08E-04 | 1.59E-04 | 6.39E-06 | 1.36E-05 | 6.12E-05 | 1.35E-04 | 4.09E-04 |
| **FRLLGQTSV** | 5.20E-06 | 5.52E-05 | nan | 1.05E-04 | 3.65E-05 | 5.11E-05 | nan | 2.35E-05 | 1.16E-04 |
| **LGQTSVDR** | 3.36E-07 | 2.48E-05 | 1.32E-05 | 1.84E-04 | 2.22E-05 | 6.17E-05 | 2.44E-05 | 2.95E-05 | 9.72E-05 |
| **GQTSVDRL** | 1.64E-07 | nan | nan | 8.39E-05 | nan | 6.39E-05 | 3.40E-05 | 3.34E-05 | nan |
| **LLQL** | 5.57E-06 | 5.62E-05 | 1.05E-03 | 8.81E-05 | 1.81E-05 | 1.24E-04 | 3.17E-04 | 9.51E-04 | 2.71E-03 |

### IPTG vs APO

| Peptide | 0 s | 30 s | 45 s | 60 s | 300 s | 1500 s | 3600 s | 7200 s | 14400 s |
|---|---|---|---|---|---|---|---|---|---|
| LIGVA | 8.88E-06 | 2.55E-05 | 6.88E-05 | 5.18E-05 | 3.29E-05 | 5.51E-06 | 3.77E-06 | 1.04E-05 | 5.68E-05 |
| ALHAP | 2.64E-07 | 2.36E-04 | 7.46E-05 | 5.21E-04 | 9.31E-04 | 5.64E-04 | 4.26E-04 | 6.68E-04 | 4.05E-04 |
| ALHAPSQIVA | 3.18E-06 | 2.41E-05 | nan | 2.94E-05 | 2.42E-05 | 2.99E-05 | 8.52E-05 | 8.56E-05 | 2.23E-05 |
| ALHAPSQIVAA | 5.12E-06 | 3.60E-05 | nan | 7.00E-05 | 9.12E-05 | 4.32E-05 | 1.02E-04 | 2.89E-04 | 1.24E-03 |
| ALHAPSQIVAAIK | 1.05E-05 | nan | 5.28E-05 | 1.97E-04 | 3.29E-05 | 2.86E-05 | 7.15E-05 | 5.01E-04 | 2.06E-04 |
| SQIVAA | 3.79E-06 | 1.13E-05 | 1.79E-06 | 8.51E-05 | 1.12E-04 | 2.52E-04 | 4.77E-06 | 7.64E-04 | 9.16E-05 |
| SQIVAAIK | 2.85E-06 | 1.76E-05 | 1.54E-05 | 1.83E-04 | 5.20E-05 | 4.55E-04 | 1.99E-05 | 2.51E-04 | 4.08E-05 |
| IVAAIK | 2.62E-06 | 1.06E-05 | 6.59E-07 | 1.46E-04 | 7.53E-06 | 2.18E-04 | 2.62E-05 | 7.97E-05 | 3.39E-04 |
| AIKSRADQLGAS | 2.41E-06 | 1.06E-04 | 1.50E-04 | 9.79E-05 | 1.36E-04 | 1.51E-04 | 2.83E-04 | 4.79E-04 | 5.11E-04 |
| ADQLGAS | 1.43E-05 | nan | 5.74E-04 | nan | 1.36E-04 | nan | 2.12E-03 | 2.30E-03 | 4.99E-03 |
| VVVSM | 3.64E-07 | 5.95E-04 | 1.88E-04 | 5.33E-05 | 1.09E-03 | 5.58E-04 | 2.98E-04 | 9.34E-04 | 9.58E-04 |
| MVERSGVE | 8.78E-07 | 1.30E-03 | nan | 1.42E-03 | 7.71E-04 | 8.01E-04 | 8.76E-04 | 1.01E-03 | 8.92E-04 |
| AAVHNL | 5.89E-05 | 2.65E-05 | 7.28E-05 | 2.57E-05 | 2.55E-04 | 1.29E-03 | 1.44E-03 | 2.96E-03 | 1.85E-03 |
| LAQRVSGL | 1.98E-06 | 2.57E-04 | 2.05E-04 | 2.04E-06 | 2.41E-04 | 3.21E-04 | 2.28E-04 | 6.19E-04 | 6.01E-04 |
| VSGLIIN | 5.11E-05 | 1.61E-05 | 5.25E-05 | 9.84E-05 | 4.98E-05 | 9.35E-06 | 1.33E-04 | 9.76E-05 | 1.28E-05 |
| IINYPLD | 3.35E-06 | 8.91E-06 | 5.50E-06 | 1.18E-05 | 3.24E-05 | 6.34E-06 | 7.56E-05 | 1.02E-04 | 2.32E-04 |
| LDDQDAIA | 2.23E-05 | 1.14E-03 | 8.53E-04 | 7.05E-04 | nan | 7.06E-04 | 6.07E-04 | 1.35E-03 | 1.30E-03 |
| IAVEAA | 9.18E-07 | 1.84E-05 | 1.14E-05 | 2.04E-04 | 2.51E-05 | 9.92E-05 | 1.97E-04 | 7.35E-04 | 1.03E-03 |
| ACTNVPAL | 1.96E-05 | 1.59E-03 | 8.39E-04 | 2.16E-04 | 1.82E-03 | 7.56E-04 | 2.02E-04 | 2.06E-03 | 1.32E-03 |
| ALFLD | 4.67E-06 | 1.31E-05 | 3.11E-05 | 2.95E-05 | 2.64E-05 | 1.10E-05 | 8.04E-05 | 6.48E-05 | 2.06E-04 |
| FLDVSDQTPINS | 8.83E-06 | 1.55E-04 | nan | 1.82E-04 | 1.04E-04 | 3.57E-04 | 3.08E-04 | 4.42E-04 | 2.14E-04 |
| DQTPIN | 7.73E-06 | 2.49E-04 | 1.08E-04 | 1.10E-04 | 6.77E-04 | 1.08E-03 | 5.05E-04 | 8.56E-04 | 6.81E-04 |
| SIIFSH | 2.77E-05 | 1.93E-04 | 1.73E-04 | 2.36E-04 | nan | 6.04E-05 | 1.56E-04 | 4.85E-04 | 3.48E-04 |
| SIIFSHEDGTRL | 5.70E-06 | 1.72E-05 | 7.16E-06 | 4.84E-06 | 4.44E-05 | 3.55E-05 | 4.31E-05 | 5.83E-05 | 1.05E-04 |
| EDGTRLGVEH | 2.19E-06 | 2.08E-05 | 2.31E-05 | 3.07E-05 | 1.32E-04 | 1.29E-05 | 5.21E-05 | 9.12E-05 | 6.11E-04 |
| LVAL | 1.14E-07 | 1.18E-06 | 2.27E-06 | 8.75E-06 | 9.66E-06 | 1.50E-05 | 4.79E-06 | 1.66E-06 | 6.39E-05 |
| LVALGHQQIAL | 1.35E-06 | 6.22E-04 | 3.66E-04 | 2.38E-04 | 4.60E-04 | 4.15E-04 | 3.61E-04 | 7.66E-04 | 1.12E-03 |
| LGHQQIALL | 4.20E-06 | 3.72E-05 | nan | 2.64E-05 | 6.38E-05 | 3.19E-05 | 3.92E-05 | 6.97E-05 | 3.56E-05 |
| IALL | 3.77E-06 | 7.50E-06 | 8.45E-06 | 9.18E-07 | 1.23E-05 | 6.68E-06 | 4.33E-06 | 6.23E-06 | 1.51E-05 |

| | | | | | | | | | |
|---|---|---|---|---|---|---|---|---|---|
| **LAGPLS** | 5.91E-07 | 5.10E-04 | 1.91E-04 | 1.46E-04 | 4.19E-04 | 8.12E-04 | 1.44E-04 | nan | 3.32E-04 |
| **LAGPLSSVSARL** | 1.55E-05 | 1.53E-04 | 1.96E-04 | 8.03E-05 | 1.82E-04 | 2.25E-04 | 7.59E-04 | 2.74E-04 | 6.19E-05 |
| **GPLSSVSAR** | 3.35E-06 | 5.70E-04 | 1.62E-04 | 1.99E-04 | nan | 1.36E-04 | 2.99E-03 | 2.88E-03 | 2.02E-03 |
| **LAGWHK** | 2.69E-06 | 2.03E-05 | 4.02E-06 | 6.92E-06 | 2.84E-05 | 3.49E-05 | 1.93E-05 | 4.21E-05 | 6.34E-05 |
| **YLTRNQIQP** | 7.04E-06 | 5.53E-04 | 1.31E-04 | 1.07E-04 | 6.95E-04 | 1.72E-03 | 4.60E-04 | 1.62E-03 | 2.45E-03 |
| **NQIQPIAEREGD** | 1.55E-06 | 7.31E-04 | 5.33E-04 | 3.01E-04 | 1.73E-03 | 8.23E-04 | 4.35E-04 | 1.25E-03 | 2.01E-03 |
| **IAEREGDWSAMSG** | 2.27E-06 | 1.12E-03 | 8.71E-04 | 9.98E-05 | 2.04E-03 | 1.93E-03 | 1.80E-03 | 2.66E-03 | 3.52E-03 |
| **REGDWSAMSGF** | 7.21E-06 | 6.85E-05 | 1.36E-04 | 6.83E-05 | 1.71E-04 | 1.73E-04 | 1.93E-04 | 2.39E-04 | 2.49E-04 |
| **DWSAM** | 1.18E-06 | 4.31E-04 | 2.67E-04 | 8.90E-05 | 8.64E-04 | 7.73E-04 | 5.74E-04 | 1.35E-03 | 1.29E-03 |
| **WSAM** | 3.11E-05 | 6.93E-04 | 5.91E-04 | 6.53E-05 | 9.89E-04 | 1.05E-03 | 5.34E-04 | nan | 1.30E-03 |
| **SAMSGFQ** | 1.49E-05 | 1.33E-03 | 2.05E-04 | 5.19E-04 | 1.17E-03 | 3.70E-05 | 1.52E-03 | nan | 2.59E-03 |
| **SGFQQTMQM** | 3.13E-06 | 1.78E-05 | 3.04E-05 | 4.95E-05 | 2.24E-05 | 2.20E-06 | 5.62E-06 | 1.14E-05 | 5.22E-05 |
| **QTMQML** | 8.91E-07 | 3.73E-06 | 3.28E-06 | 1.43E-05 | 4.26E-06 | 2.47E-06 | 2.52E-06 | 2.21E-05 | 1.18E-04 |
| **NEGIVPTAML** | 4.72E-05 | 4.45E-04 | 3.81E-04 | 3.10E-04 | 4.67E-04 | 5.15E-04 | 7.91E-04 | 1.69E-03 | 6.05E-03 |
| **LVANDQMALGAM** | 2.47E-05 | 2.17E-04 | nan | 3.52E-04 | 3.60E-04 | 3.62E-04 | 2.77E-04 | 2.98E-04 | 2.86E-04 |
| **VANDQMALGAMR** | 4.82E-07 | 1.31E-04 | 7.56E-05 | 1.25E-04 | 1.59E-04 | 5.17E-05 | 2.24E-05 | 1.43E-04 | 9.48E-04 |
| **AITESGLRVGAD** | 1.97E-06 | 9.69E-04 | 1.51E-04 | 3.84E-04 | 8.03E-04 | 6.88E-04 | 2.95E-04 | 1.18E-03 | 1.44E-03 |
| **ITESGLRVGAD** | 3.26E-06 | 2.51E-04 | 2.24E-04 | 2.01E-04 | 2.34E-04 | 1.93E-04 | 2.01E-04 | 2.78E-04 | 1.75E-04 |
| **ADISVVGY** | 8.31E-06 | 8.45E-05 | 2.56E-05 | 2.10E-04 | 1.17E-04 | 4.30E-05 | 4.65E-05 | 2.66E-05 | 4.32E-04 |
| **VVGYDD** | 2.86E-06 | 1.10E-04 | 1.20E-04 | 1.83E-04 | 1.58E-04 | 8.58E-05 | 3.27E-04 | 1.29E-04 | 3.38E-04 |
| **DDTEDSSCYIPPLTTIK** | 8.41E-07 | 2.70E-04 | 2.51E-04 | 2.08E-04 | 5.81E-04 | 5.93E-04 | 1.89E-04 | 8.19E-04 | 1.04E-03 |
| **TEDSSCYIPPLTTIK** | 3.36E-06 | 6.03E-05 | 6.72E-05 | 1.92E-04 | 2.17E-04 | 3.95E-04 | 3.33E-05 | 4.18E-04 | 3.57E-04 |
| **SSCYIPPLTTIKQD** | 1.06E-06 | 8.07E-05 | 8.33E-05 | 1.49E-04 | 2.42E-04 | 2.87E-04 | 3.54E-05 | 1.82E-04 | 3.62E-04 |
| **FRLL** | 3.36E-07 | 5.90E-06 | 9.02E-06 | 2.09E-05 | 2.73E-06 | 9.68E-05 | 3.17E-05 | 1.21E-04 | 2.23E-04 |
| **FRLLGQTSV** | 2.41E-05 | 3.33E-05 | nan | 2.93E-05 | 3.79E-05 | 3.54E-05 | 4.94E-05 | 3.65E-05 | 4.99E-05 |
| **LGQTSVDR** | 4.73E-07 | 5.74E-05 | 4.59E-05 | 7.67E-05 | 6.59E-05 | 2.51E-05 | 3.73E-05 | 7.02E-05 | 2.20E-04 |
| **GQTSVDRL** | 3.19E-06 | 2.79E-05 | 3.42E-05 | 3.82E-05 | 7.97E-05 | 5.12E-05 | 2.72E-05 | nan | nan |
| **LLQL** | 6.77E-07 | 1.49E-04 | 4.78E-05 | 1.65E-04 | 8.10E-05 | 4.83E-04 | 7.54E-05 | 6.34E-04 | 4.22E-04 |

### IPTG vs DNA

| Peptide | 0 s | 30 s | 45 s | 60 s | 300 s | 1500 s | 3600 s | 7200 s | 14400 s |
|---|---|---|---|---|---|---|---|---|---|
| LIGVA | 5.57E-06 | 1.59E-05 | 8.18E-05 | 3.85E-05 | 2.50E-05 | 4.34E-06 | 6.18E-07 | 3.30E-06 | 2.42E-05 |
| ALHAP | 1.70E-06 | 3.74E-05 | 2.47E-04 | 7.79E-05 | 1.08E-04 | 2.86E-04 | 2.66E-04 | 7.32E-04 | nan |
| ALHAPSQIVA | 1.00E-06 | 9.70E-06 | nan | 7.45E-05 | 4.57E-05 | 2.63E-05 | 1.05E-04 | 3.88E-05 | 5.84E-05 |
| ALHAPSQIVAA | 3.01E-06 | 4.60E-05 | nan | 1.38E-04 | 8.52E-05 | 5.69E-05 | 1.64E-04 | 1.85E-04 | 4.43E-04 |
| ALHAPSQIVAAIK | 8.39E-06 | nan | 2.30E-04 | 5.68E-05 | 6.04E-05 | 6.10E-05 | 5.68E-04 | 2.14E-04 | 2.04E-03 |
| SQIVAA | 5.76E-06 | 2.23E-05 | 1.35E-05 | 3.85E-05 | 3.72E-05 | 7.75E-05 | 5.68E-06 | 6.12E-05 | 4.57E-04 |
| SQIVAAIK | 5.76E-06 | 2.80E-05 | nan | 8.23E-06 | 7.33E-05 | 8.82E-07 | 1.14E-04 | 5.80E-05 | 5.35E-04 |
| IVAAIK | 3.31E-06 | 1.46E-05 | 5.08E-05 | 5.17E-06 | 9.98E-06 | 5.14E-07 | 2.69E-05 | 7.46E-06 | 1.81E-04 |
| AIKSRADQLGAS | 4.01E-07 | 8.56E-05 | 2.70E-04 | 2.43E-04 | 1.83E-04 | 1.45E-04 | 3.02E-04 | 3.88E-04 | 1.26E-03 |
| ADQLGAS | 2.70E-05 | nan | 7.18E-04 | nan | 1.39E-04 | nan | 2.03E-03 | nan | 5.35E-03 |
| VVVSM | 6.23E-07 | 2.40E-04 | 6.20E-04 | 1.75E-04 | 2.13E-04 | 3.79E-04 | 1.93E-04 | 8.29E-04 | 1.02E-03 |
| MVERSGVE | 2.46E-07 | 8.40E-04 | nan | 7.52E-04 | 1.08E-03 | 7.82E-04 | 9.84E-04 | 1.20E-03 | 5.70E-04 |
| AAVHNL | 4.34E-05 | 4.42E-05 | 1.51E-04 | 2.60E-05 | 4.09E-04 | 1.29E-03 | 1.83E-03 | 1.19E-03 | 2.24E-03 |
| LAQRVSGL | 1.84E-06 | 2.41E-04 | 2.11E-04 | 3.82E-06 | 1.43E-04 | 3.21E-04 | 2.26E-04 | 3.77E-04 | 1.12E-03 |
| VSGLIIN | 1.56E-05 | 3.39E-06 | 2.16E-05 | 1.53E-05 | 2.48E-05 | 1.10E-06 | 5.26E-05 | 6.90E-05 | 8.74E-06 |
| IINYPLD | 8.57E-06 | 1.36E-05 | 1.68E-05 | 1.42E-05 | 4.98E-05 | 2.10E-05 | 6.90E-05 | 7.86E-05 | 6.26E-04 |
| LDDQDAIA | 2.23E-05 | 7.13E-04 | 6.66E-04 | 6.94E-04 | 7.29E-04 | 8.07E-04 | 9.07E-04 | 1.46E-03 | 3.55E-03 |
| IAVEAA | 7.73E-07 | nan | 8.99E-04 | 4.45E-06 | 3.95E-05 | 1.64E-04 | 8.48E-05 | 3.75E-04 | 9.43E-04 |
| ACTNVPAL | 2.47E-05 | 9.16E-04 | 3.36E-04 | 9.09E-07 | 3.23E-04 | 5.83E-04 | 5.59E-04 | 1.82E-03 | 4.17E-03 |
| ALFLD | 4.67E-06 | 4.44E-05 | 1.81E-04 | 4.42E-05 | 7.17E-05 | 4.22E-05 | 1.39E-04 | 4.77E-05 | 1.14E-04 |
| FLDVSDQTPINS | 5.76E-07 | 1.09E-04 | nan | 2.44E-04 | 1.74E-04 | 3.96E-04 | 4.58E-04 | 6.71E-04 | 4.05E-04 |
| DQTPIN | 1.22E-05 | 2.48E-04 | 5.55E-04 | 6.68E-05 | 9.27E-05 | 1.09E-03 | 5.73E-04 | 6.93E-04 | 2.39E-03 |
| SIIFSH | 1.01E-05 | 2.80E-04 | 3.18E-04 | 3.27E-04 | 1.16E-04 | 2.80E-04 | 3.02E-04 | 1.36E-04 | 9.05E-04 |
| SIIFSHEDGTRL | 4.08E-06 | 1.75E-05 | 7.46E-06 | 2.26E-05 | 3.47E-05 | 3.34E-05 | 4.31E-05 | 5.93E-05 | 1.08E-04 |
| EDGTRLGVEH | 6.59E-06 | 1.34E-05 | 2.11E-05 | 6.66E-05 | 1.34E-04 | 9.35E-06 | 1.50E-04 | 3.25E-05 | 1.73E-04 |
| LVAL | 4.55E-09 | 2.98E-06 | 1.75E-05 | 4.73E-06 | 1.04E-05 | 7.73E-07 | 1.13E-05 | 4.71E-06 | 1.18E-04 |
| LVALGHQQIAL | 4.02E-06 | 4.22E-04 | 2.45E-04 | 1.84E-04 | 3.19E-04 | 3.89E-04 | 4.46E-04 | 7.23E-04 | 1.40E-03 |
| LGHQQIALL | 4.51E-07 | 4.08E-05 | 9.62E-05 | 3.89E-05 | 6.92E-05 | 3.99E-05 | 3.94E-05 | 7.20E-05 | 5.05E-05 |
| IALL | 3.53E-06 | 1.70E-06 | nan | 6.77E-06 | 8.75E-06 | 8.47E-06 | 5.04E-06 | 4.55E-08 | 4.79E-06 |

| | | | | | | | | | |
|---|---|---|---|---|---|---|---|---|---|
| **LAGPLS** | 5.17E-06 | 3.25E-04 | 1.97E-04 | 8.40E-05 | 4.88E-05 | nan | 3.52E-04 | 9.28E-04 | nan |
| **LAGPLSSVSARL** | 3.04E-04 | 4.21E-04 | 4.51E-04 | 8.76E-05 | 1.22E-04 | 3.60E-04 | 3.54E-04 | 4.08E-04 | 7.73E-04 |
| **GPLSSVSAR** | 1.06E-06 | 2.23E-04 | 1.14E-04 | 1.08E-04 | 1.51E-05 | 1.16E-03 | 6.48E-04 | 1.51E-03 | 4.50E-03 |
| **LAGWHK** | 9.17E-05 | 4.43E-05 | 8.03E-05 | 2.22E-06 | 7.89E-05 | 5.05E-05 | 3.13E-05 | 3.06E-05 | 2.49E-06 |
| **YLTRNQIQP** | 6.02E-06 | 4.86E-04 | 3.52E-04 | 4.20E-05 | 3.06E-04 | 7.50E-04 | 4.86E-04 | 9.99E-04 | nan |
| **NQIQPIAEREGD** | 3.96E-06 | 3.68E-04 | 1.70E-04 | 1.39E-04 | 1.07E-03 | 5.29E-04 | 5.64E-04 | 1.03E-03 | 4.66E-03 |
| **IAEREGDWSAMSG** | 7.27E-07 | 9.56E-04 | 1.04E-03 | 9.95E-05 | 8.83E-04 | 1.08E-03 | 1.84E-03 | 1.60E-03 | 5.37E-03 |
| **REGDWSAMSGF** | 8.23E-06 | 4.98E-05 | 3.62E-04 | 1.17E-04 | 1.73E-04 | 5.49E-04 | 2.68E-04 | 3.83E-04 | 9.82E-04 |
| **DWSAM** | 5.24E-06 | 4.26E-04 | 5.64E-04 | 8.15E-05 | 3.67E-04 | 5.46E-04 | 4.08E-04 | 6.74E-04 | nan |
| **WSAM** | 9.44E-05 | 1.59E-03 | 1.09E-03 | 7.30E-05 | 5.95E-04 | nan | 9.23E-04 | 1.17E-03 | nan |
| **SAMSGFQ** | 1.04E-05 | 1.30E-03 | 1.07E-03 | nan | 6.92E-04 | 3.73E-06 | 5.11E-04 | nan | nan |
| **SGFQQTMQM** | 3.72E-06 | 2.36E-05 | 2.05E-05 | 4.95E-06 | 3.35E-05 | 8.53E-06 | 1.25E-05 | 1.88E-05 | 7.87E-05 |
| **QTMQML** | 1.14E-05 | 3.62E-06 | nan | 8.19E-06 | 1.86E-06 | 1.55E-06 | 8.82E-07 | 4.67E-06 | 5.08E-05 |
| **NEGIVPTAML** | 4.63E-05 | 4.35E-04 | 3.64E-03 | 5.91E-06 | 4.75E-04 | 5.13E-04 | 2.48E-03 | 1.36E-03 | 7.47E-03 |
| **LVANDQMALGAM** | 2.17E-06 | 1.53E-04 | nan | 3.15E-04 | 2.64E-04 | 2.46E-04 | 1.58E-04 | 1.95E-04 | 1.74E-04 |
| **VANDQMALGAMR** | 1.55E-07 | 1.30E-04 | 2.84E-04 | 1.32E-04 | 2.06E-04 | 6.69E-05 | 2.74E-04 | 1.64E-04 | 7.58E-04 |
| **AITESGLRVGAD** | 6.88E-06 | 6.62E-04 | 4.53E-04 | 2.86E-04 | 4.19E-04 | 7.01E-04 | 4.52E-04 | 1.10E-03 | 1.92E-03 |
| **ITESGLRVGAD** | 8.15E-07 | 2.19E-04 | 1.83E-04 | 2.18E-04 | 2.88E-04 | 2.66E-04 | 2.96E-04 | 3.32E-04 | 4.67E-04 |
| **ADISVVGY** | 1.33E-05 | 1.60E-04 | 1.31E-04 | 3.69E-04 | 1.91E-04 | 7.87E-05 | 1.03E-04 | 3.39E-05 | 8.82E-05 |
| **VVGYDD** | 2.46E-06 | 1.63E-04 | 1.25E-04 | 2.35E-04 | 2.21E-04 | 9.14E-05 | 4.04E-04 | 1.14E-04 | 2.60E-04 |
| **DDTEDSSCYIPPLTTIK** | 7.14E-07 | 2.16E-04 | 1.77E-04 | 1.90E-04 | 4.80E-04 | 4.24E-04 | 1.18E-04 | 4.85E-04 | 1.20E-03 |
| **TEDSSCYIPPLTTIK** | 2.42E-06 | 7.45E-05 | 8.66E-05 | 6.37E-05 | 2.41E-04 | 1.14E-04 | 7.65E-05 | 2.19E-04 | 4.97E-04 |
| **SSCYIPPLTTIKQD** | 2.66E-06 | 6.70E-05 | 1.10E-04 | 1.28E-04 | 2.42E-04 | 6.57E-05 | 1.64E-04 | 1.38E-04 | 3.32E-04 |
| **FRLL** | 1.55E-07 | 7.71E-06 | 6.34E-05 | 2.27E-06 | 5.93E-06 | 2.83E-05 | 7.97E-05 | 1.15E-04 | 4.44E-04 |
| **FRLLGQTSV** | 3.84E-06 | 2.30E-05 | nan | 2.87E-06 | 3.16E-05 | 1.03E-06 | 7.72E-05 | 2.10E-05 | 1.32E-04 |
| **LGQTSVDR** | 5.68E-07 | 5.74E-05 | 3.15E-05 | 9.01E-05 | 6.59E-05 | 2.94E-05 | 4.40E-05 | 4.52E-05 | 1.00E-04 |
| **GQTSVDRL** | 3.07E-06 | 2.30E-05 | 2.96E-05 | 3.57E-05 | 8.80E-05 | 3.05E-05 | 5.98E-05 | nan | nan |
| **LLQL** | 3.79E-06 | 1.49E-04 | 7.89E-04 | 4.10E-05 | 3.09E-05 | 4.29E-04 | 2.17E-04 | 4.17E-04 | 2.70E-03 |

### IPTG vs ONPF

| Peptide | 0 s | 30 s | 45 s | 60 s | 300 s | 1500 s | 3600 s | 7200 s | 14400 s |
|---|---|---|---|---|---|---|---|---|---|
| LIGVA | 7.04E-06 | 5.69E-06 | 9.01E-05 | 3.76E-05 | 1.70E-05 | 1.76E-05 | 4.73E-07 | 7.06E-06 | 5.02E-05 |
| ALHAP | 7.73E-07 | 9.08E-05 | 3.24E-04 | 6.75E-05 | 3.65E-04 | 5.66E-04 | 2.16E-04 | 4.42E-04 | 2.12E-03 |
| ALHAPSQIVA | 7.30E-06 | 2.02E-05 | nan | 3.56E-05 | 2.51E-05 | 2.98E-05 | 1.37E-04 | 4.69E-05 | 2.78E-05 |
| ALHAPSQIVAA | 7.35E-07 | 2.76E-05 | nan | 1.64E-04 | 3.34E-05 | 9.40E-05 | 1.04E-04 | 1.61E-04 | 2.45E-05 |
| ALHAPSQIVAAIK | nan | nan | nan | nan | nan | nan | nan | nan | nan |
| SQIVAA | 5.86E-05 | nan | nan | 5.59E-05 | nan | nan | 7.21E-04 | 2.98E-04 | nan |
| SQIVAAIK | 2.31E-06 | 1.63E-05 | nan | 6.30E-06 | 1.02E-04 | 4.35E-05 | 1.11E-03 | 2.18E-04 | nan |
| IVAAIK | 4.09E-08 | 1.19E-05 | 1.79E-04 | 5.60E-06 | 2.07E-05 | 1.55E-05 | 2.29E-04 | 4.69E-05 | 1.63E-03 |
| AIKSRADQLGAS | 1.36E-06 | 6.26E-05 | 2.82E-04 | 8.93E-05 | 6.63E-05 | 1.89E-04 | 2.67E-04 | 4.16E-04 | 5.73E-04 |
| ADQLGAS | 2.57E-04 | nan | 6.40E-05 | nan | nan | nan | 2.62E-03 | nan | nan |
| VVVSM | 1.05E-05 | 3.79E-04 | 1.07E-03 | 1.11E-04 | 8.55E-04 | 7.90E-04 | 3.19E-04 | 5.33E-04 | 1.90E-03 |
| MVERSGVE | 5.16E-06 | 7.20E-04 | nan | 5.63E-04 | 5.84E-04 | 5.74E-04 | 4.81E-04 | 6.68E-04 | 3.57E-04 |
| AAVHNL | 4.44E-05 | 4.03E-05 | 5.99E-05 | 3.18E-05 | 2.09E-04 | 1.44E-03 | 1.46E-03 | 1.10E-03 | nan |
| LAQRVSGL | 2.19E-06 | 2.33E-04 | 5.82E-04 | 7.07E-06 | 2.78E-04 | 4.34E-04 | 4.09E-04 | 3.90E-04 | 4.59E-03 |
| VSGLIIN | 2.48E-05 | 4.33E-06 | 5.30E-06 | 1.68E-05 | 2.44E-05 | 2.02E-06 | 2.50E-05 | 9.60E-06 | 8.74E-06 |
| IINYPLD | 6.63E-06 | 9.84E-06 | 7.15E-06 | 1.49E-05 | 4.53E-05 | 6.34E-06 | 5.51E-05 | 4.09E-05 | 3.63E-04 |
| LDDQDAIA | 2.24E-05 | 9.58E-04 | 1.41E-03 | 3.69E-04 | 9.87E-04 | 1.01E-03 | 8.93E-04 | 1.01E-03 | nan |
| IAVEAA | 1.71E-06 | 5.60E-05 | 5.64E-05 | 4.45E-06 | 4.95E-05 | 1.14E-04 | 3.81E-04 | 3.61E-04 | 4.80E-03 |
| ACTNVPAL | 7.50E-06 | 9.59E-04 | 1.09E-03 | 1.85E-05 | 1.09E-03 | 1.14E-03 | 8.15E-04 | 8.98E-04 | nan |
| ALFLD | 5.82E-06 | 3.63E-05 | 4.30E-05 | 3.19E-05 | 2.93E-05 | 1.68E-05 | 8.35E-05 | 2.52E-04 | nan |
| FLDVSDQTPINS | 7.57E-06 | 1.57E-04 | nan | 2.26E-04 | 1.96E-04 | 4.15E-04 | 5.25E-04 | 8.03E-04 | 1.49E-03 |
| DQTPIN | 2.77E-05 | 2.62E-04 | 3.78E-04 | 7.16E-05 | 6.18E-04 | 1.68E-03 | 6.05E-04 | 5.56E-04 | nan |
| SIIFSH | nan | nan | nan | nan | nan | nan | nan | nan | nan |
| SIIFSHEDGTRL | 4.78E-06 | 1.45E-05 | nan | 1.14E-05 | 2.77E-05 | 1.74E-05 | 3.45E-05 | 4.67E-05 | 1.32E-04 |
| EDGTRLGVEH | 4.48E-06 | 1.47E-05 | 6.40E-05 | 3.06E-05 | 1.57E-04 | 3.36E-05 | 1.62E-04 | 7.40E-05 | nan |
| LVAL | 5.50E-07 | 1.16E-06 | 7.15E-06 | 4.82E-06 | 9.43E-06 | 7.86E-07 | 3.97E-06 | 2.41E-06 | 2.63E-05 |
| LVALGHQQIAL | 9.09E-07 | 4.40E-04 | 4.18E-04 | 1.77E-04 | 4.00E-04 | 4.54E-04 | 3.64E-04 | 5.34E-04 | 1.46E-03 |
| LGHQQIALL | 1.05E-05 | 4.14E-05 | 7.35E-05 | 2.66E-05 | 5.32E-05 | 4.39E-05 | 5.19E-05 | 6.20E-05 | 3.56E-05 |
| IALL | 1.62E-06 | 3.79E-06 | 1.06E-06 | 2.55E-06 | 5.02E-06 | 2.86E-06 | 7.13E-06 | 5.26E-06 | nan |

| | | | | | | | | | |
|---|---|---|---|---|---|---|---|---|---|
| **LAGPLS** | 2.06E-05 | 3.08E-04 | 1.32E-04 | 8.04E-05 | 2.17E-04 | 1.42E-03 | 1.28E-03 | 9.37E-04 | nan |
| **LAGPLSSVSARL** | 1.61E-05 | 1.48E-04 | 7.38E-04 | 1.09E-04 | 4.08E-04 | 5.54E-04 | 6.56E-04 | 6.22E-04 | 2.88E-03 |
| **GPLSSVSAR** | 1.86E-06 | 3.62E-04 | 1.77E-03 | 1.24E-04 | 2.16E-04 | 1.35E-04 | 8.90E-04 | 6.68E-04 | 3.01E-03 |
| **LAGWHK** | 2.40E-05 | 1.37E-06 | 8.82E-06 | 8.11E-07 | 2.86E-05 | 3.72E-05 | 1.90E-05 | 2.45E-05 | 2.49E-06 |
| **YLTRNQIQP** | 6.73E-06 | 4.89E-04 | 1.31E-03 | 4.34E-05 | 5.19E-04 | 7.94E-04 | 4.10E-04 | 6.54E-04 | 1.33E-03 |
| **NQIQPIAEREGD** | 1.11E-06 | 3.86E-04 | 8.25E-04 | 1.33E-04 | 2.05E-03 | 8.60E-04 | 5.09E-04 | 6.94E-04 | nan |
| **IAEREGDWSAMSG** | 3.39E-06 | 1.08E-03 | 1.41E-03 | 1.17E-04 | 1.62E-03 | 1.94E-03 | 1.60E-03 | 1.23E-03 | 8.56E-03 |
| **REGDWSAMSGF** | 6.55E-06 | 5.87E-05 | nan | 1.61E-04 | 2.23E-04 | 1.88E-04 | 5.46E-04 | 1.74E-04 | 1.40E-04 |
| **DWSAM** | 7.20E-06 | 4.36E-04 | 3.86E-04 | 7.28E-05 | 1.00E-03 | 1.16E-03 | 5.65E-04 | 5.15E-04 | nan |
| **WSAM** | 8.85E-05 | 7.36E-04 | 9.14E-04 | 6.12E-05 | 1.10E-03 | 1.50E-03 | 6.28E-04 | nan | 5.55E-04 |
| **SAMSGFQ** | 5.20E-06 | 1.53E-03 | nan | 4.78E-04 | 2.55E-03 | 2.30E-04 | 3.28E-04 | nan | nan |
| **SGFQQTMQM** | 3.26E-06 | 1.90E-05 | 3.53E-05 | 2.08E-05 | 2.25E-05 | 3.38E-06 | 1.09E-05 | 2.41E-05 | 2.98E-05 |
| **QTMQML** | 4.09E-06 | 1.11E-05 | 5.73E-06 | 7.14E-06 | 1.75E-06 | 1.55E-06 | 1.43E-06 | 6.66E-06 | 3.60E-05 |
| **NEGIVPTAML** | 4.57E-05 | 5.59E-04 | 2.50E-03 | 3.97E-06 | 7.63E-04 | 1.01E-03 | 1.76E-03 | 7.43E-04 | 1.31E-03 |
| **LVANDQMALGAM** | 3.25E-06 | 1.14E-04 | nan | 3.07E-04 | 2.29E-04 | 4.06E-04 | 2.06E-04 | 2.32E-04 | 1.86E-04 |
| **VANDQMALGAMR** | 4.82E-07 | 1.11E-04 | 1.81E-04 | 1.36E-04 | 2.04E-04 | 1.07E-04 | 1.65E-04 | 1.31E-04 | 6.99E-04 |
| **AITESGLRVGAD** | 6.59E-07 | 7.33E-04 | 7.36E-04 | 2.68E-04 | 6.96E-04 | 8.20E-04 | 4.34E-04 | 7.00E-04 | 2.41E-03 |
| **ITESGLRVGAD** | 5.33E-06 | 1.72E-04 | 3.35E-04 | 8.53E-05 | 1.88E-04 | 1.40E-04 | 1.62E-04 | 1.46E-04 | 4.83E-04 |
| **ADISVVGY** | 8.91E-06 | 9.84E-05 | 2.95E-05 | 2.15E-04 | 1.20E-04 | 2.07E-04 | 6.56E-05 | 2.86E-05 | 2.10E-04 |
| **VVGYDD** | 9.09E-07 | 1.16E-04 | 1.54E-04 | 2.13E-04 | 2.05E-04 | 1.74E-04 | 5.96E-04 | 1.79E-04 | 1.01E-03 |
| **DDTEDSSCYIPPLTTIK** | 2.19E-06 | 2.44E-04 | 7.32E-04 | 2.33E-04 | 6.61E-04 | 7.39E-04 | 8.23E-04 | 8.13E-04 | nan |
| **TEDSSCYIPPLTTIK** | 2.24E-06 | 5.75E-05 | 4.79E-04 | 1.05E-04 | 2.38E-04 | 1.93E-04 | 1.84E-04 | 1.89E-04 | nan |
| **SSCYIPPLTTIKQD** | 1.39E-06 | 8.45E-05 | 4.40E-04 | 1.57E-04 | 2.66E-04 | 1.58E-04 | 4.52E-04 | 2.58E-04 | 7.57E-03 |
| **FRLL** | 5.68E-07 | 7.36E-06 | 7.20E-05 | 3.77E-06 | 5.24E-06 | 3.19E-05 | 1.72E-04 | 1.24E-04 | 1.70E-03 |
| **FRLLGQTSV** | 4.48E-06 | 2.42E-05 | nan | 4.59E-05 | 2.72E-05 | 9.80E-06 | 5.91E-05 | 3.44E-05 | 4.75E-05 |
| **LGQTSVDR** | 4.73E-07 | 5.28E-05 | 5.55E-05 | 6.36E-05 | 6.15E-05 | 3.16E-05 | 6.26E-05 | 1.31E-04 | 5.31E-04 |
| **GQTSVDRL** | 3.53E-06 | 2.18E-05 | 2.96E-05 | 3.34E-05 | 7.68E-05 | 3.12E-05 | 4.12E-05 | nan | nan |
| **LLQL** | 6.77E-07 | 1.84E-04 | 4.85E-04 | 4.61E-05 | 2.98E-04 | 9.03E-04 | 8.73E-04 | 7.37E-04 | 4.70E-03 |

### IPTG vs ONPFDNA

| Peptide | 0 s | 30 s | 45 s | 60 s | 300 s | 1500 s | 3600 s | 7200 s | 14400 s |
|---|---|---|---|---|---|---|---|---|---|
| LIGVA | 6.73E-06 | 2.55E-05 | 6.92E-05 | 3.59E-05 | 1.43E-05 | 7.97E-06 | 9.09E-07 | 3.57E-06 | 4.21E-05 |
| ALHAP | 5.14E-07 | 2.10E-05 | 7.80E-06 | 5.89E-05 | 1.17E-04 | 2.84E-04 | 9.93E-05 | 3.37E-04 | 2.85E-04 |
| ALHAPSQIVA | 3.25E-06 | 1.09E-04 | nan | 1.22E-04 | 5.04E-05 | 2.23E-04 | 1.31E-04 | 8.17E-05 | 3.46E-04 |
| ALHAPSQIVAA | 3.14E-07 | 1.01E-04 | nan | 1.48E-04 | 6.74E-05 | 1.57E-04 | 1.95E-04 | 1.37E-04 | 2.59E-04 |
| ALHAPSQIVAAIK | 7.94E-06 | nan | 1.44E-05 | 6.68E-05 | 7.53E-05 | 4.04E-05 | 1.78E-04 | 3.56E-04 | 2.62E-05 |
| SQIVAA | 2.78E-05 | 1.53E-05 | 2.24E-06 | 2.40E-05 | nan | 1.97E-04 | 1.31E-05 | 2.27E-06 | 1.54E-05 |
| SQIVAAIK | 2.27E-06 | 1.85E-05 | 2.76E-05 | 8.42E-06 | 5.17E-05 | 4.86E-06 | 1.93E-04 | 1.21E-04 | 4.83E-05 |
| IVAAIK | 1.16E-06 | 1.12E-05 | 5.91E-06 | 5.84E-06 | 6.11E-06 | 1.25E-06 | 1.71E-05 | 4.07E-05 | 7.57E-06 |
| AIKSRADQLGAS | 3.22E-07 | 1.15E-04 | 5.77E-05 | 9.96E-05 | 1.02E-04 | 2.85E-04 | 2.11E-04 | 2.66E-04 | 6.35E-04 |
| ADQLGAS | 4.48E-05 | nan | 6.25E-05 | nan | 5.39E-04 | nan | 1.96E-03 | 4.25E-03 | 2.58E-03 |
| VVVSM | 5.27E-07 | 2.64E-04 | 3.16E-04 | 1.13E-04 | 2.71E-04 | 4.45E-04 | 4.74E-05 | 4.04E-04 | 7.38E-04 |
| MVERSGVE | 6.64E-07 | 1.39E-03 | nan | 6.11E-04 | 5.73E-04 | 6.53E-04 | 4.99E-04 | 6.94E-04 | 6.55E-04 |
| AAVHNL | 6.98E-05 | 2.56E-05 | 2.19E-06 | 4.78E-05 | 3.74E-04 | 1.29E-03 | 1.49E-03 | nan | 1.66E-03 |
| LAQRVSGL | 1.98E-06 | 2.30E-04 | 8.92E-05 | 1.98E-06 | 1.39E-04 | 3.31E-04 | 1.85E-04 | 3.25E-04 | 4.78E-04 |
| VSGLIIN | 2.42E-05 | 8.79E-06 | 1.80E-05 | 8.47E-06 | 1.29E-04 | 1.86E-06 | 2.44E-05 | 3.52E-05 | 7.44E-06 |
| IINYPLD | 6.39E-06 | 8.18E-06 | 1.05E-05 | 2.61E-05 | 3.34E-05 | 4.06E-05 | 6.34E-05 | 3.25E-05 | 6.52E-05 |
| LDDQDAIA | 2.29E-05 | 6.66E-04 | 3.33E-04 | 3.24E-04 | 6.75E-04 | 8.23E-04 | 3.22E-04 | 7.45E-04 | 1.07E-03 |
| IAVEAA | 7.14E-07 | 1.78E-05 | 6.43E-05 | 1.54E-05 | 7.90E-06 | 2.23E-04 | 6.83E-05 | 3.64E-04 | 8.60E-04 |
| ACTNVPAL | 7.93E-06 | 9.63E-04 | 2.73E-06 | 1.63E-05 | 2.96E-04 | 5.89E-04 | 7.80E-05 | 7.19E-04 | 7.57E-04 |
| ALFLD | 6.13E-06 | 3.17E-05 | nan | 8.95E-05 | 4.60E-04 | 3.32E-05 | 1.85E-04 | 1.39E-04 | 2.78E-04 |
| FLDVSDQTPINS | 9.30E-07 | 2.20E-04 | nan | 7.46E-05 | 1.60E-04 | 2.58E-04 | 4.91E-04 | 3.42E-04 | 3.55E-04 |
| DQTPIN | 1.38E-05 | 2.67E-04 | 1.52E-04 | 7.09E-05 | 1.93E-04 | 1.10E-03 | 2.92E-04 | 3.73E-04 | 3.82E-04 |
| SIIFSH | 2.47E-04 | 3.77E-04 | 8.13E-04 | 5.90E-04 | 5.56E-04 | 4.83E-04 | 1.07E-03 | 9.57E-04 | nan |
| SIIFSHEDGTRL | 3.99E-06 | 1.63E-05 | nan | 1.48E-05 | 4.88E-05 | 2.87E-05 | 4.72E-05 | 7.39E-05 | 3.68E-05 |
| EDGTRLGVEH | 2.53E-06 | 2.19E-05 | 1.89E-05 | 3.75E-05 | 1.31E-04 | 2.22E-05 | 1.87E-04 | 2.82E-05 | 4.03E-05 |
| LVAL | 4.09E-08 | 1.45E-06 | 2.75E-06 | 4.77E-06 | 9.28E-06 | 4.08E-06 | 6.39E-06 | 2.01E-06 | 3.01E-05 |
| LVALGHQQIAL | 4.95E-07 | 4.18E-04 | 1.90E-04 | 1.74E-04 | 3.31E-04 | 3.77E-04 | 3.06E-04 | 6.40E-04 | 4.43E-04 |
| LGHQQIALL | 1.23E-06 | 6.19E-05 | 1.52E-04 | 8.14E-05 | 6.51E-05 | 2.35E-05 | 2.88E-05 | 5.37E-05 | 2.23E-05 |
| IALL | 1.93E-06 | 5.14E-07 | 6.95E-07 | 7.11E-06 | 5.31E-06 | 5.27E-07 | 3.68E-06 | 3.73E-07 | 2.69E-05 |

| | | | | | | | | | |
|---|---|---|---|---|---|---|---|---|---|
| **LAGPLS** | 2.04E-06 | 3.71E-04 | 2.74E-05 | 8.88E-05 | 4.21E-05 | 6.98E-04 | 1.81E-05 | nan | 1.67E-04 |
| **LAGPLSSVSARL** | 2.12E-06 | 1.88E-04 | 2.48E-04 | 5.86E-05 | 2.17E-04 | 2.14E-04 | 2.19E-04 | 5.68E-04 | 3.21E-04 |
| **GPLSSVSAR** | 1.14E-07 | 2.66E-04 | 3.29E-04 | 1.05E-04 | 1.33E-05 | 1.42E-04 | 4.94E-04 | 9.71E-04 | 1.73E-03 |
| **LAGWHK** | 6.26E-06 | 2.76E-05 | 4.56E-06 | 6.48E-06 | 5.59E-05 | 3.94E-05 | nan | 1.41E-05 | 8.98E-05 |
| **YLTRNQIQP** | 5.59E-06 | 5.31E-04 | 1.34E-04 | 9.01E-05 | 3.00E-04 | 8.72E-04 | 2.27E-04 | 6.57E-04 | nan |
| **NQIQPIAEREGD** | 9.64E-07 | 3.56E-04 | 9.24E-05 | 1.31E-04 | 1.09E-03 | 6.24E-04 | 2.18E-04 | 7.92E-04 | nan |
| **IAEREGDWSAMSG** | 8.41E-07 | 1.01E-03 | 6.43E-04 | 1.00E-04 | 1.05E-03 | 1.15E-03 | 1.03E-03 | 8.92E-04 | 1.21E-03 |
| **REGDWSAMSGF** | 2.86E-05 | 1.15E-04 | 4.84E-04 | 1.54E-04 | 1.60E-04 | 1.67E-04 | 1.76E-04 | 1.03E-04 | 1.83E-04 |
| **DWSAM** | 7.45E-07 | 4.20E-04 | 2.26E-04 | 3.60E-05 | 4.48E-04 | 7.34E-04 | 1.01E-04 | 3.08E-04 | 3.71E-04 |
| **WSAM** | 2.82E-05 | 6.68E-04 | 4.04E-04 | 1.52E-04 | 5.20E-04 | 7.31E-04 | 7.87E-05 | 4.13E-04 | nan |
| **SAMSGFQ** | 6.39E-05 | 1.27E-03 | 1.93E-04 | 4.13E-04 | 9.60E-04 | 2.64E-05 | 3.21E-04 | nan | 7.93E-05 |
| **SGFQQTMQM** | 6.26E-06 | 1.64E-05 | 3.57E-05 | nan | 2.38E-05 | 1.09E-05 | 6.57E-06 | 2.19E-05 | 3.00E-05 |
| **QTMQML** | 7.64E-06 | 3.53E-06 | 3.65E-06 | 6.23E-06 | 2.01E-06 | 1.03E-05 | 7.68E-07 | 5.02E-06 | 6.09E-05 |
| **NEGIVPTAML** | 6.94E-05 | 5.38E-04 | 8.85E-04 | 9.45E-07 | 4.17E-04 | 7.39E-04 | 7.57E-04 | 5.08E-04 | 1.96E-03 |
| **LVANDQMALGAM** | 1.21E-04 | 1.97E-03 | nan | 3.91E-03 | 3.06E-03 | 4.73E-03 | 3.93E-03 | 2.16E-03 | 5.08E-03 |
| **VANDQMALGAMR** | 2.27E-07 | 1.16E-04 | 1.26E-04 | 1.40E-04 | 1.69E-04 | 5.21E-05 | 1.80E-04 | 1.84E-04 | 1.85E-04 |
| **AITESGLRVGAD** | 6.73E-07 | 6.39E-04 | 2.26E-04 | 2.45E-04 | 4.26E-04 | 6.73E-04 | 1.94E-04 | 6.61E-04 | 5.64E-04 |
| **ITESGLRVGAD** | 4.37E-06 | 2.68E-04 | 1.09E-04 | 1.65E-04 | 1.45E-04 | 1.06E-04 | 1.46E-04 | 1.42E-04 | 1.39E-04 |
| **ADISVVGY** | 8.67E-06 | 1.16E-04 | 8.90E-05 | 2.15E-04 | 1.29E-04 | 6.02E-05 | 7.40E-05 | 5.53E-05 | 4.68E-05 |
| **VVGYDD** | 6.77E-07 | 1.30E-04 | 1.43E-04 | 1.95E-04 | 1.57E-04 | 6.87E-05 | 3.22E-04 | 1.22E-04 | 6.21E-05 |
| **DDTEDSSCYIPPLTTIK** | 7.73E-07 | 2.09E-04 | 2.58E-04 | 2.00E-04 | 5.36E-04 | 4.22E-04 | 9.00E-05 | 3.74E-04 | 6.15E-04 |
| **TEDSSCYIPPLTTIK** | 2.42E-06 | 6.47E-05 | 7.62E-05 | 6.32E-05 | 2.68E-04 | 1.14E-04 | 7.14E-05 | 1.80E-04 | 2.32E-04 |
| **SSCYIPPLTTIKQD** | 1.25E-06 | 8.45E-05 | 7.36E-05 | 1.30E-04 | 2.33E-04 | 6.82E-05 | 2.52E-05 | 9.94E-05 | 1.81E-04 |
| **FRLL** | 5.68E-07 | 6.91E-06 | 6.99E-06 | 5.27E-07 | 2.64E-06 | 4.07E-05 | 6.05E-05 | 1.02E-04 | 8.08E-05 |
| **FRLLGQTSV** | 5.82E-05 | 3.06E-05 | nan | 2.94E-06 | 2.35E-05 | 4.45E-05 | 4.11E-05 | 1.45E-05 | 6.09E-05 |
| **LGQTSVDR** | 4.55E-07 | 4.57E-05 | 2.75E-05 | 6.30E-05 | 4.99E-05 | 1.98E-05 | 2.26E-05 | 4.78E-05 | 2.36E-05 |
| **GQTSVDRL** | 3.24E-06 | 2.13E-05 | 2.13E-05 | 3.36E-05 | 7.04E-05 | 2.50E-05 | 3.05E-05 | nan | nan |
| **LLQL** | 1.70E-06 | 1.54E-04 | 1.44E-04 | 4.55E-05 | 7.69E-05 | 4.29E-04 | 2.76E-04 | 3.94E-04 | 6.98E-04 |

### IPTG vs TMG

| Peptide | 0 s | 30 s | 45 s | 60 s | 300 s | 1500 s | 3600 s | 7200 s | 14400 s |
|---|---|---|---|---|---|---|---|---|---|
| LIGVA | 6.02E-06 | 2.78E-06 | 6.96E-05 | 2.77E-05 | 7.27E-07 | 3.33E-06 | 1.42E-05 | 3.72E-06 | 2.11E-05 |
| ALHAP | 5.91E-07 | 5.24E-05 | 1.22E-05 | 2.37E-04 | 5.39E-05 | 4.22E-04 | 3.23E-04 | 5.55E-04 | 4.77E-04 |
| ALHAPSQIVA | 1.50E-06 | 2.33E-05 | nan | 6.83E-04 | 1.07E-05 | 2.20E-04 | 3.91E-05 | 3.61E-04 | 1.82E-04 |
| ALHAPSQIVAA | 1.14E-06 | 4.27E-05 | nan | 6.73E-04 | 3.01E-05 | 1.93E-04 | 1.18E-05 | 3.64E-04 | 1.82E-04 |
| ALHAPSQIVAAIK | 1.13E-05 | nan | 1.78E-04 | nan | 1.84E-04 | 3.44E-05 | 1.12E-03 | 1.62E-04 | 2.05E-06 |
| SQIVAA | 1.68E-05 | 1.68E-05 | 3.28E-05 | 1.00E-04 | nan | 2.97E-05 | 9.50E-05 | 1.81E-05 | 1.16E-05 |
| SQIVAAIK | 3.36E-06 | 1.88E-05 | 2.60E-05 | 9.53E-05 | 5.14E-05 | 2.84E-05 | 3.78E-05 | 1.17E-06 | 5.52E-05 |
| IVAAIK | 1.82E-06 | 1.19E-05 | 5.02E-06 | 4.77E-05 | 6.78E-06 | 1.50E-05 | 1.14E-05 | 9.75E-06 | 7.57E-06 |
| AIKSRADQLGAS | 1.02E-06 | 2.42E-04 | 4.25E-05 | 1.07E-04 | 8.18E-05 | 2.27E-04 | 3.99E-04 | 1.16E-03 | 6.45E-04 |
| ADQLGAS | 2.80E-05 | nan | 8.81E-05 | nan | 1.41E-04 | nan | 3.24E-03 | 1.87E-03 | 1.13E-03 |
| VVVSM | 6.12E-05 | 4.64E-04 | 1.52E-04 | 3.52E-05 | 6.88E-05 | 3.35E-04 | 2.10E-05 | 6.96E-04 | 4.69E-04 |
| MVERSGVE | 1.07E-05 | 2.92E-03 | nan | 4.51E-04 | 5.79E-04 | 1.25E-03 | 1.12E-03 | 1.44E-03 | 8.42E-04 |
| AAVHNL | 4.20E-05 | 2.59E-05 | 3.07E-04 | 6.08E-05 | 2.40E-04 | 1.33E-03 | 1.60E-03 | 1.21E-03 | 1.73E-03 |
| LAQRVSGL | 2.04E-06 | 3.94E-04 | 1.58E-04 | 4.55E-05 | 1.59E-04 | 3.69E-04 | 2.20E-04 | 6.51E-04 | 6.34E-04 |
| VSGLIIN | 7.65E-06 | 1.79E-06 | 3.89E-05 | 1.09E-05 | 2.57E-05 | 1.33E-05 | 2.60E-05 | 1.19E-05 | 1.20E-05 |
| IINYPLD | 2.19E-05 | 1.89E-05 | 6.42E-06 | nan | 3.06E-05 | 4.14E-05 | 5.21E-05 | 3.97E-05 | 5.60E-05 |
| LDDQDAIA | 2.85E-05 | 1.34E-03 | 2.37E-04 | 3.19E-04 | 5.34E-04 | 9.22E-04 | 3.20E-04 | 9.68E-04 | 9.02E-04 |
| IAVEAA | 5.55E-07 | 2.31E-05 | 1.12E-04 | 3.60E-06 | 1.15E-05 | 6.37E-05 | 5.81E-05 | 4.68E-04 | 8.45E-04 |
| ACTNVPAL | 3.15E-05 | 2.52E-03 | 3.93E-06 | 1.03E-04 | 4.33E-04 | 1.29E-03 | 4.56E-04 | 2.43E-03 | 1.30E-03 |
| ALFLD | 1.47E-05 | 1.05E-05 | 1.38E-04 | 4.53E-05 | 3.55E-05 | 1.16E-05 | 1.03E-04 | 9.84E-05 | 1.89E-04 |
| FLDVSDQTPINS | 5.82E-06 | 2.86E-04 | nan | 7.13E-04 | 7.44E-05 | 5.68E-04 | 4.87E-04 | 7.88E-04 | 3.12E-04 |
| DQTPIN | 9.47E-06 | 2.88E-04 | 2.64E-04 | 4.82E-04 | 6.70E-05 | 1.53E-03 | 4.71E-04 | 9.69E-04 | 4.52E-04 |
| SIIFSH | 3.14E-04 | 1.67E-04 | 8.16E-05 | 8.25E-04 | 1.62E-04 | 1.78E-04 | 7.08E-05 | 1.24E-04 | 1.72E-04 |
| SIIFSHEDGTRL | 4.13E-06 | 2.00E-04 | 2.08E-06 | 2.49E-04 | 5.21E-05 | 6.12E-05 | 1.43E-05 | 6.38E-05 | 5.28E-05 |
| EDGTRLGVEH | 1.86E-05 | 6.77E-05 | 3.30E-04 | nan | 2.29E-04 | 4.68E-04 | 3.70E-05 | 5.60E-05 | 4.61E-04 |
| LVAL | 7.27E-08 | 1.28E-06 | 8.76E-06 | 1.49E-04 | 9.21E-06 | 1.82E-05 | 8.26E-06 | 1.86E-06 | 2.65E-05 |
| LVALGHQQIAL | 4.55E-06 | 6.56E-04 | 1.41E-04 | 5.01E-04 | 2.88E-04 | 5.91E-04 | 2.52E-04 | 5.03E-04 | 5.16E-04 |
| LGHQQIALL | 7.23E-06 | 5.92E-05 | 7.59E-05 | 3.62E-05 | 4.58E-05 | 6.95E-05 | 3.01E-05 | 6.27E-05 | 3.49E-05 |
| IALL | 2.10E-06 | 6.45E-06 | 1.06E-06 | 6.82E-05 | 4.88E-06 | 1.63E-05 | 4.33E-06 | 3.32E-06 | 3.97E-06 |

| | | | | | | | | | |
|---|---|---|---|---|---|---|---|---|---|
| **LAGPLS** | 2.25E-05 | 7.64E-04 | 3.15E-05 | 5.03E-04 | 6.11E-05 | 8.55E-04 | 1.33E-05 | 9.46E-04 | 2.62E-04 |
| **LAGPLSSVSARL** | 2.60E-05 | 1.48E-04 | 1.03E-04 | 2.78E-04 | 5.03E-05 | 6.31E-04 | 1.70E-04 | nan | nan |
| **GPLSSVSAR** | 3.60E-06 | 6.09E-04 | 1.90E-04 | nan | 2.21E-05 | 5.40E-04 | 7.85E-04 | 8.71E-04 | nan |
| **LAGWHK** | 1.95E-05 | 1.69E-04 | 2.46E-04 | nan | 2.85E-05 | 5.41E-04 | 3.17E-05 | 2.82E-04 | 3.67E-04 |
| **YLTRNQIQP** | 6.88E-06 | 9.01E-04 | nan | 1.34E-04 | 2.55E-04 | nan | 4.77E-04 | nan | 1.76E-03 |
| **NQIQPIAEREGD** | 1.44E-06 | 1.25E-03 | 9.05E-05 | 4.85E-04 | nan | 9.93E-04 | nan | 9.72E-04 | 9.56E-04 |
| **IAEREGDWSAMSG** | 8.41E-07 | 1.50E-03 | 4.83E-04 | 7.93E-04 | 9.48E-04 | 2.01E-03 | 1.41E-03 | 1.83E-03 | 1.70E-03 |
| **REGDWSAMSGF** | 8.56E-06 | 4.02E-04 | 3.88E-04 | 3.39E-04 | nan | 3.35E-04 | nan | 8.14E-04 | 3.64E-04 |
| **DWSAM** | 1.35E-05 | 5.59E-04 | 3.01E-04 | 7.27E-04 | 3.73E-04 | 9.42E-04 | 3.27E-04 | 1.09E-03 | 5.77E-04 |
| **WSAM** | 1.64E-05 | 6.22E-04 | 4.49E-04 | 3.16E-04 | 3.50E-04 | 8.71E-04 | 3.14E-04 | nan | 5.42E-04 |
| **SAMSGFQ** | 1.75E-05 | 1.64E-03 | 8.46E-04 | 6.77E-04 | 7.08E-04 | 1.38E-04 | 3.91E-04 | nan | 9.43E-05 |
| **SGFQQTMQM** | 5.97E-06 | 1.75E-05 | 4.08E-05 | nan | 2.47E-05 | 1.32E-05 | 8.10E-06 | 1.22E-05 | 2.16E-05 |
| **QTMQML** | 1.82E-06 | 1.44E-05 | 1.59E-05 | nan | 2.10E-06 | 6.22E-06 | 1.93E-06 | 1.02E-05 | 3.73E-05 |
| **NEGIVPTAML** | 5.68E-05 | 6.78E-04 | 1.56E-03 | 8.67E-06 | 2.22E-04 | 7.47E-04 | 3.20E-04 | 1.52E-03 | 1.53E-03 |
| **LVANDQMALGAM** | 1.83E-06 | 2.10E-04 | nan | nan | 2.39E-04 | 3.50E-04 | 1.36E-04 | 1.43E-04 | 9.36E-05 |
| **VANDQMALGAMR** | 8.82E-07 | 1.82E-04 | 7.65E-05 | nan | 1.59E-04 | 2.48E-04 | 8.30E-06 | 6.29E-05 | 1.96E-04 |
| **AITESGLRVGAD** | 9.06E-06 | 1.39E-03 | 1.55E-04 | 7.46E-04 | 5.85E-04 | 1.34E-03 | 3.62E-04 | 1.14E-03 | 8.92E-04 |
| **ITESGLRVGAD** | 6.34E-06 | 1.15E-03 | 1.07E-04 | 3.70E-04 | 2.95E-04 | 7.08E-04 | 3.23E-04 | 8.48E-04 | 3.94E-04 |
| **ADISVVGY** | 9.04E-06 | 9.78E-05 | 2.77E-05 | nan | 1.17E-04 | 2.16E-04 | 5.13E-05 | 2.33E-05 | 8.14E-05 |
| **VVGYDD** | 8.23E-07 | 1.07E-04 | 1.01E-04 | nan | 1.59E-04 | 1.46E-04 | 2.91E-04 | 4.69E-05 | 1.08E-04 |
| **DDTEDSSCYIPPLTTIK** | 7.14E-07 | 3.48E-04 | 1.90E-04 | nan | 4.67E-04 | 6.82E-04 | 7.20E-05 | 9.79E-04 | 1.36E-03 |
| **TEDSSCYIPPLTTIK** | 4.82E-06 | 1.10E-04 | 1.87E-04 | nan | 2.11E-04 | 2.08E-04 | 2.48E-05 | 4.33E-04 | 5.20E-04 |
| **SSCYIPPLTTIKQD** | 1.14E-06 | 1.12E-04 | 1.85E-04 | 3.42E-03 | 2.27E-04 | 2.98E-04 | 3.20E-05 | 1.42E-04 | 3.71E-04 |
| **FRLL** | 3.36E-07 | 7.90E-06 | 5.35E-05 | 1.58E-04 | 5.69E-06 | 3.92E-05 | 3.69E-05 | 1.96E-04 | 1.25E-04 |
| **FRLLGQTSV** | 7.95E-06 | 7.38E-05 | nan | 1.02E-04 | 5.04E-05 | 5.21E-05 | nan | 3.14E-05 | 4.87E-05 |
| **LGQTSVDR** | 6.77E-07 | 4.94E-05 | 3.01E-05 | 2.10E-04 | 5.09E-05 | 7.18E-05 | 2.00E-05 | 3.02E-05 | 4.40E-05 |
| **GQTSVDRL** | 3.24E-06 | nan | nan | 1.08E-04 | nan | 8.06E-05 | 2.81E-05 | nan | nan |
| **LLQL** | 2.23E-06 | 2.05E-04 | 3.07E-04 | 9.96E-05 | 3.30E-05 | 5.52E-04 | 1.41E-04 | 1.14E-03 | 4.22E-04 |

### ONPF vs APO

| Peptide | 0 s | 30 s | 45 s | 60 s | 300 s | 1500 s | 3600 s | 7200 s | 14400 s |
|---|---|---|---|---|---|---|---|---|---|
| LIGVA | 4.79E-06 | 2.60E-05 | 2.58E-05 | 3.69E-05 | 4.98E-05 | 1.65E-05 | 3.33E-06 | 1.27E-05 | 7.54E-05 |
| ALHAP | 5.91E-07 | 2.94E-04 | 3.84E-04 | 5.43E-04 | 1.19E-03 | 5.70E-04 | 4.48E-04 | 5.65E-04 | 2.02E-03 |
| ALHAPSQIVA | 8.79E-06 | 4.36E-05 | nan | 6.12E-05 | 2.79E-05 | 3.55E-05 | 2.05E-04 | 1.14E-04 | 2.20E-05 |
| ALHAPSQIVAA | 5.35E-06 | 5.18E-05 | nan | 1.53E-04 | 8.70E-05 | 1.01E-04 | 1.90E-04 | 4.15E-04 | 1.23E-03 |
| ALHAPSQIVAAIK | nan | nan | nan | nan | nan | nan | nan | nan | nan |
| SQIVAA | 5.52E-05 | nan | nan | 1.06E-04 | nan | nan | 7.17E-04 | 1.06E-03 | nan |
| SQIVAAIK | 7.68E-07 | 7.44E-06 | nan | 1.76E-04 | 5.20E-05 | 4.97E-04 | 1.10E-03 | 4.69E-04 | nan |
| IVAAIK | 2.66E-06 | 1.64E-06 | 1.78E-04 | 1.41E-04 | 1.64E-05 | 2.33E-04 | 2.53E-04 | 1.26E-04 | 1.96E-03 |
| AIKSRADQLGAS | 3.26E-06 | 1.13E-04 | 3.51E-04 | 1.32E-05 | 1.13E-04 | 1.94E-04 | 2.71E-04 | 4.11E-04 | 5.02E-04 |
| ADQLGAS | 2.43E-04 | nan | 5.16E-04 | 4.02E-05 | nan | 9.58E-04 | 8.97E-04 | nan | nan |
| VVVSM | 1.08E-05 | 5.01E-04 | 1.19E-03 | 1.45E-04 | 1.81E-03 | 7.05E-04 | 5.94E-04 | 9.60E-04 | 2.12E-03 |
| MVERSGVE | 5.56E-06 | 7.94E-04 | nan | 1.16E-03 | 6.62E-04 | 5.53E-04 | 6.82E-04 | 5.75E-04 | 7.04E-04 |
| AAVHNL | 2.63E-05 | 1.57E-05 | 1.29E-04 | 6.39E-06 | 5.20E-05 | 1.48E-04 | 8.49E-05 | 1.99E-03 | nan |
| LAQRVSGL | 5.32E-07 | 3.75E-05 | 6.49E-04 | 5.48E-06 | 2.50E-04 | 1.17E-04 | 3.64E-04 | 3.71E-04 | 4.23E-03 |
| VSGLIIN | 6.83E-05 | 1.89E-05 | 4.73E-05 | 9.92E-05 | 2.57E-05 | 1.03E-05 | 1.10E-04 | 9.79E-05 | 1.28E-05 |
| IINYPLD | 3.35E-06 | 2.71E-06 | 2.75E-06 | 3.07E-06 | 1.66E-05 | 2.27E-07 | 2.66E-05 | 8.06E-05 | 4.84E-04 |
| LDDQDAIA | 1.82E-07 | 7.94E-04 | 1.82E-03 | 5.38E-04 | nan | 4.30E-04 | 1.04E-03 | 1.05E-03 | nan |
| IAVEAA | 1.53E-06 | 4.28E-05 | 5.33E-05 | 2.01E-04 | 6.29E-05 | 9.52E-05 | 4.69E-04 | 4.29E-04 | 4.16E-03 |
| ACTNVPAL | 1.25E-05 | 7.17E-04 | 1.92E-03 | 2.34E-04 | 2.31E-03 | 7.46E-04 | 8.70E-04 | 1.54E-03 | nan |
| ALFLD | 1.18E-06 | 2.93E-05 | 2.30E-05 | 4.73E-06 | 3.19E-06 | 5.96E-06 | 3.08E-06 | 2.22E-04 | nan |
| FLDVSDQTPINS | 1.53E-05 | 1.44E-04 | nan | 2.67E-04 | 1.58E-04 | 3.89E-04 | 3.74E-04 | 6.52E-04 | 1.27E-03 |
| DQTPIN | 2.09E-05 | 2.16E-05 | 2.89E-04 | 5.27E-05 | 1.17E-03 | 6.02E-04 | 5.28E-04 | 7.20E-04 | nan |
| SIIFSH | nan | nan | nan | nan | nan | nan | nan | nan | nan |
| SIIFSHEDGTRL | 2.65E-06 | 1.77E-05 | nan | 6.94E-06 | 2.91E-05 | 2.33E-05 | 4.92E-05 | 6.30E-05 | 1.90E-04 |
| EDGTRLGVEH | 3.39E-06 | 2.88E-05 | 5.55E-05 | 1.55E-07 | 2.91E-05 | 3.60E-05 | 1.44E-04 | 1.46E-04 | nan |
| LVAL | 6.64E-07 | 1.82E-08 | 5.02E-06 | 4.25E-06 | 6.77E-07 | 1.43E-05 | 2.13E-06 | 7.86E-07 | 3.76E-05 |
| LVALGHQQIAL | 1.35E-06 | 2.82E-04 | 5.12E-04 | 8.99E-05 | 2.96E-04 | 1.14E-04 | 2.69E-04 | 6.03E-04 | 1.70E-03 |
| LGHQQIALL | 1.41E-05 | 1.54E-05 | nan | 6.52E-07 | 3.43E-05 | 3.51E-05 | 5.64E-05 | 6.31E-05 | 2.88E-05 |
| IALL | 4.48E-06 | 1.08E-05 | 9.43E-06 | 2.37E-06 | 8.01E-06 | 8.63E-06 | 4.33E-06 | 1.15E-05 | nan |

| | | | | | | | | | |
|---|---|---|---|---|---|---|---|---|---|
| **LAGPLS** | 2.08E-05 | 2.03E-04 | 2.72E-04 | 6.55E-05 | 5.52E-04 | 8.80E-04 | 1.41E-03 | nan | nan |
| **LAGPLSSVSARL** | 2.94E-05 | 1.56E-04 | 7.30E-04 | 1.03E-04 | 4.90E-04 | 6.06E-04 | 1.27E-03 | 7.00E-04 | 2.88E-03 |
| **GPLSSVSAR** | 5.13E-06 | 4.99E-04 | 1.73E-03 | 1.32E-04 | nan | 8.46E-06 | 2.89E-03 | 3.19E-03 | 1.57E-03 |
| **LAGWHK** | 2.14E-05 | 1.96E-05 | 6.11E-06 | 6.13E-06 | 2.00E-07 | 3.84E-06 | 3.40E-06 | 4.39E-05 | 6.09E-05 |
| **YLTRNQIQP** | 2.64E-06 | 6.96E-05 | 1.24E-03 | 6.69E-05 | 7.90E-04 | 1.01E-03 | 4.18E-04 | 9.71E-04 | 1.44E-03 |
| **NQIQPIAEREGD** | 8.77E-07 | 4.09E-04 | 1.18E-03 | 1.71E-04 | 1.71E-03 | 6.51E-04 | 6.29E-04 | 1.01E-03 | nan |
| **IAEREGDWSAMSG** | 5.51E-06 | 4.04E-04 | 1.51E-03 | 2.78E-05 | 2.10E-03 | 1.72E-03 | 1.53E-03 | 2.11E-03 | 9.73E-03 |
| **REGDWSAMSGF** | 4.29E-06 | 3.54E-05 | nan | 1.34E-04 | 1.62E-04 | 1.28E-04 | 5.79E-04 | 2.40E-04 | 2.60E-04 |
| **DWSAM** | 7.80E-06 | 3.80E-05 | 2.90E-04 | 8.98E-05 | 1.23E-03 | 8.48E-04 | 1.07E-03 | 1.25E-03 | nan |
| **WSAM** | 8.69E-05 | 1.84E-04 | 9.42E-04 | 4.13E-06 | 1.44E-03 | 1.16E-03 | 1.02E-03 | nan | 1.27E-03 |
| **SAMSGFQ** | 1.68E-05 | 3.39E-04 | nan | 3.87E-04 | 2.34E-03 | 2.67E-04 | 1.23E-03 | 8.62E-04 | nan |
| **SGFQQTMQM** | 3.76E-06 | 4.06E-06 | 2.49E-05 | 6.04E-05 | 1.66E-06 | 3.80E-06 | 6.05E-06 | 1.54E-05 | 3.88E-05 |
| **QTMQML** | 4.98E-06 | 8.67E-06 | 5.73E-06 | 1.38E-05 | 2.73E-06 | 2.71E-06 | 3.72E-06 | 1.95E-05 | 8.16E-05 |
| **NEGIVPTAML** | 1.09E-05 | 1.58E-04 | 2.44E-03 | 3.13E-04 | 7.86E-04 | 5.03E-04 | 2.07E-03 | 1.45E-03 | 5.15E-03 |
| **LVANDQMALGAM** | 2.43E-05 | 1.49E-04 | nan | 3.93E-04 | 2.58E-04 | 5.24E-04 | 3.20E-04 | 3.05E-04 | 2.91E-04 |
| **VANDQMALGAMR** | 7.36E-07 | 4.82E-05 | 1.05E-04 | 1.06E-05 | 7.81E-05 | 5.67E-05 | 1.72E-04 | 1.90E-04 | 1.36E-03 |
| **AITESGLRVGAD** | 1.32E-06 | 4.37E-04 | 5.86E-04 | 1.63E-04 | 7.45E-04 | 2.03E-04 | 3.80E-04 | 1.05E-03 | 2.72E-03 |
| **ITESGLRVGAD** | 8.34E-06 | 1.91E-04 | 4.21E-04 | 1.27E-04 | 2.34E-04 | 1.27E-04 | 2.15E-04 | 2.43E-04 | 4.57E-04 |
| **ADISVVGY** | 1.18E-06 | 2.45E-05 | 7.90E-06 | 4.99E-06 | 4.34E-06 | 1.68E-04 | 1.93E-05 | 1.19E-05 | 5.53E-04 |
| **VVGYDD** | 2.86E-06 | 3.75E-05 | 7.83E-05 | 3.97E-05 | 7.54E-05 | 1.27E-04 | 3.41E-04 | 2.23E-04 | 1.22E-03 |
| **DDTEDSSCYIPPLTTIK** | 1.93E-06 | 1.14E-04 | 7.17E-04 | 7.02E-05 | 3.68E-04 | 4.92E-04 | 9.25E-04 | 9.60E-04 | nan |
| **TEDSSCYIPPLTTIK** | 1.20E-06 | 3.77E-06 | 4.17E-04 | 1.79E-04 | 3.46E-05 | 3.61E-04 | 1.84E-04 | 2.65E-04 | nan |
| **SSCYIPPLTTIKQD** | 4.09E-07 | 6.50E-05 | 4.51E-04 | 6.44E-05 | 7.18E-05 | 3.18E-04 | 4.40E-04 | 2.98E-04 | 7.57E-03 |
| **FRLL** | 6.77E-07 | 1.48E-06 | 7.23E-05 | 2.37E-05 | 2.73E-06 | 7.48E-05 | 1.48E-04 | 6.96E-05 | 1.76E-03 |
| **FRLLGQTSV** | 2.20E-05 | 1.59E-05 | nan | 7.51E-05 | 1.96E-05 | 4.31E-05 | 3.26E-05 | 4.21E-05 | 3.26E-05 |
| **LGQTSVDR** | 3.64E-08 | 2.82E-05 | 5.30E-05 | 2.42E-05 | 3.29E-05 | 1.71E-05 | 6.02E-05 | 1.55E-04 | 7.04E-04 |
| **GQTSVDRL** | 5.68E-07 | 1.48E-05 | 2.30E-05 | 1.20E-05 | 1.89E-05 | 3.53E-05 | 1.45E-05 | 6.34E-05 | nan |
| **LLQL** | 9.09E-07 | 3.52E-05 | 4.83E-04 | 1.59E-04 | 3.33E-04 | 5.29E-04 | 9.07E-04 | 7.65E-04 | 4.71E-03 |

### ONPF vs DNA

| Peptide | 0 s | 30 s | 45 s | 60 s | 300 s | 1500 s | 3600 s | 7200 s | 14400 s |
|---|---|---|---|---|---|---|---|---|---|
| LIGVA | 1.48E-06 | 1.63E-05 | 3.89E-05 | 2.37E-05 | 4.18E-05 | 1.53E-05 | 1.82E-07 | 5.55E-06 | 4.28E-05 |
| ALHAP | 2.02E-06 | 9.55E-05 | 5.57E-04 | 9.96E-05 | 3.71E-04 | 2.93E-04 | 2.89E-04 | 6.29E-04 | nan |
| ALHAPSQIVA | 6.61E-06 | 2.92E-05 | nan | 1.06E-04 | 4.94E-05 | 3.19E-05 | 2.25E-04 | 6.70E-05 | 5.81E-05 |
| ALHAPSQIVAA | 3.24E-06 | 6.18E-05 | nan | 2.21E-04 | 8.11E-05 | 1.15E-04 | 2.51E-04 | 3.10E-04 | 4.32E-04 |
| ALHAPSQIVAAIK | nan | nan | nan | nan | nan | nan | nan | nan | nan |
| SQIVAA | 5.72E-05 | nan | nan | 5.95E-05 | nan | nan | 7.17E-04 | 3.59E-04 | nan |
| SQIVAAIK | 3.68E-06 | 1.78E-05 | nan | 2.08E-06 | 7.33E-05 | 4.29E-05 | 1.19E-03 | 2.76E-04 | nan |
| IVAAIK | 3.35E-06 | 5.56E-06 | 2.28E-04 | 8.77E-07 | 1.89E-05 | 1.56E-05 | 2.54E-04 | 5.33E-05 | 1.80E-03 |
| AIKSRADQLGAS | 1.26E-06 | 9.27E-05 | 4.71E-04 | 1.58E-04 | 1.59E-04 | 1.88E-04 | 2.89E-04 | 3.19E-04 | 1.25E-03 |
| ADQLGAS | 2.55E-04 | nan | 6.59E-04 | 3.31E-04 | nan | 1.04E-03 | 8.07E-04 | nan | nan |
| VVVSM | 1.10E-05 | 1.46E-04 | 1.62E-03 | 2.67E-04 | 9.31E-04 | 5.25E-04 | 4.89E-04 | 8.56E-04 | 2.18E-03 |
| MVERSGVE | 4.93E-06 | 3.36E-04 | nan | 4.95E-04 | 9.76E-04 | 5.34E-04 | 7.90E-04 | 7.65E-04 | 3.82E-04 |
| AAVHNL | 1.08E-05 | 3.34E-05 | 2.07E-04 | 6.68E-06 | 2.05E-04 | 1.44E-04 | 4.69E-04 | 2.19E-04 | nan |
| LAQRVSGL | 3.86E-07 | 2.17E-05 | 6.56E-04 | 7.26E-06 | 1.52E-04 | 1.16E-04 | 3.63E-04 | 1.29E-04 | 4.75E-03 |
| VSGLIIN | 3.28E-05 | 6.18E-06 | 1.64E-05 | 1.61E-05 | 7.14E-07 | 2.02E-06 | 2.91E-05 | 6.93E-05 | 8.74E-06 |
| IINYPLD | 8.57E-06 | 7.39E-06 | 1.40E-05 | 5.48E-06 | 3.40E-05 | 1.49E-05 | 2.00E-05 | 5.69E-05 | 8.77E-04 |
| LDDQDAIA | 2.05E-07 | 3.65E-04 | 1.63E-03 | 5.27E-04 | 8.31E-04 | 5.31E-04 | 1.34E-03 | 1.15E-03 | nan |
| IAVEAA | 1.39E-06 | nan | 9.41E-04 | 1.78E-06 | 7.73E-05 | 1.60E-04 | 3.56E-04 | 6.87E-05 | 4.08E-03 |
| ACTNVPAL | 1.77E-05 | 4.28E-05 | 1.42E-03 | 1.85E-05 | 8.17E-04 | 5.74E-04 | 1.23E-03 | 1.30E-03 | nan |
| ALFLD | 1.18E-06 | 6.07E-05 | 1.73E-04 | 1.94E-05 | 4.85E-05 | 3.72E-05 | 6.21E-05 | 2.05E-04 | nan |
| FLDVSDQTPINS | 7.02E-06 | 9.79E-05 | nan | 3.29E-04 | 2.27E-04 | 4.28E-04 | 5.24E-04 | 8.81E-04 | 1.47E-03 |
| DQTPIN | 2.54E-05 | 2.08E-05 | 7.36E-04 | 9.68E-06 | 5.82E-04 | 6.13E-04 | 5.96E-04 | 5.56E-04 | nan |
| SIIFSH | nan | nan | nan | nan | nan | nan | nan | nan | nan |
| SIIFSHEDGTRL | 1.03E-06 | 1.79E-05 | nan | 2.47E-05 | 1.94E-05 | 2.11E-05 | 4.92E-05 | 6.40E-05 | 1.93E-04 |
| EDGTRLGVEH | 7.79E-06 | 2.14E-05 | 5.35E-05 | 3.60E-05 | 3.08E-05 | 3.25E-05 | 2.42E-04 | 8.73E-05 | nan |
| LVAL | 5.55E-07 | 1.82E-06 | 2.02E-05 | 2.36E-07 | 1.39E-06 | 2.27E-08 | 8.67E-06 | 3.84E-06 | 9.17E-05 |
| LVALGHQQIAL | 4.02E-06 | 8.22E-05 | 3.91E-04 | 3.59E-05 | 1.55E-04 | 8.91E-05 | 3.54E-04 | 5.60E-04 | 1.97E-03 |
| LGHQQIALL | 1.03E-05 | 1.90E-05 | 2.27E-05 | 1.32E-05 | 3.96E-05 | 4.32E-05 | 5.66E-05 | 6.54E-05 | 4.37E-05 |
| IALL | 4.24E-06 | 5.04E-06 | nan | 8.23E-06 | 4.46E-06 | 1.04E-05 | 5.04E-06 | 5.30E-06 | nan |

| | | | | | | | | | |
|---|---|---|---|---|---|---|---|---|---|
| **LAGPLS** | 2.54E-05 | 1.82E-05 | 2.77E-04 | 3.60E-06 | 1.82E-04 | nan | 1.62E-03 | 9.28E-04 | nan |
| **LAGPLSSVSARL** | 3.18E-04 | 4.25E-04 | 9.85E-04 | 1.10E-04 | 4.30E-04 | 7.41E-04 | 8.63E-04 | 8.34E-04 | 3.60E-03 |
| **GPLSSVSAR** | 2.84E-06 | 1.53E-04 | 1.68E-03 | 4.11E-05 | 2.04E-04 | 1.03E-03 | 5.54E-04 | 1.82E-03 | 4.04E-03 |
| **LAGWHK** | 1.10E-04 | 4.36E-05 | 8.24E-05 | 1.43E-06 | 5.06E-05 | 1.95E-05 | 1.54E-05 | 3.24E-05 | 0 s.00E+00 |
| **YLTRNQIQP** | 1.62E-06 | 3.30E-06 | 1.46E-03 | 1.55E-06 | 4.02E-04 | 4.50E-05 | 4.43E-04 | 3.47E-04 | nan |
| **NQIQPIAEREGD** | 3.30E-06 | 4.58E-05 | 8.14E-04 | 9.28E-06 | 1.05E-03 | 3.57E-04 | 7.58E-04 | 7.89E-04 | nan |
| **IAEREGDWSAMSG** | 3.97E-06 | 2.38E-04 | 1.68E-03 | 2.75E-05 | 9.44E-04 | 8.64E-04 | 1.56E-03 | 1.05E-03 | 1.16E-02 |
| **REGDWSAMSGF** | 5.31E-06 | 1.66E-05 | nan | 1.83E-04 | 1.64E-04 | 5.03E-04 | 6.55E-04 | 3.84E-04 | 9.94E-04 |
| **DWSAM** | 1.19E-05 | 3.35E-05 | 5.87E-04 | 8.23E-05 | 7.34E-04 | 6.21E-04 | 9.00E-04 | 5.75E-04 | nan |
| **WSAM** | 1.50E-04 | 1.08E-03 | 1.44E-03 | 1.19E-05 | 1.04E-03 | nan | 1.41E-03 | nan | nan |
| **SAMSGFQ** | 1.24E-05 | 3.08E-04 | nan | nan | 1.86E-03 | 2.34E-04 | 2.14E-04 | 3.28E-04 | nan |
| **SGFQQTMQM** | 4.35E-06 | 9.86E-06 | 1.50E-05 | 1.58E-05 | 1.27E-05 | 1.01E-05 | 1.30E-05 | 2.29E-05 | 6.52E-05 |
| **QTMQML** | 1.55E-05 | 8.57E-06 | nan | 7.68E-06 | 3.36E-07 | 1.78E-06 | 2.08E-06 | 2.02E-06 | 1.48E-05 |
| **NEGIVPTAML** | 9.91E-06 | 1.48E-04 | 5.70E-03 | 8.57E-06 | 7.94E-04 | 5.01E-04 | 3.76E-03 | 1.12E-03 | 6.57E-03 |
| **LVANDQMALGAM** | 1.76E-06 | 8.53E-05 | nan | 3.56E-04 | 1.62E-04 | 4.08E-04 | 2.02E-04 | 2.03E-04 | 1.79E-04 |
| **VANDQMALGAMR** | 4.09E-07 | 4.66E-05 | 3.13E-04 | 1.74E-05 | 1.24E-04 | 7.18E-05 | 4.23E-04 | 2.11E-04 | 1.17E-03 |
| **AITESGLRVGAD** | 6.23E-06 | 1.31E-04 | 8.88E-04 | 6.46E-05 | 3.61E-04 | 2.16E-04 | 5.37E-04 | 9.80E-04 | 3.20E-03 |
| **ITESGLRVGAD** | 5.90E-06 | 1.59E-04 | 3.80E-04 | 1.44E-04 | 2.88E-04 | 1.99E-04 | 3.10E-04 | 2.96E-04 | 7.49E-04 |
| **ADISVVGY** | 6.15E-06 | 9.96E-05 | 1.14E-04 | 1.64E-04 | 7.78E-05 | 2.03E-04 | 7.62E-05 | 1.92E-05 | 2.09E-04 |
| **VVGYDD** | 2.46E-06 | 9.04E-05 | 8.30E-05 | 9.14E-05 | 1.38E-04 | 1.33E-04 | 4.18E-04 | 2.07E-04 | 1.14E-03 |
| **DDTEDSSCYIPPLTTIK** | 1.80E-06 | 5.95E-05 | 6.44E-04 | 5.22E-05 | 2.67E-04 | 3.24E-04 | 8.54E-04 | 6.26E-04 | nan |
| **TEDSSCYIPPLTTIK** | 2.64E-07 | 1.79E-05 | 4.37E-04 | 5.10E-05 | 5.82E-05 | 7.92E-05 | 2.28E-04 | 6.63E-05 | nan |
| **SSCYIPPLTTIKQD** | 2.01E-06 | 5.13E-05 | 4.78E-04 | 4.36E-05 | 7.24E-05 | 9.67E-05 | 5.69E-04 | 2.54E-04 | 7.54E-03 |
| **FRLL** | 4.95E-07 | 3.29E-06 | 1.27E-04 | 5.13E-06 | 5.93E-06 | 6.26E-06 | 1.96E-04 | 6.37E-05 | 1.98E-03 |
| **FRLLGQTSV** | 1.74E-06 | 5.61E-06 | nan | 4.87E-05 | 1.32E-05 | 8.77E-06 | 6.05E-05 | 2.65E-05 | 1.15E-04 |
| **LGQTSVDR** | 1.32E-07 | 2.82E-05 | 3.86E-05 | 3.76E-05 | 3.29E-05 | 2.14E-05 | 6.70E-05 | 1.30E-04 | 5.84E-04 |
| **GQTSVDRL** | 4.55E-07 | 9.94E-06 | 1.84E-05 | 9.45E-06 | 2.72E-05 | 1.46E-05 | 4.71E-05 | 3.62E-05 | nan |
| **LLQL** | 4.02E-06 | 3.54E-05 | 1.22E-03 | 3.46E-05 | 2.83E-04 | 4.75E-04 | 1.05E-03 | 5.49E-04 | 6.99E-03 |

### ONPF vs ONPFDNA

| Peptide | 0 s | 30 s | 45 s | 60 s | 300 s | 1500 s | 3600 s | 7200 s | 14400 s |
|---|---|---|---|---|---|---|---|---|---|
| LIGVA | 2.64E-06 | 2.60E-05 | 2.62E-05 | 2.10E-05 | 3.12E-05 | 1.89E-05 | 4.73E-07 | 5.82E-06 | 6.07E-05 |
| ALHAP | 8.41E-07 | 7.91E-05 | 3.18E-04 | 8.06E-05 | 3.80E-04 | 2.91E-04 | 1.22E-04 | 2.34E-04 | 1.90E-03 |
| ALHAPSQIVA | 8.86E-06 | 1.28E-04 | nan | 1.54E-04 | 5.41E-05 | 2.29E-04 | 2.51E-04 | 1.10E-04 | 3.45E-04 |
| ALHAPSQIVAA | 5.45E-07 | 1.17E-04 | nan | 2.31E-04 | 6.33E-05 | 2.15E-04 | 2.83E-04 | 2.63E-04 | 2.48E-04 |
| ALHAPSQIVAAIK | nan | nan | nan | nan | nan | nan | nan | nan | nan |
| SQIVAA | 7.92E-05 | nan | nan | 4.50E-05 | nan | nan | 7.25E-04 | 3.00E-04 | nan |
| SQIVAAIK | 1.86E-07 | 8.33E-06 | nan | 2.27E-06 | 5.17E-05 | 4.69E-05 | 1.27E-03 | 3.39E-04 | nan |
| IVAAIK | 1.20E-06 | 2.24E-06 | 1.83E-04 | 1.55E-06 | 1.50E-05 | 1.63E-05 | 2.44E-04 | 8.65E-05 | 1.63E-03 |
| AIKSRADQLGAS | 1.18E-06 | 1.22E-04 | 2.58E-04 | 1.49E-05 | 7.82E-05 | 3.28E-04 | 1.98E-04 | 1.97E-04 | 6.27E-04 |
| ADQLGAS | 2.73E-04 | nan | 4.15E-06 | 7.38E-05 | nan | 9.21E-04 | 7.37E-04 | nan | nan |
| VVVSM | 1.09E-05 | 1.70E-04 | 1.32E-03 | 2.05E-04 | 9.89E-04 | 5.92E-04 | 3.43E-04 | 4.30E-04 | 1.90E-03 |
| MVERSGVE | 5.35E-06 | 8.84E-04 | nan | 3.53E-04 | 4.64E-04 | 4.05E-04 | 3.05E-04 | 2.60E-04 | 4.67E-04 |
| AAVHNL | 3.72E-05 | 1.48E-05 | 5.84E-05 | 2.85E-05 | 1.70E-04 | 1.46E-04 | 1.35E-04 | nan | nan |
| LAQRVSGL | 5.32E-07 | 1.13E-05 | 5.34E-04 | 5.42E-06 | 1.49E-04 | 1.27E-04 | 3.22E-04 | 7.66E-05 | 4.11E-03 |
| VSGLIIN | 4.14E-05 | 1.16E-05 | 1.28E-05 | 9.25E-06 | 1.05E-04 | 2.79E-06 | 9.91E-07 | 3.55E-05 | 7.44E-06 |
| IINYPLD | 6.39E-06 | 1.98E-06 | 7.77E-06 | 1.73E-05 | 1.76E-05 | 3.45E-05 | 1.44E-05 | 1.08E-05 | 3.16E-04 |
| LDDQDAIA | 8.18E-07 | 3.18E-04 | 1.30E-03 | 1.57E-04 | 7.76E-04 | 5.47E-04 | 7.55E-04 | 4.38E-04 | nan |
| IAVEAA | 1.33E-06 | 4.22E-05 | 1.06E-04 | 1.27E-05 | 4.57E-05 | 2.19E-04 | 3.40E-04 | 5.82E-05 | 3.99E-03 |
| ACTNVPAL | 8.77E-07 | 8.91E-05 | 1.08E-03 | 3.39E-05 | 7.90E-04 | 5.80E-04 | 7.47E-04 | 1.99E-04 | nan |
| ALFLD | 2.64E-06 | 4.79E-05 | nan | 6.47E-05 | 4.37E-04 | 2.82E-05 | 1.08E-04 | 2.96E-04 | nan |
| FLDVSDQTPINS | 7.38E-06 | 2.09E-04 | nan | 1.60E-04 | 2.14E-04 | 2.90E-04 | 5.57E-04 | 5.52E-04 | 1.42E-03 |
| DQTPIN | 2.70E-05 | 3.97E-05 | 3.34E-04 | 1.38E-05 | 6.82E-04 | 6.24E-04 | 3.15E-04 | 2.37E-04 | nan |
| SIIFSH | nan | nan | nan | nan | nan | nan | nan | nan | nan |
| SIIFSHEDGTRL | 9.40E-07 | 1.68E-05 | nan | 1.68E-05 | 3.35E-05 | 1.64E-05 | 5.33E-05 | 7.86E-05 | 1.22E-04 |
| EDGTRLGVEH | 3.73E-06 | 3.00E-05 | 5.13E-05 | 6.95E-06 | 2.83E-05 | 4.53E-05 | 2.79E-04 | 8.30E-05 | nan |
| LVAL | 5.91E-07 | 2.91E-07 | 5.50E-06 | 2.77E-07 | 2.95E-07 | 3.33E-06 | 3.73E-06 | 1.14E-06 | 3.82E-06 |
| LVALGHQQIAL | 4.95E-07 | 7.78E-05 | 3.36E-04 | 2.61E-05 | 1.67E-04 | 7.68E-05 | 2.14E-04 | 4.76E-04 | 1.02E-03 |
| LGHQQIALL | 1.11E-05 | 4.01E-05 | 7.85E-05 | 5.56E-05 | 3.55E-05 | 2.67E-05 | 4.60E-05 | 4.71E-05 | 1.55E-05 |
| IALL | 2.64E-06 | 3.85E-06 | 1.68E-06 | 8.57E-06 | 1.02E-06 | 2.48E-06 | 3.68E-06 | 5.62E-06 | nan |

| | | | | | | | | | |
|---|---|---|---|---|---|---|---|---|---|
| **LAGPLS** | 2.22E-05 | 6.41E-05 | 1.08E-04 | 8.45E-06 | 1.75E-04 | 7.66E-04 | 1.29E-03 | nan | nan |
| **LAGPLSSVSARL** | 1.60E-05 | 1.92E-04 | 7.81E-04 | 8.12E-05 | 5.25E-04 | 5.95E-04 | 7.28E-04 | 9.95E-04 | 3.14E-03 |
| **GPLSSVSAR** | 1.89E-06 | 1.96E-04 | 1.89E-03 | 3.76E-05 | 2.02E-04 | 1.50E-05 | 4.00E-04 | 1.29E-03 | 1.28E-03 |
| **LAGWHK** | 2.50E-05 | 2.68E-05 | 6.66E-06 | 5.69E-06 | 2.77E-05 | 8.35E-06 | nan | 1.59E-05 | 8.73E-05 |
| **YLTRNQIQP** | 1.18E-06 | 4.85E-05 | 1.24E-03 | 4.97E-05 | 3.95E-04 | 1.67E-04 | 1.85E-04 | 5.02E-06 | nan |
| **NQIQPIAEREGD** | 2.95E-07 | 3.47E-05 | 7.37E-04 | 1.68E-06 | 1.07E-03 | 4.52E-04 | 4.12E-04 | 5.50E-04 | nan |
| **IAEREGDWSAMSG** | 4.08E-06 | 2.88E-04 | 1.28E-03 | 2.85E-05 | 1.11E-03 | 9.35E-04 | 7.53E-04 | 3.42E-04 | 7.43E-03 |
| **REGDWSAMSGF** | 2.57E-05 | 8.16E-05 | nan | 2.19E-04 | 1.50E-04 | 1.22E-04 | 5.63E-04 | 1.04E-04 | 1.94E-04 |
| **DWSAM** | 7.37E-06 | 2.69E-05 | 2.49E-04 | 3.68E-05 | 8.16E-04 | 8.08E-04 | 5.94E-04 | 2.08E-04 | nan |
| **WSAM** | 8.40E-05 | 1.59E-04 | 7.55E-04 | 9.04E-05 | 9.66E-04 | 8.35E-04 | 5.64E-04 | nan | nan |
| **SAMSGFQ** | 6.58E-05 | 2.78E-04 | nan | 2.81E-04 | 2.13E-03 | 2.56E-04 | 2.46E-05 | 2.47E-04 | nan |
| **SGFQQTMQM** | 6.89E-06 | 2.59E-06 | 3.01E-05 | nan | 3.09E-06 | 1.25E-05 | 6.99E-06 | 2.59E-05 | 1.66E-05 |
| **QTMQML** | 1.17E-05 | 8.47E-06 | 6.10E-06 | 5.72E-06 | 4.82E-07 | 1.05E-05 | 1.97E-06 | 2.37E-06 | 2.49E-05 |
| **NEGIVPTAML** | 3.30E-05 | 2.51E-04 | 2.94E-03 | 3.60E-06 | 7.36E-04 | 7.27E-04 | 2.04E-03 | 2.67E-04 | 1.06E-03 |
| **LVANDQMALGAM** | 1.20E-04 | 1.90E-03 | nan | 3.95E-03 | 2.96E-03 | 4.89E-03 | 3.97E-03 | 2.17E-03 | 5.08E-03 |
| **VANDQMALGAMR** | 4.82E-07 | 3.24E-05 | 1.55E-04 | 2.47E-05 | 8.74E-05 | 5.70E-05 | 3.30E-04 | 2.31E-04 | 6.00E-04 |
| **AITESGLRVGAD** | 2.27E-08 | 1.08E-04 | 6.61E-04 | 2.40E-05 | 3.68E-04 | 1.87E-04 | 2.79E-04 | 5.38E-04 | 1.84E-03 |
| **ITESGLRVGAD** | 9.46E-06 | 2.09E-04 | 3.06E-04 | 9.12E-05 | 1.45E-04 | 3.95E-05 | 1.59E-04 | 1.07E-04 | 4.21E-04 |
| **ADISVVGY** | 1.55E-06 | 5.60E-05 | 7.13E-05 | 9.60E-06 | 1.56E-05 | 1.85E-04 | 4.69E-05 | 4.05E-05 | 1.68E-04 |
| **VVGYDD** | 6.77E-07 | 5.76E-05 | 1.01E-04 | 5.08E-05 | 7.44E-05 | 1.10E-04 | 3.36E-04 | 2.15E-04 | 9.46E-04 |
| **DDTEDSSCYIPPLTTIK** | 1.86E-06 | 5.21E-05 | 7.24E-04 | 6.21E-05 | 3.24E-04 | 3.21E-04 | 8.25E-04 | 5.15E-04 | nan |
| **TEDSSCYIPPLTTIK** | 2.64E-07 | 8.10E-06 | 4.26E-04 | 5.05E-05 | 8.57E-05 | 8.02E-05 | 2.23E-04 | 2.72E-05 | nan |
| **SSCYIPPLTTIKQD** | 5.91E-07 | 6.88E-05 | 4.41E-04 | 4.52E-05 | 6.30E-05 | 9.92E-05 | 4.30E-04 | 2.16E-04 | 7.39E-03 |
| **FRLL** | 9.09E-07 | 2.50E-06 | 7.03E-05 | 3.39E-06 | 2.64E-06 | 1.87E-05 | 1.77E-04 | 4.98E-05 | 1.61E-03 |
| **FRLLGQTSV** | 5.61E-05 | 1.31E-05 | nan | 4.88E-05 | 5.22E-06 | 5.23E-05 | 2.43E-05 | 2.00E-05 | 4.36E-05 |
| **LGQTSVDR** | 1.82E-08 | 1.65E-05 | 3.46E-05 | 1.05E-05 | 1.69E-05 | 1.19E-05 | 4.56E-05 | 1.33E-04 | 5.07E-04 |
| **GQTSVDRL** | 6.18E-07 | 8.19E-06 | 1.01E-05 | 7.39E-06 | 9.66E-06 | 9.11E-06 | 1.78E-05 | 2.96E-05 | nan |
| **LLQL** | 1.93E-06 | 3.99E-05 | 5.79E-04 | 3.90E-05 | 3.29E-04 | 4.75E-04 | 1.11E-03 | 5.26E-04 | 4.98E-03 |

### ONPF vs TMG

| Peptide | 0 s | 30 s | 45 s | 60 s | 300 s | 1500 s | 3600 s | 7200 s | 14400 s |
|---|---|---|---|---|---|---|---|---|---|
| LIGVA | 1.93E-06 | 3.24E-06 | 2.66E-05 | 1.28E-05 | 1.76E-05 | 1.43E-05 | 1.38E-05 | 5.97E-06 | 3.97E-05 |
| ALHAP | 9.18E-07 | 1.10E-04 | 3.22E-04 | 2.59E-04 | 3.17E-04 | 4.29E-04 | 3.45E-04 | 4.51E-04 | 2.09E-03 |
| ALHAPSQIVA | 7.11E-06 | 4.28E-05 | nan | 7.14E-04 | 1.45E-05 | 2.26E-04 | 1.59E-04 | 3.89E-04 | 1.81E-04 |
| ALHAPSQIVAA | 1.37E-06 | 5.85E-05 | nan | 7.56E-04 | 2.60E-05 | 2.51E-04 | 9.92E-05 | 4.90E-04 | 1.71E-04 |
| ALHAPSQIVAAIK | nan | nan | nan | nan | nan | nan | nan | nan | nan |
| SQIVAA | 6.83E-05 | nan | nan | 1.21E-04 | nan | nan | 8.07E-04 | 3.16E-04 | nan |
| SQIVAAIK | 1.28E-06 | 8.64E-06 | nan | 8.92E-05 | 5.14E-05 | 7.04E-05 | 1.12E-03 | 2.19E-04 | nan |
| IVAAIK | 1.86E-06 | 2.95E-06 | 1.83E-04 | 4.34E-05 | 1.57E-05 | 3.01E-05 | 2.38E-04 | 5.56E-05 | 1.63E-03 |
| AIKSRADQLGAS | 1.88E-06 | 2.49E-04 | 2.43E-04 | 2.27E-05 | 5.81E-05 | 2.70E-04 | 3.87E-04 | 1.09E-03 | 6.36E-04 |
| ADQLGAS | 2.56E-04 | nan | 2.98E-05 | 3.17E-04 | nan | 1.52E-03 | 2.02E-03 | nan | nan |
| VVVSM | 7.16E-05 | 3.70E-04 | 1.15E-03 | 1.26E-04 | 7.87E-04 | 4.82E-04 | 3.17E-04 | 7.22E-04 | 1.63E-03 |
| MVERSGVE | 1.54E-05 | 2.41E-03 | nan | 1.93E-04 | 4.70E-04 | 1.00E-03 | 9.23E-04 | 1.01E-03 | 6.54E-04 |
| AAVHNL | 9.45E-06 | 1.51E-05 | 3.63E-04 | 4.14E-05 | 3.65E-05 | 1.85E-04 | 2.39E-04 | 2.38E-04 | nan |
| LAQRVSGL | 5.91E-07 | 1.74E-04 | 6.02E-04 | 4.89E-05 | 1.68E-04 | 1.65E-04 | 3.57E-04 | 4.02E-04 | 4.27E-03 |
| VSGLIIN | 2.48E-05 | 4.59E-06 | 3.37E-05 | 1.16E-05 | 1.64E-06 | 1.42E-05 | 2.59E-06 | 1.22E-05 | 1.20E-05 |
| IINYPLD | 2.19E-05 | 1.27E-05 | 3.67E-06 | nan | 1.48E-05 | 3.53E-05 | 3.15E-06 | 1.80E-05 | 3.07E-04 |
| LDDQDAIA | 6.39E-06 | 9.93E-04 | 1.20E-03 | 1.52E-04 | 6.36E-04 | 6.46E-04 | 7.54E-04 | 6.61E-04 | nan |
| IAVEAA | 1.17E-06 | 4.74E-05 | 1.54E-04 | 9.32E-07 | 4.92E-05 | 5.97E-05 | 3.30E-04 | 1.62E-04 | 3.98E-03 |
| ACTNVPAL | 2.44E-05 | 1.65E-03 | 1.08E-03 | 1.20E-04 | 9.28E-04 | 1.28E-03 | 1.12E-03 | 1.91E-03 | nan |
| ALFLD | 1.12E-05 | 2.67E-05 | 1.30E-04 | 2.05E-05 | 1.23E-05 | 6.55E-06 | 2.60E-05 | 2.55E-04 | nan |
| FLDVSDQTPINS | 1.23E-05 | 2.75E-04 | 7.62E-04 | 7.99E-04 | 1.28E-04 | 6.00E-04 | 5.53E-04 | 9.98E-04 | 1.37E-03 |
| DQTPIN | 2.26E-05 | 6.11E-05 | 4.45E-04 | 4.25E-04 | 5.56E-04 | 1.05E-03 | 4.94E-04 | 8.32E-04 | nan |
| SIIFSH | nan | nan | nan | nan | nan | nan | nan | nan | nan |
| SIIFSHEDGTRL | 1.09E-06 | 2.00E-04 | nan | 2.51E-04 | 3.68E-05 | 4.89E-05 | 2.04E-05 | 6.85E-05 | 1.38E-04 |
| EDGTRLGVEH | 1.98E-05 | 7.57E-05 | 3.63E-04 | nan | 1.27E-04 | 4.91E-04 | 1.29E-04 | 1.11E-04 | nan |
| LVAL | 6.23E-07 | 1.14E-07 | 1.15E-05 | 1.44E-04 | 2.27E-07 | 1.75E-05 | 5.60E-06 | 9.91E-07 | 2.91E-07 |
| LVALGHQQIAL | 4.55E-06 | 3.16E-04 | 2.86E-04 | 3.53E-04 | 1.24E-04 | 2.91E-04 | 1.60E-04 | 3.39E-04 | 1.09E-03 |
| LGHQQIALL | 1.71E-05 | 3.75E-05 | 2.40E-06 | 1.05E-05 | 1.62E-05 | 7.28E-05 | 4.73E-05 | 5.61E-05 | 2.81E-05 |
| IALL | 2.80E-06 | 9.79E-06 | 2.05E-06 | 6.97E-05 | 5.91E-07 | 1.82E-05 | 4.33E-06 | 8.57E-06 | nan |

| | | | | | | | | | |
|---|---|---|---|---|---|---|---|---|---|
| **LAGPLS** | 4.27E-05 | 4.58E-04 | 1.12E-04 | 4.23E-04 | 1.94E-04 | 9.24E-04 | 1.28E-03 | 9.46E-04 | nan |
| **LAGPLSSVSARL** | 3.99E-05 | 1.52E-04 | 6.37E-04 | 3.00E-04 | 3.59E-04 | 1.01E-03 | 6.79E-04 | nan | nan |
| **GPLSSVSAR** | 5.38E-06 | 5.39E-04 | 1.75E-03 | nan | 2.11E-04 | 4.13E-04 | 6.92E-04 | 1.19E-03 | nan |
| **LAGWHK** | 3.83E-05 | 1.68E-04 | 2.48E-04 | nan | 2.98E-07 | 5.10E-04 | 1.58E-05 | 2.84E-04 | 3.64E-04 |
| **YLTRNQIQP** | 2.48E-06 | 4.18E-04 | nan | 9.31E-05 | 3.50E-04 | nan | 4.35E-04 | nan | 7.48E-04 |
| **NQIQPIAEREGD** | 7.73E-07 | 9.32E-04 | 7.35E-04 | 3.55E-04 | nan | 8.21E-04 | nan | 7.30E-04 | nan |
| **IAEREGDWSAMSG** | 4.08E-06 | 7.77E-04 | 1.12E-03 | 7.21E-04 | 1.01E-03 | 1.79E-03 | 1.13E-03 | 1.28E-03 | 7.91E-03 |
| **REGDWSAMSGF** | 5.64E-06 | 3.69E-04 | nan | 4.04E-04 | nan | 2.89E-04 | nan | 8.15E-04 | 3.76E-04 |
| **DWSAM** | 2.02E-05 | 1.66E-04 | 3.24E-04 | 7.28E-04 | 7.40E-04 | 1.02E-03 | 8.20E-04 | 9.87E-04 | nan |
| **WSAM** | 7.22E-05 | 1.14E-04 | 8.00E-04 | 2.55E-04 | 7.96E-04 | 9.75E-04 | 7.99E-04 | nan | 5.15E-04 |
| **SAMSGFQ** | 1.94E-05 | 6.50E-04 | nan | 5.45E-04 | 1.88E-03 | 3.68E-04 | 9.50E-05 | nan | nan |
| **SGFQQTMQM** | 6.60E-06 | 3.75E-06 | 3.53E-05 | nan | 3.96E-06 | 1.48E-05 | 8.52E-06 | 1.63E-05 | 8.20E-06 |
| **QTMQML** | 5.91E-06 | 1.94E-05 | 1.83E-05 | nan | 5.68E-07 | 6.46E-06 | 3.13E-06 | 7.57E-06 | 1.31E-06 |
| **NEGIVPTAML** | 2.05E-05 | 3.91E-04 | 3.62E-03 | 1.13E-05 | 5.41E-04 | 7.35E-04 | 1.60E-03 | 1.28E-03 | 6.31E-04 |
| **LVANDQMALGAM** | 1.42E-06 | 1.42E-04 | nan | nan | 1.37E-04 | 5.12E-04 | 1.79E-04 | 1.50E-04 | 9.87E-05 |
| **VANDQMALGAMR** | 1.14E-06 | 9.91E-05 | 1.06E-04 | nan | 7.81E-05 | 2.53E-04 | 1.58E-04 | 1.10E-04 | 6.11E-04 |
| **AITESGLRVGAD** | 8.41E-06 | 8.58E-04 | 5.90E-04 | 5.25E-04 | 5.27E-04 | 8.52E-04 | 4.47E-04 | 1.02E-03 | 2.17E-03 |
| **ITESGLRVGAD** | 1.14E-05 | 1.09E-03 | 3.03E-04 | 2.97E-04 | 2.95E-04 | 6.42E-04 | 3.37E-04 | 8.12E-04 | 6.76E-04 |
| **ADISVVGY** | 1.91E-06 | 3.78E-05 | 9.98E-06 | nan | 3.60E-06 | 3.40E-04 | 2.42E-05 | 8.55E-06 | 2.03E-04 |
| **VVGYDD** | 8.23E-07 | 3.44E-05 | 5.91E-05 | nan | 7.65E-05 | 1.87E-04 | 3.05E-04 | 1.40E-04 | 9.92E-04 |
| **DDTEDSSCYIPPLTTIK** | 1.80E-06 | 1.91E-04 | 6.56E-04 | nan | 2.54E-04 | 5.81E-04 | 8.07E-04 | 1.12E-03 | nan |
| **TEDSSCYIPPLTTIK** | 2.66E-06 | 5.35E-05 | 5.37E-04 | nan | 2.85E-05 | 1.73E-04 | 1.76E-04 | 2.80E-04 | nan |
| **SSCYIPPLTTIKQD** | 4.82E-07 | 9.66E-05 | 5.53E-04 | 3.34E-03 | 5.74E-05 | 3.28E-04 | 4.37E-04 | 2.58E-04 | 7.58E-03 |
| **FRLL** | 6.77E-07 | 3.48E-06 | 1.17E-04 | 1.61E-04 | 5.69E-06 | 1.72E-05 | 1.53E-04 | 1.44E-04 | 1.66E-03 |
| **FRLLGQTSV** | 5.84E-06 | 5.64E-05 | nan | 1.48E-04 | 3.21E-05 | 5.98E-05 | nan | 3.69E-05 | 3.15E-05 |
| **LGQTSVDR** | 2.41E-07 | 2.02E-05 | 3.72E-05 | 1.58E-04 | 1.78E-05 | 6.39E-05 | 4.29E-05 | 1.15E-04 | 5.27E-04 |
| **GQTSVDRL** | 6.18E-07 | nan | nan | 8.16E-05 | nan | 6.47E-05 | 1.54E-05 | 2.37E-05 | nan |
| **LLQL** | 2.46E-06 | 9.12E-05 | 7.41E-04 | 9.31E-05 | 2.85E-04 | 5.98E-04 | 9.73E-04 | 1.27E-03 | 4.71E-03 |

### TMG vs ONPFDNA

| Peptide | 0 s | 30 s | 45 s | 60 s | 300 s | 1500 s | 3600 s | 7200 s | 14400 s |
|---|---|---|---|---|---|---|---|---|---|
| LIGVA | 1.62E-06 | 2.31E-05 | 5.69E-06 | 1.11E-05 | 1.49E-05 | 4.67E-06 | 1.42E-05 | 2.48E-06 | 3.15E-05 |
| ALHAP | 6.59E-07 | 4.07E-05 | 6.15E-06 | 2.50E-04 | 6.83E-05 | 1.47E-04 | 2.28E-04 | 3.46E-04 | 2.56E-04 |
| ALHAPSQIVA | 3.06E-06 | 1.32E-04 | nan | 8.01E-04 | 3.98E-05 | 4.19E-04 | 1.53E-04 | 4.24E-04 | 4.99E-04 |
| ALHAPSQIVAA | 9.53E-07 | 1.32E-04 | nan | 7.40E-04 | 6.00E-05 | 3.13E-04 | 1.90E-04 | 4.66E-04 | 4.06E-04 |
| ALHAPSQIVAAIK | 5.39E-06 | 1.91E-04 | 1.72E-04 | nan | 1.99E-04 | 4.93E-05 | 1.29E-03 | 5.19E-04 | 2.43E-05 |
| SQIVAA | 3.75E-05 | 1.61E-05 | 3.35E-05 | 8.94E-05 | nan | 2.26E-04 | 9.88E-05 | 2.02E-05 | 9.40E-06 |
| SQIVAAIK | 1.24E-06 | 1.08E-05 | 2.90E-05 | 9.13E-05 | 9.45E-07 | 3.17E-05 | 1.94E-04 | 1.22E-04 | 6.51E-05 |
| IVAAIK | 2.98E-06 | 2.24E-06 | 9.62E-06 | 4.37E-05 | 1.11E-06 | 1.58E-05 | 2.59E-05 | 4.94E-05 | 9.90E-06 |
| AIKSRADQLGAS | 8.36E-07 | 3.01E-04 | 1.84E-05 | 3.30E-05 | 9.37E-05 | 3.66E-04 | 3.31E-04 | 9.40E-04 | 6.98E-04 |
| ADQLGAS | 4.43E-05 | 2.22E-04 | 2.83E-05 | 3.34E-04 | 4.26E-04 | 6.28E-04 | 1.36E-03 | 4.63E-03 | 1.68E-03 |
| VVVSM | 6.16E-05 | 2.56E-04 | 4.00E-04 | 1.29E-04 | 2.02E-04 | 1.38E-04 | 4.56E-05 | 5.93E-04 | 4.69E-04 |
| MVERSGVE | 1.09E-05 | 3.08E-03 | nan | 2.41E-04 | 4.59E-04 | 1.08E-03 | 9.41E-04 | 1.04E-03 | 9.52E-04 |
| AAVHNL | 3.49E-05 | 2.95E-07 | 3.05E-04 | 5.75E-05 | 2.01E-04 | 4.27E-05 | 2.70E-04 | nan | 9.75E-05 |
| LAQRVSGL | 3.86E-07 | 1.72E-04 | 1.09E-04 | 4.38E-05 | 2.89E-05 | 6.19E-05 | 1.33E-04 | 3.37E-04 | 1.58E-04 |
| VSGLIIN | 2.42E-05 | 9.04E-06 | 4.64E-05 | 3.30E-06 | 1.06E-04 | 1.41E-05 | 2.04E-06 | 3.78E-05 | 1.07E-05 |
| IINYPLD | 2.17E-05 | 1.11E-05 | 7.04E-06 | nan | 2.84E-06 | 6.96E-05 | 1.14E-05 | 9.57E-06 | 9.21E-06 |
| LDDQDAIA | 6.88E-06 | 7.01E-04 | 1.22E-04 | 1.07E-04 | 3.24E-04 | 4.60E-04 | 1.82E-04 | 4.01E-04 | 2.84E-04 |
| IAVEAA | 1.68E-07 | 9.28E-06 | 1.62E-04 | 1.19E-05 | 7.57E-06 | 1.69E-04 | 1.63E-05 | 1.65E-04 | 3.97E-05 |
| ACTNVPAL | 2.49E-05 | 1.65E-03 | 1.43E-06 | 1.18E-04 | 1.38E-04 | 7.26E-04 | 3.87E-04 | 1.73E-03 | 6.51E-04 |
| ALFLD | 1.15E-05 | 2.21E-05 | nan | 7.81E-05 | 4.43E-04 | 2.29E-05 | 1.28E-04 | 1.43E-04 | 2.67E-04 |
| FLDVSDQTPINS | 5.62E-06 | 3.38E-04 | nan | 6.47E-04 | 9.22E-05 | 4.44E-04 | 5.19E-04 | 5.37E-04 | 2.42E-04 |
| DQTPIN | 8.76E-06 | 6.59E-05 | 2.20E-04 | 4.24E-04 | 1.31E-04 | 4.73E-04 | 1.81E-04 | 6.49E-04 | 9.56E-05 |
| SIIFSH | 5.61E-04 | 2.26E-04 | 7.31E-04 | 1.04E-03 | 4.85E-04 | 5.41E-04 | 1.03E-03 | 8.39E-04 | nan |
| SIIFSHEDGTRL | 2.97E-07 | 2.02E-04 | nan | 2.55E-04 | 5.79E-05 | 6.03E-05 | 3.30E-05 | 9.56E-05 | 4.22E-05 |
| EDGTRLGVEH | 1.78E-05 | 8.30E-05 | 3.17E-04 | nan | 1.01E-04 | 4.79E-04 | 1.54E-04 | 6.50E-05 | 4.25E-04 |
| LVAL | 1.14E-07 | 4.05E-07 | 7.11E-06 | 1.44E-04 | 7.73E-08 | 2.08E-05 | 8.02E-06 | 5.91E-07 | 4.11E-06 |
| LVALGHQQIAL | 4.13E-06 | 2.94E-04 | 5.87E-05 | 3.51E-04 | 5.54E-05 | 2.14E-04 | 1.02E-04 | 4.45E-04 | 7.34E-05 |
| LGHQQIALL | 7.83E-06 | 5.80E-05 | 8.10E-05 | 6.53E-05 | 2.81E-05 | 5.23E-05 | 2.42E-05 | 4.78E-05 | 1.48E-05 |
| IALL | 3.11E-06 | 6.51E-06 | 1.68E-06 | 7.42E-05 | 8.77E-07 | 1.59E-05 | 8.82E-07 | 3.68E-06 | 2.42E-05 |

| | | | | | | | | | |
|---|---|---|---|---|---|---|---|---|---|
| **LAGPLS** | 2.41E-05 | 5.20E-04 | 7.71E-06 | 4.31E-04 | 1.94E-05 | 1.99E-04 | 1.53E-05 | nan | 2.94E-04 |
| **LAGPLSSVSARL** | 2.59E-05 | 1.92E-04 | 1.46E-04 | 2.49E-04 | 1.67E-04 | 6.72E-04 | 2.43E-04 | nan | nan |
| **GPLSSVSAR** | 3.64E-06 | 4.43E-04 | 3.09E-04 | nan | 8.82E-06 | 4.20E-04 | 2.95E-04 | 1.49E-03 | nan |
| **LAGWHK** | 2.05E-05 | 1.94E-04 | 2.44E-04 | nan | 2.76E-05 | 5.12E-04 | nan | 2.73E-04 | 4.51E-04 |
| **YLTRNQIQP** | 1.33E-06 | 4.60E-04 | nan | 1.40E-04 | 1.31E-04 | nan | 2.52E-04 | nan | nan |
| **NQIQPIAEREGD** | 6.23E-07 | 9.03E-04 | 2.17E-06 | 3.54E-04 | nan | 5.85E-04 | nan | 8.28E-04 | nan |
| **IAEREGDWSAMSG** | 1.54E-06 | 7.02E-04 | 3.51E-04 | 7.05E-04 | 4.38E-04 | 1.00E-03 | 5.67E-04 | 9.37E-04 | 5.61E-04 |
| **REGDWSAMSGF** | 2.77E-05 | 4.25E-04 | 6.46E-04 | 3.97E-04 | nan | 2.69E-04 | nan | 7.44E-04 | 4.19E-04 |
| **DWSAM** | 1.37E-05 | 1.49E-04 | 1.64E-04 | 6.91E-04 | 1.87E-04 | 5.94E-04 | 3.56E-04 | 7.79E-04 | 3.53E-04 |
| **WSAM** | 1.18E-05 | 4.57E-05 | 2.90E-04 | 3.46E-04 | 2.12E-04 | 2.06E-04 | 2.50E-04 | nan | nan |
| **SAMSGFQ** | 7.80E-05 | 3.87E-04 | 8.60E-04 | 4.80E-04 | 2.85E-04 | 1.64E-04 | 8.80E-05 | nan | 1.71E-04 |
| **SGFQQTMQM** | 9.60E-06 | 1.17E-06 | 3.57E-05 | nan | 5.27E-06 | 2.23E-05 | 4.15E-06 | 1.40E-05 | 8.41E-06 |
| **QTMQML** | 9.46E-06 | 1.18E-05 | 1.63E-05 | nan | 8.23E-07 | 1.52E-05 | 2.47E-06 | 5.94E-06 | 2.62E-05 |
| **NEGIVPTAML** | 4.42E-05 | 3.70E-04 | 2.00E-03 | 8.31E-06 | 1.95E-04 | 4.60E-04 | 5.95E-04 | 1.04E-03 | 1.28E-03 |
| **LVANDQMALGAM** | 1.19E-04 | 2.00E-03 | nan | nan | 2.97E-03 | 4.84E-03 | 3.90E-03 | 2.08E-03 | 4.99E-03 |
| **VANDQMALGAMR** | 8.82E-07 | 1.04E-04 | 5.10E-05 | nan | 4.32E-05 | 1.98E-04 | 1.73E-04 | 1.63E-04 | 9.68E-05 |
| **AITESGLRVGAD** | 8.42E-06 | 7.64E-04 | 8.03E-05 | 5.01E-04 | 2.57E-04 | 7.04E-04 | 2.07E-04 | 9.76E-04 | 3.29E-04 |
| **ITESGLRVGAD** | 1.05E-05 | 1.19E-03 | 7.78E-05 | 3.76E-04 | 2.52E-04 | 6.08E-04 | 3.21E-04 | 8.08E-04 | 3.32E-04 |
| **ADISVVGY** | 1.68E-06 | 5.54E-05 | 6.95E-05 | nan | 1.26E-05 | 1.94E-04 | 3.26E-05 | 3.53E-05 | 3.90E-05 |
| **VVGYDD** | 5.91E-07 | 4.88E-05 | 4.76E-05 | nan | 2.85E-05 | 8.14E-05 | 3.15E-05 | 8.33E-05 | 4.82E-05 |
| **DDTEDSSCYIPPLTTIK** | 3.86E-07 | 1.56E-04 | 1.82E-04 | nan | 1.29E-04 | 2.64E-04 | 7.47E-05 | 6.80E-04 | 9.19E-04 |
| **TEDSSCYIPPLTTIK** | 2.84E-06 | 6.07E-05 | 1.34E-04 | nan | 5.89E-05 | 9.53E-05 | 6.34E-05 | 2.71E-04 | 3.36E-04 |
| **SSCYIPPLTTIKQD** | 3.36E-07 | 9.66E-05 | 1.87E-04 | 3.31E-03 | 2.40E-05 | 2.39E-04 | 1.00E-05 | 9.94E-05 | 1.92E-04 |
| **FRLL** | 6.77E-07 | 3.03E-06 | 5.18E-05 | 1.57E-04 | 3.09E-06 | 2.60E-05 | 4.20E-05 | 1.22E-04 | 4.49E-05 |
| **FRLLGQTSV** | 5.96E-05 | 6.28E-05 | 1.13E-04 | 1.05E-04 | 2.84E-05 | 9.46E-05 | nan | 1.69E-05 | 4.49E-05 |
| **LGQTSVDR** | 2.23E-07 | 1.31E-05 | 9.20E-06 | 1.57E-04 | 6.18E-06 | 5.21E-05 | 3.00E-06 | 3.22E-05 | 2.05E-05 |
| **GQTSVDRL** | 3.27E-07 | nan | nan | 8.18E-05 | nan | 5.85E-05 | 4.73E-06 | 2.68E-05 | nan |
| **LLQL** | 3.48E-06 | 6.07E-05 | 4.01E-04 | 9.25E-05 | 6.40E-05 | 1.25E-04 | 3.76E-04 | 9.28E-04 | 7.08E-04 |
