## Appendix 3 for "Ligand-induced changes in dynamics mediate long-range allostery in the *lac* repressor"

**Appendix 3. Compiled raw HDX/MS data.** Averages and standard deviations for MS centroids from HDX/MS technical replicates were calculated for all states at each timepoint. Averaged centroid values were converted to the number of deuterons exchanged given the experimentally observed centroid of the maximally labeled control sample, as described in the Methods.

### 63-67 LIGVA

**Charge** 1

**IPTG**

| Time | centroid | D |
|---|---|---|
| 0 | 472.5075 | -0.02 |
| 30 | 472.606 | 0.15 |
| 45 | 472.5805 | 0.10 |
| 60 | 472.566 | 0.08 |
| 300 | 472.549 | 0.05 |
| 1500 | 472.5895 | 0.12 |
| 3600 | 472.587 | 0.11 |
| 7200 | 472.6015 | 0.14 |
| 14400 | 472.6275 | 0.18 |

**ONPF**

| Time | centroid | D |
|---|---|---|
| 0 | 472.508 | -0.02 |
| 30 | 472.537 | 0.03 |
| 45 | 472.561 | 0.07 |
| 60 | 472.582 | 0.11 |
| 300 | 472.5505 | 0.05 |
| 1500 | 472.58 | 0.10 |
| 3600 | 472.574 | 0.09 |
| 7200 | 472.578 | 0.10 |
| 14400 | 472.58 | 0.10 |

**APO**

| Time | centroid | D |
|---|---|---|
| 0 | 472.5175 | 0.00 |
| 30 | 472.5625 | 0.07 |
| 45 | 472.576 | 0.10 |
| 60 | 472.5655 | 0.08 |
| 300 | 472.5635 | 0.07 |
| 1500 | 472.579 | 0.10 |
| 3600 | 472.5965 | 0.13 |
| 7200 | 472.584 | 0.11 |
| 14400 | 472.6515 | 0.22 |

**DNA**

| Time | centroid | D |
|---|---|---|
| 0 | 472.5095 | -0.02 |
| 30 | 472.55 | 0.05 |
| 45 | 472.545 | 0.04 |
| 60 | 472.547 | 0.05 |
| 300 | 472.543 | 0.04 |
| 1500 | 472.5265 | 0.01 |
| 3600 | 472.561 | 0.07 |
| 7200 | 472.567 | 0.08 |
| 14400 | 472.5585 | 0.07 |

**ONPFDNA**

| Time | centroid | D |
|---|---|---|
| 0 | 472.513 | -0.01 |
| 30 | 472.5485 | 0.05 |
| 45 | 472.547 | 0.05 |
| 60 | 472.564 | 0.07 |
| 300 | 472.553 | 0.06 |
| 1500 | 472.562 | 0.07 |
| 3600 | 472.562 | 0.07 |
| 7200 | 472.573 | 0.09 |
| 14400 | 472.658 | 0.24 |

**TMG**

| Time | centroid | D |
|---|---|---|
| 0 | 472.532 | 0.02 |
| 30 | 472.589 | 0.12 |
| 45 | 472.595 | 0.13 |
| 60 | 472.628 | 0.18 |
| 300 | 472.595 | 0.13 |
| 1500 | 472.598 | 0.13 |
| 3600 | 472.6225 | 0.17 |
| 7200 | 472.5925 | 0.12 |
| 14400 | 472.616 | 0.16 |

control 472.52
infinity 474.281

$$D(t) = \frac{Mt - M0}{M\infty - M0} *N$$

**IPTG**

| Time | Replicate 1 | Replicate 2 | Replicate 3 | Replicate 4 | Replicate5 | average | SD |
|---|---|---|---|---|---|---|---|
| 0 | 472.525 | 472.49 | | | | 472.51 | 0.02 |
| 30 | 472.618 | 472.594 | | | | 472.61 | 0.02 |
| 45 | 472.641 | 472.52 | | | | 472.58 | 0.09 |
| 60 | 472.604 | 472.528 | | | | 472.57 | 0.05 |
| 300 | 472.547 | 472.551 | | | | 472.55 | 0.00 |
| 1500 | 472.603 | 472.576 | | | | 472.59 | 0.02 |
| 3600 | 472.592 | 472.582 | | | | 472.59 | 0.01 |
| 7200 | 472.613 | 472.59 | | | | 472.60 | 0.02 |
| 14400 | 472.657 | 472.598 | | | | 472.63 | 0.04 |

**ONPF**

| Time | Replicate 1 | Replicate 2 | Replicate 3 | Replicate 4 | average | SD |
|---|---|---|---|---|---|---|
| 0 | 472.517 | 472.499 | | | 472.51 | 0.01 |
| 30 | 472.55 | 472.524 | | | 472.54 | 0.02 |
| 45 | 472.597 | 472.525 | | | 472.56 | 0.05 |
| 60 | 472.607 | 472.557 | | | 472.58 | 0.04 |
| 300 | 472.581 | 472.52 | | | 472.55 | 0.04 |
| 1500 | 472.608 | 472.552 | | | 472.58 | 0.04 |
| 3600 | 472.575 | 472.573 | | | 472.57 | 0.00 |
| 7200 | 472.562 | 472.594 | | | 472.58 | 0.02 |
| 14400 | 472.621 | 472.534 | | | 472.58 | 0.06 |

**APO**

| Time | Replicate 1 | Replicate 2 | Replicate 3 | Replicate 4 | Replicate 5 | average | SD |
|---|---|---|---|---|---|---|---|
| 0 | 472.531 | 472.504 | | | | 472.52 | 0.02 |
| 30 | 472.598 | 472.527 | | | | 472.56 | 0.05 |
| 45 | 472.587 | 472.565 | | | | 472.58 | 0.02 |
| 60 | 472.603 | 472.528 | | | | 472.57 | 0.05 |
| 300 | 472.606 | 472.521 | | | | 472.56 | 0.06 |
| 1500 | 472.568 | 472.59 | | | | 472.58 | 0.02 |
| 3600 | 472.583 | 472.61 | | | | 472.60 | 0.02 |
| 7200 | 472.563 | 472.605 | | | | 472.58 | 0.03 |
| 14400 | 472.604 | 472.699 | | | | 472.65 | 0.07 |

**DNA**

| Time | Replicate 1 | Replicate 2 | Replicate 3 | Replicate 4 | Replicate 5 | average | SD |
|---|---|---|---|---|---|---|---|
| 0 | 472.509 | 472.51 | | | | 472.51 | 0.00 |
| 30 | 472.577 | 472.523 | | | | 472.55 | 0.04 |
| 45 | 472.574 | 472.516 | | | | 472.55 | 0.04 |
| 60 | 472.573 | 472.521 | | | | 472.55 | 0.04 |
| 300 | 472.58 | 472.506 | | | | 472.54 | 0.05 |
| 1500 | 472.534 | 472.519 | | | | 472.53 | 0.01 |
| 3600 | 472.558 | 472.564 | | | | 472.56 | 0.00 |
| 7200 | 472.574 | 472.56 | | | | 472.57 | 0.01 |
| 14400 | 472.58 | 472.537 | | | | 472.56 | 0.03 |

**ONPFDNA**

| Time | Replicate 1 | Replicate 2 | Replicate 3 | Replicate 4 | average | SD |
|---|---|---|---|---|---|---|
| 0 | 472.521 | 472.505 | | | 472.51 | 0.01 |
| 30 | 472.584 | 472.513 | | | 472.55 | 0.05 |
| 45 | 472.559 | 472.535 | | | 472.55 | 0.02 |
| 60 | 472.587 | 472.541 | | | 472.56 | 0.03 |
| 300 | 472.581 | 472.525 | | | 472.55 | 0.04 |
| 1500 | 472.578 | 472.546 | | | 472.56 | 0.02 |
| 3600 | 472.567 | 472.557 | | | 472.56 | 0.01 |
| 7200 | 472.581 | 472.565 | | | 472.57 | 0.01 |
| 14400 | 472.696 | 472.62 | | | 472.66 | 0.05 |

**TMG**

| Time | Replicate 1 | Replicate 2 | Replicate 3 | Replicate 4 | Replicate 5 | average | SD |
|---|---|---|---|---|---|---|---|
| 0 | 472.537 | 472.527 | | | | 472.53 | 0.01 |
| 30 | 472.592 | 472.586 | | | | 472.59 | 0.00 |
| 45 | 472.608 | 472.582 | | | | 472.60 | 0.02 |
| 60 | 472.637 | 472.619 | | | | 472.63 | 0.01 |
| 300 | 472.601 | 472.589 | | | | 472.60 | 0.01 |
| 1500 | 472.599 | 472.597 | | | | 472.60 | 0.00 |
| 3600 | 472.65 | 472.595 | | | | 472.62 | 0.04 |
| 7200 | 472.601 | 472.584 | | | | 472.59 | 0.01 |
| 14400 | 472.633 | 472.599 | | | | 472.62 | 0.02 |

### 72-76 ALHAP

**Charge** 1

**IPTG**

| Time | centroid | D |
|---|---|---|
| 0 | 508.5555 | 0.00 |
| 30 | 508.7 | 0.28 |
| 45 | 508.7035 | 0.29 |
| 60 | 508.7475 | 0.38 |
| 300 | 508.893 | 0.66 |
| 1500 | 509.069 | 1.01 |
| 3600 | 509.31 | 1.48 |
| 7200 | 509.3935 | 1.65 |
| 14400 | 509.326 | 1.52 |

**ONPF**

| Time | centroid | D |
|---|---|---|
| 0 | 508.5615 | 0.01 |
| 30 | 508.938 | 0.75 |
| 45 | 509.098 | 1.07 |
| 60 | 509.1705 | 1.21 |
| 300 | 509.1495 | 1.17 |
| 1500 | 509.1875 | 1.24 |
| 3600 | 509.186 | 1.24 |
| 7200 | 509.1815 | 1.23 |
| 14400 | 509.03 | 0.93 |

**APO**

| Time | centroid | D |
|---|---|---|
| 0 | 508.5555 | 0.00 |
| 30 | 509.253 | 1.37 |
| 45 | 509.159 | 1.19 |
| 60 | 509.3405 | 1.54 |
| 300 | 509.221 | 1.31 |
| 1500 | 509.321 | 1.51 |
| 3600 | 509.3455 | 1.55 |
| 7200 | 509.2555 | 1.38 |
| 14400 | 509.3365 | 1.54 |

**DNA**

| Time | centroid | D |
|---|---|---|
| 0 | 508.563 | 0.01 |
| 30 | 509.193 | 1.25 |
| 45 | 509.274 | 1.41 |
| 60 | 509.195 | 1.26 |
| 300 | 509.24 | 1.35 |
| 1500 | 509.279 | 1.42 |
| 3600 | 509.2245 | 1.32 |
| 7200 | 509.202 | 1.27 |
| 14400 | 509.272 | 1.41 |

**ONPFDNA**

| Time | centroid | D |
|---|---|---|
| 0 | 508.555 | 0.00 |
| 30 | 509.116 | 1.10 |
| 45 | 509.247 | 1.36 |
| 60 | 509.1645 | 1.20 |
| 300 | 509.215 | 1.30 |
| 1500 | 509.0545 | 0.98 |
| 3600 | 509.2575 | 1.38 |
| 7200 | 509.1975 | 1.26 |
| 14400 | 508.971 | 0.82 |

**TMG**

| Time | centroid | D |
|---|---|---|
| 0 | 508.5645 | 0.02 |
| 30 | 508.7975 | 0.48 |
| 45 | 508.861 | 0.60 |
| 60 | 509.0495 | 0.97 |
| 300 | 509.1405 | 1.15 |
| 1500 | 509.2715 | 1.41 |
| 3600 | 509.4275 | 1.72 |
| 7200 | 509.4125 | 1.69 |
| 14400 | 509.489 | 1.84 |

control 508.5555
infinity 509.572

$$D(t) = \frac{M_t - M_0}{M_\infty - M_0} * N$$

**IPTG**

| Time | Replicate 1 | Replicate 2 | Replicate 3 | Replicate 4 | Replicate5 | average | SD |
|---|---|---|---|---|---|---|---|
| 0 | 508.559 | 508.552 | | | | 508.56 | 0.00 |
| 30 | 508.73 | 508.67 | | | | 508.70 | 0.04 |
| 45 | 508.723 | 508.684 | | | | 508.70 | 0.03 |
| 60 | 508.783 | 508.712 | | | | 508.75 | 0.05 |
| 300 | 508.946 | 508.84 | | | | 508.89 | 0.07 |
| 1500 | 509.193 | 508.945 | | | | 509.07 | 0.18 |
| 3600 | 509.383 | 509.237 | | | | 509.31 | 0.10 |
| 7200 | 509.516 | 509.271 | | | | 509.39 | 0.17 |
| 14400 | 509.444 | 509.208 | | | | 509.33 | 0.17 |

**ONPF**

| Time | Replicate 1 | Replicate 2 | Replicate 3 | Replicate 4 | average | SD |
|---|---|---|---|---|---|---|
| 0 | 508.567 | 508.556 | | | 508.56 | 0.01 |
| 30 | 509.002 | 508.874 | | | 508.94 | 0.09 |
| 45 | 509.23 | 508.966 | | | 509.10 | 0.19 |
| 60 | 509.22 | 509.121 | | | 509.17 | 0.07 |
| 300 | 509.281 | 509.018 | | | 509.15 | 0.19 |
| 1500 | 509.313 | 509.062 | | | 509.19 | 0.18 |
| 3600 | 509.267 | 509.105 | | | 509.19 | 0.11 |
| 7200 | 509.278 | 509.085 | | | 509.18 | 0.14 |
| 14400 | 509.347 | 508.706 | | | 509.03 | 0.45 |

**APO**

| Time | Replicate 1 | Replicate 2 | Replicate 3 | Replicate 4 | Replicate 5 | average | SD |
|---|---|---|---|---|---|---|---|
| 0 | 508.554 | 508.557 | | | | 508.56 | 0.00 |
| 30 | 509.363 | 509.143 | | | | 509.25 | 0.16 |
| 45 | 509.22 | 509.098 | | | | 509.16 | 0.09 |
| 60 | 509.506 | 509.175 | | | | 509.34 | 0.23 |
| 300 | 509.441 | 509.001 | | | | 509.22 | 0.31 |
| 1500 | 509.446 | 509.196 | | | | 509.32 | 0.18 |
| 3600 | 509.48 | 509.211 | | | | 509.35 | 0.19 |
| 7200 | 509.403 | 509.108 | | | | 509.26 | 0.21 |
| 14400 | 509.428 | 509.245 | | | | 509.34 | 0.13 |

**DNA**

| Time | Replicate 1 | Replicate 2 | Replicate 3 | Replicate 4 | Replicate 5 | average | SD |
|---|---|---|---|---|---|---|---|
| 0 | 508.554 | 508.572 | | | | 508.56 | 0.01 |
| 30 | 509.227 | 509.159 | | | | 509.19 | 0.05 |
| 45 | 509.159 | 509.389 | | | | 509.27 | 0.16 |
| 60 | 509.25 | 509.14 | | | | 509.20 | 0.08 |
| 300 | 509.296 | 509.184 | | | | 509.24 | 0.08 |
| 1500 | 509.298 | 509.26 | | | | 509.28 | 0.03 |
| 3600 | 509.321 | 509.128 | | | | 509.22 | 0.14 |
| 7200 | 509.361 | 509.043 | | | | 509.20 | 0.22 |
| 14400 | 509.272 | | | | | 509.27 | #DIV/0! |

**ONPFDNA**

| Time | Replicate 1 | Replicate 2 | Replicate 3 | Replicate 4 | average | SD |
|---|---|---|---|---|---|---|
| 0 | 508.559 | 508.551 | | | 508.56 | 0.01 |
| 30 | 509.1 | 509.132 | | | 509.12 | 0.02 |
| 45 | 509.254 | 509.24 | | | 509.25 | 0.01 |
| 60 | 509.209 | 509.12 | | | 509.16 | 0.06 |
| 300 | 509.275 | 509.155 | | | 509.22 | 0.08 |
| 1500 | 509.07 | 509.039 | | | 509.05 | 0.02 |
| 3600 | 509.269 | 509.246 | | | 509.26 | 0.02 |
| 7200 | 509.257 | 509.138 | | | 509.20 | 0.08 |
| 14400 | 508.929 | 509.013 | | | 508.97 | 0.06 |

**TMG**

| Time | Replicate 1 | Replicate 2 | Replicate 3 | Replicate 4 | Replicate 5 | average | SD |
|---|---|---|---|---|---|---|---|
| 0 | 508.569 | 508.56 | | | | 508.56 | 0.01 |
| 30 | 508.753 | 508.842 | | | | 508.80 | 0.06 |
| 45 | 508.878 | 508.844 | | | | 508.86 | 0.02 |
| 60 | 508.941 | 509.158 | | | | 509.05 | 0.15 |
| 300 | 509.128 | 509.153 | | | | 509.14 | 0.02 |
| 1500 | 509.183 | 509.36 | | | | 509.27 | 0.13 |
| 3600 | 509.316 | 509.539 | | | | 509.43 | 0.16 |
| 7200 | 509.288 | 509.537 | | | | 509.41 | 0.18 |
| 14400 | 509.378 | 509.6 | | | | 509.49 | 0.16 |

### 72-81 ALHAPSQIVA

**Charge** 2

**IPTG**

| Time | centroid | D |
|---|---|---|
| 0 | 504.08 | 0.00 |
| 30 | 504.24 | 0.74 |
| 45 | 504.26 | 0.83 |
| 60 | 504.31 | 1.08 |
| 300 | 504.37 | 1.39 |
| 1500 | 504.47 | 1.87 |
| 3600 | 504.53 | 2.16 |
| 7200 | 504.66 | 2.74 |
| 14400 | 504.69 | 2.89 |

**ONPF**

| Time | centroid | D |
|---|---|---|
| 0 | 504.07 | -0.06 |
| 30 | 504.36 | 1.34 |
| 45 | 504.76 | 3.23 |
| 60 | 504.63 | 2.61 |
| 300 | 504.64 | 2.67 |
| 1500 | 504.84 | 3.62 |
| 3600 | 504.98 | 4.29 |
| 7200 | 505.07 | 4.74 |
| 14400 | 505.33 | 5.97 |

**APO**

| Time | centroid | D |
|---|---|---|
| 0 | 504.09 | 0.05 |
| 30 | 504.47 | 1.86 |
| 45 | 504.52 | 2.09 |
| 60 | 504.64 | 2.69 |
| 300 | 504.72 | 3.04 |
| 1500 | 505.00 | 4.39 |
| 3600 | 505.15 | 5.11 |
| 7200 | 505.33 | 5.99 |
| 14400 | 505.39 | 6.27 |

**DNA**

| Time | centroid | D |
|---|---|---|
| 0 | 504.09 | 0.02 |
| 30 | 504.46 | 1.79 |
| 45 | | |
| 60 | 504.53 | 2.14 |
| 300 | 504.51 | 2.03 |
| 1500 | 504.59 | 2.42 |
| 3600 | 504.65 | 2.73 |
| 7200 | 504.76 | 3.25 |
| 14400 | 504.85 | 3.69 |

**ONPFDNA**

| Time | centroid | D |
|---|---|---|
| 0 | 504.07 | -0.05 |
| 30 | 504.41 | 1.59 |
| 45 | 504.55 | 2.25 |
| 60 | 504.55 | 2.23 |
| 300 | 504.54 | 2.21 |
| 1500 | 504.57 | 2.32 |
| 3600 | 504.68 | 2.88 |
| 7200 | 504.69 | 2.90 |
| 14400 | 504.73 | 3.12 |

**TMG**

| Time | centroid | D |
|---|---|---|
| 0 | 504.09 | 0.01 |
| 30 | 504.25 | 0.80 |
| 45 | 504.37 | 1.38 |
| 60 | 504.55 | 2.23 |
| 300 | 504.47 | 1.84 |
| 1500 | 504.58 | 2.37 |
| 3600 | 504.74 | 3.16 |
| 7200 | 504.82 | 3.55 |
| 14400 | 505.00 | 4.40 |

control 504.083
infinity 505.543

$$D(t) = \frac{M_t - M_0}{M_\infty - M_0} * N$$

**IPTG**

| Time | Replicate 1 | Replicate 2 | Replicate 3 | Replicate 4 | Replicate5 | average | SD |
|---|---|---|---|---|---|---|---|
| 0 | 504.083 | 504.082 | 504.084 | 504.06 | 504.103 | 504.08 | 0.02 |
| 30 | 504.235 | 504.235 | 504.229 | | 504.252 | 504.24 | 0.01 |
| 45 | | | | | 504.256 | 504.26 | |
| 60 | 504.301 | 504.319 | 504.282 | | 504.335 | 504.31 | 0.02 |
| 300 | 504.35 | 504.35 | 504.313 | 504.454 | 504.396 | 504.37 | 0.05 |
| 1500 | 504.457 | 504.457 | 504.401 | 504.559 | 504.489 | 504.47 | 0.06 |
| 3600 | 504.511 | 504.511 | 504.476 | 504.595 | 504.57 | 504.53 | 0.05 |
| 7200 | 504.607 | 504.686 | 504.595 | 504.709 | 504.678 | 504.66 | 0.05 |
| 14400 | 504.723 | 504.723 | 504.594 | | 504.706 | 504.69 | 0.06 |

**ONPF**

| Time | Replicate 1 | Replicate 2 | Replicate 3 | Replicate 4 | average | SD |
|---|---|---|---|---|---|---|
| 0 | 504.085 | 504.083 | 504.015 | 504.099 | 504.07 | 0.04 |
| 30 | 504.456 | 504.32 | 504.312 | 504.359 | 504.36 | 0.07 |
| 45 | | | | 504.757 | 504.76 | |
| 60 | 504.644 | 504.539 | 504.74 | 504.589 | 504.63 | 0.09 |
| 300 | 504.619 | | 504.596 | 504.703 | 504.64 | 0.06 |
| 1500 | 504.812 | 504.794 | | 504.91 | 504.84 | 0.06 |
| 3600 | 505.122 | 504.82 | 504.843 | 505.123 | 504.98 | 0.17 |
| 7200 | 505.07 | 504.987 | 505.199 | 505.034 | 505.07 | 0.09 |
| 14400 | 505.339 | 505.268 | | 505.376 | 505.33 | 0.05 |

**APO**

| Time | Replicate 1 | Replicate 2 | Replicate 3 | Replicate 4 | Replicate 5 | average | SD |
|---|---|---|---|---|---|---|---|
| 0 | 504.083 | 504.083 | 504.082 | 504.138 | 504.078 | 504.09 | 0.03 |
| 30 | 504.417 | 504.417 | 504.407 | 504.587 | 504.523 | 504.47 | 0.08 |
| 45 | | | | | 504.518 | 504.52 | |
| 60 | 504.595 | 504.595 | 504.58 | 504.789 | 504.661 | 504.64 | 0.09 |
| 300 | 504.702 | 504.737 | 504.635 | 504.803 | 504.708 | 504.72 | 0.06 |
| 1500 | 504.97 | 504.97 | 504.94 | 505.119 | 504.998 | 505.00 | 0.07 |
| 3600 | 505.093 | 505.093 | 504.972 | 505.242 | 505.344 | 505.15 | 0.15 |
| 7200 | 505.311 | 505.311 | 505.112 | 505.481 | 505.443 | 505.33 | 0.14 |
| 14400 | 505.359 | 505.354 | 505.355 | 505.454 | 505.429 | 505.39 | 0.05 |

**DNA**

| Time | Replicate 1 | Replicate 2 | Replicate 3 | Replicate 4 | Replicate 5 | average | SD |
|---|---|---|---|---|---|---|---|
| 0 | 504.085 | 504.097 | 504.085 | 504.089 | 504.079 | 504.09 | 0.01 |
| 30 | 504.435 | 504.534 | 504.401 | 504.477 | 504.439 | 504.46 | 0.05 |
| 45 | | | | | | | |
| 60 | 504.548 | 504.389 | 504.478 | 504.761 | 504.47 | 504.53 | 0.14 |
| 300 | 504.508 | 504.377 | 504.463 | 504.641 | 504.548 | 504.51 | 0.10 |
| 1500 | 504.602 | 504.553 | 504.509 | 504.676 | 504.602 | 504.59 | 0.06 |
| 3600 | 504.686 | 504.401 | 504.604 | 504.737 | 504.831 | 504.65 | 0.16 |
| 7200 | 504.632 | 504.772 | 504.725 | 504.803 | 504.873 | 504.76 | 0.09 |
| 14400 | 504.862 | 504.717 | 504.786 | 504.886 | 505.01 | 504.85 | 0.11 |

**ONPFDNA**

| Time | Replicate 1 | Replicate 2 | Replicate 3 | Replicate 4 | average | SD |
|---|---|---|---|---|---|---|
| 0 | 504.083 | 504.049 | 504.084 | | 504.07 | 0.02 |
| 30 | 504.286 | 504.553 | 504.402 | | 504.41 | 0.13 |
| 45 | | | 504.552 | | 504.55 | |
| 60 | 504.439 | 504.707 | 504.497 | | 504.55 | 0.14 |
| 300 | 504.473 | 504.632 | 504.526 | | 504.54 | 0.08 |
| 1500 | 504.476 | 504.782 | 504.444 | | 504.57 | 0.19 |
| 3600 | 504.528 | 504.719 | 504.806 | | 504.68 | 0.14 |
| 7200 | 504.586 | 504.672 | 504.803 | | 504.69 | 0.11 |
| 14400 | 504.668 | 504.993 | 504.539 | | 504.73 | 0.23 |

**TMG**

| Time | Replicate 1 | Replicate 2 | Replicate 3 | Replicate 4 | Replicate 5 | average | SD |
|---|---|---|---|---|---|---|---|
| 0 | 504.091 | 504.079 | | | | 504.09 | 0.01 |
| 30 | 504.214 | 504.285 | | | | 504.25 | 0.05 |
| 45 | 504.402 | 504.339 | | | | 504.37 | 0.04 |
| 60 | 504.355 | 504.742 | | | | 504.55 | 0.27 |
| 300 | 504.468 | 504.465 | | | | 504.47 | 0.00 |
| 1500 | 504.47 | 504.684 | | | | 504.58 | 0.15 |
| 3600 | 504.701 | 504.783 | | | | 504.74 | 0.06 |
| 7200 | 504.685 | 504.963 | | | | 504.82 | 0.20 |
| 14400 | 504.904 | 505.096 | | | | 505.00 | 0.14 |

### 72-82 ALHAPSQIVAA

**Charge** 2

**IPTG**

| Time | centroid | D |
|---|---|---|
| 0 | 539.6288 | 0.00 |
| 30 | 539.8333 | 0.64 |
| 45 | 539.814 | 0.58 |
| 60 | 539.9164 | 0.90 |
| 300 | 539.929 | 0.94 |
| 1500 | 540.04 | 1.29 |
| 3600 | 540.1108 | 1.52 |
| 7200 | 540.2174 | 1.85 |
| 14400 | 540.2574 | 1.98 |

**ONPF**

| Time | centroid | D |
|---|---|---|
| 0 | 539.63 | 0.00 |
| 30 | 539.98 | 1.10 |
| 45 | 540.32 | 2.16 |
| 60 | 540.27 | 2.00 |
| 300 | 540.24 | 1.92 |
| 1500 | 540.50 | 2.74 |
| 3600 | 540.67 | 3.28 |
| 7200 | 540.77 | 3.60 |
| 14400 | 540.94 | 4.13 |

**APO**

| Time | centroid | D |
|---|---|---|
| 0 | 539.6333 | 0.01 |
| 30 | 540.0194 | 1.23 |
| 45 | 539.9805 | 1.11 |
| 60 | 540.2202 | 1.86 |
| 300 | 540.2282 | 1.89 |
| 1500 | 540.5934 | 3.04 |
| 3600 | 540.713 | 3.42 |
| 7200 | 540.8326 | 3.79 |
| 14400 | 540.651 | 3.22 |

**DNA**

| Time | centroid | D |
|---|---|---|
| 0 | 539.6333 | 0.01 |
| 30 | 539.974 | 1.09 |
| 45 | 540.0755 | 1.41 |
| 60 | 540.105 | 1.50 |
| 300 | 540.0598 | 1.36 |
| 1500 | 540.1285 | 1.57 |
| 3600 | 540.219 | 1.86 |
| 7200 | 540.27 | 2.02 |
| 14400 | 540.3135 | 2.16 |

**ONPFDNA**

| Time | centroid | D |
|---|---|---|
| 0 | 539.6305 | 0.00 |
| 30 | 539.9907 | 1.14 |
| 45 | 540.092 | 1.46 |
| 60 | 540.0995 | 1.48 |
| 300 | 540.0728 | 1.40 |
| 1500 | 540.0965 | 1.47 |
| 3600 | 540.2178 | 1.85 |
| 7200 | 540.2 | 1.80 |
| 14400 | 540.2478 | 1.95 |

**TMG**

| Time | centroid | D |
|---|---|---|
| 0 | 539.62 | -0.02 |
| 30 | 539.81 | 0.56 |
| 45 | 539.93 | 0.96 |
| 60 | 540.10 | 1.49 |
| 300 | 540.01 | 1.21 |
| 1500 | 540.14 | 1.60 |
| 3600 | 540.27 | 2.01 |
| 7200 | 540.38 | 2.37 |
| 14400 | 540.56 | 2.93 |

control 539.63
infinity 542.166

$$D(t) = \frac{M_t - M_0}{M_\infty - M_0} * N$$

**IPTG**

| Time | Replicate 1 | Replicate 2 | Replicate 3 | Replicate 4 | Replicate5 | average | SD |
|---|---|---|---|---|---|---|---|
| 0 | 539.627 | 539.643 | 539.626 | 539.621 | 539.627 | 539.63 | 0.01 |
| 30 | 539.808 | | 539.892 | 539.826 | 539.807 | 539.83 | 0.04 |
| 45 | 539.814 | | | | | 539.81 | |
| 60 | 539.874 | 539.844 | 540.103 | 539.89 | 539.871 | 539.92 | 0.11 |
| 300 | 539.956 | 539.846 | 540.031 | 539.934 | 539.878 | 539.93 | 0.07 |
| 1500 | 540.061 | 539.97 | 540.145 | 540.043 | 539.981 | 540.04 | 0.07 |
| 3600 | 540.137 | 540.072 | 540.176 | 540.057 | 540.112 | 540.11 | 0.05 |
| 7200 | 540.232 | 540.163 | 540.322 | 540.226 | 540.144 | 540.22 | 0.07 |
| 14400 | 540.278 | 540.205 | 540.333 | 540.306 | 540.165 | 540.26 | 0.07 |

**ONPF**

| Time | Replicate 1 | Replicate 2 | Replicate 3 | Replicate 4 | average | SD |
|---|---|---|---|---|---|---|
| 0 | 539.643 | 539.632 | 539.625 | 539.619 | 539.63 | 0.01 |
| 30 | 539.941 | 540.036 | 539.902 | 540.04 | 539.98 | 0.07 |
| 45 | 540.315 | | | | 540.32 | |
| 60 | 540.203 | 540.5 | 540.117 | 540.241 | 540.27 | 0.16 |
| 300 | 540.305 | 540.204 | | 540.21 | 540.24 | 0.06 |
| 1500 | 540.515 | 540.673 | 540.411 | 540.39 | 540.50 | 0.13 |
| 3600 | 540.712 | 540.799 | 540.461 | 540.705 | 540.67 | 0.15 |
| 7200 | 541.008 | 540.8 | 540.614 | 540.657 | 540.77 | 0.18 |
| 14400 | | 540.911 | 540.981 | 540.921 | 540.94 | 0.04 |

**APO**

| Time | Replicate 1 | Replicate 2 | Replicate 3 | Replicate 4 | Replicate 5 | average | SD |
|---|---|---|---|---|---|---|---|
| 0 | 539.595 | | 539.683 | 539.63 | 539.625 | 539.63 | 0.04 |
| 30 | 540.108 | 539.896 | 540.11 | 540.003 | 539.98 | 540.02 | 0.09 |
| 45 | 540.069 | 539.892 | | | | 539.98 | 0.13 |
| 60 | 540.23 | 540.149 | 540.372 | 540.18 | 540.17 | 540.22 | 0.09 |
| 300 | 540.324 | 539.99 | 540.329 | 540.286 | 540.212 | 540.23 | 0.14 |
| 1500 | 540.672 | 540.525 | 540.695 | 540.548 | 540.527 | 540.59 | 0.08 |
| 3600 | 540.93 | 540.589 | 540.816 | 540.688 | 540.542 | 540.71 | 0.16 |
| 7200 | 541.065 | 540.425 | 541.069 | 540.903 | 540.701 | 540.83 | 0.27 |
| 14400 | 539.625 | 540.756 | 540.962 | 540.951 | 540.961 | 540.65 | 0.58 |

**DNA**

| Time | Replicate 1 | Replicate 2 | Replicate 3 | Replicate 4 | Replicate 5 | average | SD |
|---|---|---|---|---|---|---|---|
| 0 | 539.608 | 539.667 | 539.627 | 539.631 | | 539.63 | 0.02 |
| 30 | 540.059 | 539.866 | 540.046 | 539.925 | | 539.97 | 0.09 |
| 45 | 540.211 | 539.94 | | | | 540.08 | 0.19 |
| 60 | 540.057 | 540.01 | 540.323 | 540.03 | | 540.11 | 0.15 |
| 300 | 540.101 | 539.944 | 540.211 | 539.983 | | 540.06 | 0.12 |
| 1500 | 540.153 | 540.052 | 540.249 | 540.06 | | 540.13 | 0.09 |
| 3600 | 540.41 | 539.998 | 540.325 | 540.143 | | 540.22 | 0.18 |
| 7200 | 540.451 | 540.012 | 540.371 | 540.246 | | 540.27 | 0.19 |
| 14400 | 540.575 | 539.875 | 540.449 | 540.355 | | 540.31 | 0.31 |

**ONPFDNA**

| Time | Replicate 1 | Replicate 2 | Replicate 3 | Replicate 4 | average | SD |
|---|---|---|---|---|---|---|
| 0 | 539.627 | 539.635 | 539.632 | 539.628 | 539.63 | 0.00 |
| 30 | 539.967 | | 540.146 | 539.859 | 539.99 | 0.14 |
| 45 | 540.092 | | | | 540.09 | |
| 60 | 540.054 | 540.002 | 540.328 | 540.014 | 540.10 | 0.15 |
| 300 | 540.079 | 539.952 | 540.204 | 540.056 | 540.07 | 0.10 |
| 1500 | 539.996 | 539.985 | 540.355 | 540.05 | 540.10 | 0.17 |
| 3600 | 540.445 | 540 | 540.323 | 540.103 | 540.22 | 0.20 |
| 7200 | 540.38 | 539.995 | 540.258 | 540.167 | 540.20 | 0.16 |
| 14400 | 540.127 | 540.047 | 540.571 | 540.246 | 540.25 | 0.23 |

**TMG**

| Time | Replicate 1 | Replicate 2 | Replicate 3 | Replicate 4 | Replicate 5 | average | SD |
|---|---|---|---|---|---|---|---|
| 0 | 539.63 | 539.616 | | | | 539.62 | 0.01 |
| 30 | 539.763 | 539.853 | | | | 539.81 | 0.06 |
| 45 | 539.96 | 539.907 | | | | 539.93 | 0.04 |
| 60 | 539.915 | 540.288 | | | | 540.10 | 0.26 |
| 300 | 539.987 | 540.037 | | | | 540.01 | 0.04 |
| 1500 | 540.038 | 540.234 | | | | 540.14 | 0.14 |
| 3600 | 540.254 | 540.281 | | | | 540.27 | 0.02 |
| 7200 | 540.242 | 540.518 | | | | 540.38 | 0.20 |
| 14400 | 540.464 | 540.654 | | | | 540.56 | 0.13 |

### 72-84 ALHAPSQIVAAIK

**Charge** 2

**IPTG**

| Time | centroid | D |
|---|---|---|
| 0 | 660.2685 | -0.01 |
| 30 | 660.504 | 1.38 |
| 45 | 660.4725 | 1.19 |
| 60 | 660.6385 | 2.17 |
| 300 | 660.5255 | 1.51 |
| 1500 | 660.6485 | 2.23 |
| 3600 | 660.7385 | 2.76 |
| 7200 | 660.8125 | 3.20 |
| 14400 | 660.8715 | 3.54 |

**ONPF**

| Time | centroid | D |
|---|---|---|
| 0 | 660.278 | 0.05 |
| 30 | 660.467 | 1.16 |
| 45 | 660.952 | 4.02 |
| 60 | 660.815 | 3.21 |
| 300 | 660.926 | 3.87 |
| 1500 | 661.085 | 4.80 |
| 3600 | 661.412 | 6.73 |
| 7200 | 661.308 | 6.12 |
| 14400 | 661.64 | 8.07 |

**APO**

| Time | centroid | D |
|---|---|---|
| 0 | 660.267 | -0.02 |
| 30 | 660.655 | 2.27 |
| 45 | 660.6225 | 2.08 |
| 60 | 660.735 | 2.74 |
| 300 | 660.698 | 2.52 |
| 1500 | 661.1165 | 4.99 |
| 3600 | 661.29 | 6.01 |
| 7200 | 661.372 | 6.49 |
| 14400 | 661.522 | 7.38 |

**DNA**

| Time | centroid | D |
|---|---|---|
| 0 | 660.258 | -0.07 |
| 30 | 660.585 | 1.86 |
| 45 | 660.731 | 2.72 |
| 60 | 660.652 | 2.25 |
| 300 | 660.703 | 2.55 |
| 1500 | 660.7575 | 2.87 |
| 3600 | 660.8785 | 3.59 |
| 7200 | 660.8935 | 3.67 |
| 14400 | 660.834 | 3.32 |

**ONPFDNA**

| Time | centroid | D |
|---|---|---|
| 0 | 660.2625 | -0.04 |
| 30 | 660.6295 | 2.12 |
| 45 | 660.7215 | 2.66 |
| 60 | 660.7135 | 2.61 |
| 300 | 660.678 | 2.40 |
| 1500 | 660.697 | 2.52 |
| 3600 | 660.9735 | 4.15 |
| 7200 | 660.997 | 4.28 |
| 14400 | 660.7615 | 2.90 |

**TMG**

| Time | centroid | D |
|---|---|---|
| 0 | 660.2605 | -0.06 |
| 30 | 660.5065 | 1.39 |
| 45 | 660.708 | 2.58 |
| 60 | 660.541 | 1.60 |
| 300 | 660.706 | 2.57 |
| 1500 | 660.7045 | 2.56 |
| 3600 | 661.162 | 5.26 |
| 7200 | 661.0695 | 4.71 |
| 14400 | 661.1225 | 5.02 |

control 660.27
infinity 661.967

$$D(t) = \frac{\underline{Mt - M0}}{M\infty - M0} \ast N$$

**IPTG**

| Time | Replicate 1 | Replicate 2 | Replicate 3 | Replicate 4 | Replicate5 | average | SD |
|---|---|---|---|---|---|---|---|
| 0 | 660.288 | 660.249 | | | | 660.27 | 0.03 |
| 30 | 660.504 | | | | | 660.50 | |
| 45 | 660.496 | 660.449 | | | | 660.47 | 0.03 |
| 60 | 660.588 | 660.689 | | | | 660.64 | 0.07 |
| 300 | 660.566 | 660.485 | | | | 660.53 | 0.06 |
| 1500 | 660.675 | 660.622 | | | | 660.65 | 0.04 |
| 3600 | 660.748 | 660.729 | | | | 660.74 | 0.01 |
| 7200 | 660.813 | 660.812 | | | | 660.81 | 0.00 |
| 14400 | 660.861 | 660.882 | | | | 660.87 | 0.01 |

**ONPF**

| Time | Replicate 1 | Replicate 2 | Replicate 3 | Replicate 4 | average | SD |
|---|---|---|---|---|---|---|
| 0 | 660.278 | | | | 660.28 | |
| 30 | 660.467 | | | | 660.47 | |
| 45 | 660.952 | | | | 660.95 | |
| 60 | 660.815 | | | | 660.82 | |
| 300 | 660.926 | | | | 660.93 | |
| 1500 | 661.085 | | | | 661.09 | |
| 3600 | 661.412 | | | | 661.41 | |
| 7200 | 661.308 | | | | 661.31 | |
| 14400 | 661.64 | | | | 661.64 | |

**APO**

| Time | Replicate 1 | Replicate 2 | Replicate 3 | Replicate 4 | Replicate 5 | average | SD |
|---|---|---|---|---|---|---|---|
| 0 | 660.253 | 660.281 | | | | 660.27 | 0.02 |
| 30 | 660.655 | | | | | 660.66 | |
| 45 | 660.671 | 660.574 | | | | 660.62 | 0.07 |
| 60 | 660.644 | 660.826 | | | | 660.74 | 0.13 |
| 300 | 660.711 | 660.685 | | | | 660.70 | 0.02 |
| 1500 | 661.087 | 661.146 | | | | 661.12 | 0.04 |
| 3600 | 661.352 | 661.228 | | | | 661.29 | 0.09 |
| 7200 | 661.538 | 661.206 | | | | 661.37 | 0.23 |
| 14400 | 661.628 | 661.416 | | | | 661.52 | 0.15 |

**DNA**

| Time | Replicate 1 | Replicate 2 | Replicate 3 | Replicate 4 | Replicate 5 | average | SD |
|---|---|---|---|---|---|---|---|
| 0 | 660.267 | 660.249 | | | | 660.26 | 0.01 |
| 30 | 660.588 | 660.582 | | | | 660.59 | 0.00 |
| 45 | 660.841 | 660.621 | | | | 660.73 | 0.16 |
| 60 | 660.676 | 660.628 | | | | 660.65 | 0.03 |
| 300 | 660.744 | 660.662 | | | | 660.70 | 0.06 |
| 1500 | 660.809 | 660.706 | | | | 660.76 | 0.07 |
| 3600 | 661.055 | 660.702 | | | | 660.88 | 0.25 |
| 7200 | 661.002 | 660.785 | | | | 660.89 | 0.15 |
| 14400 | 661.169 | 660.499 | | | | 660.83 | 0.47 |

**ONPFDNA**

| Time | Replicate 1 | Replicate 2 | Replicate 3 | Replicate 4 | average | SD |
|---|---|---|---|---|---|---|
| 0 | 660.27 | 660.255 | | | 660.26 | 0.01 |
| 30 | 660.681 | 660.578 | | | 660.63 | 0.07 |
| 45 | 660.737 | 660.706 | | | 660.72 | 0.02 |
| 60 | 660.68 | 660.747 | | | 660.71 | 0.05 |
| 300 | 660.728 | 660.628 | | | 660.68 | 0.07 |
| 1500 | 660.658 | 660.736 | | | 660.70 | 0.06 |
| 3600 | 661.072 | 660.875 | | | 660.97 | 0.14 |
| 7200 | 661.137 | 660.857 | | | 661.00 | 0.20 |
| 14400 | 660.725 | 660.798 | | | 660.76 | 0.05 |

**TMG**

| Time | Replicate 1 | Replicate 2 | Replicate 3 | Replicate 4 | Replicate 5 | average | SD |
|---|---|---|---|---|---|---|---|
| 0 | 660.276 | 660.245 | | | | 660.26 | 0.02 |
| 30 | 660.418 | 660.595 | | | | 660.51 | 0.13 |
| 45 | 660.612 | 660.804 | | | | 660.71 | 0.14 |
| 60 | 660.541 | | | | | 660.54 | |
| 300 | 660.614 | 660.798 | | | | 660.71 | 0.13 |
| 1500 | 660.67 | 660.739 | | | | 660.70 | 0.05 |
| 3600 | 660.914 | 661.41 | | | | 661.16 | 0.35 |
| 7200 | 660.975 | 661.164 | | | | 661.07 | 0.13 |
| 14400 | 661.121 | 661.124 | | | | 661.12 | 0.00 |

### 77-82 SQIVAA

**Charge** 1

**IPTG**

| Time | centroid | D |
|---|---|---|
| 0 | 588.65 | 0.01 |
| 30 | 588.72 | 0.10 |
| 45 | 588.74 | 0.13 |
| 60 | 588.75 | 0.14 |
| 300 | 588.76 | 0.15 |
| 1500 | 588.73 | 0.12 |
| 3600 | 588.72 | 0.11 |
| 7200 | 588.75 | 0.13 |
| 14400 | 588.75 | 0.14 |

**ONPF**

| Time | centroid | D |
|---|---|---|
| 0 | 588.77 | 0.16 |
| 30 | 588.79 | 0.19 |
| 45 | 588.92 | 0.36 |
| 60 | 588.86 | 0.28 |
| 300 | 588.86 | 0.27 |
| 1500 | 589.11 | 0.59 |
| 3600 | 589.16 | 0.66 |
| 7200 | 589.42 | 0.99 |
| 14400 | 589.70 | 1.34 |

**APO**

| Time | centroid | D |
|---|---|---|
| 0 | 588.64 | 0.01 |
| 30 | 588.72 | 0.11 |
| 45 | 588.74 | 0.13 |
| 60 | 588.83 | 0.23 |
| 300 | 588.88 | 0.31 |
| 1500 | 589.17 | 0.67 |
| 3600 | 589.34 | 0.89 |
| 7200 | 589.50 | 1.09 |
| 14400 | 589.69 | 1.33 |

**DNA**

| Time | centroid | D |
|---|---|---|
| 0 | 588.61 | -0.04 |
| 30 | 588.76 | 0.15 |
| 45 | 588.77 | 0.17 |
| 60 | 588.73 | 0.12 |
| 300 | 588.72 | 0.11 |
| 1500 | 588.79 | 0.19 |
| 3600 | 588.81 | 0.22 |
| 7200 | 588.82 | 0.23 |
| 14400 | 588.87 | 0.29 |

**ONPFDNA**

| Time | centroid | D |
|---|---|---|
| 0 | 588.67 | 0.04 |
| 30 | 588.67 | 0.03 |
| 45 | 588.68 | 0.04 |
| 60 | 588.72 | 0.10 |
| 300 | | |
| 1500 | 588.82 | 0.23 |
| 3600 | 588.74 | 0.12 |
| 7200 | 588.74 | 0.13 |
| 14400 | 588.78 | 0.17 |

**TMG**

| Time | centroid | D |
|---|---|---|
| 0 | 588.68 | 0.05 |
| 30 | 588.76 | 0.15 |
| 45 | 588.80 | 0.20 |
| 60 | 588.89 | 0.32 |
| 300 | 588.77 | 0.16 |
| 1500 | 588.82 | 0.23 |
| 3600 | 588.83 | 0.24 |
| 7200 | 588.83 | 0.24 |
| 14400 | 588.94 | 0.38 |

| | |
|---|---|
| control | 588.64 |
| infinity | 591.793 |

$$D(t) = \frac{M_t - M_0}{M_\infty - M_0} * N$$

**IPTG**

| Time | Replicate 1 | Replicate 2 | Replicate 3 | Replicate 4 | Replicate5 | average | SD |
|---|---|---|---|---|---|---|---|
| 0 | 588.665 | 588.637 | | | | 588.65 | 0.02 |
| 30 | 588.742 | 588.7 | | | | 588.72 | 0.03 |
| 45 | 588.749 | 588.736 | | | | 588.74 | 0.01 |
| 60 | 588.78 | 588.718 | | | | 588.75 | 0.04 |
| 300 | 588.799 | 588.711 | | | | 588.76 | 0.06 |
| 1500 | 588.738 | 588.726 | | | | 588.73 | 0.01 |
| 3600 | 588.74 | 588.708 | | | | 588.72 | 0.02 |
| 7200 | 588.748 | 588.744 | | | | 588.75 | 0.00 |
| 14400 | 588.771 | 588.727 | | | | 588.75 | 0.03 |

**ONPF**

| Time | Replicate 1 | Replicate 2 | Replicate 3 | Replicate 4 | average | SD |
|---|---|---|---|---|---|---|
| 0 | 588.713 | 588.823 | | | 588.77 | 0.08 |
| 30 | 588.79 | | | | 588.79 | |
| 45 | 588.923 | | | | 588.92 | |
| 60 | 588.904 | 588.812 | | | 588.86 | 0.07 |
| 300 | 588.855 | | | | 588.86 | |
| 1500 | 589.108 | | | | 589.11 | |
| 3600 | 589.358 | 588.961 | | | 589.16 | 0.28 |
| 7200 | 589.547 | 589.291 | | | 589.42 | 0.18 |
| 14400 | 589.697 | | | | 589.70 | |

**APO**

| Time | Replicate 1 | Replicate 2 | Replicate 3 | Replicate 4 | Replicate 5 | average | SD |
|---|---|---|---|---|---|---|---|
| 0 | 588.648 | 588.641 | | | | 588.64 | 0.00 |
| 30 | 588.737 | 588.71 | | | | 588.72 | 0.02 |
| 45 | 588.75 | 588.735 | | | | 588.74 | 0.01 |
| 60 | 588.764 | 588.886 | | | | 588.83 | 0.09 |
| 300 | 588.818 | 588.948 | | | | 588.88 | 0.09 |
| 1500 | 589.053 | 589.288 | | | | 589.17 | 0.17 |
| 3600 | 589.337 | 589.342 | | | | 589.34 | 0.00 |
| 7200 | 589.704 | 589.294 | | | | 589.50 | 0.29 |
| 14400 | 589.62 | 589.755 | | | | 589.69 | 0.10 |

**DNA**

| Time | Replicate 1 | Replicate 2 | Replicate 3 | Replicate 4 | Replicate 5 | average | SD |
|---|---|---|---|---|---|---|---|
| 0 | 588.622 | 588.6 | | | | 588.61 | 0.02 |
| 30 | 588.788 | 588.732 | | | | 588.76 | 0.04 |
| 45 | 588.801 | 588.748 | | | | 588.77 | 0.04 |
| 60 | 588.767 | 588.699 | | | | 588.73 | 0.05 |
| 300 | 588.734 | 588.713 | | | | 588.72 | 0.01 |
| 1500 | 588.855 | 588.725 | | | | 588.79 | 0.09 |
| 3600 | 588.821 | 588.806 | | | | 588.81 | 0.01 |
| 7200 | 588.881 | 588.765 | | | | 588.82 | 0.08 |
| 14400 | 589.023 | 588.709 | | | | 588.87 | 0.22 |

**ONPFDNA**

| Time | Replicate 1 | Replicate 2 | Replicate 3 | Replicate 4 | average | SD |
|---|---|---|---|---|---|---|
| 0 | 588.707 | 588.634 | | | 588.67 | 0.05 |
| 30 | 588.647 | 588.687 | | | 588.67 | 0.03 |
| 45 | 588.666 | 588.684 | | | 588.68 | 0.01 |
| 60 | 588.74 | 588.702 | | | 588.72 | 0.03 |
| 300 | | | | | | |
| 1500 | 588.923 | 588.715 | | | 588.82 | 0.15 |
| 3600 | 588.757 | 588.714 | | | 588.74 | 0.03 |
| 7200 | 588.732 | 588.754 | | | 588.74 | 0.02 |
| 14400 | 588.795 | 588.757 | | | 588.78 | 0.03 |

**TMG**

| Time | Replicate 1 | Replicate 2 | Replicate 3 | Replicate 4 | Replicate 5 | average | SD |
|---|---|---|---|---|---|---|---|
| 0 | 588.704 | 588.65 | | | | 588.68 | 0.04 |
| 30 | 588.78 | 588.736 | | | | 588.76 | 0.03 |
| 45 | 588.842 | 588.758 | | | | 588.80 | 0.06 |
| 60 | 588.825 | 588.96 | | | | 588.89 | 0.10 |
| 300 | | 588.768 | | | | 588.77 | |
| 1500 | 588.78 | 588.86 | | | | 588.82 | 0.06 |
| 3600 | 588.896 | 588.755 | | | | 588.83 | 0.10 |
| 7200 | 588.859 | 588.796 | | | | 588.83 | 0.04 |
| 14400 | 588.93 | 588.955 | | | | 588.94 | 0.02 |

### 77-84 SQIVAAIK

**Charge** 1

**IPTG**

| Time | centroid | D |
|---|---|---|
| 0 | 829.99 | 0.01 |
| 30 | 830.084 | 0.12 |
| 45 | 830.092 | 0.13 |
| 60 | 830.1165 | 0.16 |
| 300 | 830.129 | 0.18 |
| 1500 | 830.1035 | 0.15 |
| 3600 | 830.0915 | 0.13 |
| 7200 | 830.1125 | 0.16 |
| 14400 | 830.1235 | 0.17 |

**ONPF**

| Time | centroid | D |
|---|---|---|
| 0 | 829.9955 | 0.02 |
| 30 | 830.07 | 0.11 |
| 45 | 830.111 | 0.16 |
| 60 | 830.247 | 0.32 |
| 300 | 830.191 | 0.25 |
| 1500 | 830.3705 | 0.47 |
| 3600 | 830.5665 | 0.70 |
| 7200 | 830.7195 | 0.88 |
| 14400 | 831.10 | 1.33 |

**APO**

| Time | centroid | D |
|---|---|---|
| 0 | 829.98 | 0.00 |
| 30 | 830.0455 | 0.08 |
| 45 | 830.096 | 0.14 |
| 60 | 830.2005 | 0.26 |
| 300 | 830.163 | 0.22 |
| 1500 | 830.521 | 0.64 |
| 3600 | 830.693 | 0.85 |
| 7200 | 830.9315 | 1.13 |
| 14400 | 831.1015 | 1.34 |

**DNA**

| Time | centroid | D |
|---|---|---|
| 0 | 829.963 | -0.02 |
| 30 | 830.0655 | 0.10 |
| 45 | 830.06 | 0.10 |
| 60 | 830.1155 | 0.16 |
| 300 | 830.113 | 0.16 |
| 1500 | 830.1225 | 0.17 |
| 3600 | 830.2015 | 0.26 |
| 7200 | 830.2135 | 0.28 |
| 14400 | 830.2695 | 0.34 |

**ONPFDNA**

| Time | centroid | D |
|---|---|---|
| 0 | 829.98 | 0.00 |
| 30 | 830.074 | 0.11 |
| 45 | 830.111 | 0.16 |
| 60 | 830.105 | 0.15 |
| 300 | 830.095 | 0.14 |
| 1500 | 830.13 | 0.18 |
| 3600 | 830.166 | 0.22 |
| 7200 | 830.1975 | 0.26 |
| 14400 | 830.178 | 0.24 |

**TMG**

| Time | centroid | D |
|---|---|---|
| 0 | 829.973 | -0.01 |
| 30 | 830.0785 | 0.12 |
| 45 | 830.1405 | 0.19 |
| 60 | 830.181 | 0.24 |
| 300 | 830.126 | 0.17 |
| 1500 | 830.146 | 0.20 |
| 3600 | 830.15 | 0.20 |
| 7200 | 830.15 | 0.20 |
| 14400 | 830.2695 | 0.34 |

control 829.98
infinity 835.018

$$D(t) = \frac{M_t - M_0}{M_\infty - M_0} *N$$

**IPTG**

| Time | Replicate 1 | Replicate 2 | Replicate 3 | Replicate 4 | Replicate5 | average | SD |
|---|---|---|---|---|---|---|---|
| 0 | 830.001 | 829.979 | | | | 829.99 | 0.02 |
| 30 | 830.111 | 830.057 | | | | 830.08 | 0.04 |
| 45 | 830.118 | 830.066 | | | | 830.09 | 0.04 |
| 60 | 830.135 | 830.098 | | | | 830.12 | 0.03 |
| 300 | 830.182 | 830.076 | | | | 830.13 | 0.07 |
| 1500 | 830.11 | 830.097 | | | | 830.10 | 0.01 |
| 3600 | 830.123 | 830.06 | | | | 830.09 | 0.04 |
| 7200 | 830.112 | 830.113 | | | | 830.11 | 0.00 |
| 14400 | 830.156 | 830.091 | | | | 830.12 | 0.05 |

**ONPF**

| Time | Replicate 1 | Replicate 2 | Replicate 3 | Replicate 4 | average | SD |
|---|---|---|---|---|---|---|
| 0 | 829.993 | 829.998 | | | 830.00 | 0.00 |
| 30 | 830.083 | 830.057 | | | 830.07 | 0.02 |
| 45 | | 830.111 | | | 830.11 | |
| 60 | 830.245 | 830.249 | | | 830.25 | 0.00 |
| 300 | 830.244 | 830.138 | | | 830.19 | 0.07 |
| 1500 | 830.419 | 830.322 | | | 830.37 | 0.07 |
| 3600 | 830.812 | 830.321 | | | 830.57 | 0.35 |
| 7200 | 830.829 | 830.61 | | | 830.72 | 0.15 |
| 14400 | 831.098 | | | | 831.10 | |

**APO**

| Time | Replicate 1 | Replicate 2 | Replicate 3 | Replicate 4 | Replicate 5 | average | SD |
|---|---|---|---|---|---|---|---|
| 0 | 829.974 | 829.986 | | | | 829.98 | 0.01 |
| 30 | 830.03 | 830.061 | | | | 830.05 | 0.02 |
| 45 | 830.109 | 830.083 | | | | 830.10 | 0.02 |
| 60 | 830.102 | 830.299 | | | | 830.20 | 0.14 |
| 300 | 830.156 | 830.17 | | | | 830.16 | 0.01 |
| 1500 | 830.363 | 830.679 | | | | 830.52 | 0.22 |
| 3600 | 830.683 | 830.703 | | | | 830.69 | 0.01 |
| 7200 | 831.049 | 830.814 | | | | 830.93 | 0.17 |
| 14400 | 831.067 | 831.136 | | | | 831.10 | 0.05 |

**DNA**

| Time | Replicate 1 | Replicate 2 | Replicate 3 | Replicate 4 | Replicate 5 | average | SD |
|---|---|---|---|---|---|---|---|
| 0 | 829.949 | 829.977 | | | | 829.96 | 0.02 |
| 30 | 830.094 | 830.037 | | | | 830.07 | 0.04 |
| 45 | | 830.06 | | | | 830.06 | |
| 60 | 830.126 | 830.105 | | | | 830.12 | 0.01 |
| 300 | 830.148 | 830.078 | | | | 830.11 | 0.05 |
| 1500 | 830.125 | 830.12 | | | | 830.12 | 0.00 |
| 3600 | 830.274 | 830.129 | | | | 830.20 | 0.10 |
| 7200 | 830.27 | 830.157 | | | | 830.21 | 0.08 |
| 14400 | 830.438 | 830.101 | | | | 830.27 | 0.24 |

**ONPFDNA**

| Time | Replicate 1 | Replicate 2 | Replicate 3 | Replicate 4 | average | SD |
|---|---|---|---|---|---|---|
| 0 | 829.982 | 829.978 | | | 829.98 | 0.00 |
| 30 | 830.091 | 830.057 | | | 830.07 | 0.02 |
| 45 | 830.14 | 830.082 | | | 830.11 | 0.04 |
| 60 | 830.116 | 830.094 | | | 830.11 | 0.02 |
| 300 | 830.101 | 830.089 | | | 830.10 | 0.01 |
| 1500 | 830.115 | 830.145 | | | 830.13 | 0.02 |
| 3600 | 830.264 | 830.068 | | | 830.17 | 0.14 |
| 7200 | 830.279 | 830.116 | | | 830.20 | 0.12 |
| 14400 | 830.218 | 830.138 | | | 830.18 | 0.06 |

**TMG**

| Time | Replicate 1 | Replicate 2 | Replicate 3 | Replicate 4 | Replicate 5 | average | SD |
|---|---|---|---|---|---|---|---|
| 0 | 829.981 | 829.965 | | | | 829.97 | 0.01 |
| 30 | 830.061 | 830.096 | | | | 830.08 | 0.02 |
| 45 | 830.168 | 830.113 | | | | 830.14 | 0.04 |
| 60 | 830.111 | 830.251 | | | | 830.18 | 0.10 |
| 300 | 830.122 | 830.13 | | | | 830.13 | 0.01 |
| 1500 | 830.107 | 830.185 | | | | 830.15 | 0.06 |
| 3600 | 830.183 | 830.117 | | | | 830.15 | 0.05 |
| 7200 | 830.158 | 830.142 | | | | 830.15 | 0.01 |
| 14400 | 830.225 | 830.314 | | | | 830.27 | 0.06 |

### 79-84 IVAAIK

**Charge** 1

**IPTG**

| Time | centroid | D |
|---|---|---|
| 0 | 614.76 | 0.01 |
| 30 | 614.84 | 0.10 |
| 45 | 614.84 | 0.10 |
| 60 | 614.84 | 0.10 |
| 300 | 614.84 | 0.11 |
| 1500 | 614.83 | 0.09 |
| 3600 | 614.83 | 0.09 |
| 7200 | 614.83 | 0.10 |
| 14400 | 614.84 | 0.11 |

**ONPF**

| Time | centroid | D |
|---|---|---|
| 0 | 614.77 | 0.02 |
| 30 | 614.83 | 0.09 |
| 45 | 614.94 | 0.21 |
| 60 | 614.90 | 0.18 |
| 300 | 614.88 | 0.15 |
| 1500 | 614.97 | 0.26 |
| 3600 | 615.07 | 0.37 |
| 7200 | 615.16 | 0.46 |
| 14400 | 615.15 | 0.45 |

**APO**

| Time | centroid | D |
|---|---|---|
| 0 | 614.75 | 0.00 |
| 30 | 614.81 | 0.07 |
| 45 | 614.83 | 0.10 |
| 60 | 614.91 | 0.19 |
| 300 | 614.86 | 0.13 |
| 1500 | 615.08 | 0.38 |
| 3600 | 615.21 | 0.52 |
| 7200 | 615.33 | 0.67 |
| 14400 | 615.39 | 0.73 |

**DNA**

| Time | centroid | D |
|---|---|---|
| 0 | 614.75 | 0.00 |
| 30 | 614.82 | 0.08 |
| 45 | 614.87 | 0.14 |
| 60 | 614.85 | 0.12 |
| 300 | 614.84 | 0.10 |
| 1500 | 614.86 | 0.12 |
| 3600 | 614.89 | 0.16 |
| 7200 | 614.91 | 0.18 |
| 14400 | 614.93 | 0.20 |

**ONPFDNA**

| Time | centroid | D |
|---|---|---|
| 0 | 614.77 | 0.02 |
| 30 | 614.82 | 0.08 |
| 45 | 614.84 | 0.11 |
| 60 | 614.84 | 0.10 |
| 300 | 614.83 | 0.10 |
| 1500 | 614.85 | 0.11 |
| 3600 | 614.86 | 0.13 |
| 7200 | 614.88 | 0.14 |
| 14400 | 614.90 | 0.17 |

**TMG**

| Time | centroid | D |
|---|---|---|
| 0 | 614.76 | 0.01 |
| 30 | 614.82 | 0.08 |
| 45 | 614.85 | 0.12 |
| 60 | 614.88 | 0.15 |
| 300 | 614.85 | 0.11 |
| 1500 | 614.86 | 0.13 |
| 3600 | 614.85 | 0.12 |
| 7200 | 614.83 | 0.09 |
| 14400 | 614.95 | 0.23 |

| | |
|---|---|
| control | 614.75 |
| infinity | 618.244 |

$$D(t) = \frac{M_t - M_0}{M_\infty - M_0} *N$$

**IPTG**

| Time | Replicate 1 | Replicate 2 | Replicate 3 | Replicate 4 | Replicate5 | average | SD |
|---|---|---|---|---|---|---|---|
| 0 | 614.761 | 614.761 | | | | 614.76 | 0.00 |
| 30 | 614.86 | 614.812 | | | | 614.84 | 0.03 |
| 45 | 614.841 | 614.829 | | | | 614.84 | 0.01 |
| 60 | 614.856 | 614.823 | | | | 614.84 | 0.02 |
| 300 | 614.86 | 614.824 | | | | 614.84 | 0.03 |
| 1500 | 614.828 | 614.835 | | | | 614.83 | 0.00 |
| 3600 | 614.836 | 614.819 | | | | 614.83 | 0.01 |
| 7200 | 614.828 | 614.839 | | | | 614.83 | 0.01 |
| 14400 | 614.856 | 614.832 | | | | 614.84 | 0.02 |

**ONPF**

| Time | Replicate 1 | Replicate 2 | Replicate 3 | Replicate 4 | average | SD |
|---|---|---|---|---|---|---|
| 0 | 614.77 | 614.773 | | | 614.77 | 0.00 |
| 30 | 614.836 | 614.818 | | | 614.83 | 0.01 |
| 45 | 615.034 | 614.836 | | | 614.94 | 0.14 |
| 60 | 614.91 | 614.898 | | | 614.90 | 0.01 |
| 300 | 614.907 | 614.85 | | | 614.88 | 0.04 |
| 1500 | 615.002 | 614.944 | | | 614.97 | 0.04 |
| 3600 | 615.186 | 614.962 | | | 615.07 | 0.16 |
| 7200 | 615.206 | 615.105 | | | 615.16 | 0.07 |
| 14400 | 615.444 | 614.846 | | | 615.15 | 0.42 |

**APO**

| Time | Replicate 1 | Replicate 2 | Replicate 3 | Replicate 4 | Replicate 5 | average | SD |
|---|---|---|---|---|---|---|---|
| 0 | 614.74 | 614.764 | | | | 614.75 | 0.02 |
| 30 | 614.805 | 614.811 | | | | 614.81 | 0.00 |
| 45 | 614.834 | 614.833 | | | | 614.83 | 0.00 |
| 60 | 614.825 | 615.001 | | | | 614.91 | 0.12 |
| 300 | 614.855 | 614.874 | | | | 614.86 | 0.01 |
| 1500 | 614.969 | 615.188 | | | | 615.08 | 0.15 |
| 3600 | 615.17 | 615.244 | | | | 615.21 | 0.05 |
| 7200 | 615.4 | 615.268 | | | | 615.33 | 0.09 |
| 14400 | 615.255 | 615.527 | | | | 615.39 | 0.19 |

**DNA**

| Time | Replicate 1 | Replicate 2 | Replicate 3 | Replicate 4 | Replicate 5 | average | SD |
|---|---|---|---|---|---|---|---|
| 0 | 614.733 | 614.76 | | | | 614.75 | 0.02 |
| 30 | 614.833 | 614.803 | | | | 614.82 | 0.02 |
| 45 | 614.923 | 614.818 | | | | 614.87 | 0.07 |
| 60 | 614.855 | 614.848 | | | | 614.85 | 0.00 |
| 300 | 614.851 | 614.821 | | | | 614.84 | 0.02 |
| 1500 | 614.861 | 614.853 | | | | 614.86 | 0.01 |
| 3600 | 614.927 | 614.852 | | | | 614.89 | 0.05 |
| 7200 | 614.928 | 614.889 | | | | 614.91 | 0.03 |
| 14400 | 615.027 | 614.829 | | | | 614.93 | 0.14 |

**ONPFDNA**

| Time | Replicate 1 | Replicate 2 | Replicate 3 | Replicate 4 | average | SD |
|---|---|---|---|---|---|---|
| 0 | 614.757 | 614.773 | | | 614.77 | 0.01 |
| 30 | 614.825 | 614.812 | | | 614.82 | 0.01 |
| 45 | 614.859 | 614.825 | | | 614.84 | 0.02 |
| 60 | 614.847 | 614.833 | | | 614.84 | 0.01 |
| 300 | 614.838 | 614.831 | | | 614.83 | 0.00 |
| 1500 | 614.84 | 614.855 | | | 614.85 | 0.01 |
| 3600 | 614.893 | 614.834 | | | 614.86 | 0.04 |
| 7200 | 614.922 | 614.828 | | | 614.88 | 0.07 |
| 14400 | 614.916 | 614.883 | | | 614.90 | 0.02 |

**TMG**

| Time | Replicate 1 | Replicate 2 | Replicate 3 | Replicate 4 | Replicate 5 | average | SD |
|---|---|---|---|---|---|---|---|
| 0 | 614.768 | 614.748 | | | | 614.76 | 0.01 |
| 30 | 614.809 | 614.827 | | | | 614.82 | 0.01 |
| 45 | 614.866 | 614.835 | | | | 614.85 | 0.02 |
| 60 | 614.834 | 614.931 | | | | 614.88 | 0.07 |
| 300 | 614.841 | 614.855 | | | | 614.85 | 0.01 |
| 1500 | 614.832 | 614.889 | | | | 614.86 | 0.04 |
| 3600 | 614.875 | 614.828 | | | | 614.85 | 0.03 |
| 7200 | 614.854 | 614.809 | | | | 614.83 | 0.03 |
| 14400 | 614.932 | 614.965 | | | | 614.95 | 0.02 |

### 82-93 AIKSRADQLGAS

**Charge** 2

**IPTG**

| Time | centroid | D |
|---|---|---|
| 0 | 609.1812 | -0.01 |
| 30 | 609.5616 | 1.29 |
| 45 | 609.651 | 1.59 |
| 60 | 609.7014 | 1.76 |
| 300 | 609.6928 | 1.73 |
| 1500 | 609.936 | 2.56 |
| 3600 | 610.0612 | 2.99 |
| 7200 | 610.2114 | 3.50 |
| 14400 | 610.27 | 3.70 |

**ONPF**

| Time | centroid | D |
|---|---|---|
| 0 | 609.1915 | 0.03 |
| 30 | 609.452 | 0.91 |
| 45 | 609.6 | 1.42 |
| 60 | 609.6 | 1.42 |
| 300 | 609.6177 | 1.48 |
| 1500 | 609.825 | 2.18 |
| 3600 | 609.9018 | 2.44 |
| 7200 | 609.9843 | 2.72 |
| 14400 | 610.20 | 3.45 |

**APO**

| Time | centroid | D |
|---|---|---|
| 0 | 609.1836 | 0.00 |
| 30 | 609.5474 | 1.24 |
| 45 | 609.5615 | 1.29 |
| 60 | 609.6413 | 1.56 |
| 300 | 609.6488 | 1.58 |
| 1500 | 609.874 | 2.35 |
| 3600 | 610.0712 | 3.02 |
| 7200 | 610.2108 | 3.49 |
| 14400 | 610.4718 | 4.38 |

**DNA**

| Time | centroid | D |
|---|---|---|
| 0 | 609.1773 | -0.02 |
| 30 | 609.5527 | 1.26 |
| 45 | 609.6475 | 1.58 |
| 60 | 609.7233 | 1.84 |
| 300 | 609.7193 | 1.82 |
| 1500 | 609.8813 | 2.37 |
| 3600 | 609.869 | 2.33 |
| 7200 | 609.9237 | 2.52 |
| 14400 | 609.949 | 2.60 |

**ONPFDNA**

| Time | centroid | D |
|---|---|---|
| 0 | 609.1755 | -0.03 |
| 30 | 609.5128 | 1.12 |
| 45 | 609.584 | 1.36 |
| 60 | 609.6305 | 1.52 |
| 300 | 609.6445 | 1.57 |
| 1500 | 609.7778 | 2.02 |
| 3600 | 609.9385 | 2.57 |
| 7200 | 609.8933 | 2.41 |
| 14400 | 609.948 | 2.60 |

**TMG**

| Time | centroid | D |
|---|---|---|
| 0 | 609.1835 | 0.00 |
| 30 | 609.5505 | 1.25 |
| 45 | 609.7125 | 1.80 |
| 60 | 609.6545 | 1.60 |
| 300 | 609.718 | 1.82 |
| 1500 | 609.846 | 2.25 |
| 3600 | 610.1915 | 3.43 |
| 7200 | 610.2625 | 3.67 |
| 14400 | 610.4825 | 4.42 |

control 609.1836
infinity 612.123

$$D(t) = \frac{Mt - M0}{M\infty - M0} *N$$

**IPTG**

| Time | Replicate 1 | Replicate 2 | Replicate 3 | Replicate 4 | Replicate5 | average | SD |
|---|---|---|---|---|---|---|---|
| 0 | 609.193 | 609.172 | 609.186 | 609.176 | 609.179 | 609.18 | 0.01 |
| 30 | 609.652 | 609.483 | 609.655 | 609.54 | 609.478 | 609.56 | 0.09 |
| 45 | 609.726 | 609.576 | | | | 609.65 | 0.11 |
| 60 | 609.734 | 609.58 | 609.957 | 609.636 | 609.6 | 609.70 | 0.15 |
| 300 | 609.756 | 609.597 | 609.836 | 609.707 | 609.568 | 609.69 | 0.11 |
| 1500 | 610.139 | 609.828 | 610.013 | 609.907 | 609.793 | 609.94 | 0.14 |
| 3600 | 610.371 | 610.092 | 610.025 | 609.976 | 609.842 | 610.06 | 0.20 |
| 7200 | 610.611 | 610.208 | 610.238 | 610.091 | 609.909 | 610.21 | 0.26 |
| 14400 | 610.704 | 610.235 | 610.093 | 610.352 | 609.966 | 610.27 | 0.28 |

**ONPF**

| Time | Replicate 1 | Replicate 2 | Replicate 3 | Replicate 4 | average | SD |
|---|---|---|---|---|---|---|
| 0 | 609.205 | 609.205 | 609.179 | 609.177 | 609.19 | 0.02 |
| 30 | 609.415 | 609.37 | 609.574 | 609.449 | 609.45 | 0.09 |
| 45 | 609.763 | 609.437 | | | 609.60 | 0.23 |
| 60 | 609.626 | 609.588 | | 609.586 | 609.60 | 0.02 |
| 300 | 609.664 | | 609.65 | 609.539 | 609.62 | 0.07 |
| 1500 | 609.819 | 609.665 | 610.044 | 609.772 | 609.83 | 0.16 |
| 3600 | 610.02 | 609.715 | 610.064 | 609.808 | 609.90 | 0.17 |
| 7200 | 609.963 | 609.835 | 610.266 | 609.873 | 609.98 | 0.20 |
| 14400 | 610.378 | 609.842 | 610.363 | 610.203 | 610.20 | 0.25 |

**APO**

| Time | Replicate 1 | Replicate 2 | Replicate 3 | Replicate 4 | Replicate 5 | average | SD |
|---|---|---|---|---|---|---|---|
| 0 | 609.167 | 609.175 | 609.226 | 609.181 | 609.169 | 609.18 | 0.02 |
| 30 | 609.568 | 609.432 | 609.792 | 609.494 | 609.451 | 609.55 | 0.15 |
| 45 | 609.684 | 609.439 | | | | 609.56 | 0.17 |
| 60 | 609.663 | 609.629 | | 609.572 | 609.701 | 609.64 | 0.05 |
| 300 | 609.639 | 609.494 | 609.912 | 609.632 | 609.567 | 609.65 | 0.16 |
| 1500 | 609.856 | 609.886 | 610.115 | 609.746 | 609.767 | 609.87 | 0.15 |
| 3600 | 610.239 | 609.995 | 610.32 | 609.946 | 609.856 | 610.07 | 0.20 |
| 7200 | 610.384 | 609.987 | 610.558 | 610.153 | 609.972 | 610.21 | 0.26 |
| 14400 | 610.841 | 610.342 | 610.607 | 610.289 | 610.28 | 610.47 | 0.25 |

**DNA**

| Time | Replicate 1 | Replicate 2 | Replicate 3 | Replicate 4 | Replicate 5 | average | SD |
|---|---|---|---|---|---|---|---|
| 0 | 609.174 | 609.175 | 609.183 | | | 609.18 | 0.00 |
| 30 | 609.545 | 609.459 | 609.654 | | | 609.55 | 0.10 |
| 45 | 609.785 | 609.51 | | | | 609.65 | 0.19 |
| 60 | 609.643 | 609.619 | 609.908 | | | 609.72 | 0.16 |
| 300 | 609.689 | 609.586 | 609.883 | | | 609.72 | 0.15 |
| 1500 | 609.813 | 609.824 | 610.007 | | | 609.88 | 0.11 |
| 3600 | 610.029 | 609.702 | 609.876 | | | 609.87 | 0.16 |
| 7200 | 610.033 | 609.746 | 609.992 | | | 609.92 | 0.16 |
| 14400 | 610.217 | 609.49 | 610.14 | | | 609.95 | 0.40 |

**ONPFDNA**

| Time | Replicate 1 | Replicate 2 | Replicate 3 | Replicate 4 | average | SD |
|---|---|---|---|---|---|---|
| 0 | 609.177 | 609.174 | 609.18 | 609.171 | 609.18 | 0.00 |
| 30 | 609.487 | 609.441 | 609.715 | 609.408 | 609.51 | 0.14 |
| 45 | 609.627 | 609.541 | | | 609.58 | 0.06 |
| 60 | 609.599 | 609.637 | 609.702 | 609.584 | 609.63 | 0.05 |
| 300 | 609.625 | 609.557 | 609.807 | 609.589 | 609.64 | 0.11 |
| 1500 | 609.599 | 609.707 | 610.092 | 609.713 | 609.78 | 0.22 |
| 3600 | 610.016 | 609.876 | 610.067 | 609.795 | 609.94 | 0.13 |
| 7200 | 609.973 | 609.831 | | 609.876 | 609.89 | 0.07 |
| 14400 | 609.71 | 609.801 | 610.334 | 609.947 | 609.95 | 0.28 |

**TMG**

| Time | Replicate 1 | Replicate 2 | Replicate 3 | Replicate 4 | Replicate 5 | average | SD |
|---|---|---|---|---|---|---|---|
| 0 | 609.19 | 609.177 | | | | 609.18 | 0.01 |
| 30 | 609.442 | 609.659 | | | | 609.55 | 0.15 |
| 45 | 609.722 | 609.703 | | | | 609.71 | 0.01 |
| 60 | 609.621 | 609.688 | | | | 609.65 | 0.05 |
| 300 | 609.673 | 609.763 | | | | 609.72 | 0.06 |
| 1500 | 609.754 | 609.938 | | | | 609.85 | 0.13 |
| 3600 | 610.072 | 610.311 | | | | 610.19 | 0.17 |
| 7200 | 610.038 | 610.487 | | | | 610.26 | 0.32 |
| 14400 | 610.343 | 610.622 | | | | 610.48 | 0.20 |

### 87-93 ADQLGAS

**Charge** 1

**IPTG**

| Time | centroid | D |
|---|---|---|
| 0 | 661.60 | -0.06 |
| 30 | 662.38 | 1.36 |
| 45 | 662.39 | 1.39 |
| 60 | 662.32 | 1.25 |
| 300 | 662.44 | 1.48 |
| 1500 | 662.65 | 1.85 |
| 3600 | 663.77 | 3.90 |
| 7200 | 663.43 | 3.28 |
| 14400 | 663.32 | 3.07 |

**ONPF**

| Time | centroid | D |
|---|---|---|
| 0 | 661.60 | -0.06 |
| 30 | 662.02 | 0.72 |
| 45 | 662.08 | 0.83 |
| 60 | 662.29 | 1.20 |
| 300 | 662.17 | 0.99 |
| 1500 | 662.65 | 1.87 |
| 3600 | 662.68 | 1.92 |
| 7200 | 662.66 | 1.87 |
| 14400 | #DIV/0! | #DIV/0! |

**APO**

| Time | centroid | D |
|---|---|---|
| 0 | 661.63 | 0.00 |
| 30 | 662.15 | 0.94 |
| 45 | 662.22 | 1.07 |
| 60 | 662.38 | 1.38 |
| 300 | 662.22 | 1.07 |
| 1500 | 662.83 | 2.19 |
| 3600 | 663.00 | 2.51 |
| 7200 | 663.14 | 2.76 |
| 14400 | 663.62 | 3.62 |

**DNA**

| Time | centroid | D |
|---|---|---|
| 0 | 661.60 | -0.05 |
| 30 | 662.16 | 0.96 |
| 45 | 662.34 | 1.29 |
| 60 | 662.46 | 1.51 |
| 300 | 662.28 | 1.18 |
| 1500 | 662.65 | 1.86 |
| 3600 | 662.62 | 1.80 |
| 7200 | 662.70 | 1.96 |
| 14400 | 662.60 | 1.77 |

**ONPFDNA**

| Time | centroid | D |
|---|---|---|
| 0 | 661.57 | -0.11 |
| 30 | 662.04 | 0.75 |
| 45 | 662.17 | 0.99 |
| 60 | 662.35 | 1.31 |
| 300 | 662.39 | 1.38 |
| 1500 | 662.44 | 1.48 |
| 3600 | 662.70 | 1.95 |
| 7200 | 662.38 | 1.38 |
| 14400 | 662.23 | 1.10 |

**TMG**

| Time | centroid | D |
|---|---|---|
| 0 | 661.67 | 0.07 |
| 30 | 662.36 | 1.33 |
| 45 | 662.57 | 1.71 |
| 60 | 662.36 | 1.32 |
| 300 | 662.58 | 1.74 |
| 1500 | 662.84 | 2.21 |
| 3600 | 663.37 | 3.18 |
| 7200 | 663.32 | 3.09 |
| 14400 | 663.50 | 3.42 |

control 661.63
infinity 664.37

$$D(t) = \frac{M_t - M_0}{M_\infty - M_0} * N$$

**IPTG**

| Time | Replicate 1 | Replicate 2 | Replicate 3 | Replicate 4 | Replicate5 | average | SD |
|---|---|---|---|---|---|---|---|
| 0 | 661.568 | 661.624 | | | | 661.60 | 0.04 |
| 30 | 662.375 | | | | | 662.38 | |
| 45 | 662.451 | 662.335 | | | | 662.39 | 0.08 |
| 60 | | 662.317 | | | | 662.32 | |
| 300 | 662.527 | 662.36 | | | | 662.44 | 0.12 |
| 1500 | | 662.646 | | | | 662.65 | |
| 3600 | 663.441 | 664.091 | | | | 663.77 | 0.46 |
| 7200 | 663.628 | 663.223 | | | | 663.43 | 0.29 |
| 14400 | 663.551 | 663.079 | | | | 663.32 | 0.33 |

**ONPF**

| Time | Replicate 1 | Replicate 2 | Replicate 3 | Replicate 4 | average | SD |
|---|---|---|---|---|---|---|
| 0 | 661.48 | 661.711 | | | 661.60 | 0.16 |
| 30 | | 662.022 | | | 662.02 | |
| 45 | 662.097 | 662.072 | | | 662.08 | 0.02 |
| 60 | 662.247 | 662.326 | | | 662.29 | 0.06 |
| 300 | | 662.171 | | | 662.17 | |
| 1500 | 662.878 | 662.431 | | | 662.65 | 0.32 |
| 3600 | 662.881 | 662.488 | | | 662.68 | 0.28 |
| 7200 | | 662.656 | | | 662.66 | |
| 14400 | | | | | | |

**APO**

| Time | Replicate 1 | Replicate 2 | Replicate 3 | Replicate 4 | Replicate 5 | average | SD |
|---|---|---|---|---|---|---|---|
| 0 | 661.634 | 661.63 | | | | 661.63 | 0.00 |
| 30 | 662.169 | 662.124 | | | | 662.15 | 0.03 |
| 45 | 662.384 | 662.048 | | | | 662.22 | 0.24 |
| 60 | 662.41 | 662.359 | | | | 662.38 | 0.04 |
| 300 | 662.239 | 662.193 | | | | 662.22 | 0.03 |
| 1500 | 662.881 | 662.776 | | | | 662.83 | 0.07 |
| 3600 | 663.107 | 662.9 | | | | 663.00 | 0.15 |
| 7200 | 663.434 | 662.85 | | | | 663.14 | 0.41 |
| 14400 | 664.084 | 663.148 | | | | 663.62 | 0.66 |

**DNA**

| Time | Replicate 1 | Replicate 2 | Replicate 3 | Replicate 4 | Replicate 5 | average | SD |
|---|---|---|---|---|---|---|---|
| 0 | 661.577 | 661.63 | | | | 661.60 | 0.04 |
| 30 | 662.118 | 662.199 | | | | 662.16 | 0.06 |
| 45 | 662.525 | 662.145 | | | | 662.34 | 0.27 |
| 60 | 662.585 | 662.327 | | | | 662.46 | 0.18 |
| 300 | 662.251 | 662.302 | | | | 662.28 | 0.04 |
| 1500 | 662.565 | 662.733 | | | | 662.65 | 0.12 |
| 3600 | 662.694 | 662.542 | | | | 662.62 | 0.11 |
| 7200 | | 662.702 | | | | 662.70 | |
| 14400 | 663.087 | 662.11 | | | | 662.60 | 0.69 |

**ONPFDNA**

| Time | Replicate 1 | Replicate 2 | Replicate 3 | Replicate 4 | average | SD |
|---|---|---|---|---|---|---|
| 0 | 661.529 | 661.611 | | | 661.57 | 0.06 |
| 30 | 662 | 662.087 | | | 662.04 | 0.06 |
| 45 | 662.179 | 662.162 | | | 662.17 | 0.01 |
| 60 | 662.397 | 662.297 | | | 662.35 | 0.07 |
| 300 | 662.537 | 662.236 | | | 662.39 | 0.21 |
| 1500 | 662.413 | 662.466 | | | 662.44 | 0.04 |
| 3600 | 662.656 | 662.744 | | | 662.70 | 0.06 |
| 7200 | 661.945 | 662.823 | | | 662.38 | 0.62 |
| 14400 | 661.941 | 662.528 | | | 662.23 | 0.42 |

**TMG**

| Time | Replicate 1 | Replicate 2 | Replicate 3 | Replicate 4 | Replicate 5 | average | SD |
|---|---|---|---|---|---|---|---|
| 0 | 661.697 | 661.642 | | | | 661.67 | 0.04 |
| 30 | 662.259 | 662.462 | | | | 662.36 | 0.14 |
| 45 | 662.527 | 662.604 | | | | 662.57 | 0.05 |
| 60 | 662.482 | 662.23 | | | | 662.36 | 0.18 |
| 300 | 662.555 | 662.61 | | | | 662.58 | 0.04 |
| 1500 | 662.657 | 663.025 | | | | 662.84 | 0.26 |
| 3600 | 663.104 | 663.643 | | | | 663.37 | 0.38 |
| 7200 | 663.073 | 663.57 | | | | 663.32 | 0.35 |
| 14400 | 663.422 | 663.582 | | | | 663.50 | 0.11 |

### 94-98 VVVSM

**Charge** 1

**IPTG**

| Time | centroid | D |
|---|---|---|
| 0 | 534.609 | 0.00 |
| 30 | 535.287 | 0.86 |
| 45 | 535.322 | 0.90 |
| 60 | 535.344 | 0.93 |
| 300 | 535.337 | 0.92 |
| 1500 | 535.649 | 1.32 |
| 3600 | 535.783 | 1.49 |
| 7200 | 535.966 | 1.72 |
| 14400 | 535.998 | 1.76 |

**ONPF**

| Time | centroid | D |
|---|---|---|
| 0 | 534.773 | 0.21 |
| 30 | 535.0925 | 0.61 |
| 45 | 535.4065 | 1.01 |
| 60 | 535.5555 | 1.20 |
| 300 | 535.61 | 1.27 |
| 1500 | 535.7455 | 1.44 |
| 3600 | 535.732 | 1.42 |
| 7200 | 535.632 | 1.30 |
| 14400 | 535.67 | 1.35 |

**APO**

| Time | centroid | D |
|---|---|---|
| 0 | 534.611 | 0.00 |
| 30 | 535.2675 | 0.83 |
| 45 | 535.293 | 0.87 |
| 60 | 535.546 | 1.19 |
| 300 | 535.69 | 1.37 |
| 1500 | 535.881 | 1.61 |
| 3600 | 535.9345 | 1.68 |
| 7200 | 535.7665 | 1.47 |
| 14400 | 535.937 | 1.68 |

**DNA**

| Time | centroid | D |
|---|---|---|
| 0 | 534.6145 | 0.00 |
| 30 | 535.3825 | 0.98 |
| 45 | 535.7515 | 1.45 |
| 60 | 535.8375 | 1.56 |
| 300 | 535.879 | 1.61 |
| 1500 | 535.945 | 1.69 |
| 3600 | 535.895 | 1.63 |
| 7200 | 535.685 | 1.36 |
| 14400 | 535.825 | 1.54 |

**ONPFDNA**

| Time | centroid | D |
|---|---|---|
| 0 | 534.613 | 0.00 |
| 30 | 535.306 | 0.88 |
| 45 | 535.7015 | 1.39 |
| 60 | 535.7595 | 1.46 |
| 300 | 535.8265 | 1.54 |
| 1500 | 535.7865 | 1.49 |
| 3600 | 535.9385 | 1.69 |
| 7200 | 535.748 | 1.44 |
| 14400 | 535.6975 | 1.38 |

**TMG**

| Time | centroid | D |
|---|---|---|
| 0 | 534.665 | 0.07 |
| 30 | 535.166 | 0.70 |
| 45 | 535.4085 | 1.01 |
| 60 | 535.3375 | 0.92 |
| 300 | 535.3735 | 0.97 |
| 1500 | 535.4975 | 1.13 |
| 3600 | 535.808 | 1.52 |
| 7200 | 535.809 | 1.52 |
| 14400 | 535.975 | 1.73 |

| | |
|---|---|
| control | 534.611 |
| infinity | 536.973 |

$$D(t) = \frac{M_t - M_0}{M_\infty - M_0} \cdot N$$

**IPTG**

| Time | Replicate 1 | Replicate 2 | Replicate 3 | Replicate 4 | Replicate5 | average | SD |
|---|---|---|---|---|---|---|---|
| 0 | 534.609 | 534.605 | | | | 534.61 | 0.00 |
| 30 | 535.287 | 535.059 | | | | 535.17 | 0.16 |
| 45 | 535.322 | 535.235 | | | | 535.28 | 0.06 |
| 60 | 535.344 | 535.298 | | | | 535.32 | 0.03 |
| 300 | 535.337 | 535.214 | | | | 535.28 | 0.09 |
| 1500 | 535.649 | 535.383 | | | | 535.52 | 0.19 |
| 3600 | 535.783 | 535.733 | | | | 535.76 | 0.04 |
| 7200 | 535.966 | 535.73 | | | | 535.85 | 0.17 |
| 14400 | 535.998 | 535.713 | | | | 535.86 | 0.20 |

**ONPF**

| Time | Replicate 1 | Replicate 2 | Replicate 3 | Replicate 4 | average | SD |
|---|---|---|---|---|---|---|
| 0 | 534.797 | 534.749 | | | 534.77 | 0.03 |
| 30 | 535.181 | 535.004 | | | 535.09 | 0.13 |
| 45 | 535.645 | 535.168 | | | 535.41 | 0.34 |
| 60 | 535.63 | 535.481 | | | 535.56 | 0.11 |
| 300 | 535.818 | 535.402 | | | 535.61 | 0.29 |
| 1500 | 535.906 | 535.585 | | | 535.75 | 0.23 |
| 3600 | 535.862 | 535.602 | | | 535.73 | 0.18 |
| 7200 | 535.756 | 535.508 | | | 535.63 | 0.18 |
| 14400 | 535.96 | 535.38 | | | 535.67 | 0.41 |

**APO**

| Time | Replicate 1 | Replicate 2 | Replicate 3 | Replicate 4 | Replicate 5 | average | SD |
|---|---|---|---|---|---|---|---|
| 0 | 534.607 | 534.615 | | | | 534.61 | 0.01 |
| 30 | 535.408 | 535.127 | | | | 535.27 | 0.20 |
| 45 | 535.385 | 535.201 | | | | 535.29 | 0.13 |
| 60 | 535.595 | 535.497 | | | | 535.55 | 0.07 |
| 300 | 535.927 | 535.453 | | | | 535.69 | 0.34 |
| 1500 | 535.995 | 535.767 | | | | 535.88 | 0.16 |
| 3600 | 536.06 | 535.809 | | | | 535.93 | 0.18 |
| 7200 | 535.96 | 535.573 | | | | 535.77 | 0.27 |
| 14400 | 536.117 | 535.757 | | | | 535.94 | 0.25 |

**DNA**

| Time | Replicate 1 | Replicate 2 | Replicate 3 | Replicate 4 | Replicate 5 | average | SD |
|---|---|---|---|---|---|---|---|
| 0 | 534.609 | 534.62 | | | | 534.61 | 0.01 |
| 30 | 535.396 | 535.369 | | | | 535.38 | 0.02 |
| 45 | 535.931 | 535.572 | | | | 535.75 | 0.25 |
| 60 | 535.933 | 535.742 | | | | 535.84 | 0.14 |
| 300 | 535.968 | 535.79 | | | | 535.88 | 0.13 |
| 1500 | 536.001 | 535.889 | | | | 535.95 | 0.08 |
| 3600 | 535.995 | 535.795 | | | | 535.90 | 0.14 |
| 7200 | 535.863 | 535.507 | | | | 535.69 | 0.25 |
| 14400 | 536.014 | 535.636 | | | | 535.83 | 0.27 |

**ONPFDNA**

| Time | Replicate 1 | Replicate 2 | Replicate 3 | Replicate 4 | average | SD |
|---|---|---|---|---|---|---|
| 0 | 534.618 | 534.608 | | | 534.61 | 0.01 |
| 30 | 535.345 | 535.267 | | | 535.31 | 0.06 |
| 45 | 535.826 | 535.577 | | | 535.70 | 0.18 |
| 60 | 535.835 | 535.684 | | | 535.76 | 0.11 |
| 300 | 535.932 | 535.721 | | | 535.83 | 0.15 |
| 1500 | 535.869 | 535.704 | | | 535.79 | 0.12 |
| 3600 | 535.983 | 535.894 | | | 535.94 | 0.06 |
| 7200 | 535.839 | 535.657 | | | 535.75 | 0.13 |
| 14400 | 535.84 | 535.555 | | | 535.70 | 0.20 |

**TMG**

| Time | Replicate 1 | Replicate 2 | Replicate 3 | Replicate 4 | Replicate 5 | average | SD |
|---|---|---|---|---|---|---|---|
| 0 | 534.723 | 534.607 | | | | 534.67 | 0.08 |
| 30 | 535.054 | 535.278 | | | | 535.17 | 0.16 |
| 45 | 535.489 | 535.328 | | | | 535.41 | 0.11 |
| 60 | 535.375 | 535.3 | | | | 535.34 | 0.05 |
| 300 | 535.372 | 535.375 | | | | 535.37 | 0.00 |
| 1500 | 535.47 | 535.525 | | | | 535.50 | 0.04 |
| 3600 | 535.785 | 535.831 | | | | 535.81 | 0.03 |
| 7200 | 535.653 | 535.965 | | | | 535.81 | 0.22 |
| 14400 | 535.901 | 536.049 | | | | 535.98 | 0.10 |

### 98-105 MVERSGVE

**Charge** 2

**IPTG**

| Time | centroid | D |
|---|---|---|
| 0 | 453.9893 | 0.00 |
| 30 | 455.1028 | 4.19 |
| 45 | | |
| 60 | 455.2693 | 4.82 |
| 300 | 455.2928 | 4.91 |
| 1500 | 455.2773 | 4.85 |
| 3600 | 455.2728 | 4.83 |
| 7200 | 455.308 | 4.97 |
| 14400 | 455.2995 | 4.94 |

**ONPF**

| Time | centroid | D |
|---|---|---|
| 0 | 454.0058 | 0.06 |
| 30 | 454.8635 | 3.29 |
| 45 | | |
| 60 | 455.15 | 4.37 |
| 300 | 455.1777 | 4.48 |
| 1500 | 455.1448 | 4.35 |
| 3600 | 455.222 | 4.64 |
| 7200 | 455.157 | 4.40 |
| 14400 | 455.27 | 4.83 |

**APO**

| Time | centroid | D |
|---|---|---|
| 0 | 453.9948 | 0.02 |
| 30 | 455.0735 | 4.08 |
| 45 | | |
| 60 | 455.2315 | 4.68 |
| 300 | 455.2135 | 4.61 |
| 1500 | 455.2643 | 4.80 |
| 3600 | 455.2575 | 4.78 |
| 7200 | 455.3 | 4.94 |
| 14400 | 455.3853 | 5.26 |

**DNA**

| Time | centroid | D |
|---|---|---|
| 0 | 453.9883 | -0.01 |
| 30 | 454.9477 | 3.61 |
| 45 | | |
| 60 | 455.21 | 4.60 |
| 300 | 455.2633 | 4.80 |
| 1500 | 455.2407 | 4.71 |
| 3600 | 455.267 | 4.81 |
| 7200 | 455.2173 | 4.63 |
| 14400 | 455.2623 | 4.80 |

**ONPFDNA**

| Time | centroid | D |
|---|---|---|
| 0 | 453.9857 | -0.02 |
| 30 | 454.944 | 3.60 |
| 45 | | |
| 60 | 455.2563 | 4.77 |
| 300 | 455.274 | 4.84 |
| 1500 | 455.2013 | 4.57 |
| 3600 | 455.3243 | 5.03 |
| 7200 | 455.2117 | 4.60 |
| 14400 | 455.179 | 4.48 |

**TMG**

| Time | centroid | D |
|---|---|---|
| 0 | 454.006 | 0.06 |
| 30 | 455.288 | 4.89 |
| 45 | 455.429 | 5.42 |
| 60 | 455.486 | 5.64 |
| 300 | 455.533 | 5.82 |
| 1500 | 455.458 | 5.53 |
| 3600 | 455.511 | 5.73 |
| 7200 | 455.4815 | 5.62 |
| 14400 | 455.564 | 5.93 |

control 453.99
infinity 455.582

$$D(t) = \frac{Mt - M0}{M\infty - M0} *N$$

**IPTG**

| Time | Replicate 1 | Replicate 2 | Replicate 3 | Replicate 4 | Replicate5 | average | SD |
|---|---|---|---|---|---|---|---|
| 0 | 453.98 | 453.995 | 453.987 | 453.995 | | 453.99 | 0.01 |
| 30 | 455.645 | 454.923 | 454.844 | 454.999 | | 455.10 | 0.37 |
| 45 | | | | | | | |
| 60 | 455.716 | 455.14 | 455.065 | 455.156 | | 455.27 | 0.30 |
| 300 | 455.694 | 455.248 | 455.148 | 455.081 | | 455.29 | 0.28 |
| 1500 | 455.701 | 455.256 | 455.148 | 455.004 | | 455.28 | 0.30 |
| 3600 | 455.67 | 455.223 | 455.129 | 455.069 | | 455.27 | 0.27 |
| 7200 | 455.701 | 455.185 | | 455.038 | | 455.31 | 0.35 |
| 14400 | 455.637 | 455.271 | 455.238 | 455.052 | | 455.30 | 0.24 |

**ONPF**

| Time | Replicate 1 | Replicate 2 | Replicate 3 | Replicate 4 | average | SD |
|---|---|---|---|---|---|---|
| 0 | 454.055 | 453.989 | 453.992 | 453.987 | 454.01 | 0.03 |
| 30 | 454.941 | 454.81 | 454.675 | 455.028 | 454.86 | 0.15 |
| 45 | | | | | | |
| 60 | 455.344 | 455.265 | 455.03 | 454.961 | 455.15 | 0.18 |
| 300 | 455.418 | 454.963 | | 455.152 | 455.18 | 0.23 |
| 1500 | 455.388 | 455.201 | 455.002 | 454.988 | 455.14 | 0.19 |
| 3600 | 455.32 | 455.336 | 454.958 | 455.274 | 455.22 | 0.18 |
| 7200 | 455.287 | 455.282 | 454.951 | 455.108 | 455.16 | 0.16 |
| 14400 | 455.472 | 455.222 | 455.171 | 455.216 | 455.27 | 0.14 |

**APO**

| Time | Replicate 1 | Replicate 2 | Replicate 3 | Replicate 4 | Replicate 5 | average | SD |
|---|---|---|---|---|---|---|---|
| 0 | 453.99 | 454.012 | 453.985 | 453.992 | | 453.99 | 0.01 |
| 30 | 455.557 | 455.218 | 454.779 | 454.74 | | 455.07 | 0.39 |
| 45 | | | | | | | |
| 60 | 455.705 | 455.558 | 454.75 | 454.913 | | 455.23 | 0.47 |
| 300 | 455.573 | 455.34 | 455.062 | 454.879 | | 455.21 | 0.31 |
| 1500 | 455.568 | 455.46 | 454.982 | 455.047 | | 455.26 | 0.29 |
| 3600 | 455.614 | 455.477 | 455.054 | 454.885 | | 455.26 | 0.34 |
| 7200 | 455.53 | 455.609 | 455.096 | 454.965 | | 455.30 | 0.32 |
| 14400 | 455.661 | 455.746 | 455.063 | 455.071 | | 455.39 | 0.37 |

**DNA**

| Time | Replicate 1 | Replicate 2 | Replicate 3 | Replicate 4 | Replicate 5 | average | SD |
|---|---|---|---|---|---|---|---|
| 0 | 453.987 | 453.989 | 453.989 | | | 453.99 | 0.00 |
| 30 | 455.115 | 454.993 | 454.735 | | | 454.95 | 0.19 |
| 45 | | | | | | | |
| 60 | 455.378 | | 455.042 | | | 455.21 | 0.24 |
| 300 | 455.496 | 455.432 | 454.862 | | | 455.26 | 0.35 |
| 1500 | 455.39 | 455.377 | 454.955 | | | 455.24 | 0.25 |
| 3600 | 455.464 | 455.447 | 454.89 | | | 455.27 | 0.33 |
| 7200 | 455.4 | 455.412 | 454.84 | | | 455.22 | 0.33 |
| 14400 | 455.403 | 455.377 | 455.007 | | | 455.26 | 0.22 |

**ONPFDNA**

| Time | Replicate 1 | Replicate 2 | Replicate 3 | Replicate 4 | average | SD |
|---|---|---|---|---|---|---|
| 0 | 453.99 | 453.976 | 453.991 | | 453.99 | 0.01 |
| 30 | 455.027 | 455.253 | 454.552 | | 454.94 | 0.36 |
| 45 | | | | | | |
| 60 | 455.321 | 455.397 | 455.051 | | 455.26 | 0.18 |
| 300 | 455.431 | 455.333 | 455.058 | | 455.27 | 0.19 |
| 1500 | 455.093 | 455.432 | 455.079 | | 455.20 | 0.20 |
| 3600 | 455.425 | 455.412 | 455.136 | | 455.32 | 0.16 |
| 7200 | 455.379 | 455.179 | 455.077 | | 455.21 | 0.15 |
| 14400 | 454.949 | 455.447 | 455.141 | | 455.18 | 0.25 |

**TMG**

| Time | Replicate 1 | Replicate 2 | Replicate 3 | Replicate 4 | Replicate 5 | average | SD |
|---|---|---|---|---|---|---|---|
| 0 | 454.03 | 453.982 | | | | 454.01 | 0.03 |
| 30 | 454.932 | 455.644 | | | | 455.29 | 0.50 |
| 45 | 455.269 | 455.589 | | | | 455.43 | 0.23 |
| 60 | 455.439 | 455.533 | | | | 455.49 | 0.07 |
| 300 | 455.42 | 455.646 | | | | 455.53 | 0.16 |
| 1500 | 455.243 | 455.673 | | | | 455.46 | 0.30 |
| 3600 | 455.304 | 455.718 | | | | 455.51 | 0.29 |
| 7200 | 455.26 | 455.703 | | | | 455.48 | 0.31 |
| 14400 | 455.387 | 455.741 | | | | 455.56 | 0.25 |

### 109-114 AAVHNL

**Charge** 1

**IPTG**

| Time | centroid | D |
|---|---|---|
| 0 | 624.73 | 0.17 |
| 30 | 624.96 | 0.60 |
| 45 | 624.94 | 0.56 |
| 60 | 624.93 | 0.53 |
| 300 | 625.01 | 0.69 |
| 1500 | 625.26 | 1.17 |
| 3600 | 625.53 | 1.67 |
| 7200 | 625.83 | 2.25 |
| 14400 | 625.93 | 2.44 |

**ONPF**

| Time | centroid | D |
|---|---|---|
| 0 | 624.69 | 0.08 |
| 30 | 624.79 | 0.28 |
| 45 | 624.86 | 0.40 |
| 60 | 624.86 | 0.41 |
| 300 | 624.95 | 0.58 |
| 1500 | 625.13 | 0.91 |
| 3600 | 625.17 | 1.00 |
| 7200 | 625.15 | 0.96 |
| 14400 | 625.43 | 1.50 |

**APO**

| Time | centroid | D |
|---|---|---|
| 0 | 624.65 | 0.00 |
| 30 | 624.78 | 0.25 |
| 45 | 624.84 | 0.37 |
| 60 | 624.84 | 0.37 |
| 300 | 624.84 | 0.37 |
| 1500 | 625.07 | 0.80 |
| 3600 | 625.10 | 0.87 |
| 7200 | 625.53 | 1.69 |
| 14400 | 625.51 | 1.65 |

**DNA**

| Time | centroid | D |
|---|---|---|
| 0 | 624.67 | 0.04 |
| 30 | 624.78 | 0.25 |
| 45 | 624.85 | 0.40 |
| 60 | 624.83 | 0.35 |
| 300 | 624.95 | 0.59 |
| 1500 | 625.01 | 0.70 |
| 3600 | 625.07 | 0.80 |
| 7200 | 625.02 | 0.71 |
| 14400 | 624.99 | 0.66 |

**ONPFDNA**

| Time | centroid | D |
|---|---|---|
| 0 | 624.71 | 0.12 |
| 30 | 624.82 | 0.34 |
| 45 | 624.84 | 0.37 |
| 60 | 624.88 | 0.43 |
| 300 | 624.96 | 0.59 |
| 1500 | 624.94 | 0.55 |
| 3600 | 625.09 | 0.85 |
| 7200 | 625.03 | 0.73 |
| 14400 | 624.95 | 0.57 |

**TMG**

| Time | centroid | D |
|---|---|---|
| 0 | 624.67 | 0.04 |
| 30 | 624.86 | 0.40 |
| 45 | 624.98 | 0.64 |
| 60 | 624.92 | 0.51 |
| 300 | 625.00 | 0.68 |
| 1500 | 625.19 | 1.04 |
| 3600 | 625.44 | 1.51 |
| 7200 | 625.56 | 1.74 |
| 14400 | 625.86 | 2.31 |

control 624.6465
infinity 626.748

$$D(t) = \frac{M_t - M_0}{M_\infty - M_0} \cdot N$$

**IPTG**

| Time | Replicate 1 | Replicate 2 | Replicate 3 | Replicate 4 | Replicate5 | average | SD |
|---|---|---|---|---|---|---|---|
| 0 | 624.688 | 624.78 | | | | 624.73 | 0.07 |
| 30 | 624.924 | 624.999 | | | | 624.96 | 0.05 |
| 45 | 624.952 | 624.932 | | | | 624.94 | 0.01 |
| 60 | 624.963 | 624.888 | | | | 624.93 | 0.05 |
| 300 | 625.113 | 624.9 | | | | 625.01 | 0.15 |
| 1500 | 625.526 | 624.993 | | | | 625.26 | 0.38 |
| 3600 | 625.805 | 625.248 | | | | 625.53 | 0.39 |
| 7200 | 626.068 | 625.591 | | | | 625.83 | 0.34 |
| 14400 | 626.227 | 625.625 | | | | 625.93 | 0.43 |

**ONPF**

| Time | Replicate 1 | Replicate 2 | Replicate 3 | Replicate 4 | average | SD |
|---|---|---|---|---|---|---|
| 0 | 624.671 | 624.707 | | | 624.69 | 0.03 |
| 30 | 624.82 | 624.763 | | | 624.79 | 0.04 |
| 45 | 624.915 | 624.802 | | | 624.86 | 0.08 |
| 60 | 624.843 | 624.88 | | | 624.86 | 0.03 |
| 300 | 624.941 | 624.966 | | | 624.95 | 0.02 |
| 1500 | 625.036 | 625.214 | | | 625.13 | 0.13 |
| 3600 | 625.225 | 625.118 | | | 625.17 | 0.08 |
| 7200 | 625.21 | 625.092 | | | 625.15 | 0.08 |
| 14400 | 625.432 | | | | 625.43 | |

**APO**

| Time | Replicate 1 | Replicate 2 | Replicate 3 | Replicate 4 | Replicate 5 | average | SD |
|---|---|---|---|---|---|---|---|
| 0 | 624.613 | 624.68 | | | | 624.65 | 0.05 |
| 30 | 624.783 | 624.769 | | | | 624.78 | 0.01 |
| 45 | 624.903 | 624.778 | | | | 624.84 | 0.09 |
| 60 | 624.839 | 624.845 | | | | 624.84 | 0.00 |
| 300 | 624.894 | 624.79 | | | | 624.84 | 0.07 |
| 1500 | 625.08 | 625.052 | | | | 625.07 | 0.02 |
| 3600 | 625.146 | 625.061 | | | | 625.10 | 0.06 |
| 7200 | 625.86 | 625.209 | | | | 625.53 | 0.46 |
| 14400 | 625.617 | 625.408 | | | | 625.51 | 0.15 |

**DNA**

| Time | Replicate 1 | Replicate 2 | Replicate 3 | Replicate 4 | Replicate 5 | average | SD |
|---|---|---|---|---|---|---|---|
| 0 | 624.652 | 624.685 | | | | 624.67 | 0.02 |
| 30 | 624.808 | 624.744 | | | | 624.78 | 0.05 |
| 45 | 624.945 | 624.764 | | | | 624.85 | 0.13 |
| 60 | 624.837 | 624.827 | | | | 624.83 | 0.01 |
| 300 | 625.06 | 624.849 | | | | 624.95 | 0.15 |
| 1500 | 625.015 | 625.012 | | | | 625.01 | 0.00 |
| 3600 | 625.218 | 624.915 | | | | 625.07 | 0.21 |
| 7200 | 625.113 | 624.928 | | | | 625.02 | 0.13 |
| 14400 | 625.175 | 624.813 | | | | 624.99 | 0.26 |

**ONPFDNA**

| Time | Replicate 1 | Replicate 2 | Replicate 3 | Replicate 4 | average | SD |
|---|---|---|---|---|---|---|
| 0 | 624.667 | 624.75 | | | 624.71 | 0.06 |
| 30 | 624.823 | 624.824 | | | 624.82 | 0.00 |
| 45 | 624.834 | 624.843 | | | 624.84 | 0.01 |
| 60 | 624.91 | 624.84 | | | 624.88 | 0.05 |
| 300 | 625.051 | 624.859 | | | 624.96 | 0.14 |
| 1500 | 624.927 | 624.946 | | | 624.94 | 0.01 |
| 3600 | 625.161 | 625.026 | | | 625.09 | 0.10 |
| 7200 | | 625.031 | | | 625.03 | |
| 14400 | 624.923 | 624.972 | | | 624.95 | 0.03 |

**TMG**

| Time | Replicate 1 | Replicate 2 | Replicate 3 | Replicate 4 | Replicate 5 | average | SD |
|---|---|---|---|---|---|---|---|
| 0 | 624.684 | 624.656 | | | | 624.67 | 0.02 |
| 30 | 624.851 | 624.859 | | | | 624.86 | 0.01 |
| 45 | 625.11 | 624.851 | | | | 624.98 | 0.18 |
| 60 | 624.96 | 624.872 | | | | 624.92 | 0.06 |
| 300 | 625.047 | 624.961 | | | | 625.00 | 0.06 |
| 1500 | 625.242 | 625.147 | | | | 625.19 | 0.07 |
| 3600 | 625.539 | 625.336 | | | | 625.44 | 0.14 |
| 7200 | 625.659 | 625.463 | | | | 625.56 | 0.14 |
| 14400 | 625.929 | 625.791 | | | | 625.86 | 0.10 |

### 115-122 LAQRVSGL

**Charge** 2

**IPTG**

| Time | centroid | D |
|---|---|---|
| 0 | 422.50 | 0.03 |
| 30 | 422.85 | 1.01 |
| 45 | 422.94 | 1.25 |
| 60 | 422.97 | 1.33 |
| 300 | 423.03 | 1.51 |
| 1500 | 423.19 | 1.94 |
| 3600 | 423.37 | 2.44 |
| 7200 | 423.49 | 2.79 |
| 14400 | 423.50 | 2.82 |

**ONPF**

| Time | centroid | D |
|---|---|---|
| 0 | 422.51 | 0.05 |
| 30 | 422.69 | 0.57 |
| 45 | 422.89 | 1.11 |
| 60 | 422.92 | 1.20 |
| 300 | 422.92 | 1.20 |
| 1500 | 423.04 | 1.54 |
| 3600 | 423.11 | 1.73 |
| 7200 | 423.13 | 1.79 |
| 14400 | 423.01 | 1.45 |

**APO**

| Time | centroid | D |
|---|---|---|
| 0 | 422.49 | 0.01 |
| 30 | 422.79 | 0.83 |
| 45 | 422.86 | 1.03 |
| 60 | 422.97 | 1.34 |
| 300 | 422.93 | 1.23 |
| 1500 | 423.18 | 1.92 |
| 3600 | 423.28 | 2.21 |
| 7200 | 423.34 | 2.36 |
| 14400 | 423.45 | 2.68 |

**DNA**

| Time | centroid | D |
|---|---|---|
| 0 | 422.49 | 0.01 |
| 30 | 422.60 | 0.29 |
| 45 | 422.68 | 0.52 |
| 60 | 422.67 | 0.49 |
| 300 | 422.66 | 0.46 |
| 1500 | 422.75 | 0.74 |
| 3600 | 422.79 | 0.84 |
| 7200 | 422.82 | 0.93 |
| 14400 | 422.83 | 0.94 |

**ONPFDNA**

| Time | centroid | D |
|---|---|---|
| 0 | 422.50 | 0.01 |
| 30 | 422.58 | 0.25 |
| 45 | 422.62 | 0.37 |
| 60 | 422.66 | 0.46 |
| 300 | 422.62 | 0.37 |
| 1500 | 422.69 | 0.57 |
| 3600 | 422.81 | 0.90 |
| 7200 | 422.82 | 0.91 |
| 14400 | 422.77 | 0.78 |

**TMG**

| Time | centroid | D |
|---|---|---|
| 0 | 422.50 | 0.02 |
| 30 | 422.84 | 0.97 |
| 45 | 423.01 | 1.44 |
| 60 | 422.92 | 1.20 |
| 300 | 423.10 | 1.70 |
| 1500 | 423.17 | 1.90 |
| 3600 | 423.38 | 2.48 |
| 7200 | 423.43 | 2.62 |
| 14400 | 423.59 | 3.07 |

control 422.49
infinity 424.647

$$D(t) = \frac{M_t - M_0}{M_\infty - M_0} * N$$

**IPTG**

| Time | Replicate 1 | Replicate 2 | Replicate 3 | Replicate 4 | Replicate5 | average | SD |
|---|---|---|---|---|---|---|---|
| 0 | 422.509 | 422.489 | | | | 422.50 | 0.01 |
| 30 | 422.966 | 422.743 | | | | 422.85 | 0.16 |
| 45 | 423.002 | 422.879 | | | | 422.94 | 0.09 |
| 60 | 422.979 | 422.959 | | | | 422.97 | 0.01 |
| 300 | 423.119 | 422.947 | | | | 423.03 | 0.12 |
| 1500 | 423.321 | 423.056 | | | | 423.19 | 0.19 |
| 3600 | 423.454 | 423.281 | | | | 423.37 | 0.12 |
| 7200 | 423.626 | 423.361 | | | | 423.49 | 0.19 |
| 14400 | 423.664 | 423.34 | | | | 423.50 | 0.23 |

**ONPF**

| Time | Replicate 1 | Replicate 2 | Replicate 3 | Replicate 4 | average | SD |
|---|---|---|---|---|---|---|
| 0 | 422.503 | 422.512 | | | 422.51 | 0.01 |
| 30 | 422.713 | 422.674 | | | 422.69 | 0.03 |
| 45 | 423.056 | 422.72 | | | 422.89 | 0.24 |
| 60 | 422.94 | 422.906 | | | 422.92 | 0.02 |
| 300 | 423.009 | 422.831 | | | 422.92 | 0.13 |
| 1500 | 423.122 | 422.963 | | | 423.04 | 0.11 |
| 3600 | 423.235 | 422.99 | | | 423.11 | 0.17 |
| 7200 | 423.195 | 423.07 | | | 423.13 | 0.09 |
| 14400 | 423.487 | 422.536 | | | 423.01 | 0.67 |

**APO**

| Time | Replicate 1 | Replicate 2 | Replicate 3 | Replicate 4 | Replicate 5 | average | SD |
|---|---|---|---|---|---|---|---|
| 0 | 422.491 | 422.497 | | | | 422.49 | 0.00 |
| 30 | 422.83 | 422.748 | | | | 422.79 | 0.06 |
| 45 | 422.946 | 422.773 | | | | 422.86 | 0.12 |
| 60 | 422.968 | 422.975 | | | | 422.97 | 0.00 |
| 300 | 423.01 | 422.857 | | | | 422.93 | 0.11 |
| 1500 | 423.189 | 423.17 | | | | 423.18 | 0.01 |
| 3600 | 423.354 | 423.212 | | | | 423.28 | 0.10 |
| 7200 | 423.468 | 423.211 | | | | 423.34 | 0.18 |
| 14400 | 423.537 | 423.372 | | | | 423.45 | 0.12 |

**DNA**

| Time | Replicate 1 | Replicate 2 | Replicate 3 | Replicate 4 | Replicate 5 | average | SD |
|---|---|---|---|---|---|---|---|
| 0 | 422.492 | 422.494 | | | | 422.49 | 0.00 |
| 30 | 422.624 | 422.567 | | | | 422.60 | 0.04 |
| 45 | 422.766 | 422.589 | | | | 422.68 | 0.13 |
| 60 | 422.657 | 422.678 | | | | 422.67 | 0.01 |
| 300 | 422.678 | 422.635 | | | | 422.66 | 0.03 |
| 1500 | 422.746 | 422.763 | | | | 422.75 | 0.01 |
| 3600 | 422.863 | 422.722 | | | | 422.79 | 0.10 |
| 7200 | 422.88 | 422.767 | | | | 422.82 | 0.08 |
| 14400 | 423.016 | 422.64 | | | | 422.83 | 0.27 |

**ONPFDNA**

| Time | Replicate 1 | Replicate 2 | Replicate 3 | Replicate 4 | average | SD |
|---|---|---|---|---|---|---|
| 0 | 422.498 | 422.492 | | | 422.50 | 0.00 |
| 30 | 422.597 | 422.566 | | | 422.58 | 0.02 |
| 45 | 422.655 | 422.588 | | | 422.62 | 0.05 |
| 60 | 422.658 | 422.652 | | | 422.66 | 0.00 |
| 300 | 422.64 | 422.608 | | | 422.62 | 0.02 |
| 1500 | 422.668 | 422.719 | | | 422.69 | 0.04 |
| 3600 | 422.865 | 422.761 | | | 422.81 | 0.07 |
| 7200 | 422.834 | 422.799 | | | 422.82 | 0.02 |
| 14400 | 422.776 | 422.767 | | | 422.77 | 0.01 |

**TMG**

| Time | Replicate 1 | Replicate 2 | Replicate 3 | Replicate 4 | Replicate 5 | average | SD |
|---|---|---|---|---|---|---|---|
| 0 | 422.499 | 422.492 | | | | 422.50 | 0.00 |
| 30 | 422.744 | 422.936 | | | | 422.84 | 0.14 |
| 45 | 423.079 | 422.939 | | | | 423.01 | 0.10 |
| 60 | 422.97 | 422.872 | | | | 422.92 | 0.07 |
| 300 | 423.063 | 423.136 | | | | 423.10 | 0.05 |
| 1500 | 423.119 | 423.224 | | | | 423.17 | 0.07 |
| 3600 | 423.314 | 423.45 | | | | 423.38 | 0.10 |
| 7200 | 423.297 | 423.567 | | | | 423.43 | 0.19 |
| 14400 | 423.499 | 423.685 | | | | 423.59 | 0.13 |

### 119-125 VSGLIIN

**Charge** 1

**IPTG**

| Time | centroid | D |
|---|---|---|
| 0 | 715.82 | 0.03 |
| 30 | 715.93 | 0.17 |
| 45 | 715.98 | 0.23 |
| 60 | 715.98 | 0.23 |
| 300 | 715.96 | 0.20 |
| 1500 | 715.97 | 0.22 |
| 3600 | 715.96 | 0.21 |
| 7200 | 715.98 | 0.22 |
| 14400 | 715.96 | 0.20 |

**ONPF**

| Time | centroid | D |
|---|---|---|
| 0 | 715.87 | 0.09 |
| 30 | 715.95 | 0.19 |
| 45 | 715.99 | 0.24 |
| 60 | 715.99 | 0.24 |
| 300 | 715.98 | 0.23 |
| 1500 | 715.98 | 0.23 |
| 3600 | 716.00 | 0.25 |
| 7200 | 715.95 | 0.20 |
| 14400 | 715.97 | 0.22 |

**APO**

| Time | centroid | D |
|---|---|---|
| 0 | 715.79 | 0.00 |
| 30 | 715.91 | 0.14 |
| 45 | 715.95 | 0.19 |
| 60 | 715.96 | 0.20 |
| 300 | 715.92 | 0.16 |
| 1500 | 715.92 | 0.15 |
| 3600 | 715.90 | 0.13 |
| 7200 | 715.88 | 0.10 |
| 14400 | 715.92 | 0.16 |

**DNA**

| Time | centroid | D |
|---|---|---|
| 0 | 715.85 | 0.06 |
| 30 | 715.94 | 0.18 |
| 45 | 715.92 | 0.16 |
| 60 | 715.95 | 0.19 |
| 300 | 715.95 | 0.19 |
| 1500 | 715.97 | 0.22 |
| 3600 | 715.99 | 0.24 |
| 7200 | 715.99 | 0.24 |
| 14400 | 716.00 | 0.25 |

**ONPFDNA**

| Time | centroid | D |
|---|---|---|
| 0 | 715.89 | 0.11 |
| 30 | 715.98 | 0.23 |
| 45 | 715.93 | 0.17 |
| 60 | 715.98 | 0.23 |
| 300 | 716.04 | 0.31 |
| 1500 | 715.97 | 0.21 |
| 3600 | 715.96 | 0.20 |
| 7200 | 716.02 | 0.27 |
| 14400 | 715.99 | 0.25 |

**TMG**

| Time | centroid | D |
|---|---|---|
| 0 | 715.81 | 0.02 |
| 30 | 715.95 | 0.19 |
| 45 | 715.95 | 0.19 |
| 60 | 716.01 | 0.26 |
| 300 | 716.00 | 0.26 |
| 1500 | 716.00 | 0.25 |
| 3600 | 715.96 | 0.21 |
| 7200 | 715.99 | 0.24 |
| 14400 | 716.01 | 0.26 |

control 715.79
infinity 719.867

$$D(t) = \frac{Mt - M0}{M\infty - M0} *N$$

**IPTG**

| Time | Replicate 1 | Replicate 2 | Replicate 3 | Replicate 4 | Replicate5 | average | SD |
|---|---|---|---|---|---|---|---|
| 0 | 715.803 | 715.832 | | | | 715.82 | 0.02 |
| 30 | 715.937 | 715.924 | | | | 715.93 | 0.01 |
| 45 | 715.996 | 715.962 | | | | 715.98 | 0.02 |
| 60 | 715.963 | 716.005 | | | | 715.98 | 0.03 |
| 300 | 715.997 | 715.924 | | | | 715.96 | 0.05 |
| 1500 | 715.969 | 715.98 | | | | 715.97 | 0.01 |
| 3600 | 716.001 | 715.928 | | | | 715.96 | 0.05 |
| 7200 | 715.959 | 715.991 | | | | 715.98 | 0.02 |
| 14400 | 715.974 | 715.943 | | | | 715.96 | 0.02 |

**ONPF**

| Time | Replicate 1 | Replicate 2 | Replicate 3 | Replicate 4 | average | SD |
|---|---|---|---|---|---|---|
| 0 | 715.837 | 715.905 | | | 715.87 | 0.05 |
| 30 | 715.931 | 715.959 | | | 715.95 | 0.02 |
| 45 | 715.984 | 715.987 | | | 715.99 | 0.00 |
| 60 | 715.971 | 716.015 | | | 715.99 | 0.03 |
| 300 | 715.975 | 715.981 | | | 715.98 | 0.00 |
| 1500 | 715.989 | 715.971 | | | 715.98 | 0.01 |
| 3600 | 715.994 | 716.007 | | | 716.00 | 0.01 |
| 7200 | 715.937 | 715.97 | | | 715.95 | 0.02 |
| 14400 | 715.988 | 715.957 | | | 715.97 | 0.02 |

**APO**

| Time | Replicate 1 | Replicate 2 | Replicate 3 | Replicate 4 | Replicate 5 | average | SD |
|---|---|---|---|---|---|---|---|
| 0 | 715.743 | 715.845 | | | | 715.79 | 0.07 |
| 30 | 715.881 | 715.939 | | | | 715.91 | 0.04 |
| 45 | 715.901 | 716.003 | | | | 715.95 | 0.07 |
| 60 | 715.887 | 716.028 | | | | 715.96 | 0.10 |
| 300 | 715.886 | 715.961 | | | | 715.92 | 0.05 |
| 1500 | 715.897 | 715.941 | | | | 715.92 | 0.03 |
| 3600 | 715.825 | 715.98 | | | | 715.90 | 0.11 |
| 7200 | 715.807 | 715.95 | | | | 715.88 | 0.10 |
| 14400 | 715.899 | 715.942 | | | | 715.92 | 0.03 |

**DNA**

| Time | Replicate 1 | Replicate 2 | Replicate 3 | Replicate 4 | Replicate 5 | average | SD |
|---|---|---|---|---|---|---|---|
| 0 | 715.82 | 715.871 | | | | 715.85 | 0.04 |
| 30 | 715.956 | 715.932 | | | | 715.94 | 0.02 |
| 45 | 715.951 | 715.891 | | | | 715.92 | 0.04 |
| 60 | 715.967 | 715.927 | | | | 715.95 | 0.03 |
| 300 | 715.94 | 715.951 | | | | 715.95 | 0.01 |
| 1500 | 715.964 | 715.975 | | | | 715.97 | 0.01 |
| 3600 | 716.029 | 715.95 | | | | 715.99 | 0.06 |
| 7200 | 715.934 | 716.053 | | | | 715.99 | 0.08 |
| 14400 | 715.984 | 716.015 | | | | 716.00 | 0.02 |

**ONPFDNA**

| Time | Replicate 1 | Replicate 2 | Replicate 3 | Replicate 4 | average | SD |
|---|---|---|---|---|---|---|
| 0 | 715.854 | 715.921 | | | 715.89 | 0.05 |
| 30 | 715.962 | 716.004 | | | 715.98 | 0.03 |
| 45 | 715.959 | 715.906 | | | 715.93 | 0.04 |
| 60 | 715.975 | 715.985 | | | 715.98 | 0.01 |
| 300 | 715.968 | 716.12 | | | 716.04 | 0.11 |
| 1500 | 715.959 | 715.976 | | | 715.97 | 0.01 |
| 3600 | 715.956 | 715.963 | | | 715.96 | 0.00 |
| 7200 | 715.977 | 716.059 | | | 716.02 | 0.06 |
| 14400 | 716.007 | 715.981 | | | 715.99 | 0.02 |

**TMG**

| Time | Replicate 1 | Replicate 2 | Replicate 3 | Replicate 4 | Replicate 5 | average | SD |
|---|---|---|---|---|---|---|---|
| 0 | 715.825 | 715.796 | | | | 715.81 | 0.02 |
| 30 | 715.939 | 715.954 | | | | 715.95 | 0.01 |
| 45 | 715.992 | 715.906 | | | | 715.95 | 0.06 |
| 60 | 715.994 | 716.019 | | | | 716.01 | 0.02 |
| 300 | 716.013 | 715.995 | | | | 716.00 | 0.01 |
| 1500 | 716.022 | 715.969 | | | | 716.00 | 0.04 |
| 3600 | 715.971 | 715.951 | | | | 715.96 | 0.01 |
| 7200 | 715.967 | 716.007 | | | | 715.99 | 0.03 |
| 14400 | 716.026 | 715.985 | | | | 716.01 | 0.03 |

### 123-129 IINYPLD

**Charge** 1

**IPTG**

| Time | centroid | D |
|---|---|---|
| 0 | 847.95 | 0.00 |
| 30 | 848.09 | 0.19 |
| 45 | 848.09 | 0.21 |
| 60 | 848.09 | 0.20 |
| 300 | 848.11 | 0.23 |
| 1500 | 848.15 | 0.29 |
| 3600 | 848.13 | 0.27 |
| 7200 | 848.27 | 0.46 |
| 14400 | 848.37 | 0.62 |

**ONPF**

| Time | centroid | D |
|---|---|---|
| 0 | 847.95 | 0.00 |
| 30 | 848.05 | 0.14 |
| 45 | 848.10 | 0.21 |
| 60 | 848.10 | 0.21 |
| 300 | 848.15 | 0.29 |
| 1500 | 848.23 | 0.41 |
| 3600 | 848.28 | 0.49 |
| 7200 | 848.35 | 0.58 |
| 14400 | 848.30 | 0.52 |

**APO**

| Time | centroid | D |
|---|---|---|
| 0 | 847.96 | 0.00 |
| 30 | 848.06 | 0.16 |
| 45 | 848.06 | 0.15 |
| 60 | 848.09 | 0.21 |
| 300 | 848.10 | 0.22 |
| 1500 | 848.22 | 0.40 |
| 3600 | 848.36 | 0.61 |
| 7200 | 848.41 | 0.68 |
| 14400 | 848.51 | 0.83 |

**DNA**

| Time | centroid | D |
|---|---|---|
| 0 | 847.97 | 0.01 |
| 30 | 848.06 | 0.16 |
| 45 | 848.09 | 0.19 |
| 60 | 848.10 | 0.22 |
| 300 | 848.09 | 0.20 |
| 1500 | 848.20 | 0.37 |
| 3600 | 848.26 | 0.45 |
| 7200 | 848.30 | 0.51 |
| 14400 | 848.23 | 0.42 |

**ONPFDNA**

| Time | centroid | D |
|---|---|---|
| 0 | 847.95 | -0.02 |
| 30 | 848.04 | 0.13 |
| 45 | 848.06 | 0.15 |
| 60 | 848.11 | 0.24 |
| 300 | 848.09 | 0.20 |
| 1500 | 848.18 | 0.34 |
| 3600 | 848.33 | 0.56 |
| 7200 | 848.35 | 0.59 |
| 14400 | 848.24 | 0.42 |

**TMG**

| Time | centroid | D |
|---|---|---|
| 0 | 847.98 | 0.03 |
| 30 | 848.11 | 0.23 |
| 45 | 848.11 | 0.23 |
| 60 | 848.11 | 0.23 |
| 300 | 848.13 | 0.26 |
| 1500 | 848.13 | 0.26 |
| 3600 | 848.20 | 0.36 |
| 7200 | 848.27 | 0.48 |
| 14400 | 848.39 | 0.66 |

| | |
|---|---|
| control | 847.96 |
| infinity | 850.616 |

$$D(t) = \frac{M_t - M_0}{M_\infty - M_0} \cdot N$$

**IPTG**

| Time | Replicate 1 | Replicate 2 | Replicate 3 | Replicate 4 | Replicate5 | average | SD |
|---|---|---|---|---|---|---|---|
| 0 | 847.941 | 847.968 | | | | 847.95 | 0.02 |
| 30 | 848.106 | 848.064 | | | | 848.09 | 0.03 |
| 45 | 848.111 | 848.078 | | | | 848.09 | 0.02 |
| 60 | 848.117 | 848.066 | | | | 848.09 | 0.04 |
| 300 | 848.151 | 848.069 | | | | 848.11 | 0.06 |
| 1500 | 848.169 | 848.132 | | | | 848.15 | 0.03 |
| 3600 | 848.08 | 848.187 | | | | 848.13 | 0.08 |
| 7200 | 848.307 | 848.224 | | | | 848.27 | 0.06 |
| 14400 | 848.421 | 848.31 | | | | 848.37 | 0.08 |

**ONPF**

| Time | Replicate 1 | Replicate 2 | Replicate 3 | Replicate 4 | average | SD |
|---|---|---|---|---|---|---|
| 0 | 847.941 | 847.968 | | | 847.95 | 0.02 |
| 30 | 848.04 | 848.06 | | | 848.05 | 0.01 |
| 45 | 848.106 | 848.084 | | | 848.10 | 0.02 |
| 60 | 848.083 | 848.109 | | | 848.10 | 0.02 |
| 300 | 848.18 | 848.123 | | | 848.15 | 0.04 |
| 1500 | 848.229 | 848.224 | | | 848.23 | 0.00 |
| 3600 | 848.294 | 848.268 | | | 848.28 | 0.02 |
| 7200 | 848.368 | 848.322 | | | 848.35 | 0.03 |
| 14400 | 848.432 | 848.172 | | | 848.30 | 0.18 |

**APO**

| Time | Replicate 1 | Replicate 2 | Replicate 3 | Replicate 4 | Replicate 5 | average | SD |
|---|---|---|---|---|---|---|---|
| 0 | 847.958 | 847.955 | | | | 847.96 | 0.00 |
| 30 | 848.07 | 848.056 | | | | 848.06 | 0.01 |
| 45 | 848.054 | 848.065 | | | | 848.06 | 0.01 |
| 60 | 848.093 | 848.093 | | | | 848.09 | 0.00 |
| 300 | 848.113 | 848.093 | | | | 848.10 | 0.01 |
| 1500 | 848.221 | 848.226 | | | | 848.22 | 0.00 |
| 3600 | 848.4 | 848.328 | | | | 848.36 | 0.05 |
| 7200 | 848.47 | 848.345 | | | | 848.41 | 0.09 |
| 14400 | 848.607 | 848.41 | | | | 848.51 | 0.14 |

**DNA**

| Time | Replicate 1 | Replicate 2 | Replicate 3 | Replicate 4 | Replicate 5 | average | SD |
|---|---|---|---|---|---|---|---|
| 0 | 847.983 | 847.949 | | | | 847.97 | 0.02 |
| 30 | 848.081 | 848.046 | | | | 848.06 | 0.02 |
| 45 | 848.111 | 848.06 | | | | 848.09 | 0.04 |
| 60 | 848.089 | 848.112 | | | | 848.10 | 0.02 |
| 300 | 848.054 | 848.119 | | | | 848.09 | 0.05 |
| 1500 | 848.176 | 848.233 | | | | 848.20 | 0.04 |
| 3600 | 848.286 | 848.225 | | | | 848.26 | 0.04 |
| 7200 | 848.346 | 848.244 | | | | 848.30 | 0.07 |
| 14400 | 848.41 | 848.056 | | | | 848.23 | 0.25 |

**ONPFDNA**

| Time | Replicate 1 | Replicate 2 | Replicate 3 | Replicate 4 | average | SD |
|---|---|---|---|---|---|---|
| 0 | 847.932 | 847.958 | | | 847.95 | 0.02 |
| 30 | 848.037 | 848.043 | | | 848.04 | 0.00 |
| 45 | 848.04 | 848.075 | | | 848.06 | 0.02 |
| 60 | 848.086 | 848.142 | | | 848.11 | 0.04 |
| 300 | 848.074 | 848.099 | | | 848.09 | 0.02 |
| 1500 | 848.139 | 848.226 | | | 848.18 | 0.06 |
| 3600 | 848.304 | 848.354 | | | 848.33 | 0.04 |
| 7200 | 848.356 | 848.34 | | | 848.35 | 0.01 |
| 14400 | 848.216 | 848.261 | | | 848.24 | 0.03 |

**TMG**

| Time | Replicate 1 | Replicate 2 | Replicate 3 | Replicate 4 | Replicate 5 | average | SD |
|---|---|---|---|---|---|---|---|
| 0 | 848.009 | 847.945 | | | | 847.98 | 0.05 |
| 30 | 848.085 | 848.134 | | | | 848.11 | 0.03 |
| 45 | 848.12 | 848.102 | | | | 848.11 | 0.01 |
| 60 | 848.108 | | | | | 848.11 | |
| 300 | 848.131 | 848.131 | | | | 848.13 | 0.00 |
| 1500 | 848.084 | 848.172 | | | | 848.13 | 0.06 |
| 3600 | 848.2 | 848.196 | | | | 848.20 | 0.00 |
| 7200 | 848.252 | 848.295 | | | | 848.27 | 0.03 |
| 14400 | 848.395 | 848.394 | | | | 848.39 | 0.00 |

### 128-135 LDDQDAIA

**Charge** 1

**IPTG**

| Time | centroid | D |
|---|---|---|
| 0 | 860.89 | 0.04 |
| 30 | 861.7285 | 1.66 |
| 45 | 861.852 | 1.90 |
| 60 | 861.8565 | 1.90 |
| 300 | 861.801 | 1.80 |
| 1500 | 861.882 | 1.95 |
| 3600 | 862.0635 | 2.30 |
| 7200 | 862.224 | 2.61 |
| 14400 | 862.3005 | 2.76 |

**ONPF**

| Time | centroid | D |
|---|---|---|
| 0 | 860.92 | 0.09 |
| 30 | 861.50 | 1.22 |
| 45 | 861.71 | 1.63 |
| 60 | 861.73 | 1.66 |
| 300 | 861.68 | 1.56 |
| 1500 | 861.79 | 1.77 |
| 3600 | 861.91 | 2.01 |
| 7200 | 861.92 | 2.03 |
| 14400 | 862.40 | 2.95 |

**APO**

| Time | centroid | D |
|---|---|---|
| 0 | 860.869 | 0.00 |
| 30 | 861.67 | 1.54 |
| 45 | 861.711 | 1.62 |
| 60 | 861.84 | 1.87 |
| 300 | 861.883 | 1.96 |
| 1500 | 861.935 | 2.06 |
| 3600 | 862.042 | 2.26 |
| 7200 | 862.133 | 2.44 |
| 14400 | 862.404 | 2.96 |

**DNA**

| Time | centroid | D |
|---|---|---|
| 0 | 860.8615 | -0.01 |
| 30 | 861.5845 | 1.38 |
| 45 | 861.766 | 1.73 |
| 60 | 861.814 | 1.82 |
| 300 | 861.7705 | 1.74 |
| 1500 | 861.815 | 1.82 |
| 3600 | 861.872 | 1.93 |
| 7200 | 861.8635 | 1.92 |
| 14400 | 861.837 | 1.87 |

**ONPFDNA**

| Time | centroid | D |
|---|---|---|
| 0 | 860.868 | 0.00 |
| 30 | 861.566 | 1.34 |
| 45 | 861.7505 | 1.70 |
| 60 | 861.7005 | 1.60 |
| 300 | 861.725 | 1.65 |
| 1500 | 861.6295 | 1.47 |
| 3600 | 861.89 | 1.97 |
| 7200 | 861.877 | 1.94 |
| 14400 | 861.6275 | 1.46 |

**TMG**

| Time | centroid | D |
|---|---|---|
| 0 | 860.88 | 0.03 |
| 30 | 861.77 | 1.73 |
| 45 | 861.90 | 1.98 |
| 60 | 861.84 | 1.87 |
| 300 | 861.93 | 2.04 |
| 1500 | 861.92 | 2.03 |
| 3600 | 862.14 | 2.45 |
| 7200 | 862.23 | 2.62 |
| 14400 | 862.45 | 3.05 |

control 860.869
infinity 863.98

$$D(t) = \frac{Mt - M0}{M\infty - M0} * N$$

**IPTG**

| Time | Replicate 1 | Replicate 2 | Replicate 3 | Replicate 4 | Replicate5 | average | SD |
|---|---|---|---|---|---|---|---|
| 0 | 860.925 | 860.855 | | | | 860.89 | 0.05 |
| 30 | 861.918 | 861.539 | | | | 861.73 | 0.27 |
| 45 | 861.963 | 861.741 | | | | 861.85 | 0.16 |
| 60 | 861.978 | 861.735 | | | | 861.86 | 0.17 |
| 300 | 861.957 | 861.645 | | | | 861.80 | 0.22 |
| 1500 | 862.07 | 861.694 | | | | 861.88 | 0.27 |
| 3600 | 862.176 | 861.951 | | | | 862.06 | 0.16 |
| 7200 | 862.414 | 862.034 | | | | 862.22 | 0.27 |
| 14400 | 862.516 | 862.085 | | | | 862.30 | 0.30 |

**ONPF**

| Time | Replicate 1 | Replicate 2 | Replicate 3 | Replicate 4 | average | SD |
|---|---|---|---|---|---|---|
| 0 | 860.913 | 860.919 | | | 860.92 | 0.00 |
| 30 | 861.629 | 861.37 | | | 861.50 | 0.18 |
| 45 | 861.969 | 861.458 | | | 861.71 | 0.36 |
| 60 | 861.806 | 861.657 | | | 861.73 | 0.11 |
| 300 | 861.852 | 861.506 | | | 861.68 | 0.24 |
| 1500 | 861.931 | 861.647 | | | 861.79 | 0.20 |
| 3600 | 862.101 | 861.719 | | | 861.91 | 0.27 |
| 7200 | 862.058 | 861.781 | | | 861.92 | 0.20 |
| 14400 | 862.398 | | | | 862.40 | |

**APO**

| Time | Replicate 1 | Replicate 2 | Replicate 3 | Replicate 4 | Replicate 5 | average | SD |
|---|---|---|---|---|---|---|---|
| 0 | 860.868 | 860.87 | | | | 860.87 | 0.00 |
| 30 | 861.834 | 861.506 | | | | 861.67 | 0.23 |
| 45 | 861.897 | 861.525 | | | | 861.71 | 0.26 |
| 60 | 861.995 | 861.685 | | | | 861.84 | 0.22 |
| 300 | 861.883 | | | | | 861.88 | |
| 1500 | 861.994 | 861.876 | | | | 861.94 | 0.08 |
| 3600 | 862.186 | 861.898 | | | | 862.04 | 0.20 |
| 7200 | 862.329 | 861.937 | | | | 862.13 | 0.28 |
| 14400 | 862.562 | 862.246 | | | | 862.40 | 0.22 |

**DNA**

| Time | Replicate 1 | Replicate 2 | Replicate 3 | Replicate 4 | Replicate 5 | average | SD |
|---|---|---|---|---|---|---|---|
| 0 | 860.86 | 860.863 | | | | 860.86 | 0.00 |
| 30 | 861.642 | 861.527 | | | | 861.58 | 0.08 |
| 45 | 861.922 | 861.61 | | | | 861.77 | 0.22 |
| 60 | 861.967 | 861.661 | | | | 861.81 | 0.22 |
| 300 | 861.896 | 861.645 | | | | 861.77 | 0.18 |
| 1500 | 861.91 | 861.72 | | | | 861.82 | 0.13 |
| 3600 | 862.065 | 861.679 | | | | 861.87 | 0.27 |
| 7200 | 862.074 | 861.653 | | | | 861.86 | 0.30 |
| 14400 | 862.223 | 861.451 | | | | 861.84 | 0.55 |

**ONPFDNA**

| Time | Replicate 1 | Replicate 2 | Replicate 3 | Replicate 4 | average | SD |
|---|---|---|---|---|---|---|
| 0 | 860.874 | 860.862 | | | 860.87 | 0.01 |
| 30 | 861.593 | 861.539 | | | 861.57 | 0.04 |
| 45 | 861.828 | 861.673 | | | 861.75 | 0.11 |
| 60 | 861.756 | 861.645 | | | 861.70 | 0.08 |
| 300 | 861.838 | 861.612 | | | 861.73 | 0.16 |
| 1500 | 861.53 | 861.729 | | | 861.63 | 0.14 |
| 3600 | 861.961 | 861.819 | | | 861.89 | 0.10 |
| 7200 | 861.947 | 861.807 | | | 861.88 | 0.10 |
| 14400 | 861.516 | 861.739 | | | 861.63 | 0.16 |

**TMG**

| Time | Replicate 1 | Replicate 2 | Replicate 3 | Replicate 4 | Replicate 5 | average | SD |
|---|---|---|---|---|---|---|---|
| 0 | 860.902 | 860.865 | | | | 860.88 | 0.03 |
| 30 | 861.574 | 861.963 | | | | 861.77 | 0.28 |
| 45 | 861.871 | 861.924 | | | | 861.90 | 0.04 |
| 60 | 861.891 | 861.785 | | | | 861.84 | 0.07 |
| 300 | 861.858 | 862 | | | | 861.93 | 0.10 |
| 1500 | 861.796 | 862.044 | | | | 861.92 | 0.18 |
| 3600 | 862.071 | 862.212 | | | | 862.14 | 0.10 |
| 7200 | 862.099 | 862.361 | | | | 862.23 | 0.19 |
| 14400 | 862.394 | 862.507 | | | | 862.45 | 0.08 |

### 134-139 IAVEAA

**Charge** 1

**IPTG**

| Time | centroid | D |
|---|---|---|
| 0 | 573.63 | 0.01 |
| 30 | 573.76 | 0.14 |
| 45 | 573.77 | 0.15 |
| 60 | 573.80 | 0.18 |
| 300 | 573.88 | 0.26 |
| 1500 | 574.17 | 0.55 |
| 3600 | 574.35 | 0.73 |
| 7200 | 574.74 | 1.12 |
| 14400 | 574.70 | 1.09 |

**ONPF**

| Time | centroid | D |
|---|---|---|
| 0 | 573.64 | 0.02 |
| 30 | 573.74 | 0.12 |
| 45 | 573.95 | 0.33 |
| 60 | 573.85 | 0.23 |
| 300 | 573.84 | 0.22 |
| 1500 | 574.05 | 0.43 |
| 3600 | 574.17 | 0.55 |
| 7200 | 574.16 | 0.54 |
| 14400 | 574.16 | 0.54 |

**APO**

| Time | centroid | D |
|---|---|---|
| 0 | 573.62 | 0.00 |
| 30 | 573.71 | 0.09 |
| 45 | 573.74 | 0.12 |
| 60 | 573.86 | 0.24 |
| 300 | 573.84 | 0.22 |
| 1500 | 574.17 | 0.56 |
| 3600 | 574.29 | 0.68 |
| 7200 | 574.38 | 0.77 |
| 14400 | 574.56 | 0.95 |

**DNA**

| Time | centroid | D |
|---|---|---|
| 0 | 573.61 | -0.01 |
| 30 | 573.77 | 0.16 |
| 45 | 573.98 | 0.36 |
| 60 | 573.83 | 0.22 |
| 300 | 573.82 | 0.20 |
| 1500 | 573.93 | 0.31 |
| 3600 | 573.97 | 0.35 |
| 7200 | 574.05 | 0.43 |
| 14400 | 574.26 | 0.64 |

**ONPFDNA**

| Time | centroid | D |
|---|---|---|
| 0 | 573.62 | 0.00 |
| 30 | 573.70 | 0.08 |
| 45 | 573.77 | 0.15 |
| 60 | 573.83 | 0.21 |
| 300 | 573.79 | 0.17 |
| 1500 | 573.93 | 0.31 |
| 3600 | 574.06 | 0.44 |
| 7200 | 574.06 | 0.44 |
| 14400 | 574.03 | 0.42 |

**TMG**

| Time | centroid | D |
|---|---|---|
| 0 | 573.63 | 0.01 |
| 30 | 573.76 | 0.14 |
| 45 | 573.88 | 0.26 |
| 60 | 573.86 | 0.24 |
| 300 | 573.89 | 0.27 |
| 1500 | 574.12 | 0.50 |
| 3600 | 574.38 | 0.77 |
| 7200 | 574.42 | 0.81 |
| 14400 | 574.69 | 1.08 |

| | |
|---|---|
| control | 573.62 |
| infinity | 577.591 |

$$D(t) = \frac{M_t - M_0}{M_\infty - M_0} \cdot N$$

**IPTG**

| Time | Replicate 1 | Replicate 2 | Replicate 3 | Replicate 4 | Replicate5 | average | SD |
|---|---|---|---|---|---|---|---|
| 0 | 573.632 | 573.621 | | | | 573.63 | 0.01 |
| 30 | 573.786 | 573.727 | | | | 573.76 | 0.04 |
| 45 | 573.787 | 573.747 | | | | 573.77 | 0.03 |
| 60 | 573.785 | 573.813 | | | | 573.80 | 0.02 |
| 300 | 573.895 | 573.859 | | | | 573.88 | 0.03 |
| 1500 | 574.223 | 574.109 | | | | 574.17 | 0.08 |
| 3600 | 574.404 | 574.294 | | | | 574.35 | 0.08 |
| 7200 | 574.6 | 574.871 | | | | 574.74 | 0.19 |
| 14400 | 574.912 | 574.484 | | | | 574.70 | 0.30 |

**ONPF**

| Time | Replicate 1 | Replicate 2 | Replicate 3 | Replicate 4 | average | SD |
|---|---|---|---|---|---|---|
| 0 | 573.644 | 573.628 | | | 573.64 | 0.01 |
| 30 | 573.784 | 573.69 | | | 573.74 | 0.07 |
| 45 | 573.894 | 573.998 | | | 573.95 | 0.07 |
| 60 | 573.856 | 573.842 | | | 573.85 | 0.01 |
| 300 | 573.887 | 573.789 | | | 573.84 | 0.07 |
| 1500 | 574.102 | 573.992 | | | 574.05 | 0.08 |
| 3600 | 574.301 | 574.033 | | | 574.17 | 0.19 |
| 7200 | 574.197 | 574.119 | | | 574.16 | 0.06 |
| 14400 | 574.626 | 573.692 | | | 574.16 | 0.66 |

**APO**

| Time | Replicate 1 | Replicate 2 | Replicate 3 | Replicate 4 | Replicate 5 | average | SD |
|---|---|---|---|---|---|---|---|
| 0 | 573.625 | 573.616 | | | | 573.62 | 0.01 |
| 30 | 573.723 | 573.699 | | | | 573.71 | 0.02 |
| 45 | 573.753 | 573.723 | | | | 573.74 | 0.02 |
| 60 | 573.753 | 573.963 | | | | 573.86 | 0.15 |
| 300 | 573.869 | 573.804 | | | | 573.84 | 0.05 |
| 1500 | 574.125 | 574.219 | | | | 574.17 | 0.07 |
| 3600 | 574.381 | 574.204 | | | | 574.29 | 0.13 |
| 7200 | 574.532 | 574.235 | | | | 574.38 | 0.21 |
| 14400 | 574.663 | 574.457 | | | | 574.56 | 0.15 |

**DNA**

| Time | Replicate 1 | Replicate 2 | Replicate 3 | Replicate 4 | Replicate 5 | average | SD |
|---|---|---|---|---|---|---|---|
| 0 | 573.611 | 573.604 | | | | 573.61 | 0.00 |
| 30 | 573.774 | | | | | 573.77 | |
| 45 | 574.197 | 573.754 | | | | 573.98 | 0.31 |
| 60 | 573.827 | 573.841 | | | | 573.83 | 0.01 |
| 300 | 573.858 | 573.772 | | | | 573.82 | 0.06 |
| 1500 | 573.853 | 574.005 | | | | 573.93 | 0.11 |
| 3600 | 574.007 | 573.926 | | | | 573.97 | 0.06 |
| 7200 | 574.096 | 574.001 | | | | 574.05 | 0.07 |
| 14400 | 574.335 | 574.179 | | | | 574.26 | 0.11 |

**ONPFDNA**

| Time | Replicate 1 | Replicate 2 | Replicate 3 | Replicate 4 | average | SD |
|---|---|---|---|---|---|---|
| 0 | 573.616 | 573.622 | | | 573.62 | 0.00 |
| 30 | 573.714 | 573.693 | | | 573.70 | 0.01 |
| 45 | 573.824 | 573.712 | | | 573.77 | 0.08 |
| 60 | 573.806 | 573.857 | | | 573.83 | 0.04 |
| 300 | 573.803 | 573.782 | | | 573.79 | 0.01 |
| 1500 | 573.833 | 574.023 | | | 573.93 | 0.13 |
| 3600 | 574.032 | 574.086 | | | 574.06 | 0.04 |
| 7200 | 574.02 | 574.102 | | | 574.06 | 0.06 |
| 14400 | 573.994 | 574.071 | | | 574.03 | 0.05 |

**TMG**

| Time | Replicate 1 | Replicate 2 | Replicate 3 | Replicate 4 | Replicate 5 | average | SD |
|---|---|---|---|---|---|---|---|
| 0 | 573.635 | 573.634 | | | | 573.63 | 0.00 |
| 30 | 573.739 | 573.779 | | | | 573.76 | 0.03 |
| 45 | 573.953 | 573.801 | | | | 573.88 | 0.11 |
| 60 | 573.861 | 573.864 | | | | 573.86 | 0.00 |
| 300 | 573.872 | 573.907 | | | | 573.89 | 0.02 |
| 1500 | 574.101 | 574.133 | | | | 574.12 | 0.02 |
| 3600 | 574.394 | 574.368 | | | | 574.38 | 0.02 |
| 7200 | 574.337 | 574.509 | | | | 574.42 | 0.12 |
| 14400 | 574.661 | 574.714 | | | | 574.69 | 0.04 |

### 139-146 ACTNVPAL

**Charge** 1

**IPTG**

| Time | centroid | D |
|---|---|---|
| 0 | 788.912 | 0.02 |
| 30 | 790.1225 | 1.79 |
| 45 | 790.226 | 1.94 |
| 60 | 790.327 | 2.09 |
| 300 | 790.4145 | 2.21 |
| 1500 | 790.5 | 2.34 |
| 3600 | 790.7925 | 2.77 |
| 7200 | 790.7875 | 2.76 |
| 14400 | 790.6875 | 2.61 |

**ONPF**

| Time | centroid | D |
|---|---|---|
| 0 | 788.9965 | 0.14 |
| 30 | 789.5825 | 1.00 |
| 45 | 789.971 | 1.57 |
| 60 | 790.1425 | 1.82 |
| 300 | 790.1555 | 1.84 |
| 1500 | 790.3315 | 2.09 |
| 3600 | 790.251 | 1.98 |
| 7200 | 790.255 | 1.98 |
| 14400 | 790.68 | 2.60 |

**APO**

| Time | centroid | D |
|---|---|---|
| 0 | 788.901 | 0.00 |
| 30 | 789.9475 | 1.53 |
| 45 | 790.0405 | 1.67 |
| 60 | 790.325 | 2.08 |
| 300 | 790.2835 | 2.02 |
| 1500 | 790.5545 | 2.42 |
| 3600 | 790.741 | 2.69 |
| 7200 | 790.481 | 2.31 |
| 14400 | 790.715 | 2.65 |

**DNA**

| Time | centroid | D |
|---|---|---|
| 0 | 788.907 | 0.01 |
| 30 | 789.774 | 1.28 |
| 45 | 790.1375 | 1.81 |
| 60 | 790.21 | 1.92 |
| 300 | 790.3365 | 2.10 |
| 1500 | 790.541 | 2.40 |
| 3600 | 790.5125 | 2.36 |
| 7200 | 790.3055 | 2.06 |
| 14400 | 790.337 | 2.10 |

**ONPFDNA**

| Time | centroid | D |
|---|---|---|
| 0 | 788.907 | 0.01 |
| 30 | 789.7045 | 1.18 |
| 45 | 790.0915 | 1.74 |
| 60 | 790.1665 | 1.85 |
| 300 | 790.208 | 1.91 |
| 1500 | 790.193 | 1.89 |
| 3600 | 790.629 | 2.53 |
| 7200 | 790.3325 | 2.09 |
| 14400 | 790.16 | 1.84 |

**TMG**

| Time | centroid | D |
|---|---|---|
| 0 | 788.9155 | 0.02 |
| 30 | 790.02 | 1.64 |
| 45 | 790.2085 | 1.91 |
| 60 | 790.202 | 1.90 |
| 300 | 790.573 | 2.45 |
| 1500 | 790.585 | 2.46 |
| 3600 | 790.707 | 2.64 |
| 7200 | 790.6715 | 2.59 |
| 14400 | 790.873 | 2.89 |

control 788.901
infinity 792.318

$$D(t) = \frac{M_t - M_0}{M_\infty - M_0} * N$$

**IPTG**

| Time | Replicate 1 | Replicate 2 | Replicate 3 | Replicate 4 | Replicate5 | average | SD |
|---|---|---|---|---|---|---|---|
| 0 | 788.892 | 788.932 | | | | 788.91 | 0.03 |
| 30 | 790.347 | 789.898 | | | | 790.12 | 0.32 |
| 45 | 790.214 | 790.238 | | | | 790.23 | 0.02 |
| 60 | 790.322 | 790.332 | | | | 790.33 | 0.01 |
| 300 | 790.542 | 790.287 | | | | 790.41 | 0.18 |
| 1500 | 790.678 | 790.322 | | | | 790.50 | 0.25 |
| 3600 | 790.856 | 790.729 | | | | 790.79 | 0.09 |
| 7200 | 790.985 | 790.59 | | | | 790.79 | 0.28 |
| 14400 | 790.884 | 790.491 | | | | 790.69 | 0.28 |

**ONPF**

| Time | Replicate 1 | Replicate 2 | Replicate 3 | Replicate 4 | average | SD |
|---|---|---|---|---|---|---|
| 0 | 788.993 | 789 | | | 789.00 | 0.00 |
| 30 | 789.631 | 789.534 | | | 789.58 | 0.07 |
| 45 | 790.215 | 789.727 | | | 789.97 | 0.35 |
| 60 | 790.174 | 790.111 | | | 790.14 | 0.04 |
| 300 | 790.364 | 789.947 | | | 790.16 | 0.29 |
| 1500 | 790.508 | 790.155 | | | 790.33 | 0.25 |
| 3600 | 790.453 | 790.049 | | | 790.25 | 0.29 |
| 7200 | 790.357 | 790.153 | | | 790.26 | 0.14 |
| 14400 | 790.675 | | | | 790.68 | |

**APO**

| Time | Replicate 1 | Replicate 2 | Replicate 3 | Replicate 4 | Replicate 5 | average | SD |
|---|---|---|---|---|---|---|---|
| 0 | 788.875 | 788.927 | | | | 788.90 | 0.04 |
| 30 | 790.14 | 789.755 | | | | 789.95 | 0.27 |
| 45 | 790.255 | 789.826 | | | | 790.04 | 0.30 |
| 60 | 790.434 | 790.216 | | | | 790.33 | 0.15 |
| 300 | 790.573 | 789.994 | | | | 790.28 | 0.41 |
| 1500 | 790.654 | 790.455 | | | | 790.55 | 0.14 |
| 3600 | 790.825 | 790.657 | | | | 790.74 | 0.12 |
| 7200 | 790.754 | 790.208 | | | | 790.48 | 0.39 |
| 14400 | 790.9 | 790.53 | | | | 790.72 | 0.26 |

**DNA**

| Time | Replicate 1 | Replicate 2 | Replicate 3 | Replicate 4 | Replicate 5 | average | SD |
|---|---|---|---|---|---|---|---|
| 0 | 788.876 | 788.938 | | | | 788.91 | 0.04 |
| 30 | 789.775 | 789.773 | | | | 789.77 | 0.00 |
| 45 | 790.273 | 790.002 | | | | 790.14 | 0.19 |
| 60 | 790.215 | 790.205 | | | | 790.21 | 0.01 |
| 300 | 790.375 | 790.298 | | | | 790.34 | 0.05 |
| 1500 | 790.561 | 790.521 | | | | 790.54 | 0.03 |
| 3600 | 790.676 | 790.349 | | | | 790.51 | 0.23 |
| 7200 | 790.553 | 790.058 | | | | 790.31 | 0.35 |
| 14400 | 790.774 | 789.9 | | | | 790.34 | 0.62 |

**ONPFDNA**

| Time | Replicate 1 | Replicate 2 | Replicate 3 | Replicate 4 | average | SD |
|---|---|---|---|---|---|---|
| 0 | 788.901 | 788.913 | | | 788.91 | 0.01 |
| 30 | 789.654 | 789.755 | | | 789.70 | 0.07 |
| 45 | 790.089 | 790.094 | | | 790.09 | 0.00 |
| 60 | 790.137 | 790.196 | | | 790.17 | 0.04 |
| 300 | 790.208 | 790.208 | | | 790.21 | 0.00 |
| 1500 | 790.166 | 790.22 | | | 790.19 | 0.04 |
| 3600 | 790.645 | 790.613 | | | 790.63 | 0.02 |
| 7200 | 790.356 | 790.309 | | | 790.33 | 0.03 |
| 14400 | 790.215 | 790.105 | | | 790.16 | 0.08 |

**TMG**

| Time | Replicate 1 | Replicate 2 | Replicate 3 | Replicate 4 | Replicate 5 | average | SD |
|---|---|---|---|---|---|---|---|
| 0 | 788.952 | 788.879 | | | | 788.92 | 0.05 |
| 30 | 789.723 | 790.317 | | | | 790.02 | 0.42 |
| 45 | 790.217 | 790.2 | | | | 790.21 | 0.01 |
| 60 | 790.277 | 790.127 | | | | 790.20 | 0.11 |
| 300 | 790.486 | 790.66 | | | | 790.57 | 0.12 |
| 1500 | 790.387 | 790.783 | | | | 790.59 | 0.28 |
| 3600 | 790.562 | 790.852 | | | | 790.71 | 0.21 |
| 7200 | 790.364 | 790.979 | | | | 790.67 | 0.43 |
| 14400 | 790.692 | 791.054 | | | | 790.87 | 0.26 |

### 145-149 ALFLD

**Charge** 1

**IPTG**

| Time | centroid | D |
|---|---|---|
| 0 | 578.692 | 0.02 |
| 30 | 578.8005 | 0.20 |
| 45 | 578.8225 | 0.23 |
| 60 | 578.8655 | 0.30 |
| 300 | 578.813 | 0.22 |
| 1500 | 578.8455 | 0.27 |
| 3600 | 578.8535 | 0.28 |
| 7200 | 578.872 | 0.31 |
| 14400 | 578.878 | 0.32 |

**ONPF**

| Time | centroid | D |
|---|---|---|
| 0 | 578.704 | 0.04 |
| 30 | 578.807 | 0.21 |
| 45 | 578.813 | 0.22 |
| 60 | 578.819 | 0.23 |
| 300 | 578.766 | 0.14 |
| 1500 | 578.839 | 0.26 |
| 3600 | 578.824 | 0.23 |
| 7200 | 578.893 | 0.35 |
| 14400 | 579.03 | 0.56 |

**APO**

| Time | centroid | D |
|---|---|---|
| 0 | 578.675 | -0.01 |
| 30 | 578.787 | 0.17 |
| 45 | 578.8055 | 0.20 |
| 60 | 578.797 | 0.19 |
| 300 | 578.8025 | 0.20 |
| 1500 | 578.815 | 0.22 |
| 3600 | 578.8355 | 0.25 |
| 7200 | 578.848 | 0.27 |
| 14400 | 578.9625 | 0.46 |

**DNA**

| Time | centroid | D |
|---|---|---|
| 0 | 578.667 | -0.02 |
| 30 | 578.8005 | 0.20 |
| 45 | 578.8745 | 0.32 |
| 60 | 578.8385 | 0.26 |
| 300 | 578.838 | 0.26 |
| 1500 | 578.8715 | 0.31 |
| 3600 | 578.853 | 0.28 |
| 7200 | 578.826 | 0.24 |
| 14400 | 578.8545 | 0.28 |

**ONPFDNA**

| Time | centroid | D |
|---|---|---|
| 0 | 578.686 | 0.01 |
| 30 | 578.8195 | 0.23 |
| 45 | 578.887 | 0.34 |
| 60 | 578.841 | 0.26 |
| 300 | 578.9315 | 0.41 |
| 1500 | 578.833 | 0.25 |
| 3600 | 578.847 | 0.27 |
| 7200 | 578.852 | 0.28 |
| 14400 | 578.836 | 0.25 |

**TMG**

| Time | centroid | D |
|---|---|---|
| 0 | 578.7005 | 0.03 |
| 30 | 578.824 | 0.23 |
| 45 | 578.8945 | 0.35 |
| 60 | 578.9155 | 0.38 |
| 300 | 578.8985 | 0.36 |
| 1500 | 578.913 | 0.38 |
| 3600 | 578.8685 | 0.31 |
| 7200 | 578.924 | 0.40 |
| 14400 | 578.909 | 0.37 |

control 578.68
infinity 580.526

$$D(t) = \frac{M_t - M_0}{M_\infty - M_0} \cdot N$$

**IPTG**

| Time | Replicate 1 | Replicate 2 | Replicate 3 | Replicate 4 | Replicate5 | average | SD |
|---|---|---|---|---|---|---|---|
| 0 | 578.708 | 578.676 | | | | 578.69 | 0.02 |
| 30 | 578.824 | 578.777 | | | | 578.80 | 0.03 |
| 45 | 578.86 | 578.785 | | | | 578.82 | 0.05 |
| 60 | 578.905 | 578.826 | | | | 578.87 | 0.06 |
| 300 | 578.851 | 578.775 | | | | 578.81 | 0.05 |
| 1500 | 578.87 | 578.821 | | | | 578.85 | 0.03 |
| 3600 | 578.92 | 578.787 | | | | 578.85 | 0.09 |
| 7200 | 578.923 | 578.821 | | | | 578.87 | 0.07 |
| 14400 | 578.952 | 578.804 | | | | 578.88 | 0.10 |

**ONPF**

| Time | Replicate 1 | Replicate 2 | Replicate 3 | Replicate 4 | average | SD |
|---|---|---|---|---|---|---|
| 0 | 578.712 | 578.696 | | | 578.70 | 0.01 |
| 30 | 578.845 | 578.769 | | | 578.81 | 0.05 |
| 45 | 578.844 | 578.782 | | | 578.81 | 0.04 |
| 60 | 578.833 | 578.805 | | | 578.82 | 0.02 |
| 300 | 578.779 | 578.753 | | | 578.77 | 0.02 |
| 1500 | 578.857 | 578.821 | | | 578.84 | 0.03 |
| 3600 | 578.837 | 578.811 | | | 578.82 | 0.02 |
| 7200 | 578.999 | 578.787 | | | 578.89 | 0.15 |
| 14400 | 579.025 | | | | 579.03 | |

**APO**

| Time | Replicate 1 | Replicate 2 | Replicate 3 | Replicate 4 | Replicate 5 | average | SD |
|---|---|---|---|---|---|---|---|
| 0 | 578.676 | 578.674 | | | | 578.68 | 0.00 |
| 30 | 578.8 | 578.774 | | | | 578.79 | 0.02 |
| 45 | 578.823 | 578.788 | | | | 578.81 | 0.02 |
| 60 | 578.789 | 578.805 | | | | 578.80 | 0.01 |
| 300 | 578.805 | 578.8 | | | | 578.80 | 0.00 |
| 1500 | 578.817 | 578.813 | | | | 578.82 | 0.00 |
| 3600 | 578.836 | 578.835 | | | | 578.84 | 0.00 |
| 7200 | 578.879 | 578.817 | | | | 578.85 | 0.04 |
| 14400 | 579.039 | 578.886 | | | | 578.96 | 0.11 |

**DNA**

| Time | Replicate 1 | Replicate 2 | Replicate 3 | Replicate 4 | Replicate 5 | average | SD |
|---|---|---|---|---|---|---|---|
| 0 | 578.668 | 578.666 | | | | 578.67 | 0.00 |
| 30 | 578.844 | 578.757 | | | | 578.80 | 0.06 |
| 45 | 578.967 | 578.782 | | | | 578.87 | 0.13 |
| 60 | 578.868 | 578.809 | | | | 578.84 | 0.04 |
| 300 | 578.888 | 578.788 | | | | 578.84 | 0.07 |
| 1500 | 578.913 | 578.83 | | | | 578.87 | 0.06 |
| 3600 | 578.91 | 578.796 | | | | 578.85 | 0.08 |
| 7200 | 578.831 | 578.821 | | | | 578.83 | 0.01 |
| 14400 | 578.883 | 578.826 | | | | 578.85 | 0.04 |

**ONPFDNA**

| Time | Replicate 1 | Replicate 2 | Replicate 3 | Replicate 4 | average | SD |
|---|---|---|---|---|---|---|
| 0 | 578.695 | 578.677 | | | 578.69 | 0.01 |
| 30 | 578.854 | 578.785 | | | 578.82 | 0.05 |
| 45 | | 578.887 | | | 578.89 | |
| 60 | 578.899 | 578.783 | | | 578.84 | 0.08 |
| 300 | 579.086 | 578.777 | | | 578.93 | 0.22 |
| 1500 | 578.868 | 578.798 | | | 578.83 | 0.05 |
| 3600 | 578.923 | 578.771 | | | 578.85 | 0.11 |
| 7200 | 578.923 | 578.781 | | | 578.85 | 0.10 |
| 14400 | 578.935 | 578.737 | | | 578.84 | 0.14 |

**TMG**

| Time | Replicate 1 | Replicate 2 | Replicate 3 | Replicate 4 | Replicate 5 | average | SD |
|---|---|---|---|---|---|---|---|
| 0 | 578.724 | 578.677 | | | | 578.70 | 0.03 |
| 30 | 578.829 | 578.819 | | | | 578.82 | 0.01 |
| 45 | 578.973 | 578.816 | | | | 578.89 | 0.11 |
| 60 | 578.885 | 578.946 | | | | 578.92 | 0.04 |
| 300 | 578.921 | 578.876 | | | | 578.90 | 0.03 |
| 1500 | 578.919 | 578.907 | | | | 578.91 | 0.01 |
| 3600 | 578.904 | 578.833 | | | | 578.87 | 0.05 |
| 7200 | 578.977 | 578.871 | | | | 578.92 | 0.07 |
| 14400 | 578.839 | 578.979 | | | | 578.91 | 0.10 |

### 147-158 FLDVSDQTPINS

**Charge** 2

**IPTG**

| Time | centroid | D |
|---|---|---|
| 0 | 668.7248 | -0.02 |
| 30 | 669.2168 | 2.08 |
| 45 | 669.435 | 3.01 |
| 60 | 669.4768 | 3.19 |
| 300 | 669.593 | 3.69 |
| 1500 | 669.8428 | 4.76 |
| 3600 | 669.8855 | 4.94 |
| 7200 | 670.0165 | 5.50 |
| 14400 | 670.032 | 5.57 |

**ONPF**

| Time | centroid | D |
|---|---|---|
| 0 | 668.72 | -0.04 |
| 30 | 669.05 | 1.38 |
| 45 | 669.27 | 2.29 |
| 60 | 669.46 | 3.10 |
| 300 | 669.46 | 3.12 |
| 1500 | 669.81 | 4.62 |
| 3600 | 669.83 | 4.68 |
| 7200 | 669.84 | 4.73 |
| 14400 | 669.94 | 5.18 |

**APO**

| Time | centroid | D |
|---|---|---|
| 0 | 668.7327 | 0.01 |
| 30 | 669.2023 | 2.02 |
| 45 | | |
| 60 | 669.4883 | 3.24 |
| 300 | 669.632 | 3.86 |
| 1500 | 669.969 | 5.30 |
| 3600 | 669.9667 | 5.29 |
| 7200 | 670.0923 | 5.82 |
| 14400 | 670.1827 | 6.21 |

**DNA**

| Time | centroid | D |
|---|---|---|
| 0 | 668.72 | -0.04 |
| 30 | 669.18 | 1.91 |
| 45 | | |
| 60 | 669.61 | 3.76 |
| 300 | 669.67 | 4.02 |
| 1500 | 669.88 | 4.92 |
| 3600 | 669.93 | 5.11 |
| 7200 | 670.00 | 5.43 |
| 14400 | 670.02 | 5.51 |

**ONPFDNA**

| Time | centroid | D |
|---|---|---|
| 0 | 668.72 | -0.05 |
| 30 | 669.11 | 1.61 |
| 45 | 669.50 | 3.28 |
| 60 | 669.51 | 3.33 |
| 300 | 669.68 | 4.06 |
| 1500 | 669.68 | 4.08 |
| 3600 | 669.96 | 5.25 |
| 7200 | 669.90 | 5.01 |
| 14400 | 669.81 | 4.61 |

**TMG**

| Time | centroid | D |
|---|---|---|
| 0 | 668.72 | -0.04 |
| 30 | 669.24 | 2.17 |
| 45 | 669.43 | 3.00 |
| 60 | 669.62 | 3.78 |
| 300 | 669.71 | 4.21 |
| 1500 | 669.93 | 5.15 |
| 3600 | 670.10 | 5.84 |
| 7200 | 670.20 | 6.30 |
| 14400 | 670.50 | 7.55 |

control 668.73
infinity 670.835

$$D(t) = \frac{Mt - M0}{M\infty - M0} *N$$

**IPTG**

| Time | Replicate 1 | Replicate 2 | Replicate 3 | Replicate 4 | Replicate5 | average | SD |
|---|---|---|---|---|---|---|---|
| 0 | 668.74 | 668.726 | 668.716 | 668.717 | | 668.72 | 0.01 |
| 30 | 669.406 | 669.225 | 669.124 | 669.112 | | 669.22 | 0.14 |
| 45 | 669.435 | | | | | 669.44 | |
| 60 | 669.503 | 669.642 | 669.384 | 669.378 | | 669.48 | 0.12 |
| 300 | 669.682 | 669.695 | 669.57 | 669.425 | | 669.59 | 0.13 |
| 1500 | 670.133 | 669.815 | 669.77 | 669.653 | | 669.84 | 0.21 |
| 3600 | 670.208 | 669.839 | 669.808 | 669.687 | | 669.89 | 0.22 |
| 7200 | 670.392 | 669.935 | 669.918 | 669.821 | | 670.02 | 0.26 |
| 14400 | 670.325 | 669.969 | 670.027 | 669.807 | | 670.03 | 0.22 |

**ONPF**

| Time | Replicate 1 | Replicate 2 | Replicate 3 | Replicate 4 | average | SD |
|---|---|---|---|---|---|---|
| 0 | 668.767 | 668.671 | 668.723 | 668.717 | 668.72 | 0.04 |
| 30 | 669.097 | 668.879 | 669.178 | 669.055 | 669.05 | 0.13 |
| 45 | 669.544 | 668.988 | | | 669.27 | 0.39 |
| 60 | 669.443 | 669.256 | 669.704 | 669.421 | 669.46 | 0.19 |
| 300 | 669.662 | 669.259 | 669.485 | 669.435 | 669.46 | 0.17 |
| 1500 | 670.063 | 669.532 | 669.878 | 669.772 | 669.81 | 0.22 |
| 3600 | 670.044 | 669.491 | 670.003 | 669.764 | 669.83 | 0.25 |
| 7200 | 670.195 | 669.396 | 669.942 | 669.813 | 669.84 | 0.33 |
| 14400 | 670.434 | 669.19 | 670.066 | 670.074 | 669.94 | 0.53 |

**APO**

| Time | Replicate 1 | Replicate 2 | Replicate 3 | Replicate 4 | Replicate 5 | average | SD |
|---|---|---|---|---|---|---|---|
| 0 | 668.775 | 668.707 | 668.716 | | | 668.73 | 0.04 |
| 30 | 669.324 | 669.167 | 669.116 | | | 669.20 | 0.11 |
| 45 | | | | | | | |
| 60 | 669.645 | 669.41 | 669.41 | | | 669.49 | 0.14 |
| 300 | 669.688 | 669.659 | 669.549 | | | 669.63 | 0.07 |
| 1500 | 669.948 | 670.144 | 669.815 | | | 669.97 | 0.17 |
| 3600 | 670.079 | 669.97 | 669.851 | | | 669.97 | 0.11 |
| 7200 | 670.245 | 670.097 | 669.935 | | | 670.09 | 0.16 |
| 14400 | 670.171 | 670.178 | 670.199 | | | 670.18 | 0.01 |

**DNA**

| Time | Replicate 1 | Replicate 2 | Replicate 3 | Replicate 4 | Replicate 5 | average | SD |
|---|---|---|---|---|---|---|---|
| 0 | 668.723 | 668.722 | 668.72 | | | 668.72 | 0.00 |
| 30 | 669.244 | 669.17 | 669.115 | | | 669.18 | 0.06 |
| 45 | | | | | | | |
| 60 | 669.542 | 669.801 | 669.483 | | | 669.61 | 0.17 |
| 300 | 669.767 | 669.72 | 669.522 | | | 669.67 | 0.13 |
| 1500 | 670.03 | 669.935 | 669.675 | | | 669.88 | 0.18 |
| 3600 | 670.109 | 669.945 | 669.722 | | | 669.93 | 0.19 |
| 7200 | 670.15 | 670.135 | 669.712 | | | 670.00 | 0.25 |
| 14400 | 670.144 | 670.096 | 669.814 | | | 670.02 | 0.18 |

**ONPFDNA**

| Time | Replicate 1 | Replicate 2 | Replicate 3 | Replicate 4 | average | SD |
|---|---|---|---|---|---|---|
| 0 | 668.722 | 668.713 | | | 668.72 | 0.01 |
| 30 | 669.193 | 669.02 | | | 669.11 | 0.12 |
| 45 | 669.496 | | | | 669.50 | |
| 60 | 669.525 | 669.494 | | | 669.51 | 0.02 |
| 300 | 669.75 | 669.61 | | | 669.68 | 0.10 |
| 1500 | 669.624 | 669.745 | | | 669.68 | 0.09 |
| 3600 | 670.078 | 669.838 | | | 669.96 | 0.17 |
| 7200 | 669.951 | 669.851 | | | 669.90 | 0.07 |
| 14400 | 669.719 | 669.896 | | | 669.81 | 0.13 |

**TMG**

| Time | Replicate 1 | Replicate 2 | Replicate 3 | Replicate 4 | Replicate 5 | average | SD |
|---|---|---|---|---|---|---|---|
| 0 | 668.738 | 668.704 | | | | 668.72 | 0.02 |
| 30 | 669.131 | 669.342 | | | | 669.24 | 0.15 |
| 45 | 669.489 | 669.375 | | | | 669.43 | 0.08 |
| 60 | 669.427 | 669.803 | | | | 669.62 | 0.27 |
| 300 | 669.701 | 669.727 | | | | 669.71 | 0.02 |
| 1500 | 669.79 | 670.078 | | | | 669.93 | 0.20 |
| 3600 | 669.976 | 670.214 | | | | 670.10 | 0.17 |
| 7200 | 670.04 | 670.369 | | | | 670.20 | 0.23 |
| 14400 | 670.423 | 670.571 | | | | 670.50 | 0.10 |

### 152-157 DQTPIN

**Charge** 1

**IPTG**

| Time | centroid | D |
|---|---|---|
| 0 | 687.698 | -0.01 |
| 30 | 688.054 | 0.46 |
| 45 | 688.1125 | 0.54 |
| 60 | 688.1975 | 0.65 |
| 300 | 688.6055 | 1.19 |
| 1500 | 688.8225 | 1.47 |
| 3600 | 689.0975 | 1.83 |
| 7200 | 688.942 | 1.63 |
| 14400 | 688.8225 | 1.47 |

**ONPF**

| Time | centroid | D |
|---|---|---|
| 0 | 687.7565 | 0.07 |
| 30 | 687.955 | 0.33 |
| 45 | 688.125 | 0.55 |
| 60 | 688.272 | 0.75 |
| 300 | 688.5025 | 1.05 |
| 1500 | 688.6785 | 1.28 |
| 3600 | 688.6785 | 1.28 |
| 7200 | 688.6445 | 1.24 |
| 14400 | 689.00 | 1.70 |

**APO**

| Time | centroid | D |
|---|---|---|
| 0 | 687.705 | 0.00 |
| 30 | 688.026 | 0.42 |
| 45 | 688.073 | 0.48 |
| 60 | 688.317 | 0.81 |
| 300 | 688.5865 | 1.16 |
| 1500 | 688.884 | 1.55 |
| 3600 | 688.9345 | 1.62 |
| 7200 | 688.8825 | 1.55 |
| 14400 | 689.054 | 1.78 |

**DNA**

| Time | centroid | D |
|---|---|---|
| 0 | 687.7015 | 0.00 |
| 30 | 688.0265 | 0.42 |
| 45 | 688.2465 | 0.71 |
| 60 | 688.3415 | 0.84 |
| 300 | 688.6115 | 1.19 |
| 1500 | 688.762 | 1.39 |
| 3600 | 688.7375 | 1.36 |
| 7200 | 688.6 | 1.18 |
| 14400 | 688.546 | 1.11 |

**ONPFDNA**

| Time | centroid | D |
|---|---|---|
| 0 | 687.712 | 0.01 |
| 30 | 687.969 | 0.35 |
| 45 | 688.1585 | 0.60 |
| 60 | 688.276 | 0.75 |
| 300 | 688.546 | 1.11 |
| 1500 | 688.5 | 1.05 |
| 3600 | 688.81 | 1.45 |
| 7200 | 688.6845 | 1.29 |
| 14400 | 688.4395 | 0.97 |

**TMG**

| Time | centroid | D |
|---|---|---|
| 0 | 687.709 | 0.01 |
| 30 | 688.068 | 0.48 |
| 45 | 688.1525 | 0.59 |
| 60 | 688.3755 | 0.88 |
| 300 | 688.69 | 1.30 |
| 1500 | 688.884 | 1.55 |
| 3600 | 688.9775 | 1.68 |
| 7200 | 688.899 | 1.57 |
| 14400 | 689.0735 | 1.80 |

control 687.705
infinity 689.984

$$D(t) = \frac{Mt - M0}{M\infty - M0} *N$$

**IPTG**

| Time | Replicate 1 | Replicate 2 | Replicate 3 | Replicate 4 | Replicate5 | average | SD |
|---|---|---|---|---|---|---|---|
| 0 | 687.718 | 687.678 | | | | 687.70 | 0.03 |
| 30 | 688.17 | 687.938 | | | | 688.05 | 0.16 |
| 45 | 688.186 | 688.039 | | | | 688.11 | 0.10 |
| 60 | 688.257 | 688.138 | | | | 688.20 | 0.08 |
| 300 | 688.665 | 688.546 | | | | 688.61 | 0.08 |
| 1500 | 689.066 | 688.579 | | | | 688.82 | 0.34 |
| 3600 | 689.224 | 688.971 | | | | 689.10 | 0.18 |
| 7200 | 689.08 | 688.804 | | | | 688.94 | 0.20 |
| 14400 | 688.965 | 688.68 | | | | 688.82 | 0.20 |

**ONPF**

| Time | Replicate 1 | Replicate 2 | Replicate 3 | Replicate 4 | average | SD |
|---|---|---|---|---|---|---|
| 0 | 687.723 | 687.79 | | | 687.76 | 0.05 |
| 30 | 687.986 | 687.924 | | | 687.96 | 0.04 |
| 45 | 688.249 | 688.001 | | | 688.13 | 0.18 |
| 60 | 688.252 | 688.292 | | | 688.27 | 0.03 |
| 300 | 688.677 | 688.328 | | | 688.50 | 0.25 |
| 1500 | 688.86 | 688.497 | | | 688.68 | 0.26 |
| 3600 | 688.81 | 688.547 | | | 688.68 | 0.19 |
| 7200 | 688.752 | 688.537 | | | 688.64 | 0.15 |
| 14400 | 688.996 | | | | 689.00 | |

**APO**

| Time | Replicate 1 | Replicate 2 | Replicate 3 | Replicate 4 | Replicate 5 | average | SD |
|---|---|---|---|---|---|---|---|
| 0 | 687.7 | 687.71 | | | | 687.71 | 0.01 |
| 30 | 688.041 | 688.011 | | | | 688.03 | 0.02 |
| 45 | 688.096 | 688.05 | | | | 688.07 | 0.03 |
| 60 | 688.267 | 688.367 | | | | 688.32 | 0.07 |
| 300 | 688.77 | 688.403 | | | | 688.59 | 0.26 |
| 1500 | 688.896 | 688.872 | | | | 688.88 | 0.02 |
| 3600 | 689.043 | 688.826 | | | | 688.93 | 0.15 |
| 7200 | 689.05 | 688.715 | | | | 688.88 | 0.24 |
| 14400 | 689.185 | 688.923 | | | | 689.05 | 0.19 |

**DNA**

| Time | Replicate 1 | Replicate 2 | Replicate 3 | Replicate 4 | Replicate 5 | average | SD |
|---|---|---|---|---|---|---|---|
| 0 | 687.685 | 687.718 | | | | 687.70 | 0.02 |
| 30 | 688.04 | 688.013 | | | | 688.03 | 0.02 |
| 45 | 688.405 | 688.088 | | | | 688.25 | 0.22 |
| 60 | 688.33 | 688.353 | | | | 688.34 | 0.02 |
| 300 | 688.651 | 688.572 | | | | 688.61 | 0.06 |
| 1500 | 688.734 | 688.79 | | | | 688.76 | 0.04 |
| 3600 | 688.862 | 688.613 | | | | 688.74 | 0.18 |
| 7200 | 688.738 | 688.462 | | | | 688.60 | 0.20 |
| 14400 | 688.879 | 688.213 | | | | 688.55 | 0.47 |

**ONPFDNA**

| Time | Replicate 1 | Replicate 2 | Replicate 3 | Replicate 4 | average | SD |
|---|---|---|---|---|---|---|
| 0 | 687.693 | 687.731 | | | 687.71 | 0.03 |
| 30 | 688.004 | 687.934 | | | 687.97 | 0.05 |
| 45 | 688.213 | 688.104 | | | 688.16 | 0.08 |
| 60 | 688.257 | 688.295 | | | 688.28 | 0.03 |
| 300 | 688.63 | 688.462 | | | 688.55 | 0.12 |
| 1500 | 688.463 | 688.537 | | | 688.50 | 0.05 |
| 3600 | 688.803 | 688.817 | | | 688.81 | 0.01 |
| 7200 | 688.723 | 688.646 | | | 688.68 | 0.05 |
| 14400 | 688.466 | 688.413 | | | 688.44 | 0.04 |

**TMG**

| Time | Replicate 1 | Replicate 2 | Replicate 3 | Replicate 4 | Replicate 5 | average | SD |
|---|---|---|---|---|---|---|---|
| 0 | 687.72 | 687.698 | | | | 687.71 | 0.02 |
| 30 | 688.019 | 688.117 | | | | 688.07 | 0.07 |
| 45 | 688.248 | 688.057 | | | | 688.15 | 0.14 |
| 60 | 688.224 | 688.527 | | | | 688.38 | 0.21 |
| 300 | 688.678 | 688.702 | | | | 688.69 | 0.02 |
| 1500 | 688.727 | 689.041 | | | | 688.88 | 0.22 |
| 3600 | 688.878 | 689.077 | | | | 688.98 | 0.14 |
| 7200 | 688.714 | 689.084 | | | | 688.90 | 0.26 |
| 14400 | 689.006 | 689.141 | | | | 689.07 | 0.10 |

### 158-163 SIIFSH

**Charge** 1

**IPTG**

| Time | centroid | D |
|---|---|---|
| 0 | 703.80 | -0.10 |
| 30 | 703.98 | 0.21 |
| 45 | 704.00 | 0.24 |
| 60 | 704.03 | 0.30 |
| 300 | 704.00 | 0.24 |
| 1500 | 704.02 | 0.28 |
| 3600 | 704.01 | 0.25 |
| 7200 | 704.02 | 0.28 |
| 14400 | 704.09 | 0.39 |

**ONPF**

| Time | centroid | D |
|---|---|---|
| 0 | 703.83 | -0.05 |
| 30 | 703.88 | 0.04 |
| 45 | 704.13 | 0.45 |
| 60 | 704.10 | 0.40 |
| 300 | 704.09 | 0.40 |
| 1500 | 704.04 | 0.31 |
| 3600 | 704.33 | 0.78 |
| 7200 | 704.19 | 0.55 |
| 14400 | 704.63 | 1.28 |

**APO**

| Time | centroid | D |
|---|---|---|
| 0 | 703.85 | 0.00 |
| 30 | 703.97 | 0.19 |
| 45 | 704.00 | 0.23 |
| 60 | 704.05 | 0.33 |
| 300 | 704.06 | 0.35 |
| 1500 | 704.13 | 0.46 |
| 3600 | 704.21 | 0.59 |
| 7200 | 704.33 | 0.78 |
| 14400 | 704.68 | 1.36 |

**DNA**

| Time | centroid | D |
|---|---|---|
| 0 | 703.93 | 0.12 |
| 30 | 703.97 | 0.19 |
| 45 | 704.00 | 0.24 |
| 60 | 704.05 | 0.32 |
| 300 | 704.07 | 0.36 |
| 1500 | 704.11 | 0.42 |
| 3600 | 704.18 | 0.54 |
| 7200 | 704.15 | 0.49 |
| 14400 | 704.21 | 0.60 |

**ONPFDNA**

| Time | centroid | D |
|---|---|---|
| 0 | 704.02 | 0.27 |
| 30 | 704.14 | 0.47 |
| 45 | 704.12 | 0.44 |
| 60 | 704.11 | 0.43 |
| 300 | 704.21 | 0.59 |
| 1500 | 704.21 | 0.58 |
| 3600 | 704.19 | 0.55 |
| 7200 | 704.30 | 0.73 |
| 14400 | 704.30 | 0.73 |

**TMG**

| Time | centroid | D |
|---|---|---|
| 0 | 703.97 | 0.20 |
| 30 | 704.13 | 0.45 |
| 45 | 704.11 | 0.42 |
| 60 | 704.25 | 0.65 |
| 300 | 704.10 | 0.41 |
| 1500 | 704.15 | 0.49 |
| 3600 | 704.13 | 0.46 |
| 7200 | 704.17 | 0.52 |
| 14400 | 704.30 | 0.74 |

control 703.854
infinity 706.272

$$D(t) = \frac{M_t - M_0}{M_\infty - M_0} * N$$

**IPTG**

| Time | Replicate 1 | Replicate 2 | Replicate 3 | Replicate 4 | Replicate5 | average | SD |
|---|---|---|---|---|---|---|---|
| 0 | 703.794 | 703.798 | | | | 703.80 | 0.00 |
| 30 | 704.076 | 703.889 | | | | 703.98 | 0.13 |
| 45 | 704.064 | 703.93 | | | | 704.00 | 0.09 |
| 60 | 704.136 | 703.932 | | | | 704.03 | 0.14 |
| 300 | 704.082 | 703.922 | | | | 704.00 | 0.11 |
| 1500 | 704.078 | 703.963 | | | | 704.02 | 0.08 |
| 3600 | 704.062 | 703.952 | | | | 704.01 | 0.08 |
| 7200 | 704.103 | 703.94 | | | | 704.02 | 0.12 |
| 14400 | 704.179 | 704.005 | | | | 704.09 | 0.12 |

**ONPF**

| Time | Replicate 1 | Replicate 2 | Replicate 3 | Replicate 4 | average | SD |
|---|---|---|---|---|---|---|
| 0 | 703.826 | | | | 703.83 | |
| 30 | 703.88 | | | | 703.88 | |
| 45 | 704.125 | | | | 704.13 | |
| 60 | 704.095 | | | | 704.10 | |
| 300 | 704.094 | | | | 704.09 | |
| 1500 | 704.039 | | | | 704.04 | |
| 3600 | 704.328 | | | | 704.33 | |
| 7200 | 704.188 | | | | 704.19 | |
| 14400 | 704.626 | | | | 704.63 | |

**APO**

| Time | Replicate 1 | Replicate 2 | Replicate 3 | Replicate 4 | Replicate 5 | average | SD |
|---|---|---|---|---|---|---|---|
| 0 | 703.815 | 703.893 | | | | 703.85 | 0.06 |
| 30 | 704.009 | 703.923 | | | | 703.97 | 0.06 |
| 45 | 704.067 | 703.925 | | | | 704.00 | 0.10 |
| 60 | 704.105 | 704.003 | | | | 704.05 | 0.07 |
| 300 | 704.063 | | | | | 704.06 | |
| 1500 | 704.127 | 704.135 | | | | 704.13 | 0.01 |
| 3600 | 704.284 | 704.135 | | | | 704.21 | 0.11 |
| 7200 | 704.467 | 704.184 | | | | 704.33 | 0.20 |
| 14400 | 704.784 | 704.569 | | | | 704.68 | 0.15 |

**DNA**

| Time | Replicate 1 | Replicate 2 | Replicate 3 | Replicate 4 | Replicate 5 | average | SD |
|---|---|---|---|---|---|---|---|
| 0 | 703.953 | 703.906 | | | | 703.93 | 0.03 |
| 30 | 704.049 | 703.886 | | | | 703.97 | 0.12 |
| 45 | 704.115 | 703.887 | | | | 704.00 | 0.16 |
| 60 | 704.136 | 703.962 | | | | 704.05 | 0.12 |
| 300 | 704.072 | 704.075 | | | | 704.07 | 0.00 |
| 1500 | 704.217 | 703.997 | | | | 704.11 | 0.16 |
| 3600 | 704.294 | 704.061 | | | | 704.18 | 0.16 |
| 7200 | 704.181 | 704.123 | | | | 704.15 | 0.04 |
| 14400 | 704.42 | 704.009 | | | | 704.21 | 0.29 |

**ONPFDNA**

| Time | Replicate 1 | Replicate 2 | Replicate 3 | Replicate 4 | average | SD |
|---|---|---|---|---|---|---|
| 0 | 704.135 | 703.902 | | | 704.02 | 0.16 |
| 30 | 704.247 | 704.028 | | | 704.14 | 0.15 |
| 45 | 704.323 | 703.922 | | | 704.12 | 0.28 |
| 60 | 704.263 | 703.966 | | | 704.11 | 0.21 |
| 300 | 704.369 | 704.058 | | | 704.21 | 0.22 |
| 1500 | 704.36 | 704.055 | | | 704.21 | 0.22 |
| 3600 | 704.423 | 703.95 | | | 704.19 | 0.33 |
| 7200 | 704.511 | 704.082 | | | 704.30 | 0.30 |
| 14400 | 704.295 | | | | 704.30 | |

**TMG**

| Time | Replicate 1 | Replicate 2 | Replicate 3 | Replicate 4 | Replicate 5 | average | SD |
|---|---|---|---|---|---|---|---|
| 0 | 704.104 | 703.841 | | | | 703.97 | 0.19 |
| 30 | 704.104 | 704.146 | | | | 704.13 | 0.03 |
| 45 | 704.106 | 704.106 | | | | 704.11 | 0.00 |
| 60 | 704.061 | 704.435 | | | | 704.25 | 0.26 |
| 300 | 704.049 | 704.149 | | | | 704.10 | 0.07 |
| 1500 | 704.067 | 704.228 | | | | 704.15 | 0.11 |
| 3600 | 704.1 | 704.159 | | | | 704.13 | 0.04 |
| 7200 | 704.157 | 704.182 | | | | 704.17 | 0.02 |
| 14400 | 704.258 | 704.345 | | | | 704.30 | 0.06 |

### 158-169 SIIFSHEDGTRL

**Charge** 3

**IPTG**

| Time | centroid | D |
|---|---|---|
| 0 | 459.1918 | 0.26 |
| 30 | 459.2418 | 0.67 |
| 45 | 459.269 | 0.89 |
| 60 | 459.263 | 0.84 |
| 300 | 459.277 | 0.95 |
| 1500 | 459.2662 | 0.87 |
| 3600 | 459.2762 | 0.95 |
| 7200 | 459.297 | 1.12 |
| 14400 | 459.3056 | 1.19 |

**ONPF**

| Time | centroid | D |
|---|---|---|
| 0 | 459.16 | -0.02 |
| 30 | 459.23 | 0.61 |
| 45 | 459.23 | 0.55 |
| 60 | 459.24 | 0.65 |
| 300 | 459.24 | 0.67 |
| 1500 | 459.28 | 0.95 |
| 3600 | 459.31 | 1.22 |
| 7200 | 459.33 | 1.39 |
| 14400 | 459.39 | 1.89 |

**APO**

| Time | centroid | D |
|---|---|---|
| 0 | 459.1638 | 0.03 |
| 30 | 459.216 | 0.46 |
| 45 | 459.252 | 0.75 |
| 60 | 459.2683 | 0.88 |
| 300 | 459.285 | 1.02 |
| 1500 | 459.3224 | 1.32 |
| 3600 | 459.35 | 1.55 |
| 7200 | 459.4662 | 2.50 |
| 14400 | 459.5616 | 3.28 |

**DNA**

| Time | centroid | D |
|---|---|---|
| 0 | 459.158 | -0.02 |
| 30 | 459.2117 | 0.42 |
| 45 | 459.2435 | 0.68 |
| 60 | 459.234 | 0.60 |
| 300 | 459.2263 | 0.54 |
| 1500 | 459.2443 | 0.69 |
| 3600 | 459.256 | 0.78 |
| 7200 | 459.2593 | 0.81 |
| 14400 | 459.2773 | 0.96 |

**ONPFDNA**

| Time | centroid | D |
|---|---|---|
| 0 | 459.1607 | 0.01 |
| 30 | 459.204 | 0.36 |
| 45 | 459.229 | 0.56 |
| 60 | 459.2337 | 0.60 |
| 300 | 459.2323 | 0.59 |
| 1500 | 459.224 | 0.52 |
| 3600 | 459.252 | 0.75 |
| 7200 | 459.2647 | 0.85 |
| 14400 | 459.2373 | 0.63 |

**TMG**

| Time | centroid | D |
|---|---|---|
| 0 | 459.16 | 0.04 |
| 30 | 459.18 | 0.13 |
| 45 | 459.29 | 1.04 |
| 60 | 459.44 | 2.24 |
| 300 | 459.26 | 0.84 |
| 1500 | 459.30 | 1.13 |
| 3600 | 459.31 | 1.22 |
| 7200 | 459.27 | 0.91 |
| 14400 | 459.36 | 1.62 |

| | |
|---|---|
| control | 459.16 |
| infinity | 460.386 |

$$D(t) = \frac{Mt - M0}{M\infty - M0} * N$$

**IPTG**

| Time | Replicate 1 | Replicate 2 | Replicate 3 | Replicate 4 | Replicate5 | average | SD |
|---|---|---|---|---|---|---|---|
| 0 | 459.241 | 459.209 | 459.168 | 459.163 | 459.178 | 459.19 | 0.03 |
| 30 | 459.213 | 459.3 | 459.215 | 459.203 | 459.278 | 459.24 | 0.04 |
| 45 | | 459.253 | | | 459.285 | 459.27 | 0.02 |
| 60 | | 459.268 | 459.243 | 459.23 | 459.311 | 459.26 | 0.04 |
| 300 | 459.394 | 459.21 | 459.233 | 459.232 | 459.316 | 459.28 | 0.08 |
| 1500 | 459.369 | 459.217 | 459.23 | 459.227 | 459.288 | 459.27 | 0.06 |
| 3600 | 459.386 | 459.263 | 459.233 | 459.243 | 459.256 | 459.28 | 0.06 |
| 7200 | 459.43 | 459.259 | 459.258 | 459.248 | 459.29 | 459.30 | 0.08 |
| 14400 | 459.439 | 459.248 | 459.26 | 459.256 | 459.325 | 459.31 | 0.08 |

**ONPF**

| Time | Replicate 1 | Replicate 2 | Replicate 3 | Replicate 4 | average | SD |
|---|---|---|---|---|---|---|
| 0 | 459.169 | 459.165 | 459.138 | 459.159 | 459.16 | 0.01 |
| 30 | 459.222 | 459.185 | 459.28 | 459.251 | 459.23 | 0.04 |
| 45 | 459.227 | | | | 459.23 | |
| 60 | 459.203 | 459.211 | 459.268 | 459.278 | 459.24 | 0.04 |
| 300 | 459.249 | 459.213 | 459.215 | 459.292 | 459.24 | 0.04 |
| 1500 | 459.274 | 459.258 | 459.265 | 459.311 | 459.28 | 0.02 |
| 3600 | 459.252 | 459.261 | 459.331 | 459.395 | 459.31 | 0.07 |
| 7200 | 459.278 | 459.305 | 459.297 | 459.442 | 459.33 | 0.08 |
| 14400 | 459.207 | 459.355 | 459.426 | 459.579 | 459.39 | 0.15 |

**APO**

| Time | Replicate 1 | Replicate 2 | Replicate 3 | Replicate 4 | Replicate 5 | average | SD |
|---|---|---|---|---|---|---|---|
| 0 | 459.201 | 459.145 | 459.163 | 459.149 | 459.161 | 459.16 | 0.02 |
| 30 | | 459.234 | 459.212 | 459.146 | 459.272 | 459.22 | 0.05 |
| 45 | | 459.225 | | | 459.279 | 459.25 | 0.04 |
| 60 | | 459.266 | 459.261 | 459.267 | 459.279 | 459.27 | 0.01 |
| 300 | 459.403 | 459.24 | 459.237 | | 459.26 | 459.29 | 0.08 |
| 1500 | 459.438 | 459.351 | 459.283 | 459.298 | 459.242 | 459.32 | 0.08 |
| 3600 | 459.491 | 459.345 | 459.312 | 459.248 | 459.354 | 459.35 | 0.09 |
| 7200 | 459.608 | 459.393 | 459.449 | 459.357 | 459.524 | 459.47 | 0.10 |
| 14400 | 459.67 | 459.599 | 459.477 | 459.347 | 459.715 | 459.56 | 0.15 |

**DNA**

| Time | Replicate 1 | Replicate 2 | Replicate 3 | Replicate 4 | Replicate 5 | average | SD |
|---|---|---|---|---|---|---|---|
| 0 | 459.164 | 459.154 | 459.156 | | | 459.16 | 0.01 |
| 30 | 459.222 | 459.166 | 459.247 | | | 459.21 | 0.04 |
| 45 | 459.222 | | 459.265 | | | 459.24 | 0.03 |
| 60 | 459.242 | 459.176 | 459.284 | | | 459.23 | 0.05 |
| 300 | 459.23 | 459.178 | 459.271 | | | 459.23 | 0.05 |
| 1500 | 459.269 | 459.181 | 459.283 | | | 459.24 | 0.06 |
| 3600 | 459.258 | 459.186 | 459.324 | | | 459.26 | 0.07 |
| 7200 | 459.282 | 459.171 | 459.325 | | | 459.26 | 0.08 |
| 14400 | 459.241 | 459.182 | 459.409 | | | 459.28 | 0.12 |

**ONPFDNA**

| Time | Replicate 1 | Replicate 2 | Replicate 3 | Replicate 4 | average | SD |
|---|---|---|---|---|---|---|
| 0 | 459.161 | 459.157 | 459.164 | | 459.16 | 0.00 |
| 30 | 459.217 | 459.16 | 459.235 | | 459.20 | 0.04 |
| 45 | 459.229 | | | | 459.23 | |
| 60 | 459.24 | 459.19 | 459.271 | | 459.23 | 0.04 |
| 300 | 459.222 | 459.171 | 459.304 | | 459.23 | 0.07 |
| 1500 | 459.231 | 459.173 | 459.268 | | 459.22 | 0.05 |
| 3600 | 459.245 | 459.182 | 459.329 | | 459.25 | 0.07 |
| 7200 | 459.243 | 459.184 | 459.367 | | 459.26 | 0.09 |
| 14400 | 459.237 | 459.191 | 459.284 | | 459.24 | 0.05 |

**TMG**

| Time | Replicate 1 | Replicate 2 | Replicate 3 | Replicate 4 | Replicate 5 | average | SD |
|---|---|---|---|---|---|---|---|
| 0 | 459.161 | 459.168 | | | | 459.16 | 0.00 |
| 30 | 459.073 | 459.279 | | | | 459.18 | 0.15 |
| 45 | 459.291 | 459.284 | | | | 459.29 | 0.00 |
| 60 | 459.319 | 459.551 | | | | 459.44 | 0.16 |
| 300 | 459.222 | 459.304 | | | | 459.26 | 0.06 |
| 1500 | 459.248 | 459.349 | | | | 459.30 | 0.07 |
| 3600 | 459.307 | 459.311 | | | | 459.31 | 0.00 |
| 7200 | 459.223 | 459.32 | | | | 459.27 | 0.07 |
| 14400 | 459.318 | 459.398 | | | | 459.36 | 0.06 |

### 164-173 EDGTRLGVEH

**Charge** 2

IPTG

| Time | centroid | D |
|---|---|---|
| 0 | 557.0895 | 0.04 |
| 30 | 557.2945 | 0.82 |
| 45 | 557.2205 | 0.54 |
| 60 | 557.29 | 0.81 |
| 300 | 557.2795 | 0.77 |
| 1500 | 557.231 | 0.58 |
| 3600 | 557.223 | 0.55 |
| 7200 | 557.236 | 0.60 |
| 14400 | 557.291 | 0.81 |

ONPF

| Time | centroid | D |
|---|---|---|
| 0 | 557.09 | 0.02 |
| 30 | 557.19 | 0.43 |
| 45 | 557.26 | 0.67 |
| 60 | 557.24 | 0.62 |
| 300 | 557.24 | 0.62 |
| 1500 | 557.25 | 0.65 |
| 3600 | 557.31 | 0.88 |
| 7200 | 557.33 | 0.97 |
| 14400 | 557.51 | 1.64 |

APO

| Time | centroid | D |
|---|---|---|
| 0 | 557.0765 | -0.01 |
| 30 | 557.182 | 0.39 |
| 45 | 557.198 | 0.45 |
| 60 | 557.2055 | 0.48 |
| 300 | 557.197 | 0.45 |
| 1500 | 557.2845 | 0.79 |
| 3600 | 557.3105 | 0.89 |
| 7200 | 557.433 | 1.36 |
| 14400 | 557.6805 | 2.31 |

DNA

| Time | centroid | D |
|---|---|---|
| 0 | 557.0715 | -0.03 |
| 30 | 557.1965 | 0.45 |
| 45 | 557.203 | 0.47 |
| 60 | 557.2325 | 0.59 |
| 300 | 557.2215 | 0.54 |
| 1500 | 557.212 | 0.51 |
| 3600 | 557.2665 | 0.72 |
| 7200 | 557.2305 | 0.58 |
| 14400 | 557.251 | 0.66 |

ONPFDNA

| Time | centroid | D |
|---|---|---|
| 0 | 557.077 | -0.01 |
| 30 | 557.216 | 0.52 |
| 45 | 557.206 | 0.48 |
| 60 | 557.2435 | 0.63 |
| 300 | 557.2085 | 0.49 |
| 1500 | 557.2095 | 0.50 |
| 3600 | 557.2635 | 0.71 |
| 7200 | 557.246 | 0.64 |
| 14400 | 557.256 | 0.68 |

TMG

| Time | centroid | D |
|---|---|---|
| 0 | 557.10 | 0.09 |
| 30 | 557.19 | 0.44 |
| 45 | 557.29 | 0.81 |
| 60 | 557.23 | 0.59 |
| 300 | 557.24 | 0.60 |
| 1500 | 557.33 | 0.96 |
| 3600 | 557.26 | 0.68 |
| 7200 | 557.23 | 0.57 |
| 14400 | 557.37 | 1.11 |

| | |
|---|---|
| control | 557.08 |
| infinity | 559.161 |

$$D(t) = \frac{M_t - M_0}{M_\infty - M_0} * N$$

IPTG

| Time | Replicate 1 | Replicate 2 | Replicate 3 | Replicate 4 | Replicate5 | average | SD |
|---|---|---|---|---|---|---|---|
| 0 | 557.08 | 557.099 | | | | 557.09 | 0.01 |
| 30 | 557.281 | 557.308 | | | | 557.29 | 0.02 |
| 45 | 557.25 | 557.191 | | | | 557.22 | 0.04 |
| 60 | 557.331 | 557.249 | | | | 557.29 | 0.06 |
| 300 | 557.364 | 557.195 | | | | 557.28 | 0.12 |
| 1500 | 557.248 | 557.214 | | | | 557.23 | 0.02 |
| 3600 | 557.267 | 557.179 | | | | 557.22 | 0.06 |
| 7200 | 557.259 | 557.213 | | | | 557.24 | 0.03 |
| 14400 | 557.337 | 557.245 | | | | 557.29 | 0.07 |

ONPF

| Time | Replicate 1 | Replicate 2 | Replicate 3 | Replicate 4 | average | SD |
|---|---|---|---|---|---|---|
| 0 | 557.074 | 557.099 | | | 557.09 | 0.02 |
| 30 | 557.218 | 557.168 | | | 557.19 | 0.04 |
| 45 | 557.307 | 557.204 | | | 557.26 | 0.07 |
| 60 | 557.242 | 557.239 | | | 557.24 | 0.00 |
| 300 | 557.279 | 557.202 | | | 557.24 | 0.05 |
| 1500 | 557.289 | 557.21 | | | 557.25 | 0.06 |
| 3600 | 557.393 | 557.226 | | | 557.31 | 0.12 |
| 7200 | 557.391 | 557.272 | | | 557.33 | 0.08 |
| 14400 | 557.507 | | | | 557.51 | |

APO

| Time | Replicate 1 | Replicate 2 | Replicate 3 | Replicate 4 | Replicate 5 | average | SD |
|---|---|---|---|---|---|---|---|
| 0 | 557.071 | 557.082 | | | | 557.08 | 0.01 |
| 30 | 557.213 | 557.151 | | | | 557.18 | 0.04 |
| 45 | 557.218 | 557.178 | | | | 557.20 | 0.03 |
| 60 | 557.208 | 557.203 | | | | 557.21 | 0.00 |
| 300 | 557.208 | 557.186 | | | | 557.20 | 0.02 |
| 1500 | 557.305 | 557.264 | | | | 557.28 | 0.03 |
| 3600 | 557.341 | 557.28 | | | | 557.31 | 0.04 |
| 7200 | 557.5 | 557.366 | | | | 557.43 | 0.09 |
| 14400 | 557.858 | 557.503 | | | | 557.68 | 0.25 |

DNA

| Time | Replicate 1 | Replicate 2 | Replicate 3 | Replicate 4 | Replicate 5 | average | SD |
|---|---|---|---|---|---|---|---|
| 0 | 557.055 | 557.088 | | | | 557.07 | 0.02 |
| 30 | 557.22 | 557.173 | | | | 557.20 | 0.03 |
| 45 | 557.22 | 557.186 | | | | 557.20 | 0.02 |
| 60 | 557.277 | 557.188 | | | | 557.23 | 0.06 |
| 300 | 557.236 | 557.207 | | | | 557.22 | 0.02 |
| 1500 | 557.227 | 557.197 | | | | 557.21 | 0.02 |
| 3600 | 557.346 | 557.187 | | | | 557.27 | 0.11 |
| 7200 | 557.266 | 557.195 | | | | 557.23 | 0.05 |
| 14400 | 557.337 | 557.165 | | | | 557.25 | 0.12 |

ONPFDNA

| Time | Replicate 1 | Replicate 2 | Replicate 3 | Replicate 4 | average | SD |
|---|---|---|---|---|---|---|
| 0 | 557.07 | 557.084 | | | 557.08 | 0.01 |
| 30 | 557.248 | 557.184 | | | 557.22 | 0.05 |
| 45 | 557.219 | 557.193 | | | 557.21 | 0.02 |
| 60 | 557.224 | 557.263 | | | 557.24 | 0.03 |
| 300 | 557.217 | 557.2 | | | 557.21 | 0.01 |
| 1500 | 557.179 | 557.24 | | | 557.21 | 0.04 |
| 3600 | 557.355 | 557.172 | | | 557.26 | 0.13 |
| 7200 | 557.278 | 557.214 | | | 557.25 | 0.05 |
| 14400 | 557.246 | 557.266 | | | 557.26 | 0.01 |

TMG

| Time | Replicate 1 | Replicate 2 | Replicate 3 | Replicate 4 | Replicate 5 | average | SD |
|---|---|---|---|---|---|---|---|
| 0 | 557.133 | 557.072 | | | | 557.10 | 0.04 |
| 30 | 557.135 | 557.254 | | | | 557.19 | 0.08 |
| 45 | 557.158 | 557.421 | | | | 557.29 | 0.19 |
| 60 | 557.233 | | | | | 557.23 | |
| 300 | 557.163 | 557.311 | | | | 557.24 | 0.10 |
| 1500 | 557.17 | 557.489 | | | | 557.33 | 0.23 |
| 3600 | 557.247 | 557.267 | | | | 557.26 | 0.01 |
| 7200 | 557.179 | 557.28 | | | | 557.23 | 0.07 |
| 14400 | 557.215 | 557.52 | | | | 557.37 | 0.22 |

### 174-177 LVAL

**Charge** 1

**IPTG**

| Time | centroid | D |
|---|---|---|
| 0 | 415.52 | 0.00 |
| 30 | 415.56 | 0.05 |
| 45 | 415.56 | 0.05 |
| 60 | 415.57 | 0.07 |
| 300 | 415.57 | 0.07 |
| 1500 | 415.58 | 0.07 |
| 3600 | 415.60 | 0.10 |
| 7200 | 415.64 | 0.16 |
| 14400 | 415.70 | 0.23 |

**ONPF**

| Time | centroid | D |
|---|---|---|
| 0 | 415.52 | 0.00 |
| 30 | 415.55 | 0.04 |
| 45 | 415.58 | 0.08 |
| 60 | 415.57 | 0.06 |
| 300 | 415.57 | 0.06 |
| 1500 | 415.57 | 0.07 |
| 3600 | 415.59 | 0.09 |
| 7200 | 415.61 | 0.12 |
| 14400 | 415.69 | 0.22 |

**APO**

| Time | centroid | D |
|---|---|---|
| 0 | 415.52 | 0.00 |
| 30 | 415.55 | 0.04 |
| 45 | 415.55 | 0.04 |
| 60 | 415.57 | 0.06 |
| 300 | 415.56 | 0.05 |
| 1500 | 415.59 | 0.09 |
| 3600 | 415.60 | 0.10 |
| 7200 | 415.64 | 0.16 |
| 14400 | 415.82 | 0.38 |

**DNA**

| Time | centroid | D |
|---|---|---|
| 0 | 415.52 | 0.00 |
| 30 | 415.55 | 0.04 |
| 45 | 415.58 | 0.07 |
| 60 | 415.56 | 0.05 |
| 300 | 415.56 | 0.05 |
| 1500 | 415.58 | 0.07 |
| 3600 | 415.60 | 0.10 |
| 7200 | 415.61 | 0.11 |
| 14400 | 415.63 | 0.14 |

**ONPFDNA**

| Time | centroid | D |
|---|---|---|
| 0 | 415.52 | 0.00 |
| 30 | 415.55 | 0.03 |
| 45 | 415.56 | 0.05 |
| 60 | 415.56 | 0.06 |
| 300 | 415.56 | 0.05 |
| 1500 | 415.57 | 0.06 |
| 3600 | 415.59 | 0.09 |
| 7200 | 415.60 | 0.11 |
| 14400 | 415.61 | 0.11 |

**TMG**

| Time | centroid | D |
|---|---|---|
| 0 | 415.52 | 0.00 |
| 30 | 415.56 | 0.05 |
| 45 | 415.59 | 0.09 |
| 60 | 415.66 | 0.18 |
| 300 | 415.58 | 0.07 |
| 1500 | 415.61 | 0.11 |
| 3600 | 415.61 | 0.12 |
| 7200 | 415.63 | 0.14 |
| 14400 | 415.71 | 0.24 |

control 415.52
infinity 417.087

$$D(t) = \frac{Mt - M0}{M\infty - M0} *N$$

**IPTG**

| Time | Replicate 1 | Replicate 2 | Replicate 3 | Replicate 4 | Replicate5 | average | SD |
|---|---|---|---|---|---|---|---|
| 0 | 415.52 | 415.52 | | | | 415.52 | 0.00 |
| 30 | 415.564 | 415.548 | | | | 415.56 | 0.01 |
| 45 | 415.57 | 415.548 | | | | 415.56 | 0.02 |
| 60 | 415.588 | 415.556 | | | | 415.57 | 0.02 |
| 300 | 415.595 | 415.55 | | | | 415.57 | 0.03 |
| 1500 | 415.583 | 415.57 | | | | 415.58 | 0.01 |
| 3600 | 415.615 | 415.588 | | | | 415.60 | 0.02 |
| 7200 | 415.654 | 415.635 | | | | 415.64 | 0.01 |
| 14400 | 415.738 | 415.662 | | | | 415.70 | 0.05 |

**ONPF**

| Time | Replicate 1 | Replicate 2 | Replicate 3 | Replicate 4 | average | SD |
|---|---|---|---|---|---|---|
| 0 | 415.516 | 415.527 | | | 415.52 | 0.01 |
| 30 | 415.55 | 415.55 | | | 415.55 | 0.00 |
| 45 | 415.598 | 415.565 | | | 415.58 | 0.02 |
| 60 | 415.567 | 415.573 | | | 415.57 | 0.00 |
| 300 | 415.569 | 415.562 | | | 415.57 | 0.00 |
| 1500 | 415.575 | 415.573 | | | 415.57 | 0.00 |
| 3600 | 415.596 | 415.584 | | | 415.59 | 0.01 |
| 7200 | 415.62 | 415.607 | | | 415.61 | 0.01 |
| 14400 | 415.689 | 415.689 | | | 415.69 | 0.00 |

**APO**

| Time | Replicate 1 | Replicate 2 | Replicate 3 | Replicate 4 | Replicate 5 | average | SD |
|---|---|---|---|---|---|---|---|
| 0 | 415.519 | 415.524 | | | | 415.52 | 0.00 |
| 30 | 415.548 | 415.55 | | | | 415.55 | 0.00 |
| 45 | 415.552 | 415.556 | | | | 415.55 | 0.00 |
| 60 | 415.55 | 415.58 | | | | 415.57 | 0.02 |
| 300 | 415.551 | 415.561 | | | | 415.56 | 0.01 |
| 1500 | 415.564 | 415.62 | | | | 415.59 | 0.04 |
| 3600 | 415.593 | 415.611 | | | | 415.60 | 0.01 |
| 7200 | 415.642 | 415.644 | | | | 415.64 | 0.00 |
| 14400 | 415.862 | 415.771 | | | | 415.82 | 0.06 |

**DNA**

| Time | Replicate 1 | Replicate 2 | Replicate 3 | Replicate 4 | Replicate 5 | average | SD |
|---|---|---|---|---|---|---|---|
| 0 | 415.519 | 415.52 | | | | 415.52 | 0.00 |
| 30 | 415.561 | 415.541 | | | | 415.55 | 0.01 |
| 45 | 415.607 | 415.549 | | | | 415.58 | 0.04 |
| 60 | 415.56 | 415.564 | | | | 415.56 | 0.00 |
| 300 | 415.57 | 415.554 | | | | 415.56 | 0.01 |
| 1500 | 415.577 | 415.578 | | | | 415.58 | 0.00 |
| 3600 | 415.619 | 415.577 | | | | 415.60 | 0.03 |
| 7200 | 415.62 | 415.594 | | | | 415.61 | 0.02 |
| 14400 | 415.7 | 415.558 | | | | 415.63 | 0.10 |

**ONPFDNA**

| Time | Replicate 1 | Replicate 2 | Replicate 3 | Replicate 4 | average | SD |
|---|---|---|---|---|---|---|
| 0 | 415.516 | 415.519 | | | 415.52 | 0.00 |
| 30 | 415.551 | 415.543 | | | 415.55 | 0.01 |
| 45 | 415.563 | 415.552 | | | 415.56 | 0.01 |
| 60 | 415.566 | 415.561 | | | 415.56 | 0.00 |
| 300 | 415.558 | 415.554 | | | 415.56 | 0.00 |
| 1500 | 415.556 | 415.583 | | | 415.57 | 0.02 |
| 3600 | 415.603 | 415.577 | | | 415.59 | 0.02 |
| 7200 | 415.607 | 415.598 | | | 415.60 | 0.01 |
| 14400 | 415.592 | 415.621 | | | 415.61 | 0.02 |

**TMG**

| Time | Replicate 1 | Replicate 2 | Replicate 3 | Replicate 4 | Replicate 5 | average | SD |
|---|---|---|---|---|---|---|---|
| 0 | 415.522 | 415.518 | | | | 415.52 | 0.00 |
| 30 | 415.558 | 415.563 | | | | 415.56 | 0.00 |
| 45 | 415.608 | 415.57 | | | | 415.59 | 0.03 |
| 60 | 415.57 | 415.748 | | | | 415.66 | 0.13 |
| 300 | 415.577 | 415.578 | | | | 415.58 | 0.00 |
| 1500 | 415.577 | 415.639 | | | | 415.61 | 0.04 |
| 3600 | 415.629 | 415.596 | | | | 415.61 | 0.02 |
| 7200 | 415.625 | 415.632 | | | | 415.63 | 0.00 |
| 14400 | 415.703 | 415.711 | | | | 415.71 | 0.01 |

### 174-184 LVALGHQQIAL

**Charge** 2

**IPTG**

| Time | centroid | D |
|---|---|---|
| 0 | 582.21 | 0.02 |
| 30 | 582.67 | 1.41 |
| 45 | 582.77 | 1.71 |
| 60 | 582.80 | 1.79 |
| 300 | 582.74 | 1.63 |
| 1500 | 582.79 | 1.78 |
| 3600 | 582.86 | 1.99 |
| 7200 | 582.95 | 2.26 |
| 14400 | 583.00 | 2.41 |

**ONPF**

| Time | centroid | D |
|---|---|---|
| 0 | 582.24 | 0.12 |
| 30 | 582.49 | 0.87 |
| 45 | 582.69 | 1.48 |
| 60 | 582.70 | 1.50 |
| 300 | 582.68 | 1.44 |
| 1500 | 582.72 | 1.55 |
| 3600 | 582.75 | 1.64 |
| 7200 | 582.75 | 1.66 |
| 14400 | 582.80 | 1.80 |

**APO**

| Time | centroid | D |
|---|---|---|
| 0 | 582.2 | 0.00 |
| 30 | 582.632 | 1.30 |
| 45 | 582.6825 | 1.45 |
| 60 | 582.7415 | 1.63 |
| 300 | 582.653 | 1.36 |
| 1500 | 582.7725 | 1.72 |
| 3600 | 582.8405 | 1.92 |
| 7200 | 582.8485 | 1.95 |
| 14400 | 583.1055 | 2.72 |

**DNA**

| Time | centroid | D |
|---|---|---|
| 0 | 582.21 | 0.03 |
| 30 | 582.61 | 1.22 |
| 45 | 582.73 | 1.58 |
| 60 | 582.73 | 1.59 |
| 300 | 582.74 | 1.62 |
| 1500 | 582.78 | 1.74 |
| 3600 | 582.82 | 1.85 |
| 7200 | 582.78 | 1.74 |
| 14400 | 582.83 | 1.88 |

**ONPFDNA**

| Time | centroid | D |
|---|---|---|
| 0 | 582.20 | 0.00 |
| 30 | 582.57 | 1.12 |
| 45 | 582.71 | 1.53 |
| 60 | 582.72 | 1.56 |
| 300 | 582.72 | 1.56 |
| 1500 | 582.68 | 1.44 |
| 3600 | 582.83 | 1.90 |
| 7200 | 582.78 | 1.74 |
| 14400 | 582.69 | 1.48 |

**TMG**

| Time | centroid | D |
|---|---|---|
| 0 | 582.21 | 0.03 |
| 30 | 582.69 | 1.46 |
| 45 | 582.78 | 1.75 |
| 60 | 582.90 | 2.09 |
| 300 | 582.82 | 1.86 |
| 1500 | 582.84 | 1.91 |
| 3600 | 582.91 | 2.13 |
| 7200 | 582.94 | 2.23 |
| 14400 | 583.07 | 2.61 |

control 582.2
infinity 585.199

$$D(t) = \frac{M_t - M_0}{M_\infty - M_0} * N$$

**IPTG**

| Time | Replicate 1 | Replicate 2 | Replicate 3 | Replicate 4 | Replicate5 | average | SD |
|---|---|---|---|---|---|---|---|
| 0 | 582.21 | 582.2 | | | | 582.21 | 0.01 |
| 30 | 582.818 | 582.525 | | | | 582.67 | 0.21 |
| 45 | 582.855 | 582.682 | | | | 582.77 | 0.12 |
| 60 | 582.892 | 582.703 | | | | 582.80 | 0.13 |
| 300 | 582.868 | 582.619 | | | | 582.74 | 0.18 |
| 1500 | 582.938 | 582.65 | | | | 582.79 | 0.20 |
| 3600 | 582.976 | 582.752 | | | | 582.86 | 0.16 |
| 7200 | 583.093 | 582.816 | | | | 582.95 | 0.20 |
| 14400 | 583.16 | 582.848 | | | | 583.00 | 0.22 |

**ONPF**

| Time | Replicate 1 | Replicate 2 | Replicate 3 | Replicate 4 | average | SD |
|---|---|---|---|---|---|---|
| 0 | 582.235 | 582.245 | | | 582.24 | 0.01 |
| 30 | 582.544 | 582.439 | | | 582.49 | 0.07 |
| 45 | 582.817 | 582.568 | | | 582.69 | 0.18 |
| 60 | 582.727 | 582.671 | | | 582.70 | 0.04 |
| 300 | 582.761 | 582.6 | | | 582.68 | 0.11 |
| 1500 | 582.783 | 582.653 | | | 582.72 | 0.09 |
| 3600 | 582.832 | 582.659 | | | 582.75 | 0.12 |
| 7200 | 582.854 | 582.652 | | | 582.75 | 0.14 |
| 14400 | 583.036 | 582.562 | | | 582.80 | 0.34 |

**APO**

| Time | Replicate 1 | Replicate 2 | Replicate 3 | Replicate 4 | Replicate 5 | average | SD |
|---|---|---|---|---|---|---|---|
| 0 | 582.193 | 582.207 | | | | 582.20 | 0.01 |
| 30 | 582.745 | 582.519 | | | | 582.63 | 0.16 |
| 45 | 582.795 | 582.57 | | | | 582.68 | 0.16 |
| 60 | 582.806 | 582.677 | | | | 582.74 | 0.09 |
| 300 | 582.752 | 582.554 | | | | 582.65 | 0.14 |
| 1500 | 582.818 | 582.727 | | | | 582.77 | 0.06 |
| 3600 | 582.926 | 582.755 | | | | 582.84 | 0.12 |
| 7200 | 583 | 582.697 | | | | 582.85 | 0.21 |
| 14400 | 583.298 | 582.913 | | | | 583.11 | 0.27 |

**DNA**

| Time | Replicate 1 | Replicate 2 | Replicate 3 | Replicate 4 | Replicate 5 | average | SD |
|---|---|---|---|---|---|---|---|
| 0 | 582.195 | 582.223 | | | | 582.21 | 0.02 |
| 30 | 582.647 | 582.563 | | | | 582.61 | 0.06 |
| 45 | 582.805 | 582.65 | | | | 582.73 | 0.11 |
| 60 | 582.764 | 582.695 | | | | 582.73 | 0.05 |
| 300 | 582.786 | 582.695 | | | | 582.74 | 0.06 |
| 1500 | 582.805 | 582.753 | | | | 582.78 | 0.04 |
| 3600 | 582.926 | 582.707 | | | | 582.82 | 0.15 |
| 7200 | 582.923 | 582.636 | | | | 582.78 | 0.20 |
| 14400 | 583.054 | 582.596 | | | | 582.83 | 0.32 |

**ONPFDNA**

| Time | Replicate 1 | Replicate 2 | Replicate 3 | Replicate 4 | average | SD |
|---|---|---|---|---|---|---|
| 0 | 582.202 | 582.199 | | | 582.20 | 0.00 |
| 30 | 582.613 | 582.535 | | | 582.57 | 0.06 |
| 45 | 582.764 | 582.655 | | | 582.71 | 0.08 |
| 60 | 582.745 | 582.694 | | | 582.72 | 0.04 |
| 300 | 582.771 | 582.667 | | | 582.72 | 0.07 |
| 1500 | 582.68 | 582.681 | | | 582.68 | 0.00 |
| 3600 | 582.9 | 582.769 | | | 582.83 | 0.09 |
| 7200 | 582.905 | 582.652 | | | 582.78 | 0.18 |
| 14400 | 582.692 | 582.697 | | | 582.69 | 0.00 |

**TMG**

| Time | Replicate 1 | Replicate 2 | Replicate 3 | Replicate 4 | Replicate 5 | average | SD |
|---|---|---|---|---|---|---|---|
| 0 | 582.225 | 582.195 | | | | 582.21 | 0.02 |
| 30 | 582.566 | 582.808 | | | | 582.69 | 0.17 |
| 45 | 582.768 | 582.8 | | | | 582.78 | 0.02 |
| 60 | 582.761 | 583.034 | | | | 582.90 | 0.19 |
| 300 | 582.8 | 582.837 | | | | 582.82 | 0.03 |
| 1500 | 582.729 | 582.946 | | | | 582.84 | 0.15 |
| 3600 | 582.874 | 582.946 | | | | 582.91 | 0.05 |
| 7200 | 582.85 | 583.034 | | | | 582.94 | 0.13 |
| 14400 | 583.006 | 583.133 | | | | 583.07 | 0.09 |

### 177-185 LGHQQIALL

**Charge** 2

**IPTG**

| Time | centroid | D |
|---|---|---|
| 0 | 497.09 | 0.05 |
| 30 | 497.35 | 0.95 |
| 45 | 497.46 | 1.34 |
| 60 | 497.42 | 1.21 |
| 300 | 497.41 | 1.19 |
| 1500 | 497.41 | 1.16 |
| 3600 | 497.42 | 1.22 |
| 7200 | 497.44 | 1.26 |
| 14400 | 497.46 | 1.33 |

**ONPF**

| Time | centroid | D |
|---|---|---|
| 0 | 497.12 | 0.17 |
| 30 | 497.30 | 0.79 |
| 45 | 497.34 | 0.94 |
| 60 | 497.38 | 1.06 |
| 300 | 497.38 | 1.07 |
| 1500 | 497.40 | 1.12 |
| 3600 | 497.41 | 1.19 |
| 7200 | 497.42 | 1.20 |
| 14400 | 497.45 | 1.30 |

**APO**

| Time | centroid | D |
|---|---|---|
| 0 | 497.10 | 0.09 |
| 30 | 497.32 | 0.86 |
| 45 | 497.33 | 0.90 |
| 60 | 497.38 | 1.06 |
| 300 | 497.39 | 1.10 |
| 1500 | 497.40 | 1.15 |
| 3600 | 497.43 | 1.23 |
| 7200 | 497.41 | 1.16 |
| 14400 | 497.44 | 1.26 |

**DNA**

| Time | centroid | D |
|---|---|---|
| 0 | 497.08 | 0.04 |
| 30 | 497.33 | 0.89 |
| 45 | 497.43 | 1.24 |
| 60 | 497.42 | 1.21 |
| 300 | 497.44 | 1.28 |
| 1500 | 497.44 | 1.29 |
| 3600 | 497.44 | 1.29 |
| 7200 | 497.45 | 1.32 |
| 14400 | 497.44 | 1.29 |

**ONPFDNA**

| Time | centroid | D |
|---|---|---|
| 0 | 497.08 | 0.02 |
| 30 | 497.30 | 0.78 |
| 45 | 497.42 | 1.21 |
| 60 | 497.45 | 1.31 |
| 300 | 497.42 | 1.22 |
| 1500 | 497.39 | 1.12 |
| 3600 | 497.44 | 1.28 |
| 7200 | 497.44 | 1.29 |
| 14400 | 497.42 | 1.19 |

**TMG**

| Time | centroid | D |
|---|---|---|
| 0 | 497.09 | 0.06 |
| 30 | 497.38 | 1.07 |
| 45 | 497.49 | 1.47 |
| 60 | 497.54 | 1.63 |
| 300 | 497.53 | 1.60 |
| 1500 | 497.50 | 1.48 |
| 3600 | 497.49 | 1.44 |
| 7200 | 497.51 | 1.53 |
| 14400 | 497.55 | 1.66 |

control 497.072
infinity 499.088

$$D(t) = \frac{Mt - M0}{M\infty - M0} *N$$

**IPTG**

| Time | Replicate 1 | Replicate 2 | Replicate 3 | Replicate 4 | Replicate5 | average | SD |
|---|---|---|---|---|---|---|---|
| 0 | 497.083 | 497.095 | 497.092 | 497.087 | 497.071 | 497.09 | 0.01 |
| 30 | 497.271 | 497.358 | 497.313 | 497.289 | 497.503 | 497.35 | 0.09 |
| 45 | 497.358 | | | | 497.559 | 497.46 | 0.14 |
| 60 | 497.385 | | 497.378 | 497.37 | 497.547 | 497.42 | 0.08 |
| 300 | 497.296 | 497.485 | 497.384 | 497.348 | 497.56 | 497.41 | 0.11 |
| 1500 | 497.315 | 497.468 | 497.407 | 497.354 | 497.493 | 497.41 | 0.07 |
| 3600 | 497.364 | 497.493 | 497.382 | 497.37 | 497.502 | 497.42 | 0.07 |
| 7200 | 497.318 | 497.537 | 497.411 | 497.378 | 497.534 | 497.44 | 0.10 |
| 14400 | 497.398 | 497.528 | 497.428 | 497.377 | 497.544 | 497.46 | 0.08 |

**ONPF**

| Time | Replicate 1 | Replicate 2 | Replicate 3 | Replicate 4 | average | SD |
|---|---|---|---|---|---|---|
| 0 | 497.111 | 497.189 | 497.086 | 497.093 | 497.12 | 0.05 |
| 30 | 497.26 | 497.338 | 497.257 | 497.34 | 497.30 | 0.05 |
| 45 | 497.343 | | | | 497.34 | 0.00 |
| 60 | 497.386 | | 497.378 | 497.367 | 497.38 | 0.01 |
| 300 | 497.339 | 497.429 | 497.334 | 497.42 | 497.38 | 0.05 |
| 1500 | 497.356 | 497.502 | 497.373 | 497.349 | 497.40 | 0.07 |
| 3600 | 497.336 | 497.53 | 497.359 | 497.43 | 497.41 | 0.09 |
| 7200 | | 497.506 | 497.355 | 497.396 | 497.42 | 0.08 |
| 14400 | | 497.512 | 497.41 | 497.42 | 497.45 | 0.06 |

**APO**

| Time | Replicate 1 | Replicate 2 | Replicate 3 | Replicate 4 | Replicate 5 | average | SD |
|---|---|---|---|---|---|---|---|
| 0 | 497.072 | 497.139 | 497.101 | 497.084 | | 497.10 | 0.03 |
| 30 | 497.311 | | 497.357 | 497.288 | | 497.32 | 0.04 |
| 45 | 497.331 | | | | | 497.33 | |
| 60 | 497.386 | | 497.373 | 497.374 | | 497.38 | 0.01 |
| 300 | 497.317 | 497.482 | 497.394 | 497.358 | | 497.39 | 0.07 |
| 1500 | 497.386 | 497.477 | 497.383 | 497.365 | | 497.40 | 0.05 |
| 3600 | 497.401 | 497.526 | 497.405 | 497.368 | | 497.43 | 0.07 |
| 7200 | 497.316 | 497.52 | 497.428 | 497.364 | | 497.41 | 0.09 |
| 14400 | 497.378 | 497.511 | 497.439 | 497.413 | | 497.44 | 0.06 |

**DNA**

| Time | Replicate 1 | Replicate 2 | Replicate 3 | Replicate 4 | Replicate 5 | average | SD |
|---|---|---|---|---|---|---|---|
| 0 | 497.083 | 497.088 | 497.09 | 497.078 | | 497.08 | 0.01 |
| 30 | 497.326 | 497.343 | 497.271 | 497.379 | | 497.33 | 0.04 |
| 45 | 497.38 | | | 497.48 | | 497.43 | 0.07 |
| 60 | 497.393 | | 497.386 | 497.481 | | 497.42 | 0.05 |
| 300 | 497.399 | 497.519 | 497.351 | 497.49 | | 497.44 | 0.08 |
| 1500 | 497.42 | 497.506 | 497.363 | 497.488 | | 497.44 | 0.07 |
| 3600 | 497.384 | 497.511 | 497.382 | 497.495 | | 497.44 | 0.07 |
| 7200 | | 497.535 | 497.355 | 497.469 | | 497.45 | 0.09 |
| 14400 | 497.347 | 497.512 | 497.407 | 497.507 | | 497.44 | 0.08 |

**ONPFDNA**

| Time | Replicate 1 | Replicate 2 | Replicate 3 | Replicate 4 | average | SD |
|---|---|---|---|---|---|---|
| 0 | 497.063 | 497.086 | 497.082 | | 497.08 | 0.01 |
| 30 | 497.281 | 497.234 | 497.373 | | 497.30 | 0.07 |
| 45 | 497.339 | | 497.5 | | 497.42 | 0.11 |
| 60 | 497.381 | | 497.516 | | 497.45 | 0.10 |
| 300 | 497.375 | 497.401 | 497.494 | | 497.42 | 0.06 |
| 1500 | 497.368 | 497.409 | 497.405 | | 497.39 | 0.02 |
| 3600 | 497.41 | 497.421 | 497.49 | | 497.44 | 0.04 |
| 7200 | | 497.404 | 497.484 | | 497.44 | 0.06 |
| 14400 | | 497.406 | 497.425 | | 497.42 | 0.01 |

**TMG**

| Time | Replicate 1 | Replicate 2 | Replicate 3 | Replicate 4 | Replicate 5 | average | SD |
|---|---|---|---|---|---|---|---|
| 0 | 497.109 | 497.07 | | | | 497.09 | 0.03 |
| 30 | 497.341 | 497.419 | | | | 497.38 | 0.06 |
| 45 | 497.483 | 497.506 | | | | 497.49 | 0.02 |
| 60 | 497.519 | 497.566 | | | | 497.54 | 0.03 |
| 300 | 497.516 | 497.547 | | | | 497.53 | 0.02 |
| 1500 | 497.445 | 497.549 | | | | 497.50 | 0.07 |
| 3600 | 497.46 | 497.513 | | | | 497.49 | 0.04 |
| 7200 | 497.473 | 497.552 | | | | 497.51 | 0.06 |
| 14400 | 497.523 | 497.578 | | | | 497.55 | 0.04 |

### 182-185 IALL

**Charge** 1

**IPTG**

| Time | centroid | D |
|---|---|---|
| 0 | 429.512 | -0.02 |
| 30 | 429.5425 | 0.01 |
| 45 | 429.5445 | 0.02 |
| 60 | 429.5615 | 0.04 |
| 300 | 429.555 | 0.03 |
| 1500 | 429.554 | 0.03 |
| 3600 | 429.546 | 0.02 |
| 7200 | 429.5515 | 0.02 |
| 14400 | 429.5425 | 0.01 |

**ONPF**

| Time | centroid | D |
|---|---|---|
| 0 | 429.502 | -0.03 |
| 30 | 429.543 | 0.01 |
| 45 | 429.5385 | 0.01 |
| 60 | 429.5355 | 0.01 |
| 300 | 429.5375 | 0.01 |
| 1500 | 429.5305 | 0.00 |
| 3600 | 429.524 | -0.01 |
| 7200 | 429.525 | -0.01 |
| 14400 | 429.56 | 0.03 |

**APO**

| Time | centroid | D |
|---|---|---|
| 0 | 429.53 | -0.01 |
| 30 | 429.55 | 0.02 |
| 45 | 429.55 | 0.02 |
| 60 | 429.56 | 0.04 |
| 300 | 429.56 | 0.04 |
| 1500 | 429.54 | 0.01 |
| 3600 | 429.55 | 0.02 |
| 7200 | 429.53 | 0.01 |
| 14400 | 429.60 | 0.08 |

**DNA**

| Time | centroid | D |
|---|---|---|
| 0 | 429.511 | -0.02 |
| 30 | 429.535 | 0.01 |
| 45 | 429.576 | 0.05 |
| 60 | 429.5405 | 0.01 |
| 300 | 429.547 | 0.02 |
| 1500 | 429.535 | 0.01 |
| 3600 | 429.544 | 0.02 |
| 7200 | 429.5435 | 0.02 |
| 14400 | 429.579 | 0.06 |

**ONPFDNA**

| Time | centroid | D |
|---|---|---|
| 0 | 429.484 | -0.05 |
| 30 | 429.526 | 0.00 |
| 45 | 429.523 | -0.01 |
| 60 | 429.541 | 0.01 |
| 300 | 429.504 | -0.03 |
| 1500 | 429.52 | -0.01 |
| 3600 | 429.5295 | 0.00 |
| 7200 | 429.5195 | -0.01 |
| 14400 | 429.514 | -0.02 |

**TMG**

| Time | centroid | D |
|---|---|---|
| 0 | 429.50 | -0.03 |
| 30 | 429.54 | 0.02 |
| 45 | 429.54 | 0.01 |
| 60 | 429.59 | 0.07 |
| 300 | 429.55 | 0.03 |
| 1500 | 429.56 | 0.04 |
| 3600 | 429.55 | 0.02 |
| 7200 | 429.54 | 0.01 |
| 14400 | 429.55 | 0.02 |

| | |
|---|---|
| control | 429.53 |
| infinity | 431.3 |

$$D(t) = \frac{M_t - M_0}{M_\infty - M_0} *N$$

**IPTG**

| Time | Replicate 1 | Replicate 2 | Replicate 3 | Replicate 4 | Replicate5 | average | SD |
|---|---|---|---|---|---|---|---|
| 0 | 429.507 | 429.517 | | | | 429.51 | 0.01 |
| 30 | 429.546 | 429.539 | | | | 429.54 | 0.00 |
| 45 | 429.543 | 429.546 | | | | 429.54 | 0.00 |
| 60 | 429.567 | 429.556 | | | | 429.56 | 0.01 |
| 300 | 429.571 | 429.539 | | | | 429.56 | 0.02 |
| 1500 | 429.549 | 429.559 | | | | 429.55 | 0.01 |
| 3600 | 429.532 | 429.56 | | | | 429.55 | 0.02 |
| 7200 | 429.552 | 429.551 | | | | 429.55 | 0.00 |
| 14400 | 429.529 | 429.556 | | | | 429.54 | 0.02 |

**ONPF**

| Time | Replicate 1 | Replicate 2 | Replicate 3 | Replicate 4 | average | SD |
|---|---|---|---|---|---|---|
| 0 | 429.51 | 429.494 | | | 429.50 | 0.01 |
| 30 | 429.557 | 429.529 | | | 429.54 | 0.02 |
| 45 | 429.546 | 429.531 | | | 429.54 | 0.01 |
| 60 | 429.546 | 429.525 | | | 429.54 | 0.01 |
| 300 | 429.542 | 429.533 | | | 429.54 | 0.01 |
| 1500 | 429.542 | 429.519 | | | 429.53 | 0.02 |
| 3600 | 429.538 | 429.51 | | | 429.52 | 0.02 |
| 7200 | 429.542 | 429.508 | | | 429.53 | 0.02 |
| 14400 | 429.56 | | | | 429.56 | |

**APO**

| Time | Replicate 1 | Replicate 2 | Replicate 3 | Replicate 4 | Replicate 5 | average | SD |
|---|---|---|---|---|---|---|---|
| 0 | 429.512 | 429.539 | | | | 429.53 | 0.02 |
| 30 | 429.527 | 429.567 | | | | 429.55 | 0.03 |
| 45 | 429.526 | 429.569 | | | | 429.55 | 0.03 |
| 60 | 429.56 | 429.569 | | | | 429.56 | 0.01 |
| 300 | 429.542 | 429.583 | | | | 429.56 | 0.03 |
| 1500 | 429.524 | 429.561 | | | | 429.54 | 0.03 |
| 3600 | 429.543 | 429.556 | | | | 429.55 | 0.01 |
| 7200 | 429.516 | 429.553 | | | | 429.53 | 0.03 |
| 14400 | 429.626 | 429.575 | | | | 429.60 | 0.04 |

**DNA**

| Time | Replicate 1 | Replicate 2 | Replicate 3 | Replicate 4 | Replicate 5 | average | SD |
|---|---|---|---|---|---|---|---|
| 0 | 429.498 | 429.524 | | | | 429.51 | 0.02 |
| 30 | 429.544 | 429.526 | | | | 429.54 | 0.01 |
| 45 | | 429.576 | | | | 429.58 | |
| 60 | 429.522 | 429.559 | | | | 429.54 | 0.03 |
| 300 | 429.532 | 429.562 | | | | 429.55 | 0.02 |
| 1500 | 429.514 | 429.556 | | | | 429.54 | 0.03 |
| 3600 | 429.535 | 429.553 | | | | 429.54 | 0.01 |
| 7200 | 429.542 | 429.545 | | | | 429.54 | 0.00 |
| 14400 | 429.57 | 429.588 | | | | 429.58 | 0.01 |

**ONPFDNA**

| Time | Replicate 1 | Replicate 2 | Replicate 3 | Replicate 4 | average | SD |
|---|---|---|---|---|---|---|
| 0 | 429.493 | 429.475 | | | 429.48 | 0.01 |
| 30 | 429.53 | 429.522 | | | 429.53 | 0.01 |
| 45 | 429.529 | 429.517 | | | 429.52 | 0.01 |
| 60 | 429.56 | 429.522 | | | 429.54 | 0.03 |
| 300 | 429.498 | 429.51 | | | 429.50 | 0.01 |
| 1500 | 429.518 | 429.522 | | | 429.52 | 0.00 |
| 3600 | 429.532 | 429.527 | | | 429.53 | 0.00 |
| 7200 | 429.524 | 429.515 | | | 429.52 | 0.01 |
| 14400 | 429.55 | 429.478 | | | 429.51 | 0.05 |

**TMG**

| Time | Replicate 1 | Replicate 2 | Replicate 3 | Replicate 4 | Replicate 5 | average | SD |
|---|---|---|---|---|---|---|---|
| 0 | 429.495 | 429.514 | | | | 429.50 | 0.01 |
| 30 | 429.525 | 429.562 | | | | 429.54 | 0.03 |
| 45 | 429.532 | 429.547 | | | | 429.54 | 0.01 |
| 60 | 429.528 | 429.65 | | | | 429.59 | 0.09 |
| 300 | 429.551 | 429.558 | | | | 429.55 | 0.00 |
| 1500 | 429.533 | 429.592 | | | | 429.56 | 0.04 |
| 3600 | 429.543 | 429.556 | | | | 429.55 | 0.01 |
| 7200 | 429.525 | 429.552 | | | | 429.54 | 0.02 |
| 14400 | 429.543 | 429.555 | | | | 429.55 | 0.01 |

### 185-190 LAGPLS

**Charge** 1

**IPTG**

| Time | centroid | D |
|---|---|---|
| 0 | 557.62 | 0.00 |
| 30 | 558.26 | 0.93 |
| 45 | 558.35 | 1.07 |
| 60 | 558.41 | 1.15 |
| 300 | 558.41 | 1.16 |
| 1500 | 558.67 | 1.53 |
| 3600 | 558.90 | 1.88 |
| 7200 | 558.89 | 1.86 |
| 14400 | 558.78 | 1.70 |

**ONPF**

| Time | centroid | D |
|---|---|---|
| 0 | 557.60 | -0.03 |
| 30 | 558.03 | 0.60 |
| 45 | 558.21 | 0.87 |
| 60 | 558.35 | 1.07 |
| 300 | 558.39 | 1.13 |
| 1500 | 558.64 | 1.49 |
| 3600 | 558.70 | 1.57 |
| 7200 | 558.55 | 1.37 |
| 14400 | 558.84 | 1.79 |

**APO**

| Time | centroid | D |
|---|---|---|
| 0 | 557.62 | 0.00 |
| 30 | 558.23 | 0.89 |
| 45 | 558.25 | 0.92 |
| 60 | 558.39 | 1.13 |
| 300 | 558.42 | 1.17 |
| 1500 | 558.74 | 1.64 |
| 3600 | 558.82 | 1.75 |
| 7200 | 558.86 | 1.81 |
| 14400 | 558.83 | 1.76 |

**DNA**

| Time | centroid | D |
|---|---|---|
| 0 | 557.61 | -0.02 |
| 30 | 558.16 | 0.79 |
| 45 | 558.30 | 0.99 |
| 60 | 558.29 | 0.97 |
| 300 | 558.47 | 1.25 |
| 1500 | 558.57 | 1.39 |
| 3600 | 558.62 | 1.46 |
| 7200 | 558.50 | 1.29 |
| 14400 | 558.74 | 1.63 |

**ONPFDNA**

| Time | centroid | D |
|---|---|---|
| 0 | 557.60 | -0.03 |
| 30 | 558.15 | 0.78 |
| 45 | 558.29 | 0.99 |
| 60 | 558.29 | 0.98 |
| 300 | 558.44 | 1.19 |
| 1500 | 558.32 | 1.02 |
| 3600 | 558.66 | 1.52 |
| 7200 | | |
| 14400 | 558.24 | 0.90 |

**TMG**

| Time | centroid | D |
|---|---|---|
| 0 | 557.60 | -0.03 |
| 30 | 558.24 | 0.90 |
| 45 | 558.34 | 1.05 |
| 60 | 558.47 | 1.24 |
| 300 | 558.51 | 1.31 |
| 1500 | 558.65 | 1.51 |
| 3600 | 558.93 | 1.91 |
| 7200 | 558.88 | 1.84 |
| 14400 | 559.02 | 2.05 |

control 557.62
infinity 559.671

$$D(t) = \frac{Mt - M0}{M\infty - M0} *N$$

**IPTG**

| Time | Replicate 1 | Replicate 2 | Replicate 3 | Replicate 4 | Replicate5 | average | SD |
|---|---|---|---|---|---|---|---|
| 0 | 557.618 | 557.625 | | | | 557.62 | 0.00 |
| 30 | 558.389 | 558.129 | | | | 558.26 | 0.18 |
| 45 | 558.391 | 558.316 | | | | 558.35 | 0.05 |
| 60 | 558.476 | 558.343 | | | | 558.41 | 0.09 |
| 300 | 558.461 | 558.365 | | | | 558.41 | 0.07 |
| 1500 | 558.861 | 558.475 | | | | 558.67 | 0.27 |
| 3600 | 558.923 | 558.881 | | | | 558.90 | 0.03 |
| 7200 | 559.05 | 558.729 | | | | 558.89 | 0.23 |
| 14400 | 558.844 | 558.722 | | | | 558.78 | 0.09 |

**ONPF**

| Time | Replicate 1 | Replicate 2 | Replicate 3 | Replicate 4 | average | SD |
|---|---|---|---|---|---|---|
| 0 | 557.566 | 557.633 | | | 557.60 | 0.05 |
| 30 | 558.036 | 558.023 | | | 558.03 | 0.01 |
| 45 | 558.289 | 558.136 | | | 558.21 | 0.11 |
| 60 | 558.351 | 558.348 | | | 558.35 | 0.00 |
| 300 | 558.49 | 558.294 | | | 558.39 | 0.14 |
| 1500 | 558.841 | 558.436 | | | 558.64 | 0.29 |
| 3600 | 558.961 | 558.431 | | | 558.70 | 0.37 |
| 7200 | 558.714 | 558.393 | | | 558.55 | 0.23 |
| 14400 | 558.841 | | | | 558.84 | |

**APO**

| Time | Replicate 1 | Replicate 2 | Replicate 3 | Replicate 4 | Replicate 5 | average | SD |
|---|---|---|---|---|---|---|---|
| 0 | 557.614 | 557.623 | | | | 557.62 | 0.01 |
| 30 | 558.336 | 558.125 | | | | 558.23 | 0.15 |
| 45 | 558.347 | 558.156 | | | | 558.25 | 0.14 |
| 60 | 558.454 | 558.334 | | | | 558.39 | 0.08 |
| 300 | 558.563 | 558.275 | | | | 558.42 | 0.20 |
| 1500 | 558.828 | 558.656 | | | | 558.74 | 0.12 |
| 3600 | 558.906 | 558.733 | | | | 558.82 | 0.12 |
| 7200 | 558.859 | | | | | 558.86 | |
| 14400 | 558.946 | 558.705 | | | | 558.83 | 0.17 |

**DNA**

| Time | Replicate 1 | Replicate 2 | Replicate 3 | Replicate 4 | Replicate 5 | average | SD |
|---|---|---|---|---|---|---|---|
| 0 | 557.593 | 557.626 | | | | 557.61 | 0.02 |
| 30 | 558.13 | 558.192 | | | | 558.16 | 0.04 |
| 45 | 558.395 | 558.201 | | | | 558.30 | 0.14 |
| 60 | 558.299 | 558.271 | | | | 558.29 | 0.02 |
| 300 | 558.494 | 558.455 | | | | 558.47 | 0.03 |
| 1500 | | 558.569 | | | | 558.57 | |
| 3600 | 558.755 | 558.48 | | | | 558.62 | 0.19 |
| 7200 | 558.659 | 558.341 | | | | 558.50 | 0.22 |
| 14400 | 558.736 | | | | | 558.74 | |

**ONPFDNA**

| Time | Replicate 1 | Replicate 2 | Replicate 3 | Replicate 4 | average | SD |
|---|---|---|---|---|---|---|
| 0 | 557.587 | 557.607 | | | 557.60 | 0.01 |
| 30 | 558.093 | 558.211 | | | 558.15 | 0.08 |
| 45 | 558.284 | 558.304 | | | 558.29 | 0.01 |
| 60 | 558.27 | 558.313 | | | 558.29 | 0.03 |
| 300 | 558.433 | 558.44 | | | 558.44 | 0.00 |
| 1500 | 558.284 | 558.351 | | | 558.32 | 0.05 |
| 3600 | 558.637 | 558.684 | | | 558.66 | 0.03 |
| 7200 | | | | | | |
| 14400 | 558.161 | 558.309 | | | 558.24 | 0.10 |

**TMG**

| Time | Replicate 1 | Replicate 2 | Replicate 3 | Replicate 4 | Replicate 5 | average | SD |
|---|---|---|---|---|---|---|---|
| 0 | 557.633 | 557.563 | | | | 557.60 | 0.05 |
| 30 | 558.08 | 558.397 | | | | 558.24 | 0.22 |
| 45 | 558.317 | 558.353 | | | | 558.34 | 0.03 |
| 60 | 558.317 | 558.622 | | | | 558.47 | 0.22 |
| 300 | 558.481 | 558.546 | | | | 558.51 | 0.05 |
| 1500 | 558.555 | 558.753 | | | | 558.65 | 0.14 |
| 3600 | 558.909 | 558.943 | | | | 558.93 | 0.02 |
| 7200 | 558.716 | 559.04 | | | | 558.88 | 0.23 |
| 14400 | 558.916 | 559.123 | | | | 559.02 | 0.15 |

### 185-196 LAGPLSSVSARL

**Charge** 2

**IPTG**

| Time | centroid | D |
|---|---|---|
| 0 | 586.1762 | -0.14 |
| 30 | 586.4296 | 0.92 |
| 45 | 586.6877 | 1.99 |
| 60 | 586.8808 | 2.80 |
| 300 | 586.9417 | 3.06 |
| 1500 | 587.2398 | 4.30 |
| 3600 | 587.3935 | 4.94 |
| 7200 | 587.6755 | 6.12 |
| 14400 | 587.6406 | 5.97 |

**ONPF**

| Time | centroid | D |
|---|---|---|
| 0 | 586.20 | -0.03 |
| 30 | 586.65 | 1.83 |
| 45 | 586.95 | 3.09 |
| 60 | 587.18 | 4.05 |
| 300 | 587.28 | 4.46 |
| 1500 | 587.61 | 5.85 |
| 3600 | 587.59 | 5.78 |
| 7200 | 587.45 | 5.18 |
| 14400 | 587.57 | 5.67 |

**APO**

| Time | centroid | D |
|---|---|---|
| 0 | 586.208 | -0.01 |
| 30 | 586.7106 | 2.09 |
| 45 | 586.958 | 3.12 |
| 60 | 587.279 | 4.46 |
| 300 | 587.2793 | 4.47 |
| 1500 | 587.4508 | 5.18 |
| 3600 | 587.5124 | 5.44 |
| 7200 | 587.6995 | 6.22 |
| 14400 | 587.742 | 6.40 |

**DNA**

| Time | centroid | D |
|---|---|---|
| 0 | 586.32 | 0.44 |
| 30 | 586.95 | 3.11 |
| 45 | 587.16 | 3.98 |
| 60 | 587.35 | 4.76 |
| 300 | 587.50 | 5.39 |
| 1500 | 587.62 | 5.89 |
| 3600 | 587.67 | 6.12 |
| 7200 | 587.52 | 5.49 |
| 14400 | 587.56 | 5.65 |

**ONPFDNA**

| Time | centroid | D |
|---|---|---|
| 0 | 586.17 | -0.18 |
| 30 | 586.73 | 2.16 |
| 45 | 587.10 | 3.73 |
| 60 | 587.21 | 4.18 |
| 300 | 587.36 | 4.79 |
| 1500 | 587.46 | 5.24 |
| 3600 | 587.74 | 6.40 |
| 7200 | 587.48 | 5.32 |
| 14400 | 587.50 | 5.37 |

**TMG**

| Time | centroid | D |
|---|---|---|
| 0 | 586.19 | -0.07 |
| 30 | 586.60 | 1.62 |
| 45 | 586.95 | 3.08 |
| 60 | 587.20 | 4.13 |
| 300 | 587.07 | 3.57 |
| 1500 | 587.30 | 4.57 |
| 3600 | 587.79 | 6.61 |
| 7200 | 588.02 | 7.58 |
| 14400 | 588.02 | 7.57 |

control 586.21
infinity 588.365

$$D(t) = \frac{Mt - M0}{M\infty - M0} *N$$

**IPTG**

| Time | Replicate 1 | Replicate 2 | Replicate 3 | Replicate 4 | Replicate5 | average | SD |
|---|---|---|---|---|---|---|---|
| 0 | 586.148 | 586.171 | 586.187 | 586.187 | 586.188 | 586.18 | 0.02 |
| 30 | 586.328 | 586.396 | 586.346 | 586.403 | 586.675 | 586.43 | 0.14 |
| 45 | | 586.599 | 586.583 | | 586.881 | 586.69 | 0.17 |
| 60 | | 586.735 | 586.993 | 586.924 | 586.871 | 586.88 | 0.11 |
| 300 | | | 587.076 | 586.86 | 586.889 | 586.94 | 0.12 |
| 1500 | | 587.122 | 587.44 | 587.115 | 587.282 | 587.24 | 0.15 |
| 3600 | 587.255 | 587.513 | 587.519 | 587.287 | | 587.39 | 0.14 |
| 7200 | 587.551 | 587.569 | 587.674 | | 587.908 | 587.68 | 0.16 |
| 14400 | 587.655 | 587.571 | 587.79 | 587.585 | 587.602 | 587.64 | 0.09 |

**ONPF**

| Time | Replicate 1 | Replicate 2 | Replicate 3 | Replicate 4 | average | SD |
|---|---|---|---|---|---|---|
| 0 | 586.129 | 586.258 | 586.187 | 586.237 | 586.20 | 0.06 |
| 30 | 586.702 | 586.502 | | 586.744 | 586.65 | 0.13 |
| 45 | | 586.685 | | 587.214 | 586.95 | 0.37 |
| 60 | | 587.052 | 587.192 | 587.292 | 587.18 | 0.12 |
| 300 | 587.193 | 586.999 | 587.249 | 587.666 | 587.28 | 0.28 |
| 1500 | 587.746 | 587.225 | 587.499 | 587.97 | 587.61 | 0.32 |
| 3600 | 587.853 | 587.186 | | 587.745 | 587.59 | 0.36 |
| 7200 | | 587.062 | 587.592 | 587.695 | 587.45 | 0.34 |
| 14400 | 587.878 | 586.437 | 587.684 | 588.27 | 587.57 | 0.79 |

**APO**

| Time | Replicate 1 | Replicate 2 | Replicate 3 | Replicate 4 | Replicate 5 | average | SD |
|---|---|---|---|---|---|---|---|
| 0 | 586.288 | 586.253 | 586.186 | 586.185 | 586.128 | 586.21 | 0.06 |
| 30 | 586.864 | 586.778 | 586.478 | 586.772 | 586.661 | 586.71 | 0.15 |
| 45 | | 586.884 | 586.848 | | 587.142 | 586.96 | 0.16 |
| 60 | 587.433 | 587.289 | 587.261 | 587.26 | 587.152 | 587.28 | 0.10 |
| 300 | 587.344 | | 587.493 | 587.241 | 587.039 | 587.28 | 0.19 |
| 1500 | | 587.49 | 587.59 | 587.558 | 587.165 | 587.45 | 0.20 |
| 3600 | 587.81 | 587.595 | 587.668 | 587.74 | 586.749 | 587.51 | 0.43 |
| 7200 | 587.938 | | 587.813 | 587.605 | 587.442 | 587.70 | 0.22 |
| 14400 | 587.696 | 587.661 | 587.85 | 587.84 | 587.663 | 587.74 | 0.10 |

**DNA**

| Time | Replicate 1 | Replicate 2 | Replicate 3 | Replicate 4 | Replicate 5 | average | SD |
|---|---|---|---|---|---|---|---|
| 0 | 586.192 | 586.192 | 586.702 | 586.174 | | 586.32 | 0.26 |
| 30 | 586.713 | 586.798 | 587.338 | 586.967 | | 586.95 | 0.28 |
| 45 | | 586.968 | | 587.36 | | 587.16 | 0.28 |
| 60 | 587.473 | 587.241 | 587.373 | 587.31 | | 587.35 | 0.10 |
| 300 | 587.397 | 587.401 | 587.658 | 587.544 | | 587.50 | 0.13 |
| 1500 | 587.649 | 587.733 | 587.271 | 587.832 | | 587.62 | 0.25 |
| 3600 | 587.703 | 587.389 | 587.616 | 587.99 | | 587.67 | 0.25 |
| 7200 | 587.497 | 587.179 | 587.625 | 587.797 | | 587.52 | 0.26 |
| 14400 | 587.625 | 586.991 | 587.692 | 587.942 | | 587.56 | 0.40 |

**ONPFDNA**

| Time | Replicate 1 | Replicate 2 | Replicate 3 | Replicate 4 | average | SD |
|---|---|---|---|---|---|---|
| 0 | 586.149 | 586.163 | 586.185 | 586.171 | 586.17 | 0.02 |
| 30 | 586.842 | 586.66 | 586.532 | 586.871 | 586.73 | 0.16 |
| 45 | | 586.976 | | 587.229 | 587.10 | 0.18 |
| 60 | | 587.147 | 587.258 | 587.23 | 587.21 | 0.06 |
| 300 | 587.303 | 587.161 | 587.342 | 587.619 | 587.36 | 0.19 |
| 1500 | 587.687 | 587.407 | 587.476 | 587.288 | 587.46 | 0.17 |
| 3600 | 587.628 | | 587.651 | 587.949 | 587.74 | 0.18 |
| 7200 | | 587.131 | 587.555 | 587.762 | 587.48 | 0.32 |
| 14400 | 587.774 | 587.225 | 587.635 | 587.346 | 587.50 | 0.25 |

**TMG**

| Time | Replicate 1 | Replicate 2 | Replicate 3 | Replicate 4 | Replicate 5 | average | SD |
|---|---|---|---|---|---|---|---|
| 0 | 586.23 | 586.156 | | | | 586.19 | 0.05 |
| 30 | 586.533 | 586.662 | | | | 586.60 | 0.09 |
| 45 | 586.955 | 586.942 | | | | 586.95 | 0.01 |
| 60 | 587.085 | 587.312 | | | | 587.20 | 0.16 |
| 300 | 587.061 | 587.069 | | | | 587.07 | 0.01 |
| 1500 | 587.131 | 587.477 | | | | 587.30 | 0.24 |
| 3600 | 587.72 | 587.866 | | | | 587.79 | 0.10 |
| 7200 | | 588.024 | | | | 588.02 | |
| 14400 | | 588.023 | | | | 588.02 | |

### 187-195 GPLSSVSAR

**Charge** 2

**IPTG**

| Time | centroid | D |
|---|---|---|
| 0 | 437.51 | 0.03 |
| 30 | 437.96 | 1.26 |
| 45 | 438.08 | 1.57 |
| 60 | 438.18 | 1.85 |
| 300 | 438.26 | 2.06 |
| 1500 | 438.67 | 3.15 |
| 3600 | 438.81 | 3.55 |
| 7200 | 439.13 | 4.41 |
| 14400 | 438.87 | 3.71 |

**ONPF**

| Time | centroid | D |
|---|---|---|
| 0 | 437.54 | 0.11 |
| 30 | 438.03 | 1.43 |
| 45 | 438.40 | 2.43 |
| 60 | 438.54 | 2.80 |
| 300 | 438.68 | 3.18 |
| 1500 | 438.60 | 2.97 |
| 3600 | 438.87 | 3.70 |
| 7200 | 438.82 | 3.57 |
| 14400 | 438.38 | 2.37 |

**APO**

| Time | centroid | D |
|---|---|---|
| 0 | 437.50 | 0.01 |
| 30 | 438.31 | 2.19 |
| 45 | 438.31 | 2.18 |
| 60 | 438.66 | 3.13 |
| 300 | 438.49 | 2.68 |
| 1500 | 438.60 | 2.98 |
| 3600 | 438.94 | 3.89 |
| 7200 | 438.94 | 3.90 |
| 14400 | 439.18 | 4.54 |

**DNA**

| Time | centroid | D |
|---|---|---|
| 0 | 437.51 | 0.02 |
| 30 | 438.19 | 1.85 |
| 45 | 438.58 | 2.92 |
| 60 | 438.58 | 2.91 |
| 300 | 438.77 | 3.42 |
| 1500 | 438.86 | 3.67 |
| 3600 | 438.74 | 3.34 |
| 7200 | 438.82 | 3.57 |
| 14400 | 438.67 | 3.15 |

**ONPFDNA**

| Time | centroid | D |
|---|---|---|
| 0 | 437.51 | 0.02 |
| 30 | 438.11 | 1.66 |
| 45 | 438.37 | 2.35 |
| 60 | 438.43 | 2.52 |
| 300 | 438.59 | 2.94 |
| 1500 | 438.59 | 2.95 |
| 3600 | 439.11 | 4.36 |
| 7200 | 438.77 | 3.43 |
| 14400 | 438.54 | 2.80 |

**TMG**

| Time | centroid | D |
|---|---|---|
| 0 | 437.52 | 0.04 |
| 30 | 437.97 | 1.27 |
| 45 | 438.19 | 1.86 |
| 60 | 438.13 | 1.71 |
| 300 | 438.37 | 2.36 |
| 1500 | 438.65 | 3.10 |
| 3600 | 439.07 | 4.25 |
| 7200 | 439.23 | 4.68 |
| 14400 | 439.51 | 5.45 |

control 437.5
infinity 440.088

$$D(t) = \frac{M_t - M_0}{M_\infty - M_0} * N$$

**IPTG**

| Time | Replicate 1 | Replicate 2 | Replicate 3 | Replicate 4 | Replicate5 | average | SD |
|---|---|---|---|---|---|---|---|
| 0 | 437.512 | 437.509 | | | | 437.51 | 0.00 |
| 30 | 438.073 | 437.855 | | | | 437.96 | 0.15 |
| 45 | 438.156 | 438.004 | | | | 438.08 | 0.11 |
| 60 | 438.255 | 438.11 | | | | 438.18 | 0.10 |
| 300 | 438.287 | 438.233 | | | | 438.26 | 0.04 |
| 1500 | 438.75 | 438.58 | | | | 438.67 | 0.12 |
| 3600 | 438.649 | 438.978 | | | | 438.81 | 0.23 |
| 7200 | 439.228 | 439.031 | | | | 439.13 | 0.14 |
| 14400 | 439.179 | 438.562 | | | | 438.87 | 0.44 |

**ONPF**

| Time | Replicate 1 | Replicate 2 | Replicate 3 | Replicate 4 | average | SD |
|---|---|---|---|---|---|---|
| 0 | 437.55 | 437.53 | | | 437.54 | 0.01 |
| 30 | 438.117 | 437.938 | | | 438.03 | 0.13 |
| 45 | 438.702 | 438.096 | | | 438.40 | 0.43 |
| 60 | 438.575 | 438.496 | | | 438.54 | 0.06 |
| 300 | 438.781 | 438.57 | | | 438.68 | 0.15 |
| 1500 | 438.613 | 438.583 | | | 438.60 | 0.02 |
| 3600 | 439.017 | 438.721 | | | 438.87 | 0.21 |
| 7200 | 438.983 | 438.654 | | | 438.82 | 0.23 |
| 14400 | 438.642 | 438.112 | | | 438.38 | 0.37 |

**APO**

| Time | Replicate 1 | Replicate 2 | Replicate 3 | Replicate 4 | Replicate 5 | average | SD |
|---|---|---|---|---|---|---|---|
| 0 | 437.489 | 437.516 | | | | 437.50 | 0.02 |
| 30 | 438.448 | 438.169 | | | | 438.31 | 0.20 |
| 45 | 438.363 | 438.251 | | | | 438.31 | 0.08 |
| 60 | 438.732 | 438.581 | | | | 438.66 | 0.11 |
| 300 | | 438.489 | | | | 438.49 | |
| 1500 | 438.618 | 438.587 | | | | 438.60 | 0.02 |
| 3600 | 439.308 | 438.567 | | | | 438.94 | 0.52 |
| 7200 | 439.328 | 438.557 | | | | 438.94 | 0.55 |
| 14400 | 439.306 | 439.053 | | | | 439.18 | 0.18 |

**DNA**

| Time | Replicate 1 | Replicate 2 | Replicate 3 | Replicate 4 | Replicate 5 | average | SD |
|---|---|---|---|---|---|---|---|
| 0 | 437.501 | 437.516 | | | | 437.51 | 0.01 |
| 30 | 438.166 | 438.205 | | | | 438.19 | 0.03 |
| 45 | 438.6 | 438.556 | | | | 438.58 | 0.03 |
| 60 | 438.604 | 438.551 | | | | 438.58 | 0.04 |
| 300 | 438.755 | 438.775 | | | | 438.77 | 0.01 |
| 1500 | 438.619 | 439.095 | | | | 438.86 | 0.34 |
| 3600 | 438.644 | 438.829 | | | | 438.74 | 0.13 |
| 7200 | 439.092 | 438.551 | | | | 438.82 | 0.38 |
| 14400 | 439.056 | 438.276 | | | | 438.67 | 0.55 |

**ONPFDNA**

| Time | Replicate 1 | Replicate 2 | Replicate 3 | Replicate 4 | average | SD |
|---|---|---|---|---|---|---|
| 0 | 437.505 | 437.509 | | | 437.51 | 0.00 |
| 30 | 438.166 | 438.061 | | | 438.11 | 0.07 |
| 45 | 438.48 | 438.258 | | | 438.37 | 0.16 |
| 60 | 438.411 | 438.456 | | | 438.43 | 0.03 |
| 300 | 438.587 | 438.585 | | | 438.59 | 0.00 |
| 1500 | 438.614 | 438.565 | | | 438.59 | 0.03 |
| 3600 | 439.122 | 439.1 | | | 439.11 | 0.02 |
| 7200 | 438.976 | 438.558 | | | 438.77 | 0.30 |
| 14400 | 438.54 | 438.533 | | | 438.54 | 0.00 |

**TMG**

| Time | Replicate 1 | Replicate 2 | Replicate 3 | Replicate 4 | Replicate 5 | average | SD |
|---|---|---|---|---|---|---|---|
| 0 | 437.529 | 437.501 | | | | 437.52 | 0.02 |
| 30 | 437.824 | 438.118 | | | | 437.97 | 0.21 |
| 45 | 438.258 | 438.121 | | | | 438.19 | 0.10 |
| 60 | 438.133 | | | | | 438.13 | |
| 300 | 438.35 | 438.394 | | | | 438.37 | 0.03 |
| 1500 | 438.495 | 438.795 | | | | 438.65 | 0.21 |
| 3600 | 438.944 | 439.198 | | | | 439.07 | 0.18 |
| 7200 | 439.035 | 439.426 | | | | 439.23 | 0.28 |
| 14400 | | 439.514 | | | | 439.51 | |

### 198-203 LAGWHK

**Charge** 1

**IPTG**

| Time | centroid | D |
|---|---|---|
| 0 | 711.87 | 0.04 |
| 30 | 712.05 | 0.41 |
| 45 | 712.03 | 0.38 |
| 60 | 712.10 | 0.52 |
| 300 | 712.03 | 0.37 |
| 1500 | 712.04 | 0.40 |
| 3600 | 711.97 | 0.25 |
| 7200 | 711.98 | 0.26 |
| 14400 | 712.03 | 0.38 |

**ONPF**

| Time | centroid | D |
|---|---|---|
| 0 | 711.87 | 0.04 |
| 30 | 711.95 | 0.20 |
| 45 | 711.99 | 0.29 |
| 60 | 711.99 | 0.28 |
| 300 | 711.99 | 0.29 |
| 1500 | 711.97 | 0.25 |
| 3600 | 711.96 | 0.24 |
| 7200 | 712.03 | 0.36 |
| 14400 | 712.03 | 0.37 |

**APO**

| Time | centroid | D |
|---|---|---|
| 0 | 711.85 | 0.01 |
| 30 | 711.95 | 0.21 |
| 45 | 712.02 | 0.35 |
| 60 | 711.96 | 0.22 |
| 300 | 712.02 | 0.35 |
| 1500 | 711.98 | 0.26 |
| 3600 | 711.98 | 0.26 |
| 7200 | 711.98 | 0.27 |
| 14400 | 712.13 | 0.58 |

**DNA**

| Time | centroid | D |
|---|---|---|
| 0 | 711.92 | 0.15 |
| 30 | 712.05 | 0.42 |
| 45 | 712.06 | 0.42 |
| 60 | 711.97 | 0.25 |
| 300 | 712.04 | 0.40 |
| 1500 | 712.03 | 0.37 |
| 3600 | 712.07 | 0.45 |
| 7200 | 712.02 | 0.36 |
| 14400 | 712.01 | 0.32 |

**ONPFDNA**

| Time | centroid | D |
|---|---|---|
| 0 | 711.81 | -0.08 |
| 30 | 712.01 | 0.33 |
| 45 | 711.97 | 0.25 |
| 60 | 711.97 | 0.24 |
| 300 | 712.03 | 0.37 |
| 1500 | 711.95 | 0.22 |
| 3600 | 711.97 | 0.24 |
| 7200 | 711.96 | 0.23 |
| 14400 | 712.02 | 0.35 |

**TMG**

| Time | centroid | D |
|---|---|---|
| 0 | 711.85 | -0.01 |
| 30 | 711.98 | 0.26 |
| 45 | 712.01 | 0.32 |
| 60 | 711.99 | 0.30 |
| 300 | 712.11 | 0.53 |
| 1500 | 712.03 | 0.36 |
| 3600 | 712.06 | 0.44 |
| 7200 | 711.99 | 0.29 |
| 14400 | 712.06 | 0.44 |

control 711.85
infinity 713.783

$$D(t) = \frac{M_t - M_0}{M_\infty - M_0} \cdot N$$

**IPTG**

| Time | Replicate 1 | Replicate 2 | Replicate 3 | Replicate 4 | Replicate5 | average | SD |
|---|---|---|---|---|---|---|---|
| 0 | 711.849 | 711.887 | | | | 711.87 | 0.03 |
| 30 | 712.058 | 712.034 | | | | 712.05 | 0.02 |
| 45 | 712.053 | 712.01 | | | | 712.03 | 0.03 |
| 60 | 712.092 | 712.113 | | | | 712.10 | 0.01 |
| 300 | 712.093 | 711.968 | | | | 712.03 | 0.09 |
| 1500 | 711.974 | 712.111 | | | | 712.04 | 0.10 |
| 3600 | 712.019 | 711.921 | | | | 711.97 | 0.07 |
| 7200 | 712.016 | 711.937 | | | | 711.98 | 0.06 |
| 14400 | 712.052 | 712.015 | | | | 712.03 | 0.03 |

**ONPF**

| Time | Replicate 1 | Replicate 2 | Replicate 3 | Replicate 4 | average | SD |
|---|---|---|---|---|---|---|
| 0 | 711.822 | 711.919 | | | 711.87 | 0.07 |
| 30 | 711.954 | 711.942 | | | 711.95 | 0.01 |
| 45 | 712.017 | 711.968 | | | 711.99 | 0.03 |
| 60 | 711.985 | 711.987 | | | 711.99 | 0.00 |
| 300 | 711.988 | 711.997 | | | 711.99 | 0.01 |
| 1500 | 711.99 | 711.953 | | | 711.97 | 0.03 |
| 3600 | 711.977 | 711.951 | | | 711.96 | 0.02 |
| 7200 | 711.988 | 712.064 | | | 712.03 | 0.05 |
| 14400 | 712.027 | | | | 712.03 | 0.00 |

**APO**

| Time | Replicate 1 | Replicate 2 | Replicate 3 | Replicate 4 | Replicate 5 | average | SD |
|---|---|---|---|---|---|---|---|
| 0 | 711.851 | 711.857 | | | | 711.85 | 0.00 |
| 30 | 712.001 | 711.898 | | | | 711.95 | 0.07 |
| 45 | 712.028 | 712.009 | | | | 712.02 | 0.01 |
| 60 | 711.985 | 711.927 | | | | 711.96 | 0.04 |
| 300 | 712.019 | 712.016 | | | | 712.02 | 0.00 |
| 1500 | 711.966 | 711.986 | | | | 711.98 | 0.01 |
| 3600 | 711.991 | 711.959 | | | | 711.98 | 0.02 |
| 7200 | 712.047 | 711.917 | | | | 711.98 | 0.09 |
| 14400 | 712.222 | 712.039 | | | | 712.13 | 0.13 |

**DNA**

| Time | Replicate 1 | Replicate 2 | Replicate 3 | Replicate 4 | Replicate 5 | average | SD |
|---|---|---|---|---|---|---|---|
| 0 | 711.824 | 712.022 | | | | 711.92 | 0.14 |
| 30 | 711.984 | 712.122 | | | | 712.05 | 0.10 |
| 45 | 711.963 | 712.147 | | | | 712.06 | 0.13 |
| 60 | 711.959 | 711.984 | | | | 711.97 | 0.02 |
| 300 | 711.97 | 712.119 | | | | 712.04 | 0.11 |
| 1500 | 712.07 | 711.985 | | | | 712.03 | 0.06 |
| 3600 | 712.027 | 712.105 | | | | 712.07 | 0.06 |
| 7200 | 711.978 | 712.07 | | | | 712.02 | 0.07 |
| 14400 | 712.006 | 712.006 | | | | 712.01 | 0.00 |

**ONPFDNA**

| Time | Replicate 1 | Replicate 2 | Replicate 3 | Replicate 4 | average | SD |
|---|---|---|---|---|---|---|
| 0 | 711.832 | 711.792 | | | 711.81 | 0.03 |
| 30 | 711.957 | 712.065 | | | 712.01 | 0.08 |
| 45 | 711.984 | 711.961 | | | 711.97 | 0.02 |
| 60 | 711.993 | 711.943 | | | 711.97 | 0.04 |
| 300 | 711.973 | 712.083 | | | 712.03 | 0.08 |
| 1500 | 711.93 | 711.978 | | | 711.95 | 0.03 |
| 3600 | 711.967 | | | | 711.97 | |
| 7200 | 711.979 | 711.944 | | | 711.96 | 0.02 |
| 14400 | 711.922 | 712.118 | | | 712.02 | 0.14 |

**TMG**

| Time | Replicate 1 | Replicate 2 | Replicate 3 | Replicate 4 | Replicate 5 | average | SD |
|---|---|---|---|---|---|---|---|
| 0 | 711.815 | 711.876 | | | | 711.85 | 0.04 |
| 30 | 711.881 | 712.073 | | | | 711.98 | 0.14 |
| 45 | 711.89 | 712.121 | | | | 712.01 | 0.16 |
| 60 | 711.994 | | | | | 711.99 | |
| 300 | 712.106 | 712.111 | | | | 712.11 | 0.00 |
| 1500 | 711.859 | 712.193 | | | | 712.03 | 0.24 |
| 3600 | 712.034 | 712.09 | | | | 712.06 | 0.04 |
| 7200 | 711.868 | 712.112 | | | | 711.99 | 0.17 |
| 14400 | 711.922 | 712.205 | | | | 712.06 | 0.20 |

### 204-212 YLTRNQIQP

**Charge** 2

**IPTG**

| Time | centroid | D |
|---|---|---|
| 0 | 567.1345 | 0.01 |
| 30 | 567.5975 | 1.15 |
| 45 | 567.7285 | 1.48 |
| 60 | 567.768 | 1.58 |
| 300 | 567.845 | 1.77 |
| 1500 | 568.062 | 2.30 |
| 3600 | 568.3575 | 3.03 |
| 7200 | 568.5455 | 3.50 |
| 14400 | 568.4985 | 3.38 |

**ONPF**

| Time | centroid | D |
|---|---|---|
| 0 | 567.142 | 0.03 |
| 30 | 567.382 | 0.62 |
| 45 | 567.6465 | 1.28 |
| 60 | 567.732 | 1.49 |
| 300 | 567.76 | 1.56 |
| 1500 | 567.9385 | 2.00 |
| 3600 | 568.0275 | 2.22 |
| 7200 | 568.108 | 2.41 |
| 14400 | 568.32 | 2.93 |

**APO**

| Time | centroid | D |
|---|---|---|
| 0 | 567.131 | 0.00 |
| 30 | 567.5225 | 0.97 |
| 45 | 567.592 | 1.14 |
| 60 | 567.775 | 1.59 |
| 300 | 567.724 | 1.47 |
| 1500 | 568.243 | 2.75 |
| 3600 | 568.2395 | 2.74 |
| 7200 | 568.434 | 3.22 |
| 14400 | 568.5435 | 3.49 |

**DNA**

| Time | centroid | D |
|---|---|---|
| 0 | 567.125 | -0.01 |
| 30 | 567.5345 | 1.00 |
| 45 | 567.6395 | 1.26 |
| 60 | 567.668 | 1.33 |
| 300 | 567.773 | 1.59 |
| 1500 | 568.024 | 2.21 |
| 3600 | 568.0925 | 2.38 |
| 7200 | 568.301 | 2.89 |
| 14400 | 568.557 | 3.52 |

**ONPFDNA**

| Time | centroid | D |
|---|---|---|
| 0 | 567.121 | -0.02 |
| 30 | 567.538 | 1.01 |
| 45 | 567.6065 | 1.18 |
| 60 | 567.6745 | 1.34 |
| 300 | 567.7655 | 1.57 |
| 1500 | 567.796 | 1.64 |
| 3600 | 568.141 | 2.50 |
| 7200 | 568.3155 | 2.93 |
| 14400 | | |

**TMG**

| Time | centroid | D |
|---|---|---|
| 0 | 567.1315 | 0.00 |
| 30 | 567.59 | 1.14 |
| 45 | 567.713 | 1.44 |
| 60 | 567.794 | 1.64 |
| 300 | 567.8815 | 1.86 |
| 1500 | 568.155 | 2.53 |
| 3600 | 568.4125 | 3.17 |
| 7200 | 568.568 | 3.55 |
| 14400 | 568.7 | 3.88 |

control 567.13
infinity 569.56

$$D(t) = \frac{M_t - M_0}{M_\infty - M_0} * N$$

**IPTG**

| Time | Replicate 1 | Replicate 2 | Replicate 3 | Replicate 4 | Replicate5 | average | SD |
|---|---|---|---|---|---|---|---|
| 0 | 567.152 | 567.117 | | | | 567.13 | 0.02 |
| 30 | 567.761 | 567.434 | | | | 567.60 | 0.23 |
| 45 | 567.803 | 567.654 | | | | 567.73 | 0.11 |
| 60 | 567.816 | 567.72 | | | | 567.77 | 0.07 |
| 300 | 567.953 | 567.737 | | | | 567.85 | 0.15 |
| 1500 | 568.265 | 567.859 | | | | 568.06 | 0.29 |
| 3600 | 568.469 | 568.246 | | | | 568.36 | 0.16 |
| 7200 | 568.735 | 568.356 | | | | 568.55 | 0.27 |
| 14400 | 568.752 | 568.245 | | | | 568.50 | 0.36 |

**ONPF**

| Time | Replicate 1 | Replicate 2 | Replicate 3 | Replicate 4 | average | SD |
|---|---|---|---|---|---|---|
| 0 | 567.134 | 567.15 | | | 567.14 | 0.01 |
| 30 | 567.395 | 567.369 | | | 567.38 | 0.02 |
| 45 | 567.904 | 567.389 | | | 567.65 | 0.36 |
| 60 | 567.741 | 567.723 | | | 567.73 | 0.01 |
| 300 | 567.89 | 567.63 | | | 567.76 | 0.18 |
| 1500 | 567.988 | 567.889 | | | 567.94 | 0.07 |
| 3600 | 568.128 | 567.927 | | | 568.03 | 0.14 |
| 7200 | 568.114 | 568.102 | | | 568.11 | 0.01 |
| 14400 | 568.412 | 568.225 | | | 568.32 | 0.13 |

**APO**

| Time | Replicate 1 | Replicate 2 | Replicate 3 | Replicate 4 | Replicate 5 | average | SD |
|---|---|---|---|---|---|---|---|
| 0 | 567.14 | 567.122 | | | | 567.13 | 0.01 |
| 30 | 567.583 | 567.462 | | | | 567.52 | 0.09 |
| 45 | 567.633 | 567.551 | | | | 567.59 | 0.06 |
| 60 | 567.835 | 567.715 | | | | 567.78 | 0.08 |
| 300 | 567.887 | 567.561 | | | | 567.72 | 0.23 |
| 1500 | 568.474 | 568.012 | | | | 568.24 | 0.33 |
| 3600 | 568.353 | 568.126 | | | | 568.24 | 0.16 |
| 7200 | 568.665 | 568.203 | | | | 568.43 | 0.33 |
| 14400 | 568.809 | 568.278 | | | | 568.54 | 0.38 |

**DNA**

| Time | Replicate 1 | Replicate 2 | Replicate 3 | Replicate 4 | Replicate 5 | average | SD |
|---|---|---|---|---|---|---|---|
| 0 | 567.13 | 567.12 | | | | 567.13 | 0.01 |
| 30 | 567.538 | 567.531 | | | | 567.53 | 0.00 |
| 45 | 567.757 | 567.522 | | | | 567.64 | 0.17 |
| 60 | 567.666 | 567.67 | | | | 567.67 | 0.00 |
| 300 | 567.845 | 567.701 | | | | 567.77 | 0.10 |
| 1500 | 568.019 | 568.029 | | | | 568.02 | 0.01 |
| 3600 | 568.212 | 567.973 | | | | 568.09 | 0.17 |
| 7200 | 568.439 | 568.163 | | | | 568.30 | 0.20 |
| 14400 | 568.557 | | | | | 568.56 | |

**ONPFDNA**

| Time | Replicate 1 | Replicate 2 | Replicate 3 | Replicate 4 | average | SD |
|---|---|---|---|---|---|---|
| 0 | 567.122 | 567.12 | | | 567.12 | 0.00 |
| 30 | 567.488 | 567.588 | | | 567.54 | 0.07 |
| 45 | 567.649 | 567.564 | | | 567.61 | 0.06 |
| 60 | 567.726 | 567.623 | | | 567.67 | 0.07 |
| 300 | 567.835 | 567.696 | | | 567.77 | 0.10 |
| 1500 | 567.714 | 567.878 | | | 567.80 | 0.12 |
| 3600 | 568.149 | 568.133 | | | 568.14 | 0.01 |
| 7200 | 568.3 | 568.331 | | | 568.32 | 0.02 |
| 14400 | | | | | #DIV/0! | |

**TMG**

| Time | Replicate 1 | Replicate 2 | Replicate 3 | Replicate 4 | Replicate 5 | average | SD |
|---|---|---|---|---|---|---|---|
| 0 | 567.14 | 567.123 | | | | 567.13 | 0.01 |
| 30 | 567.439 | 567.741 | | | | 567.59 | 0.21 |
| 45 | | 567.713 | | | | 567.71 | |
| 60 | 567.723 | 567.865 | | | | 567.79 | 0.10 |
| 300 | 567.833 | 567.93 | | | | 567.88 | 0.07 |
| 1500 | | 568.155 | | | | 568.16 | |
| 3600 | 568.295 | 568.53 | | | | 568.41 | 0.17 |
| 7200 | | 568.568 | | | | 568.57 | |
| 14400 | 568.52 | 568.88 | | | | 568.70 | 0.25 |

#### 208-219 NQIQPIAEREGD

**Charge** 2

**IPTG**

| Time | centroid | D |
|---|---|---|
| 0 | 685.732 | 0.04 |
| 30 | 686.5615 | 2.47 |
| 45 | 686.7235 | 2.94 |
| 60 | 686.768 | 3.07 |
| 300 | 686.8305 | 3.25 |
| 1500 | 686.7885 | 3.13 |
| 3600 | 686.993 | 3.73 |
| 7200 | 687.1135 | 4.08 |
| 14400 | 687.2465 | 4.47 |

**ONPF**

| Time | centroid | D |
|---|---|---|
| 0 | 685.74 | 0.06 |
| 30 | 686.29 | 1.66 |
| 45 | 686.58 | 2.52 |
| 60 | 686.59 | 2.56 |
| 300 | 686.66 | 2.75 |
| 1500 | 686.67 | 2.78 |
| 3600 | 686.72 | 2.93 |
| 7200 | 686.78 | 3.10 |
| 14400 | 687.23 | 4.43 |

**APO**

| Time | centroid | D |
|---|---|---|
| 0 | 685.723 | 0.01 |
| 30 | 686.516 | 2.33 |
| 45 | 686.563 | 2.47 |
| 60 | 686.7005 | 2.87 |
| 300 | 686.6115 | 2.61 |
| 1500 | 686.872 | 3.38 |
| 3600 | 686.9435 | 3.59 |
| 7200 | 687.029 | 3.84 |
| 14400 | 687.334 | 4.73 |

**DNA**

| Time | centroid | D |
|---|---|---|
| 0 | 685.72 | 0.00 |
| 30 | 686.44 | 2.12 |
| 45 | 686.53 | 2.37 |
| 60 | 686.53 | 2.38 |
| 300 | 686.60 | 2.57 |
| 1500 | 686.70 | 2.86 |
| 3600 | 686.73 | 2.95 |
| 7200 | 686.77 | 3.08 |
| 14400 | 686.66 | 2.76 |

**ONPFDNA**

| Time | centroid | D |
|---|---|---|
| 0 | 685.73 | 0.02 |
| 30 | 686.41 | 2.03 |
| 45 | 686.58 | 2.52 |
| 60 | 686.51 | 2.32 |
| 300 | 686.57 | 2.49 |
| 1500 | 686.46 | 2.17 |
| 3600 | 686.84 | 3.28 |
| 7200 | 686.91 | 3.47 |
| 14400 | 686.67 | 2.77 |

**TMG**

| Time | centroid | D |
|---|---|---|
| 0 | 685.73 | 0.02 |
| 30 | 686.53 | 2.37 |
| 45 | 686.71 | 2.89 |
| 60 | 686.89 | 3.42 |
| 300 | 686.82 | 3.23 |
| 1500 | 686.85 | 3.31 |
| 3600 | 687.06 | 3.93 |
| 7200 | 687.04 | 3.88 |
| 14400 | 687.27 | 4.55 |

control 685.72
infinity 688.791

$$D(t) = \frac{Mt - M0}{M\infty - M0} *N$$

**IPTG**

| Time | Replicate 1 | Replicate 2 | Replicate 3 | Replicate 4 | Replicate5 | average | SD |
|---|---|---|---|---|---|---|---|
| 0 | 685.739 | 685.725 | | | | 685.73 | 0.01 |
| 30 | 686.701 | 686.422 | | | | 686.56 | 0.20 |
| 45 | 686.794 | 686.653 | | | | 686.72 | 0.10 |
| 60 | 686.853 | 686.683 | | | | 686.77 | 0.12 |
| 300 | 687.069 | 686.592 | | | | 686.83 | 0.34 |
| 1500 | 686.957 | 686.62 | | | | 686.79 | 0.24 |
| 3600 | 687.086 | 686.9 | | | | 686.99 | 0.13 |
| 7200 | 687.274 | 686.953 | | | | 687.11 | 0.23 |
| 14400 | 687.454 | 687.039 | | | | 687.25 | 0.29 |

**ONPF**

| Time | Replicate 1 | Replicate 2 | Replicate 3 | Replicate 4 | average | SD |
|---|---|---|---|---|---|---|
| 0 | 685.737 | 685.744 | | | 685.74 | 0.00 |
| 30 | 686.329 | 686.245 | | | 686.29 | 0.06 |
| 45 | 686.781 | 686.379 | | | 686.58 | 0.28 |
| 60 | 686.603 | 686.584 | | | 686.59 | 0.01 |
| 300 | 686.894 | 686.421 | | | 686.66 | 0.33 |
| 1500 | 686.807 | 686.532 | | | 686.67 | 0.19 |
| 3600 | 686.86 | 686.582 | | | 686.72 | 0.20 |
| 7200 | 686.889 | 686.666 | | | 686.78 | 0.16 |
| 14400 | 687.231 | | | | 687.23 | |

**APO**

| Time | Replicate 1 | Replicate 2 | Replicate 3 | Replicate 4 | Replicate 5 | average | SD |
|---|---|---|---|---|---|---|---|
| 0 | 685.717 | 685.729 | | | | 685.72 | 0.01 |
| 30 | 686.66 | 686.372 | | | | 686.52 | 0.20 |
| 45 | 686.719 | 686.407 | | | | 686.56 | 0.22 |
| 60 | 686.797 | 686.604 | | | | 686.70 | 0.14 |
| 300 | 686.807 | 686.416 | | | | 686.61 | 0.28 |
| 1500 | 687.002 | 686.742 | | | | 686.87 | 0.18 |
| 3600 | 687.067 | 686.82 | | | | 686.94 | 0.17 |
| 7200 | 687.237 | 686.821 | | | | 687.03 | 0.29 |
| 14400 | 687.594 | 687.074 | | | | 687.33 | 0.37 |

**DNA**

| Time | Replicate 1 | Replicate 2 | Replicate 3 | Replicate 4 | Replicate 5 | average | SD |
|---|---|---|---|---|---|---|---|
| 0 | 685.706 | 685.732 | | | | 685.72 | 0.02 |
| 30 | 686.472 | 686.417 | | | | 686.44 | 0.04 |
| 45 | 686.596 | 686.464 | | | | 686.53 | 0.09 |
| 60 | 686.552 | 686.511 | | | | 686.53 | 0.03 |
| 300 | 686.64 | 686.553 | | | | 686.60 | 0.06 |
| 1500 | 686.669 | 686.723 | | | | 686.70 | 0.04 |
| 3600 | 686.876 | 686.577 | | | | 686.73 | 0.21 |
| 7200 | 686.947 | 686.595 | | | | 686.77 | 0.25 |
| 14400 | 687.122 | 686.199 | | | | 686.66 | 0.65 |

**ONPFDNA**

| Time | Replicate 1 | Replicate 2 | Replicate 3 | Replicate 4 | average | SD |
|---|---|---|---|---|---|---|
| 0 | 685.725 | 685.729 | | | 685.73 | 0.00 |
| 30 | 686.4 | 686.424 | | | 686.41 | 0.02 |
| 45 | 686.592 | 686.571 | | | 686.58 | 0.01 |
| 60 | 686.51 | 686.513 | | | 686.51 | 0.00 |
| 300 | 686.623 | 686.515 | | | 686.57 | 0.08 |
| 1500 | 686.382 | 686.536 | | | 686.46 | 0.11 |
| 3600 | 686.897 | 686.781 | | | 686.84 | 0.08 |
| 7200 | 687.039 | 686.772 | | | 686.91 | 0.19 |
| 14400 | | 686.666 | | | 686.67 | |

**TMG**

| Time | Replicate 1 | Replicate 2 | Replicate 3 | Replicate 4 | Replicate 5 | average | SD |
|---|---|---|---|---|---|---|---|
| 0 | 685.734 | 685.723 | | | | 685.73 | 0.01 |
| 30 | 686.306 | 686.751 | | | | 686.53 | 0.31 |
| 45 | 686.702 | 686.708 | | | | 686.71 | 0.00 |
| 60 | 686.747 | 687.026 | | | | 686.89 | 0.20 |
| 300 | | 686.821 | | | | 686.82 | |
| 1500 | 686.686 | 687.01 | | | | 686.85 | 0.23 |
| 3600 | | 687.062 | | | | 687.06 | |
| 7200 | 686.877 | 687.21 | | | | 687.04 | 0.24 |
| 14400 | 687.174 | 687.369 | | | | 687.27 | 0.14 |

### 213-225 IAEREGDWSAMSG

**Charge** 2

**IPTG**

| Time | centroid | D |
|---|---|---|
| 0 | 705.219 | -0.04 |
| 30 | 705.9905 | 2.67 |
| 45 | 706.129 | 3.15 |
| 60 | 706.229 | 3.50 |
| 300 | 706.506 | 4.48 |
| 1500 | 706.5925 | 4.78 |
| 3600 | 706.63 | 4.91 |
| 7200 | 706.6635 | 5.03 |
| 14400 | 706.619 | 4.87 |

**ONPF**

| Time | centroid | D |
|---|---|---|
| 0 | 705.2605 | 0.11 |
| 30 | 705.61 | 1.33 |
| 45 | 705.9085 | 2.38 |
| 60 | 706.047 | 2.87 |
| 300 | 706.136 | 3.18 |
| 1500 | 706.367 | 3.99 |
| 3600 | 706.4005 | 4.11 |
| 7200 | 706.298 | 3.75 |
| 14400 | 706.23 | 3.51 |

**APO**

| Time | centroid | D |
|---|---|---|
| 0 | 705.229 | 0.00 |
| 30 | 705.7945 | 1.98 |
| 45 | 705.906 | 2.37 |
| 60 | 706.0675 | 2.94 |
| 300 | 706.233 | 3.52 |
| 1500 | 706.5695 | 4.70 |
| 3600 | 706.6575 | 5.01 |
| 7200 | 706.6065 | 4.83 |
| 14400 | 706.862 | 5.72 |

**DNA**

| Time | centroid | D |
|---|---|---|
| 0 | 705.23 | 0.00 |
| 30 | 705.7125 | 1.69 |
| 45 | 705.916 | 2.41 |
| 60 | 705.983 | 2.64 |
| 300 | 706.1335 | 3.17 |
| 1500 | 706.442 | 4.25 |
| 3600 | 706.378 | 4.03 |
| 7200 | 706.319 | 3.82 |
| 14400 | 706.2385 | 3.54 |

**ONPFDNA**

| Time | centroid | D |
|---|---|---|
| 0 | 705.2265 | -0.01 |
| 30 | 705.6585 | 1.50 |
| 45 | 705.8425 | 2.15 |
| 60 | 705.9445 | 2.51 |
| 300 | 706.0315 | 2.81 |
| 1500 | 706.0985 | 3.05 |
| 3600 | 706.4995 | 4.45 |
| 7200 | 706.462 | 4.32 |
| 14400 | 706.0775 | 2.97 |

**TMG**

| Time | centroid | D |
|---|---|---|
| 0 | 705.2365 | 0.02 |
| 30 | 705.962 | 2.57 |
| 45 | 706.0965 | 3.04 |
| 60 | 706.407 | 4.13 |
| 300 | 706.5195 | 4.52 |
| 1500 | 706.633 | 4.92 |
| 3600 | 706.6935 | 5.13 |
| 7200 | 706.645 | 4.96 |
| 14400 | 706.793 | 5.48 |

control 705.23
infinity 708.366

$$D(t) = \frac{M_t - M_0}{M_\infty - M_0} * N$$

**IPTG**

| Time | Replicate 1 | Replicate 2 | Replicate 3 | Replicate 4 | Replicate5 | average | SD |
|---|---|---|---|---|---|---|---|
| 0 | 705.221 | 705.217 | | | | 705.22 | 0.00 |
| 30 | 706.213 | 705.768 | | | | 705.99 | 0.31 |
| 45 | 706.275 | 705.983 | | | | 706.13 | 0.21 |
| 60 | 706.301 | 706.157 | | | | 706.23 | 0.10 |
| 300 | 706.713 | 706.299 | | | | 706.51 | 0.29 |
| 1500 | 706.836 | 706.349 | | | | 706.59 | 0.34 |
| 3600 | 706.857 | 706.403 | | | | 706.63 | 0.32 |
| 7200 | 706.885 | 706.442 | | | | 706.66 | 0.31 |
| 14400 | 706.873 | 706.365 | | | | 706.62 | 0.36 |

**ONPF**

| Time | Replicate 1 | Replicate 2 | Replicate 3 | Replicate 4 | average | SD |
|---|---|---|---|---|---|---|
| 0 | 705.247 | 705.274 | | | 705.26 | 0.02 |
| 30 | 705.71 | 705.51 | | | 705.61 | 0.14 |
| 45 | 706.146 | 705.671 | | | 705.91 | 0.34 |
| 60 | 706.082 | 706.012 | | | 706.05 | 0.05 |
| 300 | 706.351 | 705.921 | | | 706.14 | 0.30 |
| 1500 | 706.585 | 706.149 | | | 706.37 | 0.31 |
| 3600 | 706.591 | 706.21 | | | 706.40 | 0.27 |
| 7200 | 706.435 | 706.161 | | | 706.30 | 0.19 |
| 14400 | 706.867 | 705.592 | | | 706.23 | 0.90 |

**APO**

| Time | Replicate 1 | Replicate 2 | Replicate 3 | Replicate 4 | Replicate 5 | average | SD |
|---|---|---|---|---|---|---|---|
| 0 | 705.218 | 705.24 | | | | 705.23 | 0.02 |
| 30 | 705.905 | 705.684 | | | | 705.79 | 0.16 |
| 45 | 706.069 | 705.743 | | | | 705.91 | 0.23 |
| 60 | 706.085 | 706.05 | | | | 706.07 | 0.02 |
| 300 | 706.496 | 705.97 | | | | 706.23 | 0.37 |
| 1500 | 706.786 | 706.353 | | | | 706.57 | 0.31 |
| 3600 | 706.876 | 706.439 | | | | 706.66 | 0.31 |
| 7200 | 706.918 | 706.295 | | | | 706.61 | 0.44 |
| 14400 | 707.221 | 706.503 | | | | 706.86 | 0.51 |

**DNA**

| Time | Replicate 1 | Replicate 2 | Replicate 3 | Replicate 4 | Replicate 5 | average | SD |
|---|---|---|---|---|---|---|---|
| 0 | 705.224 | 705.236 | | | | 705.23 | 0.01 |
| 30 | 705.768 | 705.657 | | | | 705.71 | 0.08 |
| 45 | 706.105 | 705.727 | | | | 705.92 | 0.27 |
| 60 | 706 | 705.966 | | | | 705.98 | 0.02 |
| 300 | 706.209 | 706.058 | | | | 706.13 | 0.11 |
| 1500 | 706.438 | 706.446 | | | | 706.44 | 0.01 |
| 3600 | 706.601 | 706.155 | | | | 706.38 | 0.32 |
| 7200 | 706.516 | 706.122 | | | | 706.32 | 0.28 |
| 14400 | 706.719 | 705.758 | | | | 706.24 | 0.68 |

**ONPFDNA**

| Time | Replicate 1 | Replicate 2 | Replicate 3 | Replicate 4 | average | SD |
|---|---|---|---|---|---|---|
| 0 | 705.233 | 705.22 | | | 705.23 | 0.01 |
| 30 | 705.735 | 705.582 | | | 705.66 | 0.11 |
| 45 | 705.961 | 705.724 | | | 705.84 | 0.17 |
| 60 | 705.963 | 705.926 | | | 705.94 | 0.03 |
| 300 | 706.153 | 705.91 | | | 706.03 | 0.17 |
| 1500 | 706.036 | 706.161 | | | 706.10 | 0.09 |
| 3600 | 706.571 | 706.428 | | | 706.50 | 0.10 |
| 7200 | 706.466 | 706.458 | | | 706.46 | 0.01 |
| 14400 | 706.122 | 706.033 | | | 706.08 | 0.06 |

**TMG**

| Time | Replicate 1 | Replicate 2 | Replicate 3 | Replicate 4 | Replicate 5 | average | SD |
|---|---|---|---|---|---|---|---|
| 0 | 705.243 | 705.23 | | | | 705.24 | 0.01 |
| 30 | 705.781 | 706.143 | | | | 705.96 | 0.26 |
| 45 | 706.169 | 706.024 | | | | 706.10 | 0.10 |
| 60 | 706.211 | 706.603 | | | | 706.41 | 0.28 |
| 300 | 706.423 | 706.616 | | | | 706.52 | 0.14 |
| 1500 | 706.407 | 706.859 | | | | 706.63 | 0.32 |
| 3600 | 706.532 | 706.855 | | | | 706.69 | 0.23 |
| 7200 | 706.418 | 706.872 | | | | 706.65 | 0.32 |
| 14400 | 706.623 | 706.963 | | | | 706.79 | 0.24 |

### 216-226 REGDWSAMSGF

**Charge** 2

**IPTG**

| Time | centroid | D |
|---|---|---|
| 0 | 622.13 | -0.05 |
| 30 | 622.40 | 0.91 |
| 45 | 622.52 | 1.32 |
| 60 | 622.69 | 1.93 |
| 300 | 622.82 | 2.36 |
| 1500 | 622.98 | 2.93 |
| 3600 | 622.98 | 2.93 |
| 7200 | 622.99 | 2.98 |
| 14400 | 623.01 | 3.04 |

**ONPF**

| Time | centroid | D |
|---|---|---|
| 0 | 622.16 | 0.06 |
| 30 | 622.33 | 0.65 |
| 45 | 622.63 | 1.73 |
| 60 | 622.62 | 1.69 |
| 300 | 622.63 | 1.72 |
| 1500 | 622.92 | 2.73 |
| 3600 | 623.20 | 3.71 |
| 7200 | 622.85 | 2.50 |
| 14400 | 623.07 | 3.27 |

**APO**

| Time | centroid | D |
|---|---|---|
| 0 | 622.1428 | 0.01 |
| 30 | 622.3478 | 0.73 |
| 45 | 622.415 | 0.96 |
| 60 | 622.5756 | 1.53 |
| 300 | 622.7176 | 2.02 |
| 1500 | 622.925 | 2.75 |
| 3600 | 622.9746 | 2.92 |
| 7200 | 622.971 | 2.91 |
| 14400 | 623.1148 | 3.41 |

**DNA**

| Time | centroid | D |
|---|---|---|
| 0 | 622.13 | -0.04 |
| 30 | 622.30 | 0.56 |
| 45 | 622.45 | 1.07 |
| 60 | 622.55 | 1.43 |
| 300 | 622.60 | 1.62 |
| 1500 | 622.80 | 2.30 |
| 3600 | 622.84 | 2.45 |
| 7200 | 622.74 | 2.11 |
| 14400 | 622.76 | 2.17 |

**ONPFDNA**

| Time | centroid | D |
|---|---|---|
| 0 | 622.06 | -0.28 |
| 30 | 622.24 | 0.36 |
| 45 | 622.35 | 0.72 |
| 60 | 622.49 | 1.23 |
| 300 | 622.55 | 1.44 |
| 1500 | 622.68 | 1.89 |
| 3600 | 622.91 | 2.70 |
| 7200 | 622.90 | 2.65 |
| 14400 | 622.77 | 2.21 |

**TMG**

| Time | centroid | D |
|---|---|---|
| 0 | 622.09 | -0.16 |
| 30 | 622.45 | 1.08 |
| 45 | 622.60 | 1.59 |
| 60 | 622.80 | 2.30 |
| 300 | 622.95 | 2.82 |
| 1500 | 623.09 | 3.34 |
| 3600 | 623.20 | 3.72 |
| 7200 | 623.01 | 3.04 |
| 14400 | 623.24 | 3.85 |

control 622.14
infinity 624.71

$$D(t) = \frac{Mt - M0}{M\infty - M0} * N$$

**IPTG**

| Time | Replicate 1 | Replicate 2 | Replicate 3 | Replicate 4 | Replicate5 | average | SD |
|---|---|---|---|---|---|---|---|
| 0 | 622.101 | 622.1 | 622.188 | 622.119 | 622.122 | 622.13 | 0.04 |
| 30 | 622.597 | 622.324 | 622.35 | 622.39 | 622.342 | 622.40 | 0.11 |
| 45 | 622.641 | 622.392 | | | | 622.52 | 0.18 |
| 60 | 622.679 | 622.566 | 622.879 | 622.665 | 622.661 | 622.69 | 0.11 |
| 300 | 623.032 | 622.648 | 622.949 | 622.819 | 622.628 | 622.82 | 0.18 |
| 1500 | 623.233 | 622.828 | 623.094 | 622.903 | 622.831 | 622.98 | 0.18 |
| 3600 | 623.17 | 623.014 | | 622.883 | 622.842 | 622.98 | 0.15 |
| 7200 | 623.215 | | 622.967 | 622.923 | 622.864 | 622.99 | 0.15 |
| 14400 | 623.198 | 622.936 | 623.084 | 622.952 | 622.865 | 623.01 | 0.13 |

**ONPF**

| Time | Replicate 1 | Replicate 2 | Replicate 3 | Replicate 4 | average | SD |
|---|---|---|---|---|---|---|
| 0 | 622.128 | 622.168 | 622.168 | 622.168 | 622.16 | 0.02 |
| 30 | 622.285 | 622.369 | 622.374 | 622.275 | 622.33 | 0.05 |
| 45 | 622.633 | | | | 622.63 | |
| 60 | 622.574 | 622.614 | 622.837 | 622.46 | 622.62 | 0.16 |
| 300 | 622.775 | 622.748 | 622.537 | 622.466 | 622.63 | 0.15 |
| 1500 | 623.018 | 622.862 | 623.026 | 622.769 | 622.92 | 0.13 |
| 3600 | 623.01 | 623.034 | 623.072 | 623.677 | 623.20 | 0.32 |
| 7200 | 622.909 | 622.897 | 622.958 | 622.649 | 622.85 | 0.14 |
| 14400 | 623.209 | 623.09 | 623.094 | 622.898 | 623.07 | 0.13 |

**APO**

| Time | Replicate 1 | Replicate 2 | Replicate 3 | Replicate 4 | Replicate 5 | average | SD |
|---|---|---|---|---|---|---|---|
| 0 | 622.116 | 622.116 | 622.146 | 622.168 | 622.168 | 622.14 | 0.03 |
| 30 | 622.283 | 622.314 | 622.462 | 622.332 | | 622.35 | 0.08 |
| 45 | 622.471 | 622.359 | | | | 622.42 | 0.08 |
| 60 | 622.464 | 622.573 | 622.674 | 622.593 | 622.574 | 622.58 | 0.07 |
| 300 | 622.792 | 622.605 | 622.871 | 622.737 | 622.583 | 622.72 | 0.12 |
| 1500 | 623.105 | 622.903 | 622.988 | 622.837 | 622.792 | 622.93 | 0.12 |
| 3600 | 623.211 | 622.931 | 623.089 | 622.878 | 622.764 | 622.97 | 0.18 |
| 7200 | 623.237 | 622.812 | | 623.027 | 622.808 | 622.97 | 0.20 |
| 14400 | 623.45 | 622.999 | | 623.039 | 622.971 | 623.11 | 0.23 |

**DNA**

| Time | Replicate 1 | Replicate 2 | Replicate 3 | Replicate 4 | Replicate 5 | average | SD |
|---|---|---|---|---|---|---|---|
| 0 | 622.115 | 622.105 | 622.125 | 622.168 | | 622.13 | 0.03 |
| 30 | 622.311 | 622.275 | 622.334 | 622.275 | | 622.30 | 0.03 |
| 45 | 622.611 | 622.28 | | | | 622.45 | 0.23 |
| 60 | 622.538 | 622.499 | 622.721 | 622.433 | | 622.55 | 0.12 |
| 300 | 622.675 | 622.564 | 622.71 | 622.463 | | 622.60 | 0.11 |
| 1500 | 622.935 | 622.912 | 623.005 | 622.339 | | 622.80 | 0.31 |
| 3600 | | 622.697 | 622.985 | | | 622.84 | 0.20 |
| 7200 | 622.952 | 622.74 | 622.893 | 622.383 | | 622.74 | 0.26 |
| 14400 | 623.222 | 622.427 | 623.061 | 622.323 | | 622.76 | 0.45 |

**ONPFDNA**

| Time | Replicate 1 | Replicate 2 | Replicate 3 | Replicate 4 | average | SD |
|---|---|---|---|---|---|---|
| 0 | 622.123 | 621.96 | 622.054 | 622.102 | 622.06 | 0.07 |
| 30 | 622.308 | 622.091 | 622.372 | 622.205 | 622.24 | 0.12 |
| 45 | 622.547 | 622.143 | | | 622.35 | 0.29 |
| 60 | 622.489 | 622.298 | 622.671 | 622.509 | 622.49 | 0.15 |
| 300 | 622.627 | 622.419 | 622.623 | 622.532 | 622.55 | 0.10 |
| 1500 | 622.615 | 622.585 | 622.821 | 622.698 | 622.68 | 0.11 |
| 3600 | 623.065 | | 622.891 | 622.775 | 622.91 | 0.15 |
| 7200 | 622.941 | | | 622.855 | 622.90 | 0.06 |
| 14400 | 622.74 | 622.625 | | 622.944 | 622.77 | 0.16 |

**TMG**

| Time | Replicate 1 | Replicate 2 | Replicate 3 | Replicate 4 | Replicate 5 | average | SD |
|---|---|---|---|---|---|---|---|
| 0 | 622.079 | 622.108 | | | | 622.09 | 0.02 |
| 30 | 622.309 | 622.589 | | | | 622.45 | 0.20 |
| 45 | 622.718 | 622.472 | | | | 622.60 | 0.17 |
| 60 | 622.67 | 622.923 | | | | 622.80 | 0.18 |
| 300 | | 622.945 | | | | 622.95 | |
| 1500 | 622.983 | 623.202 | | | | 623.09 | 0.15 |
| 3600 | | 623.201 | | | | 623.20 | |
| 7200 | 622.808 | 623.208 | | | | 623.01 | 0.28 |
| 14400 | 623.111 | 623.368 | | | | 623.24 | 0.18 |

### 219-223 DWSAM

**Charge** 1

**IPTG**

| Time | centroid | D |
|---|---|---|
| 0 | 609.56 | -0.03 |
| 30 | 610.01 | 0.63 |
| 45 | 610.09 | 0.75 |
| 60 | 610.17 | 0.87 |
| 300 | 610.60 | 1.49 |
| 1500 | 610.67 | 1.60 |
| 3600 | 610.79 | 1.77 |
| 7200 | 610.75 | 1.72 |
| 14400 | 610.75 | 1.71 |

**ONPF**

| Time | centroid | D |
|---|---|---|
| 0 | 609.56 | -0.02 |
| 30 | 609.71 | 0.20 |
| 45 | 609.88 | 0.44 |
| 60 | 610.01 | 0.63 |
| 300 | 610.24 | 0.97 |
| 1500 | 610.51 | 1.37 |
| 3600 | 610.55 | 1.42 |
| 7200 | 610.50 | 1.35 |
| 14400 | 610.88 | 1.90 |

**APO**

| Time | centroid | D |
|---|---|---|
| 0 | 609.57 | 0.00 |
| 30 | 609.82 | 0.35 |
| 45 | 609.91 | 0.49 |
| 60 | 610.04 | 0.68 |
| 300 | 610.37 | 1.16 |
| 1500 | 610.72 | 1.66 |
| 3600 | 610.78 | 1.75 |
| 7200 | 610.75 | 1.72 |
| 14400 | 610.94 | 1.99 |

**DNA**

| Time | centroid | D |
|---|---|---|
| 0 | 609.5455 | -0.04 |
| 30 | 609.7255 | 0.22 |
| 45 | 609.883 | 0.45 |
| 60 | 609.955 | 0.56 |
| 300 | 610.2135 | 0.93 |
| 1500 | 610.5075 | 1.36 |
| 3600 | 610.507 | 1.36 |
| 7200 | 610.458 | 1.29 |
| 14400 | 610.812 | 1.80 |

**ONPFDNA**

| Time | centroid | D |
|---|---|---|
| 0 | 609.55 | -0.04 |
| 30 | 609.68 | 0.15 |
| 45 | 609.76 | 0.27 |
| 60 | 609.92 | 0.50 |
| 300 | 610.08 | 0.74 |
| 1500 | 610.22 | 0.95 |
| 3600 | 610.60 | 1.50 |
| 7200 | 610.51 | 1.37 |
| 14400 | 610.23 | 0.95 |

**TMG**

| Time | centroid | D |
|---|---|---|
| 0 | 609.55 | -0.03 |
| 30 | 609.97 | 0.57 |
| 45 | 610.04 | 0.68 |
| 60 | 610.33 | 1.10 |
| 300 | 610.58 | 1.46 |
| 1500 | 610.71 | 1.66 |
| 3600 | 610.75 | 1.72 |
| 7200 | 610.74 | 1.70 |
| 14400 | 610.87 | 1.89 |

control 609.574
infinity 611.632

$$D(t) = \frac{Mt - M0}{M\infty - M0} *N$$

**IPTG**

| Time | Replicate 1 | Replicate 2 | Replicate 3 | Replicate 4 | Replicate5 | average | SD |
|---|---|---|---|---|---|---|---|
| 0 | 609.551 | 609.559 | | | | 609.56 | 0.01 |
| 30 | 610.16 | 609.858 | | | | 610.01 | 0.21 |
| 45 | 610.186 | 609.986 | | | | 610.09 | 0.14 |
| 60 | 610.216 | 610.127 | | | | 610.17 | 0.06 |
| 300 | 610.728 | 610.464 | | | | 610.60 | 0.19 |
| 1500 | 610.846 | 610.501 | | | | 610.67 | 0.24 |
| 3600 | 610.832 | 610.743 | | | | 610.79 | 0.06 |
| 7200 | 610.883 | 610.623 | | | | 610.75 | 0.18 |
| 14400 | 610.878 | 610.622 | | | | 610.75 | 0.18 |

**ONPF**

| Time | Replicate 1 | Replicate 2 | Replicate 3 | Replicate 4 | average | SD |
|---|---|---|---|---|---|---|
| 0 | 609.54 | 609.579 | | | 609.56 | 0.03 |
| 30 | 609.746 | 609.677 | | | 609.71 | 0.05 |
| 45 | 609.984 | 609.772 | | | 609.88 | 0.15 |
| 60 | 609.963 | 610.053 | | | 610.01 | 0.06 |
| 300 | 610.43 | 610.042 | | | 610.24 | 0.27 |
| 1500 | 610.698 | 610.33 | | | 610.51 | 0.26 |
| 3600 | 610.718 | 610.377 | | | 610.55 | 0.24 |
| 7200 | 610.607 | 610.393 | | | 610.50 | 0.15 |
| 14400 | 610.879 | | | | 610.88 | |

**APO**

| Time | Replicate 1 | Replicate 2 | Replicate 3 | Replicate 4 | Replicate 5 | average | SD |
|---|---|---|---|---|---|---|---|
| 0 | 609.567 | 609.581 | | | | 609.57 | 0.01 |
| 30 | 609.847 | 609.787 | | | | 609.82 | 0.04 |
| 45 | 609.979 | 609.842 | | | | 609.91 | 0.10 |
| 60 | 609.984 | 610.092 | | | | 610.04 | 0.08 |
| 300 | 610.546 | 610.199 | | | | 610.37 | 0.25 |
| 1500 | 610.828 | 610.602 | | | | 610.72 | 0.16 |
| 3600 | 610.949 | 610.605 | | | | 610.78 | 0.24 |
| 7200 | 610.991 | 610.513 | | | | 610.75 | 0.34 |
| 14400 | 611.171 | 610.703 | | | | 610.94 | 0.33 |

**DNA**

| Time | Replicate 1 | Replicate 2 | Replicate 3 | Replicate 4 | Replicate 5 | average | SD |
|---|---|---|---|---|---|---|---|
| 0 | 609.529 | 609.562 | | | | 609.55 | 0.02 |
| 30 | 609.751 | 609.7 | | | | 609.73 | 0.04 |
| 45 | 610.028 | 609.738 | | | | 609.88 | 0.21 |
| 60 | 609.905 | 610.005 | | | | 609.96 | 0.07 |
| 300 | 610.266 | 610.161 | | | | 610.21 | 0.07 |
| 1500 | 610.491 | 610.524 | | | | 610.51 | 0.02 |
| 3600 | 610.65 | 610.364 | | | | 610.51 | 0.20 |
| 7200 | 610.6 | 610.316 | | | | 610.46 | 0.20 |
| 14400 | 610.812 | | | | | 610.81 | |

**ONPFDNA**

| Time | Replicate 1 | Replicate 2 | Replicate 3 | Replicate 4 | average | SD |
|---|---|---|---|---|---|---|
| 0 | 609.541 | 609.551 | | | 609.55 | 0.01 |
| 30 | 609.695 | 609.661 | | | 609.68 | 0.02 |
| 45 | 609.812 | 609.713 | | | 609.76 | 0.07 |
| 60 | 609.915 | 609.917 | | | 609.92 | 0.00 |
| 300 | 610.168 | 609.998 | | | 610.08 | 0.12 |
| 1500 | 610.12 | 610.326 | | | 610.22 | 0.15 |
| 3600 | 610.66 | 610.54 | | | 610.60 | 0.08 |
| 7200 | 610.509 | 610.517 | | | 610.51 | 0.01 |
| 14400 | 610.162 | 610.289 | | | 610.23 | 0.09 |

**TMG**

| Time | Replicate 1 | Replicate 2 | Replicate 3 | Replicate 4 | Replicate 5 | average | SD |
|---|---|---|---|---|---|---|---|
| 0 | 609.524 | 609.578 | | | | 609.55 | 0.04 |
| 30 | 609.877 | 610.055 | | | | 609.97 | 0.13 |
| 45 | 610.123 | 609.961 | | | | 610.04 | 0.11 |
| 60 | 610.136 | 610.526 | | | | 610.33 | 0.28 |
| 300 | 610.523 | 610.634 | | | | 610.58 | 0.08 |
| 1500 | 610.564 | 610.861 | | | | 610.71 | 0.21 |
| 3600 | 610.626 | 610.879 | | | | 610.75 | 0.18 |
| 7200 | 610.536 | 610.95 | | | | 610.74 | 0.29 |
| 14400 | 610.744 | 610.992 | | | | 610.87 | 0.18 |

### 220-223 WSAM

**Charge** 1

**IPTG**

| Time | centroid | D |
|---|---|---|
| 0 | 494.48 | 0.00 |
| 30 | 494.90 | 0.56 |
| 45 | 494.98 | 0.67 |
| 60 | 495.06 | 0.77 |
| 300 | 495.47 | 1.31 |
| 1500 | 495.54 | 1.39 |
| 3600 | 495.66 | 1.55 |
| 7200 | 495.61 | 1.49 |
| 14400 | 495.59 | 1.46 |

**ONPF**

| Time | centroid | D |
|---|---|---|
| 0 | 494.52 | 0.05 |
| 30 | 494.63 | 0.20 |
| 45 | 494.82 | 0.45 |
| 60 | 494.93 | 0.59 |
| 300 | 495.12 | 0.84 |
| 1500 | 495.38 | 1.19 |
| 3600 | 495.47 | 1.30 |
| 7200 | 495.48 | 1.32 |
| 14400 | 495.52 | 1.37 |

**APO**

| Time | centroid | D |
|---|---|---|
| 0 | 494.48 | 0.00 |
| 30 | 494.72 | 0.32 |
| 45 | 494.82 | 0.45 |
| 60 | 494.95 | 0.62 |
| 300 | 495.27 | 1.05 |
| 1500 | 495.62 | 1.51 |
| 3600 | 495.63 | 1.51 |
| 7200 | 495.63 | 1.51 |
| 14400 | 495.60 | 1.48 |

**DNA**

| Time | centroid | D |
|---|---|---|
| 0 | 494.51 | 0.04 |
| 30 | 494.81 | 0.43 |
| 45 | 494.82 | 0.45 |
| 60 | 494.89 | 0.54 |
| 300 | 495.14 | 0.87 |
| 1500 | 495.49 | 1.33 |
| 3600 | 495.33 | 1.13 |
| 7200 | 495.35 | 1.14 |
| 14400 | 495.75 | 1.68 |

**ONPFDNA**

| Time | centroid | D |
|---|---|---|
| 0 | 494.47 | -0.01 |
| 30 | 494.58 | 0.14 |
| 45 | 494.67 | 0.26 |
| 60 | 494.87 | 0.52 |
| 300 | 494.94 | 0.61 |
| 1500 | 495.13 | 0.86 |
| 3600 | 495.43 | 1.25 |
| 7200 | 495.38 | 1.19 |
| 14400 | 495.25 | 1.01 |

**TMG**

| Time | centroid | D |
|---|---|---|
| 0 | 494.50 | 0.03 |
| 30 | 494.95 | 0.62 |
| 45 | 495.00 | 0.68 |
| 60 | 495.18 | 0.92 |
| 300 | 495.47 | 1.30 |
| 1500 | 495.60 | 1.48 |
| 3600 | 495.65 | 1.54 |
| 7200 | 495.68 | 1.58 |
| 14400 | 495.72 | 1.64 |

control 494.4775
infinity 495.999

$$D(t) = \frac{M_t - M_0}{M_\infty - M_0} \cdot N$$

**IPTG**

| Time | Replicate 1 | Replicate 2 | Replicate 3 | Replicate 4 | Replicate5 | average | SD |
|---|---|---|---|---|---|---|---|
| 0 | 494.508 | 494.448 | | | | 494.48 | 0.04 |
| 30 | 495.085 | 494.715 | | | | 494.90 | 0.26 |
| 45 | 495.108 | 494.859 | | | | 494.98 | 0.18 |
| 60 | 495.122 | 495.006 | | | | 495.06 | 0.08 |
| 300 | 495.607 | 495.338 | | | | 495.47 | 0.19 |
| 1500 | 495.732 | 495.34 | | | | 495.54 | 0.28 |
| 3600 | 495.72 | 495.595 | | | | 495.66 | 0.09 |
| 7200 | 495.76 | 495.461 | | | | 495.61 | 0.21 |
| 14400 | 495.712 | 495.459 | | | | 495.59 | 0.18 |

**ONPF**

| Time | Replicate 1 | Replicate 2 | Replicate 3 | Replicate 4 | average | SD |
|---|---|---|---|---|---|---|
| 0 | 494.578 | 494.452 | | | 494.52 | 0.09 |
| 30 | 494.711 | 494.553 | | | 494.63 | 0.11 |
| 45 | 495.003 | 494.63 | | | 494.82 | 0.26 |
| 60 | 494.925 | 494.928 | | | 494.93 | 0.00 |
| 300 | 495.324 | 494.911 | | | 495.12 | 0.29 |
| 1500 | 495.594 | 495.174 | | | 495.38 | 0.30 |
| 3600 | 495.644 | 495.294 | | | 495.47 | 0.25 |
| 7200 | 495.479 | | | | 495.48 | |
| 14400 | 495.638 | 495.397 | | | 495.52 | 0.17 |

**APO**

| Time | Replicate 1 | Replicate 2 | Replicate 3 | Replicate 4 | Replicate 5 | average | SD |
|---|---|---|---|---|---|---|---|
| 0 | 494.506 | 494.449 | | | | 494.48 | 0.04 |
| 30 | 494.784 | 494.659 | | | | 494.72 | 0.09 |
| 45 | 494.954 | 494.693 | | | | 494.82 | 0.18 |
| 60 | 494.934 | 494.964 | | | | 494.95 | 0.02 |
| 300 | 495.464 | 495.083 | | | | 495.27 | 0.27 |
| 1500 | 495.763 | 495.484 | | | | 495.62 | 0.20 |
| 3600 | 495.787 | 495.468 | | | | 495.63 | 0.23 |
| 7200 | 495.625 | | | | | 495.63 | |
| 14400 | 495.838 | 495.367 | | | | 495.60 | 0.33 |

**DNA**

| Time | Replicate 1 | Replicate 2 | Replicate 3 | Replicate 4 | Replicate 5 | average | SD |
|---|---|---|---|---|---|---|---|
| 0 | 494.573 | 494.442 | | | | 494.51 | 0.09 |
| 30 | 495.039 | 494.577 | | | | 494.81 | 0.33 |
| 45 | 495.03 | 494.608 | | | | 494.82 | 0.30 |
| 60 | 494.913 | 494.862 | | | | 494.89 | 0.04 |
| 300 | 495.261 | 495.019 | | | | 495.14 | 0.17 |
| 1500 | 495.487 | | | | | 495.49 | |
| 3600 | 495.55 | 495.117 | | | | 495.33 | 0.31 |
| 7200 | 495.55 | 495.14 | | | | 495.35 | 0.29 |
| 14400 | 495.754 | | | | | 495.75 | |

**ONPFDNA**

| Time | Replicate 1 | Replicate 2 | Replicate 3 | Replicate 4 | average | SD |
|---|---|---|---|---|---|---|
| 0 | 494.492 | 494.441 | | | 494.47 | 0.04 |
| 30 | 494.634 | 494.534 | | | 494.58 | 0.07 |
| 45 | 494.756 | 494.592 | | | 494.67 | 0.12 |
| 60 | 494.941 | 494.8 | | | 494.87 | 0.10 |
| 300 | 495.045 | 494.84 | | | 494.94 | 0.14 |
| 1500 | 495.088 | 495.173 | | | 495.13 | 0.06 |
| 3600 | 495.449 | 495.408 | | | 495.43 | 0.03 |
| 7200 | 495.404 | 495.365 | | | 495.38 | 0.03 |
| 14400 | 495.247 | | | | 495.25 | |

**TMG**

| Time | Replicate 1 | Replicate 2 | Replicate 3 | Replicate 4 | Replicate 5 | average | SD |
|---|---|---|---|---|---|---|---|
| 0 | 494.502 | 494.5 | | | | 494.50 | 0.00 |
| 30 | 494.946 | 494.953 | | | | 494.95 | 0.00 |
| 45 | 495.092 | 494.9 | | | | 495.00 | 0.14 |
| 60 | 495.06 | 495.297 | | | | 495.18 | 0.17 |
| 300 | 495.432 | 495.5 | | | | 495.47 | 0.05 |
| 1500 | 495.503 | 495.698 | | | | 495.60 | 0.14 |
| 3600 | 495.535 | 495.766 | | | | 495.65 | 0.16 |
| 7200 | | 495.678 | | | | 495.68 | |
| 14400 | 495.604 | 495.839 | | | | 495.72 | 0.17 |

### 221-227 SAMSGFQ

**Charge** 1

**IPTG**

| Time | centroid | D |
|---|---|---|
| 0 | 727.74 | -0.06 |
| 30 | 728.45 | 0.88 |
| 45 | 728.65 | 1.15 |
| 60 | 728.95 | 1.55 |
| 300 | 729.18 | 1.85 |
| 1500 | 729.55 | 2.35 |
| 3600 | 729.41 | 2.17 |
| 7200 | 729.47 | 2.24 |
| 14400 | 729.42 | 2.18 |

**ONPF**

| Time | centroid | D |
|---|---|---|
| 0 | 727.83 | 0.05 |
| 30 | 727.94 | 0.20 |
| 45 | #DIV/0! | #DIV/0! |
| 60 | 728.65 | 1.15 |
| 300 | 728.66 | 1.17 |
| 1500 | 729.22 | 1.91 |
| 3600 | 729.14 | 1.80 |
| 7200 | 729.04 | 1.66 |
| 14400 | 729.41 | 2.17 |

**APO**

| Time | centroid | D |
|---|---|---|
| 0 | 727.79 | -0.01 |
| 30 | 728.23 | 0.59 |
| 45 | 728.39 | 0.80 |
| 60 | 728.43 | 0.86 |
| 300 | 728.93 | 1.52 |
| 1500 | 729.32 | 2.04 |
| 3600 | 729.41 | 2.16 |
| 7200 | 729.36 | 2.09 |
| 14400 | 729.54 | 2.33 |

**DNA**

| Time | centroid | D |
|---|---|---|
| 0 | 727.80 | 0.01 |
| 30 | 728.10 | 0.41 |
| 45 | 728.38 | 0.78 |
| 60 | 728.70 | 1.21 |
| 300 | 728.60 | 1.08 |
| 1500 | 729.12 | 1.77 |
| 3600 | 729.21 | 1.90 |
| 7200 | 729.01 | 1.63 |
| 14400 | 729.30 | 2.01 |

**ONPFDNA**

| Time | centroid | D |
|---|---|---|
| 0 | 727.79 | 0.00 |
| 30 | 728.10 | 0.41 |
| 45 | 728.28 | 0.65 |
| 60 | 728.48 | 0.92 |
| 300 | 728.54 | 1.00 |
| 1500 | 728.85 | 1.41 |
| 3600 | 729.22 | 1.91 |
| 7200 | 729.20 | 1.88 |
| 14400 | 729.00 | 1.61 |

**TMG**

| Time | centroid | D |
|---|---|---|
| 0 | 727.85 | 0.09 |
| 30 | 728.49 | 0.94 |
| 45 | 728.73 | 1.26 |
| 60 | 729.13 | 1.79 |
| 300 | 729.21 | 1.89 |
| 1500 | 729.38 | 2.13 |
| 3600 | 729.43 | 2.19 |
| 7200 | 729.49 | 2.27 |
| 14400 | 729.51 | 2.29 |

control 727.79
infinity 731.537

$$D(t) = \frac{M_t - M_0}{M_\infty - M_0} * N$$

**IPTG**

| Time | Replicate 1 | Replicate 2 | Replicate 3 | Replicate 4 | Replicate5 | average | SD |
|---|---|---|---|---|---|---|---|
| 0 | 727.734 | 727.753 | | | | 727.74 | 0.01 |
| 30 | 728.715 | 728.188 | | | | 728.45 | 0.37 |
| 45 | 728.723 | 728.583 | | | | 728.65 | 0.10 |
| 60 | 728.824 | 729.083 | | | | 728.95 | 0.18 |
| 300 | 729.372 | 728.982 | | | | 729.18 | 0.28 |
| 1500 | 729.555 | 729.549 | | | | 729.55 | 0.00 |
| 3600 | 729.544 | 729.282 | | | | 729.41 | 0.19 |
| 7200 | 729.467 | | | | | 729.47 | |
| 14400 | 729.429 | 729.412 | | | | 729.42 | 0.01 |

**ONPF**

| Time | Replicate 1 | Replicate 2 | Replicate 3 | Replicate 4 | average | SD |
|---|---|---|---|---|---|---|
| 0 | 727.815 | 727.843 | | | 727.83 | 0.02 |
| 30 | 728.06 | 727.816 | | | 727.94 | 0.17 |
| 45 | | | | | | |
| 60 | 728.557 | 728.752 | | | 728.65 | 0.14 |
| 300 | 728.984 | 728.344 | | | 728.66 | 0.45 |
| 1500 | 729.332 | 729.107 | | | 729.22 | 0.16 |
| 3600 | 729.167 | 729.108 | | | 729.14 | 0.04 |
| 7200 | 729.152 | 728.919 | | | 729.04 | 0.16 |
| 14400 | 729.414 | | | | 729.41 | |

**APO**

| Time | Replicate 1 | Replicate 2 | Replicate 3 | Replicate 4 | Replicate 5 | average | SD |
|---|---|---|---|---|---|---|---|
| 0 | 727.759 | 727.813 | | | | 727.79 | 0.04 |
| 30 | 728.168 | 728.291 | | | | 728.23 | 0.09 |
| 45 | 728.473 | 728.313 | | | | 728.39 | 0.11 |
| 60 | 728.324 | 728.541 | | | | 728.43 | 0.15 |
| 300 | 729.091 | 728.765 | | | | 728.93 | 0.23 |
| 1500 | 729.362 | 729.272 | | | | 729.32 | 0.06 |
| 3600 | 729.667 | 729.151 | | | | 729.41 | 0.36 |
| 7200 | 729.54 | 729.172 | | | | 729.36 | 0.26 |
| 14400 | 729.916 | 729.162 | | | | 729.54 | 0.53 |

**DNA**

| Time | Replicate 1 | Replicate 2 | Replicate 3 | Replicate 4 | Replicate 5 | average | SD |
|---|---|---|---|---|---|---|---|
| 0 | 727.817 | 727.773 | | | | 727.80 | 0.03 |
| 30 | 728.055 | 728.146 | | | | 728.10 | 0.06 |
| 45 | 728.61 | 728.145 | | | | 728.38 | 0.33 |
| 60 | | 728.695 | | | | 728.70 | |
| 300 | 728.594 | 728.6 | | | | 728.60 | 0.00 |
| 1500 | 729.103 | 729.131 | | | | 729.12 | 0.02 |
| 3600 | 729.318 | 729.109 | | | | 729.21 | 0.15 |
| 7200 | 729.075 | 728.941 | | | | 729.01 | 0.09 |
| 14400 | 729.299 | | | | | 729.30 | |

**ONPFDNA**

| Time | Replicate 1 | Replicate 2 | Replicate 3 | Replicate 4 | average | SD |
|---|---|---|---|---|---|---|
| 0 | 727.846 | 727.729 | | | 727.79 | 0.08 |
| 30 | 728.079 | 728.118 | | | 728.10 | 0.03 |
| 45 | 728.355 | 728.204 | | | 728.28 | 0.11 |
| 60 | 728.555 | 728.401 | | | 728.48 | 0.11 |
| 300 | 728.658 | 728.415 | | | 728.54 | 0.17 |
| 1500 | 728.807 | 728.883 | | | 728.85 | 0.05 |
| 3600 | 729.24 | 729.196 | | | 729.22 | 0.03 |
| 7200 | 729.202 | 729.196 | | | 729.20 | 0.00 |
| 14400 | 728.934 | 729.065 | | | 729.00 | 0.09 |

**TMG**

| Time | Replicate 1 | Replicate 2 | Replicate 3 | Replicate 4 | Replicate 5 | average | SD |
|---|---|---|---|---|---|---|---|
| 0 | 727.884 | 727.825 | | | | 727.85 | 0.04 |
| 30 | 728.347 | 728.636 | | | | 728.49 | 0.20 |
| 45 | 728.936 | 728.528 | | | | 728.73 | 0.29 |
| 60 | 728.987 | 729.273 | | | | 729.13 | 0.20 |
| 300 | 729.236 | 729.176 | | | | 729.21 | 0.04 |
| 1500 | 729.296 | 729.47 | | | | 729.38 | 0.12 |
| 3600 | 729.367 | 729.499 | | | | 729.43 | 0.09 |
| 7200 | | 729.492 | | | | 729.49 | |
| 14400 | 729.436 | 729.579 | | | | 729.51 | 0.10 |

### 224-232 SGFQQTMQM

**Charge** 2

**IPTG**

| Time | centroid | D |
|---|---|---|
| 0 | 529.54 | 0.00 |
| 30 | 529.60 | 0.19 |
| 45 | 529.61 | 0.22 |
| 60 | 529.61 | 0.23 |
| 300 | 529.61 | 0.21 |
| 1500 | 529.61 | 0.22 |
| 3600 | 529.62 | 0.26 |
| 7200 | 529.65 | 0.34 |
| 14400 | 529.71 | 0.54 |

**ONPF**

| Time | centroid | D |
|---|---|---|
| 0 | 529.56 | 0.06 |
| 30 | 529.61 | 0.23 |
| 45 | 529.65 | 0.34 |
| 60 | 529.65 | 0.36 |
| 300 | 529.62 | 0.25 |
| 1500 | 529.64 | 0.30 |
| 3600 | 529.66 | 0.39 |
| 7200 | 529.66 | 0.38 |
| 14400 | 529.71 | 0.54 |

**APO**

| Time | centroid | D |
|---|---|---|
| 0 | 529.54 | -0.01 |
| 30 | 529.58 | 0.12 |
| 45 | 529.61 | 0.21 |
| 60 | 529.63 | 0.29 |
| 300 | 529.60 | 0.18 |
| 1500 | 529.62 | 0.24 |
| 3600 | 529.62 | 0.26 |
| 7200 | 529.63 | 0.29 |
| 14400 | 529.75 | 0.67 |

**DNA**

| Time | centroid | D |
|---|---|---|
| 0 | 529.55 | 0.04 |
| 30 | 529.60 | 0.19 |
| 45 | 529.58 | 0.13 |
| 60 | 529.60 | 0.19 |
| 300 | 529.61 | 0.21 |
| 1500 | 529.63 | 0.27 |
| 3600 | 529.61 | 0.22 |
| 7200 | 529.63 | 0.29 |
| 14400 | 529.68 | 0.44 |

**ONPFDNA**

| Time | centroid | D |
|---|---|---|
| 0 | 529.55 | 0.02 |
| 30 | 529.59 | 0.15 |
| 45 | 529.61 | 0.22 |
| 60 | 529.63 | 0.29 |
| 300 | 529.58 | 0.13 |
| 1500 | 529.61 | 0.23 |
| 3600 | 529.61 | 0.23 |
| 7200 | 529.64 | 0.33 |
| 14400 | 529.62 | 0.25 |

**TMG**

| Time | centroid | D |
|---|---|---|
| 0 | 529.55 | 0.04 |
| 30 | 529.63 | 0.27 |
| 45 | 529.64 | 0.31 |
| 60 | 529.65 | 0.35 |
| 300 | 529.66 | 0.37 |
| 1500 | 529.65 | 0.35 |
| 3600 | 529.67 | 0.42 |
| 7200 | 529.68 | 0.44 |
| 14400 | 529.75 | 0.65 |

control 529.54
infinity 531.777

$$D(t) = \frac{M_t - M_0}{M_\infty - M_0} * N$$

**IPTG**

| Time | Replicate 1 | Replicate 2 | Replicate 3 | Replicate 4 | Replicate5 | average | SD |
|---|---|---|---|---|---|---|---|
| 0 | 529.532 | 529.549 | | | | 529.54 | 0.01 |
| 30 | 529.632 | 529.572 | | | | 529.60 | 0.04 |
| 45 | 529.644 | 529.577 | | | | 529.61 | 0.05 |
| 60 | 529.631 | 529.598 | | | | 529.61 | 0.02 |
| 300 | 529.643 | 529.574 | | | | 529.61 | 0.05 |
| 1500 | 529.617 | 529.603 | | | | 529.61 | 0.01 |
| 3600 | 529.64 | 529.606 | | | | 529.62 | 0.02 |
| 7200 | 529.671 | 529.624 | | | | 529.65 | 0.03 |
| 14400 | 529.746 | 529.677 | | | | 529.71 | 0.05 |

**ONPF**

| Time | Replicate 1 | Replicate 2 | Replicate 3 | Replicate 4 | average | SD |
|---|---|---|---|---|---|---|
| 0 | 529.565 | 529.576 | 529.541 | | 529.56 | 0.02 |
| 30 | 529.632 | 529.591 | 529.616 | | 529.61 | 0.02 |
| 45 | 529.683 | 529.613 | | | 529.65 | 0.05 |
| 60 | 529.626 | 529.623 | 529.713 | | 529.65 | 0.05 |
| 300 | 529.614 | 529.609 | 529.632 | | 529.62 | 0.01 |
| 1500 | 529.629 | 529.619 | 529.658 | | 529.64 | 0.02 |
| 3600 | 529.687 | 529.629 | 529.675 | | 529.66 | 0.03 |
| 7200 | 529.666 | 529.61 | 529.706 | | 529.66 | 0.05 |
| 14400 | 529.685 | | 529.737 | | 529.71 | 0.04 |

**APO**

| Time | Replicate 1 | Replicate 2 | Replicate 3 | Replicate 4 | Replicate 5 | average | SD |
|---|---|---|---|---|---|---|---|
| 0 | 529.528 | 529.548 | | | | 529.54 | 0.01 |
| 30 | 529.57 | 529.588 | | | | 529.58 | 0.01 |
| 45 | 529.583 | 529.63 | | | | 529.61 | 0.03 |
| 60 | 529.583 | 529.682 | | | | 529.63 | 0.07 |
| 300 | 529.592 | 529.605 | | | | 529.60 | 0.01 |
| 1500 | 529.607 | 529.624 | | | | 529.62 | 0.01 |
| 3600 | 529.628 | 529.619 | | | | 529.62 | 0.01 |
| 7200 | 529.642 | 529.625 | | | | 529.63 | 0.01 |
| 14400 | 529.794 | 529.712 | | | | 529.75 | 0.06 |

**DNA**

| Time | Replicate 1 | Replicate 2 | Replicate 3 | Replicate 4 | Replicate 5 | average | SD |
|---|---|---|---|---|---|---|---|
| 0 | 529.541 | 529.564 | | | | 529.55 | 0.02 |
| 30 | 529.622 | 529.582 | | | | 529.60 | 0.03 |
| 45 | 529.58 | 529.585 | | | | 529.58 | 0.00 |
| 60 | 529.6 | 529.6 | | | | 529.60 | 0.00 |
| 300 | 529.633 | 529.582 | | | | 529.61 | 0.04 |
| 1500 | 529.606 | 529.647 | | | | 529.63 | 0.03 |
| 3600 | 529.591 | 529.631 | | | | 529.61 | 0.03 |
| 7200 | 529.656 | 529.612 | | | | 529.63 | 0.03 |
| 14400 | 529.737 | 529.625 | | | | 529.68 | 0.08 |

**ONPFDNA**

| Time | Replicate 1 | Replicate 2 | Replicate 3 | Replicate 4 | average | SD |
|---|---|---|---|---|---|---|
| 0 | 529.564 | 529.531 | | | 529.55 | 0.02 |
| 30 | 529.587 | 529.586 | | | 529.59 | 0.00 |
| 45 | 529.639 | 529.581 | | | 529.61 | 0.04 |
| 60 | 529.634 | | | | 529.63 | |
| 300 | 529.592 | 529.57 | | | 529.58 | 0.02 |
| 1500 | 529.591 | 529.638 | | | 529.61 | 0.03 |
| 3600 | 529.623 | 529.606 | | | 529.61 | 0.01 |
| 7200 | 529.67 | 529.619 | | | 529.64 | 0.04 |
| 14400 | 529.597 | 529.64 | | | 529.62 | 0.03 |

**TMG**

| Time | Replicate 1 | Replicate 2 | Replicate 3 | Replicate 4 | Replicate 5 | average | SD |
|---|---|---|---|---|---|---|---|
| 0 | 529.569 | 529.537 | | | | 529.55 | 0.02 |
| 30 | 529.617 | 529.633 | | | | 529.63 | 0.01 |
| 45 | 529.674 | 529.607 | | | | 529.64 | 0.05 |
| 60 | 529.652 | | | | | 529.65 | |
| 300 | 529.67 | 529.644 | | | | 529.66 | 0.02 |
| 1500 | 529.627 | 529.679 | | | | 529.65 | 0.04 |
| 3600 | 529.686 | 529.661 | | | | 529.67 | 0.02 |
| 7200 | 529.691 | 529.669 | | | | 529.68 | 0.02 |
| 14400 | 529.749 | 529.748 | | | | 529.75 | 0.00 |

### 228-233 QTMQML

**Charge** 1

**IPTG**

| Time | centroid | D |
|---|---|---|
| 0 | 751.85 | -0.01 |
| 30 | 751.90 | 0.05 |
| 45 | 751.91 | 0.06 |
| 60 | 751.92 | 0.07 |
| 300 | 751.90 | 0.06 |
| 1500 | 751.93 | 0.09 |
| 3600 | 751.95 | 0.11 |
| 7200 | 752.01 | 0.19 |
| 14400 | 752.09 | 0.29 |

**ONPF**

| Time | centroid | D |
|---|---|---|
| 0 | 751.86 | 0.00 |
| 30 | 751.88 | 0.02 |
| 45 | 751.94 | 0.10 |
| 60 | 751.92 | 0.07 |
| 300 | 751.91 | 0.07 |
| 1500 | 751.92 | 0.07 |
| 3600 | 751.94 | 0.11 |
| 7200 | 751.97 | 0.14 |
| 14400 | 752.05 | 0.24 |

**APO**

| Time | centroid | D |
|---|---|---|
| 0 | 751.86 | -0.01 |
| 30 | 751.90 | 0.05 |
| 45 | 751.89 | 0.04 |
| 60 | 751.91 | 0.06 |
| 300 | 751.91 | 0.06 |
| 1500 | 751.96 | 0.13 |
| 3600 | 751.95 | 0.11 |
| 7200 | 752.02 | 0.20 |
| 14400 | 752.15 | 0.37 |

**DNA**

| Time | centroid | D |
|---|---|---|
| 0 | 751.83 | -0.04 |
| 30 | 751.89 | 0.04 |
| 45 | 751.91 | 0.06 |
| 60 | 751.90 | 0.06 |
| 300 | 751.91 | 0.06 |
| 1500 | 751.93 | 0.08 |
| 3600 | 751.91 | 0.06 |
| 7200 | 751.96 | 0.12 |
| 14400 | 751.94 | 0.11 |

**ONPFDNA**

| Time | centroid | D |
|---|---|---|
| 0 | 751.84 | -0.03 |
| 30 | 751.89 | 0.03 |
| 45 | 751.89 | 0.04 |
| 60 | 751.92 | 0.08 |
| 300 | 751.90 | 0.05 |
| 1500 | 751.93 | 0.09 |
| 3600 | 751.93 | 0.09 |
| 7200 | 751.95 | 0.11 |
| 14400 | 751.95 | 0.11 |

**TMG**

| Time | centroid | D |
|---|---|---|
| 0 | 751.85 | -0.02 |
| 30 | 751.90 | 0.06 |
| 45 | 751.94 | 0.10 |
| 60 | 751.91 | 0.06 |
| 300 | 751.92 | 0.08 |
| 1500 | 751.95 | 0.11 |
| 3600 | 751.95 | 0.12 |
| 7200 | 752.00 | 0.18 |
| 14400 | 752.08 | 0.28 |

control 751.86
infinity 755.01

$$D(t) = \frac{M_t - M_0}{M_\infty - M_0} * N$$

**IPTG**

| Time | Replicate 1 | Replicate 2 | Replicate 3 | Replicate 4 | Replicate5 | average | SD |
|---|---|---|---|---|---|---|---|
| 0 | 751.854 | 751.854 | | | | 751.85 | 0.00 |
| 30 | 751.913 | 751.887 | | | | 751.90 | 0.02 |
| 45 | 751.916 | 751.897 | | | | 751.91 | 0.01 |
| 60 | 751.93 | 751.901 | | | | 751.92 | 0.02 |
| 300 | 751.914 | 751.895 | | | | 751.90 | 0.01 |
| 1500 | 751.928 | 751.94 | | | | 751.93 | 0.01 |
| 3600 | 751.944 | 751.949 | | | | 751.95 | 0.00 |
| 7200 | 752.027 | 751.995 | | | | 752.01 | 0.02 |
| 14400 | 752.134 | 752.045 | | | | 752.09 | 0.06 |

**ONPF**

| Time | Replicate 1 | Replicate 2 | Replicate 3 | Replicate 4 | average | SD |
|---|---|---|---|---|---|---|
| 0 | 751.845 | 751.875 | | | 751.86 | 0.02 |
| 30 | 751.857 | 751.899 | | | 751.88 | 0.03 |
| 45 | 751.951 | 751.921 | | | 751.94 | 0.02 |
| 60 | 751.904 | 751.931 | | | 751.92 | 0.02 |
| 300 | 751.916 | 751.911 | | | 751.91 | 0.00 |
| 1500 | 751.911 | 751.925 | | | 751.92 | 0.01 |
| 3600 | 751.952 | 751.935 | | | 751.94 | 0.01 |
| 7200 | 751.978 | 751.957 | | | 751.97 | 0.01 |
| 14400 | 752.047 | | | | 752.05 | 0.00 |

**APO**

| Time | Replicate 1 | Replicate 2 | Replicate 3 | Replicate 4 | Replicate 5 | average | SD |
|---|---|---|---|---|---|---|---|
| 0 | 751.849 | 751.863 | | | | 751.86 | 0.01 |
| 30 | 751.892 | 751.904 | | | | 751.90 | 0.01 |
| 45 | 751.881 | 751.9 | | | | 751.89 | 0.01 |
| 60 | 751.883 | 751.931 | | | | 751.91 | 0.03 |
| 300 | 751.897 | 751.921 | | | | 751.91 | 0.02 |
| 1500 | 751.953 | 751.973 | | | | 751.96 | 0.01 |
| 3600 | 751.934 | 751.957 | | | | 751.95 | 0.02 |
| 7200 | 752.05 | 751.988 | | | | 752.02 | 0.04 |
| 14400 | 752.217 | 752.083 | | | | 752.15 | 0.09 |

**DNA**

| Time | Replicate 1 | Replicate 2 | Replicate 3 | Replicate 4 | Replicate 5 | average | SD |
|---|---|---|---|---|---|---|---|
| 0 | 751.806 | 751.856 | | | | 751.83 | 0.04 |
| 30 | 751.887 | 751.898 | | | | 751.89 | 0.01 |
| 45 | | 751.909 | | | | 751.91 | |
| 60 | 751.888 | 751.919 | | | | 751.90 | 0.02 |
| 300 | 751.904 | 751.911 | | | | 751.91 | 0.00 |
| 1500 | 751.918 | 751.932 | | | | 751.93 | 0.01 |
| 3600 | 751.904 | 751.917 | | | | 751.91 | 0.01 |
| 7200 | 751.954 | 751.956 | | | | 751.96 | 0.00 |
| 14400 | 751.973 | 751.916 | | | | 751.94 | 0.04 |

**ONPFDNA**

| Time | Replicate 1 | Replicate 2 | Replicate 3 | Replicate 4 | average | SD |
|---|---|---|---|---|---|---|
| 0 | 751.819 | 751.86 | | | 751.84 | 0.03 |
| 30 | 751.88 | 751.89 | | | 751.89 | 0.01 |
| 45 | 751.88 | 751.901 | | | 751.89 | 0.01 |
| 60 | 751.934 | 751.911 | | | 751.92 | 0.02 |
| 300 | 751.907 | 751.898 | | | 751.90 | 0.01 |
| 1500 | 751.905 | 751.951 | | | 751.93 | 0.03 |
| 3600 | 751.937 | 751.925 | | | 751.93 | 0.01 |
| 7200 | 751.945 | 751.954 | | | 751.95 | 0.01 |
| 14400 | 751.913 | 751.987 | | | 751.95 | 0.05 |

**TMG**

| Time | Replicate 1 | Replicate 2 | Replicate 3 | Replicate 4 | Replicate 5 | average | SD |
|---|---|---|---|---|---|---|---|
| 0 | 751.836 | 751.856 | | | | 751.85 | 0.01 |
| 30 | 751.879 | 751.929 | | | | 751.90 | 0.04 |
| 45 | 751.964 | 751.908 | | | | 751.94 | 0.04 |
| 60 | 751.905 | | | | | 751.91 | |
| 300 | 751.927 | 751.917 | | | | 751.92 | 0.01 |
| 1500 | 751.933 | 751.968 | | | | 751.95 | 0.02 |
| 3600 | 751.963 | 751.943 | | | | 751.95 | 0.01 |
| 7200 | 752.017 | 751.982 | | | | 752.00 | 0.02 |
| 14400 | 752.089 | 752.072 | | | | 752.08 | 0.01 |

### 234-243 NEGIVPTAML

**Charge** 1

**IPTG**

| Time | centroid | D |
|---|---|---|
| 0 | 1045.24 | 0.03 |
| 30 | 1045.75 | 0.67 |
| 45 | 1045.89 | 0.84 |
| 60 | 1046.00 | 0.98 |
| 300 | 1046.22 | 1.26 |
| 1500 | 1046.82 | 2.01 |
| 3600 | 1047.17 | 2.45 |
| 7200 | 1047.22 | 2.52 |
| 14400 | 1047.20 | 2.49 |

**ONPF**

| Time | centroid | D |
|---|---|---|
| 0 | 1045.35 | 0.16 |
| 30 | 1045.56 | 0.43 |
| 45 | 1045.94 | 0.90 |
| 60 | 1046.12 | 1.13 |
| 300 | 1046.08 | 1.08 |
| 1500 | 1046.57 | 1.71 |
| 3600 | 1046.76 | 1.94 |
| 7200 | 1046.74 | 1.92 |
| 14400 | 1047.20 | 2.49 |

**APO**

| Time | centroid | D |
|---|---|---|
| 0 | 1045.22 | 0.00 |
| 30 | 1045.64 | 0.53 |
| 45 | 1045.80 | 0.73 |
| 60 | 1046.03 | 1.03 |
| 300 | 1046.15 | 1.17 |
| 1500 | 1046.87 | 2.08 |
| 3600 | 1047.07 | 2.33 |
| 7200 | 1047.08 | 2.34 |
| 14400 | 1047.47 | 2.83 |

**DNA**

| Time | centroid | D |
|---|---|---|
| 0 | 1045.27 | 0.07 |
| 30 | 1045.62 | 0.51 |
| 45 | 1046.12 | 1.13 |
| 60 | 1046.19 | 1.23 |
| 300 | 1046.21 | 1.25 |
| 1500 | 1046.66 | 1.82 |
| 3600 | 1046.81 | 2.01 |
| 7200 | 1046.71 | 1.88 |
| 14400 | 1046.68 | 1.84 |

**ONPFDNA**

| Time | centroid | D |
|---|---|---|
| 0 | 1045.23 | 0.01 |
| 30 | 1045.59 | 0.47 |
| 45 | 1045.87 | 0.81 |
| 60 | 1046.11 | 1.12 |
| 300 | 1046.09 | 1.09 |
| 1500 | 1046.41 | 1.50 |
| 3600 | 1046.96 | 2.19 |
| 7200 | 1046.85 | 2.06 |
| 14400 | 1046.62 | 1.76 |

**TMG**

| Time | centroid | D |
|---|---|---|
| 0 | 1045.25 | 0.04 |
| 30 | 1045.74 | 0.66 |
| 45 | 1046.12 | 1.13 |
| 60 | 1046.17 | 1.20 |
| 300 | 1046.30 | 1.36 |
| 1500 | 1046.75 | 1.93 |
| 3600 | 1047.18 | 2.47 |
| 7200 | 1047.12 | 2.40 |
| 14400 | 1047.43 | 2.79 |

control 1045.22
infinity 1050.776

$$D(t) = \frac{Mt - M0}{M\infty - M0} *N$$

**IPTG**

| Time | Replicate 1 | Replicate 2 | Replicate 3 | Replicate 4 | Replicate5 | average | SD |
|---|---|---|---|---|---|---|---|
| 0 | 1045.292 | 1045.197 | | | | 1045.24 | 0.07 |
| 30 | 1045.906 | 1045.601 | | | | 1045.75 | 0.22 |
| 45 | 1046.001 | 1045.78 | | | | 1045.89 | 0.16 |
| 60 | 1045.992 | 1046.004 | | | | 1046.00 | 0.01 |
| 300 | 1046.334 | 1046.113 | | | | 1046.22 | 0.16 |
| 1500 | 1046.987 | 1046.651 | | | | 1046.82 | 0.24 |
| 3600 | 1047.28 | 1047.05 | | | | 1047.17 | 0.16 |
| 7200 | 1047.384 | 1047.055 | | | | 1047.22 | 0.23 |
| 14400 | 1047.442 | 1046.949 | | | | 1047.20 | 0.35 |

**ONPF**

| Time | Replicate 1 | Replicate 2 | Replicate 3 | Replicate 4 | average | SD |
|---|---|---|---|---|---|---|
| 0 | 1045.361 | 1045.329 | | | 1045.35 | 0.02 |
| 30 | 1045.647 | 1045.474 | | | 1045.56 | 0.12 |
| 45 | 1046.29 | 1045.582 | | | 1045.94 | 0.50 |
| 60 | 1046.105 | 1046.132 | | | 1046.12 | 0.02 |
| 300 | 1046.248 | 1045.903 | | | 1046.08 | 0.24 |
| 1500 | 1046.74 | 1046.408 | | | 1046.57 | 0.23 |
| 3600 | 1047.053 | 1046.474 | | | 1046.76 | 0.41 |
| 7200 | 1046.862 | 1046.627 | | | 1046.74 | 0.17 |
| 14400 | 1047.305 | 1047.091 | | | 1047.20 | 0.15 |

**APO**

| Time | Replicate 1 | Replicate 2 | Replicate 3 | Replicate 4 | Replicate 5 | average | SD |
|---|---|---|---|---|---|---|---|
| 0 | 1045.198 | 1045.235 | | | | 1045.22 | 0.03 |
| 30 | 1045.678 | 1045.608 | | | | 1045.64 | 0.05 |
| 45 | 1045.893 | 1045.706 | | | | 1045.80 | 0.13 |
| 60 | 1045.904 | 1046.165 | | | | 1046.03 | 0.18 |
| 300 | 1046.268 | 1046.036 | | | | 1046.15 | 0.16 |
| 1500 | 1046.878 | 1046.858 | | | | 1046.87 | 0.01 |
| 3600 | 1047.244 | 1046.896 | | | | 1047.07 | 0.25 |
| 7200 | 1047.334 | 1046.821 | | | | 1047.08 | 0.36 |
| 14400 | 1047.99 | 1046.947 | | | | 1047.47 | 0.74 |

**DNA**

| Time | Replicate 1 | Replicate 2 | Replicate 3 | Replicate 4 | Replicate 5 | average | SD |
|---|---|---|---|---|---|---|---|
| 0 | 1045.29 | 1045.256 | | | | 1045.27 | 0.02 |
| 30 | 1045.65 | 1045.599 | | | | 1045.62 | 0.04 |
| 45 | 1046.551 | 1045.684 | | | | 1046.12 | 0.61 |
| 60 | 1046.211 | 1046.177 | | | | 1046.19 | 0.02 |
| 300 | 1046.328 | 1046.092 | | | | 1046.21 | 0.17 |
| 1500 | 1046.665 | 1046.657 | | | | 1046.66 | 0.01 |
| 3600 | 1047.165 | 1046.463 | | | | 1046.81 | 0.50 |
| 7200 | 1046.929 | 1046.493 | | | | 1046.71 | 0.31 |
| 14400 | 1047.272 | 1046.089 | | | | 1046.68 | 0.84 |

**ONPFDNA**

| Time | Replicate 1 | Replicate 2 | Replicate 3 | Replicate 4 | average | SD |
|---|---|---|---|---|---|---|
| 0 | 1045.27 | 1045.191 | | | 1045.23 | 0.06 |
| 30 | 1045.672 | 1045.513 | | | 1045.59 | 0.11 |
| 45 | 1046.056 | 1045.674 | | | 1045.87 | 0.27 |
| 60 | 1046.116 | 1046.108 | | | 1046.11 | 0.01 |
| 300 | 1046.191 | 1045.984 | | | 1046.09 | 0.15 |
| 1500 | 1046.303 | 1046.526 | | | 1046.41 | 0.16 |
| 3600 | 1047.13 | 1046.793 | | | 1046.96 | 0.24 |
| 7200 | 1046.881 | 1046.822 | | | 1046.85 | 0.04 |
| 14400 | 1046.832 | 1046.398 | | | 1046.62 | 0.31 |

**TMG**

| Time | Replicate 1 | Replicate 2 | Replicate 3 | Replicate 4 | Replicate 5 | average | SD |
|---|---|---|---|---|---|---|---|
| 0 | 1045.284 | 1045.225 | | | | 1045.25 | 0.04 |
| 30 | 1045.625 | 1045.862 | | | | 1045.74 | 0.17 |
| 45 | 1046.387 | 1045.844 | | | | 1046.12 | 0.38 |
| 60 | 1046.153 | 1046.195 | | | | 1046.17 | 0.03 |
| 300 | 1046.3 | 1046.297 | | | | 1046.30 | 0.00 |
| 1500 | 1046.638 | 1046.865 | | | | 1046.75 | 0.16 |
| 3600 | 1047.111 | 1047.243 | | | | 1047.18 | 0.09 |
| 7200 | 1046.887 | 1047.362 | | | | 1047.12 | 0.34 |
| 14400 | 1047.281 | 1047.586 | | | | 1047.43 | 0.22 |

### 243-254 LVANDQMALGAM

**Charge** 2

**IPTG**

| Time | centroid | D |
|---|---|---|
| 0 | 617.6978 | -0.01 |
| 30 | 617.928 | 1.00 |
| 45 | 617.828 | 0.56 |
| 60 | 618.0675 | 1.61 |
| 300 | 617.9755 | 1.21 |
| 1500 | 618.0183 | 1.40 |
| 3600 | 618.0295 | 1.44 |
| 7200 | 618.1115 | 1.80 |
| 14400 | 618.1148 | 1.82 |

**ONPF**

| Time | centroid | D |
|---|---|---|
| 0 | 617.727 | 0.12 |
| 30 | 617.973 | 1.20 |
| 45 | 617.879 | 0.78 |
| 60 | 618.1175 | 1.83 |
| 300 | 618.0243 | 1.42 |
| 1500 | 618.0948 | 1.73 |
| 3600 | 618.1773 | 2.09 |
| 7200 | 618.1995 | 2.19 |
| 14400 | 618.26 | 2.45 |

**APO**

| Time | centroid | D |
|---|---|---|
| 0 | 617.7045 | 0.02 |
| 30 | 617.9255 | 0.99 |
| 45 | 617.882 | 0.80 |
| 60 | 618.037 | 1.48 |
| 300 | 617.9868 | 1.26 |
| 1500 | 618.084 | 1.68 |
| 3600 | 618.1265 | 1.87 |
| 7200 | 618.2123 | 2.25 |
| 14400 | 618.2335 | 2.34 |

**DNA**

| Time | centroid | D |
|---|---|---|
| 0 | 617.7147 | 0.06 |
| 30 | 617.9227 | 0.98 |
| 45 | 618.06 | 1.58 |
| 60 | 618.1287 | 1.88 |
| 300 | 618.018 | 1.39 |
| 1500 | 618.129 | 1.88 |
| 3600 | 618.1537 | 1.99 |
| 7200 | 618.2137 | 2.25 |
| 14400 | 618.2507 | 2.41 |

**ONPFDNA**

| Time | centroid | D |
|---|---|---|
| 0 | 617.80 | 0.44 |
| 30 | 618.21 | 2.22 |
| 45 | 617.95 | 1.11 |
| 60 | 618.43 | 3.22 |
| 300 | 618.34 | 2.79 |
| 1500 | 618.47 | 3.38 |
| 3600 | 618.51 | 3.57 |
| 7200 | 618.43 | 3.22 |
| 14400 | 618.55 | 3.73 |

**TMG**

| Time | centroid | D |
|---|---|---|
| 0 | 617.682 | -0.08 |
| 30 | 617.823 | 0.54 |
| 45 | 617.8655 | 0.73 |
| 60 | 617.819 | 0.52 |
| 300 | 617.8135 | 0.50 |
| 1500 | 617.8 | 0.44 |
| 3600 | 617.836 | 0.60 |
| 7200 | 617.896 | 0.86 |
| 14400 | 617.8235 | 0.54 |

control 617.7
infinity 619.981

$$D(t) = \frac{M_t - M_0}{M_\infty - M_0} * N$$

**IPTG**

| Time | Replicate 1 | Replicate 2 | Replicate 3 | Replicate 4 | Replicate5 | average | SD |
|---|---|---|---|---|---|---|---|
| 0 | 617.687 | 617.675 | 617.712 | 617.717 | | 617.70 | 0.02 |
| 30 | 617.75 | 618.096 | 617.936 | 617.93 | | 617.93 | 0.14 |
| 45 | 617.828 | | | | | 617.83 | |
| 60 | 617.877 | 618.292 | 618.066 | 618.035 | | 618.07 | 0.17 |
| 300 | 617.726 | 618.19 | 618.006 | 617.98 | | 617.98 | 0.19 |
| 1500 | 617.811 | 618.21 | 618.045 | 618.007 | | 618.02 | 0.16 |
| 3600 | 617.867 | 618.194 | 618.036 | 618.021 | | 618.03 | 0.13 |
| 7200 | 617.923 | 618.306 | 618.126 | 618.091 | | 618.11 | 0.16 |
| 14400 | 617.946 | 618.287 | 618.14 | 618.086 | | 618.11 | 0.14 |

**ONPF**

| Time | Replicate 1 | Replicate 2 | Replicate 3 | Replicate 4 | average | SD |
|---|---|---|---|---|---|---|
| 0 | 617.752 | 617.715 | 617.727 | 617.714 | 617.73 | 0.02 |
| 30 | 617.959 | 618.064 | 617.891 | 617.978 | 617.97 | 0.07 |
| 45 | 617.879 | | | | 617.88 | |
| 60 | 617.865 | 618.33 | 618.087 | 618.188 | 618.12 | 0.20 |
| 300 | 617.914 | 618.181 | 617.954 | 618.048 | 618.02 | 0.12 |
| 1500 | 617.759 | 618.362 | 618.107 | 618.151 | 618.09 | 0.25 |
| 3600 | 618.004 | 618.382 | 618.092 | 618.231 | 618.18 | 0.17 |
| 7200 | 617.99 | 618.381 | 618.184 | 618.243 | 618.20 | 0.16 |
| 14400 | 618.074 | 618.424 | 618.241 | 618.293 | 618.26 | 0.14 |

**APO**

| Time | Replicate 1 | Replicate 2 | Replicate 3 | Replicate 4 | Replicate 5 | average | SD |
|---|---|---|---|---|---|---|---|
| 0 | 617.61 | 617.782 | 617.711 | 617.715 | | 617.70 | 0.07 |
| 30 | 617.743 | 618.146 | 617.893 | 617.92 | | 617.93 | 0.17 |
| 45 | 617.882 | | | | | 617.88 | 0.00 |
| 60 | 617.716 | 618.21 | 618.125 | 618.097 | | 618.04 | 0.22 |
| 300 | 617.718 | 618.222 | 618.013 | 617.994 | | 617.99 | 0.21 |
| 1500 | 617.756 | 618.293 | 618.149 | 618.138 | | 618.08 | 0.23 |
| 3600 | 617.856 | 618.362 | 618.151 | 618.137 | | 618.13 | 0.21 |
| 7200 | 617.945 | 618.432 | 618.265 | 618.207 | | 618.21 | 0.20 |
| 14400 | 617.96 | 618.464 | 618.256 | 618.254 | | 618.23 | 0.21 |

**DNA**

| Time | Replicate 1 | Replicate 2 | Replicate 3 | Replicate 4 | Replicate 5 | average | SD |
|---|---|---|---|---|---|---|---|
| 0 | 617.707 | 617.722 | 617.715 | | | 617.71 | 0.01 |
| 30 | 617.854 | 618.039 | 617.875 | | | 617.92 | 0.10 |
| 45 | 618.06 | | | | | 618.06 | |
| 60 | 617.997 | 618.325 | 618.064 | | | 618.13 | 0.17 |
| 300 | 617.95 | 618.165 | 617.939 | | | 618.02 | 0.13 |
| 1500 | 618.01 | 618.288 | 618.089 | | | 618.13 | 0.14 |
| 3600 | 618.088 | 618.284 | 618.089 | | | 618.15 | 0.11 |
| 7200 | 618.104 | 618.337 | 618.2 | | | 618.21 | 0.12 |
| 14400 | 618.166 | 618.385 | 618.201 | | | 618.25 | 0.12 |

**ONPFDNA**

| Time | Replicate 1 | Replicate 2 | Replicate 3 | Replicate 4 | average | SD |
|---|---|---|---|---|---|---|
| 0 | 617.727 | 617.963 | 617.714 | | 617.80 | 0.14 |
| 30 | 617.897 | 618.849 | 617.873 | | 618.21 | 0.56 |
| 45 | 617.954 | | | | 617.95 | |
| 60 | 617.948 | 619.345 | 618.01 | | 618.43 | 0.79 |
| 300 | 617.866 | 619.131 | 618.014 | | 618.34 | 0.69 |
| 1500 | 617.926 | 619.477 | 618.01 | | 618.47 | 0.87 |
| 3600 | 618.097 | 619.433 | 618.012 | | 618.51 | 0.80 |
| 7200 | 618.103 | 619.106 | 618.095 | | 618.43 | 0.58 |
| 14400 | 617.964 | 619.596 | 618.093 | | 618.55 | 0.91 |

**TMG**

| Time | Replicate 1 | Replicate 2 | Replicate 3 | Replicate 4 | Replicate 5 | average | SD |
|---|---|---|---|---|---|---|---|
| 0 | 617.682 | 617.682 | | | | 617.68 | 0.00 |
| 30 | 617.742 | 617.904 | | | | 617.82 | 0.11 |
| 45 | 617.847 | 617.884 | | | | 617.87 | 0.03 |
| 60 | 617.819 | | | | | 617.82 | |
| 300 | 617.75 | 617.877 | | | | 617.81 | 0.09 |
| 1500 | 617.688 | 617.912 | | | | 617.80 | 0.16 |
| 3600 | 617.781 | 617.891 | | | | 617.84 | 0.08 |
| 7200 | 617.855 | 617.937 | | | | 617.90 | 0.06 |
| 14400 | 617.81 | 617.837 | | | | 617.82 | 0.02 |

### 244-255 VANDQMALGAMR

**Charge** 2

**IPTG**

| Time | centroid | D |
|---|---|---|
| 0 | 639.2315 | -0.01 |
| 30 | 639.488 | 0.73 |
| 45 | 639.5035 | 0.77 |
| 60 | 639.521 | 0.82 |
| 300 | 639.5005 | 0.76 |
| 1500 | 639.513 | 0.80 |
| 3600 | 639.6055 | 1.07 |
| 7200 | 639.562 | 0.94 |
| 14400 | 639.5935 | 1.03 |

**ONPF**

| Time | centroid | D |
|---|---|---|
| 0 | 639.2575 | 0.07 |
| 30 | 639.4205 | 0.54 |
| 45 | 639.507 | 0.78 |
| 60 | 639.518 | 0.81 |
| 300 | 639.502 | 0.77 |
| 1500 | 639.5615 | 0.94 |
| 3600 | 639.602 | 1.06 |
| 7200 | 639.614 | 1.09 |
| 14400 | 639.62 | 1.10 |

**APO**

| Time | centroid | D |
|---|---|---|
| 0 | 639.2295 | -0.01 |
| 30 | 639.4315 | 0.57 |
| 45 | 639.4505 | 0.62 |
| 60 | 639.4895 | 0.73 |
| 300 | 639.4635 | 0.66 |
| 1500 | 639.59 | 1.02 |
| 3600 | 639.6215 | 1.11 |
| 7200 | 639.6595 | 1.22 |
| 14400 | 639.8585 | 1.79 |

**DNA**

| Time | centroid | D |
|---|---|---|
| 0 | 639.2305 | -0.01 |
| 30 | 639.4375 | 0.58 |
| 45 | 639.516 | 0.81 |
| 60 | 639.5035 | 0.77 |
| 300 | 639.492 | 0.74 |
| 1500 | 639.5335 | 0.86 |
| 3600 | 639.606 | 1.07 |
| 7200 | 639.603 | 1.06 |
| 14400 | 639.623 | 1.12 |

**ONPFDNA**

| Time | centroid | D |
|---|---|---|
| 0 | 639.23 | -0.01 |
| 30 | 639.42 | 0.53 |
| 45 | 639.48 | 0.70 |
| 60 | 639.48 | 0.71 |
| 300 | 639.46 | 0.65 |
| 1500 | 639.47 | 0.67 |
| 3600 | 639.58 | 0.98 |
| 7200 | 639.60 | 1.04 |
| 14400 | 639.57 | 0.96 |

**TMG**

| Time | centroid | D |
|---|---|---|
| 0 | 639.2375 | 0.01 |
| 30 | 639.4845 | 0.72 |
| 45 | 639.541 | 0.88 |
| 60 | 639.493 | 0.74 |
| 300 | 639.5345 | 0.86 |
| 1500 | 639.613 | 1.09 |
| 3600 | 639.563 | 0.94 |
| 7200 | 639.576 | 0.98 |
| 14400 | 639.6835 | 1.29 |

control 639.234
infinity 642.719

$$D(t) = \frac{M_t - M_0}{M_\infty - M_0} \cdot N$$

**IPTG**

| Time | Replicate 1 | Replicate 2 | Replicate 3 | Replicate 4 | Replicate5 | average | SD |
|---|---|---|---|---|---|---|---|
| 0 | 639.234 | 639.229 | | | | 639.23 | 0.00 |
| 30 | 639.561 | 639.415 | | | | 639.49 | 0.10 |
| 45 | 639.568 | 639.439 | | | | 639.50 | 0.09 |
| 60 | 639.604 | 639.438 | | | | 639.52 | 0.12 |
| 300 | 639.589 | 639.412 | | | | 639.50 | 0.13 |
| 1500 | 639.566 | 639.46 | | | | 639.51 | 0.07 |
| 3600 | 639.585 | 639.626 | | | | 639.61 | 0.03 |
| 7200 | 639.61 | 639.514 | | | | 639.56 | 0.07 |
| 14400 | 639.682 | 639.505 | | | | 639.59 | 0.13 |

**ONPF**

| Time | Replicate 1 | Replicate 2 | Replicate 3 | Replicate 4 | average | SD |
|---|---|---|---|---|---|---|
| 0 | 639.253 | 639.262 | | | 639.26 | 0.01 |
| 30 | 639.448 | 639.393 | | | 639.42 | 0.04 |
| 45 | 639.583 | 639.431 | | | 639.51 | 0.11 |
| 60 | 639.542 | 639.494 | | | 639.52 | 0.03 |
| 300 | 639.56 | 639.444 | | | 639.50 | 0.08 |
| 1500 | 639.617 | 639.506 | | | 639.56 | 0.08 |
| 3600 | 639.695 | 639.509 | | | 639.60 | 0.13 |
| 7200 | 639.684 | 639.544 | | | 639.61 | 0.10 |
| 14400 | 639.792 | 639.442 | | | 639.62 | 0.25 |

**APO**

| Time | Replicate 1 | Replicate 2 | Replicate 3 | Replicate 4 | Replicate 5 | average | SD |
|---|---|---|---|---|---|---|---|
| 0 | 639.225 | 639.234 | | | | 639.23 | 0.01 |
| 30 | 639.475 | 639.388 | | | | 639.43 | 0.06 |
| 45 | 639.498 | 639.403 | | | | 639.45 | 0.00 |
| 60 | 639.492 | 639.487 | | | | 639.49 | 0.00 |
| 300 | 639.494 | 639.433 | | | | 639.46 | 0.04 |
| 1500 | 639.596 | 639.584 | | | | 639.59 | 0.01 |
| 3600 | 639.65 | 639.593 | | | | 639.62 | 0.04 |
| 7200 | 639.734 | 639.585 | | | | 639.66 | 0.11 |
| 14400 | 640.069 | 639.648 | | | | 639.86 | 0.30 |

**DNA**

| Time | Replicate 1 | Replicate 2 | Replicate 3 | Replicate 4 | Replicate 5 | average | SD |
|---|---|---|---|---|---|---|---|
| 0 | 639.229 | 639.232 | | | | 639.23 | 0.00 |
| 30 | 639.48 | 639.395 | | | | 639.44 | 0.06 |
| 45 | 639.623 | 639.409 | | | | 639.52 | 0.15 |
| 60 | 639.523 | 639.484 | | | | 639.50 | 0.03 |
| 300 | 639.551 | 639.433 | | | | 639.49 | 0.08 |
| 1500 | 639.563 | 639.504 | | | | 639.53 | 0.04 |
| 3600 | 639.727 | 639.485 | | | | 639.61 | 0.17 |
| 7200 | 639.685 | 639.521 | | | | 639.60 | 0.12 |
| 14400 | 639.807 | 639.439 | | | | 639.62 | 0.26 |

**ONPFDNA**

| Time | Replicate 1 | Replicate 2 | Replicate 3 | Replicate 4 | average | SD |
|---|---|---|---|---|---|---|
| 0 | 639.234 | 639.229 | | | 639.23 | 0.00 |
| 30 | 639.452 | 639.388 | | | 639.42 | 0.05 |
| 45 | 639.531 | 639.426 | | | 639.48 | 0.07 |
| 60 | 639.511 | 639.455 | | | 639.48 | 0.04 |
| 300 | 639.498 | 639.422 | | | 639.46 | 0.05 |
| 1500 | 639.476 | 639.461 | | | 639.47 | 0.01 |
| 3600 | 639.673 | 639.478 | | | 639.58 | 0.14 |
| 7200 | 639.686 | 639.509 | | | 639.60 | 0.13 |
| 14400 | 639.616 | 639.519 | | | 639.57 | 0.07 |

**TMG**

| Time | Replicate 1 | Replicate 2 | Replicate 3 | Replicate 4 | Replicate 5 | average | SD |
|---|---|---|---|---|---|---|---|
| 0 | 639.244 | 639.231 | | | | 639.24 | 0.01 |
| 30 | 639.416 | 639.553 | | | | 639.48 | 0.10 |
| 45 | 639.534 | 639.548 | | | | 639.54 | 0.01 |
| 60 | 639.493 | | | | | 639.49 | |
| 300 | 639.504 | 639.565 | | | | 639.53 | 0.04 |
| 1500 | 639.509 | 639.717 | | | | 639.61 | 0.15 |
| 3600 | 639.569 | 639.557 | | | | 639.56 | 0.01 |
| 7200 | 639.542 | 639.61 | | | | 639.58 | 0.05 |
| 14400 | 639.629 | 639.738 | | | | 639.68 | 0.08 |

### 256-267 AITESGLRVGAD

**Charge** 2

**IPTG**

| Time | centroid | D |
|---|---|---|
| 0 | 595.162 | 0.01 |
| 30 | 595.9025 | 2.26 |
| 45 | 596.053 | 2.72 |
| 60 | 596.132 | 2.96 |
| 300 | 596.073 | 2.78 |
| 1500 | 596.1225 | 2.94 |
| 3600 | 596.257 | 3.35 |
| 7200 | 596.3055 | 3.49 |
| 14400 | 596.409 | 3.81 |

**ONPF**

| Time | centroid | D |
|---|---|---|
| 0 | 595.2155 | 0.17 |
| 30 | 595.5995 | 1.34 |
| 45 | 595.8665 | 2.15 |
| 60 | 595.949 | 2.41 |
| 300 | 595.9175 | 2.31 |
| 1500 | 596.006 | 2.58 |
| 3600 | 596.0065 | 2.58 |
| 7200 | 595.992 | 2.54 |
| 14400 | 596.07 | 2.76 |

**APO**

| Time | centroid | D |
|---|---|---|
| 0 | 595.1605 | 0.00 |
| 30 | 595.809 | 1.98 |
| 45 | 595.9 | 2.26 |
| 60 | 596.0435 | 2.69 |
| 300 | 595.94 | 2.38 |
| 1500 | 596.088 | 2.83 |
| 3600 | 596.1715 | 3.08 |
| 7200 | 596.164 | 3.06 |
| 14400 | 596.518 | 4.14 |

**DNA**

| Time | centroid | D |
|---|---|---|
| 0 | 595.1715 | 0.04 |
| 30 | 595.7555 | 1.82 |
| 45 | 595.934 | 2.36 |
| 60 | 595.9285 | 2.34 |
| 300 | 595.968 | 2.46 |
| 1500 | 596.0335 | 2.66 |
| 3600 | 596.0405 | 2.69 |
| 7200 | 595.998 | 2.56 |
| 14400 | 596.022 | 2.63 |

**ONPFDNA**

| Time | centroid | D |
|---|---|---|
| 0 | 595.16 | 0.01 |
| 30 | 595.71 | 1.68 |
| 45 | 595.92 | 2.32 |
| 60 | 595.89 | 2.24 |
| 300 | 595.95 | 2.41 |
| 1500 | 595.87 | 2.15 |
| 3600 | 596.08 | 2.81 |
| 7200 | 596.05 | 2.72 |
| 14400 | 595.91 | 2.29 |

**TMG**

| Time | centroid | D |
|---|---|---|
| 0 | 595.1755 | 0.05 |
| 30 | 595.865 | 2.15 |
| 45 | 596.017 | 2.61 |
| 60 | 596.14 | 2.99 |
| 300 | 596.141 | 2.99 |
| 1500 | 596.172 | 3.09 |
| 3600 | 596.2425 | 3.30 |
| 7200 | 596.269 | 3.38 |
| 14400 | 596.4835 | 4.04 |

| | |
|---|---|
| control | 595.16 |
| infinity | 598.439 |

$$D(t) = \frac{M_t - M_0}{M_\infty - M_0} * N$$

**IPTG**

| Time | Replicate 1 | Replicate 2 | Replicate 3 | Replicate 4 | Replicate5 | average | SD |
|---|---|---|---|---|---|---|---|
| 0 | 595.168 | 595.156 | | | | 595.16 | 0.01 |
| 30 | 596.089 | 595.716 | | | | 595.90 | 0.26 |
| 45 | 596.144 | 595.962 | | | | 596.05 | 0.13 |
| 60 | 596.248 | 596.016 | | | | 596.13 | 0.16 |
| 300 | 596.217 | 595.929 | | | | 596.07 | 0.20 |
| 1500 | 596.312 | 595.933 | | | | 596.12 | 0.27 |
| 3600 | 596.355 | 596.159 | | | | 596.26 | 0.14 |
| 7200 | 596.456 | 596.155 | | | | 596.31 | 0.21 |
| 14400 | 596.585 | 596.233 | | | | 596.41 | 0.25 |

**ONPF**

| Time | Replicate 1 | Replicate 2 | Replicate 3 | Replicate 4 | average | SD |
|---|---|---|---|---|---|---|
| 0 | 595.216 | 595.215 | | | 595.22 | 0.00 |
| 30 | 595.674 | 595.525 | | | 595.60 | 0.11 |
| 45 | 596.046 | 595.687 | | | 595.87 | 0.25 |
| 60 | 595.985 | 595.913 | | | 595.95 | 0.05 |
| 300 | 596.05 | 595.785 | | | 595.92 | 0.19 |
| 1500 | 596.102 | 595.91 | | | 596.01 | 0.14 |
| 3600 | 596.126 | 595.887 | | | 596.01 | 0.17 |
| 7200 | 596.118 | 595.866 | | | 595.99 | 0.18 |
| 14400 | 596.385 | 595.748 | | | 596.07 | 0.45 |

**APO**

| Time | Replicate 1 | Replicate 2 | Replicate 3 | Replicate 4 | Replicate 5 | average | SD |
|---|---|---|---|---|---|---|---|
| 0 | 595.152 | 595.169 | | | | 595.16 | 0.01 |
| 30 | 595.945 | 595.673 | | | | 595.81 | 0.19 |
| 45 | 596.034 | 595.766 | | | | 595.90 | 0.00 |
| 60 | 596.131 | 595.956 | | | | 596.04 | 0.12 |
| 300 | 596.093 | 595.787 | | | | 595.94 | 0.22 |
| 1500 | 596.132 | 596.044 | | | | 596.09 | 0.06 |
| 3600 | 596.253 | 596.09 | | | | 596.17 | 0.12 |
| 7200 | 596.369 | 595.959 | | | | 596.16 | 0.29 |
| 14400 | 596.737 | 596.299 | | | | 596.52 | 0.31 |

**DNA**

| Time | Replicate 1 | Replicate 2 | Replicate 3 | Replicate 4 | Replicate 5 | average | SD |
|---|---|---|---|---|---|---|---|
| 0 | 595.153 | 595.19 | | | | 595.17 | 0.03 |
| 30 | 595.796 | 595.715 | | | | 595.76 | 0.06 |
| 45 | 596.063 | 595.805 | | | | 595.93 | 0.18 |
| 60 | 595.976 | 595.881 | | | | 595.93 | 0.07 |
| 300 | 596.016 | 595.92 | | | | 595.97 | 0.07 |
| 1500 | 596.085 | 595.982 | | | | 596.03 | 0.07 |
| 3600 | 596.164 | 595.917 | | | | 596.04 | 0.17 |
| 7200 | 596.193 | 595.803 | | | | 596.00 | 0.28 |
| 14400 | 596.295 | 595.749 | | | | 596.02 | 0.39 |

**ONPFDNA**

| Time | Replicate 1 | Replicate 2 | Replicate 3 | Replicate 4 | average | SD |
|---|---|---|---|---|---|---|
| 0 | 595.161 | 595.163 | | | 595.16 | 0.00 |
| 30 | 595.73 | 595.691 | | | 595.71 | 0.03 |
| 45 | 595.984 | 595.855 | | | 595.92 | 0.09 |
| 60 | 595.898 | 595.888 | | | 595.89 | 0.01 |
| 300 | 596.003 | 595.899 | | | 595.95 | 0.07 |
| 1500 | 595.833 | 595.899 | | | 595.87 | 0.05 |
| 3600 | 596.116 | 596.05 | | | 596.08 | 0.05 |
| 7200 | 596.17 | 595.936 | | | 596.05 | 0.17 |
| 14400 | 595.908 | 595.917 | | | 595.91 | 0.01 |

**TMG**

| Time | Replicate 1 | Replicate 2 | Replicate 3 | Replicate 4 | Replicate 5 | average | SD |
|---|---|---|---|---|---|---|---|
| 0 | 595.197 | 595.154 | | | | 595.18 | 0.03 |
| 30 | 595.661 | 596.069 | | | | 595.87 | 0.29 |
| 45 | 596.001 | 596.033 | | | | 596.02 | 0.02 |
| 60 | 595.974 | 596.306 | | | | 596.14 | 0.23 |
| 300 | 596.034 | 596.248 | | | | 596.14 | 0.15 |
| 1500 | 595.978 | 596.366 | | | | 596.17 | 0.27 |
| 3600 | 596.141 | 596.344 | | | | 596.24 | 0.14 |
| 7200 | 596.069 | 596.469 | | | | 596.27 | 0.28 |
| 14400 | 596.349 | 596.618 | | | | 596.48 | 0.19 |

### 257-267 ITESGLRVGAD

**Charge** 2

**IPTG**

| Time | centroid | D |
|---|---|---|
| 0 | 559.62 | -0.04 |
| 30 | 560.23 | 1.91 |
| 45 | 560.49 | 2.73 |
| 60 | 560.51 | 2.78 |
| 300 | 560.45 | 2.59 |
| 1500 | 560.51 | 2.78 |
| 3600 | 560.57 | 2.99 |
| 7200 | 560.68 | 3.33 |
| 14400 | 560.75 | 3.56 |

**ONPF**

| Time | centroid | D |
|---|---|---|
| 0 | 559.65 | 0.06 |
| 30 | 560.09 | 1.44 |
| 45 | 560.30 | 2.13 |
| 60 | 560.37 | 2.33 |
| 300 | 560.32 | 2.18 |
| 1500 | 560.40 | 2.43 |
| 3600 | 560.45 | 2.59 |
| 7200 | 560.45 | 2.59 |
| 14400 | 560.59 | 3.05 |

**APO**

| Time | centroid | D |
|---|---|---|
| 0 | 559.6262 | -0.01 |
| 30 | 560.2032 | 1.81 |
| 45 | 560.349 | 2.28 |
| 60 | 560.4542 | 2.61 |
| 300 | 560.371 | 2.34 |
| 1500 | 560.4782 | 2.68 |
| 3600 | 560.5496 | 2.91 |
| 7200 | 560.6272 | 3.16 |
| 14400 | 560.7978 | 3.70 |

**DNA**

| Time | centroid | D |
|---|---|---|
| 0 | 559.62 | -0.03 |
| 30 | 560.14 | 1.63 |
| 45 | 560.35 | 2.28 |
| 60 | 560.39 | 2.41 |
| 300 | 560.39 | 2.41 |
| 1500 | 560.44 | 2.57 |
| 3600 | 560.49 | 2.71 |
| 7200 | 560.41 | 2.48 |
| 14400 | 560.49 | 2.72 |

**ONPFDNA**

| Time | centroid | D |
|---|---|---|
| 0 | 559.60 | -0.09 |
| 30 | 560.15 | 1.63 |
| 45 | 560.36 | 2.32 |
| 60 | 560.38 | 2.36 |
| 300 | 560.40 | 2.43 |
| 1500 | 560.29 | 2.08 |
| 3600 | 560.50 | 2.76 |
| 7200 | 560.42 | 2.49 |
| 14400 | 560.39 | 2.42 |

**TMG**

| Time | centroid | D |
|---|---|---|
| 0 | 559.64 | 0.04 |
| 30 | 560.33 | 2.23 |
| 45 | 560.47 | 2.66 |
| 60 | 560.56 | 2.95 |
| 300 | 560.57 | 2.97 |
| 1500 | 560.60 | 3.08 |
| 3600 | 560.67 | 3.28 |
| 7200 | 560.74 | 3.51 |
| 14400 | 560.88 | 3.97 |

control 559.63
infinity 562.474

$$D(t) = \frac{M_t - M_0}{M_\infty - M_0} * N$$

**IPTG**

| Time | Replicate 1 | Replicate 2 | Replicate 3 | Replicate 4 | Replicate5 | average | SD |
|---|---|---|---|---|---|---|---|
| 0 | 559.622 | 559.61 | 559.623 | 559.613 | 559.614 | 559.62 | 0.01 |
| 30 | 560.531 | 560.138 | 560.262 | 560.136 | 560.094 | 560.23 | 0.18 |
| 45 | 560.589 | 560.394 | | | | 560.49 | 0.14 |
| 60 | 560.637 | 560.448 | 560.693 | 560.4 | 560.362 | 560.51 | 0.15 |
| 300 | 560.666 | 560.367 | 560.563 | 560.373 | 560.273 | 560.45 | 0.16 |
| 1500 | 560.727 | 560.375 | 560.646 | 560.429 | 560.359 | 560.51 | 0.17 |
| 3600 | 560.769 | 560.602 | 560.629 | 560.461 | 560.41 | 560.57 | 0.14 |
| 7200 | 560.915 | 560.627 | 560.769 | 560.587 | 560.52 | 560.68 | 0.16 |
| 14400 | 561.021 | 560.662 | 560.753 | 560.753 | 560.578 | 560.75 | 0.17 |

**ONPF**

| Time | Replicate 1 | Replicate 2 | Replicate 3 | Replicate 4 | average | SD |
|---|---|---|---|---|---|---|
| 0 | 559.683 | 559.671 | 559.619 | 559.619 | 559.65 | 0.03 |
| 30 | 560.112 | 559.981 | 560.23 | 560.017 | 560.09 | 0.11 |
| 45 | 560.474 | 560.132 | | | 560.30 | 0.24 |
| 60 | 560.407 | 560.332 | 560.386 | 560.342 | 560.37 | 0.04 |
| 300 | 560.496 | 560.205 | 560.379 | 560.2 | 560.32 | 0.14 |
| 1500 | 560.526 | 560.317 | 560.391 | 560.363 | 560.40 | 0.09 |
| 3600 | 560.537 | 560.313 | 560.596 | 560.349 | 560.45 | 0.14 |
| 7200 | 560.535 | 560.299 | 560.529 | 560.435 | 560.45 | 0.11 |
| 14400 | 560.811 | 560.167 | 560.716 | 560.684 | 560.59 | 0.29 |

**APO**

| Time | Replicate 1 | Replicate 2 | Replicate 3 | Replicate 4 | Replicate 5 | average | SD |
|---|---|---|---|---|---|---|---|
| 0 | 559.605 | 559.624 | 559.677 | 559.617 | 559.608 | 559.63 | 0.03 |
| 30 | 560.364 | 560.117 | 560.453 | 560.03 | 560.052 | 560.20 | 0.19 |
| 45 | 560.495 | 560.203 | | | | 560.35 | 0.21 |
| 60 | 560.551 | 560.382 | 560.727 | 560.306 | 560.305 | 560.45 | 0.18 |
| 300 | 560.532 | 560.204 | 560.626 | 560.289 | 560.204 | 560.37 | 0.20 |
| 1500 | 560.568 | 560.466 | 560.696 | 560.33 | 560.331 | 560.48 | 0.16 |
| 3600 | 560.667 | 560.484 | 560.816 | 560.403 | 560.378 | 560.55 | 0.19 |
| 7200 | 560.799 | 560.381 | 560.916 | 560.589 | 560.451 | 560.63 | 0.23 |
| 14400 | 561.002 | 560.737 | 560.891 | 560.685 | 560.674 | 560.80 | 0.14 |

**DNA**

| Time | Replicate 1 | Replicate 2 | Replicate 3 | Replicate 4 | Replicate 5 | average | SD |
|---|---|---|---|---|---|---|---|
| 0 | 559.613 | 559.638 | 559.618 | 559.611 | | 559.62 | 0.01 |
| 30 | 560.282 | 560.15 | 560.212 | 559.934 | | 560.14 | 0.15 |
| 45 | 560.463 | 560.239 | | | | 560.35 | 0.16 |
| 60 | 560.415 | 560.311 | 560.624 | 560.218 | | 560.39 | 0.17 |
| 300 | 560.5 | 560.353 | 560.597 | 560.122 | | 560.39 | 0.21 |
| 1500 | 560.526 | 560.427 | 560.632 | 560.189 | | 560.44 | 0.19 |
| 3600 | 560.68 | 560.343 | 560.67 | 560.252 | | 560.49 | 0.22 |
| 7200 | 560.601 | 560.189 | 560.625 | 560.244 | | 560.41 | 0.23 |
| 14400 | 560.742 | 560.154 | 560.708 | 560.357 | | 560.49 | 0.28 |

**ONPFDNA**

| Time | Replicate 1 | Replicate 2 | Replicate 3 | Replicate 4 | average | SD |
|---|---|---|---|---|---|---|
| 0 | 559.619 | 559.618 | 559.556 | 559.614 | 559.60 | 0.03 |
| 30 | 560.178 | 560.103 | 560.371 | 559.929 | 560.15 | 0.18 |
| 45 | 560.429 | 560.296 | | | 560.36 | 0.09 |
| 60 | 560.33 | 560.319 | 560.58 | 560.279 | 560.38 | 0.14 |
| 300 | 560.446 | 560.327 | 560.521 | 560.292 | 560.40 | 0.11 |
| 1500 | 560.265 | 560.321 | 560.273 | 560.293 | 560.29 | 0.02 |
| 3600 | 560.562 | 560.486 | 560.627 | 560.335 | 560.50 | 0.13 |
| 7200 | 560.527 | 560.315 | | 560.409 | 560.42 | 0.11 |
| 14400 | 560.294 | 560.34 | 560.479 | 560.466 | 560.39 | 0.09 |

**TMG**

| Time | Replicate 1 | Replicate 2 | Replicate 3 | Replicate 4 | Replicate 5 | average | SD |
|---|---|---|---|---|---|---|---|
| 0 | 559.661 | 559.624 | | | | 559.64 | 0.03 |
| 30 | 560.095 | 560.573 | | | | 560.33 | 0.34 |
| 45 | 560.425 | 560.516 | | | | 560.47 | 0.06 |
| 60 | 560.437 | 560.69 | | | | 560.56 | 0.18 |
| 300 | 560.462 | 560.672 | | | | 560.57 | 0.15 |
| 1500 | 560.42 | 560.785 | | | | 560.60 | 0.26 |
| 3600 | 560.55 | 560.784 | | | | 560.67 | 0.17 |
| 7200 | 560.535 | 560.943 | | | | 560.74 | 0.29 |
| 14400 | 560.757 | 561.011 | | | | 560.88 | 0.18 |

### 266-273 ADISVVGY

**Charge** 1

**IPTG**

| Time | centroid | D |
|---|---|---|
| 0 | 823.917 | 0.02 |
| 30 | 824.122 | 0.27 |
| 45 | 824.093 | 0.23 |
| 60 | 824.1735 | 0.33 |
| 300 | 824.152 | 0.31 |
| 1500 | 824.0955 | 0.24 |
| 3600 | 824.1005 | 0.24 |
| 7200 | 824.1275 | 0.28 |
| 14400 | 824.1135 | 0.26 |

**ONPF**

| Time | centroid | D |
|---|---|---|
| 0 | 823.92 | 0.03 |
| 30 | 824.08 | 0.22 |
| 45 | 824.13 | 0.28 |
| 60 | 824.14 | 0.29 |
| 300 | 824.12 | 0.27 |
| 1500 | 824.21 | 0.37 |
| 3600 | 824.13 | 0.28 |
| 7200 | 824.08 | 0.22 |
| 14400 | 824.08 | 0.22 |

**APO**

| Time | centroid | D |
|---|---|---|
| 0 | 823.904 | 0.00 |
| 30 | 824.062 | 0.20 |
| 45 | 824.0585 | 0.19 |
| 60 | 824.0815 | 0.22 |
| 300 | 824.0875 | 0.23 |
| 1500 | 824.0905 | 0.23 |
| 3600 | 824.0915 | 0.23 |
| 7200 | 824.0835 | 0.22 |
| 14400 | 824.314 | 0.50 |

**DNA**

| Time | centroid | D |
|---|---|---|
| 0 | 823.909 | 0.01 |
| 30 | 824.0855 | 0.23 |
| 45 | 824.127 | 0.28 |
| 60 | 824.1505 | 0.30 |
| 300 | 824.112 | 0.26 |
| 1500 | 824.0855 | 0.23 |
| 3600 | 824.094 | 0.24 |
| 7200 | 824.07 | 0.21 |
| 14400 | 824.093 | 0.23 |

**ONPFDNA**

| Time | centroid | D |
|---|---|---|
| 0 | 823.897 | 0.00 |
| 30 | 824.091 | 0.23 |
| 45 | 824.123 | 0.27 |
| 60 | 824.115 | 0.26 |
| 300 | 824.075 | 0.21 |
| 1500 | 824.1015 | 0.24 |
| 3600 | 824.076 | 0.21 |
| 7200 | 824.078 | 0.22 |
| 14400 | 824.099 | 0.24 |

**TMG**

| Time | centroid | D |
|---|---|---|
| 0 | 823.91 | 0.02 |
| 30 | 824.11 | 0.26 |
| 45 | 824.15 | 0.30 |
| 60 | 824.18 | 0.34 |
| 300 | 824.17 | 0.32 |
| 1500 | 824.24 | 0.41 |
| 3600 | 824.16 | 0.32 |
| 7200 | 824.14 | 0.29 |
| 14400 | 824.20 | 0.37 |

control 823.9
infinity 828.844

$$D(t) = \frac{Mt - M0}{M\infty - M0} *N$$

**IPTG**

| Time | Replicate 1 | Replicate 2 | Replicate 3 | Replicate 4 | Replicate5 | average | SD |
|---|---|---|---|---|---|---|---|
| 0 | 823.938 | 823.896 | | | | 823.92 | 0.03 |
| 30 | 824.188 | 824.056 | | | | 824.12 | 0.09 |
| 45 | 824.129 | 824.057 | | | | 824.09 | 0.05 |
| 60 | 824.281 | 824.066 | | | | 824.17 | 0.15 |
| 300 | 824.232 | 824.072 | | | | 824.15 | 0.11 |
| 1500 | 824.143 | 824.048 | | | | 824.10 | 0.07 |
| 3600 | 824.151 | 824.05 | | | | 824.10 | 0.07 |
| 7200 | 824.162 | 824.093 | | | | 824.13 | 0.05 |
| 14400 | 824.163 | 824.064 | | | | 824.11 | 0.07 |

**ONPF**

| Time | Replicate 1 | Replicate 2 | Replicate 3 | Replicate 4 | average | SD |
|---|---|---|---|---|---|---|
| 0 | 823.928 | 823.914 | | | 823.92 | 0.01 |
| 30 | 824.112 | 824.047 | | | 824.08 | 0.05 |
| 45 | 824.146 | 824.11 | | | 824.13 | 0.03 |
| 60 | 824.152 | 824.119 | | | 824.14 | 0.02 |
| 300 | 824.133 | 824.106 | | | 824.12 | 0.02 |
| 1500 | 824.303 | 824.112 | | | 824.21 | 0.14 |
| 3600 | 824.161 | 824.096 | | | 824.13 | 0.05 |
| 7200 | 824.1 | 824.061 | | | 824.08 | 0.03 |
| 14400 | 824.174 | 823.983 | | | 824.08 | 0.14 |

**APO**

| Time | Replicate 1 | Replicate 2 | Replicate 3 | Replicate 4 | Replicate 5 | average | SD |
|---|---|---|---|---|---|---|---|
| 0 | 823.908 | 823.9 | | | | 823.90 | 0.01 |
| 30 | 824.079 | 824.045 | | | | 824.06 | 0.02 |
| 45 | 824.048 | 824.069 | | | | 824.06 | 0.01 |
| 60 | 824.083 | 824.08 | | | | 824.08 | 0.00 |
| 300 | 824.08 | 824.095 | | | | 824.09 | 0.01 |
| 1500 | 824.08 | 824.101 | | | | 824.09 | 0.01 |
| 3600 | 824.089 | 824.094 | | | | 824.09 | 0.00 |
| 7200 | 824.1 | 824.067 | | | | 824.08 | 0.02 |
| 14400 | 824.46 | 824.168 | | | | 824.31 | 0.21 |

**DNA**

| Time | Replicate 1 | Replicate 2 | Replicate 3 | Replicate 4 | Replicate 5 | average | SD |
|---|---|---|---|---|---|---|---|
| 0 | 823.926 | 823.892 | | | | 823.91 | 0.02 |
| 30 | 824.152 | 824.019 | | | | 824.09 | 0.09 |
| 45 | 824.204 | 824.05 | | | | 824.13 | 0.11 |
| 60 | 824.244 | 824.057 | | | | 824.15 | 0.13 |
| 300 | 824.176 | 824.048 | | | | 824.11 | 0.09 |
| 1500 | 824.131 | 824.04 | | | | 824.09 | 0.06 |
| 3600 | 824.15 | 824.038 | | | | 824.09 | 0.08 |
| 7200 | 824.096 | 824.044 | | | | 824.07 | 0.04 |
| 14400 | 824.142 | 824.044 | | | | 824.09 | 0.07 |

**ONPFDNA**

| Time | Replicate 1 | Replicate 2 | Replicate 3 | Replicate 4 | average | SD |
|---|---|---|---|---|---|---|
| 0 | 823.903 | 823.891 | | | 823.90 | 0.01 |
| 30 | 824.136 | 824.046 | | | 824.09 | 0.06 |
| 45 | 824.183 | 824.063 | | | 824.12 | 0.08 |
| 60 | 824.131 | 824.099 | | | 824.12 | 0.02 |
| 300 | 824.101 | 824.049 | | | 824.08 | 0.04 |
| 1500 | 824.134 | 824.069 | | | 824.10 | 0.05 |
| 3600 | 824.115 | 824.037 | | | 824.08 | 0.06 |
| 7200 | 824.121 | 824.035 | | | 824.08 | 0.06 |
| 14400 | 824.11 | 824.088 | | | 824.10 | 0.02 |

**TMG**

| Time | Replicate 1 | Replicate 2 | Replicate 3 | Replicate 4 | Replicate 5 | average | SD |
|---|---|---|---|---|---|---|---|
| 0 | 823.906 | 823.921 | | | | 823.91 | 0.01 |
| 30 | 824.082 | 824.146 | | | | 824.11 | 0.05 |
| 45 | 824.166 | 824.136 | | | | 824.15 | 0.02 |
| 60 | 824.181 | | | | | 824.18 | |
| 300 | 824.162 | 824.17 | | | | 824.17 | 0.01 |
| 1500 | 824.137 | 824.333 | | | | 824.24 | 0.14 |
| 3600 | 824.177 | 824.144 | | | | 824.16 | 0.02 |
| 7200 | 824.131 | 824.15 | | | | 824.14 | 0.01 |
| 14400 | 824.159 | 824.249 | | | | 824.20 | 0.06 |

### 270-275 VVGYDD

**Charge** 1

**IPTG**

| Time | centroid | D |
|---|---|---|
| 0 | 667.66 | 0.01 |
| 30 | 667.89 | 0.52 |
| 45 | 667.87 | 0.47 |
| 60 | 667.91 | 0.57 |
| 300 | 667.88 | 0.51 |
| 1500 | 667.86 | 0.47 |
| 3600 | 667.94 | 0.63 |
| 7200 | 667.87 | 0.48 |
| 14400 | 667.87 | 0.48 |

**ONPF**

| Time | centroid | D |
|---|---|---|
| 0 | 667.65 | 0.00 |
| 30 | 667.82 | 0.38 |
| 45 | 667.87 | 0.48 |
| 60 | 667.86 | 0.45 |
| 300 | 667.84 | 0.42 |
| 1500 | 667.87 | 0.48 |
| 3600 | 667.87 | 0.49 |
| 7200 | 667.90 | 0.55 |
| 14400 | 667.84 | 0.42 |

**APO**

| Time | centroid | D |
|---|---|---|
| 0 | 667.65 | 0.01 |
| 30 | 667.83 | 0.38 |
| 45 | 667.82 | 0.36 |
| 60 | 667.83 | 0.39 |
| 300 | 667.80 | 0.33 |
| 1500 | 667.86 | 0.47 |
| 3600 | 667.90 | 0.55 |
| 7200 | 667.93 | 0.62 |
| 14400 | 668.10 | 0.99 |

**DNA**

| Time | centroid | D |
|---|---|---|
| 0 | 667.64 | -0.01 |
| 30 | 667.83 | 0.40 |
| 45 | 667.82 | 0.38 |
| 60 | 667.82 | 0.36 |
| 300 | 667.83 | 0.39 |
| 1500 | 667.84 | 0.42 |
| 3600 | 667.84 | 0.42 |
| 7200 | 667.85 | 0.43 |
| 14400 | 667.83 | 0.40 |

**ONPFDNA**

| Time | centroid | D |
|---|---|---|
| 0 | 667.64 | -0.01 |
| 30 | 667.83 | 0.40 |
| 45 | 667.86 | 0.45 |
| 60 | 667.83 | 0.39 |
| 300 | 667.80 | 0.33 |
| 1500 | 667.78 | 0.28 |
| 3600 | 667.83 | 0.39 |
| 7200 | 667.82 | 0.38 |
| 14400 | 667.80 | 0.32 |

**TMG**

| Time | centroid | D |
|---|---|---|
| 0 | 667.65 | 0.00 |
| 30 | 667.89 | 0.52 |
| 45 | 667.92 | 0.59 |
| 60 | 667.91 | 0.57 |
| 300 | 667.94 | 0.63 |
| 1500 | 667.97 | 0.69 |
| 3600 | 667.91 | 0.58 |
| 7200 | 667.90 | 0.56 |
| 14400 | 667.98 | 0.73 |

control 667.65
infinity 669.477

$$D(t) = \frac{Mt - M0}{M\infty - M0} * N$$

**IPTG**

| Time | Replicate 1 | Replicate 2 | Replicate 3 | Replicate 4 | Replicate5 | average | SD |
|---|---|---|---|---|---|---|---|
| 0 | 667.66 | 667.65 | | | | 667.66 | 0.01 |
| 30 | 667.959 | 667.815 | | | | 667.89 | 0.10 |
| 45 | 667.94 | 667.793 | | | | 667.87 | 0.10 |
| 60 | 668.011 | 667.813 | | | | 667.91 | 0.14 |
| 300 | 667.971 | 667.793 | | | | 667.88 | 0.13 |
| 1500 | 667.925 | 667.804 | | | | 667.86 | 0.09 |
| 3600 | 668.065 | 667.812 | | | | 667.94 | 0.18 |
| 7200 | 667.92 | 667.823 | | | | 667.87 | 0.07 |
| 14400 | 667.925 | 667.809 | | | | 667.87 | 0.08 |

**ONPF**

| Time | Replicate 1 | Replicate 2 | Replicate 3 | Replicate 4 | average | SD |
|---|---|---|---|---|---|---|
| 0 | 667.647 | 667.657 | | | 667.65 | 0.01 |
| 30 | 667.858 | 667.789 | | | 667.82 | 0.05 |
| 45 | 667.923 | 667.812 | | | 667.87 | 0.08 |
| 60 | 667.9 | 667.813 | | | 667.86 | 0.06 |
| 300 | 667.898 | 667.782 | | | 667.84 | 0.08 |
| 1500 | 667.948 | 667.794 | | | 667.87 | 0.11 |
| 3600 | 668.002 | 667.743 | | | 667.87 | 0.18 |
| 7200 | 667.987 | 667.814 | | | 667.90 | 0.12 |
| 14400 | 668.068 | 667.612 | | | 667.84 | 0.32 |

**APO**

| Time | Replicate 1 | Replicate 2 | Replicate 3 | Replicate 4 | Replicate 5 | average | SD |
|---|---|---|---|---|---|---|---|
| 0 | 667.664 | 667.641 | | | | 667.65 | 0.02 |
| 30 | 667.855 | 667.796 | | | | 667.83 | 0.04 |
| 45 | 667.851 | 667.781 | | | | 667.82 | 0.05 |
| 60 | 667.843 | 667.809 | | | | 667.83 | 0.02 |
| 300 | 667.831 | 667.775 | | | | 667.80 | 0.04 |
| 1500 | 667.895 | 667.83 | | | | 667.86 | 0.05 |
| 3600 | 667.947 | 667.858 | | | | 667.90 | 0.06 |
| 7200 | 668.002 | 667.864 | | | | 667.93 | 0.10 |
| 14400 | 668.226 | 667.979 | | | | 668.10 | 0.17 |

**DNA**

| Time | Replicate 1 | Replicate 2 | Replicate 3 | Replicate 4 | Replicate 5 | average | SD |
|---|---|---|---|---|---|---|---|
| 0 | 667.654 | 667.633 | | | | 667.64 | 0.01 |
| 30 | 667.893 | 667.77 | | | | 667.83 | 0.09 |
| 45 | 667.86 | 667.783 | | | | 667.82 | 0.05 |
| 60 | 667.871 | 667.759 | | | | 667.82 | 0.08 |
| 300 | 667.893 | 667.763 | | | | 667.83 | 0.09 |
| 1500 | 667.878 | 667.804 | | | | 667.84 | 0.05 |
| 3600 | 667.921 | 667.763 | | | | 667.84 | 0.11 |
| 7200 | 667.909 | 667.784 | | | | 667.85 | 0.09 |
| 14400 | 667.936 | 667.727 | | | | 667.83 | 0.15 |

**ONPFDNA**

| Time | Replicate 1 | Replicate 2 | Replicate 3 | Replicate 4 | average | SD |
|---|---|---|---|---|---|---|
| 0 | 667.641 | 667.648 | | | 667.64 | 0.00 |
| 30 | 667.875 | 667.786 | | | 667.83 | 0.06 |
| 45 | 667.905 | 667.806 | | | 667.86 | 0.07 |
| 60 | 667.858 | 667.798 | | | 667.83 | 0.04 |
| 300 | 667.828 | 667.774 | | | 667.80 | 0.04 |
| 1500 | 667.787 | 667.765 | | | 667.78 | 0.02 |
| 3600 | 667.871 | 667.788 | | | 667.83 | 0.06 |
| 7200 | 667.89 | 667.758 | | | 667.82 | 0.09 |
| 14400 | 667.79 | 667.804 | | | 667.80 | 0.01 |

**TMG**

| Time | Replicate 1 | Replicate 2 | Replicate 3 | Replicate 4 | Replicate 5 | average | SD |
|---|---|---|---|---|---|---|---|
| 0 | 667.645 | 667.654 | | | | 667.65 | 0.01 |
| 30 | 667.863 | 667.916 | | | | 667.89 | 0.04 |
| 45 | 667.907 | 667.933 | | | | 667.92 | 0.02 |
| 60 | 667.91 | | | | | 667.91 | |
| 300 | 667.909 | 667.967 | | | | 667.94 | 0.04 |
| 1500 | 667.9 | 668.032 | | | | 667.97 | 0.09 |
| 3600 | 667.917 | 667.91 | | | | 667.91 | 0.00 |
| 7200 | 667.889 | 667.919 | | | | 667.90 | 0.02 |
| 14400 | 667.931 | 668.033 | | | | 667.98 | 0.07 |

### 274-290 DDTEDSSCYIPPLTTIK

**Charge** 2

**IPTG**

| Time | centroid | D |
|---|---|---|
| 0 | 950.0085 | -0.03 |
| 30 | 950.424 | 1.02 |
| 45 | 950.4475 | 1.08 |
| 60 | 950.567 | 1.39 |
| 300 | 950.804 | 1.99 |
| 1500 | 951.166 | 2.90 |
| 3600 | 951.407 | 3.51 |
| 7200 | 951.574 | 3.94 |
| 14400 | 951.6605 | 4.16 |

**ONPF**

| Time | centroid | D |
|---|---|---|
| 0 | 950.0145 | -0.01 |
| 30 | 950.335 | 0.80 |
| 45 | 950.5905 | 1.45 |
| 60 | 950.598 | 1.46 |
| 300 | 950.61 | 1.49 |
| 1500 | 950.9635 | 2.39 |
| 3600 | 951.247 | 3.11 |
| 7200 | 951.6 | 4.00 |
| 14400 | 952.08 | 5.22 |

**APO**

| Time | centroid | D |
|---|---|---|
| 0 | 950.02 | 0.00 |
| 30 | 950.381 | 0.91 |
| 45 | 950.4145 | 1.00 |
| 60 | 950.5705 | 1.39 |
| 300 | 950.613 | 1.50 |
| 1500 | 951.1745 | 2.92 |
| 3600 | 951.4885 | 3.72 |
| 7200 | 951.784 | 4.47 |
| 14400 | 952.248 | 5.64 |

**DNA**

| Time | centroid | D |
|---|---|---|
| 0 | 950.001 | -0.05 |
| 30 | 950.3295 | 0.78 |
| 45 | 950.3615 | 0.87 |
| 60 | 950.4085 | 0.98 |
| 300 | 950.4155 | 1.00 |
| 1500 | 950.5445 | 1.33 |
| 3600 | 950.519 | 1.26 |
| 7200 | 950.4885 | 1.19 |
| 14400 | 950.4555 | 1.10 |

**ONPFDNA**

| Time | centroid | D |
|---|---|---|
| 0 | 949.9975 | -0.06 |
| 30 | 950.3205 | 0.76 |
| 45 | 950.398 | 0.96 |
| 60 | 950.3985 | 0.96 |
| 300 | 950.381 | 0.91 |
| 1500 | 950.417 | 1.01 |
| 3600 | 950.5495 | 1.34 |
| 7200 | 950.5345 | 1.30 |
| 14400 | 950.416 | 1.00 |

**TMG**

| Time | centroid | D |
|---|---|---|
| 0 | 950.007 | -0.03 |
| 30 | 950.444 | 1.07 |
| 45 | 950.554 | 1.35 |
| 60 | 950.636 | 1.56 |
| 300 | 950.7855 | 1.94 |
| 1500 | 951.149 | 2.86 |
| 3600 | 951.4065 | 3.51 |
| 7200 | 951.494 | 3.73 |
| 14400 | 951.763 | 4.42 |

| | |
|---|---|
| control | 950.02 |
| infinity | 955.152 |

$$D(t) = \frac{Mt - M0}{M\infty - M0} *N$$

**IPTG**

| Time | Replicate 1 | Replicate 2 | Replicate 3 | Replicate 4 | Replicate5 | average | SD |
|---|---|---|---|---|---|---|---|
| 0 | 950.003 | 950.014 | | | | 950.01 | 0.01 |
| 30 | 950.319 | 950.529 | | | | 950.42 | 0.15 |
| 45 | 950.362 | 950.533 | | | | 950.45 | 0.12 |
| 60 | 950.466 | 950.668 | | | | 950.57 | 0.14 |
| 300 | 950.649 | 950.959 | | | | 950.80 | 0.22 |
| 1500 | 951.014 | 951.318 | | | | 951.17 | 0.21 |
| 3600 | 951.358 | 951.456 | | | | 951.41 | 0.07 |
| 7200 | 951.438 | 951.71 | | | | 951.57 | 0.19 |
| 14400 | 951.49 | 951.831 | | | | 951.66 | 0.24 |

**ONPF**

| Time | Replicate 1 | Replicate 2 | Replicate 3 | Replicate 4 | average | SD |
|---|---|---|---|---|---|---|
| 0 | 950.024 | 950.005 | | | 950.01 | 0.01 |
| 30 | 950.286 | 950.384 | | | 950.34 | 0.07 |
| 45 | 950.409 | 950.772 | | | 950.59 | 0.26 |
| 60 | 950.649 | 950.547 | | | 950.60 | 0.07 |
| 300 | 950.499 | 950.721 | | | 950.61 | 0.16 |
| 1500 | 950.831 | 951.096 | | | 950.96 | 0.19 |
| 3600 | 951.04 | 951.454 | | | 951.25 | 0.29 |
| 7200 | 951.438 | 951.762 | | | 951.60 | 0.23 |
| 14400 | | 952.082 | | | 952.08 | |

**APO**

| Time | Replicate 1 | Replicate 2 | Replicate 3 | Replicate 4 | Replicate 5 | average | SD |
|---|---|---|---|---|---|---|---|
| 0 | 950.016 | 950.024 | | | | 950.02 | 0.01 |
| 30 | 950.319 | 950.443 | | | | 950.38 | 0.09 |
| 45 | 950.334 | 950.495 | | | | 950.41 | 0.11 |
| 60 | 950.606 | 950.535 | | | | 950.57 | 0.05 |
| 300 | 950.524 | 950.702 | | | | 950.61 | 0.13 |
| 1500 | 951.272 | 951.077 | | | | 951.17 | 0.14 |
| 3600 | 951.399 | 951.578 | | | | 951.49 | 0.13 |
| 7200 | 951.621 | 951.947 | | | | 951.78 | 0.23 |
| 14400 | 952.081 | 952.415 | | | | 952.25 | 0.24 |

**DNA**

| Time | Replicate 1 | Replicate 2 | Replicate 3 | Replicate 4 | Replicate 5 | average | SD |
|---|---|---|---|---|---|---|---|
| 0 | 949.998 | 950.004 | | | | 950.00 | 0.00 |
| 30 | 950.3 | 950.359 | | | | 950.33 | 0.04 |
| 45 | 950.312 | 950.411 | | | | 950.36 | 0.07 |
| 60 | 950.392 | 950.425 | | | | 950.41 | 0.02 |
| 300 | 950.367 | 950.464 | | | | 950.42 | 0.07 |
| 1500 | 950.529 | 950.56 | | | | 950.54 | 0.02 |
| 3600 | 950.455 | 950.583 | | | | 950.52 | 0.09 |
| 7200 | 950.398 | 950.579 | | | | 950.49 | 0.13 |
| 14400 | 950.264 | 950.647 | | | | 950.46 | 0.27 |

**ONPFDNA**

| Time | Replicate 1 | Replicate 2 | Replicate 3 | Replicate 4 | average | SD |
|---|---|---|---|---|---|---|
| 0 | 949.994 | 950.001 | | | 950.00 | 0.00 |
| 30 | 950.299 | 950.342 | | | 950.32 | 0.03 |
| 45 | 950.315 | 950.481 | | | 950.40 | 0.12 |
| 60 | 950.37 | 950.427 | | | 950.40 | 0.04 |
| 300 | 950.307 | 950.455 | | | 950.38 | 0.10 |
| 1500 | 950.428 | 950.406 | | | 950.42 | 0.02 |
| 3600 | 950.499 | 950.6 | | | 950.55 | 0.07 |
| 7200 | 950.489 | 950.58 | | | 950.53 | 0.06 |
| 14400 | 950.485 | 950.347 | | | 950.42 | 0.10 |

**TMG**

| Time | Replicate 1 | Replicate 2 | Replicate 3 | Replicate 4 | Replicate 5 | average | SD |
|---|---|---|---|---|---|---|---|
| 0 | 950.004 | 950.01 | | | | 950.01 | 0.00 |
| 30 | 950.354 | 950.534 | | | | 950.44 | 0.13 |
| 45 | 950.61 | 950.498 | | | | 950.55 | 0.08 |
| 60 | 950.636 | | | | | 950.64 | |
| 300 | 950.745 | 950.826 | | | | 950.79 | 0.06 |
| 1500 | 951.029 | 951.269 | | | | 951.15 | 0.17 |
| 3600 | 951.367 | 951.446 | | | | 951.41 | 0.06 |
| 7200 | 951.306 | 951.682 | | | | 951.49 | 0.27 |
| 14400 | 951.549 | 951.977 | | | | 951.76 | 0.30 |

### 276-290 TEDSSCYIPPLTTIK

**Charge** 2

**IPTG**

| Time | centroid | D |
|---|---|---|
| 0 | 834.895 | -0.07 |
| 30 | 835.118 | 1.00 |
| 45 | 835.1525 | 1.16 |
| 60 | 835.243 | 1.60 |
| 300 | 835.3665 | 2.19 |
| 1500 | 835.758 | 4.07 |
| 3600 | 835.968 | 5.07 |
| 7200 | 836.202 | 6.20 |
| 14400 | 836.175 | 6.07 |

**ONPF**

| Time | centroid | D |
|---|---|---|
| 0 | 834.9085 | -0.01 |
| 30 | 835.056 | 0.70 |
| 45 | 835.23 | 1.53 |
| 60 | 835.2755 | 1.75 |
| 300 | 835.259 | 1.67 |
| 1500 | 835.538 | 3.01 |
| 3600 | 835.81 | 4.32 |
| 7200 | 835.9305 | 4.89 |
| 14400 | 836.35 | 6.92 |

**APO**

| Time | centroid | D |
|---|---|---|
| 0 | 834.911 | 0.00 |
| 30 | 835.0665 | 0.75 |
| 45 | 835.1105 | 0.96 |
| 60 | 835.2145 | 1.46 |
| 300 | 835.2545 | 1.65 |
| 1500 | 835.7565 | 4.06 |
| 3600 | 836.0035 | 5.24 |
| 7200 | 836.2905 | 6.62 |
| 14400 | 836.5885 | 8.05 |

**DNA**

| Time | centroid | D |
|---|---|---|
| 0 | 834.8885 | -0.10 |
| 30 | 835.039 | 0.62 |
| 45 | 835.067 | 0.75 |
| 60 | 835.096 | 0.89 |
| 300 | 835.116 | 0.99 |
| 1500 | 835.2025 | 1.40 |
| 3600 | 835.1715 | 1.25 |
| 7200 | 835.1755 | 1.27 |
| 14400 | 835.167 | 1.23 |

**ONPFDNA**

| Time | centroid | D |
|---|---|---|
| 0 | 834.9055 | -0.02 |
| 30 | 835.0335 | 0.59 |
| 45 | 835.0745 | 0.79 |
| 60 | 835.092 | 0.87 |
| 300 | 835.0595 | 0.72 |
| 1500 | 835.1375 | 1.09 |
| 3600 | 835.212 | 1.45 |
| 7200 | 835.1955 | 1.37 |
| 14400 | 835.1395 | 1.10 |

**TMG**

| Time | centroid | D |
|---|---|---|
| 0 | 834.906 | -0.02 |
| 30 | 835.155 | 1.17 |
| 45 | 835.355 | 2.13 |
| 60 | 835.277 | 1.76 |
| 300 | 835.39 | 2.30 |
| 1500 | 835.685 | 3.72 |
| 3600 | 835.9305 | 4.89 |
| 7200 | 836.083 | 5.62 |
| 14400 | 836.317 | 6.75 |

control 834.91
infinity 837.204

$D(t) = \frac{Mt - M0}{M\infty - M0} *N$

**IPTG**

| Time | Replicate 1 | Replicate 2 | Replicate 3 | Replicate 4 | Replicate5 | average | SD |
|---|---|---|---|---|---|---|---|
| 0 | 834.906 | 834.884 | | | | 834.90 | 0.02 |
| 30 | 835.174 | 835.062 | | | | 835.12 | 0.08 |
| 45 | 835.212 | 835.093 | | | | 835.15 | 0.08 |
| 60 | 835.3 | 835.186 | | | | 835.24 | 0.08 |
| 300 | 835.474 | 835.259 | | | | 835.37 | 0.15 |
| 1500 | 835.837 | 835.679 | | | | 835.76 | 0.11 |
| 3600 | 835.998 | 835.938 | | | | 835.97 | 0.04 |
| 7200 | 836.299 | 836.105 | | | | 836.20 | 0.14 |
| 14400 | 836.282 | 836.068 | | | | 836.18 | 0.15 |

**ONPF**

| Time | Replicate 1 | Replicate 2 | Replicate 3 | Replicate 4 | average | SD |
|---|---|---|---|---|---|---|
| 0 | 834.907 | 834.91 | | | 834.91 | 0.00 |
| 30 | 835.061 | 835.051 | | | 835.06 | 0.01 |
| 45 | 835.381 | 835.079 | | | 835.23 | 0.21 |
| 60 | 835.225 | 835.326 | | | 835.28 | 0.07 |
| 300 | 835.298 | 835.22 | | | 835.26 | 0.06 |
| 1500 | 835.604 | 835.472 | | | 835.54 | 0.09 |
| 3600 | 835.906 | 835.714 | | | 835.81 | 0.14 |
| 7200 | 835.899 | 835.962 | | | 835.93 | 0.04 |
| 14400 | 836.354 | | | | 836.35 | |

**APO**

| Time | Replicate 1 | Replicate 2 | Replicate 3 | Replicate 4 | Replicate 5 | average | SD |
|---|---|---|---|---|---|---|---|
| 0 | 834.903 | 834.919 | | | | 834.91 | 0.01 |
| 30 | 835.08 | 835.053 | | | | 835.07 | 0.02 |
| 45 | 835.123 | 835.098 | | | | 835.11 | 0.02 |
| 60 | 835.129 | 835.3 | | | | 835.21 | 0.12 |
| 300 | 835.274 | 835.235 | | | | 835.25 | 0.03 |
| 1500 | 835.632 | 835.881 | | | | 835.76 | 0.18 |
| 3600 | 836.034 | 835.973 | | | | 836.00 | 0.04 |
| 7200 | 836.407 | 836.174 | | | | 836.29 | 0.16 |
| 14400 | 836.679 | 836.498 | | | | 836.59 | 0.13 |

**DNA**

| Time | Replicate 1 | Replicate 2 | Replicate 3 | Replicate 4 | Replicate 5 | average | SD |
|---|---|---|---|---|---|---|---|
| 0 | 834.892 | 834.885 | | | | 834.89 | 0.00 |
| 30 | 835.07 | 835.008 | | | | 835.04 | 0.04 |
| 45 | 835.102 | 835.032 | | | | 835.07 | 0.05 |
| 60 | 835.112 | 835.08 | | | | 835.10 | 0.02 |
| 300 | 835.157 | 835.075 | | | | 835.12 | 0.06 |
| 1500 | 835.204 | 835.201 | | | | 835.20 | 0.00 |
| 3600 | 835.229 | 835.114 | | | | 835.17 | 0.08 |
| 7200 | 835.227 | 835.124 | | | | 835.18 | 0.07 |
| 14400 | 835.293 | 835.041 | | | | 835.17 | 0.18 |

**ONPFDNA**

| Time | Replicate 1 | Replicate 2 | Replicate 3 | Replicate 4 | average | SD |
|---|---|---|---|---|---|---|
| 0 | 834.902 | 834.909 | | | 834.91 | 0.00 |
| 30 | 835.054 | 835.013 | | | 835.03 | 0.03 |
| 45 | 835.1 | 835.049 | | | 835.07 | 0.04 |
| 60 | 835.107 | 835.077 | | | 835.09 | 0.02 |
| 300 | 835.116 | 835.003 | | | 835.06 | 0.08 |
| 1500 | 835.13 | 835.145 | | | 835.14 | 0.01 |
| 3600 | 835.267 | 835.157 | | | 835.21 | 0.08 |
| 7200 | 835.218 | 835.173 | | | 835.20 | 0.03 |
| 14400 | 835.103 | 835.176 | | | 835.14 | 0.05 |

**TMG**

| Time | Replicate 1 | Replicate 2 | Replicate 3 | Replicate 4 | Replicate 5 | average | SD |
|---|---|---|---|---|---|---|---|
| 0 | 834.918 | 834.894 | | | | 834.91 | 0.02 |
| 30 | 835.101 | 835.209 | | | | 835.16 | 0.08 |
| 45 | 835.437 | 835.273 | | | | 835.36 | 0.12 |
| 60 | 835.277 | | | | | 835.28 | |
| 300 | 835.383 | 835.397 | | | | 835.39 | 0.01 |
| 1500 | 835.613 | 835.757 | | | | 835.69 | 0.10 |
| 3600 | 835.909 | 835.952 | | | | 835.93 | 0.03 |
| 7200 | 835.963 | 836.203 | | | | 836.08 | 0.17 |
| 14400 | 836.186 | 836.448 | | | | 836.32 | 0.19 |

### 279-292 SSCYIPPLTTIKQD

**Charge** 2

**IPTG**

| Time | centroid | D |
|---|---|---|
| 0 | 783.9025 | 0.01 |
| 30 | 784.2165 | 0.84 |
| 45 | 784.2365 | 0.90 |
| 60 | 784.3165 | 1.11 |
| 300 | 784.3625 | 1.23 |
| 1500 | 784.568 | 1.77 |
| 3600 | 784.726 | 2.19 |
| 7200 | 784.8975 | 2.65 |
| 14400 | 784.9435 | 2.77 |

**ONPF**

| Time | centroid | D |
|---|---|---|
| 0 | 783.9145 | 0.04 |
| 30 | 784.1165 | 0.58 |
| 45 | 784.317 | 1.11 |
| 60 | 784.3105 | 1.09 |
| 300 | 784.2805 | 1.01 |
| 1500 | 784.473 | 1.52 |
| 3600 | 784.6905 | 2.10 |
| 7200 | 784.8995 | 2.65 |
| 14400 | 784.74 | 2.24 |

**APO**

| Time | centroid | D |
|---|---|---|
| 0 | 783.8985 | 0.00 |
| 30 | 784.169 | 0.72 |
| 45 | 784.195 | 0.79 |
| 60 | 784.2895 | 1.04 |
| 300 | 784.262 | 0.96 |
| 1500 | 784.658 | 2.01 |
| 3600 | 784.8805 | 2.60 |
| 7200 | 785.204 | 3.46 |
| 14400 | 785.599 | 4.51 |

**DNA**

| Time | centroid | D |
|---|---|---|
| 0 | 783.9085 | 0.03 |
| 30 | 784.1215 | 0.59 |
| 45 | 784.188 | 0.77 |
| 60 | 784.1835 | 0.76 |
| 300 | 784.1945 | 0.78 |
| 1500 | 784.2055 | 0.81 |
| 3600 | 784.257 | 0.95 |
| 7200 | 784.1985 | 0.80 |
| 14400 | 784.1935 | 0.78 |

**ONPFDNA**

| Time | centroid | D |
|---|---|---|
| 0 | 783.9045 | 0.02 |
| 30 | 784.1065 | 0.55 |
| 45 | 784.1455 | 0.65 |
| 60 | 784.1535 | 0.68 |
| 300 | 784.1705 | 0.72 |
| 1500 | 784.1485 | 0.66 |
| 3600 | 784.2355 | 0.89 |
| 7200 | 784.2195 | 0.85 |
| 14400 | 784.167 | 0.71 |

**TMG**

| Time | centroid | D |
|---|---|---|
| 0 | 783.8995 | 0.00 |
| 30 | 784.2005 | 0.80 |
| 45 | 784.3605 | 1.22 |
| 60 | 784.716 | 2.17 |
| 300 | 784.3595 | 1.22 |
| 1500 | 784.5615 | 1.76 |
| 3600 | 784.6805 | 2.07 |
| 7200 | 784.8465 | 2.51 |
| 14400 | 785.0945 | 3.17 |

control 783.8985
infinity 787.671

$$D(t) = \frac{Mt - M0}{M\infty - M0} * N$$

**IPTG**

| Time | Replicate 1 | Replicate 2 | Replicate 3 | Replicate 4 | Replicate5 | average | SD |
|---|---|---|---|---|---|---|---|
| 0 | 783.91 | 783.895 | | | | 783.90 | 0.01 |
| 30 | 784.269 | 784.164 | | | | 784.22 | 0.07 |
| 45 | 784.281 | 784.192 | | | | 784.24 | 0.06 |
| 60 | 784.398 | 784.235 | | | | 784.32 | 0.12 |
| 300 | 784.472 | 784.253 | | | | 784.36 | 0.15 |
| 1500 | 784.627 | 784.509 | | | | 784.57 | 0.08 |
| 3600 | 784.762 | 784.69 | | | | 784.73 | 0.05 |
| 7200 | 784.96 | 784.835 | | | | 784.90 | 0.09 |
| 14400 | 785.043 | 784.844 | | | | 784.94 | 0.14 |

**ONPF**

| Time | Replicate 1 | Replicate 2 | Replicate 3 | Replicate 4 | average | SD |
|---|---|---|---|---|---|---|
| 0 | 783.919 | 783.91 | | | 783.91 | 0.01 |
| 30 | 784.16 | 784.073 | | | 784.12 | 0.06 |
| 45 | 784.466 | 784.168 | | | 784.32 | 0.21 |
| 60 | 784.266 | 784.355 | | | 784.31 | 0.06 |
| 300 | 784.332 | 784.229 | | | 784.28 | 0.07 |
| 1500 | 784.545 | 784.401 | | | 784.47 | 0.10 |
| 3600 | 784.844 | 784.537 | | | 784.69 | 0.22 |
| 7200 | 785.001 | 784.798 | | | 784.90 | 0.14 |
| 14400 | 785.381 | 784.106 | | | 784.74 | 0.90 |

**APO**

| Time | Replicate 1 | Replicate 2 | Replicate 3 | Replicate 4 | Replicate 5 | average | SD |
|---|---|---|---|---|---|---|---|
| 0 | 783.897 | 783.9 | | | | 783.90 | 0.00 |
| 30 | 784.21 | 784.128 | | | | 784.17 | 0.06 |
| 45 | 784.246 | 784.144 | | | | 784.20 | 0.07 |
| 60 | 784.25 | 784.329 | | | | 784.29 | 0.06 |
| 300 | 784.298 | 784.226 | | | | 784.26 | 0.05 |
| 1500 | 784.547 | 784.769 | | | | 784.66 | 0.16 |
| 3600 | 784.906 | 784.855 | | | | 784.88 | 0.04 |
| 7200 | 785.282 | 785.126 | | | | 785.20 | 0.11 |
| 14400 | 785.699 | 785.499 | | | | 785.60 | 0.14 |

**DNA**

| Time | Replicate 1 | Replicate 2 | Replicate 3 | Replicate 4 | Replicate 5 | average | SD |
|---|---|---|---|---|---|---|---|
| 0 | 783.918 | 783.899 | | | | 783.91 | 0.01 |
| 30 | 784.152 | 784.091 | | | | 784.12 | 0.04 |
| 45 | 784.252 | 784.124 | | | | 784.19 | 0.09 |
| 60 | 784.204 | 784.163 | | | | 784.18 | 0.03 |
| 300 | 784.231 | 784.158 | | | | 784.19 | 0.05 |
| 1500 | 784.217 | 784.194 | | | | 784.21 | 0.02 |
| 3600 | 784.345 | 784.169 | | | | 784.26 | 0.12 |
| 7200 | 784.259 | 784.138 | | | | 784.20 | 0.09 |
| 14400 | 784.285 | 784.102 | | | | 784.19 | 0.13 |

**ONPFDNA**

| Time | Replicate 1 | Replicate 2 | Replicate 3 | Replicate 4 | average | SD |
|---|---|---|---|---|---|---|
| 0 | 783.908 | 783.901 | | | 783.90 | 0.00 |
| 30 | 784.15 | 784.063 | | | 784.11 | 0.06 |
| 45 | 784.191 | 784.1 | | | 784.15 | 0.06 |
| 60 | 784.176 | 784.131 | | | 784.15 | 0.03 |
| 300 | 784.199 | 784.142 | | | 784.17 | 0.04 |
| 1500 | 784.132 | 784.165 | | | 784.15 | 0.02 |
| 3600 | 784.245 | 784.226 | | | 784.24 | 0.01 |
| 7200 | 784.259 | 784.18 | | | 784.22 | 0.06 |
| 14400 | 784.158 | 784.176 | | | 784.17 | 0.01 |

**TMG**

| Time | Replicate 1 | Replicate 2 | Replicate 3 | Replicate 4 | Replicate 5 | average | SD |
|---|---|---|---|---|---|---|---|
| 0 | 783.902 | 783.897 | | | | 783.90 | 0.00 |
| 30 | 784.142 | 784.259 | | | | 784.20 | 0.08 |
| 45 | 784.451 | 784.27 | | | | 784.36 | 0.13 |
| 60 | 784.29 | 785.142 | | | | 784.72 | 0.60 |
| 300 | 784.337 | 784.382 | | | | 784.36 | 0.03 |
| 1500 | 784.448 | 784.675 | | | | 784.56 | 0.16 |
| 3600 | 784.659 | 784.702 | | | | 784.68 | 0.03 |
| 7200 | 784.784 | 784.909 | | | | 784.85 | 0.09 |
| 14400 | 784.992 | 785.197 | | | | 785.09 | 0.14 |

### 293-296 FRLL

**Charge** 1

**IPTG**

| Time | centroid | D |
|---|---|---|
| 0 | 548.6745 | 0.01 |
| 30 | 548.722 | 0.07 |
| 45 | 548.7295 | 0.08 |
| 60 | 548.767 | 0.12 |
| 300 | 548.774 | 0.13 |
| 1500 | 548.9555 | 0.36 |
| 3600 | 549.154 | 0.62 |
| 7200 | 549.2785 | 0.77 |
| 14400 | 549.2585 | 0.75 |

**ONPF**

| Time | centroid | D |
|---|---|---|
| 0 | 548.676 | 0.01 |
| 30 | 548.712 | 0.05 |
| 45 | 548.793 | 0.16 |
| 60 | 548.7725 | 0.13 |
| 300 | 548.753 | 0.11 |
| 1500 | 548.8265 | 0.20 |
| 3600 | 548.921 | 0.32 |
| 7200 | 548.9855 | 0.40 |
| 14400 | 548.94 | 0.34 |

**APO**

| Time | centroid | D |
|---|---|---|
| 0 | 548.6695 | 0.00 |
| 30 | 548.7085 | 0.05 |
| 45 | 548.735 | 0.08 |
| 60 | 548.7745 | 0.13 |
| 300 | 548.7455 | 0.10 |
| 1500 | 548.928 | 0.33 |
| 3600 | 549.03 | 0.46 |
| 7200 | 549.112 | 0.56 |
| 14400 | 549.3015 | 0.80 |

**DNA**

| Time | centroid | D |
|---|---|---|
| 0 | 548.6675 | 0.00 |
| 30 | 548.704 | 0.04 |
| 45 | 548.764 | 0.12 |
| 60 | 548.739 | 0.09 |
| 300 | 548.7295 | 0.08 |
| 1500 | 548.7685 | 0.13 |
| 3600 | 548.8355 | 0.21 |
| 7200 | 548.871 | 0.26 |
| 14400 | 548.8855 | 0.27 |

**ONPFDNA**

| Time | centroid | D |
|---|---|---|
| 0 | 548.668 | 0.00 |
| 30 | 548.7055 | 0.05 |
| 45 | 548.722 | 0.07 |
| 60 | 548.731 | 0.08 |
| 300 | 548.721 | 0.07 |
| 1500 | 548.7665 | 0.12 |
| 3600 | 548.8195 | 0.19 |
| 7200 | 548.8335 | 0.21 |
| 14400 | 548.8385 | 0.21 |

**TMG**

| Time | centroid | D |
|---|---|---|
| 0 | 548.6645 | -0.01 |
| 30 | 548.7195 | 0.06 |
| 45 | 548.799 | 0.16 |
| 60 | 548.851 | 0.23 |
| 300 | 548.788 | 0.15 |
| 1500 | 548.864 | 0.25 |
| 3600 | 549.0725 | 0.51 |
| 7200 | 549.139 | 0.60 |
| 14400 | 549.3005 | 0.80 |

control 548.6695
infinity 550.244

$$D(t) = \frac{M_t - M_0}{M_\infty - M_0} *N$$

**IPTG**

| Time | Replicate 1 | Replicate 2 | Replicate 3 | Replicate 4 | Replicate5 | average | SD |
|---|---|---|---|---|---|---|---|
| 0 | 548.677 | 548.672 | | | | 548.67 | 0.00 |
| 30 | 548.74 | 548.704 | | | | 548.72 | 0.03 |
| 45 | 548.745 | 548.714 | | | | 548.73 | 0.02 |
| 60 | 548.772 | 548.762 | | | | 548.77 | 0.01 |
| 300 | 548.786 | 548.762 | | | | 548.77 | 0.02 |
| 1500 | 548.994 | 548.917 | | | | 548.96 | 0.05 |
| 3600 | 549.193 | 549.115 | | | | 549.15 | 0.06 |
| 7200 | 549.348 | 549.209 | | | | 549.28 | 0.10 |
| 14400 | 549.325 | 549.192 | | | | 549.26 | 0.09 |

**ONPF**

| Time | Replicate 1 | Replicate 2 | Replicate 3 | Replicate 4 | average | SD |
|---|---|---|---|---|---|---|
| 0 | 548.671 | 548.681 | | | 548.68 | 0.01 |
| 30 | 548.721 | 548.703 | | | 548.71 | 0.01 |
| 45 | 548.854 | 548.732 | | | 548.79 | 0.09 |
| 60 | 548.759 | 548.786 | | | 548.77 | 0.02 |
| 300 | 548.765 | 548.741 | | | 548.75 | 0.02 |
| 1500 | 548.843 | 548.81 | | | 548.83 | 0.02 |
| 3600 | 549.01 | 548.832 | | | 548.92 | 0.13 |
| 7200 | 549.03 | 548.941 | | | 548.99 | 0.06 |
| 14400 | 549.239 | 548.643 | | | 548.94 | 0.42 |

**APO**

| Time | Replicate 1 | Replicate 2 | Replicate 3 | Replicate 4 | Replicate 5 | average | SD |
|---|---|---|---|---|---|---|---|
| 0 | 548.666 | 548.673 | | | | 548.67 | 0.00 |
| 30 | 548.709 | 548.708 | | | | 548.71 | 0.00 |
| 45 | 548.751 | 548.719 | | | | 548.74 | 0.02 |
| 60 | 548.741 | 548.808 | | | | 548.77 | 0.05 |
| 300 | 548.743 | 548.748 | | | | 548.75 | 0.00 |
| 1500 | 548.866 | 548.99 | | | | 548.93 | 0.09 |
| 3600 | 549.045 | 549.015 | | | | 549.03 | 0.02 |
| 7200 | 549.155 | 549.069 | | | | 549.11 | 0.06 |
| 14400 | 549.39 | 549.213 | | | | 549.30 | 0.13 |

**DNA**

| Time | Replicate 1 | Replicate 2 | Replicate 3 | Replicate 4 | Replicate 5 | average | SD |
|---|---|---|---|---|---|---|---|
| 0 | 548.666 | 548.669 | | | | 548.67 | 0.00 |
| 30 | 548.714 | 548.694 | | | | 548.70 | 0.01 |
| 45 | 548.821 | 548.707 | | | | 548.76 | 0.08 |
| 60 | 548.729 | 548.749 | | | | 548.74 | 0.01 |
| 300 | 548.743 | 548.716 | | | | 548.73 | 0.02 |
| 1500 | 548.777 | 548.76 | | | | 548.77 | 0.01 |
| 3600 | 548.889 | 548.782 | | | | 548.84 | 0.08 |
| 7200 | 548.91 | 548.832 | | | | 548.87 | 0.06 |
| 14400 | 549.027 | 548.744 | | | | 548.89 | 0.20 |

**ONPFDNA**

| Time | Replicate 1 | Replicate 2 | Replicate 3 | Replicate 4 | average | SD |
|---|---|---|---|---|---|---|
| 0 | 548.663 | 548.673 | | | 548.67 | 0.01 |
| 30 | 548.713 | 548.698 | | | 548.71 | 0.01 |
| 45 | 548.734 | 548.71 | | | 548.72 | 0.02 |
| 60 | 548.729 | 548.733 | | | 548.73 | 0.00 |
| 300 | 548.72 | 548.722 | | | 548.72 | 0.00 |
| 1500 | 548.739 | 548.794 | | | 548.77 | 0.04 |
| 3600 | 548.862 | 548.777 | | | 548.82 | 0.06 |
| 7200 | 548.861 | 548.806 | | | 548.83 | 0.04 |
| 14400 | 548.834 | 548.843 | | | 548.84 | 0.01 |

**TMG**

| Time | Replicate 1 | Replicate 2 | Replicate 3 | Replicate 4 | Replicate 5 | average | SD |
|---|---|---|---|---|---|---|---|
| 0 | 548.661 | 548.668 | | | | 548.66 | 0.00 |
| 30 | 548.709 | 548.73 | | | | 548.72 | 0.01 |
| 45 | 548.851 | 548.747 | | | | 548.80 | 0.07 |
| 60 | 548.758 | 548.944 | | | | 548.85 | 0.13 |
| 300 | 548.801 | 548.775 | | | | 548.79 | 0.02 |
| 1500 | 548.838 | 548.89 | | | | 548.86 | 0.04 |
| 3600 | 549.095 | 549.05 | | | | 549.07 | 0.03 |
| 7200 | 549.062 | 549.216 | | | | 549.14 | 0.11 |
| 14400 | 549.251 | 549.35 | | | | 549.30 | 0.07 |

#### 293-301 FRLLGQTSV

**Charge** 1

**IPTG**

| Time | centroid | D |
|---|---|---|
| 0 | 511.09 | -0.03 |
| 30 | 511.23 | 0.33 |
| 45 | 511.22 | 0.31 |
| 60 | 511.27 | 0.43 |
| 300 | 511.28 | 0.47 |
| 1500 | 511.35 | 0.63 |
| 3600 | 511.43 | 0.83 |
| 7200 | 511.51 | 1.03 |
| 14400 | 511.52 | 1.07 |

**ONPF**

| Time | centroid | D |
|---|---|---|
| 0 | 511.09 | -0.03 |
| 30 | 511.18 | 0.21 |
| 45 | | |
| 60 | 511.25 | 0.39 |
| 300 | 511.24 | 0.35 |
| 1500 | 511.28 | 0.45 |
| 3600 | 511.37 | 0.68 |
| 7200 | 511.42 | 0.81 |
| 14400 | 511.60 | 1.28 |

**APO**

| Time | centroid | D |
|---|---|---|
| 0 | 511.10 | 0.00 |
| 30 | 511.19 | 0.24 |
| 45 | 511.27 | 0.42 |
| 60 | 511.29 | 0.48 |
| 300 | 511.27 | 0.43 |
| 1500 | 511.33 | 0.58 |
| 3600 | 511.42 | 0.82 |
| 7200 | 511.51 | 1.05 |
| 14400 | 511.69 | 1.50 |

**DNA**

| Time | centroid | D |
|---|---|---|
| 0 | 511.07 | -0.09 |
| 30 | 511.16 | 0.16 |
| 45 | | |
| 60 | 511.19 | 0.24 |
| 300 | 511.19 | 0.24 |
| 1500 | 511.21 | 0.28 |
| 3600 | 511.26 | 0.40 |
| 7200 | 511.32 | 0.56 |
| 14400 | 511.38 | 0.72 |

**ONPFDNA**

| Time | centroid | D |
|---|---|---|
| 0 | 511.15 | 0.12 |
| 30 | 511.18 | 0.20 |
| 45 | 511.23 | 0.33 |
| 60 | 511.19 | 0.22 |
| 300 | 511.18 | 0.21 |
| 1500 | 511.21 | 0.29 |
| 3600 | 511.26 | 0.40 |
| 7200 | 511.25 | 0.38 |
| 14400 | 511.26 | 0.42 |

**TMG**

| Time | centroid | D |
|---|---|---|
| 0 | 511.07 | -0.07 |
| 30 | 511.21 | 0.27 |
| 45 | 511.25 | 0.37 |
| 60 | 511.29 | 0.47 |
| 300 | 511.27 | 0.42 |
| 1500 | 511.23 | 0.33 |
| 3600 | 511.39 | 0.74 |
| 7200 | 511.40 | 0.77 |
| 14400 | 511.56 | 1.17 |

control 511.1
infinity 513.855

$$D(t) = \frac{M_t - M_0}{M_\infty - M_0} * N$$

**IPTG**

| Time | Replicate 1 | Replicate 2 | Replicate 3 | Replicate 4 | Replicate5 | average | SD |
|---|---|---|---|---|---|---|---|
| 0 | 511.114 | 511.069 | 511.081 | | | 511.09 | 0.02 |
| 30 | 511.296 | 511.201 | 511.189 | | | 511.23 | 0.06 |
| 45 | 511.222 | | | | | 511.22 | |
| 60 | 511.27 | 511.27 | 511.266 | | | 511.27 | 0.00 |
| 300 | 511.355 | 511.246 | 511.252 | | | 511.28 | 0.06 |
| 1500 | 511.353 | 511.359 | 511.334 | | | 511.35 | 0.01 |
| 3600 | 511.518 | 511.382 | 511.38 | | | 511.43 | 0.08 |
| 7200 | 511.56 | 511.493 | 511.465 | | | 511.51 | 0.05 |
| 14400 | 511.594 | 511.527 | 511.448 | | | 511.52 | 0.07 |

**ONPF**

| Time | Replicate 1 | Replicate 2 | Replicate 3 | Replicate 4 | average | SD |
|---|---|---|---|---|---|---|
| 0 | 511.102 | 511.082 | 511.075 | | 511.09 | 0.01 |
| 30 | 511.162 | 511.175 | 511.208 | | 511.18 | 0.02 |
| 45 | | | | | | |
| 60 | 511.174 | 511.237 | 511.346 | | 511.25 | 0.09 |
| 300 | 511.255 | 511.205 | 511.248 | | 511.24 | 0.03 |
| 1500 | 511.242 | 511.277 | 511.318 | | 511.28 | 0.04 |
| 3600 | 511.371 | 511.307 | 511.425 | | 511.37 | 0.06 |
| 7200 | 511.355 | 511.429 | 511.468 | | 511.42 | 0.06 |
| 14400 | 511.626 | 511.545 | 511.636 | | 511.60 | 0.05 |

**APO**

| Time | Replicate 1 | Replicate 2 | Replicate 3 | Replicate 4 | Replicate 5 | average | SD |
|---|---|---|---|---|---|---|---|
| 0 | 511.029 | 511.228 | 511.094 | 511.066 | 511.085 | 511.10 | 0.08 |
| 30 | 511.165 | | 511.282 | 511.174 | 511.158 | 511.19 | 0.06 |
| 45 | 511.27 | 511.261 | | | | 511.27 | 0.01 |
| 60 | 511.186 | 511.326 | 511.423 | 511.247 | 511.267 | 511.29 | 0.09 |
| 300 | 511.354 | 511.241 | 511.323 | 511.223 | 511.209 | 511.27 | 0.06 |
| 1500 | 511.167 | 511.421 | 511.381 | 511.348 | 511.328 | 511.33 | 0.10 |
| 3600 | | 511.434 | 511.499 | 511.389 | 511.374 | 511.42 | 0.06 |
| 7200 | | 511.4 | 511.563 | 511.566 | 511.525 | 511.51 | 0.08 |
| 14400 | 511.705 | 511.651 | 511.805 | 511.639 | 511.649 | 511.69 | 0.07 |

**DNA**

| Time | Replicate 1 | Replicate 2 | Replicate 3 | Replicate 4 | Replicate 5 | average | SD |
|---|---|---|---|---|---|---|---|
| 0 | 511.061 | 511.072 | | | | 511.07 | 0.01 |
| 30 | 511.174 | 511.152 | | | | 511.16 | 0.02 |
| 45 | | | | | | | |
| 60 | 511.207 | 511.182 | | | | 511.19 | 0.02 |
| 300 | 511.216 | 511.172 | | | | 511.19 | 0.03 |
| 1500 | 511.212 | 511.212 | | | | 511.21 | 0.00 |
| 3600 | 511.305 | 511.212 | | | | 511.26 | 0.07 |
| 7200 | 511.341 | 511.303 | | | | 511.32 | 0.03 |
| 14400 | 511.457 | 511.309 | | | | 511.38 | 0.10 |

**ONPFDNA**

| Time | Replicate 1 | Replicate 2 | Replicate 3 | Replicate 4 | average | SD |
|---|---|---|---|---|---|---|
| 0 | 511.104 | 511.255 | 511.079 | | 511.15 | 0.10 |
| 30 | 511.169 | 511.224 | 511.146 | | 511.18 | 0.04 |
| 45 | 511.194 | 511.264 | | | 511.23 | 0.05 |
| 60 | 511.172 | | 511.203 | | 511.19 | 0.02 |
| 300 | 511.176 | | 511.192 | | 511.18 | 0.01 |
| 1500 | 511.143 | 511.308 | 511.192 | | 511.21 | 0.08 |
| 3600 | 511.273 | 511.268 | 511.231 | | 511.26 | 0.02 |
| 7200 | 511.251 | 511.248 | 511.252 | | 511.25 | 0.00 |
| 14400 | 511.193 | 511.268 | 511.33 | | 511.26 | 0.07 |

**TMG**

| Time | Replicate 1 | Replicate 2 | Replicate 3 | Replicate 4 | Replicate 5 | average | SD |
|---|---|---|---|---|---|---|---|
| 0 | 511.09 | 511.058 | | | | 511.07 | 0.02 |
| 30 | 511.151 | 511.259 | | | | 511.21 | 0.08 |
| 45 | 511.32 | 511.173 | | | | 511.25 | 0.10 |
| 60 | 511.21 | 511.36 | | | | 511.29 | 0.11 |
| 300 | 511.228 | 511.306 | | | | 511.27 | 0.06 |
| 1500 | 511.282 | 511.176 | | | | 511.23 | 0.07 |
| 3600 | 511.39 | | | | | 511.39 | |
| 7200 | 511.371 | 511.432 | | | | 511.40 | 0.04 |
| 14400 | 511.529 | 511.589 | | | | 511.56 | 0.04 |

### 296-303 LGQTSVDR

**Charge** 2

**IPTG**

| Time | centroid | D |
|---|---|---|
| 0 | 438.474 | 0.02 |
| 30 | 438.6195 | 0.48 |
| 45 | 438.6285 | 0.51 |
| 60 | 438.6435 | 0.56 |
| 300 | 438.634 | 0.53 |
| 1500 | 438.625 | 0.50 |
| 3600 | 438.615 | 0.47 |
| 7200 | 438.6485 | 0.58 |
| 14400 | 438.636 | 0.54 |

**ONPF**

| Time | centroid | D |
|---|---|---|
| 0 | 438.476 | 0.03 |
| 30 | 438.5825 | 0.37 |
| 45 | 438.6215 | 0.49 |
| 60 | 438.6125 | 0.46 |
| 300 | 438.612 | 0.46 |
| 1500 | 438.6185 | 0.48 |
| 3600 | 438.6265 | 0.51 |
| 7200 | 438.671 | 0.65 |
| 14400 | 438.63 | 0.50 |

**APO**

| Time | centroid | D |
|---|---|---|
| 0 | 438.467 | 0.00 |
| 30 | 438.59 | 0.39 |
| 45 | 438.5975 | 0.41 |
| 60 | 438.611 | 0.46 |
| 300 | 438.586 | 0.38 |
| 1500 | 438.626 | 0.50 |
| 3600 | 438.638 | 0.54 |
| 7200 | 438.669 | 0.64 |
| 14400 | 438.803 | 1.07 |

**DNA**

| Time | centroid | D |
|---|---|---|
| 0 | 438.4635 | -0.01 |
| 30 | 438.587 | 0.38 |
| 45 | 438.584 | 0.37 |
| 60 | 438.605 | 0.44 |
| 300 | 438.6 | 0.42 |
| 1500 | 438.597 | 0.41 |
| 3600 | 438.6025 | 0.43 |
| 7200 | 438.612 | 0.46 |
| 14400 | 438.605 | 0.44 |

**ONPFDNA**

| Time | centroid | D |
|---|---|---|
| 0 | 438.465 | -0.01 |
| 30 | 438.581 | 0.36 |
| 45 | 438.5985 | 0.42 |
| 60 | 438.5955 | 0.41 |
| 300 | 438.581 | 0.36 |
| 1500 | 438.5805 | 0.36 |
| 3600 | 438.5995 | 0.42 |
| 7200 | 438.618 | 0.48 |
| 14400 | 438.589 | 0.39 |

**TMG**

| Time | centroid | D |
|---|---|---|
| 0 | 438.4695 | 0.01 |
| 30 | 438.6425 | 0.56 |
| 45 | 438.646 | 0.57 |
| 60 | 438.7505 | 0.90 |
| 300 | 438.662 | 0.62 |
| 1500 | 438.6745 | 0.66 |
| 3600 | 438.662 | 0.62 |
| 7200 | 438.657 | 0.60 |
| 14400 | 438.7105 | 0.77 |

control 438.467
infinity 440.358

$$D(t) = \frac{Mt - M0}{M\infty - M0} *N$$

**IPTG**

| Time | Replicate 1 | Replicate 2 | Replicate 3 | Replicate 4 | Replicate5 | average | SD |
|---|---|---|---|---|---|---|---|
| 0 | 438.479 | 438.469 | | | | 438.47 | 0.01 |
| 30 | 438.667 | 438.572 | | | | 438.62 | 0.07 |
| 45 | 438.665 | 438.592 | | | | 438.63 | 0.05 |
| 60 | 438.7 | 438.587 | | | | 438.64 | 0.08 |
| 300 | 438.685 | 438.583 | | | | 438.63 | 0.07 |
| 1500 | 438.658 | 438.592 | | | | 438.63 | 0.05 |
| 3600 | 438.648 | 438.582 | | | | 438.62 | 0.05 |
| 7200 | 438.684 | 438.613 | | | | 438.65 | 0.05 |
| 14400 | 438.672 | 438.6 | | | | 438.64 | 0.05 |

**ONPF**

| Time | Replicate 1 | Replicate 2 | Replicate 3 | Replicate 4 | average | SD |
|---|---|---|---|---|---|---|
| 0 | 438.475 | 438.477 | | | 438.48 | 0.00 |
| 30 | 438.608 | 438.557 | | | 438.58 | 0.04 |
| 45 | 438.663 | 438.58 | | | 438.62 | 0.06 |
| 60 | 438.63 | 438.595 | | | 438.61 | 0.02 |
| 300 | 438.64 | 438.584 | | | 438.61 | 0.04 |
| 1500 | 438.644 | 438.593 | | | 438.62 | 0.04 |
| 3600 | 438.675 | 438.578 | | | 438.63 | 0.07 |
| 7200 | 438.748 | 438.594 | | | 438.67 | 0.11 |
| 14400 | 438.793 | 438.459 | | | 438.63 | 0.24 |

**APO**

| Time | Replicate 1 | Replicate 2 | Replicate 3 | Replicate 4 | Replicate 5 | average | SD |
|---|---|---|---|---|---|---|---|
| 0 | 438.466 | 438.468 | | | | 438.47 | 0.00 |
| 30 | 438.62 | 438.56 | | | | 438.59 | 0.04 |
| 45 | 438.632 | 438.563 | | | | 438.60 | 0.05 |
| 60 | 438.643 | 438.579 | | | | 438.61 | 0.05 |
| 300 | 438.618 | 438.554 | | | | 438.59 | 0.05 |
| 1500 | 438.643 | 438.609 | | | | 438.63 | 0.02 |
| 3600 | 438.669 | 438.607 | | | | 438.64 | 0.04 |
| 7200 | 438.72 | 438.618 | | | | 438.67 | 0.07 |
| 14400 | 438.907 | 438.699 | | | | 438.80 | 0.15 |

**DNA**

| Time | Replicate 1 | Replicate 2 | Replicate 3 | Replicate 4 | Replicate 5 | average | SD |
|---|---|---|---|---|---|---|---|
| 0 | 438.461 | 438.466 | | | | 438.46 | 0.00 |
| 30 | 438.617 | 438.557 | | | | 438.59 | 0.04 |
| 45 | 438.604 | 438.564 | | | | 438.58 | 0.03 |
| 60 | 438.647 | 438.563 | | | | 438.61 | 0.06 |
| 300 | 438.632 | 438.568 | | | | 438.60 | 0.05 |
| 1500 | 438.62 | 438.574 | | | | 438.60 | 0.03 |
| 3600 | 438.639 | 438.566 | | | | 438.60 | 0.05 |
| 7200 | 438.647 | 438.577 | | | | 438.61 | 0.05 |
| 14400 | 438.67 | 438.54 | | | | 438.61 | 0.09 |

**ONPFDNA**

| Time | Replicate 1 | Replicate 2 | Replicate 3 | Replicate 4 | average | SD |
|---|---|---|---|---|---|---|
| 0 | 438.465 | 438.465 | | | 438.47 | 0.00 |
| 30 | 438.597 | 438.565 | | | 438.58 | 0.02 |
| 45 | 438.612 | 438.585 | | | 438.60 | 0.02 |
| 60 | 438.612 | 438.579 | | | 438.60 | 0.02 |
| 300 | 438.593 | 438.569 | | | 438.58 | 0.02 |
| 1500 | 438.582 | 438.579 | | | 438.58 | 0.00 |
| 3600 | 438.612 | 438.587 | | | 438.60 | 0.02 |
| 7200 | 438.655 | 438.581 | | | 438.62 | 0.05 |
| 14400 | 438.591 | 438.587 | | | 438.59 | 0.00 |

**TMG**

| Time | Replicate 1 | Replicate 2 | Replicate 3 | Replicate 4 | Replicate 5 | average | SD |
|---|---|---|---|---|---|---|---|
| 0 | 438.473 | 438.466 | | | | 438.47 | 0.00 |
| 30 | 438.621 | 438.664 | | | | 438.64 | 0.03 |
| 45 | 438.628 | 438.664 | | | | 438.65 | 0.03 |
| 60 | 438.659 | 438.842 | | | | 438.75 | 0.13 |
| 300 | 438.648 | 438.676 | | | | 438.66 | 0.02 |
| 1500 | 438.621 | 438.728 | | | | 438.67 | 0.08 |
| 3600 | 438.665 | 438.659 | | | | 438.66 | 0.00 |
| 7200 | 438.637 | 438.677 | | | | 438.66 | 0.03 |
| 14400 | 438.677 | 438.744 | | | | 438.71 | 0.05 |

### 297-304 GQTSVDRL

**Charge** 1

**IPTG**

| Time | centroid | D |
|---|---|---|
| 0 | 438.474 | 0.04 |
| 30 | 438.574 | 0.34 |
| 45 | 438.5835 | 0.37 |
| 60 | 438.5875 | 0.38 |
| 300 | 438.6045 | 0.43 |
| 1500 | 438.586 | 0.38 |
| 3600 | 438.5955 | 0.40 |
| 7200 | 438.636 | 0.52 |
| 14400 | 438.556 | 0.29 |

**ONPF**

| Time | centroid | D |
|---|---|---|
| 0 | 438.47 | 0.04 |
| 30 | 438.55 | 0.27 |
| 45 | 438.57 | 0.34 |
| 60 | 438.58 | 0.35 |
| 300 | 438.58 | 0.34 |
| 1500 | 438.59 | 0.39 |
| 3600 | 438.58 | 0.35 |
| 7200 | 438.58 | 0.36 |
| 14400 | | |

**APO**

| Time | centroid | D |
|---|---|---|
| 0 | 438.4625 | 0.01 |
| 30 | 438.568 | 0.32 |
| 45 | 438.5625 | 0.31 |
| 60 | 438.5735 | 0.34 |
| 300 | 438.5585 | 0.29 |
| 1500 | 438.614 | 0.46 |
| 3600 | 438.579 | 0.35 |
| 7200 | 438.6085 | 0.44 |
| 14400 | 438.685 | 0.67 |

**DNA**

| Time | centroid | D |
|---|---|---|
| 0 | 438.458 | -0.01 |
| 30 | 438.5475 | 0.26 |
| 45 | 438.5675 | 0.32 |
| 60 | 438.563 | 0.31 |
| 300 | 438.5745 | 0.34 |
| 1500 | 438.5855 | 0.37 |
| 3600 | 438.5885 | 0.38 |
| 7200 | 438.5655 | 0.31 |
| 14400 | 438.5725 | 0.33 |

**ONPFDNA**

| Time | centroid | D |
|---|---|---|
| 0 | 438.462 | 0.01 |
| 30 | 438.5545 | 0.28 |
| 45 | 438.561 | 0.30 |
| 60 | 438.5595 | 0.30 |
| 300 | 438.5555 | 0.28 |
| 1500 | 438.549 | 0.27 |
| 3600 | 438.564 | 0.31 |
| 7200 | 438.563 | 0.31 |
| 14400 | 438.5505 | 0.27 |

**TMG**

| Time | centroid | D |
|---|---|---|
| 0 | 438.46 | -0.01 |
| 30 | | |
| 45 | 438.58 | 0.35 |
| 60 | 438.65 | 0.56 |
| 300 | 438.59 | 0.38 |
| 1500 | 438.62 | 0.46 |
| 3600 | 438.61 | 0.43 |
| 7200 | 438.60 | 0.42 |
| 14400 | | |

| | |
|---|---|
| control | 438.46 |
| infinity | 440.475 |

$$D(t) = \frac{M_t - M_0}{M_\infty - M_0} \cdot N$$

**IPTG**

| Time | Replicate 1 | Replicate 2 | Replicate 3 | Replicate 4 | Replicate5 | average | SD |
|---|---|---|---|---|---|---|---|
| 0 | 438.487 | 438.461 | | | | 438.47 | 0.02 |
| 30 | 438.605 | 438.543 | | | | 438.57 | 0.04 |
| 45 | 438.617 | 438.55 | | | | 438.58 | 0.05 |
| 60 | 438.628 | 438.547 | | | | 438.59 | 0.06 |
| 300 | 438.666 | 438.543 | | | | 438.60 | 0.09 |
| 1500 | 438.622 | 438.55 | | | | 438.59 | 0.05 |
| 3600 | 438.634 | 438.557 | | | | 438.60 | 0.05 |
| 7200 | 438.636 | | | | | 438.64 | |
| 14400 | | 438.556 | | | | 438.56 | |

**ONPF**

| Time | Replicate 1 | Replicate 2 | Replicate 3 | Replicate 4 | average | SD |
|---|---|---|---|---|---|---|
| 0 | 438.467 | 438.477 | | | 438.47 | 0.01 |
| 30 | 438.565 | 438.534 | | | 438.55 | 0.02 |
| 45 | 438.596 | 438.551 | | | 438.57 | 0.03 |
| 60 | 438.59 | 438.562 | | | 438.58 | 0.02 |
| 300 | 438.596 | 438.554 | | | 438.58 | 0.03 |
| 1500 | 438.611 | 438.57 | | | 438.59 | 0.03 |
| 3600 | 438.607 | 438.551 | | | 438.58 | 0.04 |
| 7200 | 438.608 | 438.554 | | | 438.58 | 0.04 |
| 14400 | | | | | #DIV/0! | |

**APO**

| Time | Replicate 1 | Replicate 2 | Replicate 3 | Replicate 4 | Replicate 5 | average | SD |
|---|---|---|---|---|---|---|---|
| 0 | 438.465 | 438.46 | | | | 438.46 | 0.00 |
| 30 | 438.592 | 438.544 | | | | 438.57 | 0.03 |
| 45 | 438.59 | 438.535 | | | | 438.56 | 0.04 |
| 60 | 438.595 | 438.552 | | | | 438.57 | 0.03 |
| 300 | 438.583 | 438.534 | | | | 438.56 | 0.03 |
| 1500 | 438.653 | 438.575 | | | | 438.61 | 0.06 |
| 3600 | 438.583 | 438.575 | | | | 438.58 | 0.01 |
| 7200 | 438.661 | 438.556 | | | | 438.61 | 0.07 |
| 14400 | 438.745 | 438.625 | | | | 438.69 | 0.08 |

**DNA**

| Time | Replicate 1 | Replicate 2 | Replicate 3 | Replicate 4 | Replicate 5 | average | SD |
|---|---|---|---|---|---|---|---|
| 0 | 438.458 | 438.458 | | | | 438.46 | 0.00 |
| 30 | 438.565 | 438.53 | | | | 438.55 | 0.02 |
| 45 | 438.59 | 438.545 | | | | 438.57 | 0.03 |
| 60 | 438.581 | 438.545 | | | | 438.56 | 0.03 |
| 300 | 438.607 | 438.542 | | | | 438.57 | 0.05 |
| 1500 | 438.605 | 438.566 | | | | 438.59 | 0.03 |
| 3600 | 438.631 | 438.546 | | | | 438.59 | 0.06 |
| 7200 | 438.601 | 438.53 | | | | 438.57 | 0.05 |
| 14400 | 438.619 | 438.526 | | | | 438.57 | 0.07 |

**ONPFDNA**

| Time | Replicate 1 | Replicate 2 | Replicate 3 | Replicate 4 | average | SD |
|---|---|---|---|---|---|---|
| 0 | 438.465 | 438.459 | | | 438.46 | 0.00 |
| 30 | 438.569 | 438.54 | | | 438.55 | 0.02 |
| 45 | 438.568 | 438.554 | | | 438.56 | 0.01 |
| 60 | 438.574 | 438.545 | | | 438.56 | 0.02 |
| 300 | 438.565 | 438.546 | | | 438.56 | 0.01 |
| 1500 | 438.54 | 438.558 | | | 438.55 | 0.01 |
| 3600 | 438.578 | 438.55 | | | 438.56 | 0.02 |
| 7200 | 438.593 | 438.533 | | | 438.56 | 0.04 |
| 14400 | 438.559 | 438.542 | | | 438.55 | 0.01 |

**TMG**

| Time | Replicate 1 | Replicate 2 | Replicate 3 | Replicate 4 | Replicate 5 | average | SD |
|---|---|---|---|---|---|---|---|
| 0 | 438.459 | 438.453 | | | | 438.46 | 0.00 |
| 30 | | | | | | | |
| 45 | 438.576 | | | | | 438.58 | |
| 60 | 438.584 | 438.715 | | | | 438.65 | 0.09 |
| 300 | 438.586 | | | | | 438.59 | |
| 1500 | 438.559 | 438.671 | | | | 438.62 | 0.08 |
| 3600 | 438.597 | 438.613 | | | | 438.61 | 0.01 |
| 7200 | 438.578 | 438.626 | | | | 438.60 | 0.03 |
| 14400 | | | | | | | |

### 304-307 LLQL

**Charge** 1

**IPTG**

| Time | centroid | D |
|---|---|---|
| 0 | 486.6065 | 0.00 |
| 30 | 486.8875 | 0.36 |
| 45 | 486.963 | 0.46 |
| 60 | 487.026 | 0.54 |
| 300 | 487.2025 | 0.77 |
| 1500 | 487.4555 | 1.10 |
| 3600 | 487.7605 | 1.50 |
| 7200 | 487.803 | 1.56 |
| 14400 | 487.7145 | 1.44 |

**ONPF**

| Time | centroid | D |
|---|---|---|
| 0 | 486.629 | 0.02 |
| 30 | 486.722 | 0.15 |
| 45 | 486.867 | 0.34 |
| 60 | 486.951 | 0.44 |
| 300 | 487.041 | 0.56 |
| 1500 | 487.2275 | 0.81 |
| 3600 | 487.3195 | 0.93 |
| 7200 | 487.3185 | 0.92 |
| 14400 | 487.24 | 0.82 |

**APO**

| Time | centroid | D |
|---|---|---|
| 0 | 486.607 | 0.00 |
| 30 | 486.7885 | 0.23 |
| 45 | 486.8695 | 0.34 |
| 60 | 487.0185 | 0.53 |
| 300 | 487.1405 | 0.69 |
| 1500 | 487.443 | 1.09 |
| 3600 | 487.639 | 1.34 |
| 7200 | 487.605 | 1.30 |
| 14400 | 487.73 | 1.46 |

**DNA**

| Time | centroid | D |
|---|---|---|
| 0 | 486.607 | 0.00 |
| 30 | 486.726 | 0.15 |
| 45 | 486.964 | 0.46 |
| 60 | 487.0085 | 0.52 |
| 300 | 487.094 | 0.63 |
| 1500 | 487.373 | 0.99 |
| 3600 | 487.423 | 1.06 |
| 7200 | 487.4135 | 1.05 |
| 14400 | 487.3465 | 0.96 |

**ONPFDNA**

| Time | centroid | D |
|---|---|---|
| 0 | 486.594 | -0.02 |
| 30 | 486.704 | 0.12 |
| 45 | 486.803 | 0.25 |
| 60 | 486.9125 | 0.39 |
| 300 | 486.9915 | 0.50 |
| 1500 | 487.1525 | 0.71 |
| 3600 | 487.4065 | 1.04 |
| 7200 | 487.31 | 0.91 |
| 14400 | 487.2065 | 0.78 |

**TMG**

| Time | centroid | D |
|---|---|---|
| 0 | 486.6125 | 0.00 |
| 30 | 486.9065 | 0.39 |
| 45 | 487.0195 | 0.53 |
| 60 | 486.9685 | 0.47 |
| 300 | 487.3065 | 0.91 |
| 1500 | 487.4165 | 1.05 |
| 3600 | 487.6965 | 1.42 |
| 7200 | 487.6925 | 1.41 |
| 14400 | 487.879 | 1.65 |

control 486.61
infinity 488.144

$$D(t) = \frac{M_t - M_0}{M_\infty - M_0} * N$$

**IPTG**

| Time | Replicate 1 | Replicate 2 | Replicate 3 | Replicate 4 | Replicate5 | average | SD |
|---|---|---|---|---|---|---|---|
| 0 | 486.61 | 486.603 | | | | 486.61 | 0.00 |
| 30 | 486.978 | 486.797 | | | | 486.89 | 0.13 |
| 45 | 487 | 486.926 | | | | 486.96 | 0.05 |
| 60 | 486.988 | 487.064 | | | | 487.03 | 0.05 |
| 300 | 487.238 | 487.167 | | | | 487.20 | 0.05 |
| 1500 | 487.609 | 487.302 | | | | 487.46 | 0.22 |
| 3600 | 487.794 | 487.727 | | | | 487.76 | 0.05 |
| 7200 | 487.932 | 487.674 | | | | 487.80 | 0.18 |
| 14400 | 487.821 | 487.608 | | | | 487.71 | 0.15 |

**ONPF**

| Time | Replicate 1 | Replicate 2 | Replicate 3 | Replicate 4 | average | SD |
|---|---|---|---|---|---|---|
| 0 | 486.634 | 486.624 | | | 486.63 | 0.01 |
| 30 | 486.766 | 486.678 | | | 486.72 | 0.06 |
| 45 | 487.026 | 486.708 | | | 486.87 | 0.22 |
| 60 | 486.984 | 486.918 | | | 486.95 | 0.05 |
| 300 | 487.164 | 486.918 | | | 487.04 | 0.17 |
| 1500 | 487.389 | 487.066 | | | 487.23 | 0.23 |
| 3600 | 487.536 | 487.103 | | | 487.32 | 0.31 |
| 7200 | 487.473 | 487.164 | | | 487.32 | 0.22 |
| 14400 | 487.733 | 486.739 | | | 487.24 | 0.70 |

**APO**

| Time | Replicate 1 | Replicate 2 | Replicate 3 | Replicate 4 | Replicate 5 | average | SD |
|---|---|---|---|---|---|---|---|
| 0 | 486.602 | 486.612 | | | | 486.61 | 0.01 |
| 30 | 486.787 | 486.79 | | | | 486.79 | 0.00 |
| 45 | 486.905 | 486.834 | | | | 486.87 | 0.05 |
| 60 | 486.931 | 487.106 | | | | 487.02 | 0.12 |
| 300 | 487.197 | 487.084 | | | | 487.14 | 0.08 |
| 1500 | 487.388 | 487.498 | | | | 487.44 | 0.08 |
| 3600 | 487.694 | 487.584 | | | | 487.64 | 0.08 |
| 7200 | 487.74 | 487.47 | | | | 487.61 | 0.19 |
| 14400 | 487.839 | 487.621 | | | | 487.73 | 0.15 |

**DNA**

| Time | Replicate 1 | Replicate 2 | Replicate 3 | Replicate 4 | Replicate 5 | average | SD |
|---|---|---|---|---|---|---|---|
| 0 | 486.593 | 486.621 | | | | 486.61 | 0.02 |
| 30 | 486.729 | 486.723 | | | | 486.73 | 0.00 |
| 45 | 487.169 | 486.759 | | | | 486.96 | 0.29 |
| 60 | 486.98 | 487.037 | | | | 487.01 | 0.04 |
| 300 | 487.115 | 487.073 | | | | 487.09 | 0.03 |
| 1500 | 487.369 | 487.377 | | | | 487.37 | 0.01 |
| 3600 | 487.527 | 487.319 | | | | 487.42 | 0.15 |
| 7200 | 487.493 | 487.334 | | | | 487.41 | 0.11 |
| 14400 | 487.717 | 486.976 | | | | 487.35 | 0.52 |

**ONPFDNA**

| Time | Replicate 1 | Replicate 2 | Replicate 3 | Replicate 4 | average | SD |
|---|---|---|---|---|---|---|
| 0 | 486.603 | 486.585 | | | 486.59 | 0.01 |
| 30 | 486.72 | 486.688 | | | 486.70 | 0.02 |
| 45 | 486.884 | 486.722 | | | 486.80 | 0.11 |
| 60 | 486.945 | 486.88 | | | 486.91 | 0.05 |
| 300 | 487.046 | 486.937 | | | 486.99 | 0.08 |
| 1500 | 487.146 | 487.159 | | | 487.15 | 0.01 |
| 3600 | 487.525 | 487.288 | | | 487.41 | 0.17 |
| 7200 | 487.381 | 487.239 | | | 487.31 | 0.10 |
| 14400 | 487.371 | 487.042 | | | 487.21 | 0.23 |

**TMG**

| Time | Replicate 1 | Replicate 2 | Replicate 3 | Replicate 4 | Replicate 5 | average | SD |
|---|---|---|---|---|---|---|---|
| 0 | 486.623 | 486.602 | | | | 486.61 | 0.01 |
| 30 | 486.851 | 486.962 | | | | 486.91 | 0.08 |
| 45 | 487.144 | 486.895 | | | | 487.02 | 0.18 |
| 60 | 487.032 | 486.905 | | | | 486.97 | 0.09 |
| 300 | 487.283 | 487.33 | | | | 487.31 | 0.03 |
| 1500 | 487.334 | 487.499 | | | | 487.42 | 0.12 |
| 3600 | 487.615 | 487.778 | | | | 487.70 | 0.12 |
| 7200 | 487.478 | 487.907 | | | | 487.69 | 0.30 |
| 14400 | 487.77 | 487.988 | | | | 487.88 | 0.15 |
